# DNMT3A^R882H^ Is Not Required for Disease Maintenance in Primary Human AML, but Is Associated With Increased Leukemia Stem Cell Frequency

**DOI:** 10.1101/2024.10.26.620318

**Authors:** Thomas Köhnke, Daiki Karigane, Eleanor Hilgart, Amy C. Fan, Kensuke Kayamori, Masashi Miyauchi, Cailin T. Collins, Fabian P. Suchy, Athreya Rangavajhula, Yang Feng, Yusuke Nakauchi, Eduardo Martinez-Montes, Jonas L. Fowler, Kyle M. Loh, Hiromitsu Nakauchi, Michael A. Koldobskiy, Andrew P. Feinberg, Ravindra Majeti

**Affiliations:** Department of Medicine, Division of Hematology, Cancer Institute, and Institute for Stem Cell Biology and Regenerative Medicine, Stanford University; Stanford, CA, 94305, USA; Center for Epigenetics and Department of Medicine, Johns Hopkins University School of Medicine; Baltimore, MD, 21205, USA; Department of Developmental Biology and Institute for Stem Cell Biology and Regenerative Medicine, Stanford University; Stanford, CA, 94305, USA

## Abstract

Genetic mutations are being thoroughly mapped in human cancers, yet a fundamental question in cancer biology is whether such mutations are functionally required for cancer initiation, maintenance of established cancer, or both. Here, we study this question in the context of human acute myeloid leukemia (AML), where *DNMT3A^R882^* missense mutations often arise early, in pre-leukemic clonal hematopoiesis, and corrupt the DNA methylation landscape to initiate leukemia. We developed CRISPR-based methods to directly correct *DNMT3A^R882^* mutations in leukemic cells obtained from patients. Surprisingly, *DNMT3A^R882^* mutations were largely dispensable for disease maintenance. Replacing *DNMT3A^R882^* mutants with wild-type *DNMT3A* did not impair the ability of AML cells to engraft *in vivo*, and minimally altered DNA methylation. Taken together, *DNMT3A^R882^* mutations are initially necessary for AML initiation, but are largely dispensable for disease maintenance. The notion that initiating oncogenes differ from those that maintain cancer has important implications for cancer evolution and therapy.

**STATEMENT OF SIGNIFICANCE:** Understanding which driver mutations are required for cancer initiation, maintenance, or both phases remains poorly understood. Here, we uncover that highly prevalent pre-leukemic *DNMT3A* mutations are only required during disease initiation, but become dispensable after leukemic transformation, uncovering the context-specific role of this driver mutation with important therapeutic implications.

## INTRODUCTION

An important and understudied question in cancer pathogenesis is whether a given genetic mutation is required for disease initiation, maintenance of established disease, or both. In myeloid malignancies, previous studies in mouse models interrogating the role of the driver mutations *IDH2^R140Q^* and *JAK2^V617F^* have suggested that their continued presence is required for both disease initiation and maintenance.^1,2^ Likewise, within *in vivo* mouse models of colorectal cancer, correction of the *Apc* driver mutation in established cancer cells leads to disease regression, again suggesting that the driver mutation is required for both cancer initiation and maintenance.^3^ However, whether this principle extends to all genetic mutations found in cancer, especially those acquired early during pre-malignant evolution, is unclear. Delineating the specific contribution of individual mutations across the disease course directly in patient specimens is hampered by substantial variability in co-mutations, germline background, inter-patient variability, and technical challenges in molecularly manipulating primary human cells. Thus, prospective, functional interrogation of how specific somatic mutations affect these cellular phenotypes, as well as the biology of cancer stem cells, have been performed in surrogate systems such as mouse disease models and cell lines. To overcome these limitations and explore this question in human acute myeloid leukemia (AML), we developed an approach to perform functional genetic experiments directly in AML patient specimens by leveraging CRISPR/Cas9 to selectively and specifically correct individual somatic mutations and assay the effects on disease initiation, disease maintenance and leukemia stem cell phenotypes.

We focused on *DNMT3A^R882^* missense mutations, which are among the most frequently occurring founding mutations in human clonal hematopoiesis and acute myeloid leukemia (AML).^4–6^ Mutations in *DNMT3A* impair its ability to methylate DNA CpGs^7–9^, leading to increased self-renewal of hematopoietic stem cells (HSCs),^10^ and have been associated with poor treatment outcomes in AML.^4,11^ During disease evolution, *DNMT3A^R882^* missense mutations are frequently acquired early, preceding leukemic transformation,^12^ suggesting a critical role in disease initiation. However, it is possible that such mutations might be required to provide the epigenetic background necessary for disease initiation but may no longer be required for maintenance of established disease in the presence of other somatic driver mutations. Since current approaches are unable to discern whether *DNMT3A^R882^* missense mutations contribute to pre-leukemic and leukemic phases of human AML equally, we aimed to functionally interrogate the disease-phase specific contribution directly in patient specimens.

## RESULTS

### Correction of DNMT3A^R882H^ in pre-leukemic HSCs results in normalization of aberrant self-renewal

Functional studies in murine models^10,13,14^ and descriptive studies in human clonal hematopoiesis^15^ suggest that the *DNMT3A^R882H^* missense mutation induces increased self-renewal in HSCs. To prospectively interrogate whether *DNMT3A^R882H^* is continuously required for increased self-renewal in patient-derived HSCs, we utilized CRISPR/Cas9 in combination with recombinant AAV6 to perform allele-specific correction of the *DNMT3A^R882H^* missense mutation in pre-leukemic HSCs isolated from AML patient specimens. We hypothesized that correction of the mutant allele would promptly normalize HSC self-renewal, thus demonstrating that *DNMT3A^R882H^* directly drives and is continuously required for this phenotype. Building on our previous approaches to *introduce* mutations into primary human hematopoietic stem and progenitor cells (HSPCs),^16–21^ we developed an approach to *revert* existing mutations directly in mutant, patient-derived cells. First, we used our published sorting strategy to isolate rare pre-leukemic HSCs from AML patient samples that contain the *DNMT3A^R882H^* mutation, but lack the leukemic *NPM1c* and/or *FLT3-ITD* mutations.^12,22^ After isolation by flow cytometry, cells were electroporated with an R882H-specific single guide RNA (sgRNA) pre-complexed with Cas9 protein and transduced with recombinant adeno-associated vector (rAAV6) encoding either the wildtype *DNMT3A^R882^* codon (correction) or, as a control, the mutant *DNMT3A^R882H^* codon (re-mutation, **Figure 1A, S1**). Each donor vector also encodes for a fluorescent marker driven by a separate promoter to allow for purification and tracking of successfully engineered cells (**Figure 1B**). Importantly, since we employ an R882H-specific sgRNA, only cells containing the DNMT3A^R882H^ mutation are subjected to Cas9-induced double stranded DNA break and repair, and only on the mutant allele. We achieved editing efficiencies of ∼5-20% in cells from several primary samples (**Figure 1C**) and single-cell derived colonies from either re-mutated (*DNMT3A^WT/R882H^*) or corrected (*DNMT3A^WT/WT^*) HSPCs expressed the appropriate fluorescent marker (**Figure 1D**). We validated on-target integration (**Figure S2A**), in-frame expression of the corrected or re-mutated cDNA (**Figure S2B**), and that our edited pre-leukemic HSCs lacked mutations found in the leukemic fraction (*NPM1c*, *FLT3-ITD*; **Figure S2C**). Taken together, this approach demonstrates that *DNMT3A^R882H^* can be corrected in pre-leukemic HSCs, and we sought to functional characterize the corrected cells.

**Fig. 1:**
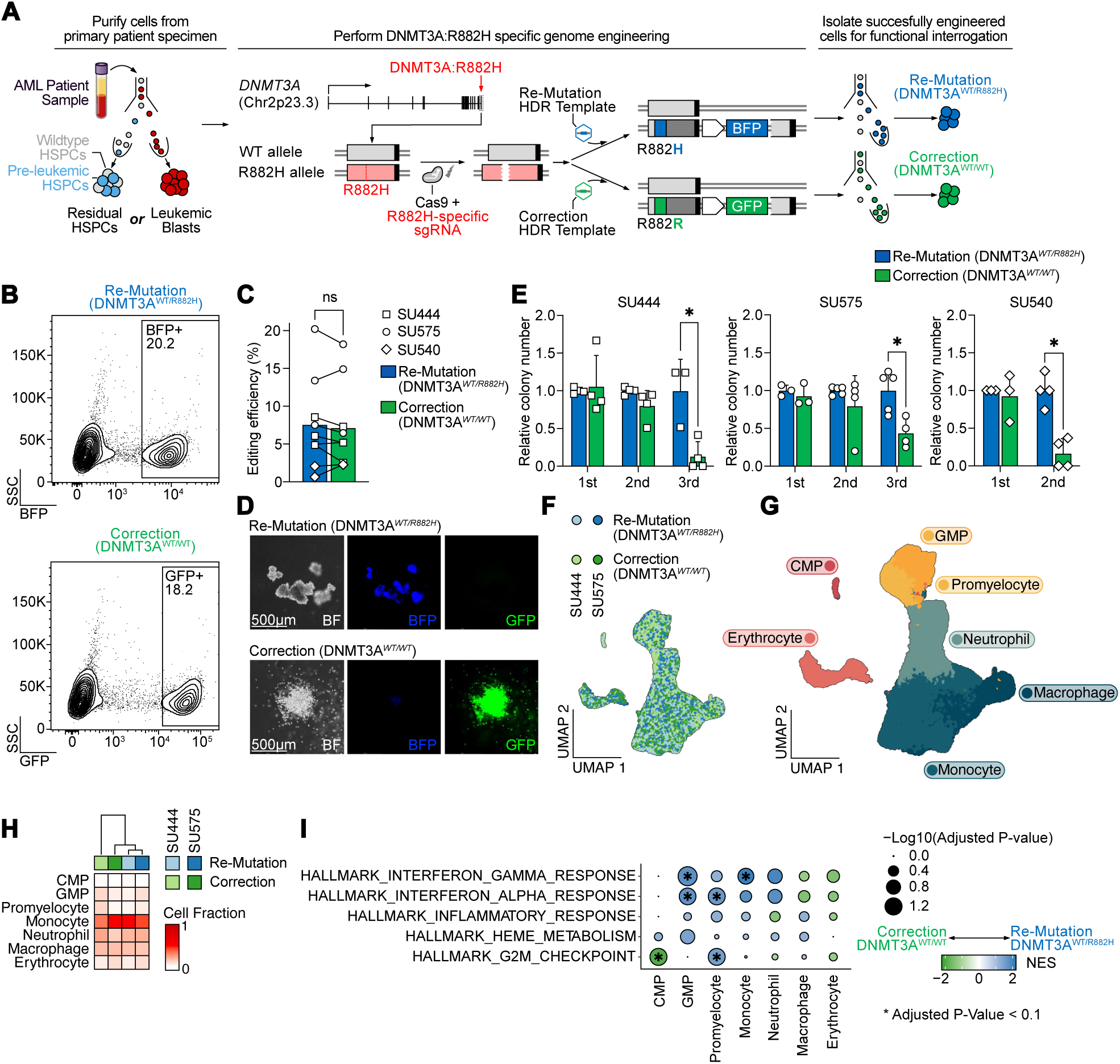
Correction of *DNMT3A^R882H^* in pre-leukemic HSCs results in normalization of aberrant self-renewal. A: Schematic of *DNMT3A^R882H^* correction approach. Cell populations of interest (either residual HSPCs containing pre-leukemic HSPCs or leukemic blasts) are purified by flow cytometry. Subsequently, pre-complexed Cas9 and sgRNA targeting the mutant, R882H allele, are electroporated into the target cells and cells are transduced with rAAV6 containing either the codon-optimized mutant (R882H) or wildtype (R882R) sequence, and a selectable fluorescent marker (BFP or GFP). Successfully engineered cells are then isolated for functional experiments. B: Isolation by FACS of successfully edited residual HSPCs (from patient SU575) for either re-mutation (top panel) or correction (bottom panel) of *DNMT3A^R882H^*. C: Editing efficiency across all experiments in residual HSPCs as percentage of fluorescent protein positive cells (n=3 independent patient specimens). D: Representative hematopoietic colonies 14 days after plating into semisolid media (Methocult). E: Serial replating assay of gene-edited residual HSPCs derived from three independent patient specimens (n=3-4 technical replicates per patient), * P < 0.05. F: UMAP embedding of single-cell RNA-Seq from gene-edited residual HSPCs differentiated in vitro for 14 days after editing colored by genotype and donor. G: UMAP embedding as in F colored by assigned cell type. H: Relative abundance of cell types within each sample. Sample identities and *DNMT3A* genotypes are indicated at the top. I: Gene set enrichment analysis of Hallmark gene sets between re-mutation (*DNMT3A^WT/R882H^*, blue) and correction (*DNMT3A^WT/WT^*, green) across cell types. * P < 0.1.

We first performed a serial colony replating assay to determine whether correction of *DNMT3A^R882H^* alters pre-leukemic HSC self-renewal. During the first plating, gene-edited pre-leukemic cells showed no difference in colony number. However, after serial replating, cells with *DNMT3A^R882H^* showed a clear and consistent replating advantage compared to gene-corrected *DNMT3A^WT/WT^*HSPCs (**Figure 1E**). Next, to prospectively elucidate the effect of *DNMT3A^R882H^* on mature hematopoietic cells, we performed single-cell RNA-sequencing on the progeny of corrected or re-mutated pre-leukemic HSCs from two patients differentiated *in vitro* for 14 days into the erythroid and myeloid lineages. After quality control, 58,036 cells were available for downstream analysis (**Figure 1F**). We assigned cell types based on known marker genes (**Figure 1G**, Methods) and quantified relative abundance of genotypes within each cell type. Here, we noted no major differences in cell abundances between corrected and re-mutated *DNMT3A^R882H^* (**Figure 1H**), consistent with the lack of large effects on hematopoietic differentiation observed in mouse models^10,11^ and individuals with clonal hematopoiesis harboring this mutation. However, gene set enrichment analysis within individual cell types revealed several differences. First, we observed an enrichment across early myelo-erythroid progenitors for heme metabolism in re-mutated *DNMT3A^R882H^* cells, in line with previous observations of erythroid lineage bias of *DNMT3A*-mutant HPSCs.^15,23^ Next, we observed a strong enrichment of gene sets linked to inflammation, particularly response to interferon both in myeloid-lineage committed progenitors (granulocyte/monocyte progenitors and promyelocytes) and in mature monocytes and neutrophils (**Figure 1I**), as has been observed in patients with clonal hematopoiesis and zebrafish models.^24–27^ Notably, however, this was not the case in the earliest progenitors detected in this assay (common myeloid progenitors, CMP), which instead showed a marked de-enrichment of the G2M checkpoint signature in re-mutated (*DNMT3A^WT/R882H^*) cells, in line with observations in murine models^11^ and our functional observation that stem cell features, such as increased replating capacity, are lost upon correction of *DNMT3A^R882H^*.

Taken together, these data support the idea that the *DNMT3A^R882H^*mutation both drives and is continuously required for increased self-renewal in human pre-leukemic HSCs. Importantly, our approach demonstrates that correction of this mutation rapidly eliminates this phenotype, within weeks. Furthermore, this approach prospectively demonstrates the relationship between *DNMT3A^R882H^* and aberrant HSC self-renewal, increased expression of erythroid programs in HSPCs, as well as a hyperinflammatory state in late progenitors and mature myeloid cells which previously had been noted in descriptive or murine studies.

### Correction of DNMT3A^R882H^ in primary AML blasts does not eliminate leukemogenicity

Next, we aimed to determine whether *DNMT3A^R882H^* is still required for the maintenance of established leukemia. We isolated leukemic blasts from six patient specimens with *DNMT3A^R882H^*, as well as one specimen with *DNMT3A^R882C^*, and performed either re-mutation or correction as described above (**Figure 2A**). Importantly, unlike pre-leukemic HSCs, these cells carry additional mutations recurrently found in AML, such as in *NPM1*, *FLT3*, *IDH1/2*, and *TET2* (**Figure 2B**). Notably, the correction efficiency varied between patient samples and ranged from 2-38% (**Figure 2C-D**). Next, we performed primary transplantation of FACS-purified, edited leukemic blasts into irradiated NSGS mice and assessed human leukemic engraftment 12-16 weeks later (**Figure 2E**). Strikingly, correction of *DNMT3A^R882H^* did not eliminate leukemogenicity, with equal levels of human leukemic engraftment regardless of *DNMT3A* genotype (**Figure 2F**). We confirmed that co-mutation patterns were preserved in the engrafted, edited cells, determining that all mutations, except for *DNMT3A^R882H^*, were present (**Figure 2G, S3A**). Finally, we confirmed that our editing approach correctly replaced the endogenous, mutant allele with either the inserted wildtype *DNMT3A^R882^* codon (correction) or the inserted mutant *DNMT3A^R882H^* codon (re-mutation) while leaving the endogenous wildtype-allele unaltered (**Figure S3B-D**).

**Fig. 2:**
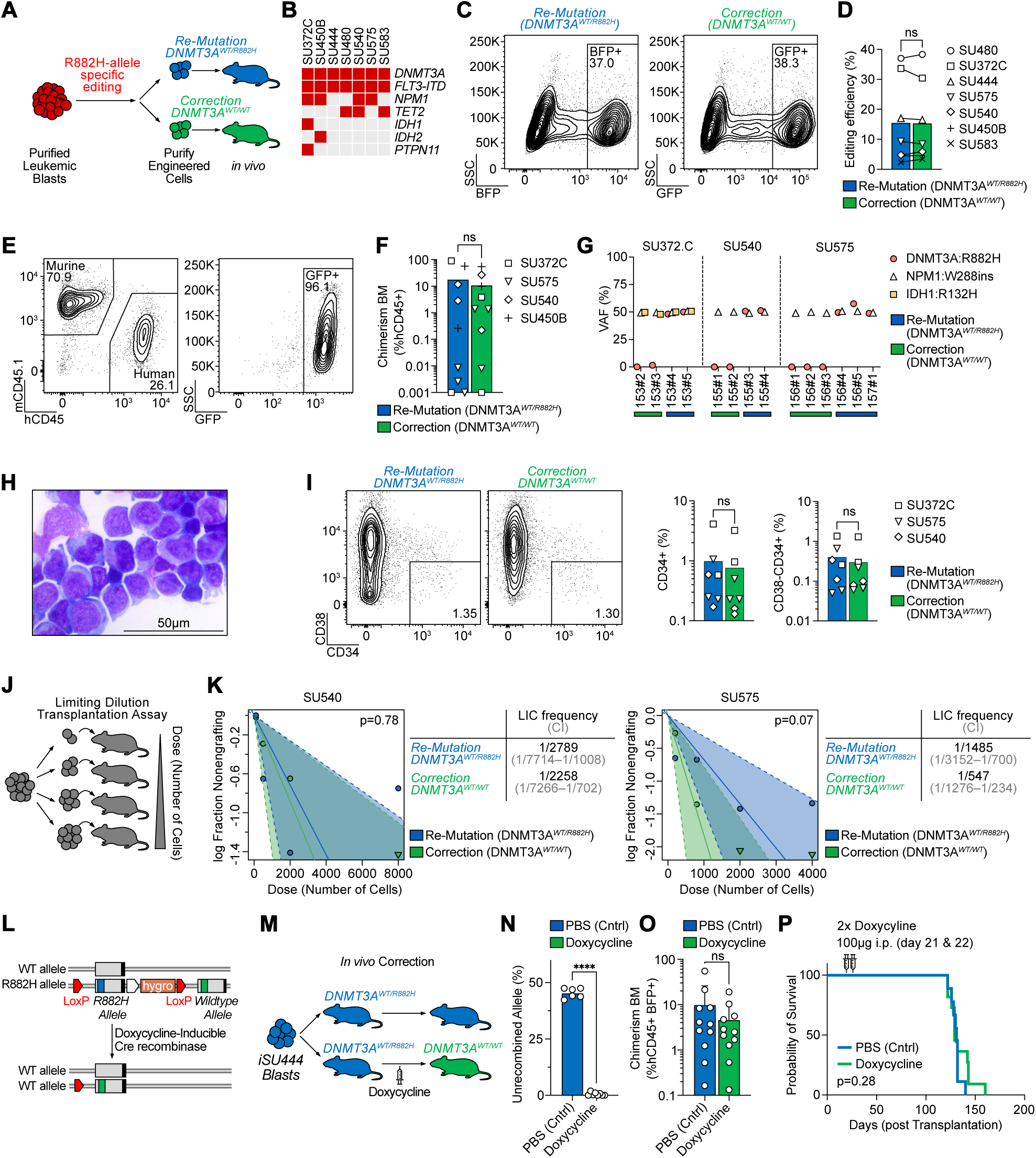
Correction of *DNMT3A^R882H^*in AML blasts has no effect on leukemogenicity or leukemia disease maintenance. A: Schematic of leukemic blast editing and xenotransplantation. B: Oncoprint of samples included in this study. C: Isolation of successfully edited leukemic blasts for either re-mutation (left panel) or correction (right panel) of *DNMT3A^R882H^*from patient SU480. D: Editing efficiency across patient specimens quantified as percentage of fluorescent protein positive cells 3 days after gene-editing. E: Representative flow cytometry analysis of bone marrow aspirate 12 weeks after injection of human, gene corrected (*DNMT3A^WT/WT^*) leukemic blasts showing human engraftment with retention of the GFP fluorescent protein. F: Human chimerism in bone marrow aspirates at 12 weeks after injection of the indicated patient sample with either *DNMT3A* re-mutation or correction. G: Variant allele frequency (VAF) of somatic mutations found in the indicated patient specimens in human genomic DNA extracted from mice 4 months after engraftment indicating clonal retention of all somatic variants, except for *DNMT3A* in *DNMT3A*-corrected specimens. H: Representative Giemsa staining of cytospin of human leukemic cells isolated 4 months after *DNMT3A^R882H^*correction. I: Representative flow cytometry plots (left) and quantification (right) of CD34+ and CD34+CD38- leukemic subpopulations 4 months after re-mutation or correction of *DNMT3A^R882H^*. J: Schematic of limiting dilution transplantation assay. K: Quantification of leukemia initiating cell (LIC) frequency for patient specimens SU540 (left) and SU575 (right) after re-mutation or correction of *DNMT3A^R882H^*. Estimated frequency of LIC and test statistics are indicated. L: Schematic of doxycycline-inducible Cre recombinase driven correction of the endogenous *DNMT3A^R882H^* allele in iPSC-derived AML blasts (iSU444). M: Schematic of *in vivo* correction experiment. Mice are engrafted with mutant (*DNMT3A^WT/R882H^*) leukemic blasts, and administration of doxycycline 3 weeks after engraftment leads to correction of *DNMT3A^WT/R882H^* to *DNMT3A^WT/WT^ in vivo*. N: Quantification of successful *in vivo* recombination of *DNMT3A^WT/R882H^*, P < 0.0001. O: Human Chimerism 5 weeks after *in vivo DNMT3A^R882H^*correction. P: Kaplan-Meier survival curve of mice with or without *in vivo DNMT3A^R882H^* correction.

Phenotypically, correction of *DNMT3A^R882H^* in leukemic blasts did not alter the morphological appearance (**Figure 2H**) or immunophenotype (**Figure 2I**) assessed in engrafted cells 4 months after gene-correction. To further assess the ability of *DNMT3A*- corrected cells to initiate leukemia, we determined the leukemia-initiating cell (LIC) frequency from two patient samples by performing limiting dilution transplantation assays with re-mutated or corrected leukemic blasts (**Figure 2J**). Similar to engraftment levels, we found no difference in LIC frequency between the re-mutated (*DNMT3A^WT/R882H^*) and corrected (*DNMT3A^WT/WT^*) genotypes (**Figure 2K**). Taken together, our approach allows for the selective and specific correction of *DNMT3A^R882H^*missense mutation in primary human AML blasts and demonstrates that correction of this mutation does not impair leukemic engraftment potential, human chimerism, immunophenotype, or leukemia-initiating cell frequency upon primary transplantation.

### In vivo correction of DNMT3A^R882H^ shows no effect on leukemia disease maintenance

In our experiments above, gene-correction of the *DNMT3A^R882H^* allele was performed *ex vivo*, and re-mutated or corrected leukemic blasts were transplanted into NSGS mice. Since this approach cannot delineate between the effect of *DNMT3A^R882H^* on initial engraftment and subsequent maintenance/proliferation, we established an approach where gene correction of *DNMT3A^R882H^* was inducibly performed *in vivo* after the engraftment of human leukemia. For this approach, we utilized an AML-derived induced pluripotent stem cell (iPSC) model, which allows for more extensive genome engineering. We and others have previously demonstrated that iPSCs derived from human AML can be re-differentiated into a transplantable human leukemia that re-acquires both phenotypic and epigenetic features of the original patient sample.^28,29^ To assess the role of *DNMT3A^R882H^* in leukemic maintenance, we established an AML-derived iPSC line (iSU444) that carries the *DNMT3A^R882H^* mutation along with *FLT3-ITD*, but otherwise has a normal karyotype (**Figure S4A**). These AML-derived iPSCs were then differentiated into HSC-like cells,^30^ and then xenotransplanted into mice, whereupon they developed into leukemic cells *in vivo* (**Figure S4B**). Using this line, we engineered LoxP-sites flanking the mutant (R882H) exon 23 of *DNMT3A* and introduced a corrected (WT) exon 23 downstream. Further, we engineered these cells to express doxycycline-inducible Cre recombinase. These cells are mutant (*DNMT3A^WT/R882H^*) at baseline, but upon treatment with doxycycline will recombine to correct the endogenous *DNMT3A* mutant allele to wildtype (*DNMT3A^WT/WT^*, **Figure 2L**). We transplanted leukemic blasts derived from iSU444 engineered with this *DNMT3A* correction cassette into NSGS mice and waited for 3 weeks to allow for initial engraftment. Then, half of the mice were injected with two doses of doxycycline and recombination rates, human chimerism, immunophenotype, and survival were monitored (**Figure 2M**). 5 weeks after injection of doxycycline, we obtained bone marrow aspirates from these mice. Using a droplet digital PCR (ddPCR) strategy to quantify the recombination rate (**Figure S5A**), we found near complete recombination rates of the *DNMT3A* allele in the human leukemic cells (**Figure 2N**) while observing no difference in human leukemic engraftment (**Figure 2O**) or difference in the percentage of CD34+ human leukemic cells (**Figure S5B**). 14-18 weeks after *in vivo* correction, mice became moribund from fulminant leukemia, showing no difference in survival compared to mice carrying mutant control (*DNMT3A^WT/R882H^*) cells (**Figure 2P**). Of note, we confirmed the presence of *FLT3-ITD* (**Figure S5C**) and lack of the mutant *DNMT3A^R882H^* allele in most of these mice upon sacrifice (**Figure S5D**). Taken together, *in vivo* correction of *DNMT3A^R882H^* showed no difference in human leukemic engraftment, phenotype, or overall survival, further demonstrating that *DNMT3A^R882H^*is not required for maintenance of established AML.

### Secondary transplantation reveals a reduction of leukemic stem cell frequency upon DNMT3A^R882H^ correction

Leukemic stem cells (LSCs) are a fraction of leukemia cells that exhibit stem cell features, including the ability to self-renew and to generate downstream progeny that are detected through secondary transplantation.^31^ In **Figure 2**, we assessed the ability of leukemia cells to initiate disease in primary recipient mice and saw no difference between re-mutated (*DNMT3A^WT/R882H^*) and corrected (*DNMT3A^WT/WT^*) primary leukemic blasts, including in leukemia-initiating cell (LIC) frequency. To assess whether *DNMT3A^R882H^*influences the frequency of LSCs, we performed secondary transplantation of leukemic cells from primary recipient mice 4 months after transplantation (**Figure 3A**). Specifically, we performed secondary limiting dilution transplantation assays of re-mutated (*DNMT3A^WT/R882H^*) and corrected (*DNMT3A^WT/WT^*) leukemic blasts for two cases. Mice were injected with decreasing doses of either re-mutated or corrected leukemic cells harvested from primary recipient mice and engraftment was assessed 4 months later. Strikingly, the LSC frequency was significantly reduced in cells that had been corrected (*DNMT3A^WT/WT^*) compared to the re-mutated control (LSC frequency 1/216 in re-mutated versus 1/23,572 in corrected, **Figure 3B**, and 1/10,879 in re-mutated versus undetected in corrected, **Figure 3C**). We confirmed presence of the other clonal mutations in the human leukemia cells engrafted in these mice (**Figure 3D-E**). Taken together, we demonstrate that the frequency of leukemia stem cells is decreased upon correction of the *DNMT3A^R882H^*mutation in established leukemia, while leukemia engraftment, leukemia-initiating cell frequency, and disease maintenance are not dependent on the persistence of the *DNMT3A^R882H^* mutation.

**Fig. 3:**
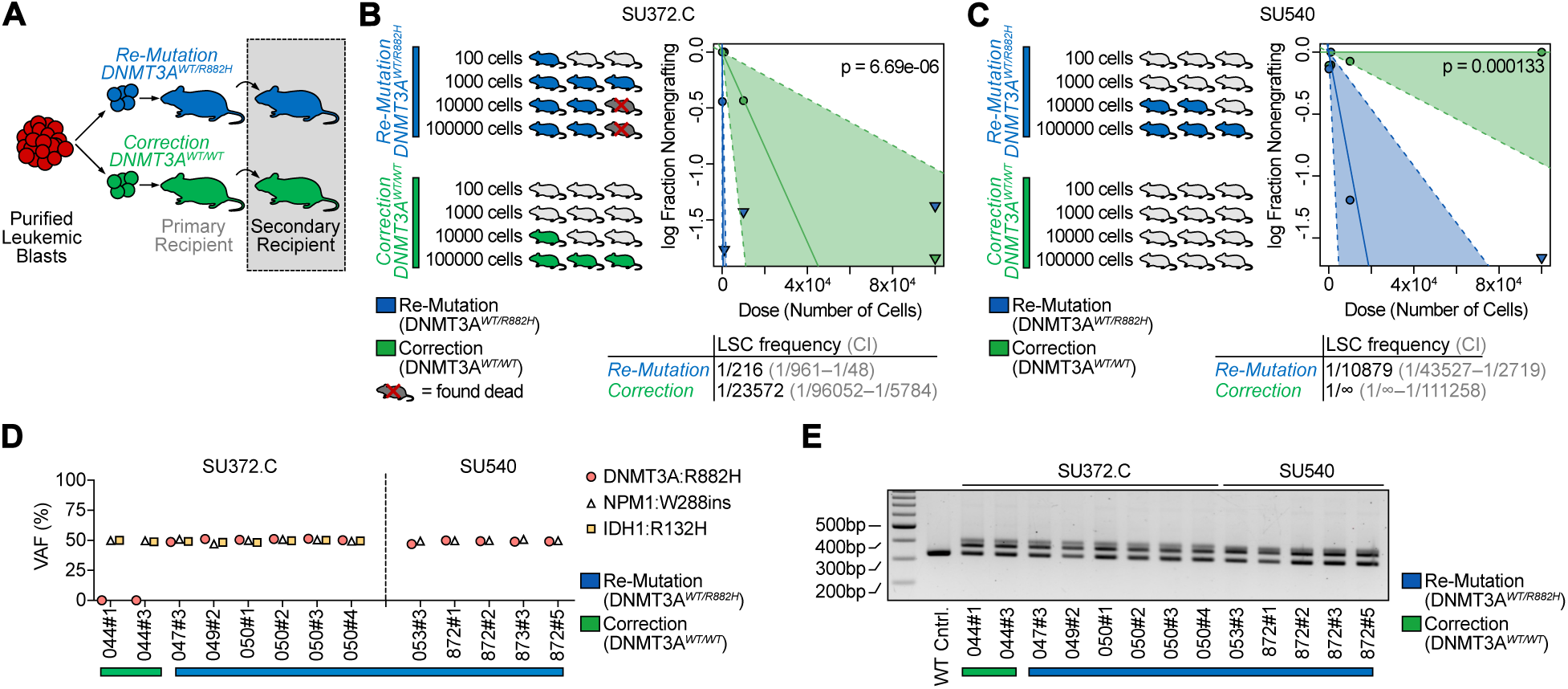
Secondary transplantation reveals a reduction of leukemic stem frequency upon *DNMT3A^R882H^* correction. A: Schematic of secondary transplantation experiments. Human leukemic cells are harvested from primary recipients 4 months after gene-editing and then transplanted in limiting dilution into secondary host animals. B: Quantification of secondary limiting dilution assay results for patient specimen SU372.C. Estimated LSC frequencies and test statistics are indicated. C: Quantification of secondary limiting dilution assay results for patient specimen SU540. None of the mice receiving corrected (*DNMT3A^WT/WT^*) leukemic cells showed engraftment at the doses indicated. Estimated LSC frequencies and test statistics are shown. D: VAF in human genomic DNA isolated from secondary recipients for recurrent somatic mutations found in patient samples utilized for secondary transplantation studies. E: *FLT3-ITD* PCR from human genomic DNA isolated from secondary recipients.

### Epigenetic and transcriptional identity is highly patient-specific with only minimal changes upon correction of DNMT3A^R882H^

The R882H missense mutation in *DNMT3A* is thought to exert a dominant-negative effect on DNA methyltransferase activity, resulting in CpG hypomethylation.^15,32^ However, recent murine studies have suggested that the DNA methylation landscape can be restored upon overexpression of *DNMT3A* in *DNMT3A*-mutant cells.^33,34^ Inspired by this previous research, we performed whole-genome bisulfite sequencing (WGBS) and transcriptomic analysis (RNA-Seq) in leukemic blasts 4 months after re-mutation or correction of *DNMT3A^R882H^* (**Figure 4A**). Global genome-wide CpG methylation levels were similar across three patient specimens between *DNMT3A* corrected or *DNMT3A* re-mutated cells (**Figure 4B**). Indeed, when assessing methylation patterns across the genome by principal component analysis (PCA), we found that samples from each patient grouped closely, with no detectable influence of *DNMT3A* genotype in the first two dimensions, although there was partial separation in PC3, which accounts for 4.7% of the variance (**Figure 4C**). Next, we identified differentially methylated regions (DMRs) between *DNMT3A* re-mutated and *DNMT3A* corrected samples within each patient specimen using dmrseq^35^ (Methods). We detected DMRs in the three patients analyzed based on 2-3 biological replicates in each, with the number of significant DMRs between *DNMT3A* re-mutated and corrected samples in a given patient ranging from 5611 to 13,331 and encompassing 4.0 to 12.4 Mb of the genome. Correction of *DNMT3A^R882H^* led to increased methylation at 94-99% of these loci (**Table S3-6, Data S1-3**), an effect that was consistent across patient samples and replicates (**Figure 4D**). DMRs were highly patient-specific, with only 89 genes containing DMRs in all three patients; however, this set included genes involved in leukemia, hematopoiesis, and multipotency, such as *FLI1*, *EGFL7*, and *TCF15* (**Table S7**).^36–40^ We then analyzed the genomic features associated with DMRs across our cohort (**Figure 4E, S6A**). DMRs showed enrichment over CpG islands, shores, and shelves (**Figure S6B**).

**Fig. 4:**
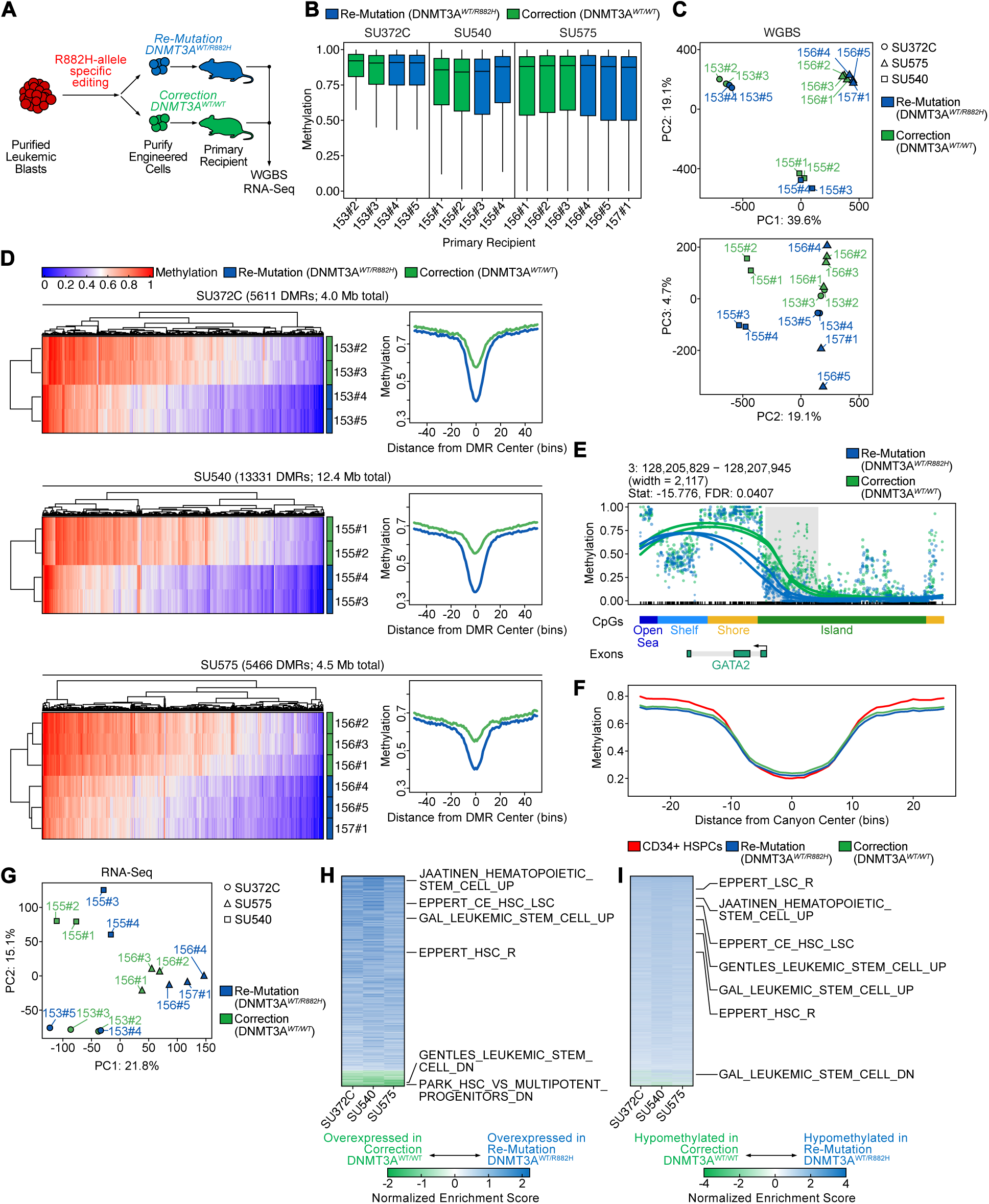
Epigenetic and transcriptional identity is highly patient-specific with only minimal changes upon correction of *DNMT3A^R882H^*. A: Schematic of sample generation for whole-genome bisulfite (WGBS) and RNA-Sequencing (RNA-Seq). Human cells were harvested from recipient mice 4 months after gene-editing. B: Average CpG methylation across the human genome for three patient specimens with either correction (green) or re-mutation (blue). Each bar represents an individual sample harvested from a primary recipient mouse (mouse ID indicated on x-axis). C: Principal component analysis of global CpG methylation from engrafted mice. Patient ID is indicated by shape and *DNMT3A* genotype is indicated by color. D: Methylation at differentially methylated loci (heatmaps, left) for all DMRs identified within each patient sample as well as a summary meta-region plot of methylation at DMRs normalized for width (right). DMRs in meta-region plots are extended by 2 kb for visualization of flanking regions. E: Methylation at a representative DMR (adjacent to *GATA2*) for patient sample SU372.C. Individual lines indicate cells harvested from separate mice, color indicates the *DNMT3A* genotype. F: Summary meta-region plot of methylation at methylation canyon loci from Jeong et al.^41^ for re-mutated (blue) or corrected (green) patient samples, as well as healthy CD34+ HSPC control (red). G: Principal component analysis of global RNA- Seq patterns from engrafted mice. Patient ID is indicated by shape and *DNMT3A* genotype is indicated by color. H: Gene set enrichment analysis of differential gene expression within each patient plotted as a heatmap of normalized enrichment scores (NES) sorted by descending average. Gene sets enriched in re-mutated (*DNMT3A^WT/R882H^*) are colored in blue, gene sets enriched in corrected (*DNMT3A^WT/WT^*) are colored in green. Representative gene sets relating to hematopoietic or leukemic stem cell function are highlighted. I: Gene set enrichment analysis of differential promoter methylation within each patient as a heatmap of NES sorted by descending average. Gene sets enriched in hypomethylated DMRs in re-mutated (*DNMT3A^WT/R882H^*) samples are colored in blue, and gene sets enriched in hypermethylated DMRs in re-mutated samples are colored in red. Representative gene sets relating to hematopoietic or leukemic stem cell function are highlighted.

Since *DNMT3A* mutations have previously been associated with erosion of the borders of large, normally hypomethylated genomic regions (“methylation canyons”),^41^ we next interrogated whether we could detect changes at these loci. We first identified human genomic regions analogous to these murine canyons (Methods). Next, we assessed the methylation levels over canyon regions between leukemic patient samples with re-mutated or corrected *DNMT3A^R882H^*, as well as healthy donor-derived normal CD34+ HSPCs (**Figure 4F, S6C**). Over these regions, correction of *DNMT3A^R882H^* did not consistently induce re-normalization of methylation levels towards the healthy-donor derived CD34+ HSPC reference. Only 6-21% of canyons displayed differential methylation between re-mutation and correction of *DNMT3A^R882H^*, indicated by presence of a DMR in the annotated canyon, and only 1-4% of all DMRs lay within canyon regions (**Figure S6D, Table S8, Data S4-7**). These results are in stark contrast to the methylation changes seen in normal HSCs upon knockout of *DNMT3A*, where 75% of canyons display altered DNA methylation, and 30% of DMRs are located within canyons or canyon edges.^41^

We previously reported that hypomethylated blocks are a nearly universal feature across many cancer types, including colon, lung, breast, thyroid, and pancreatic cancer, as well as in premalignant lesions and EBV-transformed B-cells, and can range in size from 5 kb to over 2 Mb.^42–44^ CpG islands contained in blocks show flattening of methylation profiles, resulting in both the hypermethylation of islands and hypomethylation of shores and shelves characteristic of altered DNA methylation in cancer, suggesting that this disruption of methylation around CpG islands is associated with the loss of structural integrity of heterochromatin in blocks.^42^ We observed a striking relationship between these previously defined blocks and canyons, where canyon regions corresponded with the edges of blocks identified in EBV-transformed B-cells (**Figure S6E, Data S7**). Blocks of hypomethylation have not yet been reported in myeloid tissues, so we selected blocks identified in EBV-transformed B-cells, another hematopoietic cell type, for this comparison. Next, we searched for blocks between leukemic blasts with re-mutated or corrected *DNMT3A^R882H^*. Putative blocks were identified using minimum sizes of 5 kb, as originally defined, and 3.5 kb, to encompass smaller canyon regions (Methods). Correction of *DNMT3A^R882H^* did not induce large-scale restoration of methylation, with no putative block regions having significant differences in methylation (**Table S9-14**).

Finally, we determined whether correction of *DNMT3A^R882H^*affects the transcriptome of AML by performing RNA-Seq on the same specimens utilized for methylome analysis. Similarly to DNA methylation, the global transcriptome primarily clustered by patient identity, rather than *DNMT3A* genotype (**Figure 4G**). To elucidate whether common transcriptional programs were modified upon *DNMT3A* correction, we identified differentially expressed genes and performed gene set enrichment analysis on a per-patient basis (**Figure 4H**). Here, we found that correction of *DNMT3A^R882H^* consistently reduced the expression of gene sets associated with hematopoietic and leukemic stem cell function, in line with our functional observation of reduced LSC frequency with correction of *DNMT3A^R882H^*. We then aimed to correlate changes in DNA methylation and gene expression between *DNMT3A^R882H^* re-mutation and correction. We plotted the fold change in gene expression against the differences in methylation over putative promoter DMRs between re-mutated and corrected *DNMT3A^R882H^* (**Figure S6F**). While correlation between gene expression and methylation was modest (Pearson’s R −0.02 to −0.04 per patient), correction of *DNMT3A^R882H^* consistently led to increased methylation over genes associated with HSC and LSC function, linking DNA methylation to gene expression (**Figure 4I**).

Our analysis of the methylome and transcriptome in leukemic blasts revealed that correction of *DNMT3A^R882H^* shows relatively modest influence on epigenetic maintenance in established AML. Both the methylome and transcriptome were highly patient-specific and showed retention of leukemia-specific patterns 4 months after correction of the *DNMT3A^R882H^*mutation. However, in line with our functional experiments in secondary transplantation studies, we observed an enrichment of hematopoietic and leukemic stem cell programs in re-mutated versus corrected leukemic blasts.

## DISCUSSION

During pre-malignant evolution, driver mutations confer a competitive advantage to individual clones within healthy tissue,^45^ which then may acquire additional mutations to progress into frank malignancy. Several of these driver mutations found in pre-malignant stages of cancers have been identified,^46^ such as biallelic inactivation of *APC* in colorectal cancer,^47^ mutations in *TP53* and *CDKN2A* in Barrett’s esophagus, melanoma, and pancreatic cancer,^48,49^ and *BRCA1* and *BRCA2* in breast and ovarian cancer.^50^ In pre-leukemic clonal hematopoiesis, the expansion of clones carrying driver mutations has similarly been associated with an increased risk of developing hematologic cancers.^51–53^ However, our understanding whether mutations acquired early during cancer evolution are required for disease initiation, maintenance of established disease, or both, is limited. While mouse models of driver mutations such as *JAK2^V617F^*, *IDH2^R140Q^* or *APC* loss have suggested that these drivers are continuously required for disease maintenance,^1–3^ it is unclear whether this extends to other pre-malignant drivers, particularly those associated with clonal hematopoiesis.

We focus on the *DNMT3A^R882H^* missense mutation, which is the most frequent *DNMT3A* missense mutation in human AML,^4,54^ and is also the most frequently affected gene in clonal hematopoiesis.^6,55,56^ The effects of mutant *DNMT3A* have been intensely studied in the context of hematopoiesis, where mutant *DNMT3A* confers increased self-renewal in HSCs^13,23^ and a hyperinflammatory phenotype in mature myeloid cells.^25^ However, in leukemic cells, the role of *DNMT3A^R882H^* is less clear. Overall, mutations in DNMT3A, including *DNMT3A^R882H^*, are associated with worse prognosis in AML,^4,57^ however, both co-mutation patterns, as well as clinical attributes such as age, are also associated with *DNMT3A* mutations. Therefore, despite being one of the most frequently mutated genes in human AML, current clinical risk assessment does not take presence or absence of *DNMT3A* mutations into account for treatment decisions^58^ and further investigation to delineate the specific contribution of *DNMT3A* mutations is warranted.

Cancer cell lines and model organisms have proven to be useful tools to investigate cancer biology and develop novel treatments. However, prospective, functional experiments with primary human cancers and cancer cells are critical to advance our understanding of human cancer biology. In cancer genetics, interrogation of individual somatic mutations in primary patient specimens is inherently challenging, as it is difficult to control for the combination of co-mutation patterns and germline background. In this study, we approach this challenge by employing tools derived from advances in gene therapy to cancer genetics. We aim to delineate the specific contribution of *DNMT3A^R882H^*, clarifying whether the presence of this mutation is necessary once leukemia has been established through prospective, controlled genetic experiments directly in primary human leukemia specimens, avoiding surrogate systems for human cancer genetics. Using primary patient samples, we selectively and specifically revert *DNMT3A^R882H^* back to wildtype, and demonstrate that correction has a pronounced effect in pre-leukemic HSPCs that rapidly lose features of aberrant self-renewal and mature myeloid cells that lose a hyperinflammatory transcriptional profile. However, in established disease, correction of *DNMT3A^R882H^* does not affect leukemic engraftment potential, human chimerism, immunophenotype, or leukemia-initiating cell frequency. Nor does it affect leukemia disease maintenance when inducibly corrected in engrafted leukemia cells. Notably, correction of *DNMT3A^R882H^* does result in a decrease in leukemia stem cell frequency, indicating more subtle effects in established disease. Finally, we demonstrate that reversion of the methylation landscape upon correction of *DNMT3A^R882H^* is minimal, with global methylation levels unchanged 4 months after gene correction. Our results additionally clarify the relationship between DNA methylation canyons identified in normal hematopoietic tissues and blocks of hypomethylation identified in cancer, demonstrating that canyons correspond to the edges of blocks.

This data supports the concept that *DNMT3A^R882^* mutations are only required during disease initiation, acting as landscaping mutations upon which secondary mutations can act. Importantly, however, once disease transformation has occurred, *DNMT3A^R882^* mutations become non-essential, where correction does not immediately affect disease maintenance. This stands in stark contrast to other mutations such as *JAK2^V617F^* or *IDH2^R140Q^*in leukemia, or *APC* in colorectal cancer, where previous studies have suggested a continued requirement for disease maintenance.^1–3^

Taken together, our data highlights that prospective, genetic experiments in primary patient samples are feasible, and reveal a surprising stability of leukemic phenotypes and methylation patterns months after reversion of a pre-leukemic driver mutation of human AML. By demonstrating the context-specific contribution of a driver mutation in human cancer, our experiments also have important therapeutic implications. Specifically, therapeutic targeting of DNMT3A^R882^ missense mutations might be most effective during pre-leukemic evolution and less effective in established disease. We envision that similar approaches to elucidate the interplay of driver mutations in cancer and improve the discovery of novel therapies.

## METHODS

### Primary Human Samples

AML patient samples were obtained from patients with AML, with informed consent according to institutional guidelines (Stanford University Institutional Review Board No. 6453). Mononuclear cells were isolated by Ficoll-Paque PLUS (GE Healthcare) density gradient centrifugation, red blood cells were removed by ACK lysis (ACK Lysing Buffer, Thermo Fisher Scientific. Samples were resuspended in 90% FBS + 10% DMSO (Sigma-Aldrich) and cryopreserved in liquid nitrogen for future use.

### Purification of cell populations from primary human AML samples

Purification of residual HSCs from AML specimens was performed as previously described.^12^ Briefly, AML specimens were thawed, washed, stained and sorted on a FACSAria II SORP (BD). The residual HSC fraction was defined as CD3-/CD19-/CD34+/CD38-/CD99-/TIM3- and AML blasts were sorted as CD3-/CD19-/CD14-. FACS staining panels can be found in Supplementary Table 1.

### iPSC culture and derivation

The iSU444 iPSC line was established from purified AML patient blasts as described previously,^28^ with some modifications. AML blasts were cultured at 37°C, 5% CO2 and 5% O2 in AML pre-culture media comprised of MyeloCult H5100 (STEMCELL Technologies) base media supplemented with 20 ng/mL each of TPO, SCF, FLT3L, G-CSF, GM-CSF, IL-3 and IL-6 (Peprotech) for three days. Cell viability was confirmed to be >70% at the time of transduction with reprograming factors. AML blast transgene-free reprogramming was performed using Sendai virus containing SeVdp(KOSM)302L as previously described,^59^ with some modifications. Cells were cultured in AML pre-culture media at a density of 0.5×10^6^ cells/mL (0.5×10^5^ cells per 100 μl) in a 96 well plate. On day 0, SeVdp(KOSM)302L Sendai virus was added at an MOI of 10 with cells kept at 37°C at 5% O2 for 12 hr. After infection, the cell suspensions were collected, diluted with 900 µl of pre-culture medium, and centrifuged at 300 xg for 5 min at room temperature. Cells were then resuspended in a mixture of 375 µl AML pre-culture media and 375 μl of StemFit Basic04 Complete Type medium iPSCs media (Ajinomoto Co.) in a 12-well plate coated with iMatrix-511 basement membrane matrix (ReproCell). On day 3, 375 µl AML pre-culture media and 375 µl iPSCs media were gently added. On day 5, 1500 µl iPSCs media was gently added. From day 7, half media changes with 1 ml of fresh iPSCs media were performed until formation of colonies with iPSC morphology could be observed. iPSC-like colonies were manually picked and expanded individually for further analyses. iPSCs and their derivative subclones were propagated feeder-free in StemFit Basic04 Complete Type medium (Ajinomoto Co.) on cell culture plastics with iMatrix-511 basement membrane matrix (ReproCell).^60^ HSPC Differentiation from iPSCs was performed using a non-transgene 10-day-culture differentiation method as previously described ^30^, with slight modifications. Specifically, on day 4, artery endothelial cells were dissociated into a single-cell suspension (Accutase) and densely re-seeded at 500,000 cells/cm2 onto plates precoated with 10 µg/mL Vitronectin and 20 nM of high-affinity NOTCH agonist DLL4-E12 (Vincent Luca’s laboratory, Moffitt Cancer Center and Chris Garcia’s laboratory, Stanford).^61^ Cells were then further differentiated towards hemogenic endothelium in CDM3 media supplemented with Forskolin (10 µM), LIF (20 ng/mL [R&D Systems, 7734-LF-025]), OSM (10 ng/mL [R&D Systems, 295-OM-010]) and SB505124 (2 µM) for 72 hours. Hemogenic endothelium induction media was refreshed every 24 hours with a complete media change. Day 7 hemogenic endothelium cells were differentiated towards HLF+ HOXA+ hematopoietic progenitors in CDM3 media supplemented with Forskolin (10 µM), SB505124 (2 µM), OSM (10 ng/mL [R&D Systems, 295-OM-010]), LIF (20 ng/mL [R&D Systems, 7734-LF-025]), SB505124 (2 µM), SR1 (750 nM [Cellagen, C7710-5]), UM171 (75 nM [ApexBio, A8950]) and IL-1β (5 ng/mL [PeproTech, 200-01B]) for 72 hours. Media changes were performed gently to avoid disturbing the emerging semi-adherent cells. Hematopoietic progenitor induction media was refreshed every 24 hours with a complete media change on days 8 and 9; on day 10, fresh media was added without media change. On day 11, CD45+ cells were isolated by FACS (CD45 Mouse anti-Human, APC, Clone: 2D1 [BD, 340943]).

### AAV vector production

AAV vector plasmids were cloned in the pAAV-MCS plasmid containing ITRs from AAV serotype 2 (AAV2).^16^ Between the ITRs, we inserted two homology arms of 400 bp flanking the remaining, codon-optimized corrected (R882R) or re-mutated (R882H) coding sequence of the final DNMT3A exon, as well as a cassette containing the UBC promoter and either turboGFP (for DNMT3A correction) or mTagBFP2 (for DNMT3A re-mutation). For experiments with iPSC-AML, we increased the length of two homology arms to 600-729 bp and inserted a loxP site 81bp upstream of the final, mutant DNMT3A exon, followed by a UBC-Hygromycin selection cassette, the second loxP site, followed by the final, corrected DNMT3A exon. Finally, all vectors were engineered to contain distal cut sites for ITR removal contained the same sequences as the genomic sgRNA target sequences, with the PAM sequence facing inward between the ITR and homology arms.^21^ AAV6 particles were produced in 293FT cells (Thermo Fisher Scientific) transfected using standard PEI transfection with ITR-containing plasmids and pDGM6 containing the AAV6 cap genes, AAV2 rep genes and adenovirus serotype 5 helper genes (gift from David Russell - Addgene plasmid #110660).^62^ Particles were harvested after 72 h, purified using the AAVpro Purification Kit (Takara Bio) according to the manufacturer’s instructions and then stored at −80 °C until further use. Viral vector titers measured as vector genomes per microliter (vg/µl) were determined by ddPCR analysis of serially diluted vector samples using primers and probes that target the vector ITR as previously described.^63^

### CRISPR/Cas9 Editing

CRISPR/Cas9 nucleofection and rAAV6-mediated HDR were performed as previously described,^16^ with some modifications. For experiments with residual HSPCs, cells were cultured at 37°C, 5% CO2 and 5% O2 in in growth media comprised of StemSpan SFEM II (STEMCELL Technologies) base media supplemented with 20 ng/mL of TPO, SCF, FLT3L, and IL-6 (Peprotech) and 35nM of UM-171 (Selleckchem) for 48h. Synthetic, chemically modified sgRNAs targeting DNMT3A:R882H (5’- CGTCTCCAACATGAGCCACT-3’) or DNMT3A:R882C (5’-CGTCTCCAACATGAGCTGCT-3’) and Cas9 protein (Alt-R HiFi CRISPR-Cas9) were purchased from IDT. The appropriate sgRNA was precomplexed with Cas9 protein at a molar ratio of 1:2.5 at 25°C for 10 min immediately prior to electroporation. Cells were electroporated using the Lonza Nucleofector 4D (program DZ-100) using 150 µg/mL Cas9 protein, with the addition of 20 pmol siTP53^64^ in P3 Primary Cell Nucleofector Solution (Lonza). Immediately after electroporation, cells were plated with rAAV6 particles at a multiplicity of infection of 5,000 for 8h in growth media as above, supplemented with 0.5 µM AZD7648.^65^ Cells were washed and cultured in growth media for a total of 3 days after electroporation before isolation of fluorescent protein positive populations by FACS. For experiments with purified AML blasts, cells were cultured at 37°C, 5% CO2 and 21% O2 in AML media comprised of StemSpan SFEM II (STEMCELL Technologies) base media supplemented with 20 ng/mL of TPO, SCF, FLT3L, GM-CSF and IL-6 (Peprotech) and editing was performed as described above. For experiments with iPSC, editing was performed immediately after dissociation using program CB-150 (Lonza Nucleofector 4D), but otherwise as described above.

### Genotyping

Droplet Digital PCR (ddPCR) assays were performed using custom TaqMan SNP Genotyping Assays (Thermo Fisher) and ddPCR Supermix for Probes (Bio-Rad). Droplet generation and analysis were performed on the Bio-Rad QX200 Droplet Digital PCR System (Bio-Rad). FLT3-ITD and In-Out-PCR were performed using target-specific primers and SeqAmp DNA Polymerase (Takara). For RT-PCR, cDNA was generated using SuperScript VILO cDNA Synthesis Kit (Thermo Fisher), and PCR was performed using cDNA-specific primers and SeqAmp DNA Polymerase (Takara). Primer sequences can be found in Supplementary Table 2.

### Colony Formation Assays

For colony formation assays, 200-500, edited residual HSCs were added to 4 mL of MethoCult H4435 (Stemcell Technologies) and plated in triplicate. The number of colonies in each sample was scored at 12-16 days. For replating assays, cells were washed and replated 100,000 cells/plate.

### Animal Studies

All mouse experiments were conducted in accordance with a protocol by the Institutional Animal Care and Use Committee (Stanford Administrative Panel on Laboratory Animal Care #22264) and in adherence with the U.S. National Institutes of Health’s Guide for the Care and Use of Laboratory Animals.

NOD.Cg-Prkdcscid Il2rgtm1WjlTg(CMV-IL3,CSF2,KITLG)1Eav/MloySzJ (NSGS) mice were purchased from Jackson laboratory and bred in-house. Mice were housed in specific-pathogen-free animal facilities in microisolator cages. Human cells were engrafted into 6 to 8 week old mice 2 to 24h after sublethal irradiation (200 rad) by intravenous (iPSC-AML studies) or intrafemoral (primary AML studies) injection of 10- 50,000 cells. Bone marrow aspirates were obtained at the indicated time points from the right femur. For terminal engraftment analysis, mice were humanely euthanized, and femurs, tibias, hip bones, sternum, and spine were harvested and crushed. Mononuclear cells were isolated by Ficoll-Paque PLUS (GE Healthcare) density gradient centrifugation. For both bone marrow aspirates and terminal engraftment analysis, red blood cells were removed with ACK lysis (ACK Lysing Buffer, ThermoFisher Scientific), and stained for flow cytometry. For secondary transplantation, cells were sorted for human CD45 and transplanted into recipients as indicated.

### Whole-genome bisulfite sequencing

Cells were sorted from crushed whole bone marrow for Whole Genome Bisulfite Sequencing (WGBS) at the indicate timepoints. Genomic DNA was extracted from >100,000 cells using DNA/RNA microprep plus kit (Zymo) following manufacturer’s protocol. The generation of dual-indexed libraries for WGBS followed a modified protocol based on the NEBNext Ultra DNA Library Prep Kit for Illumina (New England BioLabs), in accordance with the manufacturer’s instructions. Library input was 200 ng genomic DNA, quantified by the Qubit BR assay (Invitrogen), and 1% unmethylated Lambda DNA (Promega) was added as a control for bisulfite conversion efficiency. Fragmentation of input gDNA to achieve an average insert size of 350 bp was done with a Covaris S220 Focused-ultrasonicator. Size selection was performed using SPRIselect beads, isolating fragments within the 300-400 bp range. Bisulfite conversion post-size selection was done with the EZ DNA Methylation-Lightning Kit (Zymo), following the manufacturer’s guidelines. For amplification post-bisulfite conversion, Kapa HiFi Uracil+ (Kapa Biosystems) polymerase was used under the following cycling conditions: 98°C for 45 s, followed by 9 cycles of 98°C for 15 s, 65°C for 30 s, and 72°C for 30 s, with a final extension step at 72°C for 1 min. WGBS libraries were sequenced on an Illumina NovaSeq 6000 instrument using an S4 flowcell and a 150 bp paired-end dual indexed run, with 5% PhiX library control (Illumina). Autosmal ranged from 23X to 29X. The bisulfite conversion rate for unmethylated Lambda DNA averaged 99.8%.

Sequencing reads were trimmed and quality control was performed using TrimGalore v.0.6.6 (https://doi.org/10.5281/zenodo.5127899) with default parameters. Alignment of reads to the hg19 genome was performed using Bismark v.0.23.0^66^ with default parameters. Resulting BAM files were processed with samtools v.1.18^67^ for merging, sorting, deduplication, and indexing. Finally, Bismark’s methylation_extractor was used to extract methylation calls and generate CpG_report files. Bismark CpG_report files were imported into bsseq v.1.36.0^68^ for downstream analysis.

### Methylation analysis

Global methylation levels were assessed using raw CpG methylation over chromosomes 1-22. Principal component analysis (PCA) was performed on these raw methylation values using the prcomp function from the R stats package. Feature-level CpG methylation was calculated over genomic annotations generated by the Bioconductor package annotatr v.1.22.0.^69^ Differentially methylated regions (DMRs) were identified between re-mutated and corrected samples within each patient. CpGs were filtered to include only autosomal CpG sites with at least 1x coverage in at least one sample per group. DMRs were called using Bioconductor package dmrseq v.1.16.0^35^ with cutoff set to 0.05, and significant DMRs were determined with a Benjamini-Hochberg q-value (q ≤ 0.2). Block DMRs were identified using the dmrseq() function with minInSpan=500, bpSpan=2000, and block=TRUE; blockSize was set to 5000 for identification of blocks as originally defined (5 kb and up), or set to 3500 to include detection of canyon regions (3.5 kb and up). Meta-region plots over DMRs were generated using the Bioconductor package genomation v.1.28.0,^70^ and heatmaps were generated using the ComplexHeatmap package v.2.15.4^71^ with default clustering.

For genomic annotation of DMR regions, promoters were defined as ± 2 kb from transcriptional start sites generated by the Bioconductor package TxDb.Hsapiens.UCSC.hg19.knownGene. DMRs were assigned to genes based on their presence within annotated promoter regions. Annotations for K562 chromatin states from chromHMM^72^ and genic features were generated using annotatr’s built-in hg19_K562- chromatin and hg19_basicgenes. Annotations for hg19 CpG islands were obtained from Wu et al.^73^ CpG island shores were defined as 2 kb regions flanking islands, CpG island shelves were defined as 2 kb regions flanking shores, and open seas were defined as everything else. Enrichment analysis of DMRs overlapping genomic features was performed using the odds ratio statistic and Fisher’s two-sided exact test on a 2 x 2 contingency table containing the number of CpGs within and outside of DMRs and the genomic feature (such as inside DMR and inside CpG island, inside DMR and outside CpG island, and so on). Gene set enrichment analysis (GSEA) of genes associated with DMRs was performed using gene sets from the GSEA database.^74,75^

For analysis of methylation over canyon regions, the UCSC LiftOver tool^76^ was used to identify hg19 genomic regions orthologous to murine canyons identified in DNMT3A- null HSCs obtained from Jeong et al.^41^ WGBS data from a healthy CD34+ HSPC control was included for visualization of a normal comparison. Meta-region plots were generated using genomation, and dmrseq’s plotDMRs was used for visualization of individual canyon regions. Genomic coordinates for blocks identified in EBV-transformed B-cells were obtained from Hansen et al.^44^ For enrichment analysis of canyons overlapping block edges, block edges were defined as 1 kb regions flanking blocks. Enrichment analysis was performed as above, with the 2 x 2 contingency table containing the number of CpGs inside canyons and inside block edges, inside canyons and outside block edges, and so on.

### RNA-Sequencing

Cells were sorted from crushed whole bone marrow for RNA-seq at the indicate timepoints. Total RNA was extracted from >100,000 cells using DNA/RNA microprep plus kit (Zymo) following manufacturer’s protocol. RNA quality control, SMART-Seq v4 library preparation, Illumina Next Generation Sequencing, and data quality control were performed at the MedGenome commercial sequencing facility. Sequencing reads were trimmed and quality control was performed using TrimGalore v.0.6.6 (https://doi.org/10.5281/zenodo.5127899). The cDNA index was built using Kallisto Bustools v.0.40.0^77^ using gene annotations downloaded from Ensembl (release 110) and the GRCh37 reference genome. Pseudo-alignment was performed using Kallisto v0.44.0^78^. Data was imported into R using tximport v.1.30.0^79^ using gene mode. Differential testing was performed using DESeq2 v1.42.1.^80^ Gene set enrichment analysis (GSEA) was performed using gene sets from the GSEA database.^74,75^ Methylation and expression were correlated by associating the methylation difference of putative DMRs in promoter regions to the fold change in expression of the corresponding genes. Pearson’s correlation coefficient was calculated using cor.test in R.

### Single-cell RNA-Sequencing and data processing

200-500 edited residual HSCs were added to 4 mL of MethoCult H4435 (Stemcell Technologies) and differentiated for 14 days. Cells were washed and live cells were sorted by FACS. Samples were then fixed using the Parse Biosciences Cell Fixation kit (Parse Biosciences); barcoding and library prep was then performed using the the Evercode Whole Transcriptome (v2) kit (Parse Biosciences). Resulting libraries were sequenced on a NovaSeq Illumina instrument using 150 cycle kits. Read trimming, quality control, alignment and count matrix generation was performed using the Parse Pipeline v.1.1.2. Data was processed using Seurat v5.0.3,^81^ pre-processing and normalization was performed using the SCTransformv2, putative doublets were removed using DoubletFinder v2.0.4^82^ and samples were integrated using HarmonyIntegration.^83^ Cell types were assigned using Cell Marker Accordion^84^ and GSEA was performed using fgsea.^85^

### Statistical analysis

Statistical analyses were performed in R version 4.3.1 or Prism10 (GraphPad Software). Paired/unpaired t-test was used to define statistical significance (*P < 0.05, **P < 0.01, ***P < 0.001 and ****P < 0.0001). Enrichment analyses of genomic regions were performed as described above.

## Acknowledgments

We would like to thank all patients and their families for providing AML specimens for this study. We thank the Stanford Veterinary Service Center for animal husbandry services. We thank the Flow Cytometry core at the Stanford Institute for Stem Cell Biology and Regenerative Medicine for providing flow cytometry training, equipment, and support. Some of the computing for this project was performed on the Sherlock cluster, and we thank Stanford University and the Stanford Research Computing Center for providing computational resources and support that contributed to these research results. We thank Kasper Hansen for his thoughtful review and suggestions. We also thank all members of the Majeti and Feinberg labs for helpful input, comments, and discussion.

## Author contributions

Conceived and performed experiments and/or analyzed the data: T.K., D.K., E.H., A.C.F., K.K., M.M., C.T.C., F.P.S., A.R., Y.F., Y.N., E.M.M., M.K., J.L.F., K.M.L., A.P.F., and R.M.; contributed reagents, materials, and analysis tools, J.L.F., and K.M.L.; wrote the paper, T.K., E.H., M.K., A.P.F., and R.M.

## Data and materials availability

All unique reagents generated in this study are available from the lead contact without restriction. Sequencing data will be uploaded to GEO. This paper does not report original code. Any additional information required to reanalyze the data reported in this paper is available from the lead contact upon request.

## Supplementary Materials

Figs. S1 to S6 Tables S1 to S14 Data S1 to S7

