## Supplemental Information for "DNMT3A^R882H^ Is Not Required for Disease Maintenance in Primary Human AML, but Is Associated With Increased Leukemia Stem Cell Frequency"

**Fig. S1. Gene editing of primary human patient specimens.**

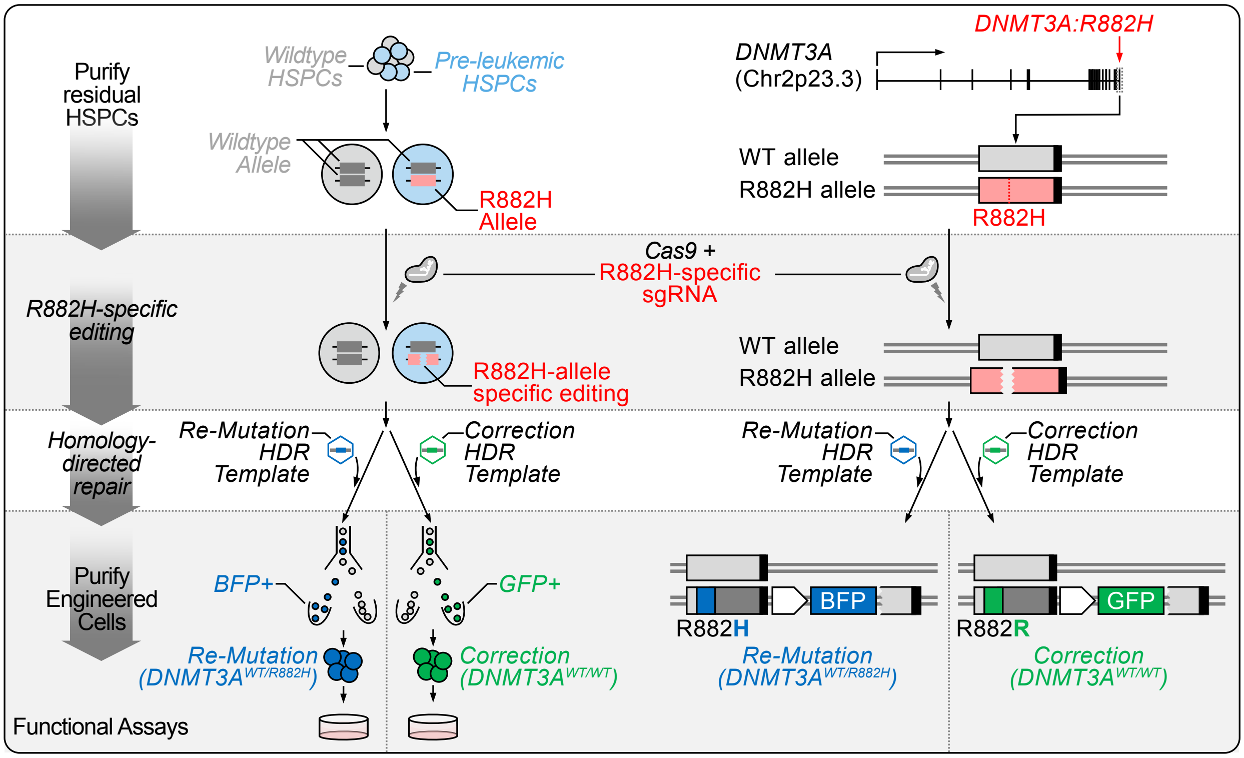


Primary human cells (either residual HSPCs as shown, or leukemic blasts) carrying the *DNMT3A*$^{R882H}$ missense mutation are isolated by flow cytometry from cryopreserved peripheral blood or bone marrow AML specimens. After culturing cells for 48 hours, pre-complexed Cas9 and sgRNA targeting the mutant, R882H allele, are electroporated into the target cells and induce a double-strand break of the mutant allele. Next, cells are transduced with rAAV6 containing either the codon-optimized mutant (R882H) or wildtype (R882R) sequence, a selectable marker (BFP or GFP) driven by a UBC promoter as well as two homology arms (400bp each) flanking the cutsite. 72h after gene editing, cells expressing either BFP (Re-Mutation – *DNMT3A*$^{WT/R882H}$) or GFP (Correction – *DNMT3A*$^{WT/WT}$) are purified by FACS and subjected to functional assays.

**Fig. S2. Genotyping of gene-edited pre-leukemic HSCs.**

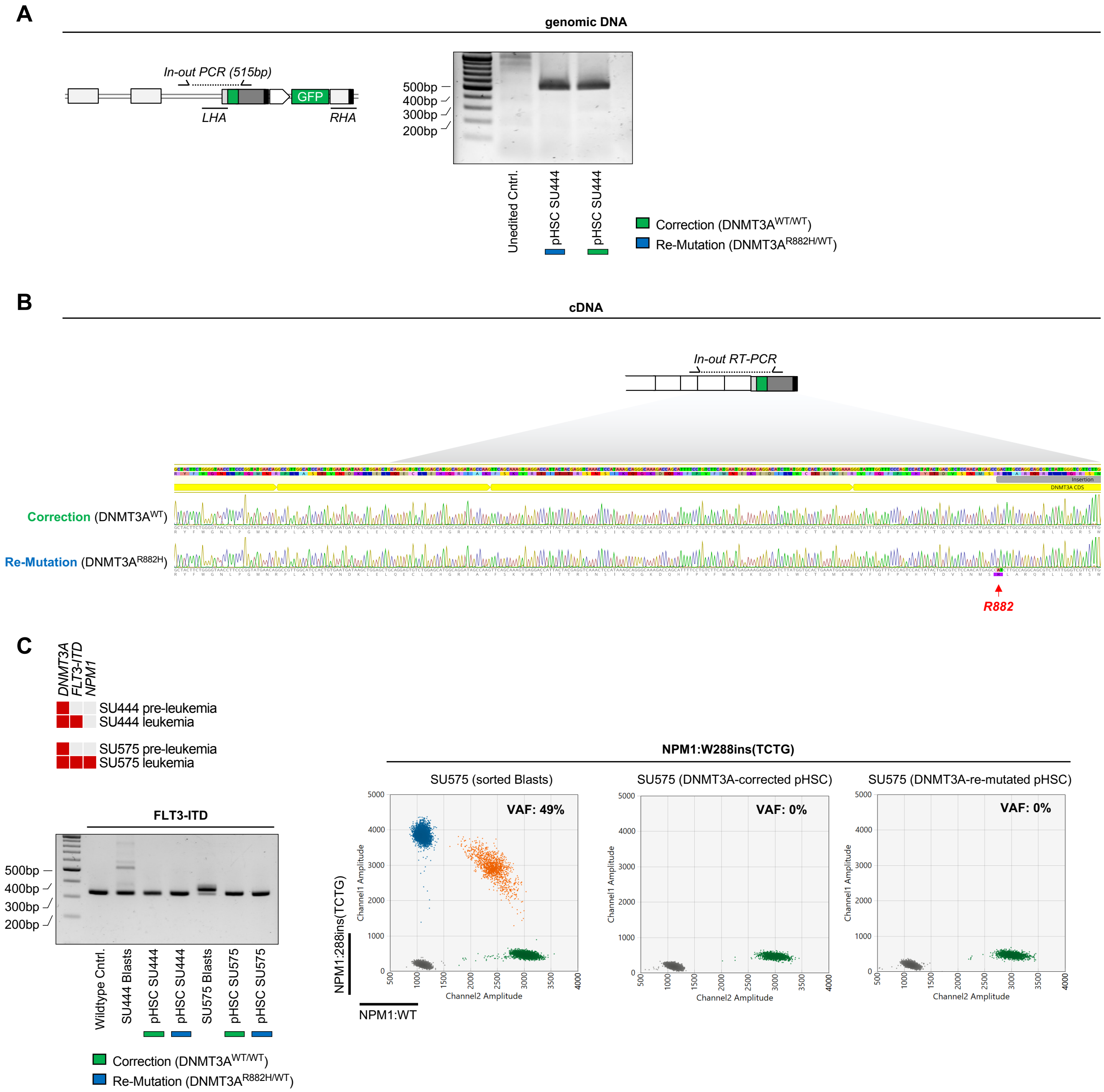


A: In-out PCR with a forward primer placed in the intron preceding exon 23 of *DNMT3A*, upstream of the left arm of homology. The reverse primer is placed in the codon-optimized coding sequence. Presence of a 515bp band indicates on-target integration of the repair template. B: Sanger sequencing of cDNA derived from the edited allele either for cells with corrected (*DNMT3A*$^{WT/WT}$, top) or re-mutated (*DNMT3A*$^{WT/R882H}$,bottom) genotype. A forward primer is placed in the native exon 20 of the *DNMT3A* cDNA and a reverse primer is placed in the inserted, codon-optimized coding sequence of exon 23. C: *FLT3-ITD* (left) and *NPM1* (right) genotyping for gene-edited pre-leukemic HSCs showing no detection of *FLT3-ITD* or *NPM1c* in edited, pre-leukemic cells.

### Fig. S3. Genotyping of gene-edited leukemic blasts.

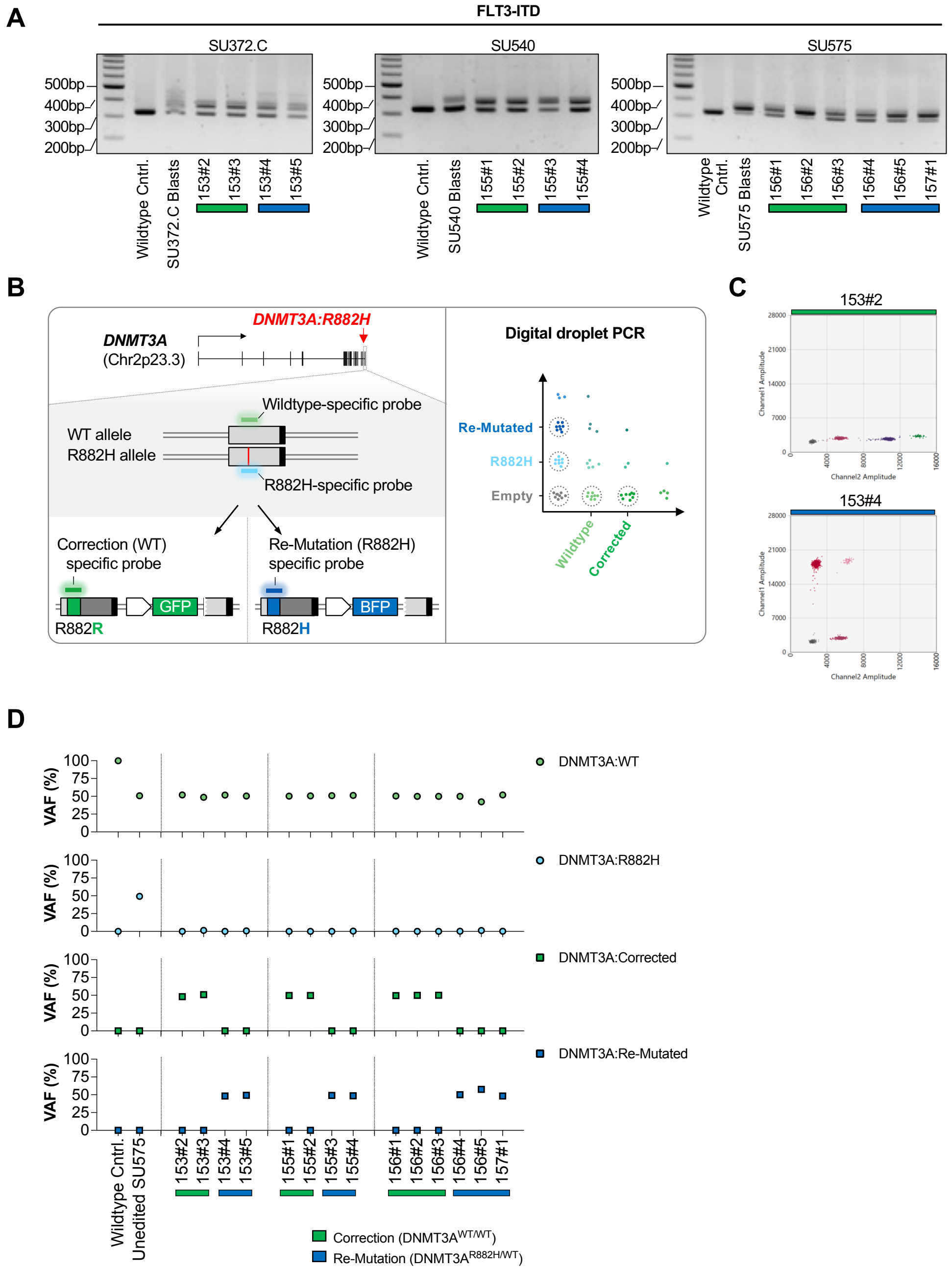


A: *FLT3-ITD* PCR from genomic DNA. Wildtype control DNA (negative control) and DNA from freshly isolated leukemic blasts (positive control) are shown alongside with *DNMT3A* corrected (*DNMT3A*$^{WT/WT}$ – green) or *DNMT3A* re-mutated (*DNMT3A*$^{WT/R882H}$ – blue) human leukemic cells isolated from NSGS mice. B: Multiplexed digital droplet PCR (ddPCR) assays showing probe targets for the quantification of endogenous wildtype, endogenous mutant, corrected, or re-mutated *DNMT3A* alleles (left). Quantification strategy for multiplexed assay (right). C: Representative ddPCR plot showing *DNMT3A* corrected (*DNMT3A*$^{WT/WT}$, top) cells with detection of the endogenous wildtype and corrected alleles, and *DNMT3A* re-mutated (*DNMT3A*$^{WT/R882H}$, bottom) with detection of the endogenous wildtype and re-mutated alleles. D: Quantification of the frequency of endogenous wildtype, endogenous mutant, corrected, and re-mutated alleles in control (wildtype and unedited leukemic cells) as well as in cells isolated from xenografted mice.

**Fig. S4 . Features of AML-derived iPSC line iSU444.**

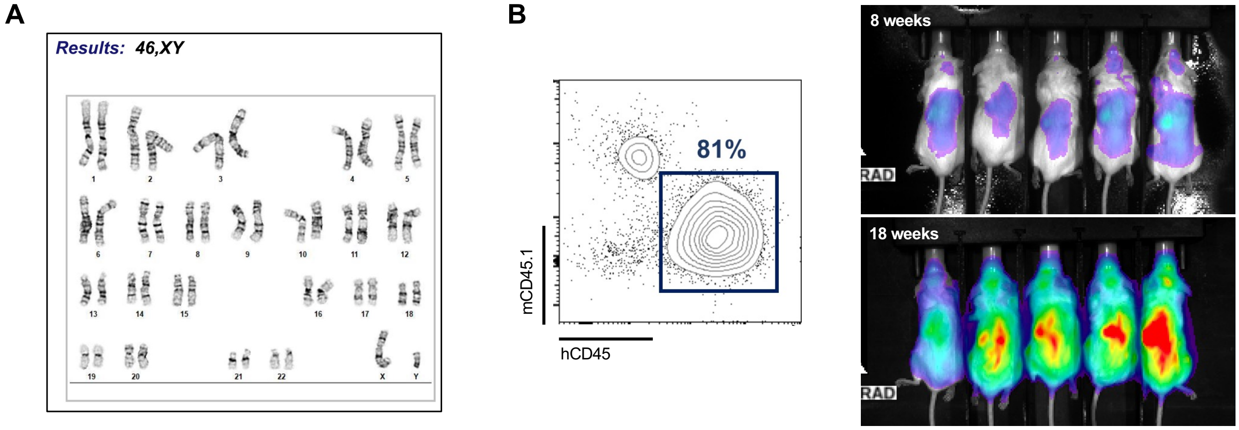


A: Karyotyping of AML-derived iPSC line iSU444 showing a normal (46,XY) karyotype. B: Representative engraftment chimerism (left) and *in vivo* bioluminescence imaging (right) of xenografted leukemic blasts derived from AkaLuc$^{+}$ iPSC line iSU444 5 minutes after injection of 30 µM AkaLumine-HCl at 8 weeks (top right) and 18 weeks (bottom right) after transplantation into NSGS mice.

### Fig. S5. In vivo correction of DNMT3A$^{R882H}$.

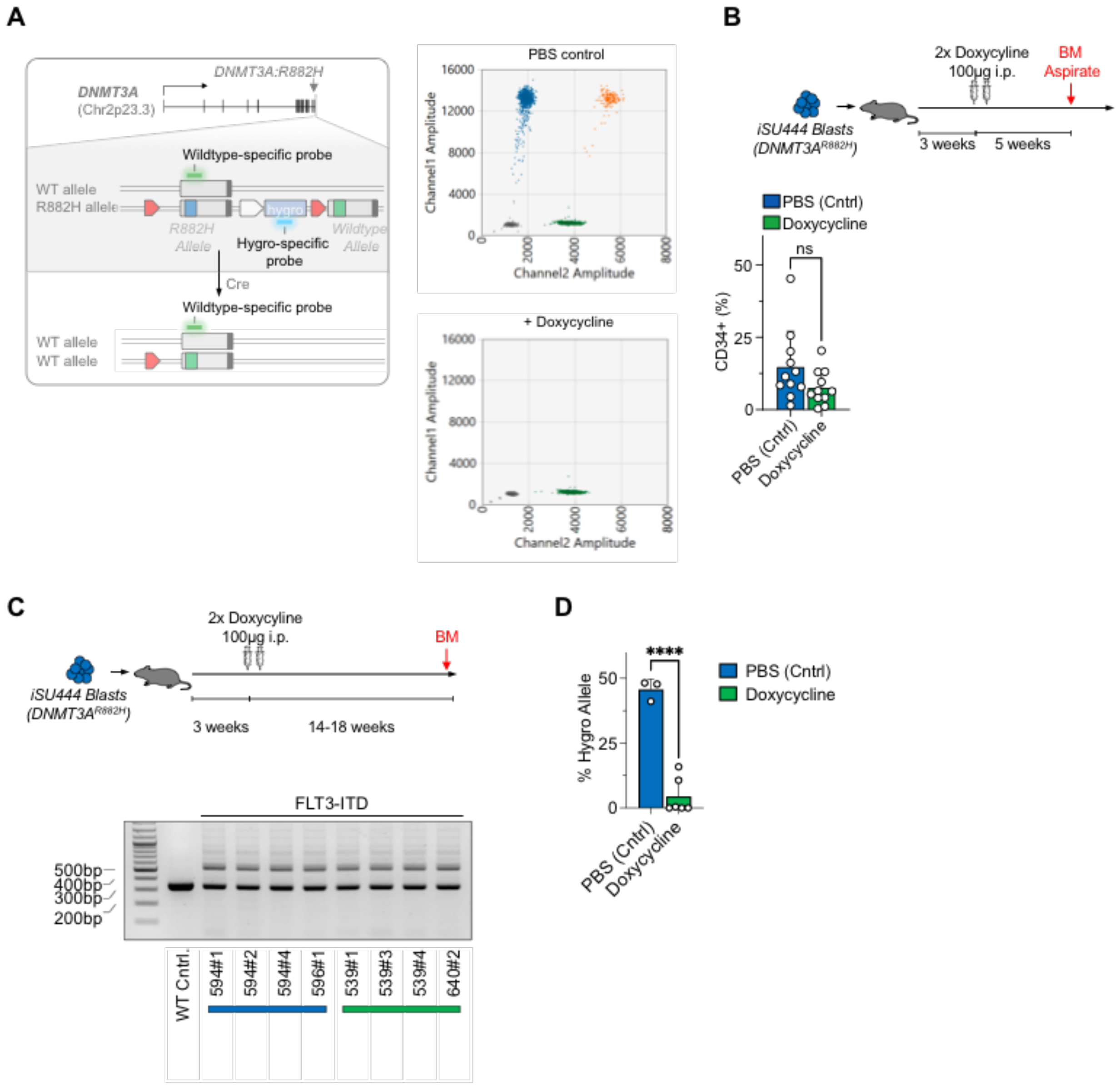


A: ddPCR strategy to quantify the frequency of recombination in iSU444. One probe (FAM) binds to the Hygromycin resistance cassette which is between the LoxP sites, the second probe binds the *DNMT3A* wildtype allele (VIC) and serves as a reference. Upon successful recombination, the Hygromycin binding probe is lost. Representative ddPCR plots for control (PBS) mouse (top right) and mouse receiving Doxycycline (bottom right). B: Bone marrow aspirates were obtained 5 weeks after injection of two doses of Doxycycline (top) and percentage of CD34+ cells was determined (bottom). C: Moribund mice were sacrificed between 14-18 weeks after injection of Doxycycline (top) and *FLT3-ITD* status was determined (bottom). D: Recombination rates of cells recovered at sacrifice as determined by the ddPCR assay, ****, $P < 0.0001$.

**Fig. S6 . Whole genome bisulfite sequencing of *DNMT3A* corrected leukemic cells**

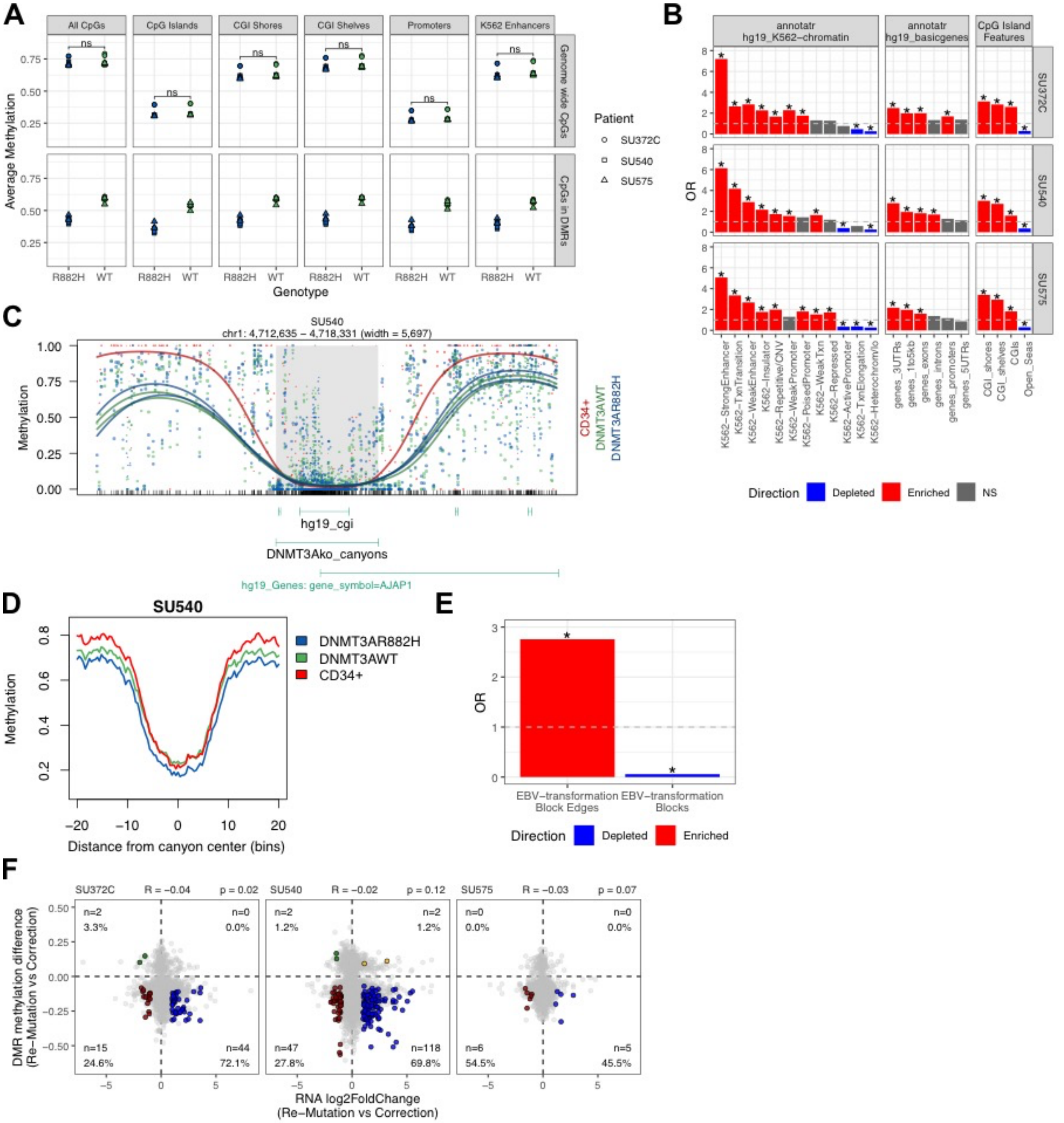


A: Average methylation for each patient-derived specimen grouped by genomic feature (columns), location of CpGs either genome-wide or within DMR (rows). Each block is further subdivided by genotype (*DNMT3A* re-mutation: “R882H” versus *DNMT3A* correction: “WT”). Patient ID is indicated by the symbol shape. B: Enrichment of DMRs over annotated genomic features, grouped by patient (rows) and annotation type (columns). C: Representative locus at methylation canyons as defined by Jeong et al., Nature Genetics 2014. D: Meta-region plot of methylation over canyon regions containing

DMRs in *DNMT3A* re-mutation versus *DNMT3A* correction for patient SU540. E: Enrichment of methylation canyons in EBV-transformation blocks and block edges. Block edges are defined as 1 kb regions flanking blocks. F: Scatterplot of fold change in gene expression (x-axis) versus methylation change over putative promoter DMRs (y-axis) for each of the three patient specimens. Genes containing significant DMRs (q £ 0.2) and absolute RNA log2 fold change ³ 1 are highlighted. *: Odds ratio ≥ 1.5 (enrichment) or ≤ 0.5 (depletion) and Fisher’s p-value < 2e-16.

**Table S1. FACS staining panels used in this study**

**Table S2. sgRNA sequences and PCR Primers used in this study**

**Table S3. Summary of DMRs between DNMT3A$^{R882H}$ re-mutated and DNMT3A$^{WT}$ corrected blasts for patients SU372.C, SU540, and SU575**

DMRs were identified between DNMT3AR882H re-mutated and DNMT3AWT corrected samples within each patient using dmrseq (Methods). Summary statistics for DMRs within each patient are reported here. The contents of the tables are as follows (column name = description):

- Patient comparison = Patient identifier for DMR comparison (SU372.C, SU540, or SU575)
- Total # of DMRs = Number of significant DMRs identified
- Average # CpGs in DMR = Average number of CpGs in a DMR
- Average width of DMR (bp) = Average number of base pairs in a DMR
- # hypomethylated (in DNMT3A$^{R882H/WT}$) = Number of hypomethylated DMRs (raw difference < 0) in DNMT3A$^{R882H}$ versus DNMT3A$^{WT}$
- Proportion hypomethylated (in DNMT3A$^{R882H/WT}$) = Proportion of total DMRs that are hypomethylated in DNMT3A$^{R882H}$ versus DNMT3A$^{WT}$ (raw difference < 0), calculated as number hypomethylated DMRs divided by total number of DMRs

**Table S4-6: DMRs between DNMT3A$^{R882H}$ re-mutated and DNMT3A$^{WT}$ corrected blasts for patients SU372.C, SU540, and SU575**

DMRs were identified between DNMT3AR882H re-mutated and DNMT3AWT corrected samples within each patient using dmrseq (Methods). DMRs are reported for each patient comparison (SU372.C, **Table S4**; SU540, **Table S5**; SU575, **Table S6**). The contents of the tables are as follows (column name = description):

- DMR_coordinates (chr:start-end) = Genomic location of DMR, chr:start-end (hg19)
- DMR_width (bp) = Width of DMR (bp)
- num_CpGs = Number of CpGs within DMR
- areaStat = dmrseq area statistic - the sum of the smoothed beta values for the DMR
- rawMethDiff = Raw methylation difference (DNMT3AR882H - DNMT3AWT)
- testStat = dmrseq test statistic
- pval = dmrseq permutation p-value for the significance of the test statistic
- qval = dmrseq false discovery rate-adjusted q-value
- DMRdirection (relative to DNMT3AWT) = Direction of DMR relative to DNMT3AWT. "Hypo" = hypomethylated (in DNMT3AR882H), "Hyper" = hypermethylated (in DNMT3AR882H)

**Table S7: Genes containing promoter DMRs between DNMT3A$^{R882H}$ re-mutated and DNMT3A$^{WT}$ corrected blasts in all patients (SU372.C, SU540, and SU575)**

DMRs were identified between DNMT3AR882H re-mutated and DNMT3AWT corrected samples within each patient using dmrseq, and mapped to genes by presence within annotated promoter regions (Methods). The intersection of genes with DMRs in all patients is reported.

**Table S8. Overlap between DMRs (DNMT3A$^{R882H}$ versus DNMT3A$^{WT}$) and DNA methylation canyons, stratified by patient and canyon type.**
The overlaps between DMRs in each patient and DNA methylation canyon regions (as defined by Jeong et al., Nature Genetics 2014) are reported here. The contents of the tables are as follows (column name = description):

- Patient (for DMRs) = Patient identifier for DMR comparison (SU372.C, SU540, or SU575)
- # of DMRs = Number of significant DMRs identified (DNMT3A$^{R882H}$ versus DNMT3A$^{WT}$) for given patient
- Canyon Type (Jeong et al., 2014) = Genotype of murine HSCs in which DNA methylation canyons were defined by Jeong et al., Nature Genetics 2014.
- # of canyons (hg19 liftover) = Number of analogous hg19 DNA methylation canyons, identified by applying UCSC LiftOver tool to murine canyons and filtering by width $\geq$ 3500 base pairs
- # DMRs overlapping canyons = Number of DMRs overlapping the given canyon type, calculated by counting overlaps between DMRs and canyon regions
- Proportion DMRs overlapping canyons (out of total # of DMRs) = Proportion of DMRs overlapping the given canyon type, calculated as number of DMRs overlapping canyons divided by total number of DMRs
- # canyons overlapping DMRs = Number of canyons overlapping DMRs for the given patient, calculated by counting overlaps between canyon regions and DMRs
- Proportion canyons overlapping DMRs (out of total # of canyons) = Proportion of canyons overlapping DMRs, calculated as number of canyons overlapping DMRs divided by total number of canyons

**Table S9-14: Putative blocks between DNMT3A$^{R882H}$ re-mutated and DNMT3A$^{WT}$ corrected blasts for patients SU372.C, SU540, and SU575**
The R package dmrseq was used to search for blocks between DNMT3AR882H re-mutated and DNMT3AWT corrected samples within each patient (Methods). Putative blocks are reported for each patient comparison with dmrseq blockSize=3500 or 5000 (SU372.C, blockSize=3500, **Table S9**; SU372.C, blockSize=5000, **Table S10**; SU540, blockSize=3500, **Table S11**; SU540, blockSize=5000, **Table S12;** SU575, blockSize=3500, **Table S13**; SU540, blockSize=5000, **Table S14**). The contents of the tables are as follows (column name = description):

- Block_coordinates (chr:start-end) = Genomic location of block, chr:start-end (hg19)
- Block_width (bp) = Width of block (bp)
- num_CpGs = Number of CpGs within block
- areaStat = dmrseq area statistic - the sum of the smoothed beta values for the block
- rawMethDiff = Raw methylation difference (DNMT3AR882H - DNMT3AWT)
- testStat = dmrseq test statistic

- pval = dmrseq permutation p-value for the significance of the test statistic
- qval = dmrseq false discovery rate-adjusted q-value
- Direction_relativeToD3AWT = Direction of block relative to DNMT3AWT. "Hypo_NS" = hypomethylated (in DNMT3AR882H), qval > 0.2; "Hyper_NS" = hypermethylated (in DNMT3AR882H), qval > 0.2

**Data S1-S3. (separate file)**
Plots of top 100 DMRs by p-value between *DNMT3A*$^{R882H}$ re-mutated and DNMT3A$^{WT}$ samples for patient SU372.C (Data S1), SU540 (Data S2), and SU575 (Data S3). Annotations for gene and CpG island locations are included using dmrseq's getAnnot(). Color represents genotype (green, *DNMT3A*$^{WT}$ corrected; blue, *DNMT3A*$^{R882H}$ re-mutated). Stat: dmrseq test statistic. FDR: false discovery rate-adjusted p-value.

**Data S4-S6. (separate file)**
Plots of top 50 DMRs between *DNMT3A*$^{R882H}$ re-mutated and *DNMT3A*$^{WT}$ samples overlapping DNA methylation canyons as defined by Jeong et al., Nature Genetics 2014. Individual files contain plots for each patient: patient SU372.C (Data S4), SU540 (Data S5), and SU575 (Data S6). Color represents genotype (green, *DNMT3A*$^{WT}$ corrected; blue, *DNMT3A*$^{R882H}$ re-mutated). Stat: dmrseq test statistic. FDR: false discovery rate-adjusted p-value.

**Data S7. (separate file)**
Plots of methylation profiles over DNA methylation canyons as defined by Jeong et al., Nature Genetics 2014. Color represents genotype (red, healthy donor CD34+ reference; green, *DNMT3A* corrected; blue, *DNMT3A* re-mutated), and line type represents patient (solid, SU372.C; dashed, SU540; dotted, SU575). Genomic annotation of EBV-transformation blocks as defined by Hansen et al., Genome Res. 2014, are included for visualization of the correspondence between block edges and canyons.
