## Supplemental Data S1-S7 for "DNMT3A^R882H^ Is Not Required for Disease Maintenance in Primary Human AML, but Is Associated With Increased Leukemia Stem Cell Frequency": DataS1_dmrSeq_SU372_top100DMRs.pdf

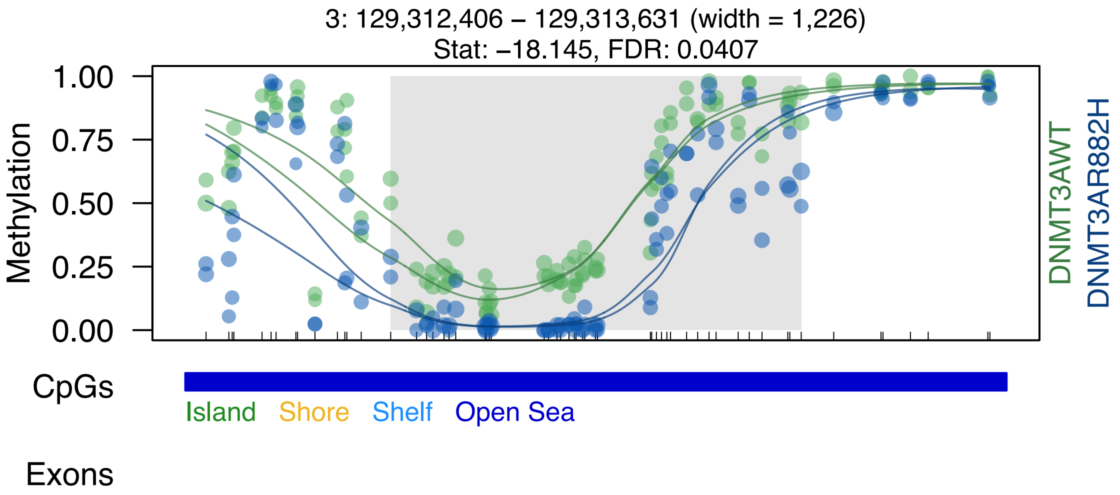
3: 129,312,406 – 129,313,631 (width = 1,226)
Stat: −18.145, FDR: 0.0407
1.00
0.75
0.50
0.25
0.00
Methylation
DNMT3AWT
DNMT3AR882H
CpGs
Island
Shore
Shelf
Open Sea
Exons

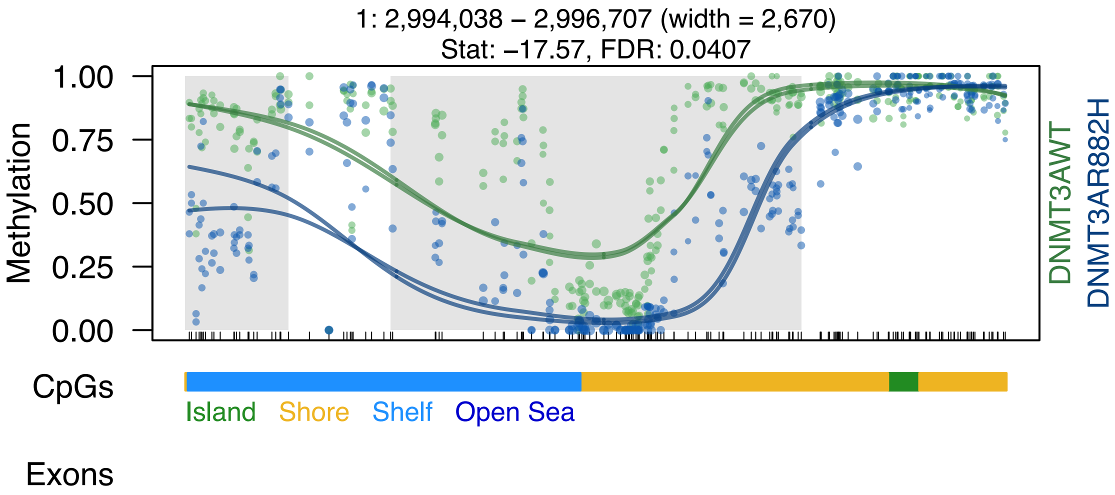
1: 2,994,038 – 2,996,707 (width = 2,670)
Stat: –17.57, FDR: 0.0407
Methylation
1.00
0.75
0.50
0.25
0.00
DNMT3AWT
DNMT3AR882H
CpGs
Island
Shore
Shelf
Open Sea
Exons

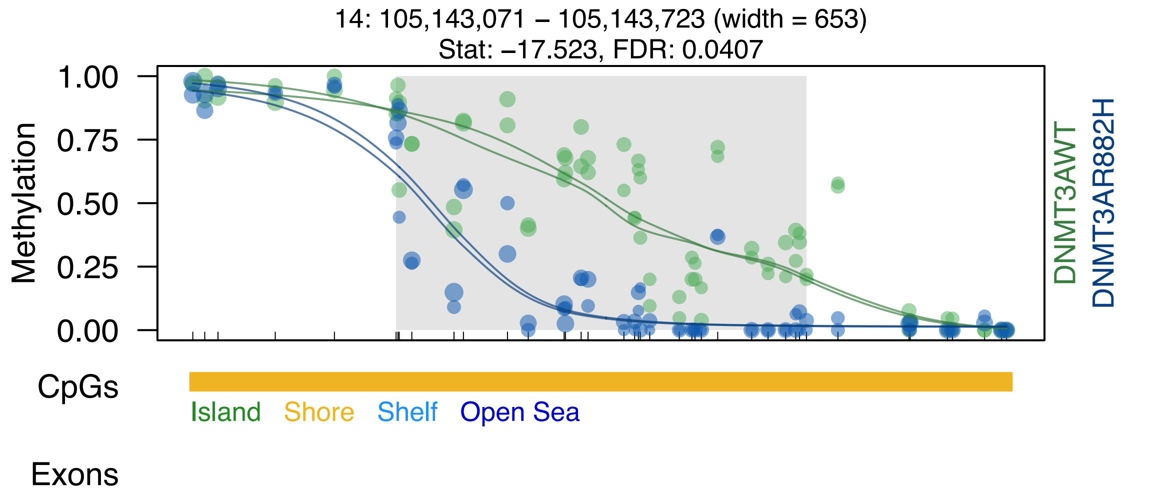
14: 105,143,071 – 105,143,723 (width = 653)
Stat: −17.523, FDR: 0.0407
Methylation
1.00
0.75
0.50
0.25
0.00
DNMT3AWT
DNMT3AR882H
CpGs
Island
Shore
Shelf
Open Sea
Exons

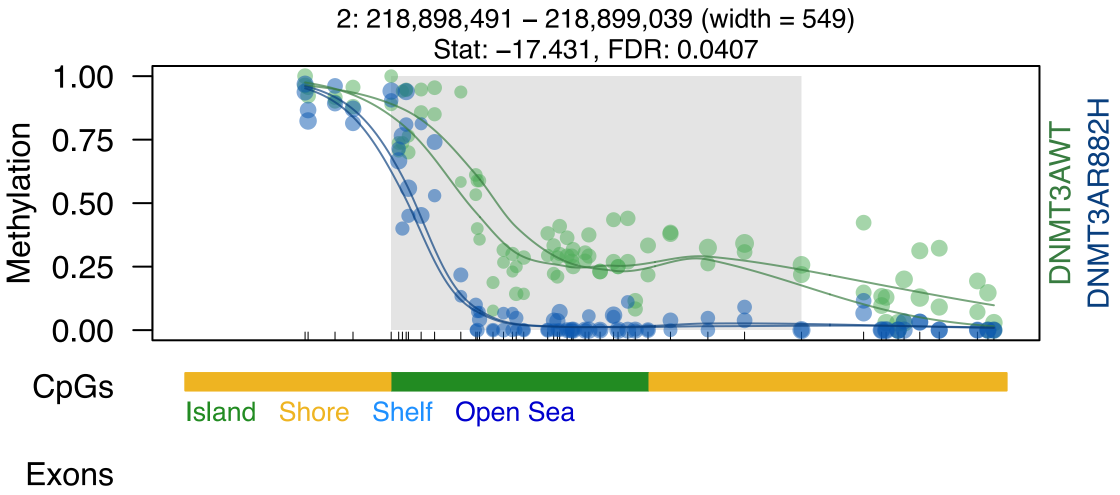

2: 218,898,491 – 218,899,039 (width = 549)
Stat: –17.431, FDR: 0.0407
Methylation
1.00
0.75
0.50
0.25
0.00
DNMT3AWT
DNMT3AR882H
CpGs
Island
Shore
Shelf
Open Sea
Exons

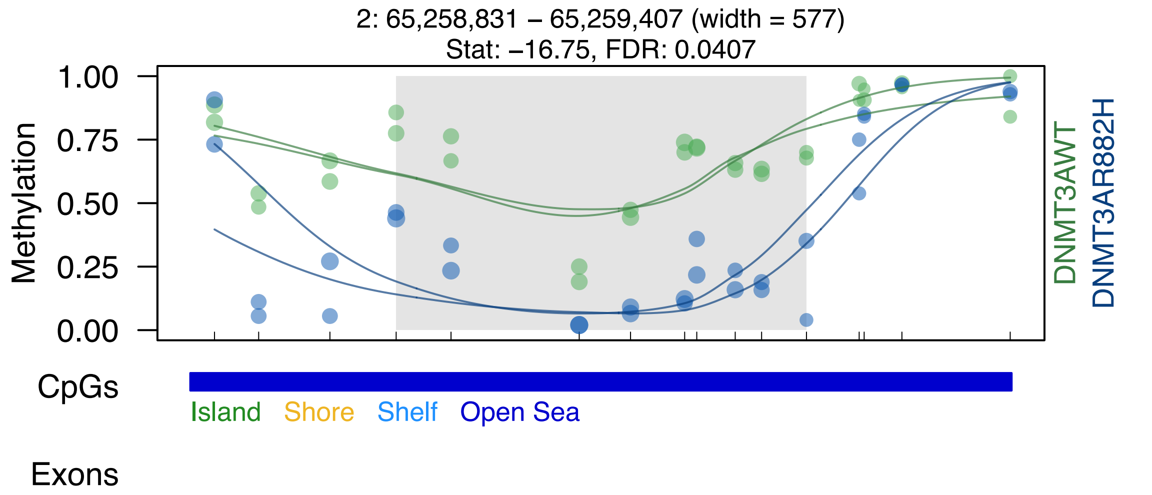

2: 65,258,831 – 65,259,407 (width = 577)
Stat: –16.75, FDR: 0.0407
Methylation
1.00
0.75
0.50
0.25
0.00
DNMT3AWT
DNMT3AR882H
CpGs
Island
Shore
Shelf
Open Sea
Exons

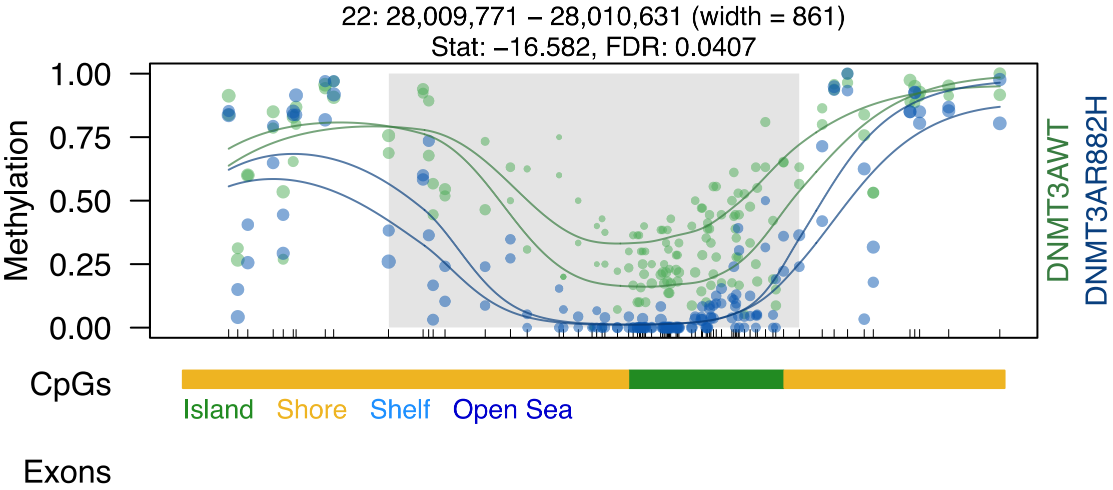

22: 28,009,771 – 28,010,631 (width = 861)
Stat: –16.582, FDR: 0.0407
Methylation
1.00
0.75
0.50
0.25
0.00
DNMT3AWT
DNMT3AR882H
CpGs
Island
Shore
Shelf
Open Sea
Exons

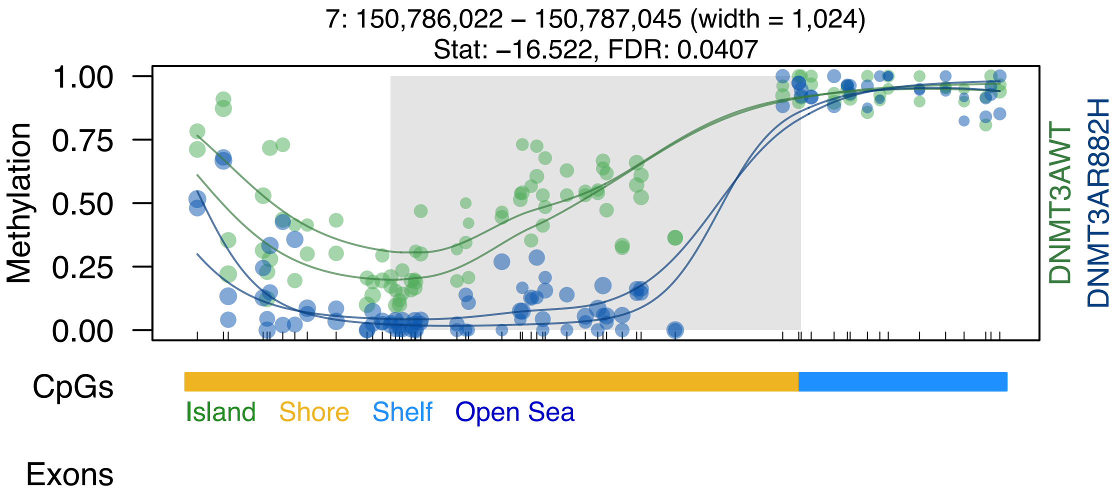
7: 150,786,022 – 150,787,045 (width = 1,024)
Stat: −16.522, FDR: 0.0407
Methylation
1.00
0.75
0.50
0.25
0.00
DNMT3AWT
DNMT3AR882H
CpGs
Island
Shore
Shelf
Open Sea
Exons

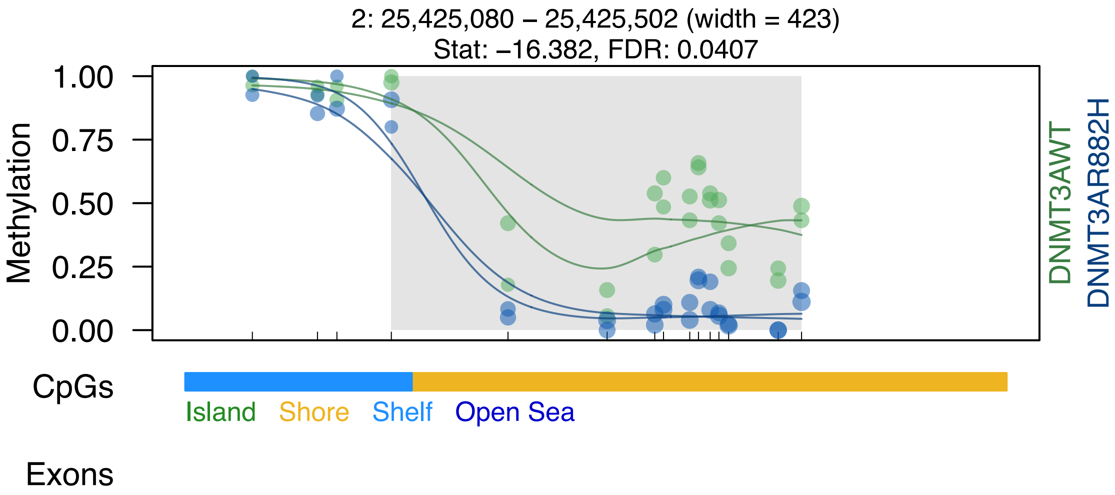

2: 25,425,080 – 25,425,502 (width = 423)
Stat: –16.382, FDR: 0.0407
Methylation
1.00
0.75
0.50
0.25
0.00
DNMT3AWT
DNMT3AR882H
CpGs
Island
Shore
Shelf
Open Sea
Exons

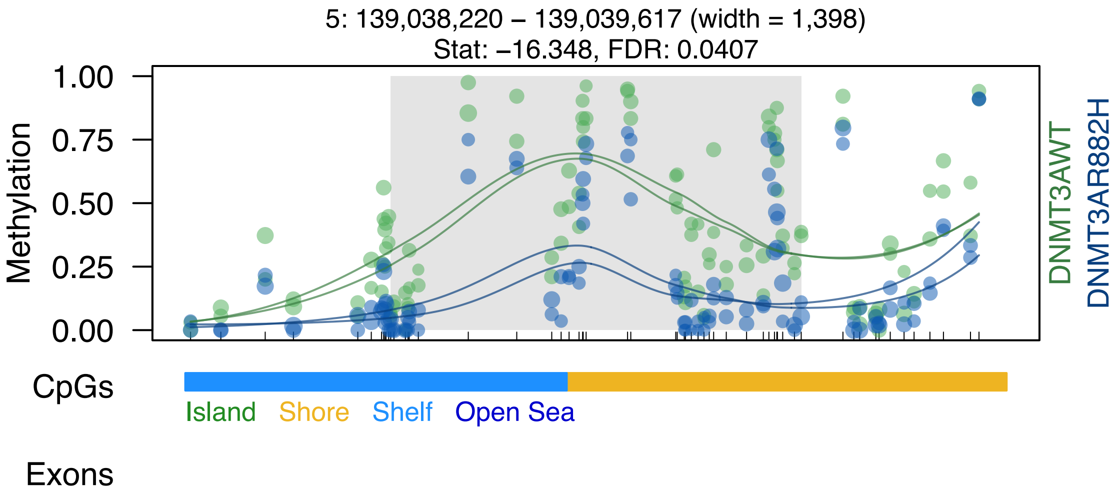

5: 139,038,220 – 139,039,617 (width = 1,398)
Stat: −16.348, FDR: 0.0407
Methylation
1.00
0.75
0.50
0.25
0.00
DNMT3AWT
DNMT3AR882H
CpGs
Island
Shore
Shelf
Open Sea
Exons

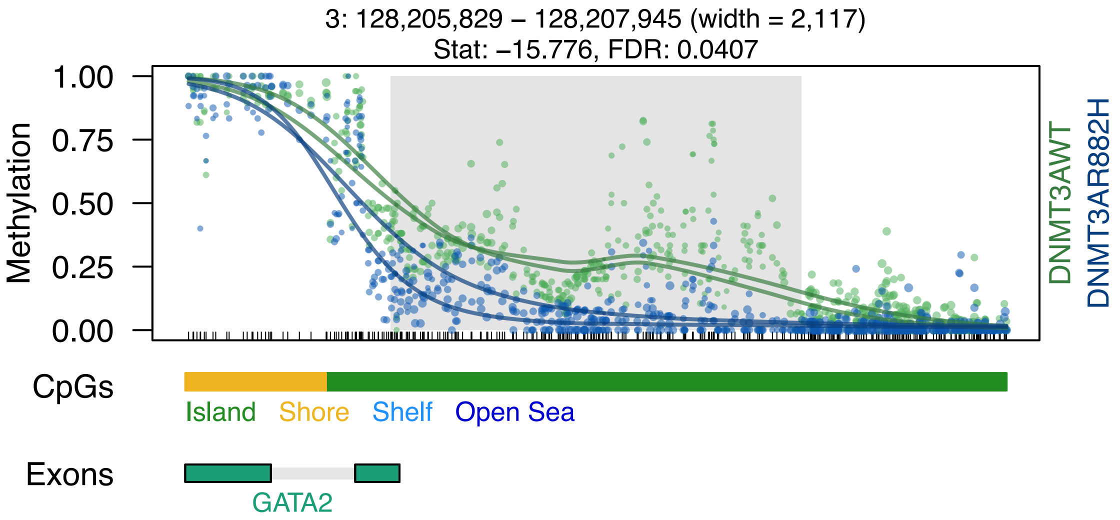

3: 128,205,829 – 128,207,945 (width = 2,117)
Stat: –15.776, FDR: 0.0407
Methylation
1.00
0.75
0.50
0.25
0.00
DNMT3AWT
DNMT3AR882H
CpGs
Island
Shore
Shelf
Open Sea
Exons
GATA2

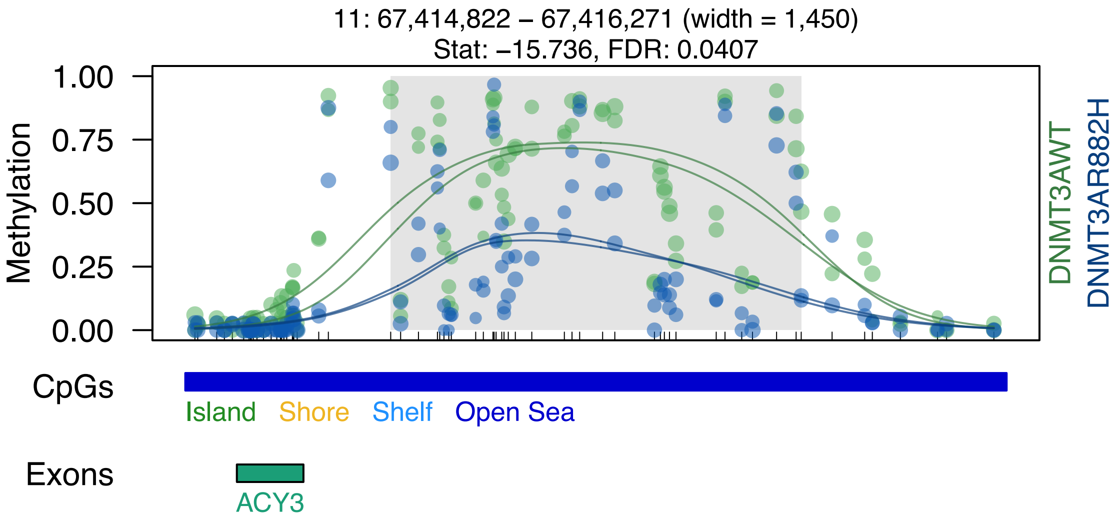

11: 67,414,822 – 67,416,271 (width = 1,450)
Stat: –15.736, FDR: 0.0407
Methylation
1.00
0.75
0.50
0.25
0.00
DNMT3AWT
DNMT3AR882H
CpGs
Island
Shore
Shelf
Open Sea
Exons
ACY3

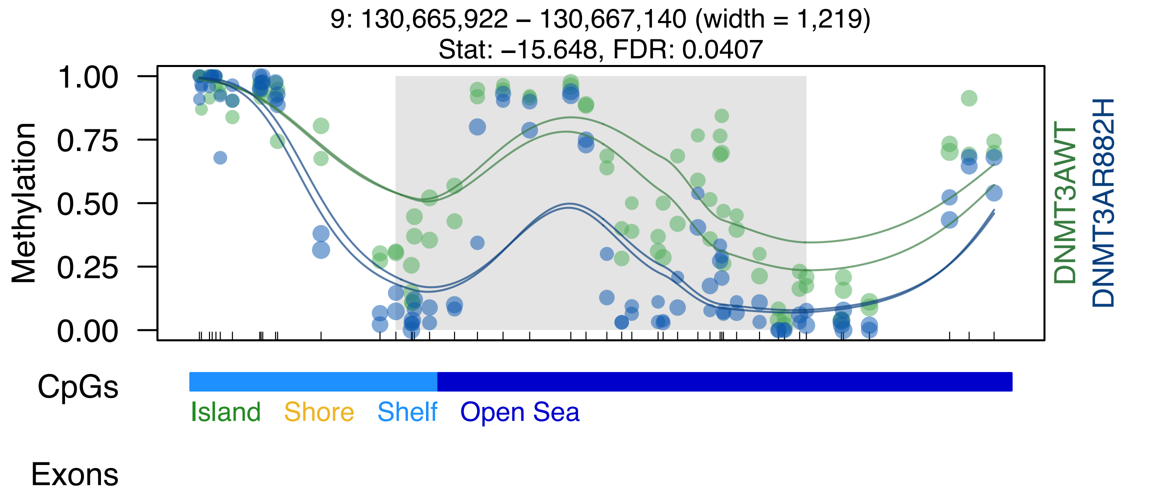

9: 130,665,922 – 130,667,140 (width = 1,219)
Stat: –15.648, FDR: 0.0407
1.00
0.75
0.50
0.25
0.00
Methylation
DNMT3AWT
DNMT3AR882H
CpGs
Island
Shore
Shelf
Open Sea
Exons

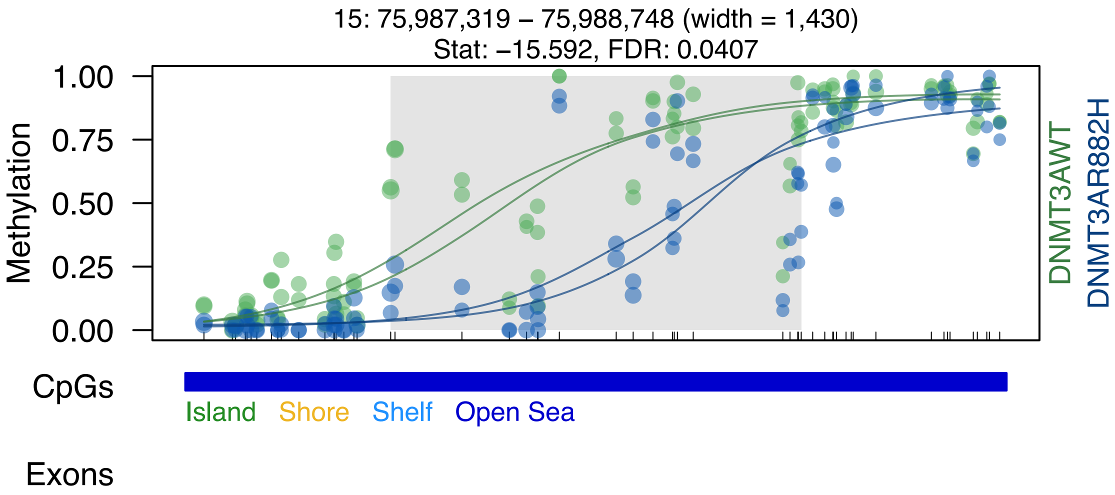
15: 75,987,319 – 75,988,748 (width = 1,430)
Stat: –15.592, FDR: 0.0407
Methylation
1.00
0.75
0.50
0.25
0.00
DNMT3AWT
DNMT3AR882H
CpGs
Island
Shore
Shelf
Open Sea
Exons

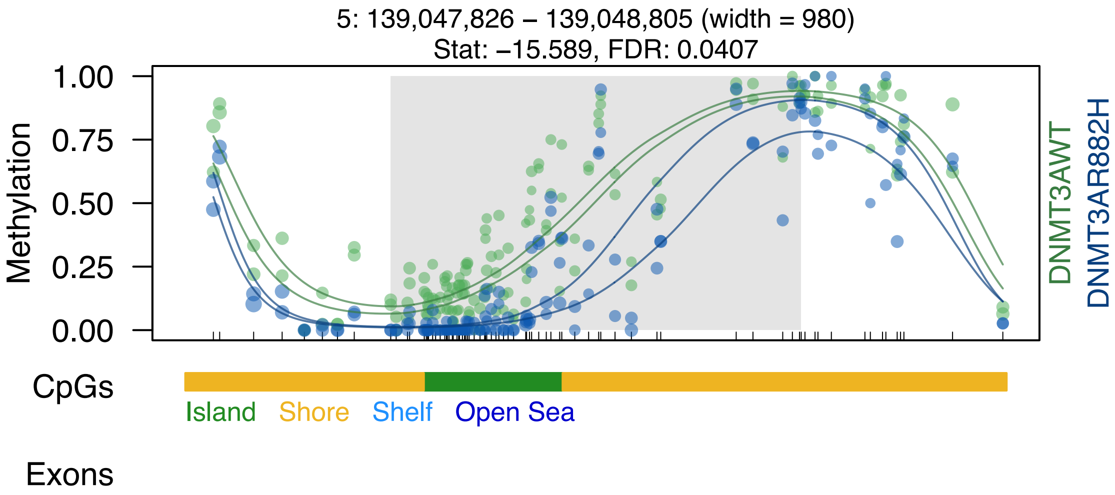

5: 139,047,826 – 139,048,805 (width = 980)
Stat: –15.589, FDR: 0.0407
Methylation
1.00
0.75
0.50
0.25
0.00
DNMT3AWT
DNMT3AR882H
CpGs
Island
Shore
Shelf
Open Sea
Exons

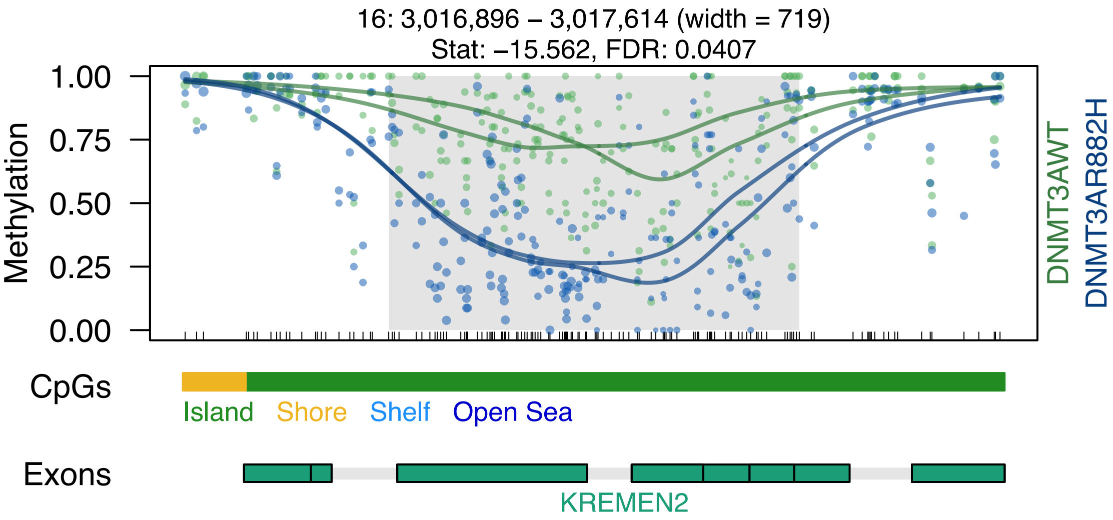

16: 3,016,896 – 3,017,614 (width = 719)
Stat: –15.562, FDR: 0.0407
Methylation
1.00
0.75
0.50
0.25
0.00
DNMT3AWT
DNMT3AR882H
CpGs
Island
Shore
Shelf
Open Sea
Exons
KREMEN2

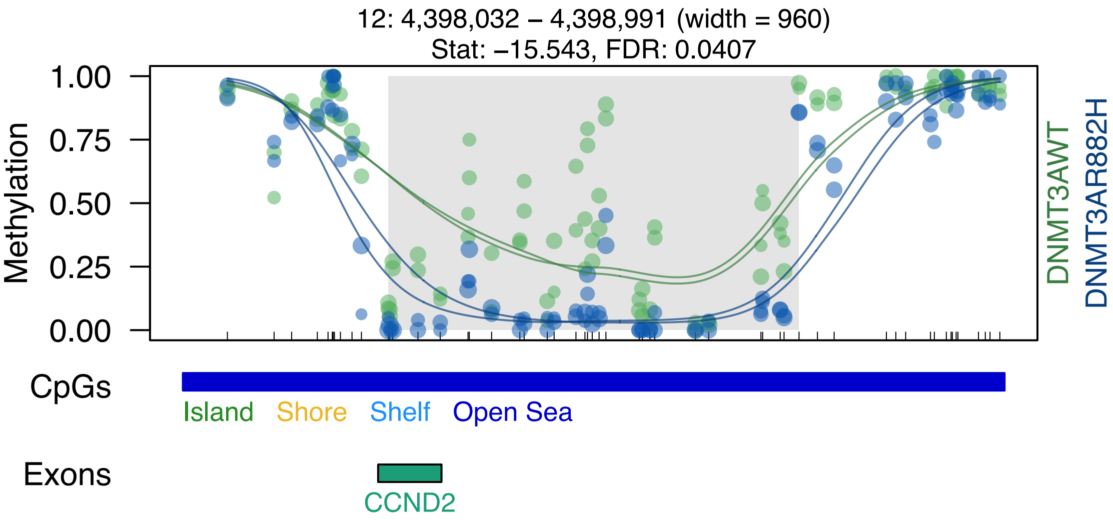

12: 4,398,032 – 4,398,991 (width = 960)
Stat: –15.543, FDR: 0.0407
Methylation
1.00
0.75
0.50
0.25
0.00
DNMT3AWT
DNMT3AR882H
CpGs
Island
Shore
Shelf
Open Sea
Exons
CCND2

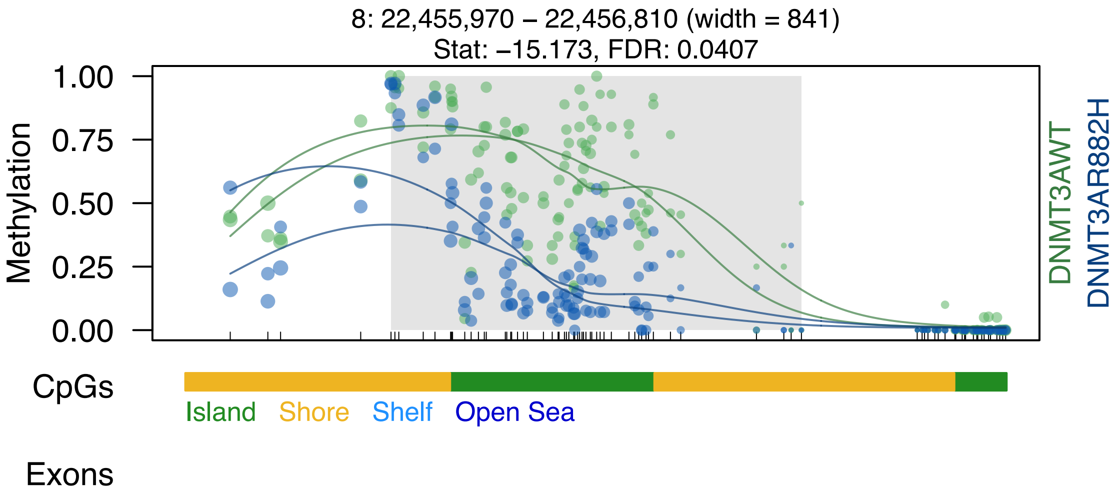

8: 22,455,970 – 22,456,810 (width = 841)
Stat: –15.173, FDR: 0.0407
Methylation
1.00
0.75
0.50
0.25
0.00
DNMT3AWT
DNMT3AR882H
CpGs
Island
Shore
Shelf
Open Sea
Exons

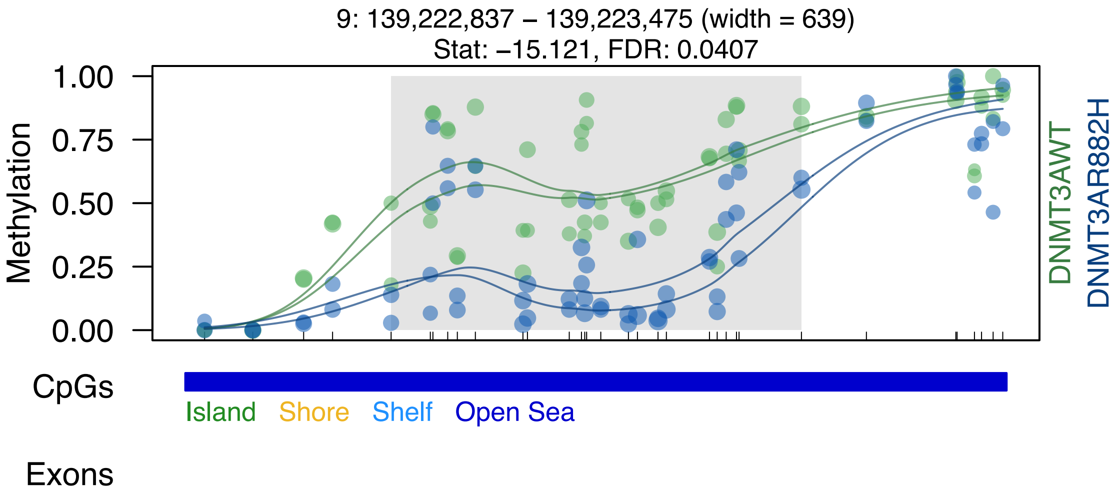

9: 139,222,837 – 139,223,475 (width = 639)
Stat: –15.121, FDR: 0.0407
Methylation
1.00
0.75
0.50
0.25
0.00
DNMT3AWT
DNMT3AR882H
CpGs
Island
Shore
Shelf
Open Sea
Exons

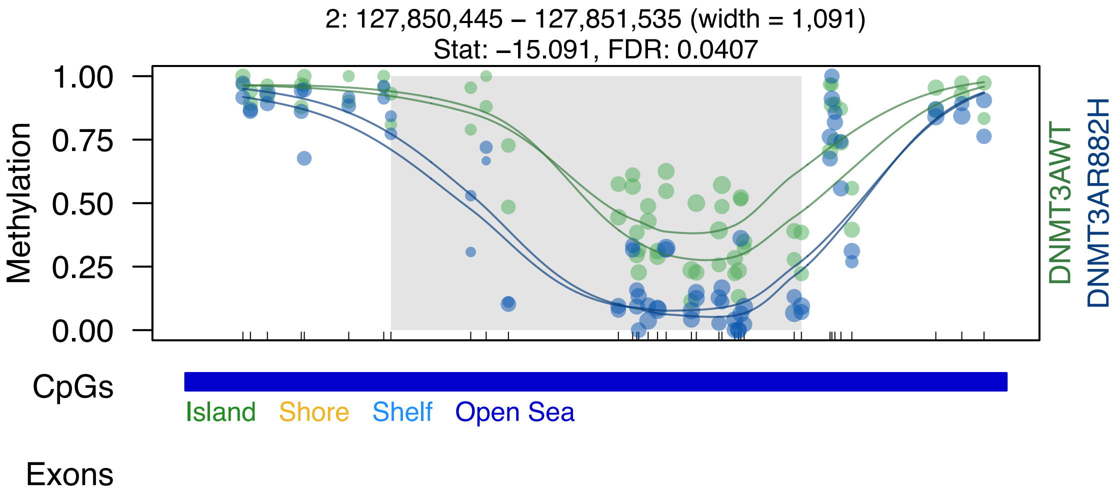

2: 127,850,445 – 127,851,535 (width = 1,091)
Stat: −15.091, FDR: 0.0407
Methylation
1.00
0.75
0.50
0.25
0.00
DNMT3AWT
DNMT3AR882H
CpGs
Island
Shore
Shelf
Open Sea
Exons

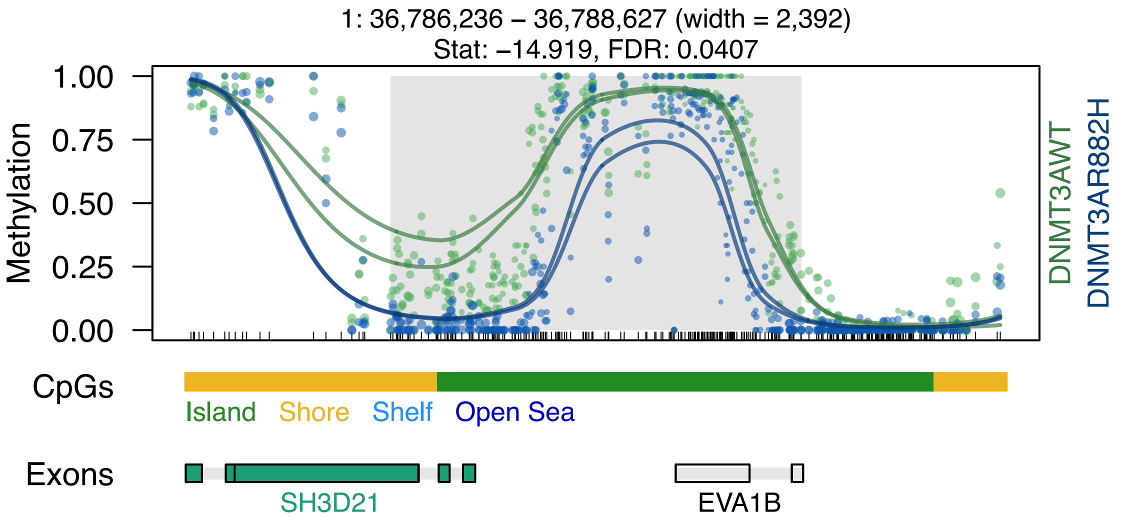
1: 36,786,236 – 36,788,627 (width = 2,392)
Stat: –14.919, FDR: 0.0407
1.00
0.75
0.50
0.25
0.00
Methylation
DNMT3AWT
DNMT3AR882H
CpGs
Island
Shore
Shelf
Open Sea
Exons
SH3D21
EVA1B

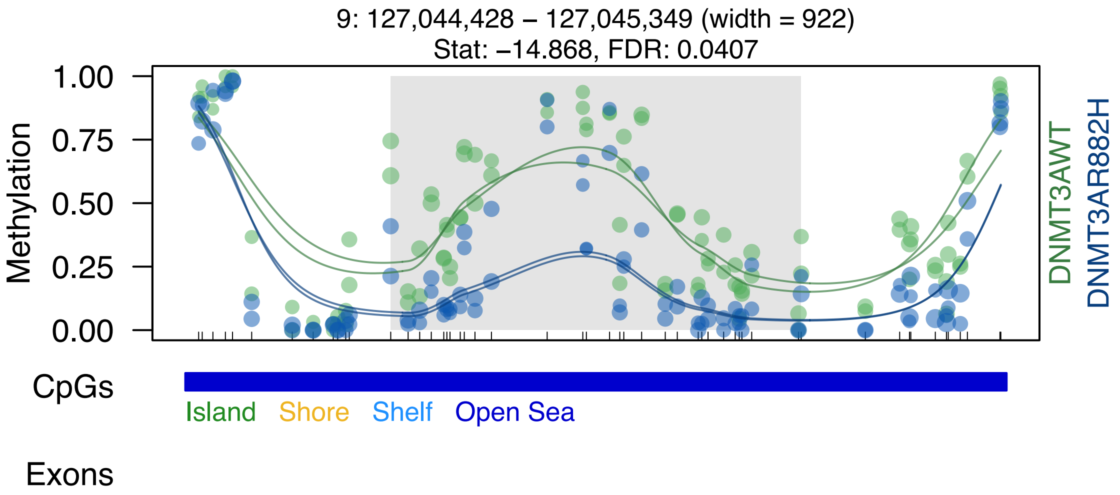

9: 127,044,428 – 127,045,349 (width = 922)
Stat: –14.868, FDR: 0.0407
1.00
0.75
0.50
0.25
0.00
Methylation
DNMT3AWT
DNMT3AR882H
CpGs
Island
Shore
Shelf
Open Sea
Exons

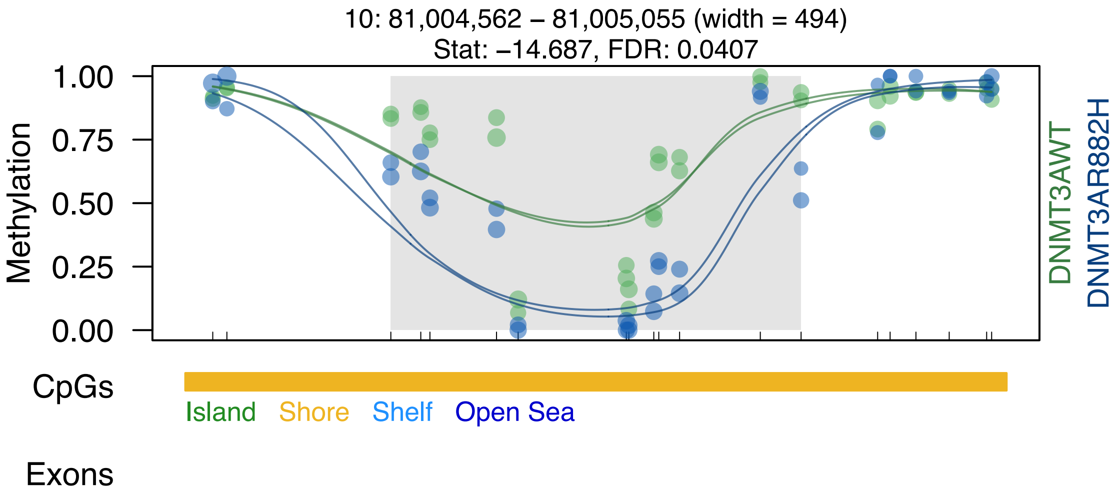

10: 81,004,562 – 81,005,055 (width = 494)
Stat: –14.687, FDR: 0.0407
Methylation
1.00
0.75
0.50
0.25
0.00
DNMT3AWT
DNMT3AR882H
CpGs
Island
Shore
Shelf
Open Sea
Exons

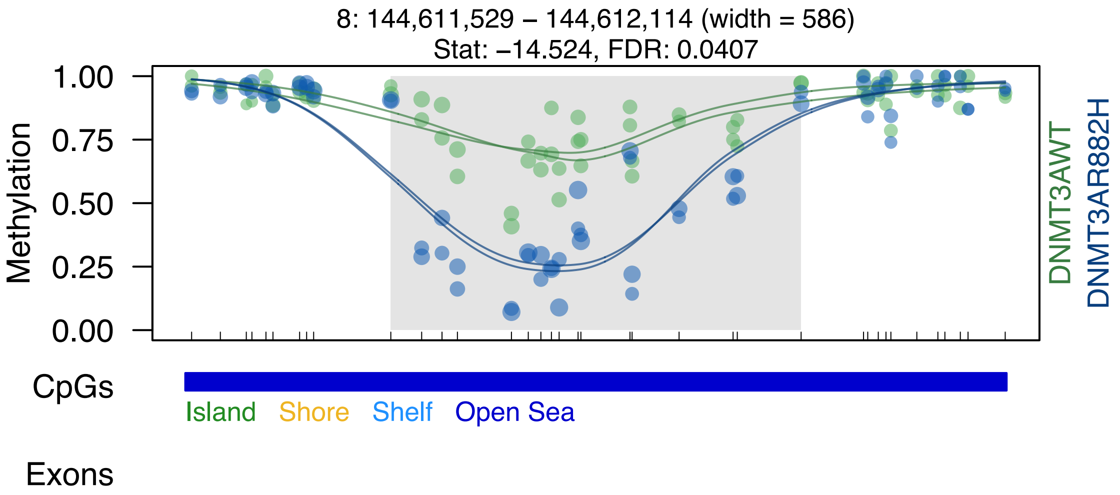

8: 144,611,529 – 144,612,114 (width = 586)
Stat: –14.524, FDR: 0.0407
Methylation
1.00
0.75
0.50
0.25
0.00
DNMT3AWT
DNMT3AR882H
CpGs
Island
Shore
Shelf
Open Sea
Exons

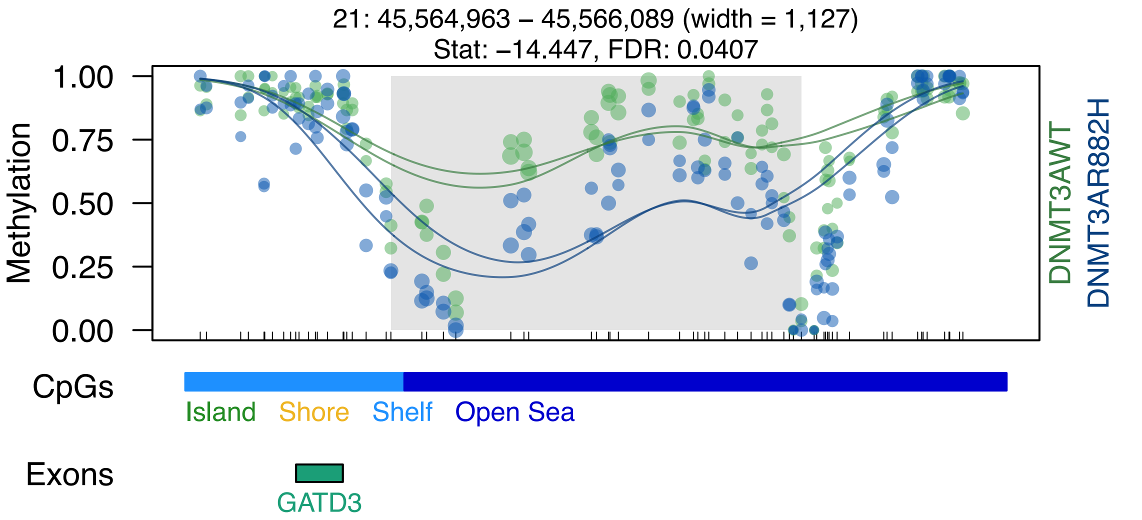
21: 45,564,963 – 45,566,089 (width = 1,127)
Stat: –14.447, FDR: 0.0407
Methylation
1.00
0.75
0.50
0.25
0.00
DNMT3AWT
DNMT3AR882H
CpGs
Island
Shore
Shelf
Open Sea
Exons
GATD3

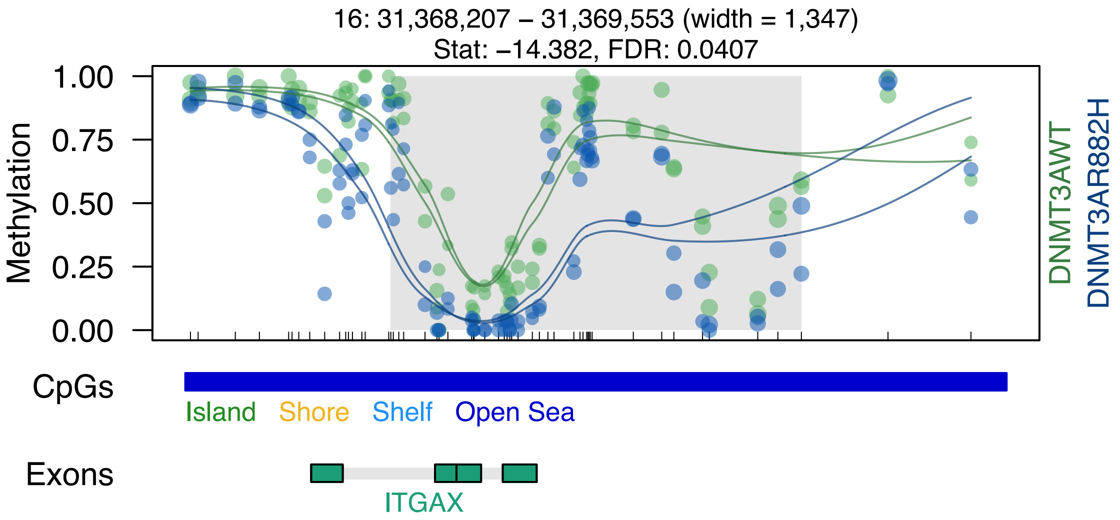
16: 31,368,207 – 31,369,553 (width = 1,347)
Stat: –14.382, FDR: 0.0407
Methylation
1.00
0.75
0.50
0.25
0.00
DNMT3AWT
DNMT3AR882H
CpGs
Island
Shore
Shelf
Open Sea
Exons
ITGAX

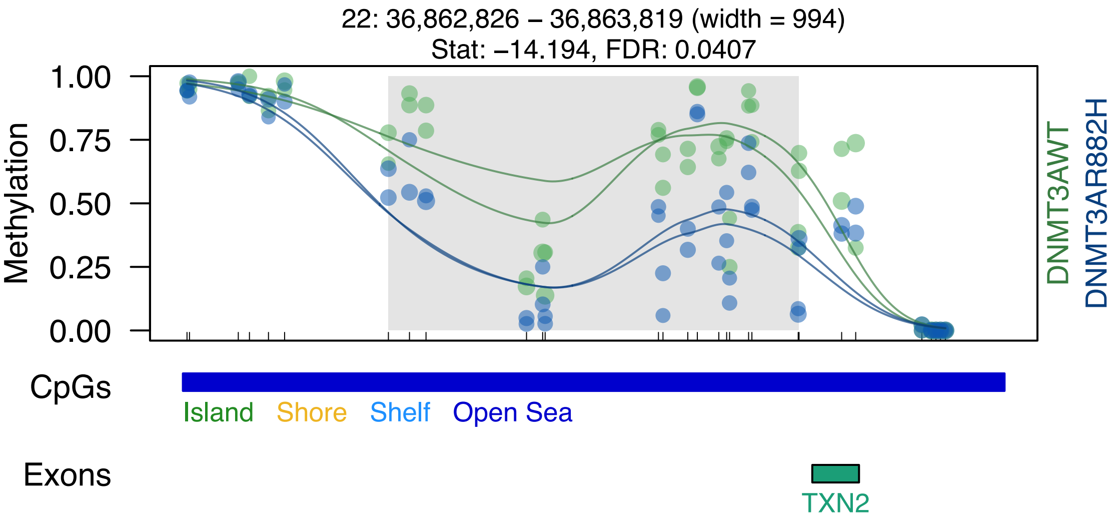
22: 36,862,826 – 36,863,819 (width = 994)
Stat: −14.194, FDR: 0.0407
1.00
0.75
0.50
0.25
0.00
Methylation
DNMT3AWT
DNMT3AR882H
CpGs
Island
Shore
Shelf
Open Sea
Exons
TXN2

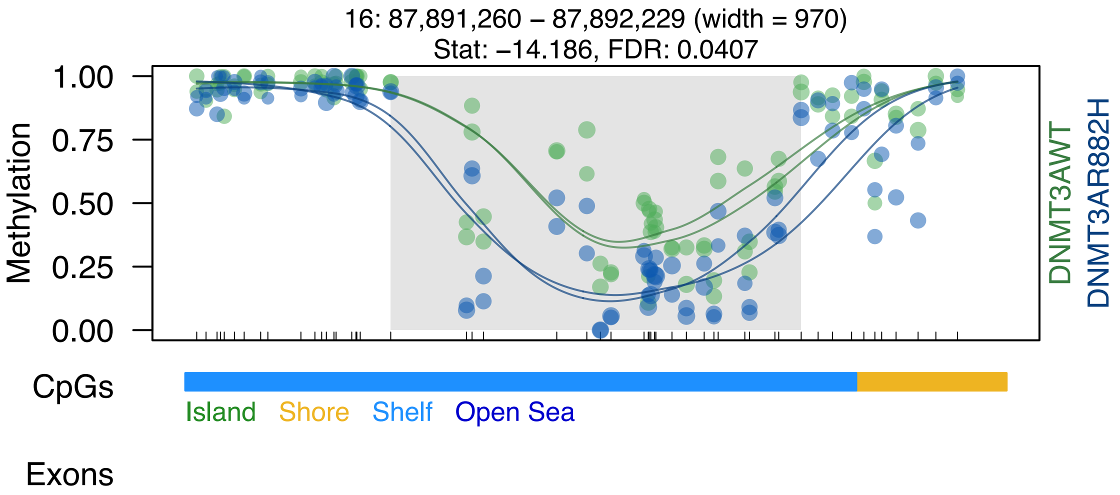
16: 87,891,260 – 87,892,229 (width = 970)
Stat: –14.186, FDR: 0.0407
Methylation
1.00
0.75
0.50
0.25
0.00
DNMT3AWT
DNMT3AR882H
CpGs
Island
Shore
Shelf
Open Sea
Exons

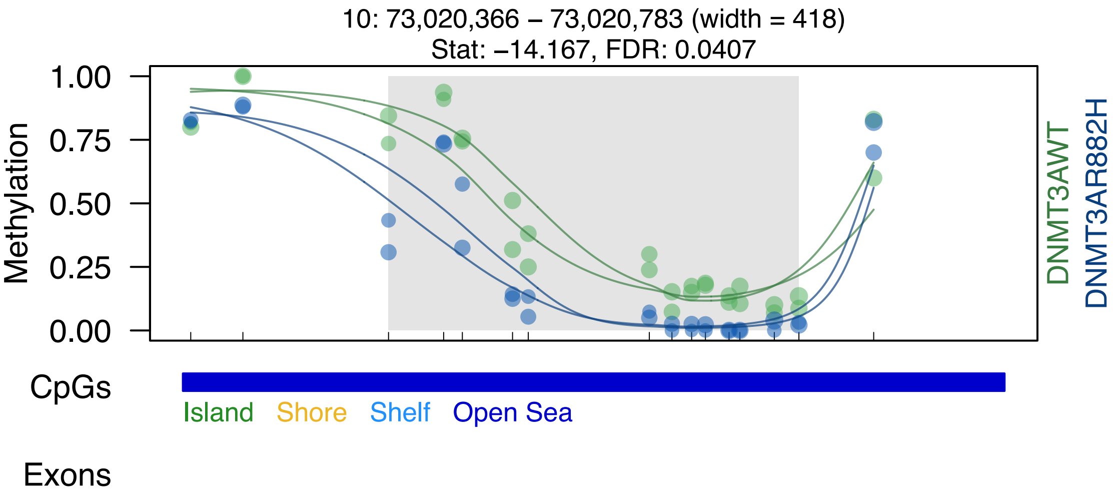
10: 73,020,366 – 73,020,783 (width = 418)
Stat: –14.167, FDR: 0.0407
1.00
0.75
0.50
0.25
0.00
Methylation
DNMT3AWT
DNMT3AR882H
CpGs
Island
Shore
Shelf
Open Sea
Exons

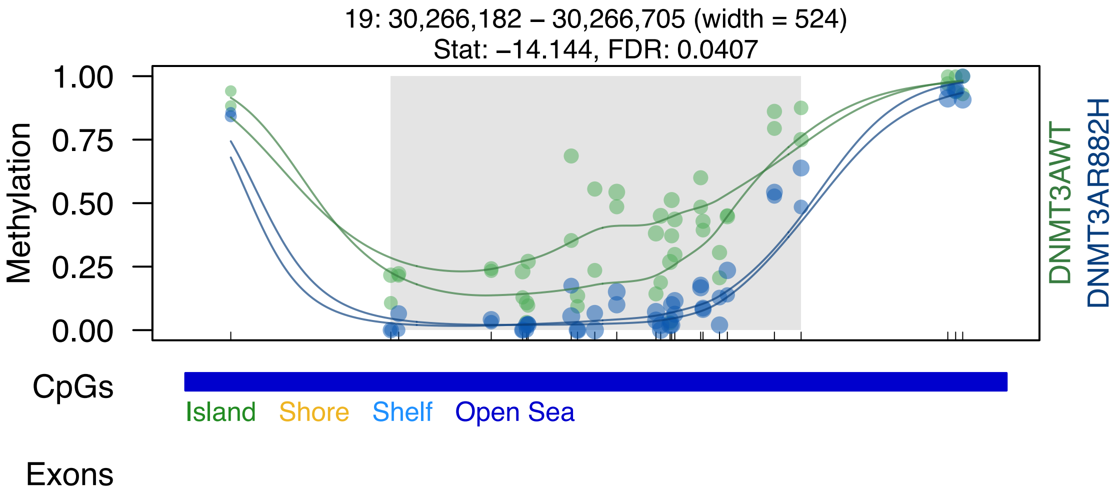

19: 30,266,182 – 30,266,705 (width = 524)
Stat: –14.144, FDR: 0.0407
1.00
0.75
0.50
0.25
0.00
Methylation
DNMT3AWT
DNMT3AR882H
CpGs
Island
Shore
Shelf
Open Sea
Exons

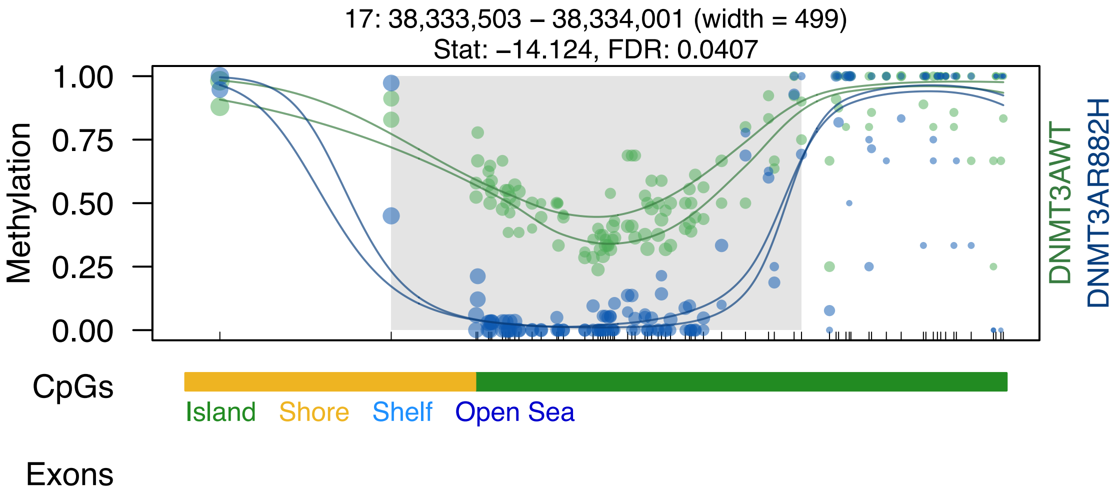
17: 38,333,503 – 38,334,001 (width = 499)
Stat: −14.124, FDR: 0.0407
Methylation
1.00
0.75
0.50
0.25
0.00
DNMT3AWT
DNMT3AR882H
CpGs
Island
Shore
Shelf
Open Sea
Exons

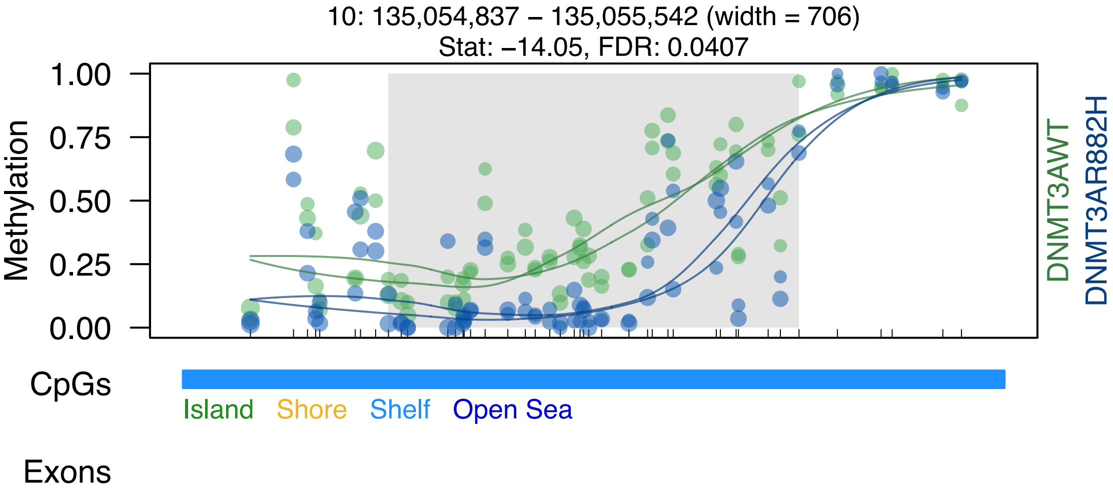
10: 135,054,837 – 135,055,542 (width = 706)
Stat: –14.05, FDR: 0.0407
Methylation
1.00
0.75
0.50
0.25
0.00
DNMT3AWT
DNMT3AR882H
CpGs
Island
Shore
Shelf
Open Sea
Exons

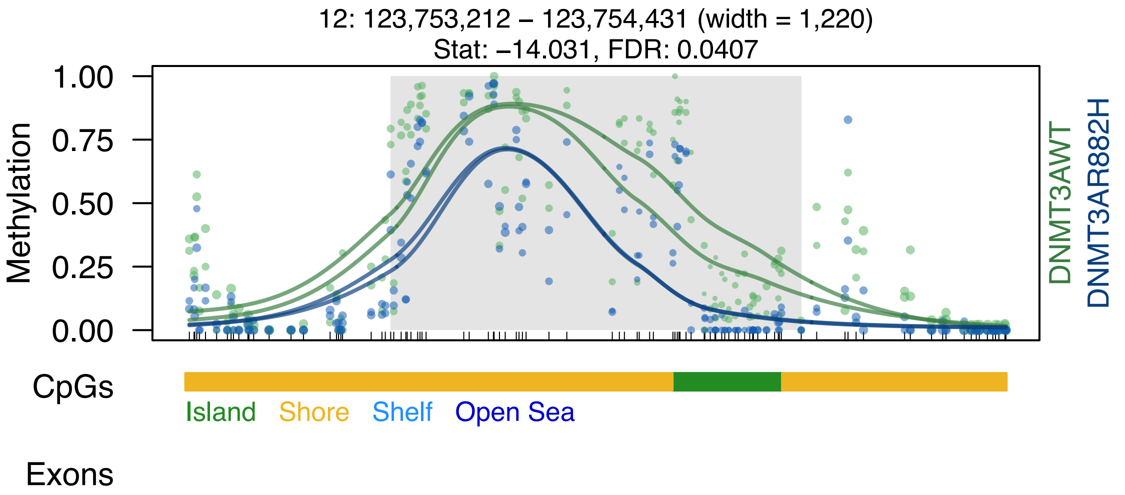

12: 123,753,212 – 123,754,431 (width = 1,220)
Stat: –14.031, FDR: 0.0407
Methylation
1.00
0.75
0.50
0.25
0.00
DNMT3AWT
DNMT3AR882H
CpGs
Island
Shore
Shelf
Open Sea
Exons

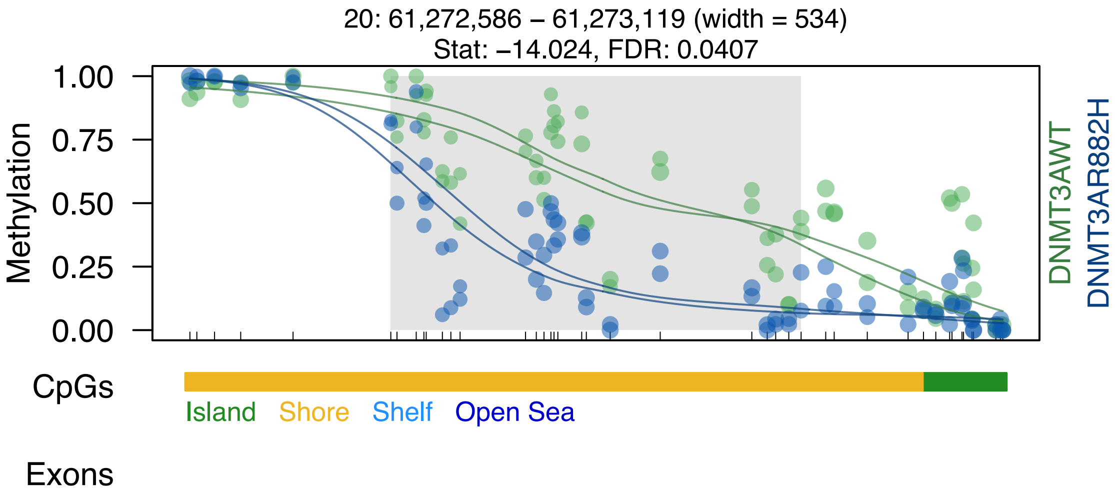

20: 61,272,586 – 61,273,119 (width = 534)
Stat: –14.024, FDR: 0.0407
Methylation
1.00
0.75
0.50
0.25
0.00
DNMT3AWT
DNMT3AR882H
CpGs
Island
Shore
Shelf
Open Sea
Exons

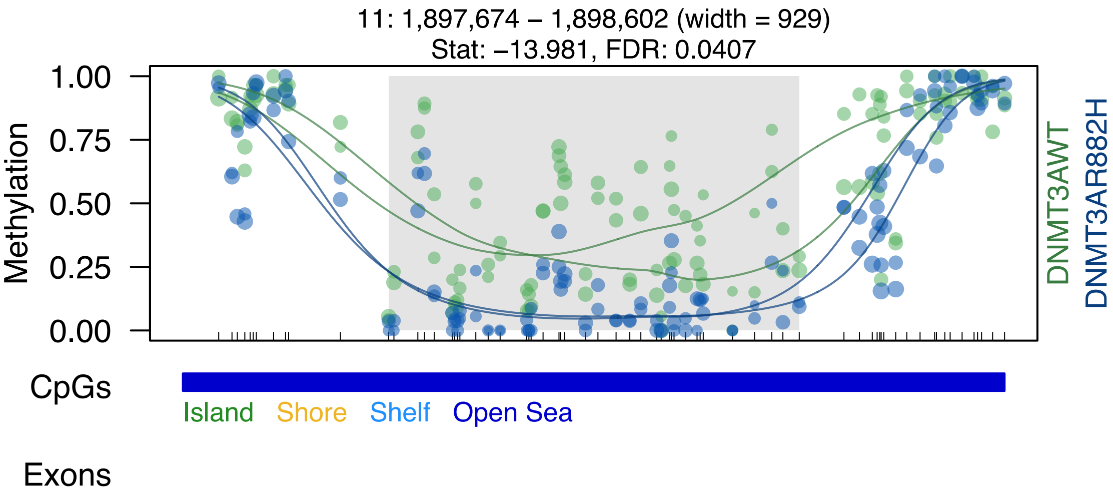

11: 1,897,674 – 1,898,602 (width = 929)
Stat: –13.981, FDR: 0.0407
Methylation
1.00
0.75
0.50
0.25
0.00
DNMT3AWT
DNMT3AR882H
CpGs
Island
Shore
Shelf
Open Sea
Exons

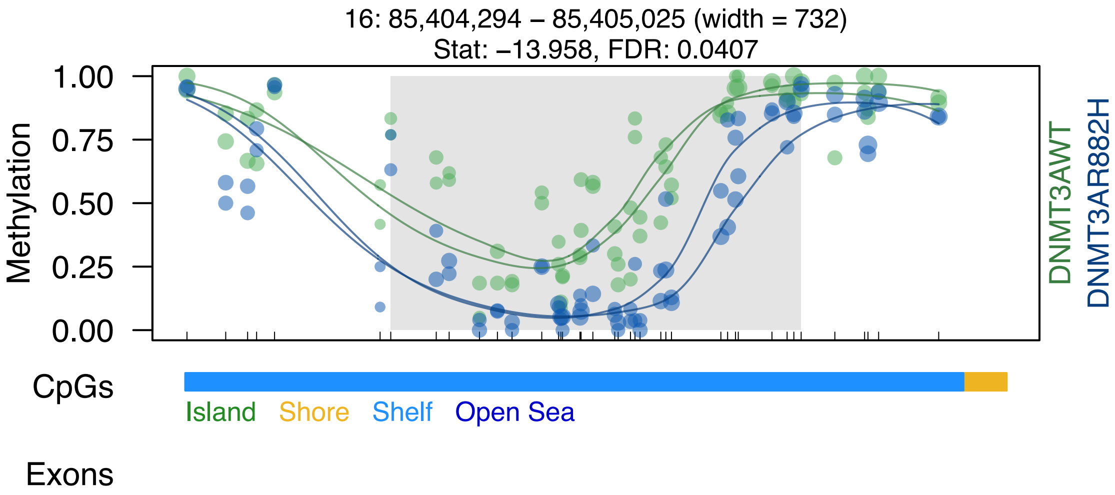
16: 85,404,294 – 85,405,025 (width = 732)
Stat: −13.958, FDR: 0.0407
Methylation
1.00
0.75
0.50
0.25
0.00
DNMT3AWT
DNMT3AR882H
CpGs
Island
Shore
Shelf
Open Sea
Exons

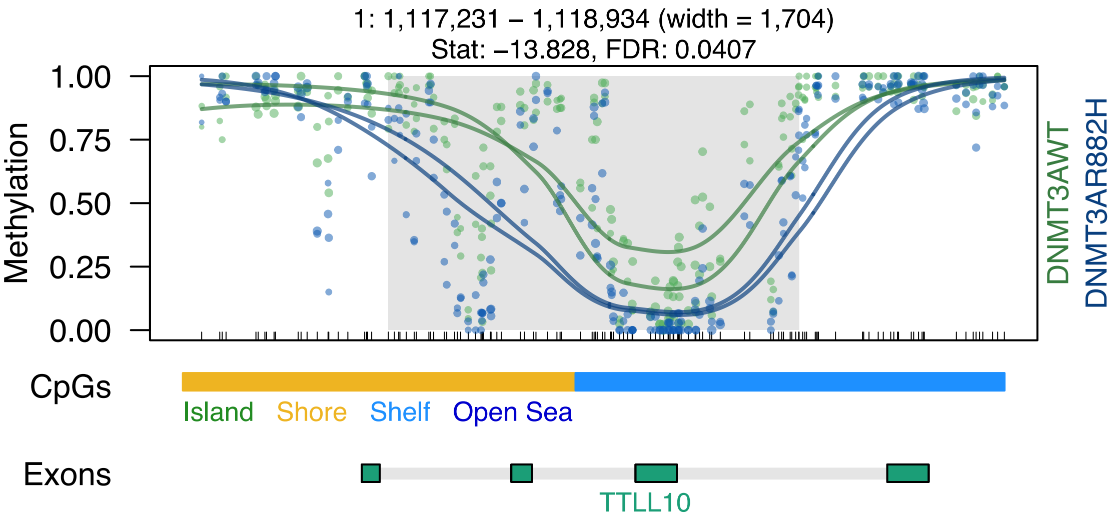

1: 1,117,231 – 1,118,934 (width = 1,704)
Stat: –13.828, FDR: 0.0407
Methylation
1.00
0.75
0.50
0.25
0.00
DNMT3AWT
DNMT3AR882H
CpGs
Island
Shore
Shelf
Open Sea
Exons
TTLL10

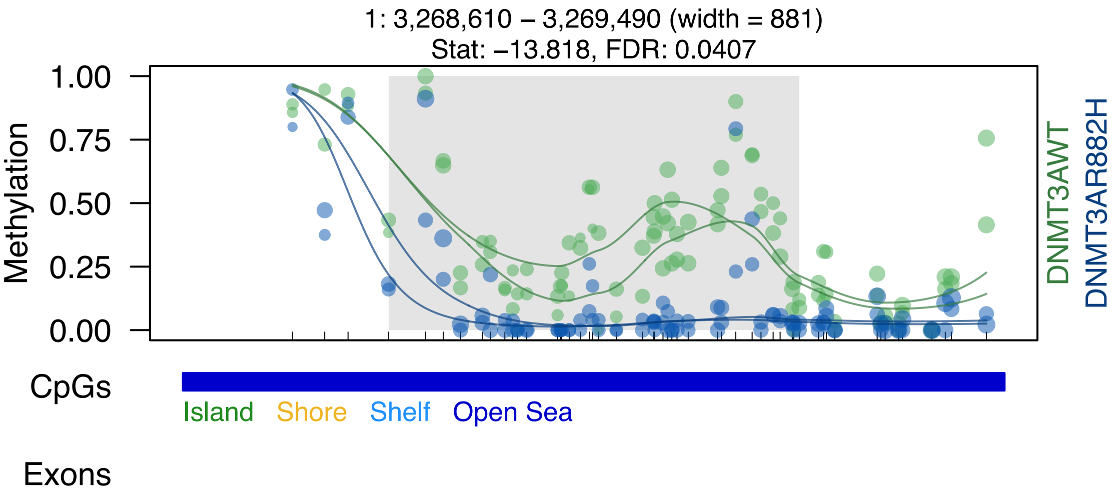
1: 3,268,610 – 3,269,490 (width = 881)
Stat: –13.818, FDR: 0.0407
Methylation
1.00
0.75
0.50
0.25
0.00
DNMT3AWT
DNMT3AR882H
CpGs
Island
Shore
Shelf
Open Sea
Exons

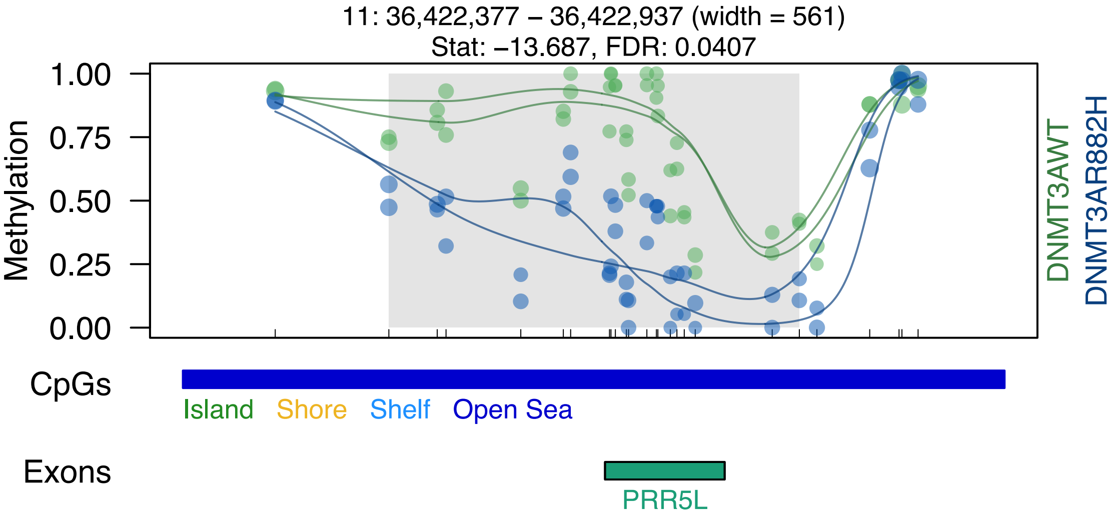
11: 36,422,377 – 36,422,937 (width = 561)
Stat: −13.687, FDR: 0.0407
Methylation
1.00
0.75
0.50
0.25
0.00
DNMT3AWT
DNMT3AR882H
CpGs
Island
Shore
Shelf
Open Sea
Exons
PRR5L

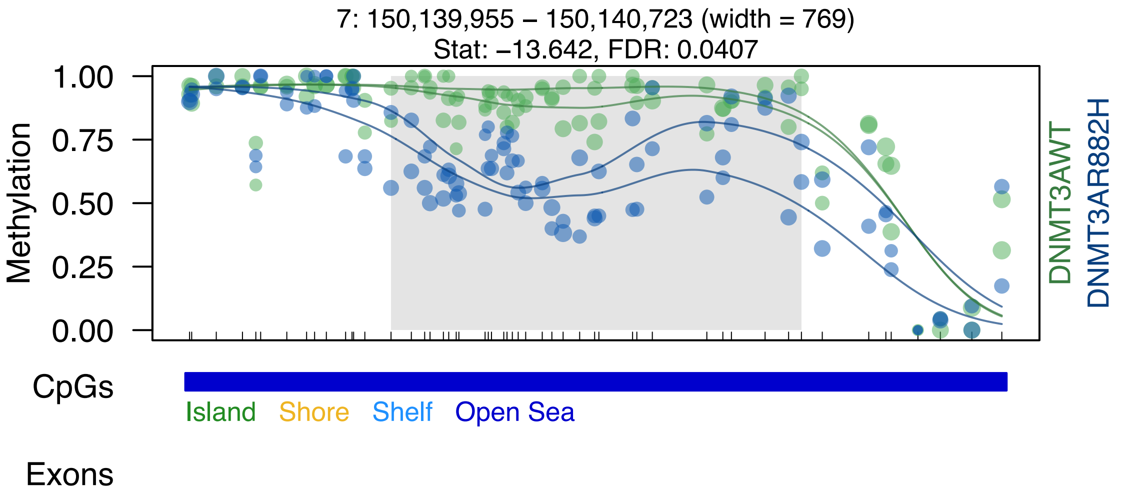

7: 150,139,955 – 150,140,723 (width = 769)
Stat: −13.642, FDR: 0.0407
Methylation
1.00
0.75
0.50
0.25
0.00
DNMT3AWT
DNMT3AR882H
CpGs
Island
Shore
Shelf
Open Sea
Exons

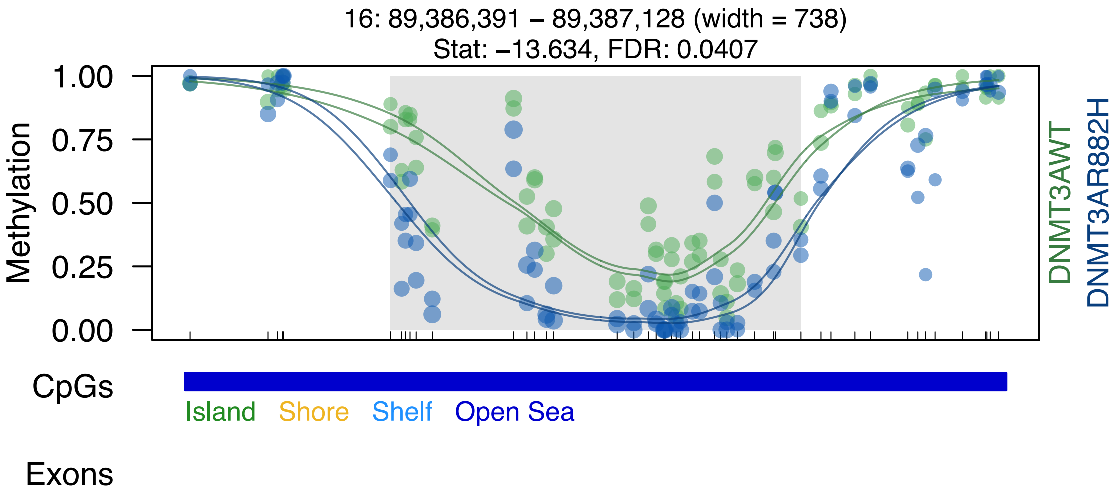
16: 89,386,391 – 89,387,128 (width = 738)
Stat: –13.634, FDR: 0.0407
Methylation
1.00
0.75
0.50
0.25
0.00
DNMT3AWT
DNMT3AR882H
CpGs
Island
Shore
Shelf
Open Sea
Exons

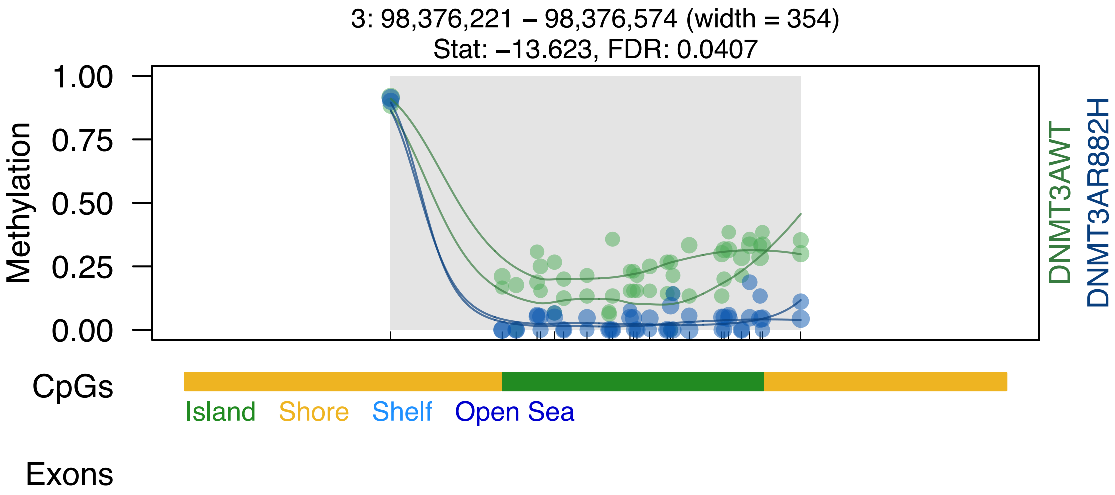
3: 98,376,221 – 98,376,574 (width = 354)
Stat: –13.623, FDR: 0.0407
Methylation
1.00
0.75
0.50
0.25
0.00
DNMT3AWT
DNMT3AR882H
CpGs
Island
Shore
Shelf
Open Sea
Exons

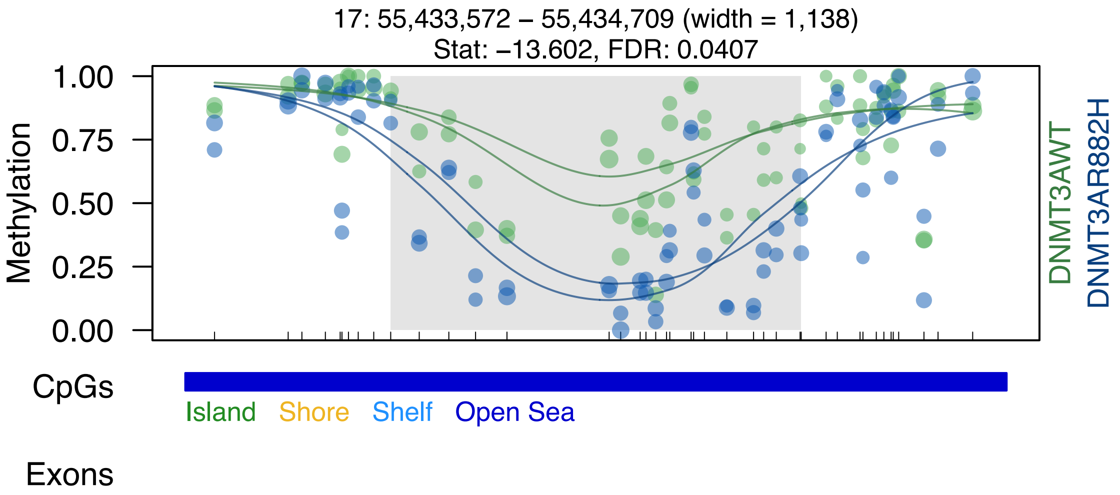
17: 55,433,572 – 55,434,709 (width = 1,138)
Stat: −13.602, FDR: 0.0407
Methylation
1.00
0.75
0.50
0.25
0.00
DNMT3AWT
DNMT3AR882H
CpGs
Island
Shore
Shelf
Open Sea
Exons

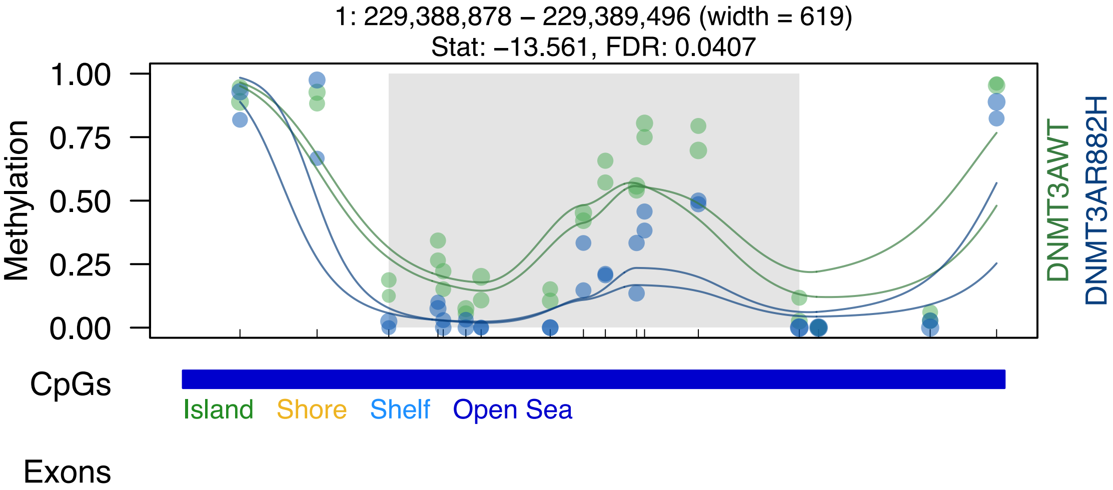
1: 229,388,878 – 229,389,496 (width = 619)
Stat: –13.561, FDR: 0.0407
Methylation
1.00
0.75
0.50
0.25
0.00
DNMT3AWT
DNMT3AR882H
CpGs
Island
Shore
Shelf
Open Sea
Exons

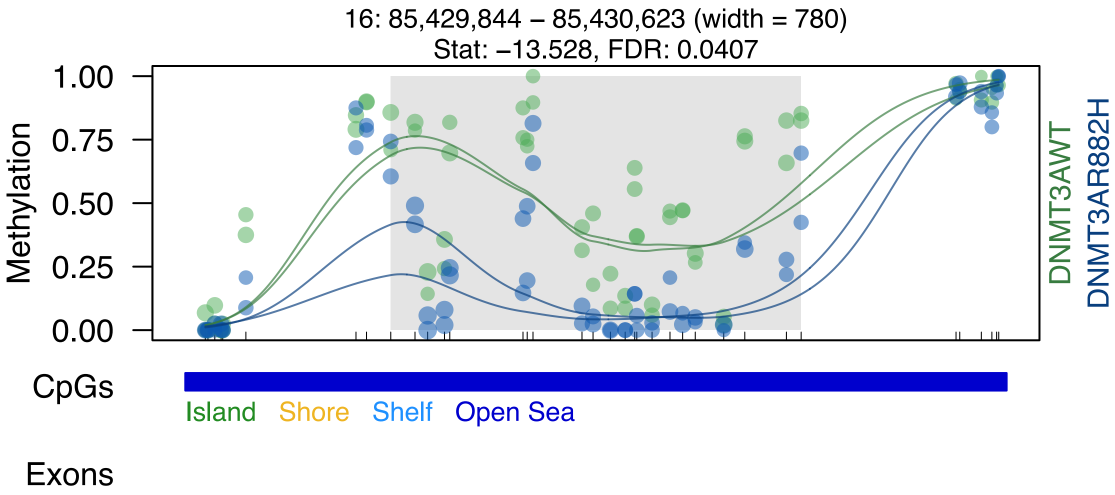
16: 85,429,844 – 85,430,623 (width = 780)
Stat: –13.528, FDR: 0.0407
Methylation
1.00
0.75
0.50
0.25
0.00
DNMT3AWT
DNMT3AR882H
CpGs
Island
Shore
Shelf
Open Sea
Exons

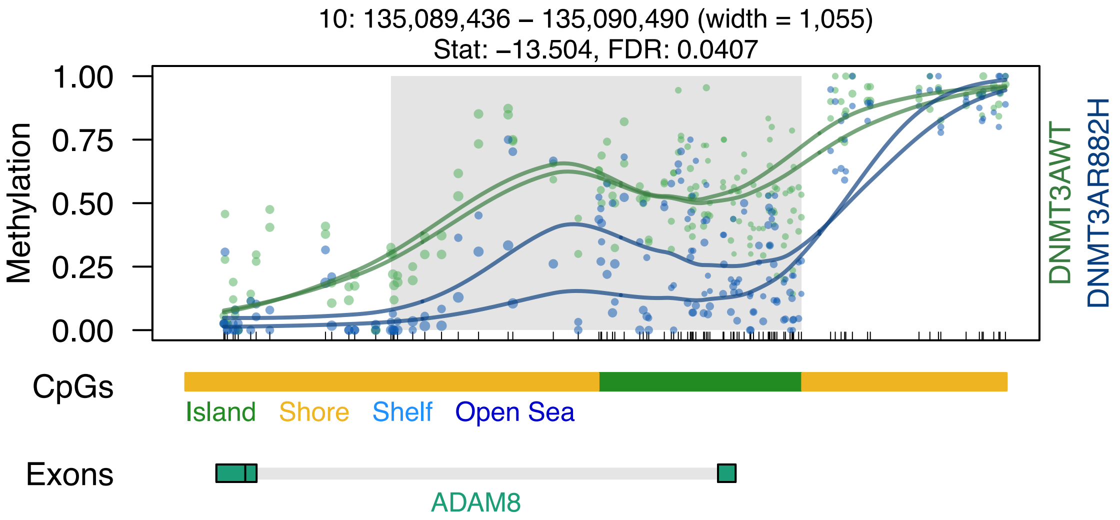

10: 135,089,436 – 135,090,490 (width = 1,055)
Stat: −13.504, FDR: 0.0407
Methylation
1.00
0.75
0.50
0.25
0.00
DNMT3AWT
DNMT3AR882H
CpGs
Island
Shore
Shelf
Open Sea
Exons
ADAM8

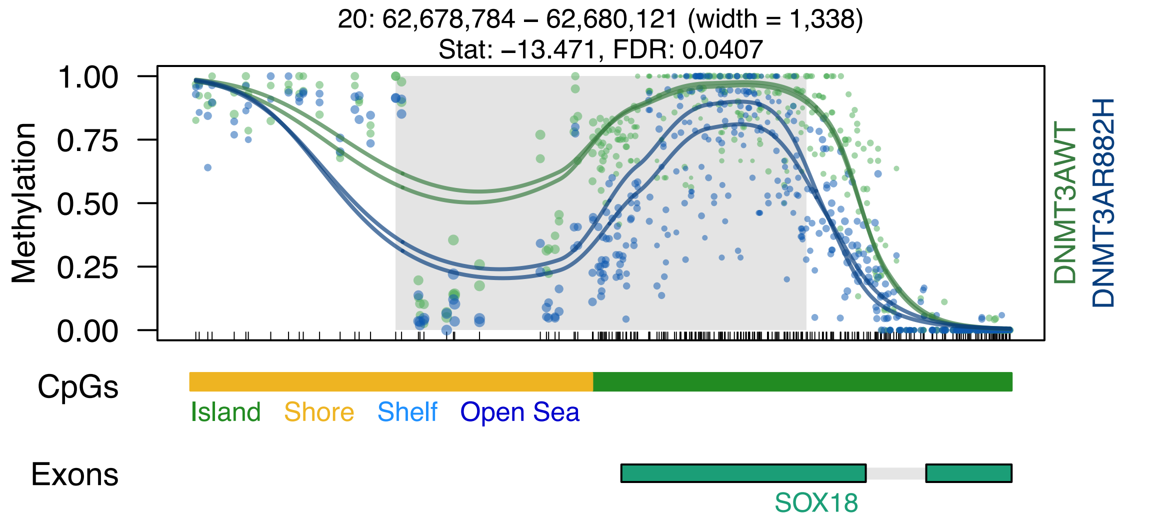

20: 62,678,784 – 62,680,121 (width = 1,338)
Stat: –13.471, FDR: 0.0407
1.00
0.75
0.50
0.25
0.00
Methylation
DNMT3AWT
DNMT3AR882H
CpGs
Island
Shore
Shelf
Open Sea
Exons
SOX18

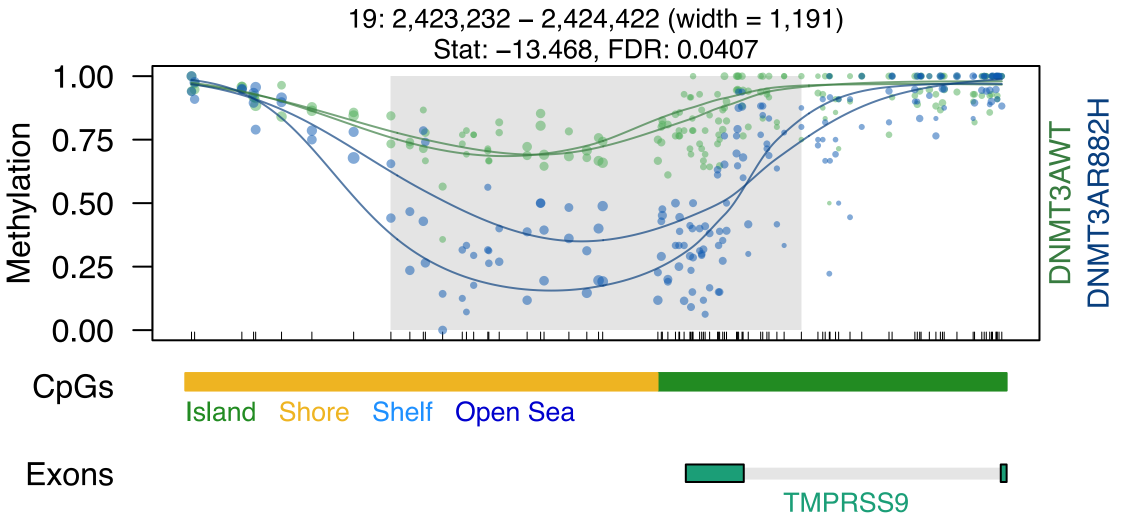
19: 2,423,232 – 2,424,422 (width = 1,191)
Stat: −13.468, FDR: 0.0407
Methylation
1.00
0.75
0.50
0.25
0.00
DNMT3AWT
DNMT3AR882H
CpGs
Island
Shore
Shelf
Open Sea
Exons
TMPRSS9

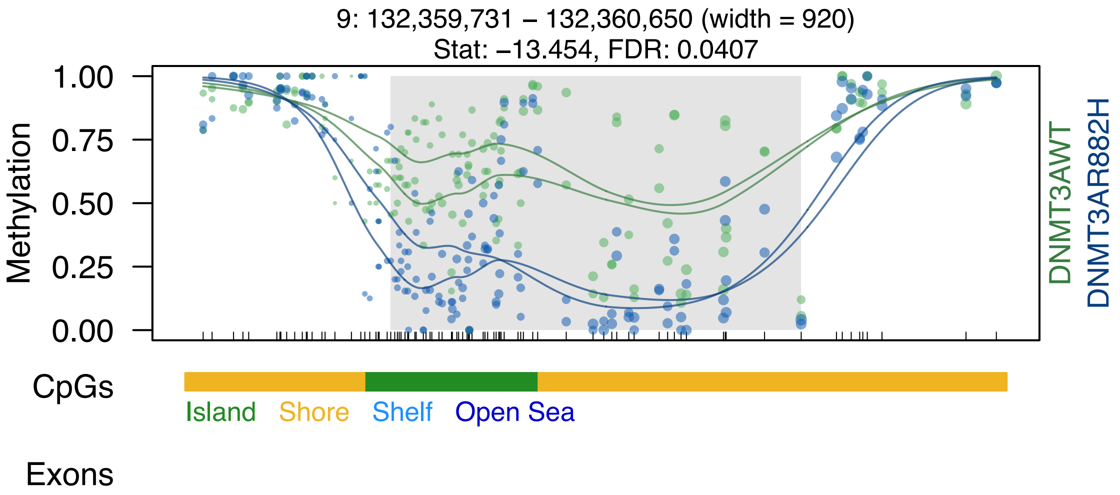

9: 132,359,731 – 132,360,650 (width = 920)
Stat: –13.454, FDR: 0.0407
Methylation
1.00
0.75
0.50
0.25
0.00
DNMT3AWT
DNMT3AR882H
CpGs
Island
Shore
Shelf
Open Sea
Exons

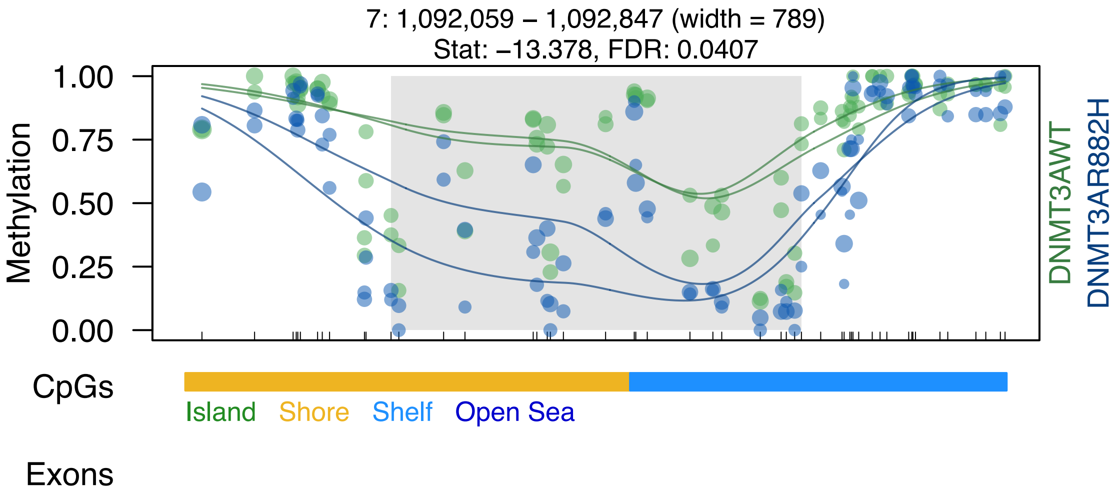
7: 1,092,059 – 1,092,847 (width = 789)
Stat: −13.378, FDR: 0.0407
Methylation
1.00
0.75
0.50
0.25
0.00
DNMT3AWT
DNMT3AR882H
CpGs
Island
Shore
Shelf
Open Sea
Exons

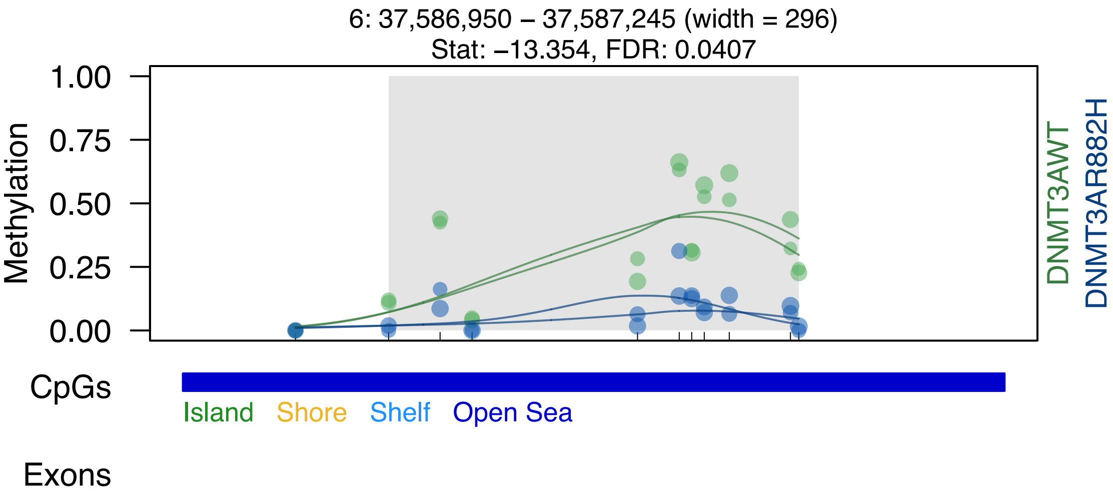
6: 37,586,950 – 37,587,245 (width = 296)
Stat: –13.354, FDR: 0.0407
Methylation
1.00
0.75
0.50
0.25
0.00
DNMT3AWT
DNMT3AR882H
CpGs
Island
Shore
Shelf
Open Sea
Exons

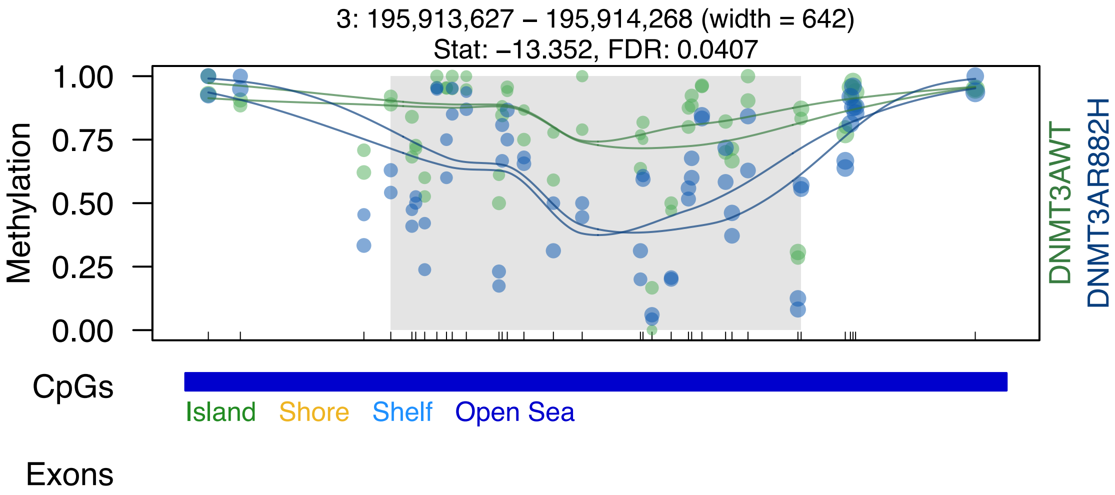

3: 195,913,627 – 195,914,268 (width = 642)
Stat: –13.352, FDR: 0.0407
Methylation
1.00
0.75
0.50
0.25
0.00
DNMT3AWT
DNMT3AR882H
CpGs
Island
Shore
Shelf
Open Sea
Exons

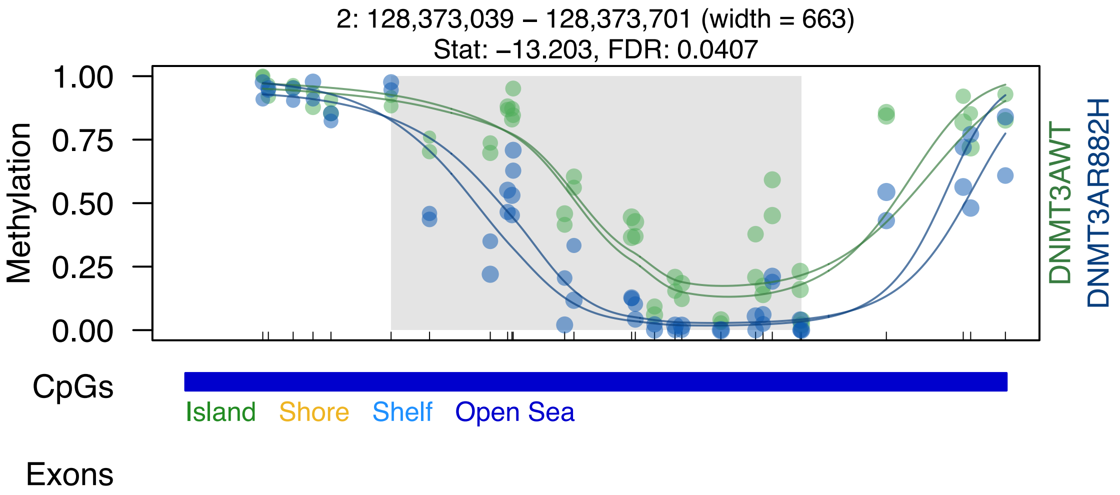
2: 128,373,039 – 128,373,701 (width = 663)
Stat: –13.203, FDR: 0.0407
Methylation
1.00
0.75
0.50
0.25
0.00
DNMT3AWT
DNMT3AR882H
CpGs
Island
Shore
Shelf
Open Sea
Exons

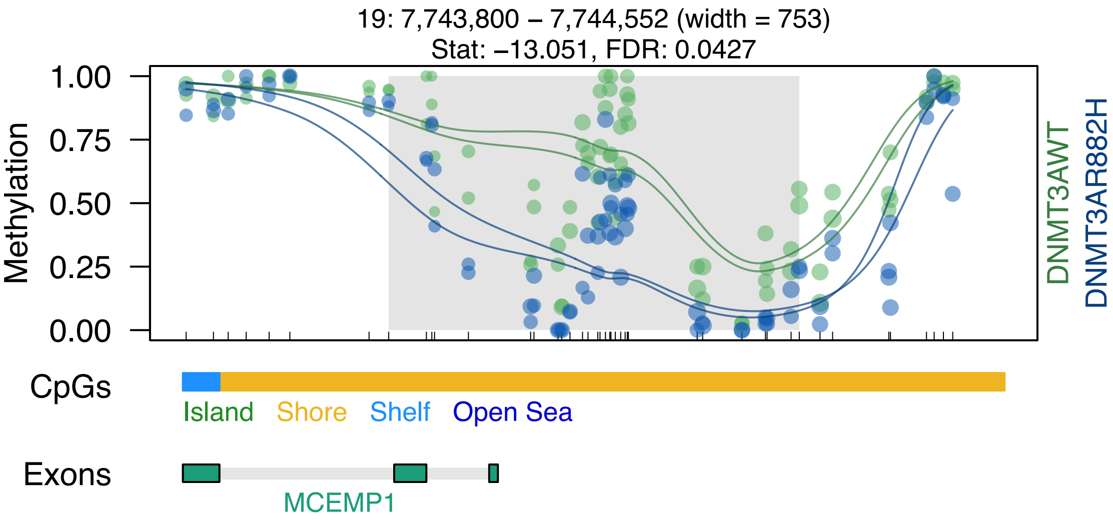

19: 7,743,800 – 7,744,552 (width = 753)
Stat: −13.051, FDR: 0.0427
Methylation
1.00
0.75
0.50
0.25
0.00
DNMT3AWT
DNMT3AR882H
CpGs
Island
Shore
Shelf
Open Sea
Exons
MCEMP1

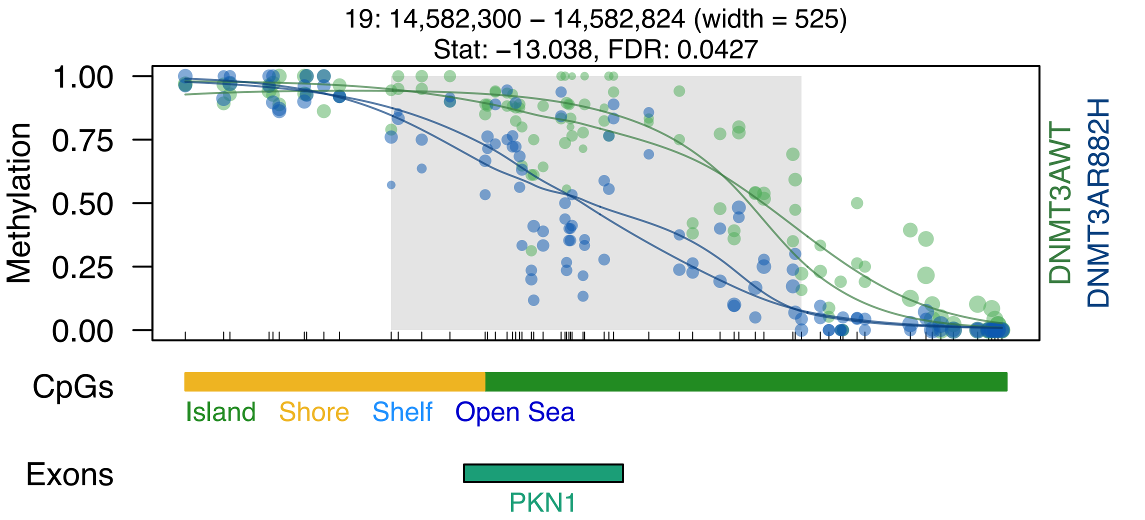
19: 14,582,300 – 14,582,824 (width = 525)
Stat: –13.038, FDR: 0.0427
Methylation
1.00
0.75
0.50
0.25
0.00
DNMT3AWT
DNMT3AR882H
CpGs
Island
Shore
Shelf
Open Sea
Exons
PKN1

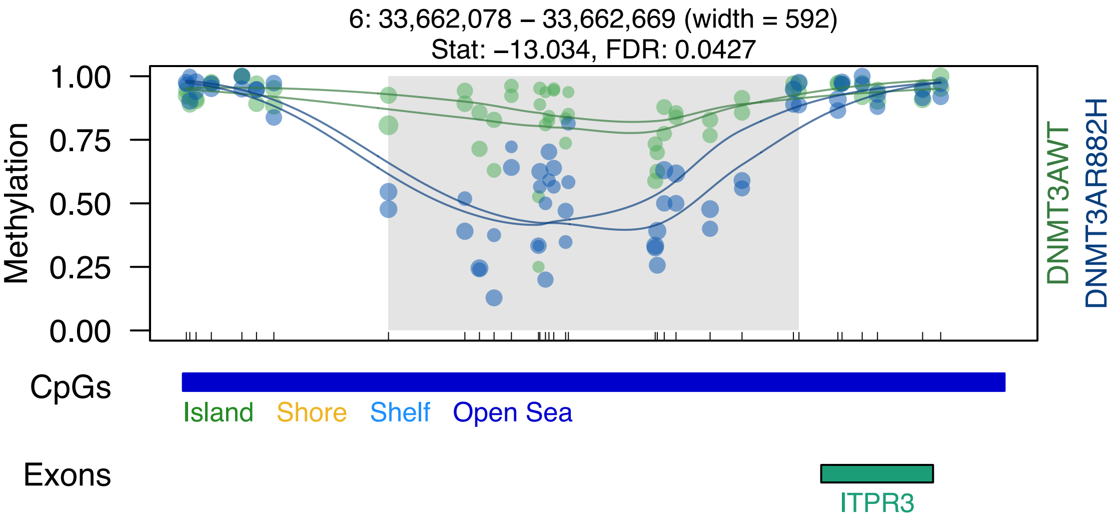
6: 33,662,078 – 33,662,669 (width = 592)
Stat: –13.034, FDR: 0.0427
1.00
0.75
0.50
0.25
0.00
Methylation
DNMT3AWT
DNMT3AR882H
CpGs
Island
Shore
Shelf
Open Sea
Exons
ITPR3

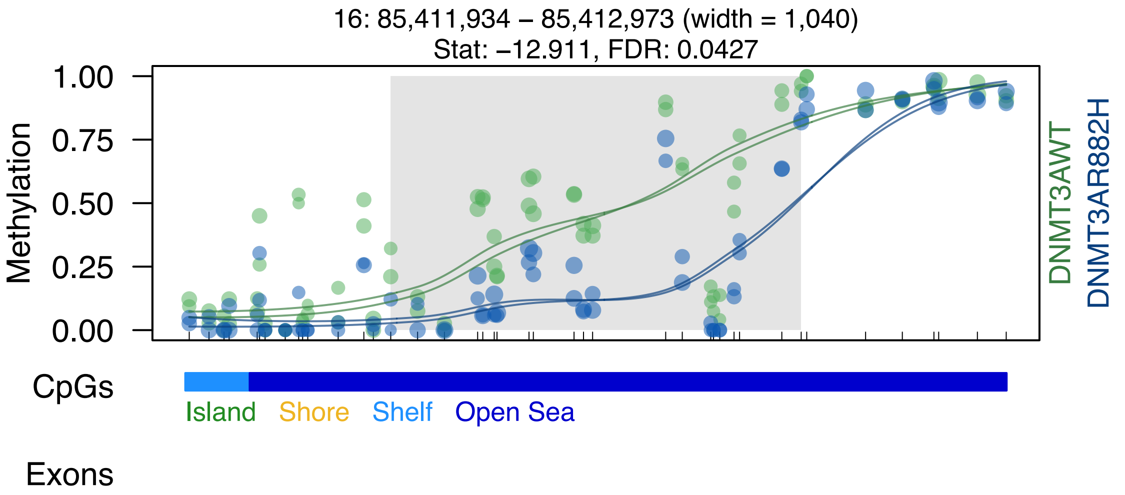
16: 85,411,934 – 85,412,973 (width = 1,040)
Stat: –12.911, FDR: 0.0427
1.00
0.75
0.50
0.25
0.00
Methylation
DNMT3AWT
DNMT3AR882H
CpGs
Island
Shore
Shelf
Open Sea
Exons

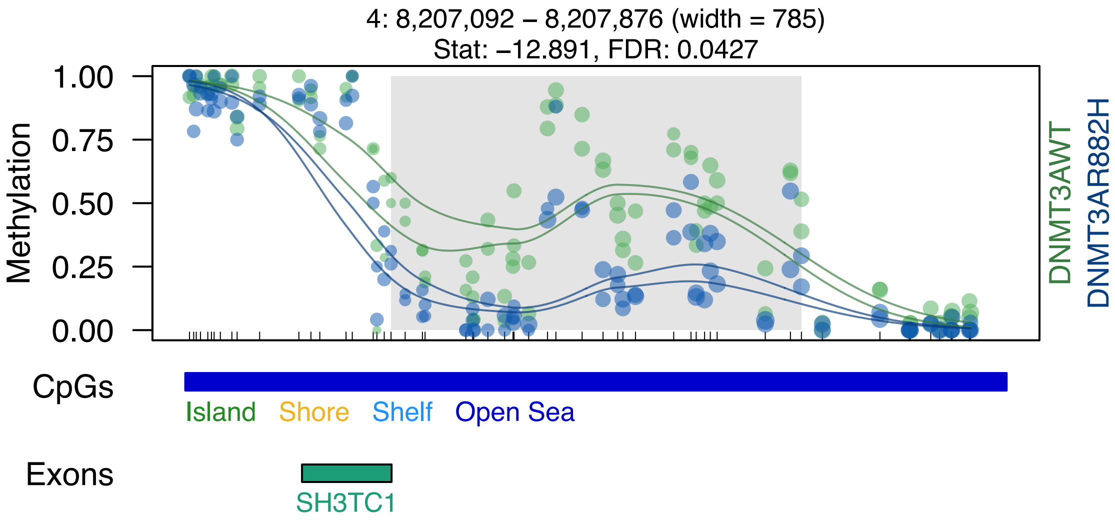

4: 8,207,092 – 8,207,876 (width = 785)
Stat: –12.891, FDR: 0.0427
Methylation
1.00
0.75
0.50
0.25
0.00
DNMT3AWT
DNMT3AR882H
CpGs
Island
Shore
Shelf
Open Sea
Exons
SH3TC1

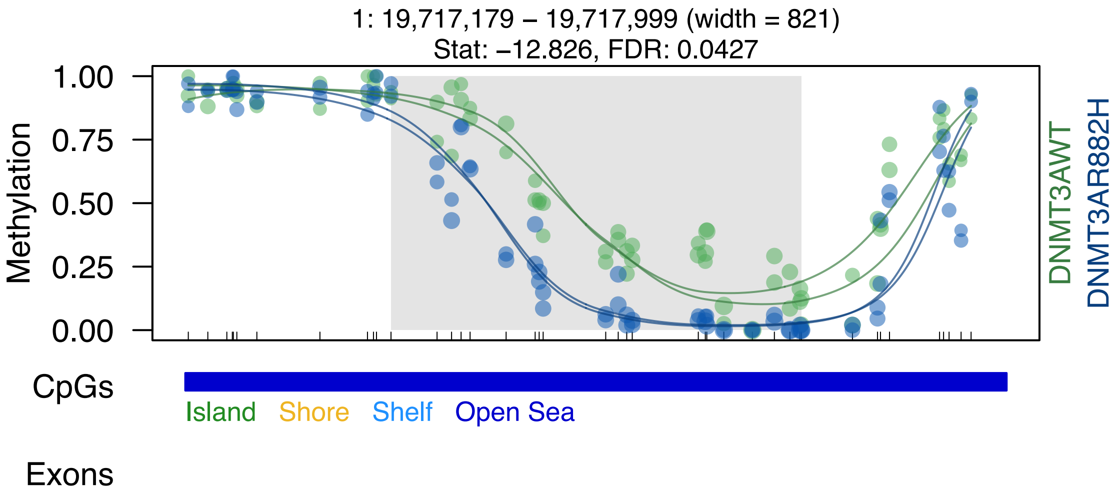

1: 19,717,179 – 19,717,999 (width = 821)
Stat: –12.826, FDR: 0.0427
1.00
0.75
0.50
0.25
0.00
Methylation
DNMT3AWT
DNMT3AR882H
CpGs
Island
Shore
Shelf
Open Sea
Exons

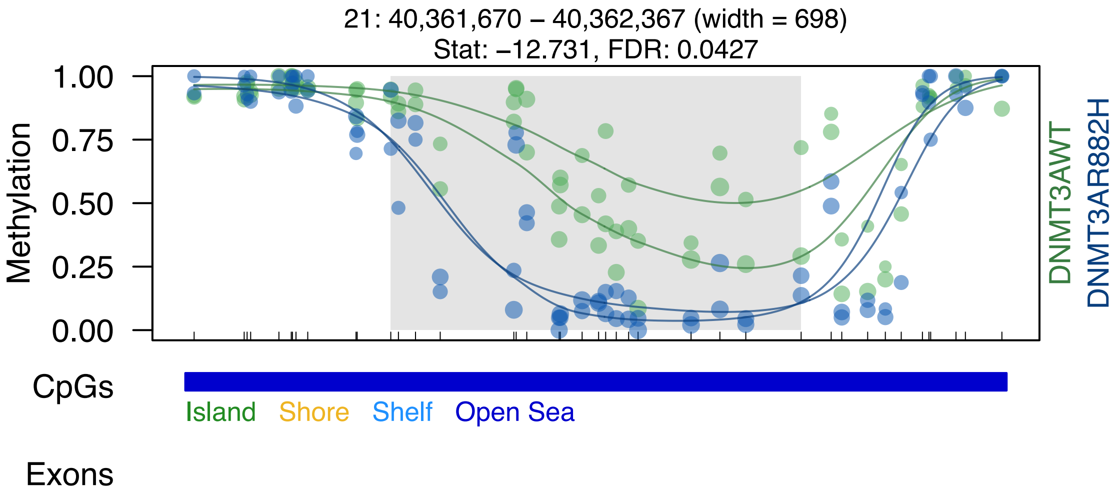
21: 40,361,670 – 40,362,367 (width = 698)
Stat: –12.731, FDR: 0.0427
Methylation
1.00
0.75
0.50
0.25
0.00
DNMT3AWT
DNMT3AR882H
CpGs
Island
Shore
Shelf
Open Sea
Exons

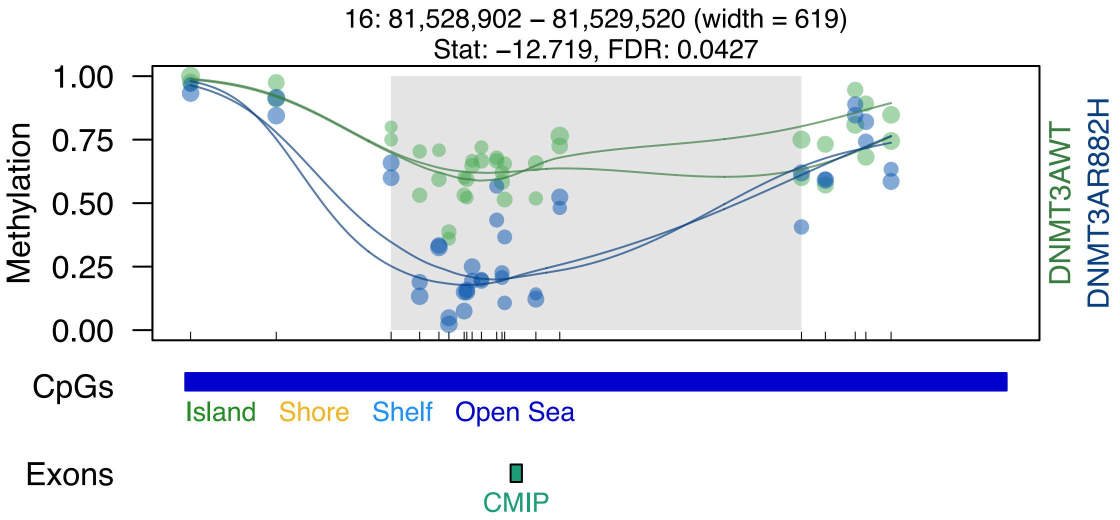
16: 81,528,902 – 81,529,520 (width = 619)
Stat: –12.719, FDR: 0.0427
1.00
0.75
0.50
0.25
0.00
Methylation
DNMT3AWT
DNMT3AR882H
CpGs
Island
Shore
Shelf
Open Sea
Exons
CMIP

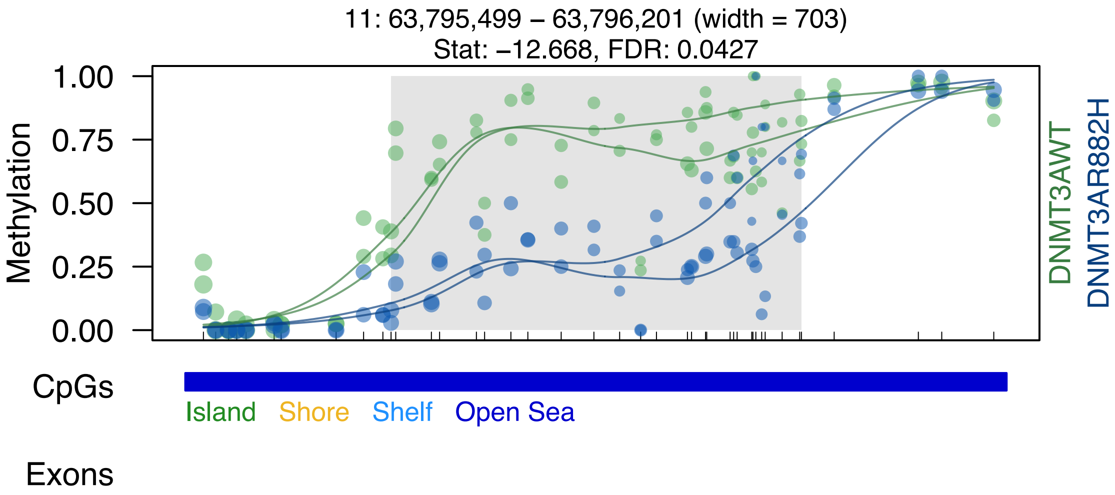
11: 63,795,499 – 63,796,201 (width = 703)
Stat: –12.668, FDR: 0.0427
1.00
0.75
0.50
0.25
0.00
Methylation
DNMT3AWT
DNMT3AR882H
CpGs
Island
Shore
Shelf
Open Sea
Exons

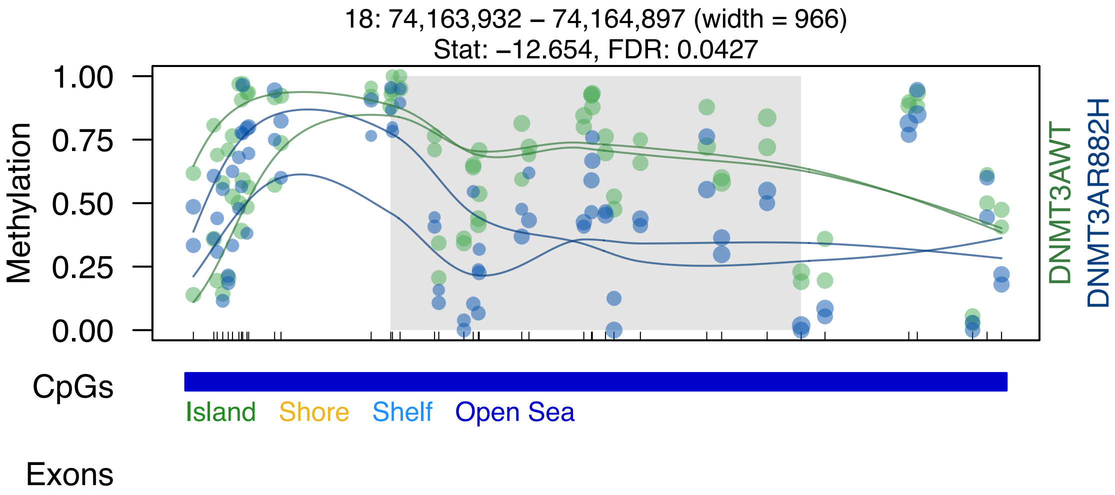

18: 74,163,932 – 74,164,897 (width = 966)
Stat: −12.654, FDR: 0.0427
1.00
0.75
0.50
0.25
0.00
Methylation
DNMT3AWT
DNMT3AR882H
CpGs
Island
Shore
Shelf
Open Sea
Exons

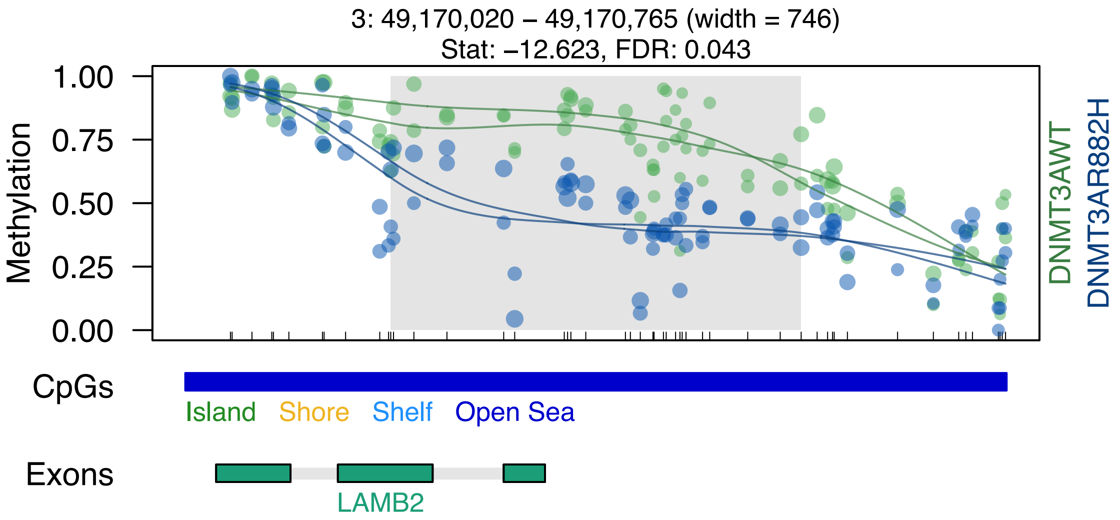
3: 49,170,020 – 49,170,765 (width = 746)
Stat: –12.623, FDR: 0.043
Methylation
1.00
0.75
0.50
0.25
0.00
DNMT3AWT
DNMT3AR882H
CpGs
Island
Shore
Shelf
Open Sea
Exons
LAMB2

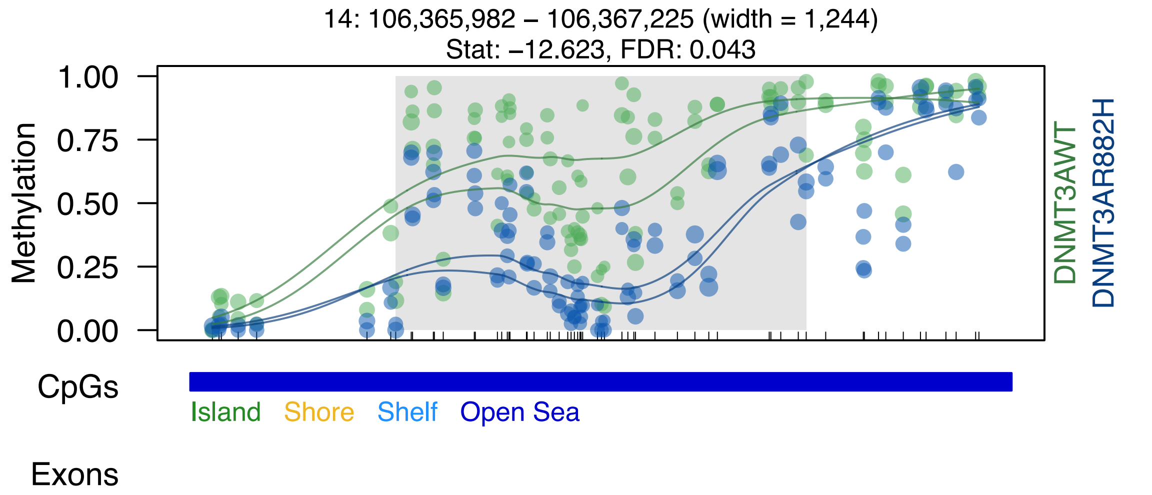
14: 106,365,982 – 106,367,225 (width = 1,244)
Stat: −12.623, FDR: 0.043
1.00
0.75
0.50
0.25
0.00
Methylation
DNMT3AWT
DNMT3AR882H
CpGs
Island
Shore
Shelf
Open Sea
Exons

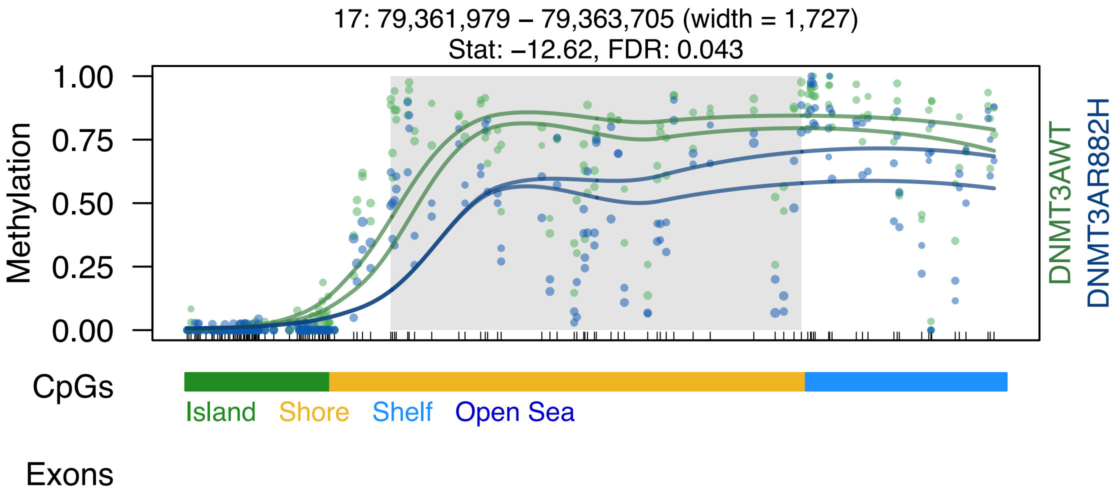
17: 79,361,979 – 79,363,705 (width = 1,727)
Stat: –12.62, FDR: 0.043
Methylation
1.00
0.75
0.50
0.25
0.00
DNMT3AWT
DNMT3AR882H
CpGs
Island
Shore
Shelf
Open Sea
Exons

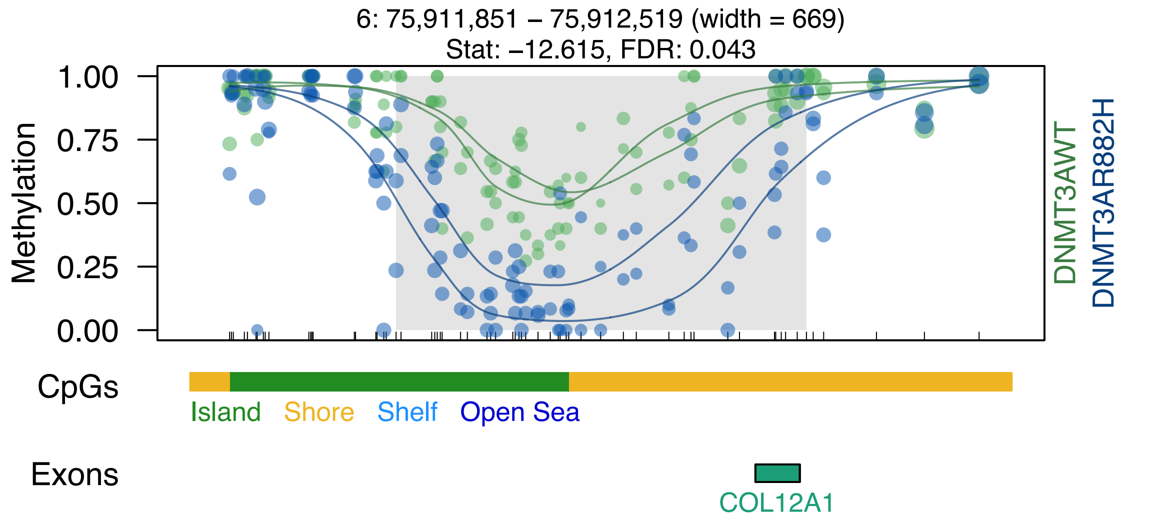

6: 75,911,851 – 75,912,519 (width = 669)
Stat: –12.615, FDR: 0.043
Methylation
1.00
0.75
0.50
0.25
0.00
DNMT3AWT
DNMT3AR882H
CpGs
Island
Shore
Shelf
Open Sea
Exons
COL12A1

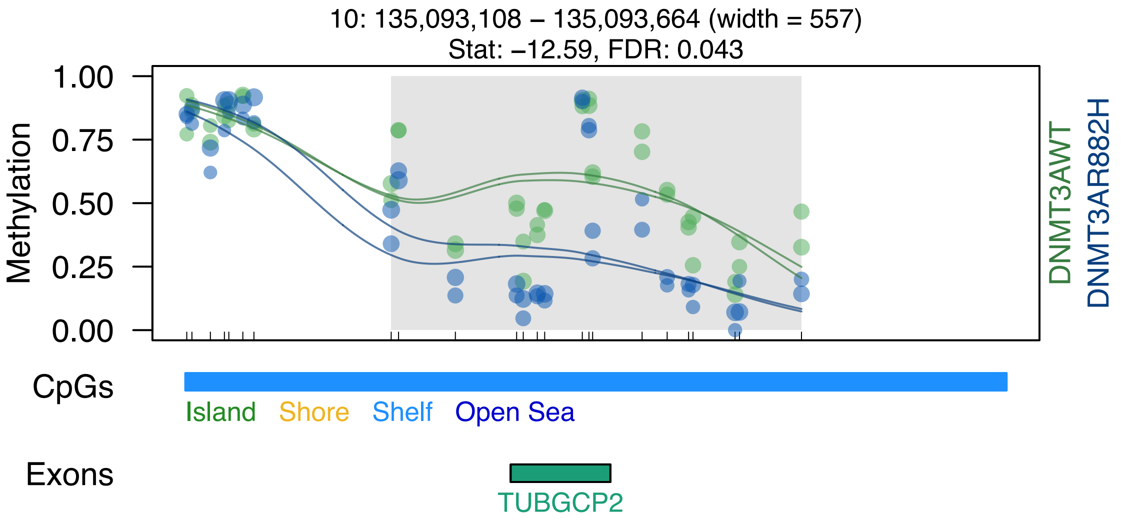

10: 135,093,108 – 135,093,664 (width = 557)
Stat: –12.59, FDR: 0.043
Methylation
1.00
0.75
0.50
0.25
0.00
DNMT3AWT
DNMT3AR882H
CpGs
Island
Shore
Shelf
Open Sea
Exons
TUBGCP2

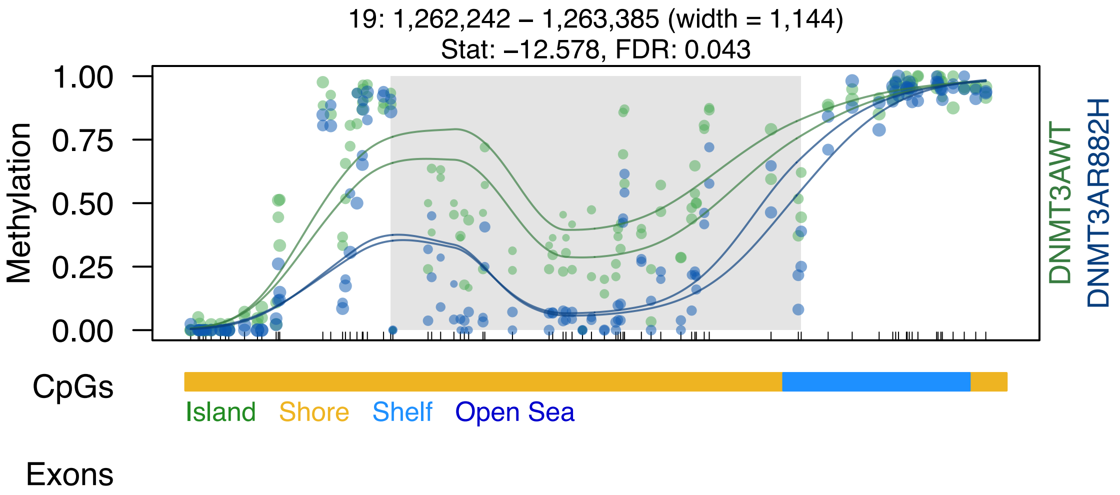

19: 1,262,242 – 1,263,385 (width = 1,144)
Stat: –12.578, FDR: 0.043
Methylation
1.00
0.75
0.50
0.25
0.00
DNMT3AWT
DNMT3AR882H
CpGs
Island
Shore
Shelf
Open Sea
Exons

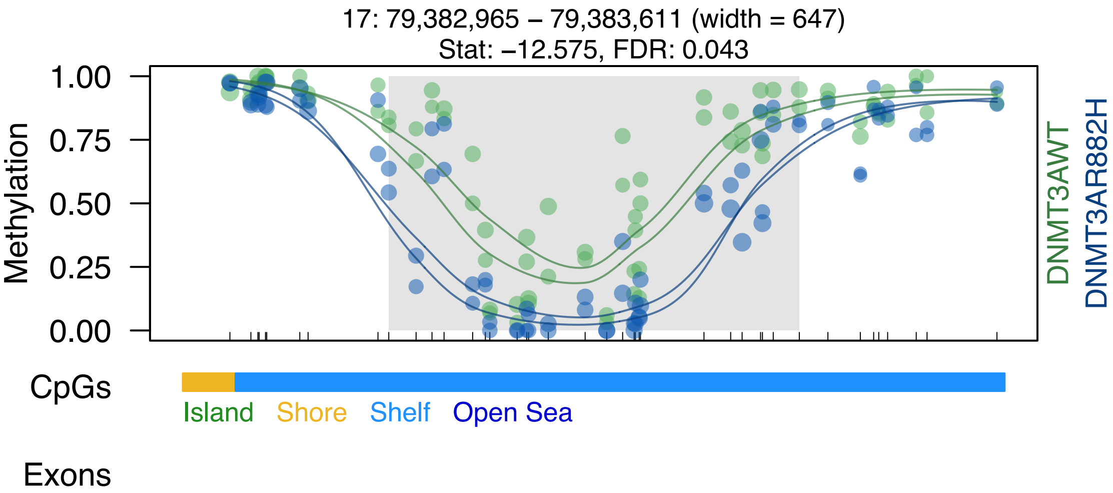

17: 79,382,965 – 79,383,611 (width = 647)
Stat: –12.575, FDR: 0.043
1.00
0.75
0.50
0.25
0.00
Methylation
DNMT3AWT
DNMT3AR882H
CpGs
Island
Shore
Shelf
Open Sea
Exons

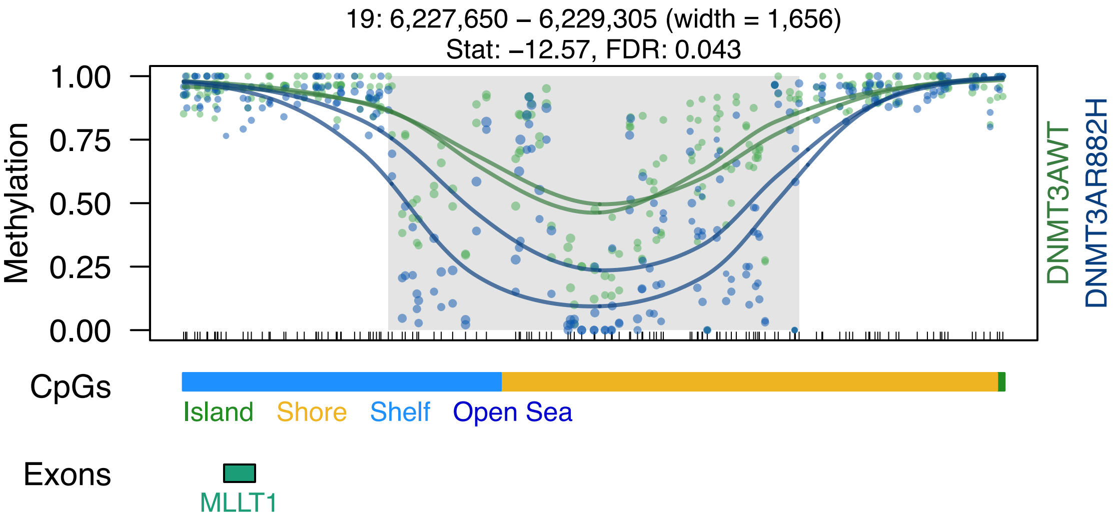
19: 6,227,650 – 6,229,305 (width = 1,656)
Stat: –12.57, FDR: 0.043
Methylation
1.00
0.75
0.50
0.25
0.00
DNMT3AWT
DNMT3AR882H
CpGs
Island
Shore
Shelf
Open Sea
Exons
MLLT1

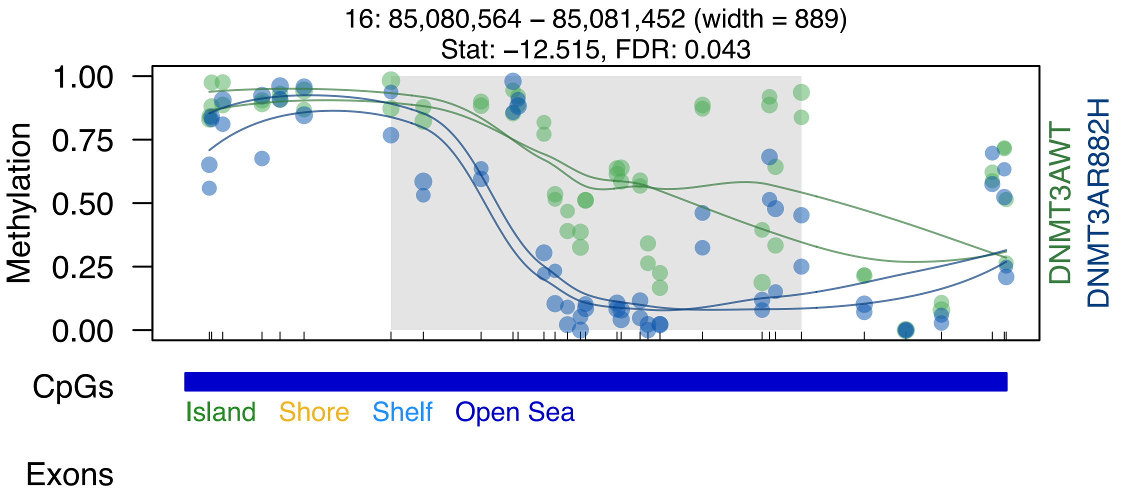
16: 85,080,564 – 85,081,452 (width = 889)
Stat: –12.515, FDR: 0.043
Methylation
1.00
0.75
0.50
0.25
0.00
DNMT3AWT
DNMT3AR882H
CpGs
Island
Shore
Shelf
Open Sea
Exons

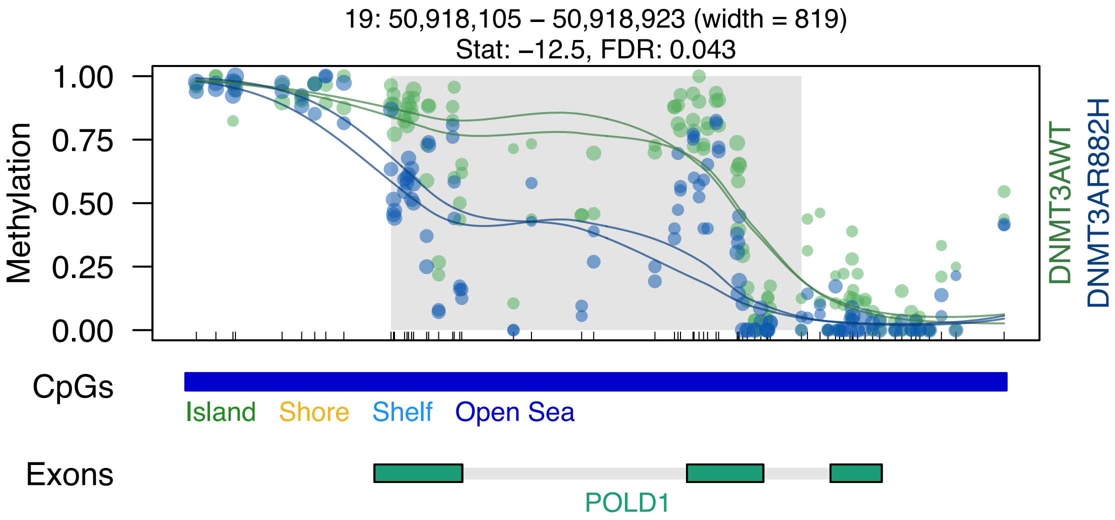

19: 50,918,105 – 50,918,923 (width = 819)
Stat: –12.5, FDR: 0.043
Methylation
1.00
0.75
0.50
0.25
0.00
DNMT3AWT
DNMT3AR882H
CpGs
Island
Shore
Shelf
Open Sea
Exons
POLD1

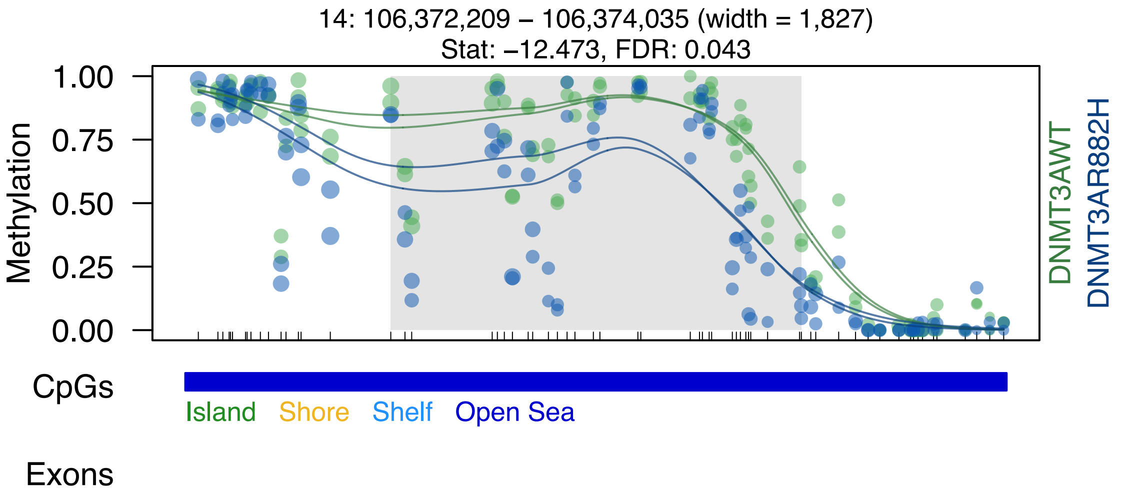
14: 106,372,209 – 106,374,035 (width = 1,827)
Stat: –12.473, FDR: 0.043
Methylation
1.00
0.75
0.50
0.25
0.00
DNMT3AWT
DNMT3AR882H
CpGs
Island
Shore
Shelf
Open Sea
Exons

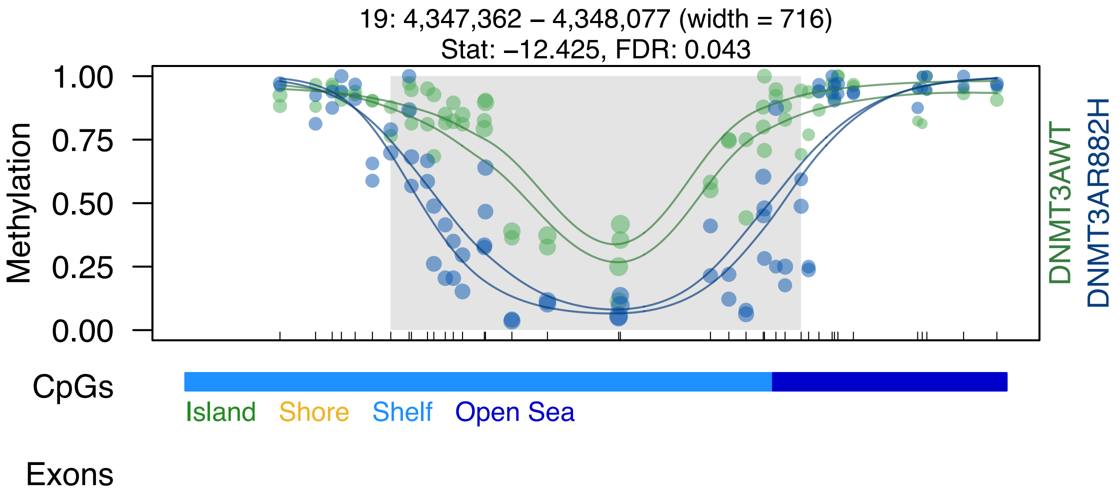
19: 4,347,362 – 4,348,077 (width = 716)
Stat: −12.425, FDR: 0.043
Methylation
1.00
0.75
0.50
0.25
0.00
DNMT3AWT
DNMT3AR882H
CpGs
Island
Shore
Shelf
Open Sea
Exons

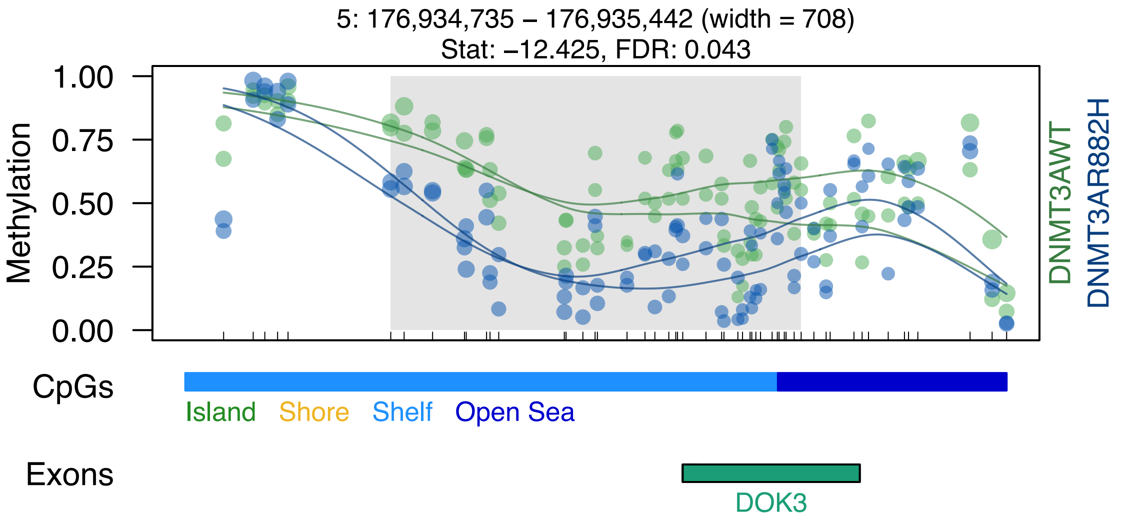
5: 176,934,735 – 176,935,442 (width = 708)
Stat: –12.425, FDR: 0.043
Methylation
1.00
0.75
0.50
0.25
0.00
DNMT3AWT
DNMT3AR882H
CpGs
Island
Shore
Shelf
Open Sea
Exons
DOK3

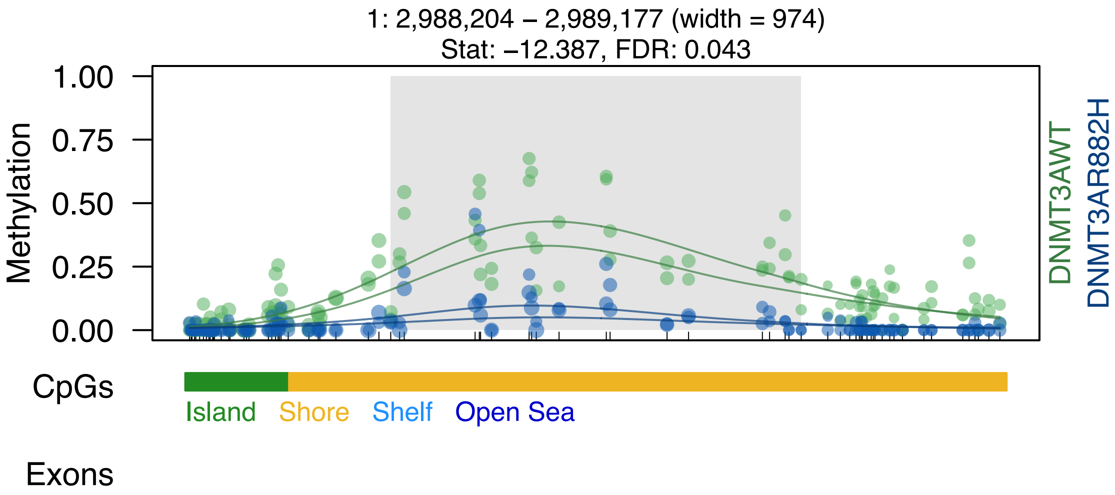
1: 2,988,204 – 2,989,177 (width = 974)
Stat: –12.387, FDR: 0.043
Methylation
1.00
0.75
0.50
0.25
0.00
DNMT3AWT
DNMT3AR882H
CpGs
Island
Shore
Shelf
Open Sea
Exons

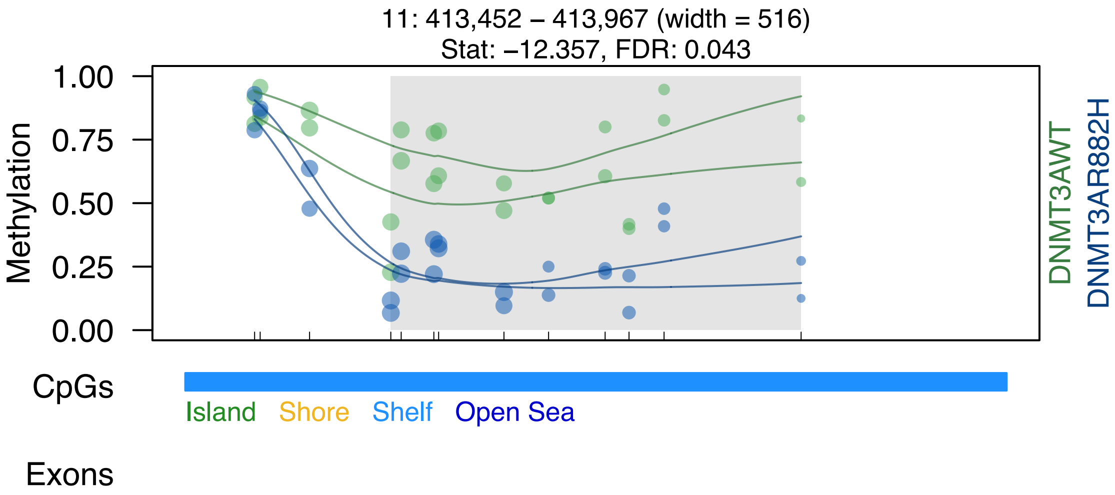

11: 413,452 – 413,967 (width = 516)
Stat: –12.357, FDR: 0.043
Methylation
1.00
0.75
0.50
0.25
0.00
DNMT3AWT
DNMT3AR882H
CpGs
Island
Shore
Shelf
Open Sea
Exons

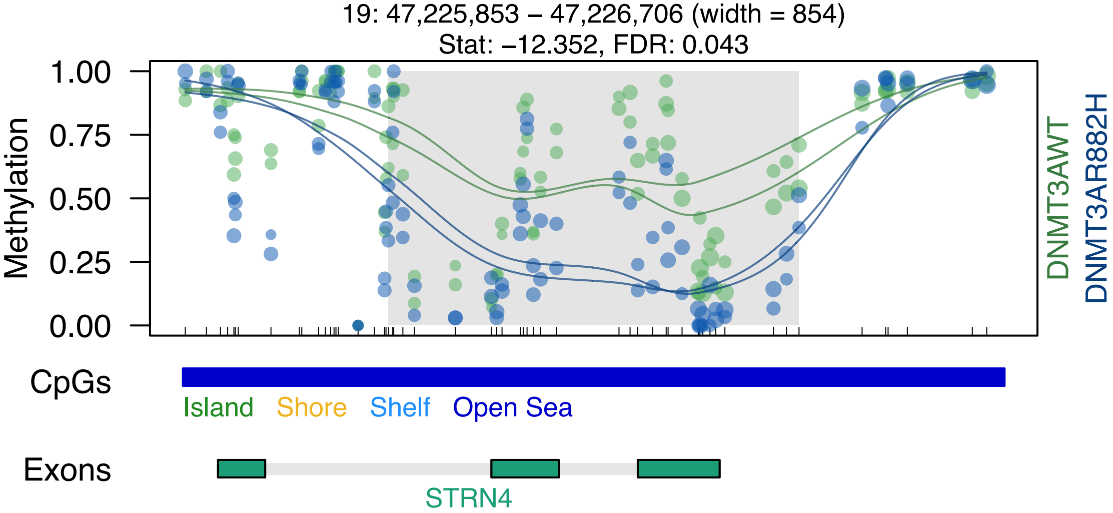
19: 47,225,853 – 47,226,706 (width = 854)
Stat: –12.352, FDR: 0.043
Methylation
1.00
0.75
0.50
0.25
0.00
DNMT3AWT
DNMT3AR882H
CpGs
Island
Shore
Shelf
Open Sea
Exons
STRN4

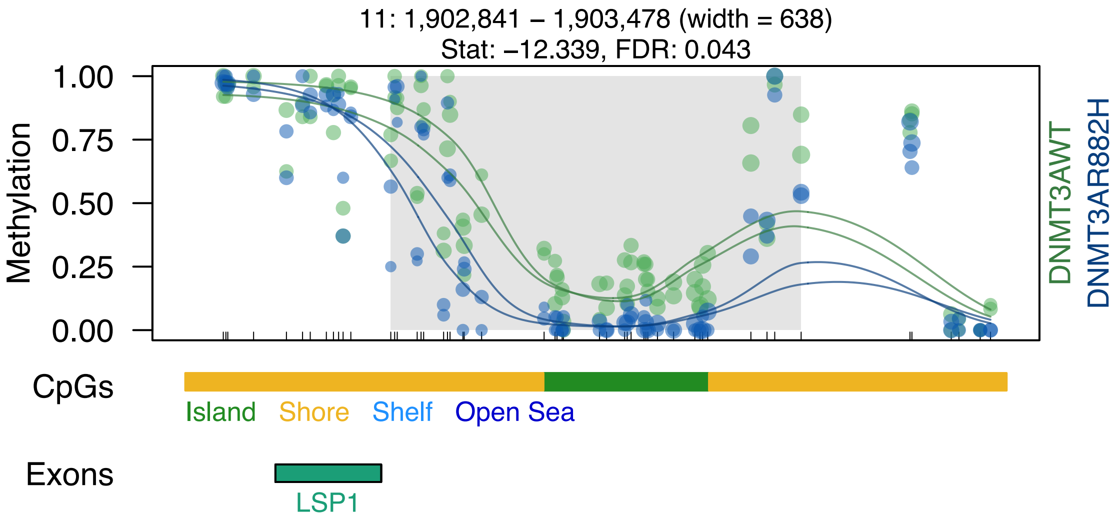

11: 1,902,841 – 1,903,478 (width = 638)
Stat: −12.339, FDR: 0.043
Methylation
1.00
0.75
0.50
0.25
0.00
DNMT3AWT
DNMT3AR882H
CpGs
Island
Shore
Shelf
Open Sea
Exons
LSP1

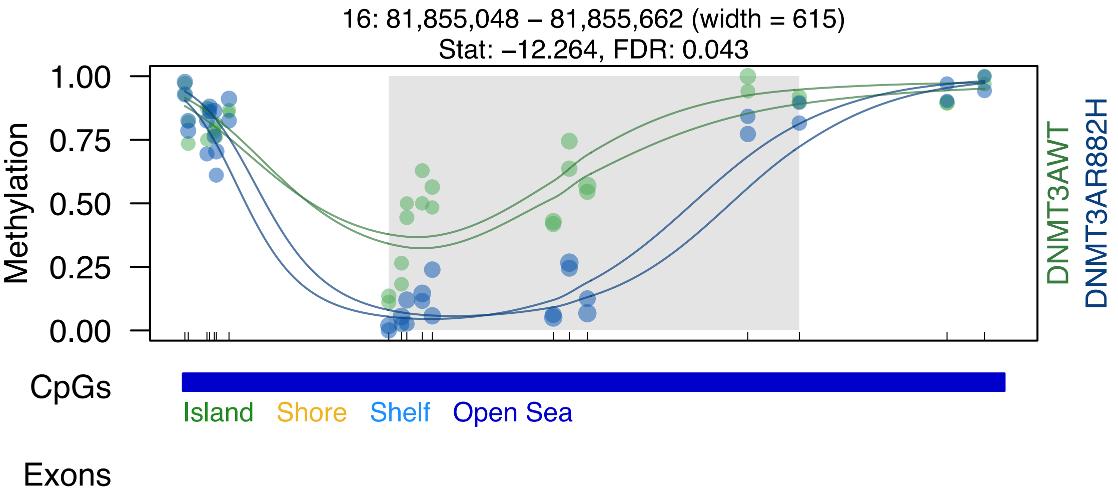
16: 81,855,048 – 81,855,662 (width = 615)
Stat: –12.264, FDR: 0.043
Methylation
1.00
0.75
0.50
0.25
0.00
DNMT3AWT
DNMT3AR882H
CpGs
Island
Shore
Shelf
Open Sea
Exons

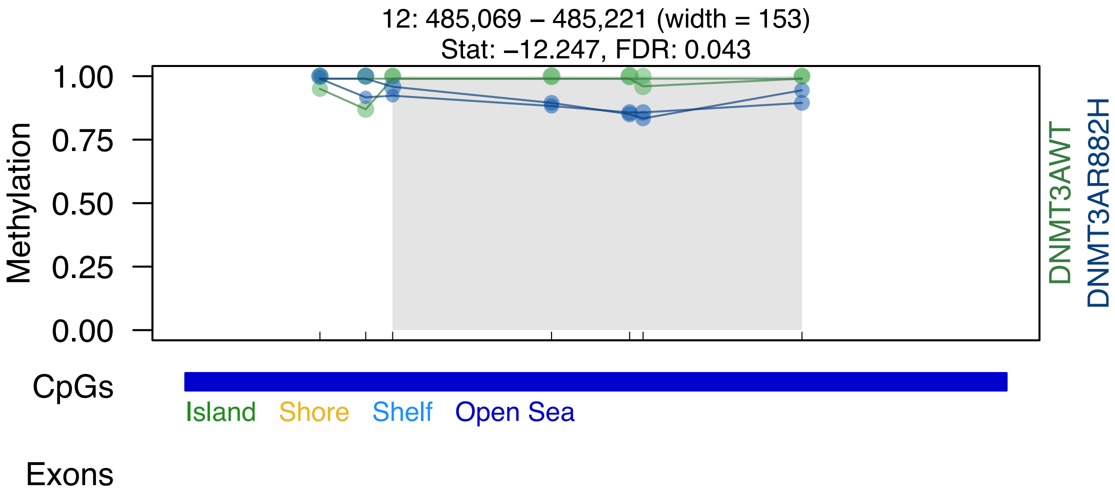

12: 485,069 – 485,221 (width = 153)
Stat: −12.247, FDR: 0.043
Methylation
1.00
0.75
0.50
0.25
0.00
DNMT3AWT
DNMT3AR882H
CpGs
Island
Shore
Shelf
Open Sea
Exons

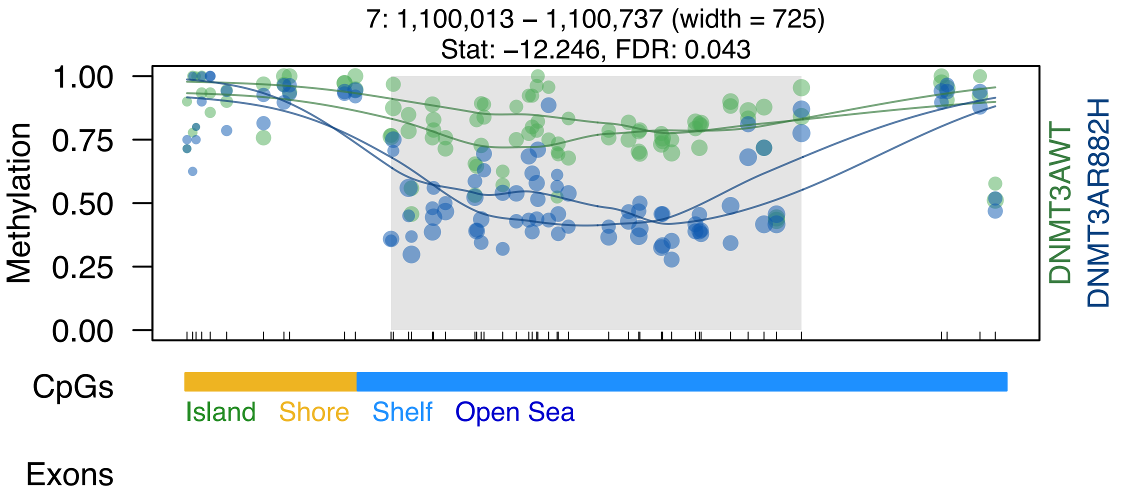

7: 1,100,013 – 1,100,737 (width = 725)
Stat: –12.246, FDR: 0.043
Methylation
1.00
0.75
0.50
0.25
0.00
DNMT3AWT
DNMT3AR882H
CpGs
Island
Shore
Shelf
Open Sea
Exons

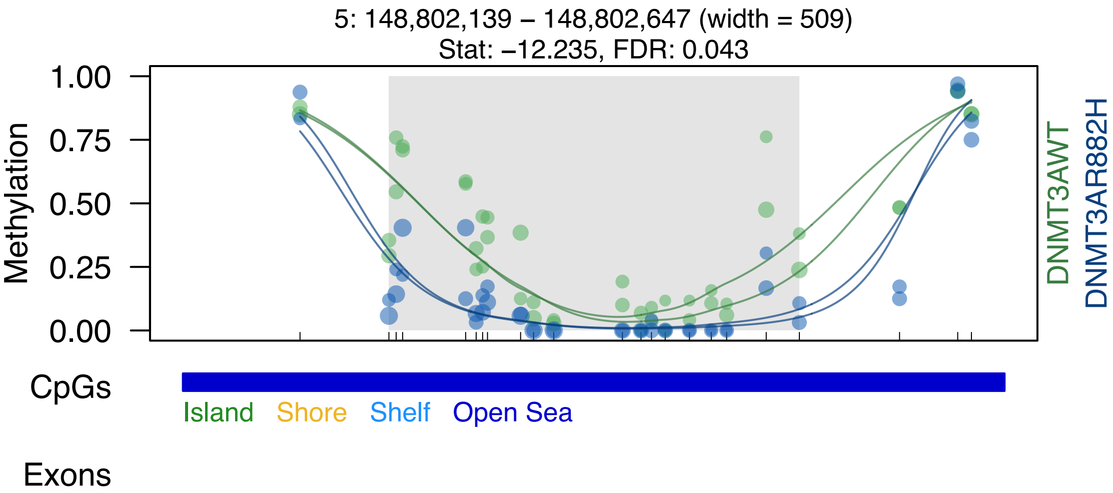

5: 148,802,139 – 148,802,647 (width = 509)
Stat: –12.235, FDR: 0.043
Methylation
1.00
0.75
0.50
0.25
0.00
DNMT3AWT
DNMT3AR882H
CpGs
Island
Shore
Shelf
Open Sea
Exons

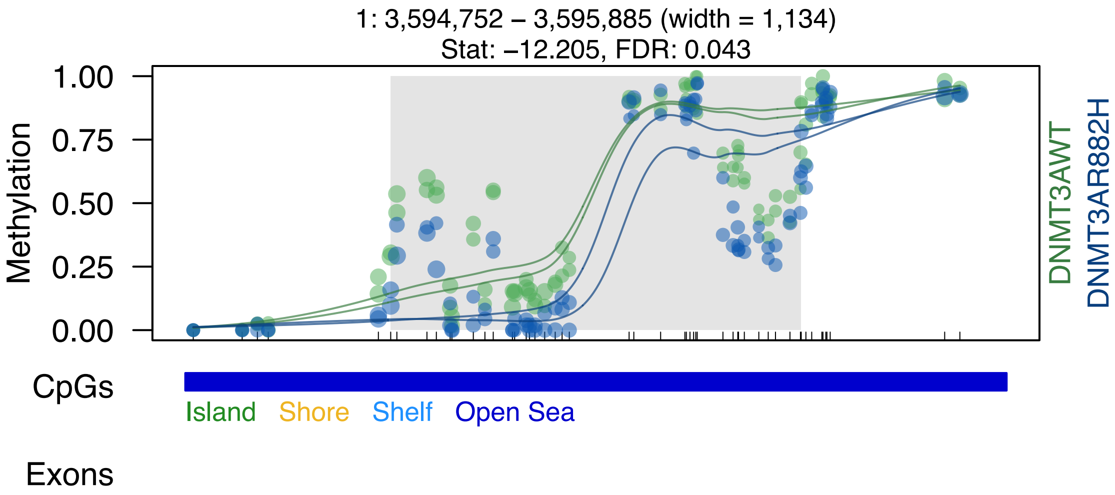

1: 3,594,752 – 3,595,885 (width = 1,134)
Stat: –12.205, FDR: 0.043
Methylation
1.00
0.75
0.50
0.25
0.00
DNMT3AWT
DNMT3AR882H
CpGs
Island
Shore
Shelf
Open Sea
Exons

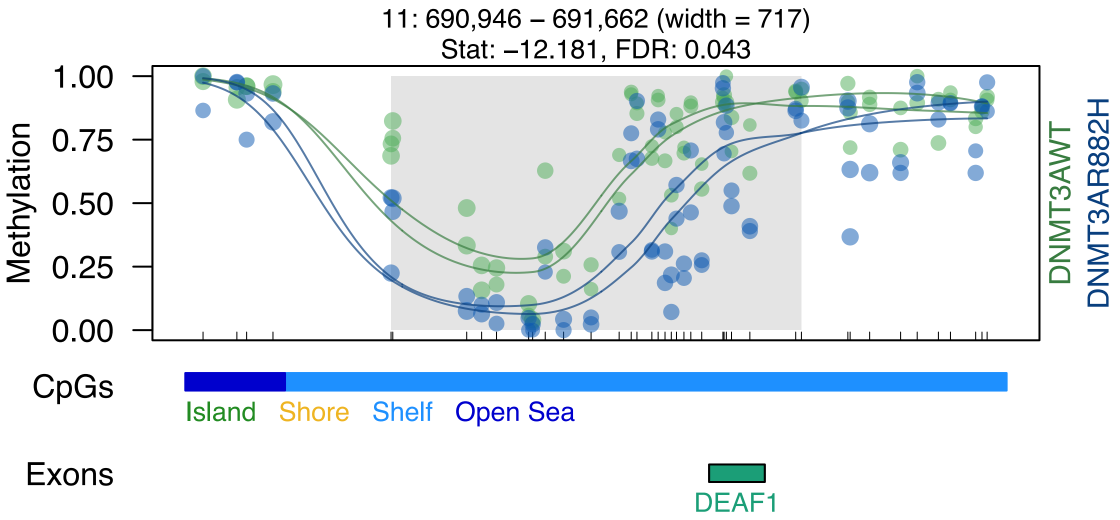
11: 690,946 – 691,662 (width = 717)
Stat: –12.181, FDR: 0.043
Methylation
1.00
0.75
0.50
0.25
0.00
DNMT3AWT
DNMT3AR882H
CpGs
Island
Shore
Shelf
Open Sea
Exons
DEAF1

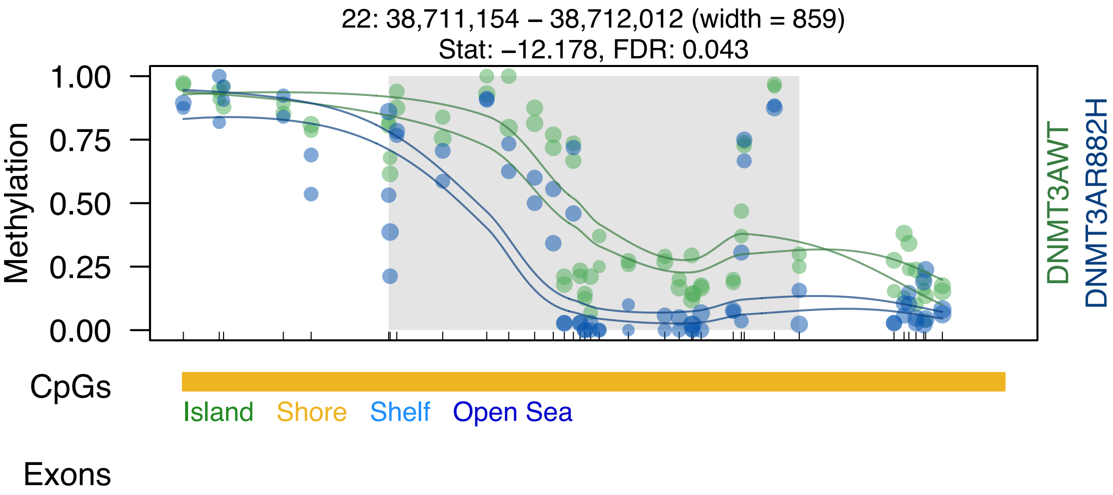

22: 38,711,154 – 38,712,012 (width = 859)
Stat: –12.178, FDR: 0.043
1.00
0.75
0.50
0.25
0.00
Methylation
DNMT3AWT
DNMT3AR882H
CpGs
Island
Shore
Shelf
Open Sea
Exons

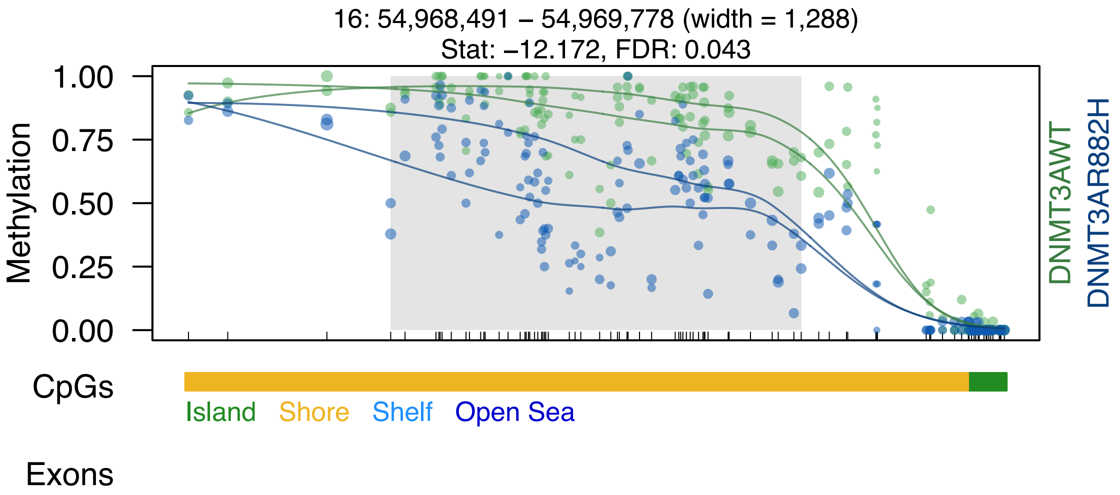

16: 54,968,491 – 54,969,778 (width = 1,288)
Stat: –12.172, FDR: 0.043
Methylation
1.00
0.75
0.50
0.25
0.00
DNMT3AWT
DNMT3AR882H
CpGs
Island
Shore
Shelf
Open Sea
Exons

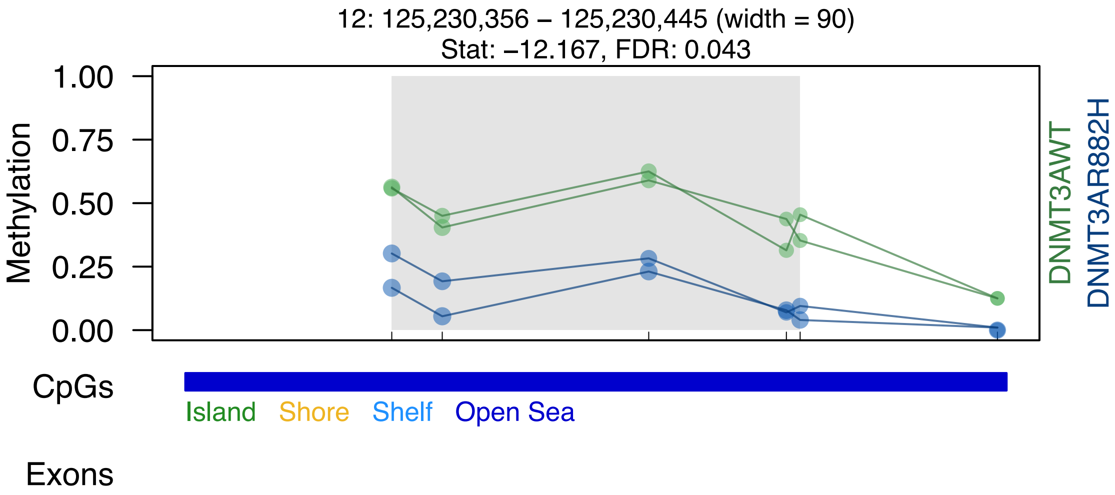

12: 125,230,356 – 125,230,445 (width = 90)
Stat: –12.167, FDR: 0.043
Methylation
1.00
0.75
0.50
0.25
0.00
DNMT3AWT
DNMT3AR882H
CpGs
Island
Shore
Shelf
Open Sea
Exons

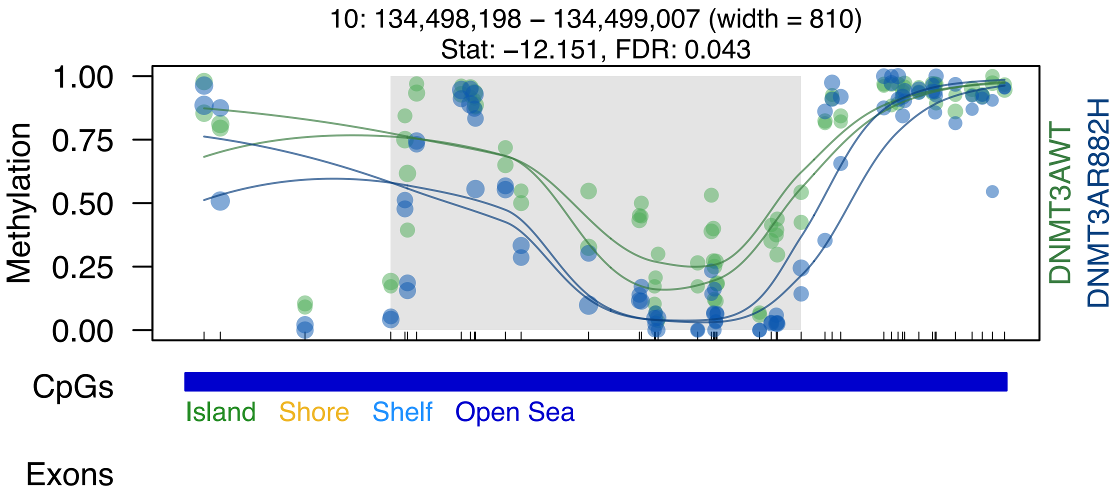
10: 134,498,198 – 134,499,007 (width = 810)
Stat: –12.151, FDR: 0.043
Methylation
1.00
0.75
0.50
0.25
0.00
DNMT3AWT
DNMT3AR882H
CpGs
Island
Shore
Shelf
Open Sea
Exons

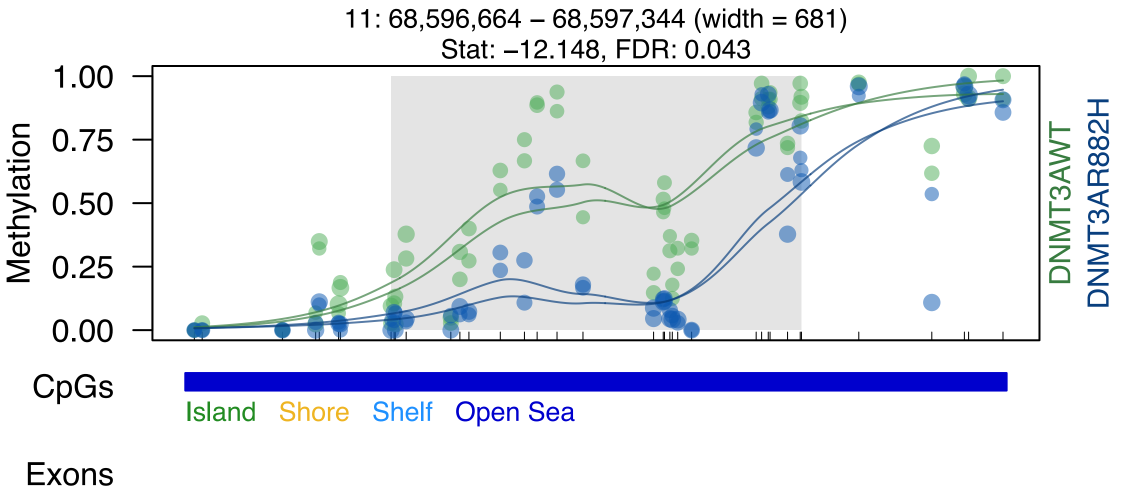

11: 68,596,664 – 68,597,344 (width = 681)
Stat: –12.148, FDR: 0.043
Methylation
1.00
0.75
0.50
0.25
0.00
DNMT3AWT
DNMT3AR882H
CpGs
Island
Shore
Shelf
Open Sea
Exons

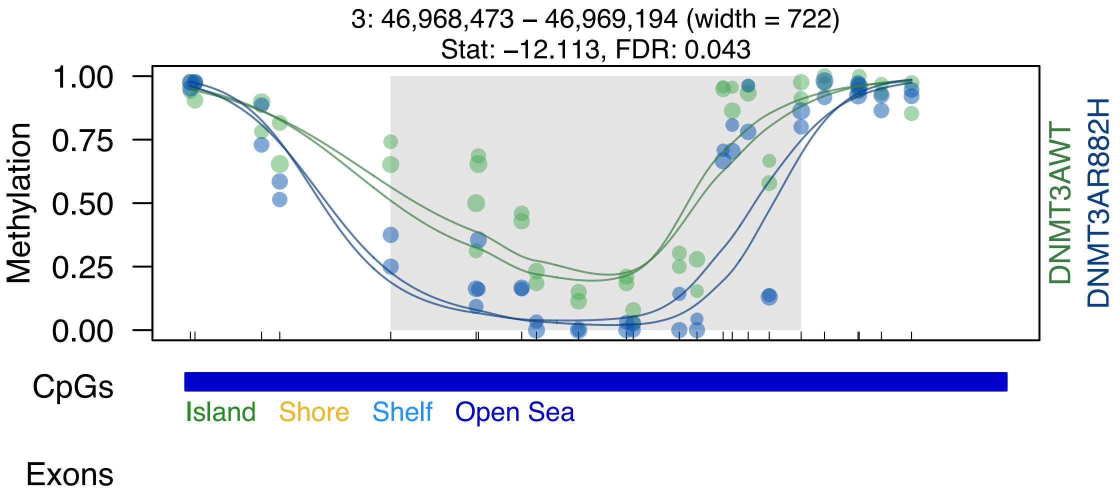
3: 46,968,473 – 46,969,194 (width = 722)
Stat: –12.113, FDR: 0.043
Methylation
1.00
0.75
0.50
0.25
0.00
DNMT3AWT
DNMT3AR882H
CpGs
Island
Shore
Shelf
Open Sea
Exons

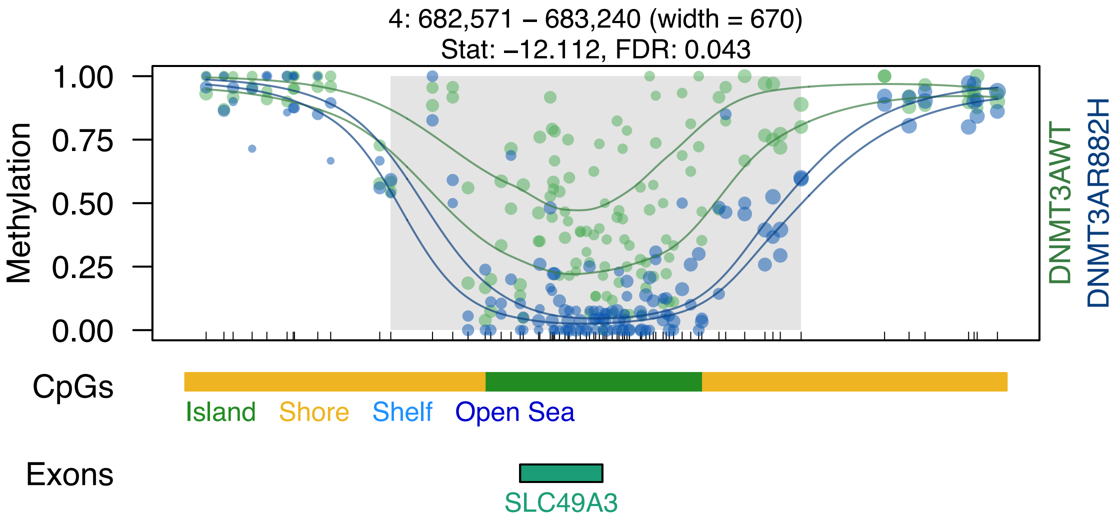

4: 682,571 – 683,240 (width = 670)
Stat: –12.112, FDR: 0.043
1.00
0.75
0.50
0.25
0.00
Methylation
DNMT3AWT
DNMT3AR882H
CpGs
Island
Shore
Shelf
Open Sea
Exons
SLC49A3

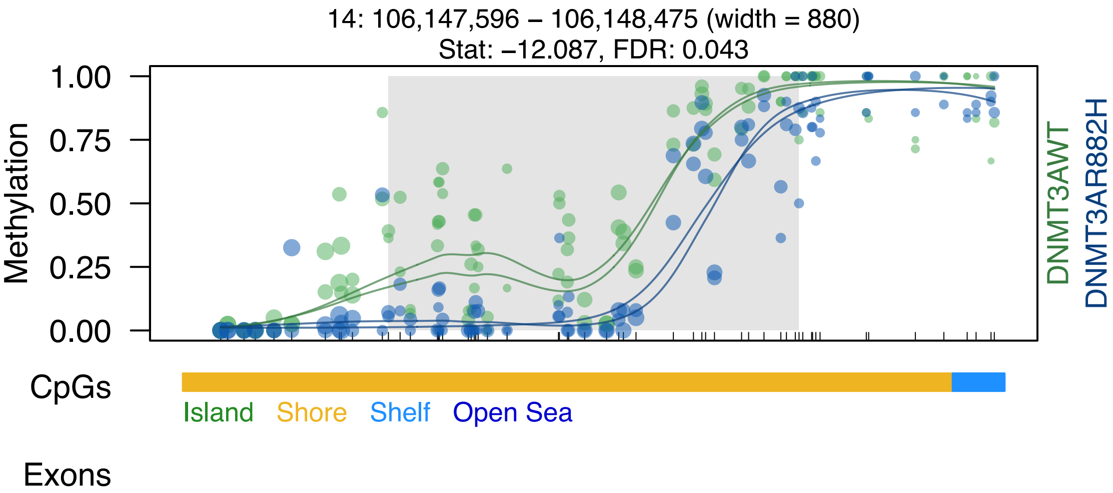

14: 106,147,596 – 106,148,475 (width = 880)
Stat: –12.087, FDR: 0.043
Methylation
1.00
0.75
0.50
0.25
0.00
DNMT3AWT
DNMT3AR882H
CpGs
Island
Shore
Shelf
Open Sea
Exons

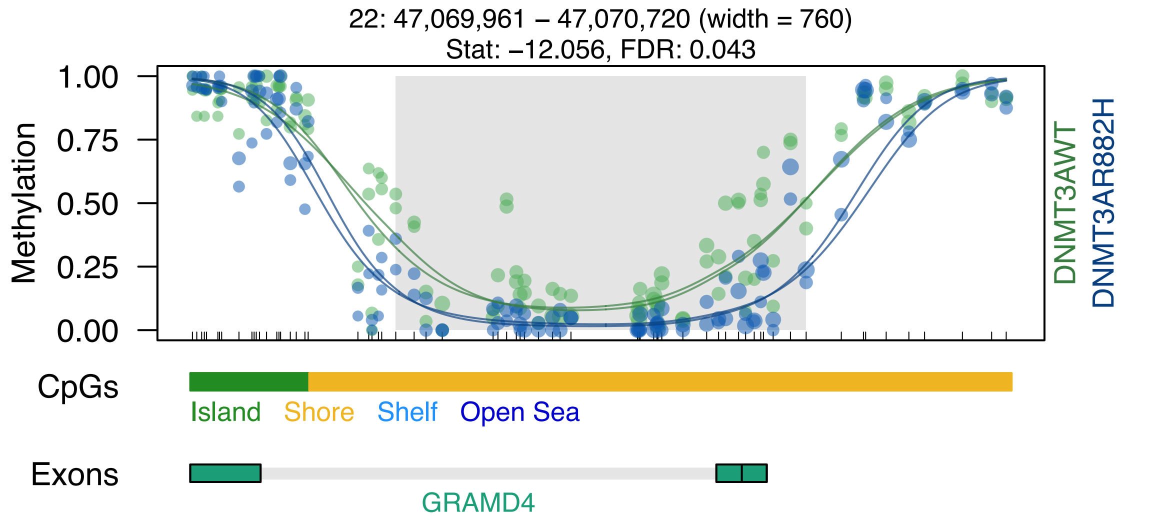

22: 47,069,961 – 47,070,720 (width = 760)
Stat: –12.056, FDR: 0.043
1.00
0.75
0.50
0.25
0.00
Methylation
DNMT3AWT
DNMT3AR882H
CpGs
Island
Shore
Shelf
Open Sea
Exons
GRAMD4

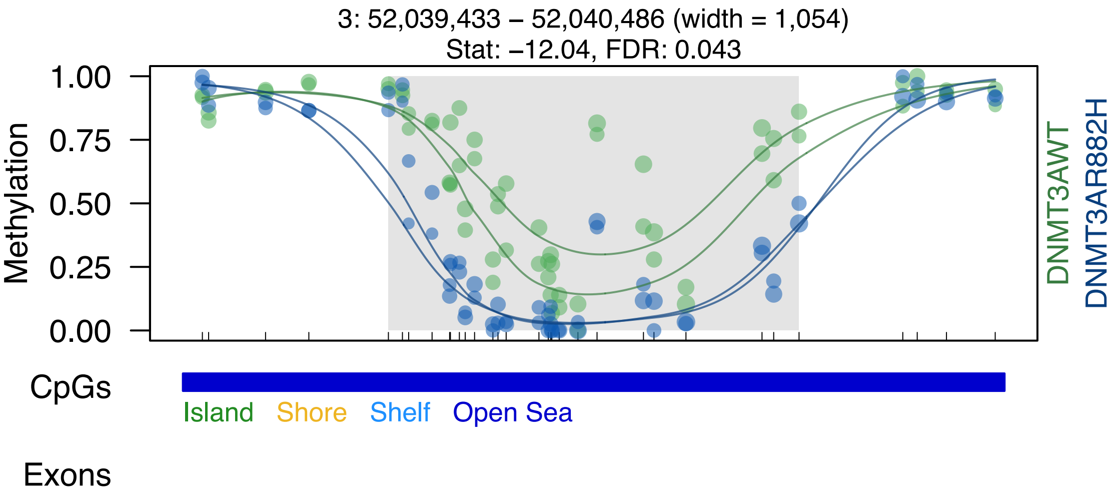

3: 52,039,433 – 52,040,486 (width = 1,054)
Stat: −12.04, FDR: 0.043
Methylation
1.00
0.75
0.50
0.25
0.00
DNMT3AWT
DNMT3AR882H
CpGs
Island
Shore
Shelf
Open Sea
Exons

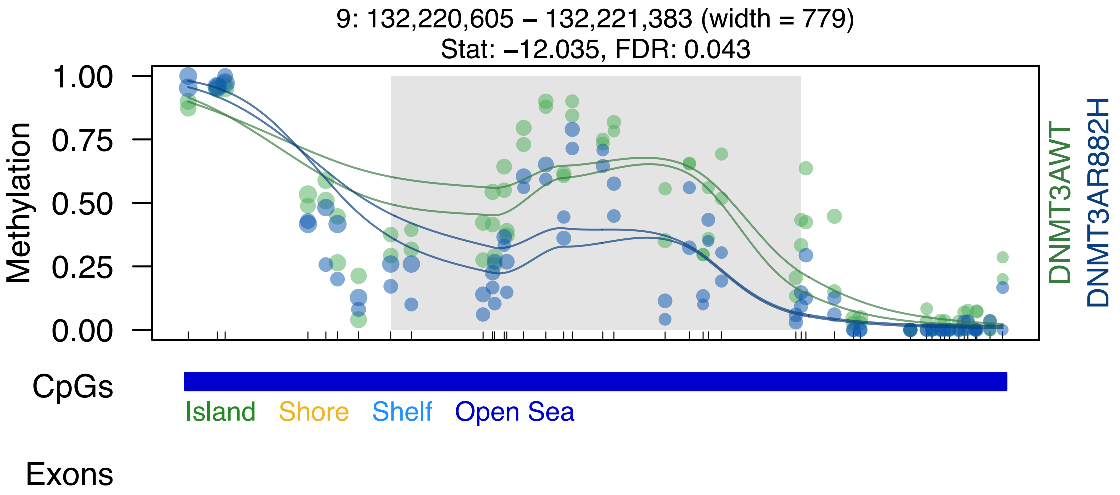

9: 132,220,605 – 132,221,383 (width = 779)
Stat: –12.035, FDR: 0.043
Methylation
1.00
0.75
0.50
0.25
0.00
DNMT3AWT
DNMT3AR882H
CpGs
Island
Shore
Shelf
Open Sea
Exons

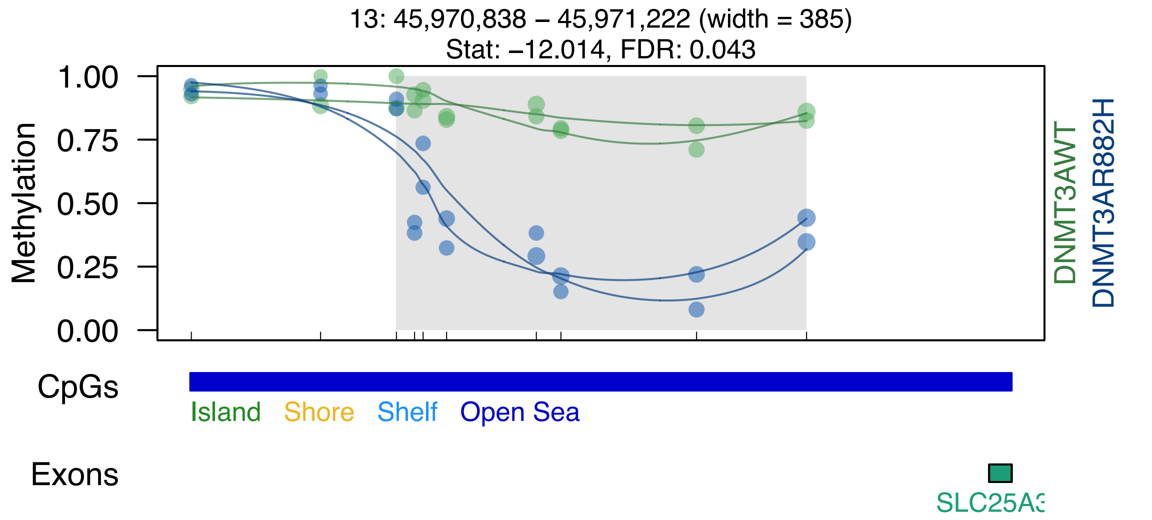

13: 45,970,838 – 45,971,222 (width = 385)
Stat: –12.014, FDR: 0.043
1.00
0.75
0.50
0.25
0.00
Methylation
DNMT3AWT
DNMT3AR882H
CpGs
Island
Shore
Shelf
Open Sea
Exons
SLC25A3

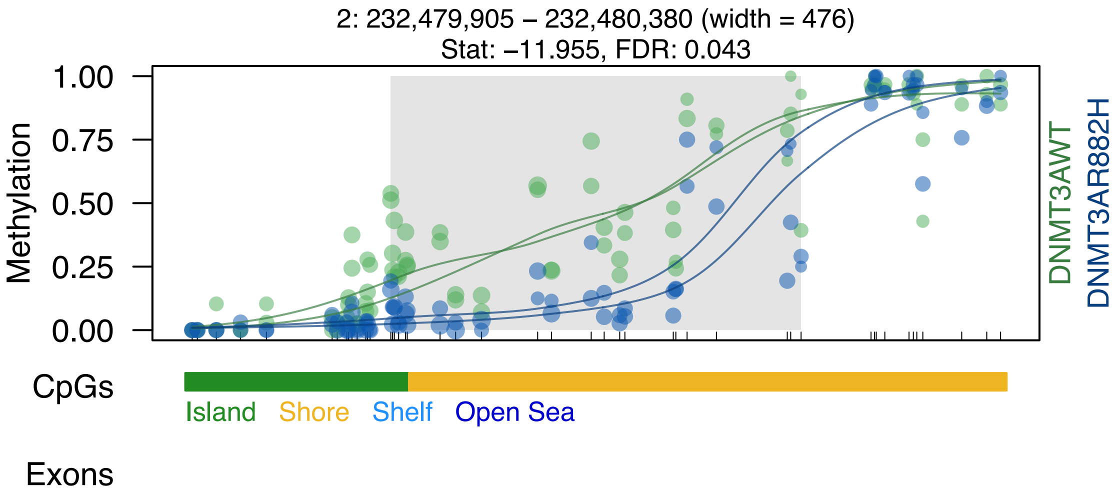
2: 232,479,905 – 232,480,380 (width = 476)
Stat: –11.955, FDR: 0.043
Methylation
1.00
0.75
0.50
0.25
0.00
DNMT3AWT
DNMT3AR882H
CpGs
Island
Shore
Shelf
Open Sea
Exons

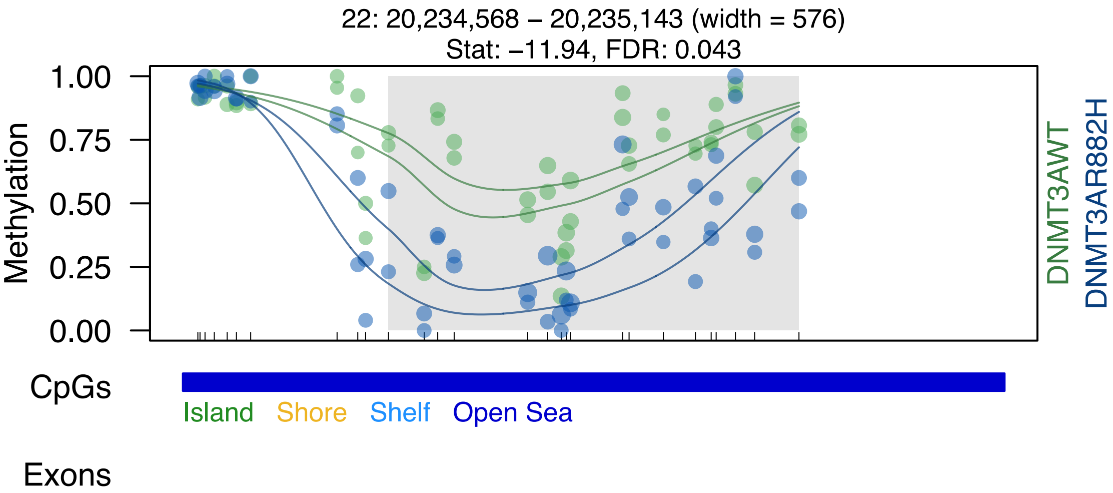

22: 20,234,568 – 20,235,143 (width = 576)
Stat: –11.94, FDR: 0.043
1.00
0.75
0.50
0.25
0.00
Methylation
DNMT3AWT
DNMT3AR882H
CpGs
Island
Shore
Shelf
Open Sea
Exons

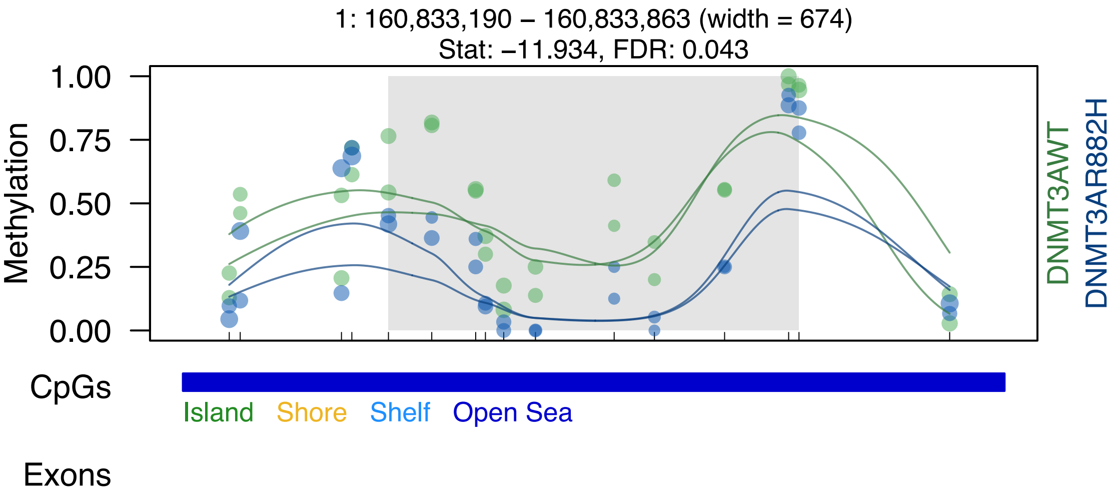

1: 160,833,190 – 160,833,863 (width = 674)
Stat: −11.934, FDR: 0.043
1.00
0.75
0.50
0.25
0.00
Methylation
DNMT3AWT
DNMT3AR882H
CpGs
Island
Shore
Shelf
Open Sea
Exons
