## Supplemental Data S1-S7 for "DNMT3A^R882H^ Is Not Required for Disease Maintenance in Primary Human AML, but Is Associated With Increased Leukemia Stem Cell Frequency": DataS2_dmrSeq_SU540_top100DMRs.pdf

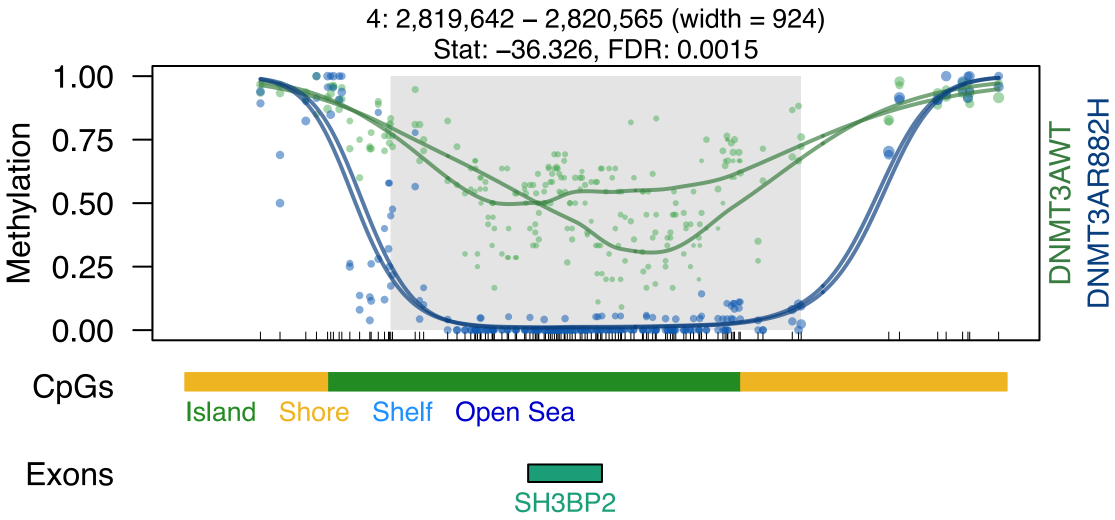

4: 2,819,642 – 2,820,565 (width = 924)
Stat: –36.326, FDR: 0.0015
1.00
0.75
0.50
0.25
0.00
Methylation
DNMT3AWT
DNMT3AR882H
CpGs
Island
Shore
Shelf
Open Sea
Exons
SH3BP2

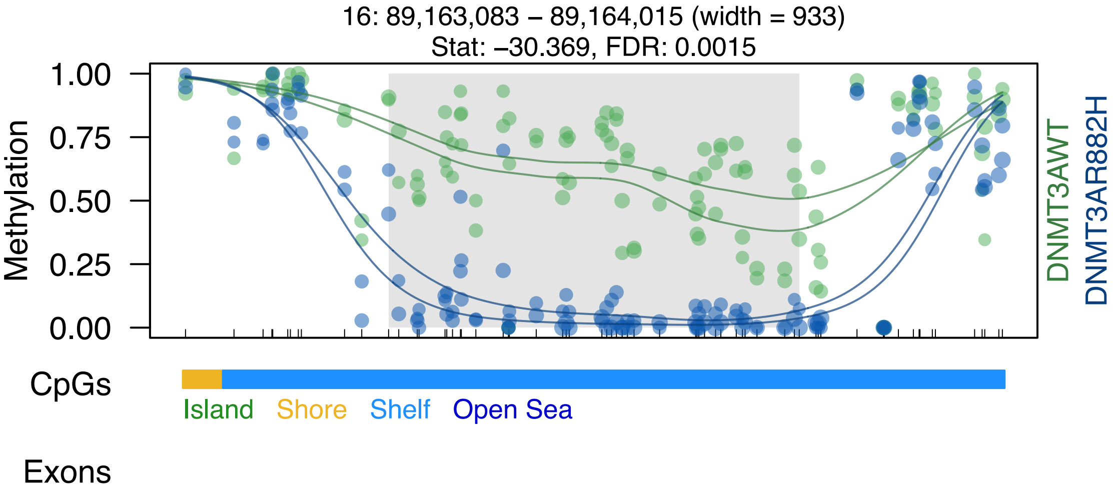
16: 89,163,083 – 89,164,015 (width = 933)
Stat: –30.369, FDR: 0.0015
1.00
0.75
0.50
0.25
0.00
Methylation
DNMT3AWT
DNMT3AR882H
CpGs
Island
Shore
Shelf
Open Sea
Exons

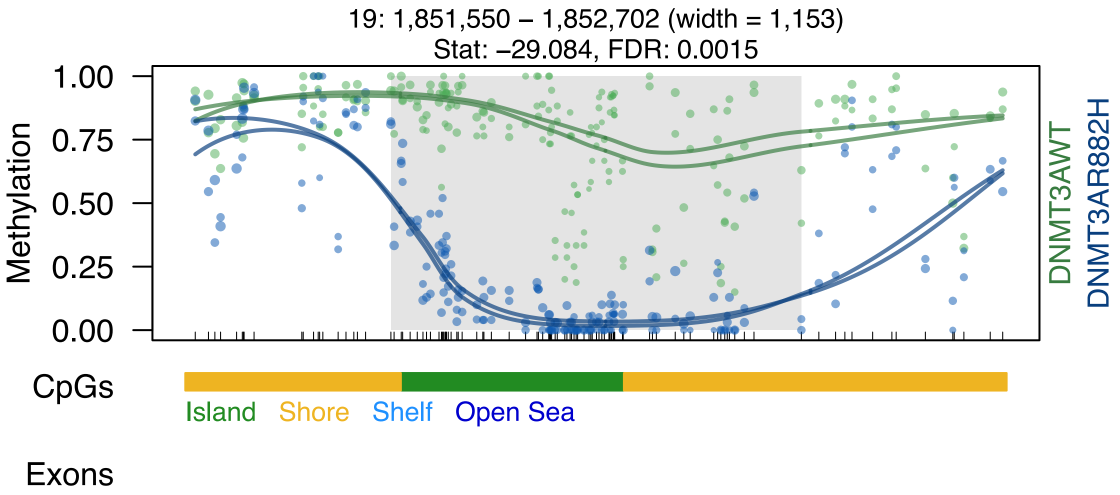

19: 1,851,550 – 1,852,702 (width = 1,153)
Stat: −29.084, FDR: 0.0015
1.00
0.75
0.50
0.25
0.00
Methylation
DNMT3AWT
DNMT3AR882H
CpGs
Island
Shore
Shelf
Open Sea
Exons

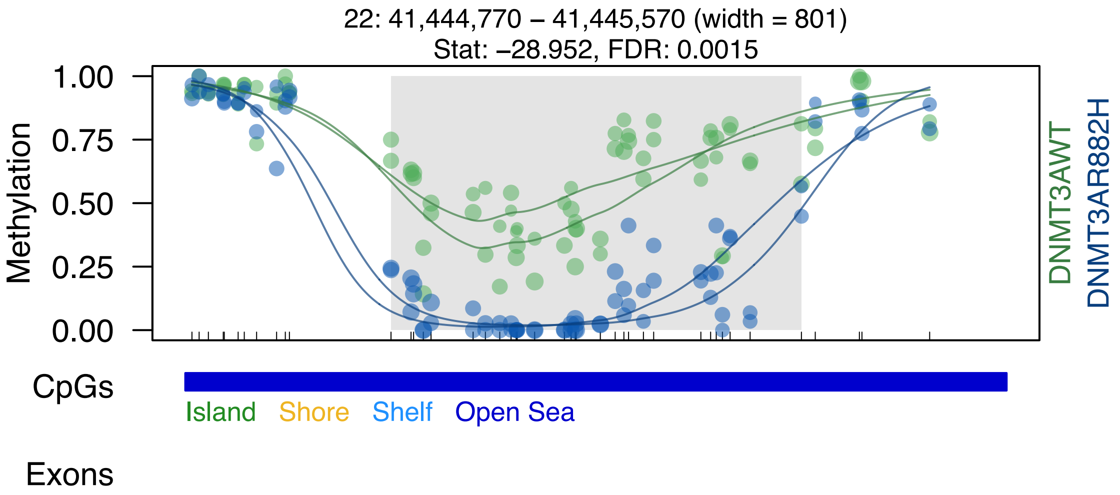
22: 41,444,770 – 41,445,570 (width = 801)
Stat: –28.952, FDR: 0.0015
Methylation
1.00
0.75
0.50
0.25
0.00
DNMT3AWT
DNMT3AR882H
CpGs
Island
Shore
Shelf
Open Sea
Exons

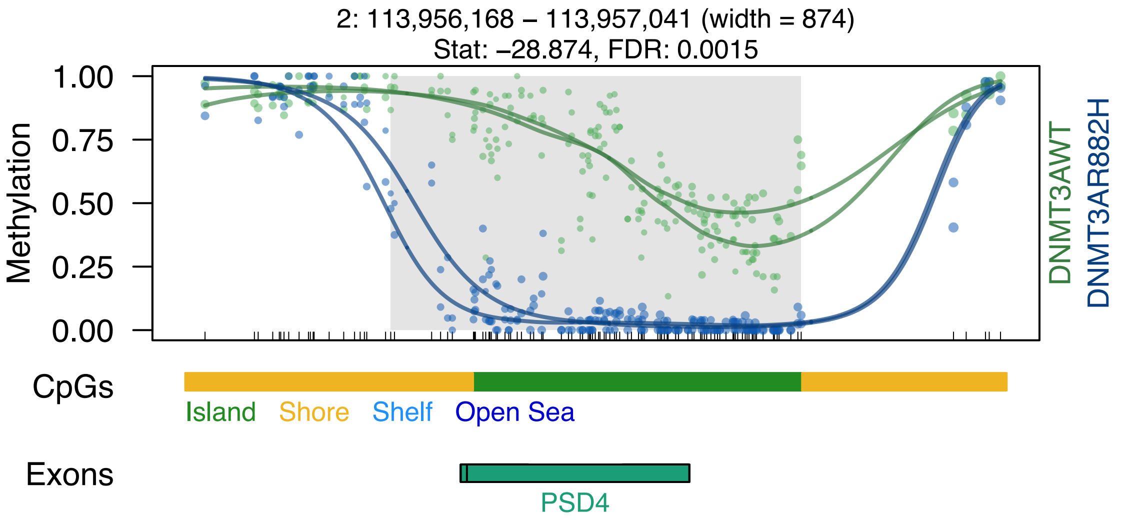
2: 113,956,168 – 113,957,041 (width = 874)
Stat: –28.874, FDR: 0.0015
Methylation
1.00
0.75
0.50
0.25
0.00
DNMT3AWT
DNMT3AR882H
CpGs
Island
Shore
Shelf
Open Sea
Exons
PSD4

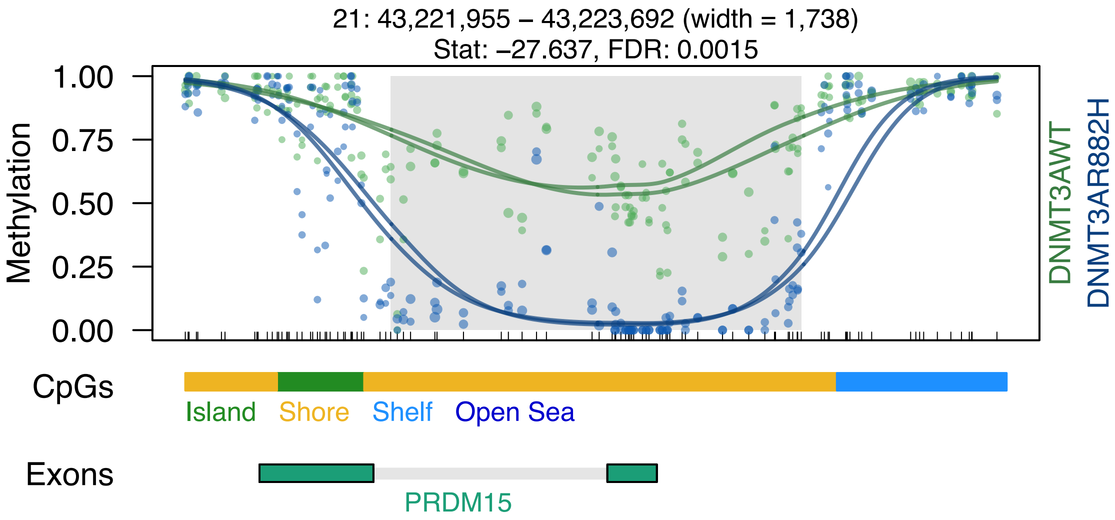

21: 43,221,955 – 43,223,692 (width = 1,738)
Stat: –27.637, FDR: 0.0015
Methylation
1.00
0.75
0.50
0.25
0.00
DNMT3AWT
DNMT3AR882H
CpGs
Island
Shore
Shelf
Open Sea
Exons
PRDM15

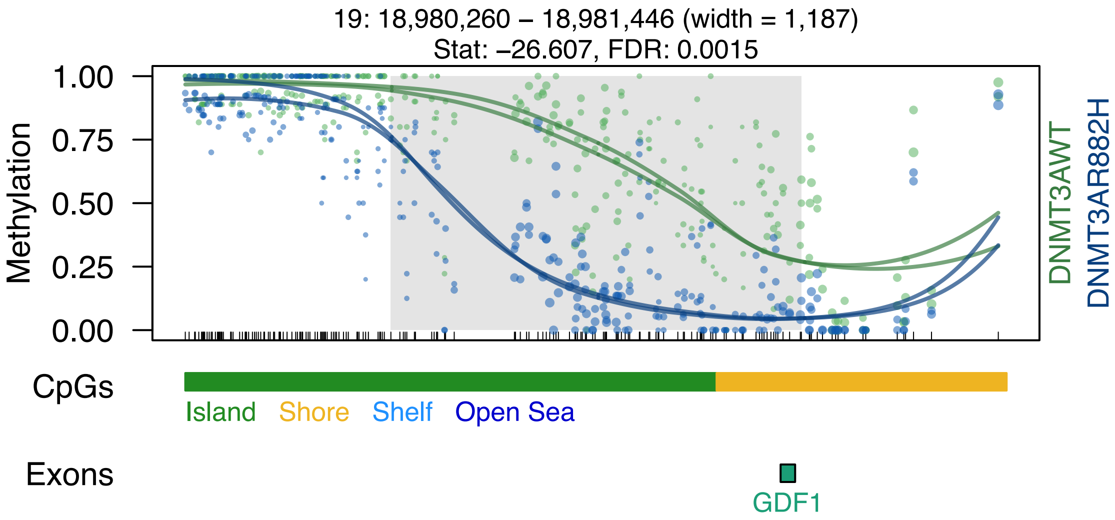

19: 18,980,260 – 18,981,446 (width = 1,187)
Stat: –26.607, FDR: 0.0015
1.00
0.75
0.50
0.25
0.00
Methylation
DNMT3AWT
DNMT3AR882H
CpGs
Island
Shore
Shelf
Open Sea
Exons
GDF1

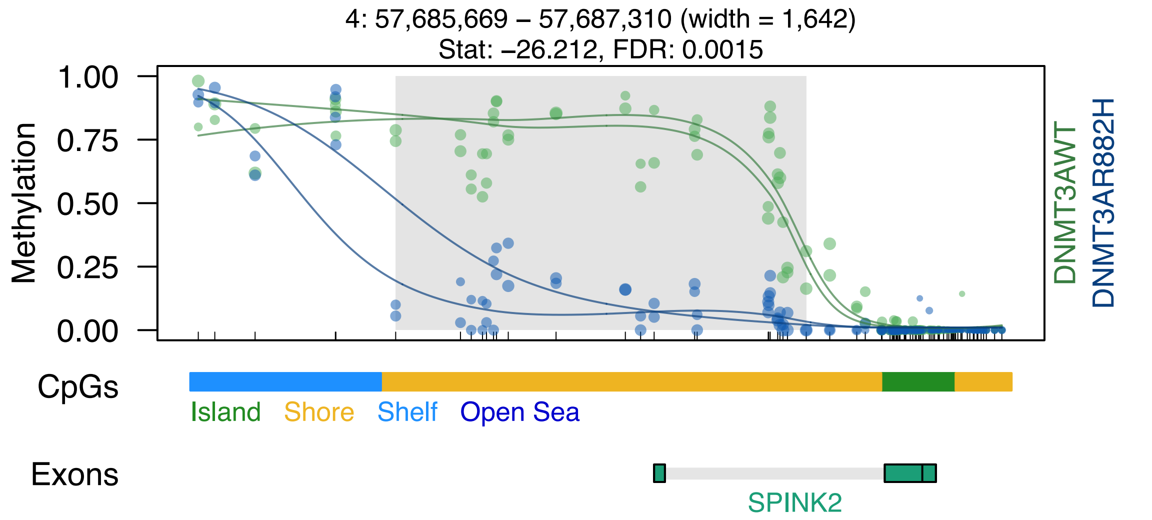
4: 57,685,669 – 57,687,310 (width = 1,642)
Stat: –26.212, FDR: 0.0015
1.00
0.75
0.50
0.25
0.00
Methylation
DNMT3AWT
DNMT3AR882H
CpGs
Island
Shore
Shelf
Open Sea
Exons
SPINK2

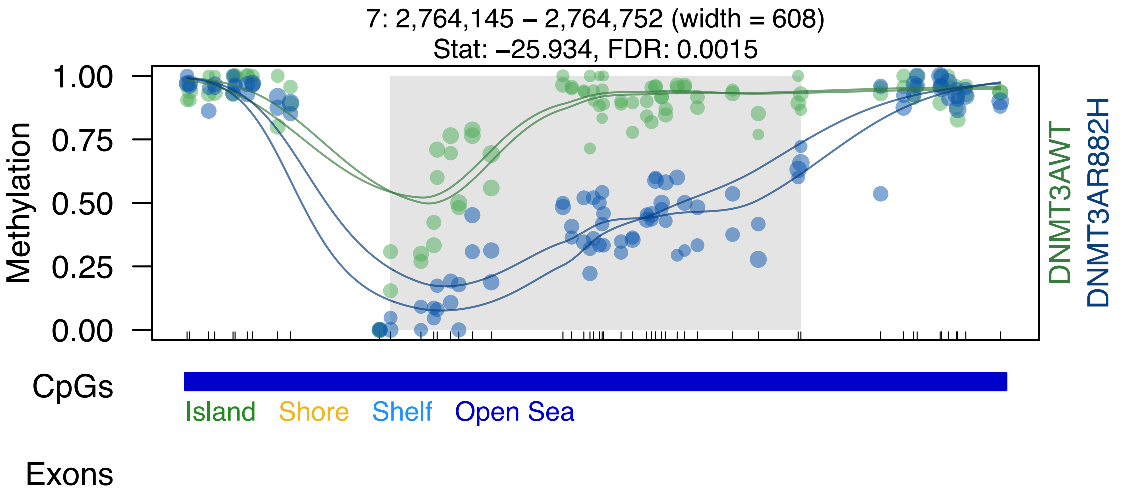
7: 2,764,145 – 2,764,752 (width = 608)
Stat: –25.934, FDR: 0.0015
Methylation
1.00
0.75
0.50
0.25
0.00
DNMT3AWT
DNMT3AR882H
CpGs
Island
Shore
Shelf
Open Sea
Exons

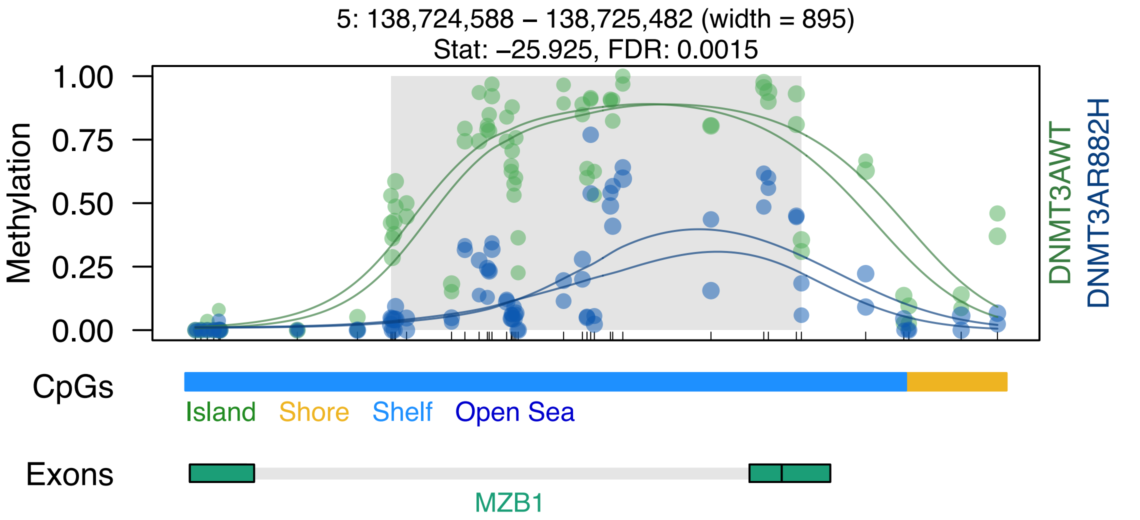

5: 138,724,588 – 138,725,482 (width = 895)
Stat: –25.925, FDR: 0.0015
Methylation
1.00
0.75
0.50
0.25
0.00
DNMT3AWT
DNMT3AR882H
CpGs
Island
Shore
Shelf
Open Sea
Exons
MZB1

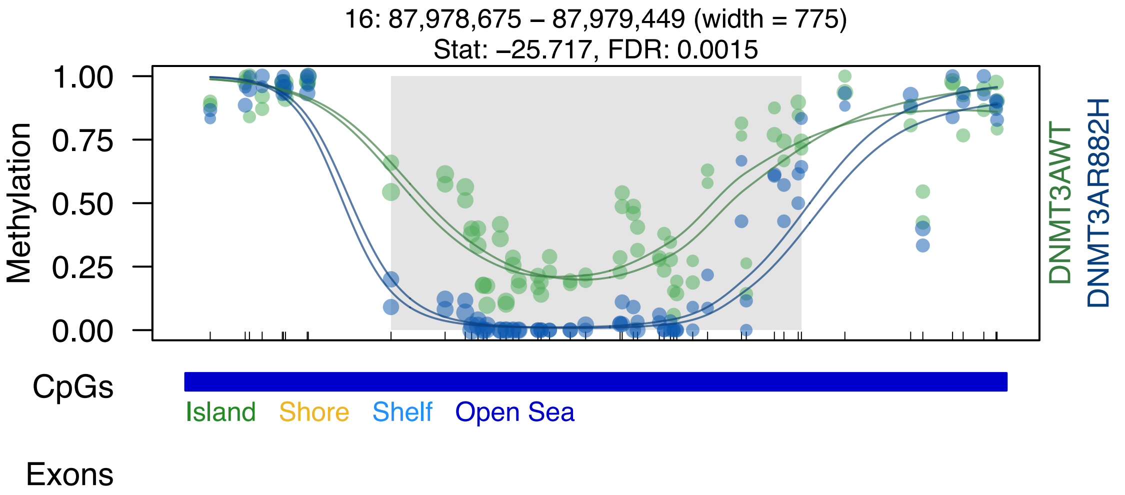
16: 87,978,675 – 87,979,449 (width = 775)
Stat: –25.717, FDR: 0.0015
Methylation
1.00
0.75
0.50
0.25
0.00
DNMT3AWT
DNMT3AR882H
CpGs
Island
Shore
Shelf
Open Sea
Exons

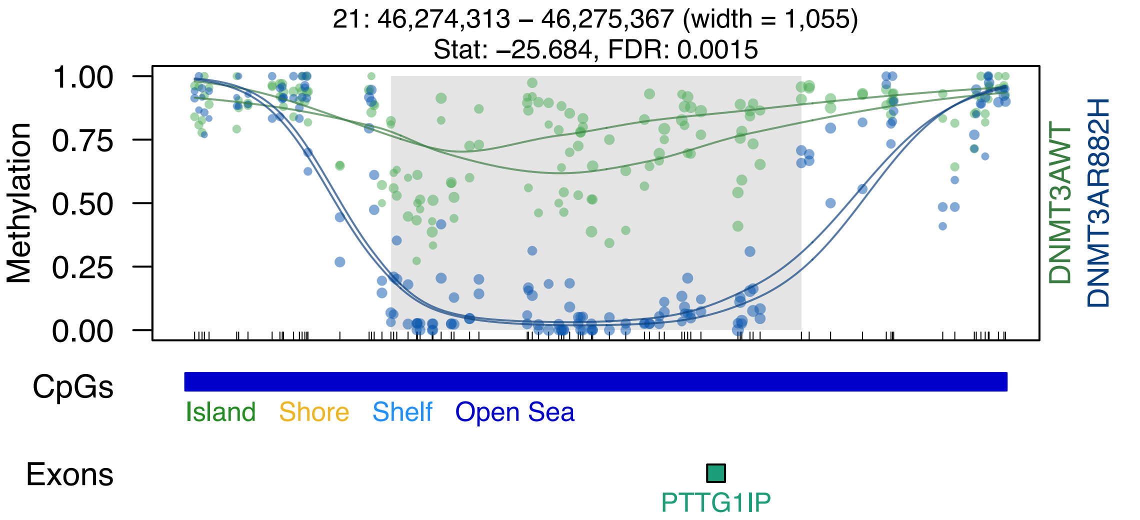

21: 46,274,313 – 46,275,367 (width = 1,055)
Stat: −25.684, FDR: 0.0015
Methylation
1.00
0.75
0.50
0.25
0.00
DNMT3AWT
DNMT3AR882H
CpGs
Island
Shore
Shelf
Open Sea
Exons
PTTG1IP

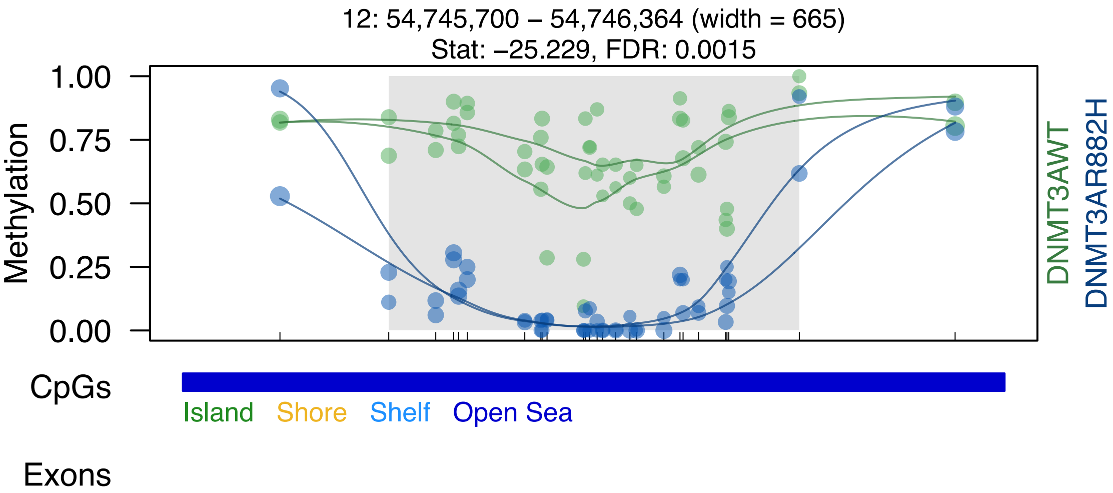
12: 54,745,700 – 54,746,364 (width = 665)
Stat: –25.229, FDR: 0.0015
Methylation
1.00
0.75
0.50
0.25
0.00
DNMT3AWT
DNMT3AR882H
CpGs
Island
Shore
Shelf
Open Sea
Exons

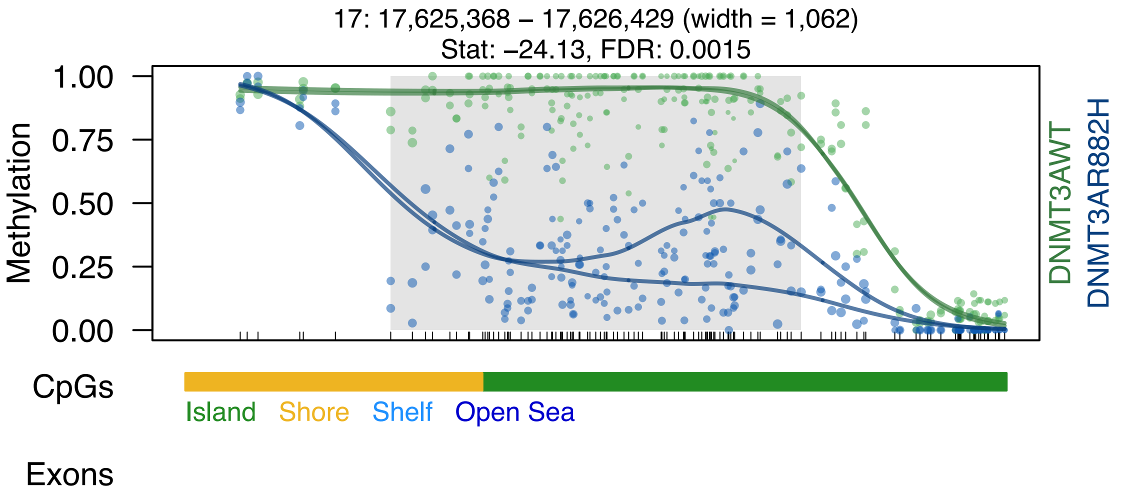

17: 17,625,368 – 17,626,429 (width = 1,062)
Stat: –24.13, FDR: 0.0015
1.00
0.75
0.50
0.25
0.00
Methylation
DNMT3AWT
DNMT3AR882H
CpGs
Island
Shore
Shelf
Open Sea
Exons

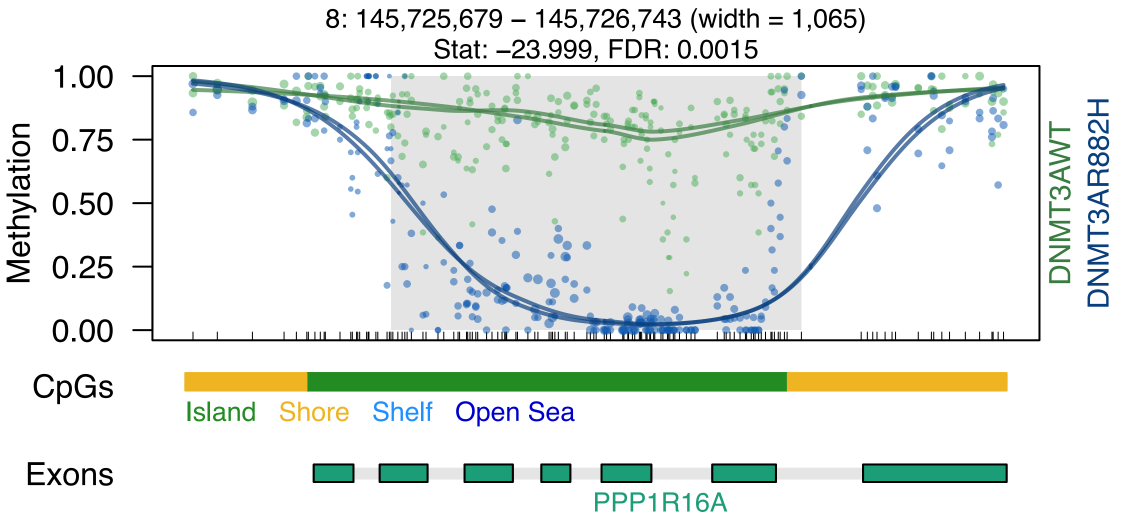

8: 145,725,679 – 145,726,743 (width = 1,065)
Stat: −23.999, FDR: 0.0015
Methylation
1.00
0.75
0.50
0.25
0.00
DNMT3AWT
DNMT3AR882H
CpGs
Island
Shore
Shelf
Open Sea
Exons
PPP1R16A

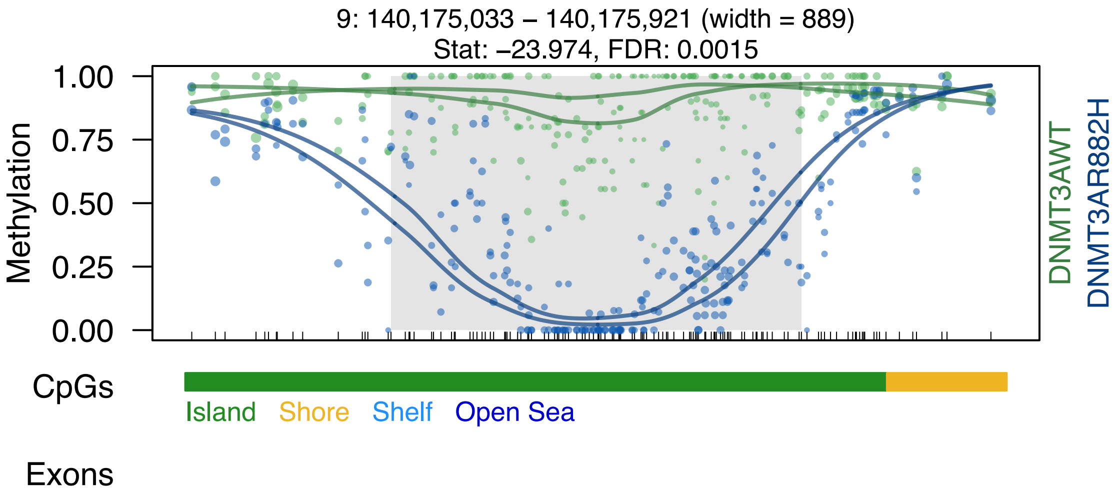

9: 140,175,033 – 140,175,921 (width = 889)
Stat: –23.974, FDR: 0.0015
1.00
0.75
0.50
0.25
0.00
Methylation
DNMT3AWT
DNMT3AR882H
CpGs
Island
Shore
Shelf
Open Sea
Exons

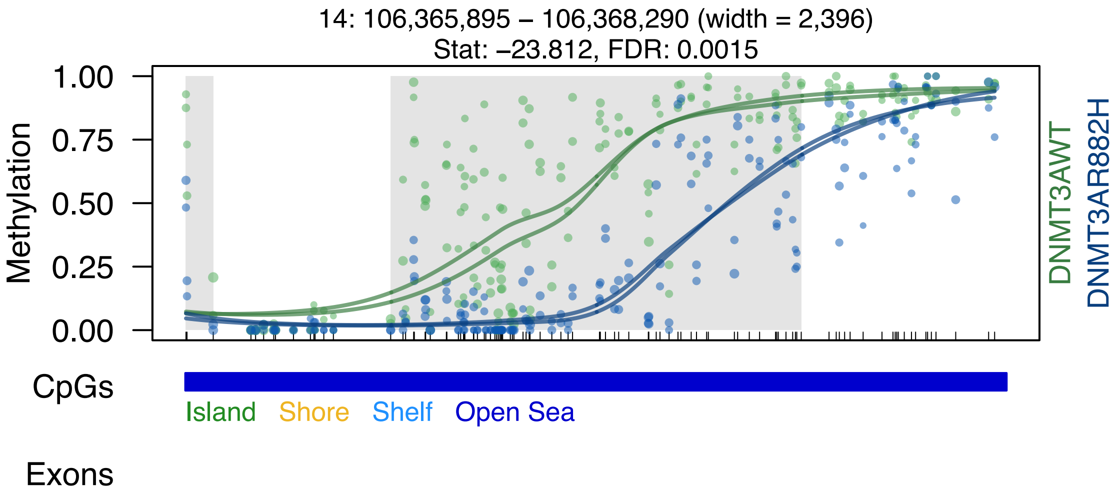

14: 106,365,895 – 106,368,290 (width = 2,396)
Stat: –23.812, FDR: 0.0015
Methylation
1.00
0.75
0.50
0.25
0.00
DNMT3AWT
DNMT3AR882H
CpGs
Island
Shore
Shelf
Open Sea
Exons

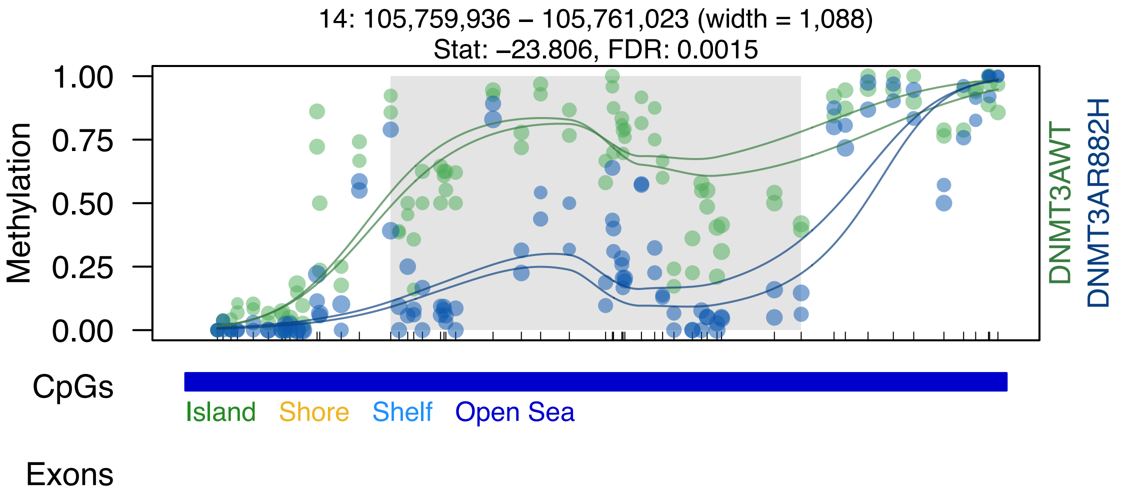
14: 105,759,936 – 105,761,023 (width = 1,088)
Stat: –23.806, FDR: 0.0015
Methylation
1.00
0.75
0.50
0.25
0.00
DNMT3AWT
DNMT3AR882H
CpGs
Island
Shore
Shelf
Open Sea
Exons

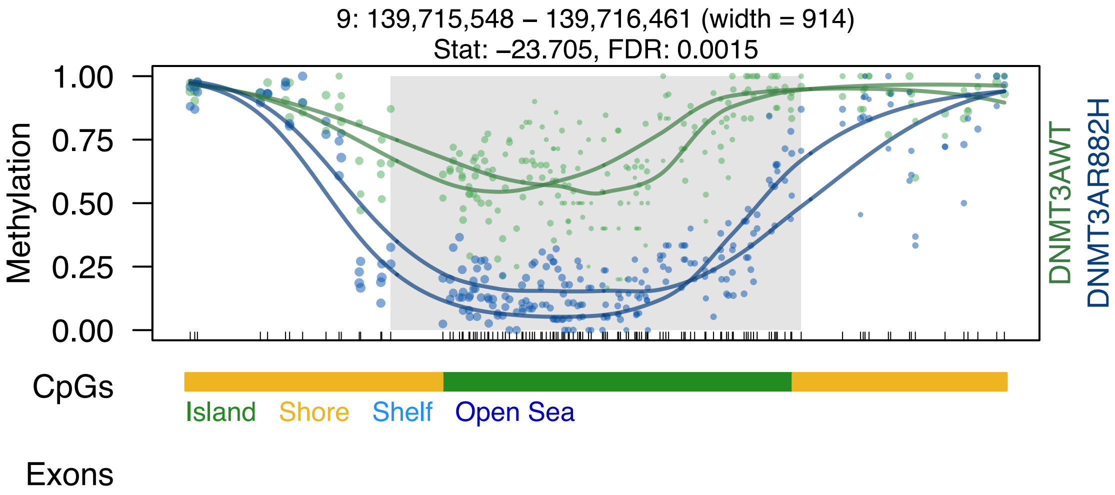

9: 139,715,548 – 139,716,461 (width = 914)
Stat: –23.705, FDR: 0.0015
Methylation
1.00
0.75
0.50
0.25
0.00
DNMT3AWT
DNMT3AR882H
CpGs
Island
Shore
Shelf
Open Sea
Exons

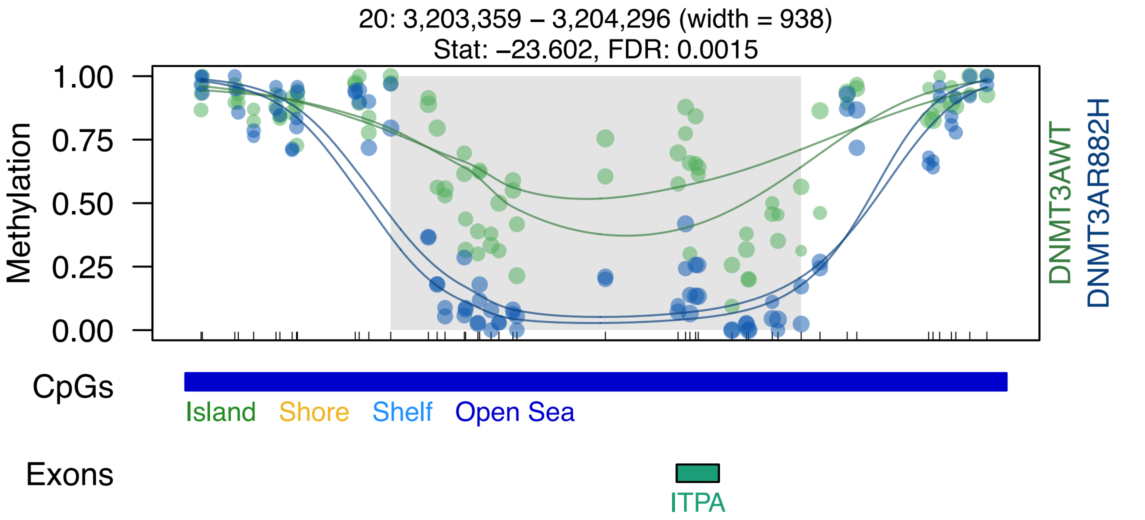

20: 3,203,359 – 3,204,296 (width = 938)
Stat: –23.602, FDR: 0.0015
1.00
0.75
0.50
0.25
0.00
Methylation
DNMT3AWT
DNMT3AR882H
CpGs
Island
Shore
Shelf
Open Sea
Exons
ITPA

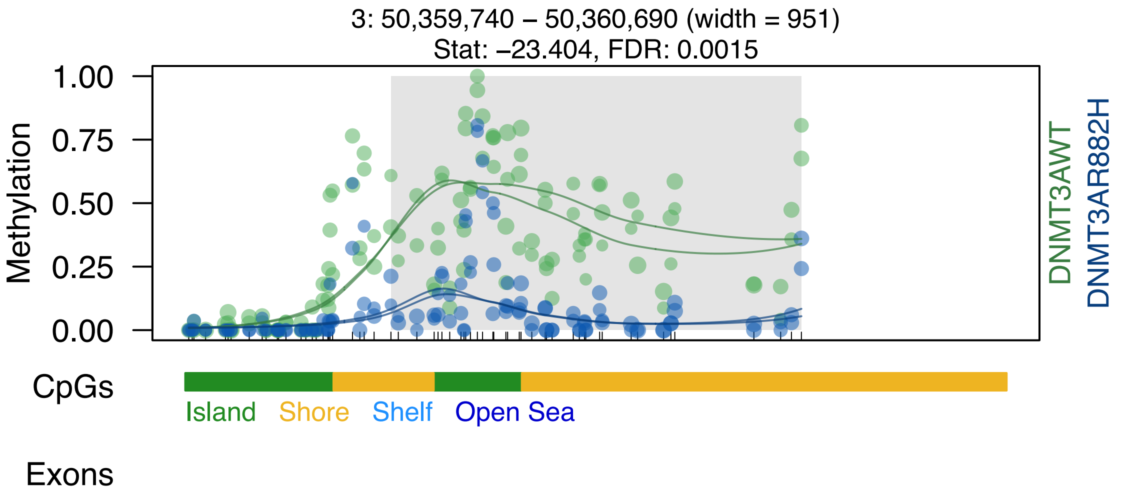
3: 50,359,740 – 50,360,690 (width = 951)
Stat: –23.404, FDR: 0.0015
Methylation
1.00
0.75
0.50
0.25
0.00
DNMT3AWT
DNMT3AR882H
CpGs
Island
Shore
Shelf
Open Sea
Exons

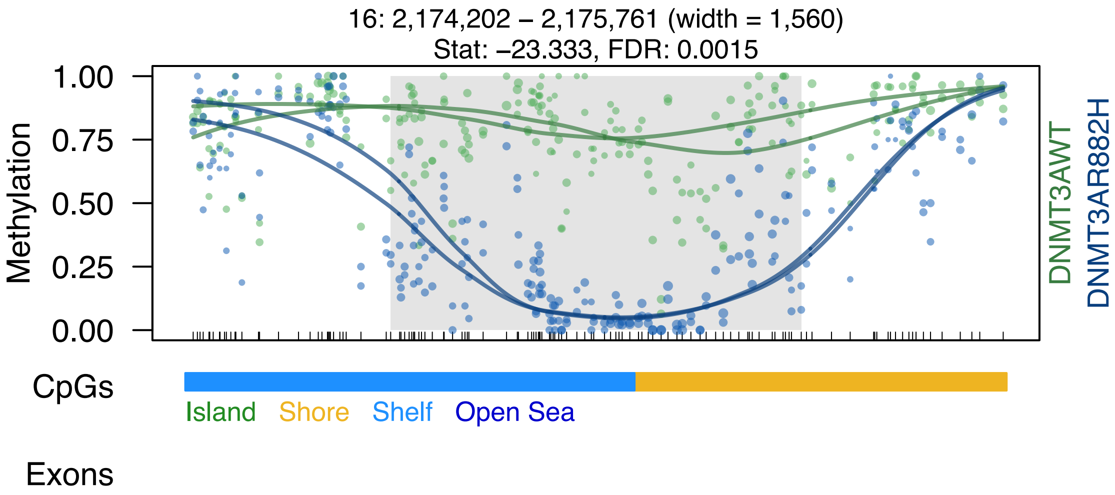

16: 2,174,202 – 2,175,761 (width = 1,560)
Stat: –23.333, FDR: 0.0015
Methylation
1.00
0.75
0.50
0.25
0.00
DNMT3AWT
DNMT3AR882H
CpGs
Island
Shore
Shelf
Open Sea
Exons

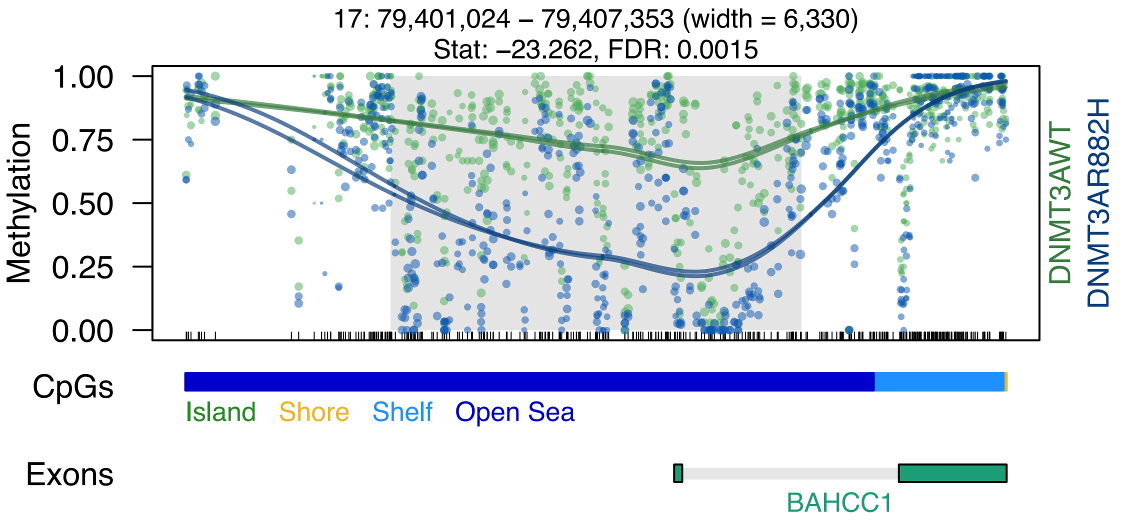
17: 79,401,024 – 79,407,353 (width = 6,330)
Stat: –23.262, FDR: 0.0015
Methylation
1.00
0.75
0.50
0.25
0.00
DNMT3AWT
DNMT3AR882H
CpGs
Island
Shore
Shelf
Open Sea
Exons
BAHCC1

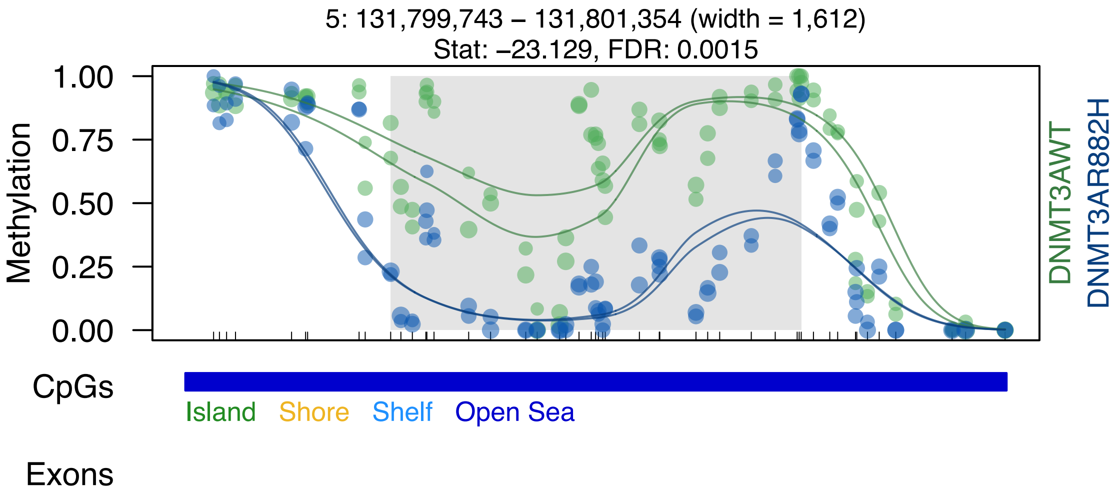

5: 131,799,743 – 131,801,354 (width = 1,612)
Stat: –23.129, FDR: 0.0015
1.00
0.75
0.50
0.25
0.00
Methylation
DNMT3AWT
DNMT3AR882H
CpGs
Island
Shore
Shelf
Open Sea
Exons

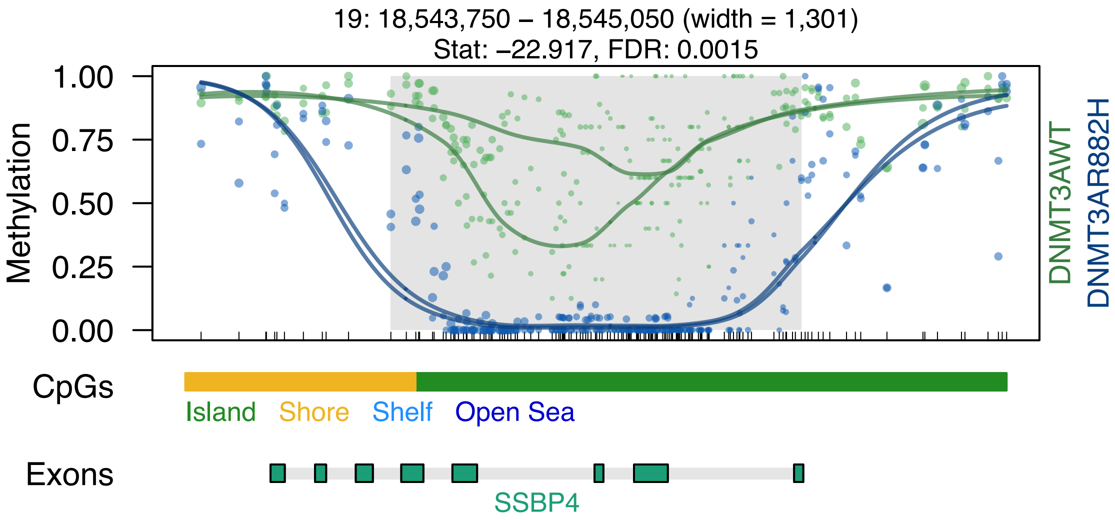

19: 18,543,750 – 18,545,050 (width = 1,301)
Stat: –22.917, FDR: 0.0015
1.00
0.75
0.50
0.25
0.00
Methylation
DNMT3AWT
DNMT3AR882H
CpGs
Island
Shore
Shelf
Open Sea
Exons
SSBP4

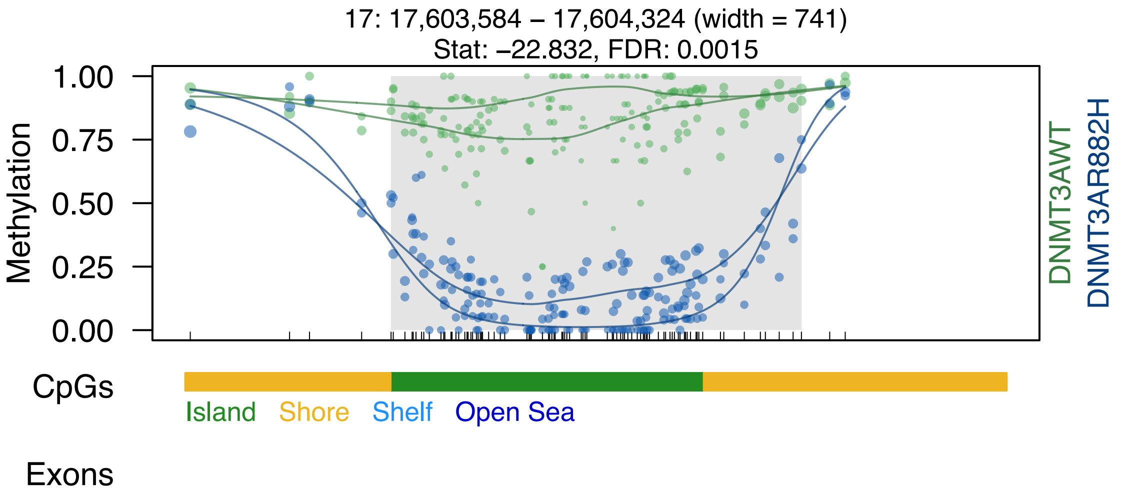
17: 17,603,584 – 17,604,324 (width = 741)
Stat: –22.832, FDR: 0.0015
Methylation
1.00
0.75
0.50
0.25
0.00
DNMT3AWT
DNMT3AR882H
CpGs
Island
Shore
Shelf
Open Sea
Exons

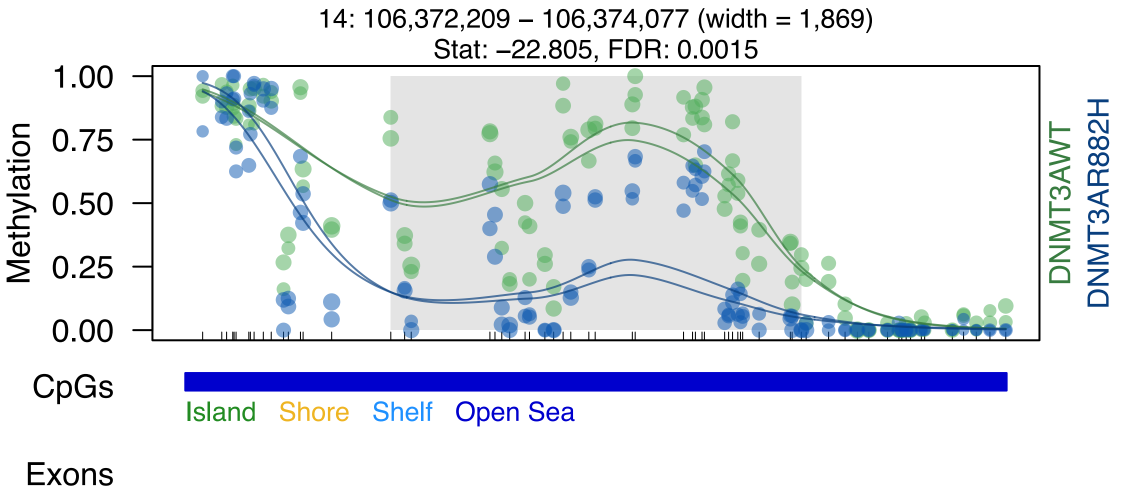

14: 106,372,209 – 106,374,077 (width = 1,869)
Stat: –22.805, FDR: 0.0015
Methylation
1.00
0.75
0.50
0.25
0.00
DNMT3AWT
DNMT3AR882H
CpGs
Island
Shore
Shelf
Open Sea
Exons

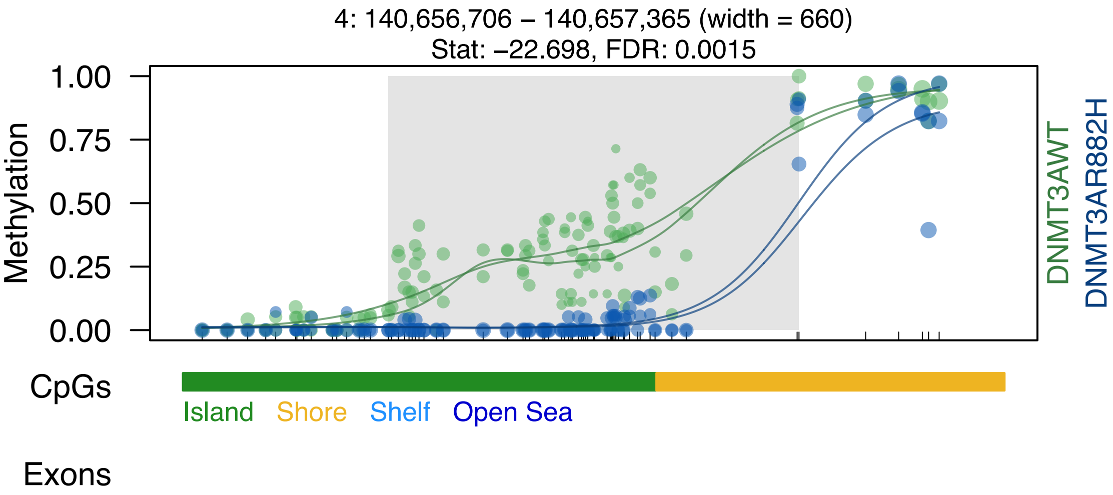

4: 140,656,706 – 140,657,365 (width = 660)
Stat: –22.698, FDR: 0.0015
Methylation
1.00
0.75
0.50
0.25
0.00
DNMT3AWT
DNMT3AR882H
CpGs
Island
Shore
Shelf
Open Sea
Exons

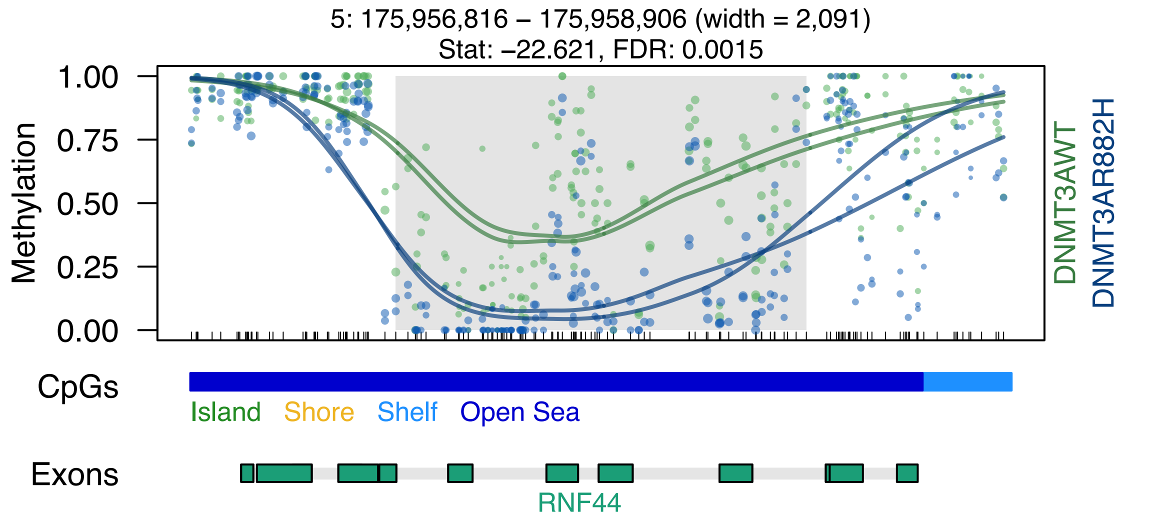
5: 175,956,816 – 175,958,906 (width = 2,091)
Stat: –22.621, FDR: 0.0015
Methylation
1.00
0.75
0.50
0.25
0.00
DNMT3AWT
DNMT3AR882H
CpGs
Island
Shore
Shelf
Open Sea
Exons
RNF44

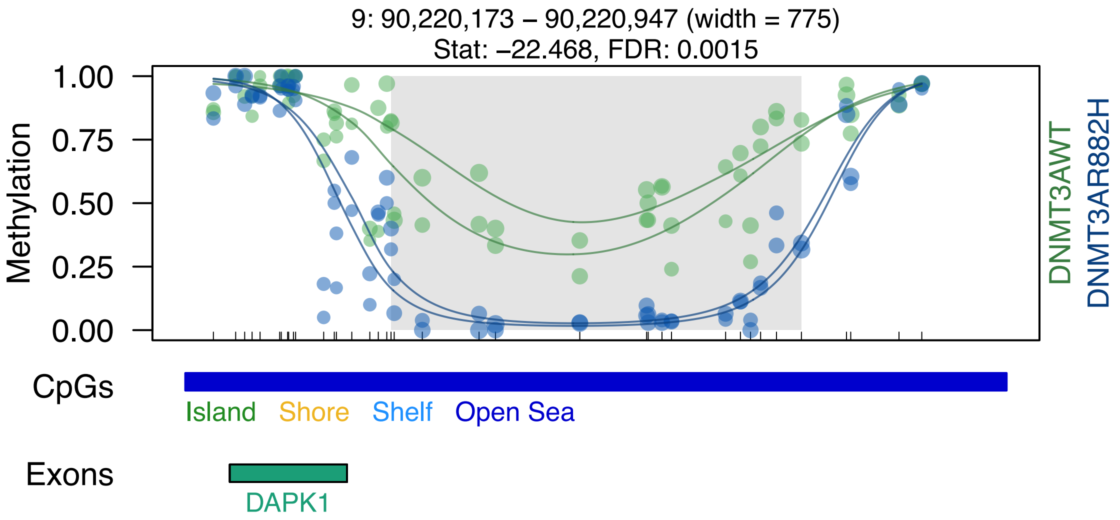

9: 90,220,173 – 90,220,947 (width = 775)
Stat: –22.468, FDR: 0.0015
Methylation
1.00
0.75
0.50
0.25
0.00
DNMT3AWT
DNMT3AR882H
CpGs
Island
Shore
Shelf
Open Sea
Exons
DAPK1

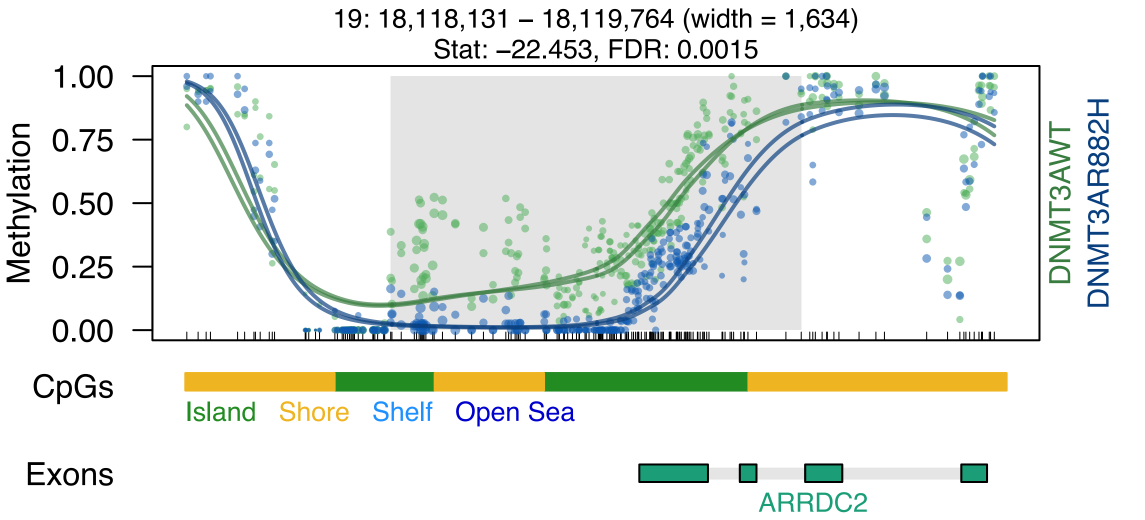

19: 18,118,131 – 18,119,764 (width = 1,634)
Stat: −22.453, FDR: 0.0015
Methylation
1.00
0.75
0.50
0.25
0.00
DNMT3AWT
DNMT3AR882H
CpGs
Island
Shore
Shelf
Open Sea
Exons
ARRDC2

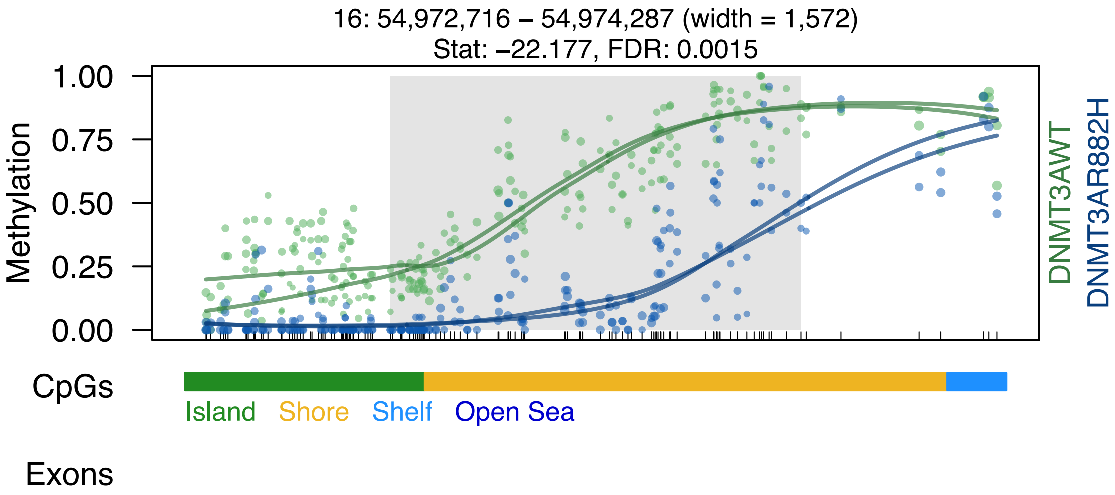
16: 54,972,716 – 54,974,287 (width = 1,572)
Stat: –22.177, FDR: 0.0015
Methylation
1.00
0.75
0.50
0.25
0.00
DNMT3AWT
DNMT3AR882H
CpGs
Island
Shore
Shelf
Open Sea
Exons

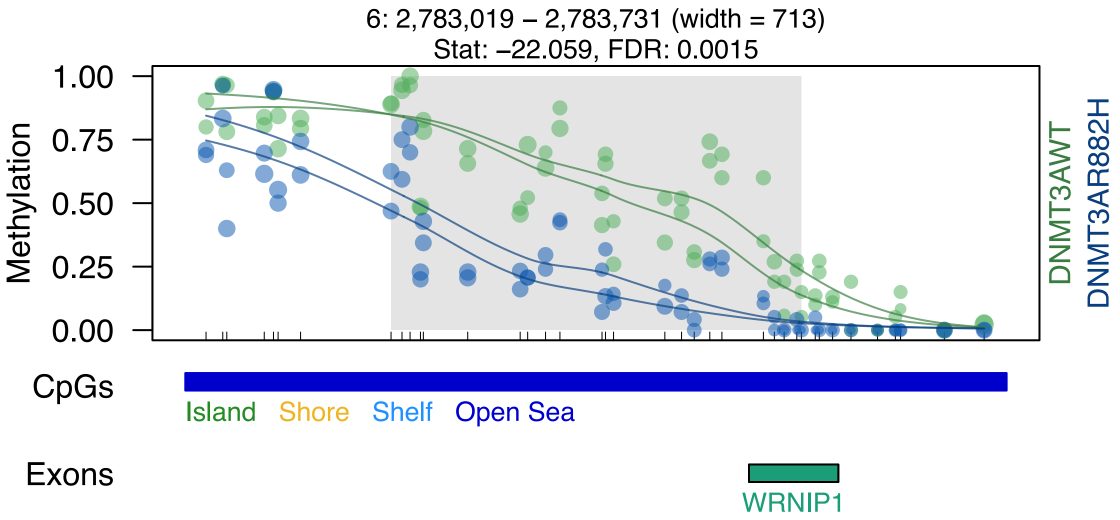
6: 2,783,019 – 2,783,731 (width = 713)
Stat: –22.059, FDR: 0.0015
Methylation
1.00
0.75
0.50
0.25
0.00
DNMT3AWT
DNMT3AR882H
CpGs
Island
Shore
Shelf
Open Sea
Exons
WRNIP1

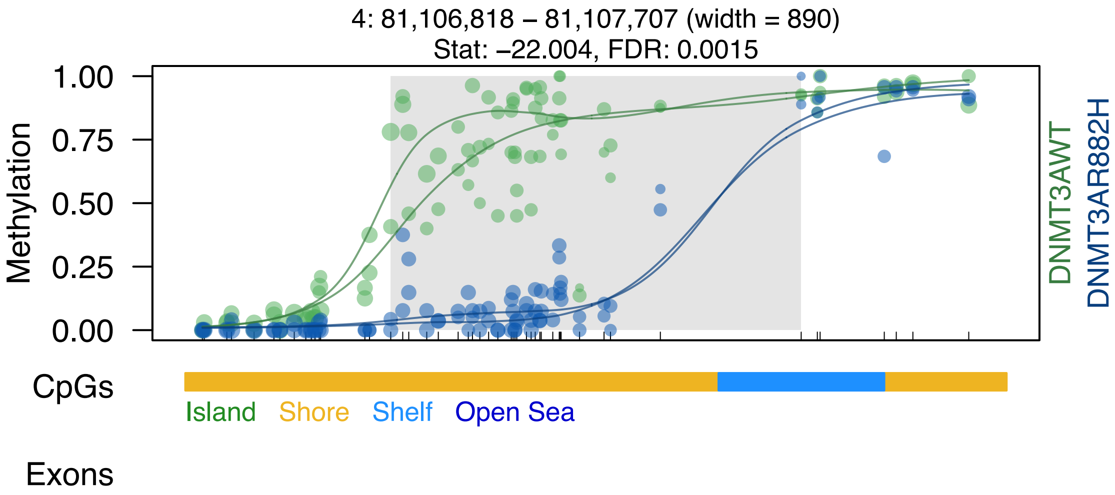

4: 81,106,818 – 81,107,707 (width = 890)
Stat: –22.004, FDR: 0.0015
Methylation
1.00
0.75
0.50
0.25
0.00
DNMT3AWT
DNMT3AR882H
CpGs
Island
Shore
Shelf
Open Sea
Exons

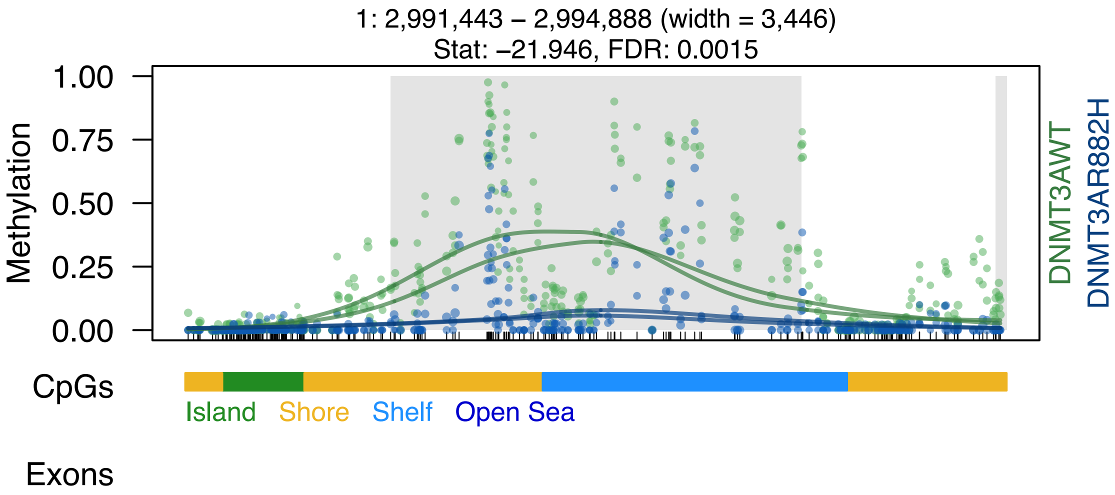

1: 2,991,443 – 2,994,888 (width = 3,446)
Stat: −21.946, FDR: 0.0015
Methylation
1.00
0.75
0.50
0.25
0.00
DNMT3AWT
DNMT3AR882H
CpGs
Island
Shore
Shelf
Open Sea
Exons

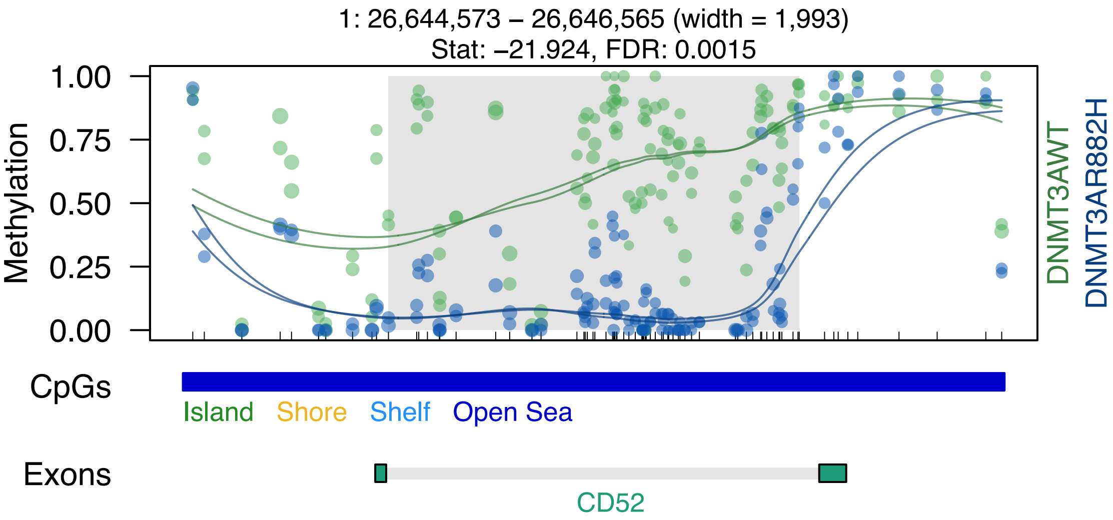

1: 26,644,573 – 26,646,565 (width = 1,993)
Stat: –21.924, FDR: 0.0015
Methylation
1.00
0.75
0.50
0.25
0.00
DNMT3AWT
DNMT3AR882H
CpGs
Island
Shore
Shelf
Open Sea
Exons
CD52

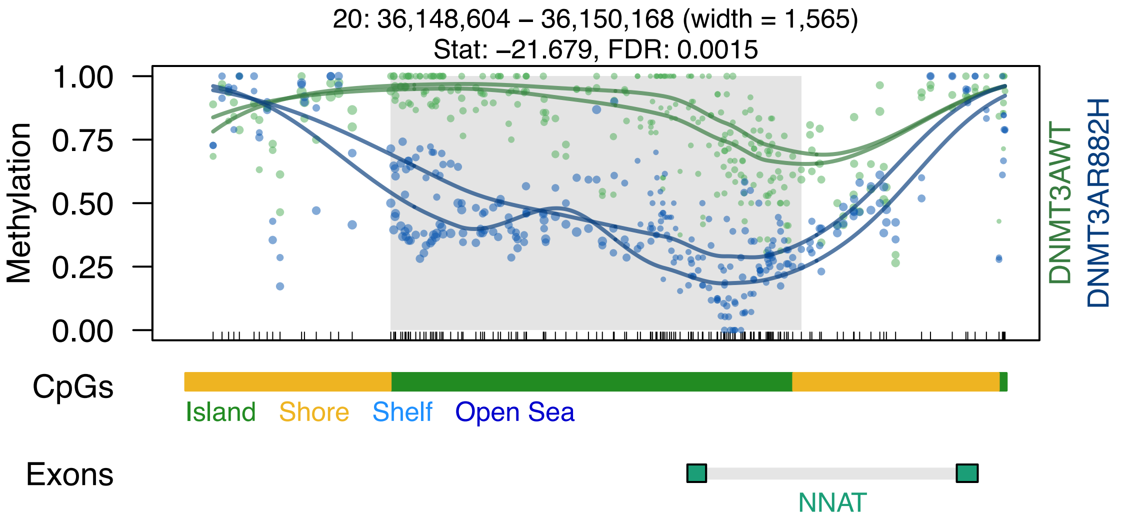

20: 36,148,604 – 36,150,168 (width = 1,565)
Stat: –21.679, FDR: 0.0015
1.00
0.75
0.50
0.25
0.00
Methylation
DNMT3AWT
DNMT3AR882H
CpGs
Island
Shore
Shelf
Open Sea
Exons
NNAT

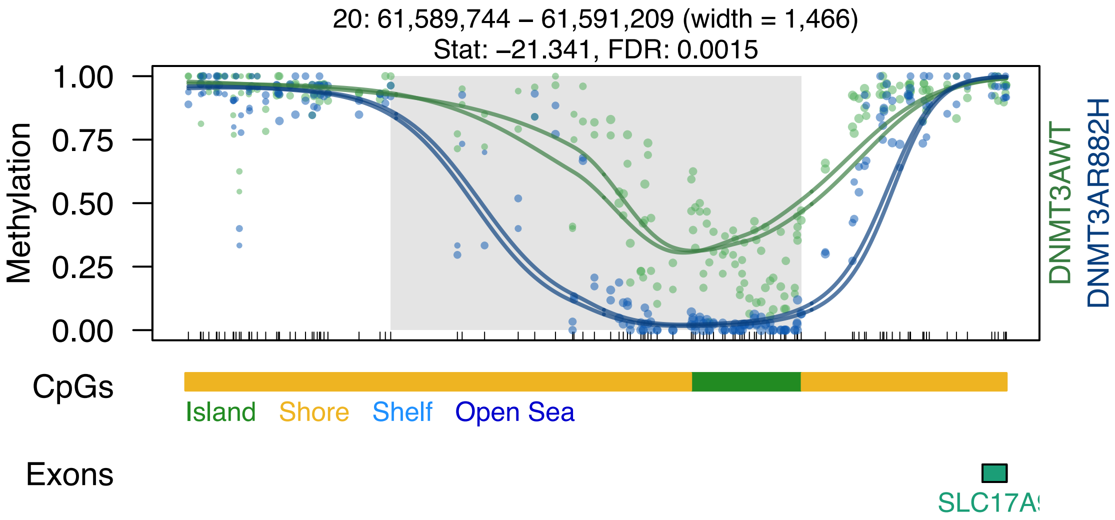

20: 61,589,744 – 61,591,209 (width = 1,466)
Stat: –21.341, FDR: 0.0015
Methylation
1.00
0.75
0.50
0.25
0.00
DNMT3AWT
DNMT3AR882H
CpGs
Island
Shore
Shelf
Open Sea
Exons
SLC17A

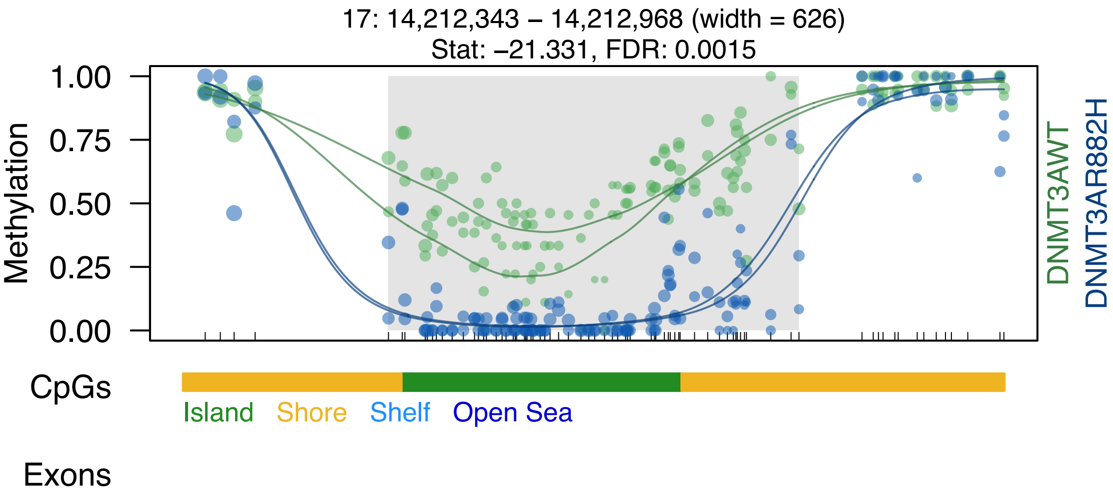
17: 14,212,343 – 14,212,968 (width = 626)
Stat: –21.331, FDR: 0.0015
Methylation
1.00
0.75
0.50
0.25
0.00
DNMT3AWT
DNMT3AR882H
CpGs
Island
Shore
Shelf
Open Sea
Exons

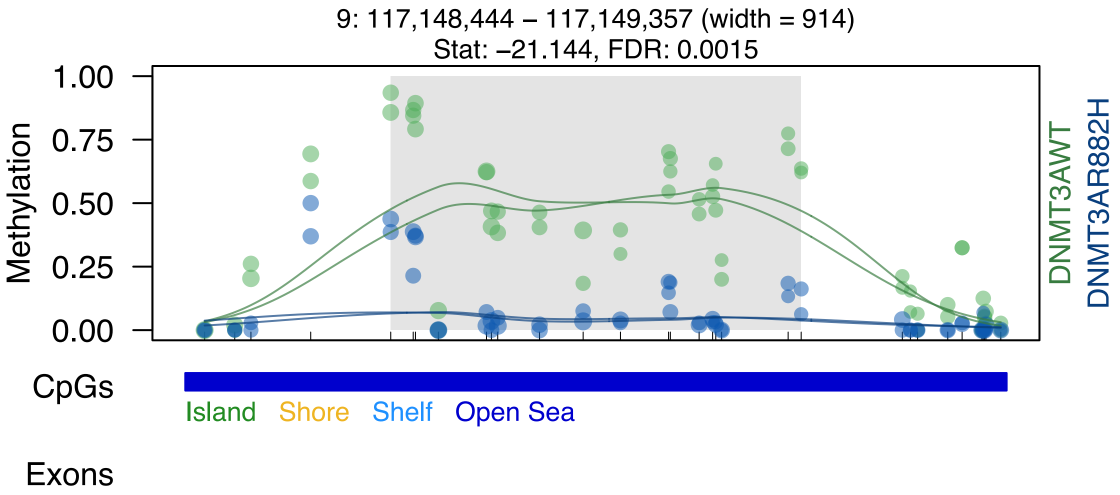

9: 117,148,444 – 117,149,357 (width = 914)
Stat: –21.144, FDR: 0.0015
Methylation
1.00
0.75
0.50
0.25
0.00
DNMT3AWT
DNMT3AR882H
CpGs
Island
Shore
Shelf
Open Sea
Exons

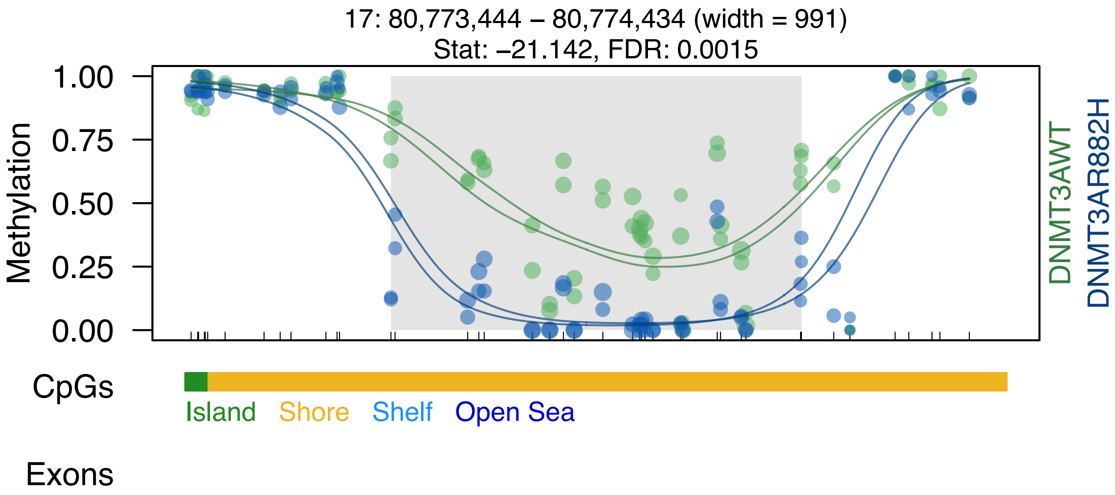
17: 80,773,444 – 80,774,434 (width = 991)
Stat: –21.142, FDR: 0.0015
1.00
0.75
0.50
0.25
0.00
Methylation
DNMT3AWT
DNMT3AR882H
CpGs
Island
Shore
Shelf
Open Sea
Exons

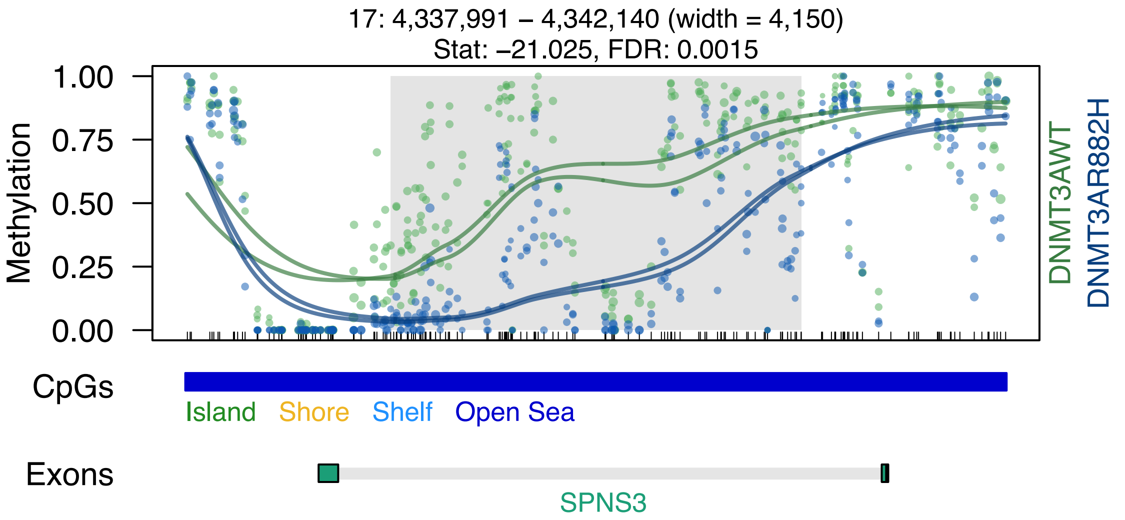
17: 4,337,991 – 4,342,140 (width = 4,150)
Stat: –21.025, FDR: 0.0015
Methylation
1.00
0.75
0.50
0.25
0.00
DNMT3AWT
DNMT3AR882H
CpGs
Island
Shore
Shelf
Open Sea
Exons
SPNS3

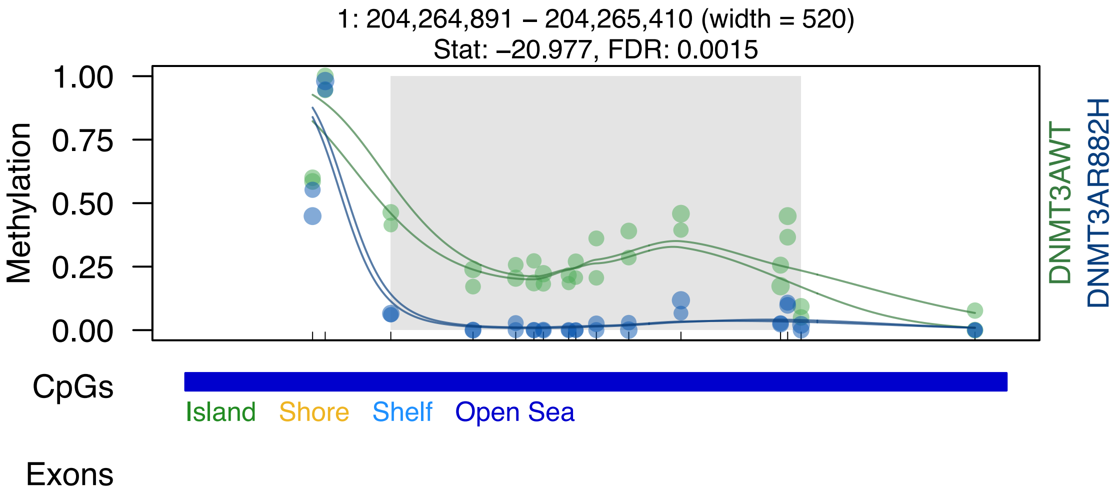
1: 204,264,891 – 204,265,410 (width = 520)
Stat: –20.977, FDR: 0.0015
1.00
0.75
0.50
0.25
0.00
Methylation
DNMT3AWT
DNMT3AR882H
CpGs
Island
Shore
Shelf
Open Sea
Exons

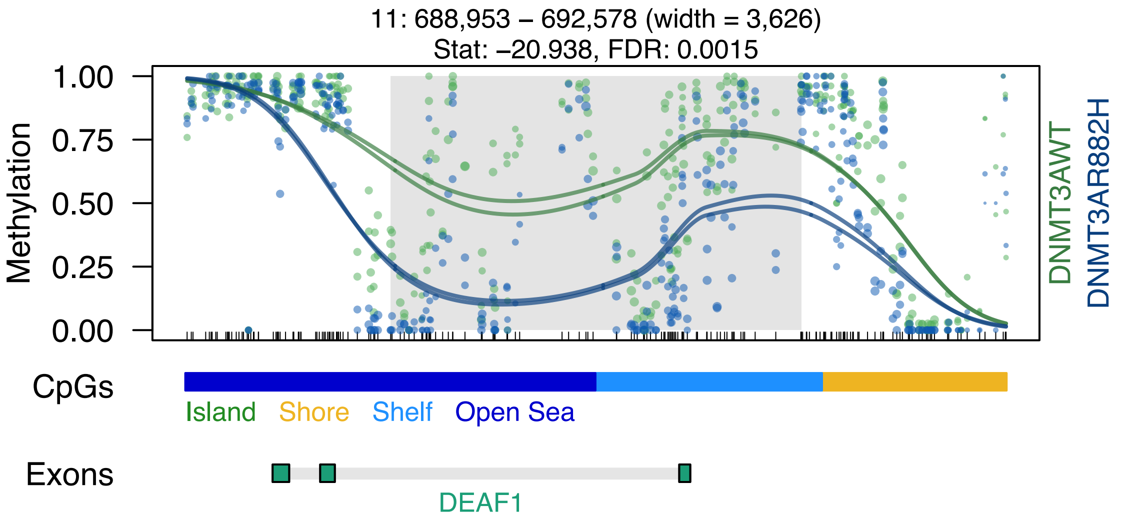

11: 688,953 – 692,578 (width = 3,626)
Stat: –20.938, FDR: 0.0015
1.00
0.75
0.50
0.25
0.00
Methylation
DNMT3AWT
DNMT3AR882H
CpGs
Island
Shore
Shelf
Open Sea
Exons
DEAF1

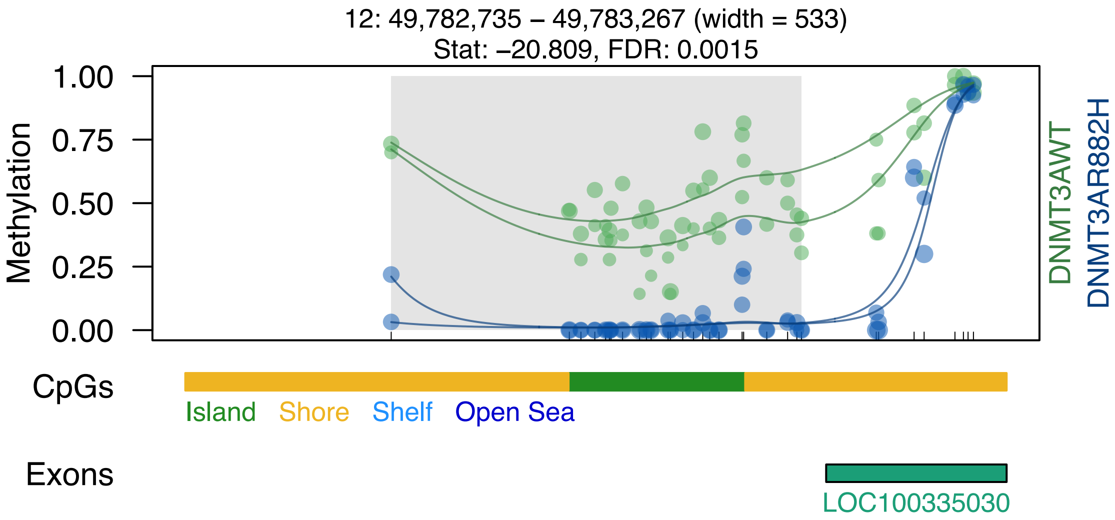

12: 49,782,735 – 49,783,267 (width = 533)
Stat: –20.809, FDR: 0.0015
Methylation
1.00
0.75
0.50
0.25
0.00
DNMT3AWT
DNMT3AR882H
CpGs
Island
Shore
Shelf
Open Sea
Exons
LOC100335030

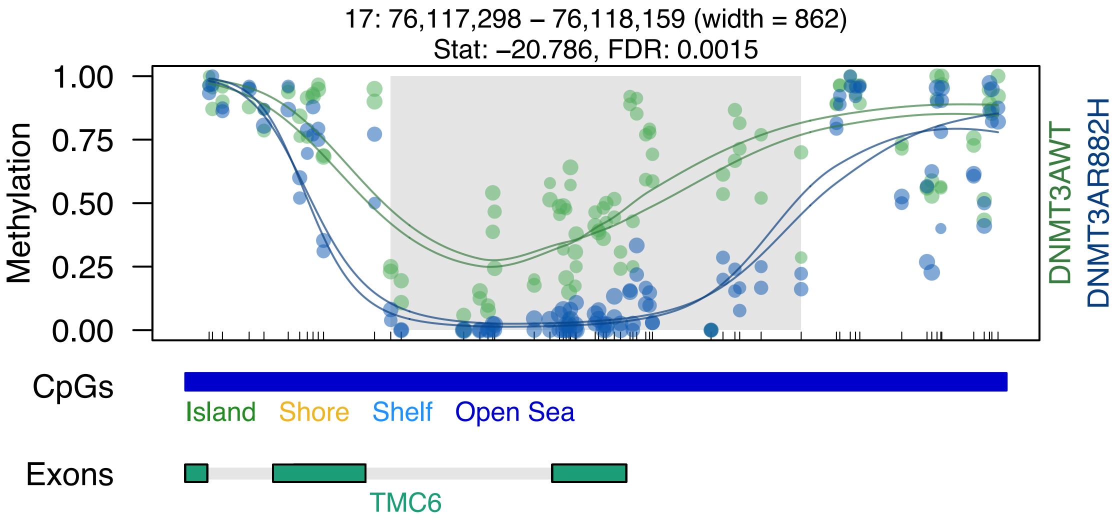

17: 76,117,298 – 76,118,159 (width = 862)
Stat: –20.786, FDR: 0.0015
Methylation
1.00
0.75
0.50
0.25
0.00
DNMT3AWT
DNMT3AR882H
CpGs
Island
Shore
Shelf
Open Sea
Exons
TMC6

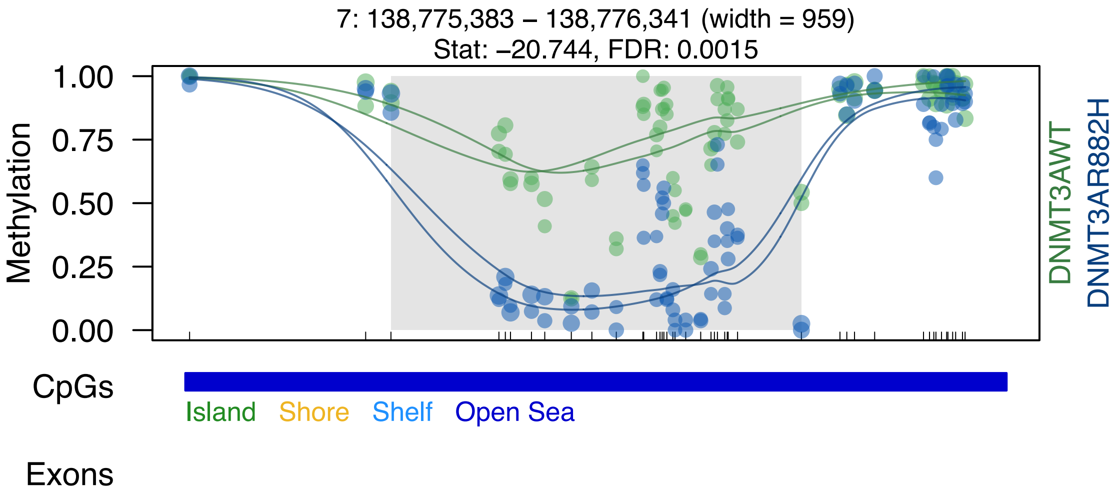

7: 138,775,383 – 138,776,341 (width = 959)
Stat: –20.744, FDR: 0.0015
Methylation
1.00
0.75
0.50
0.25
0.00
DNMT3AWT
DNMT3AR882H
CpGs
Island
Shore
Shelf
Open Sea
Exons

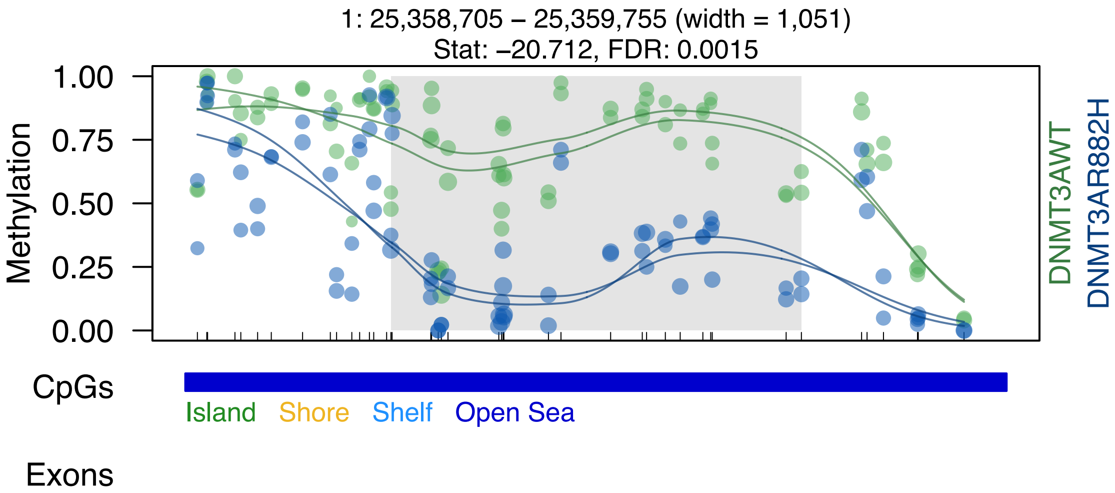
1: 25,358,705 – 25,359,755 (width = 1,051)
Stat: –20.712, FDR: 0.0015
1.00
0.75
0.50
0.25
0.00
Methylation
DNMT3AWT
DNMT3AR882H
CpGs
Island
Shore
Shelf
Open Sea
Exons

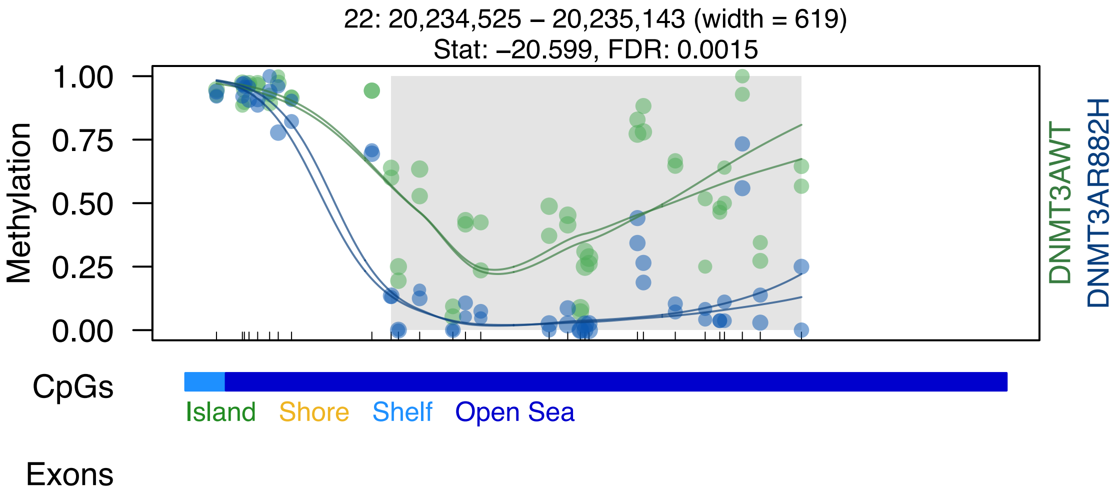
22: 20,234,525 – 20,235,143 (width = 619)
Stat: −20.599, FDR: 0.0015
Methylation
1.00
0.75
0.50
0.25
0.00
DNMT3AWT
DNMT3AR882H
CpGs
Island
Shore
Shelf
Open Sea
Exons

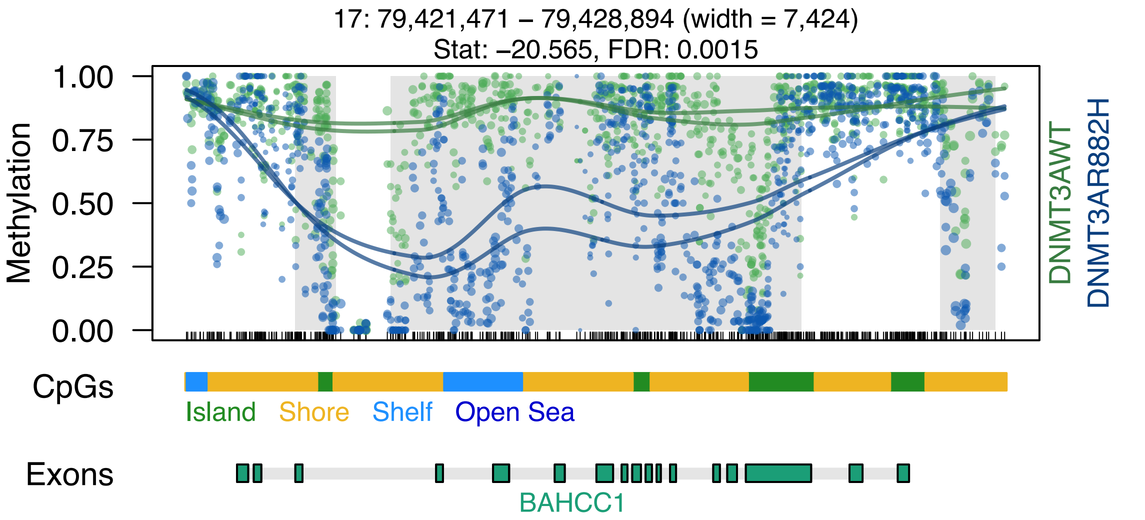

17: 79,421,471 – 79,428,894 (width = 7,424)
Stat: –20.565, FDR: 0.0015
Methylation
1.00
0.75
0.50
0.25
0.00
DNMT3AWT
DNMT3AR882H
CpGs
Island
Shore
Shelf
Open Sea
Exons
BAHCC1

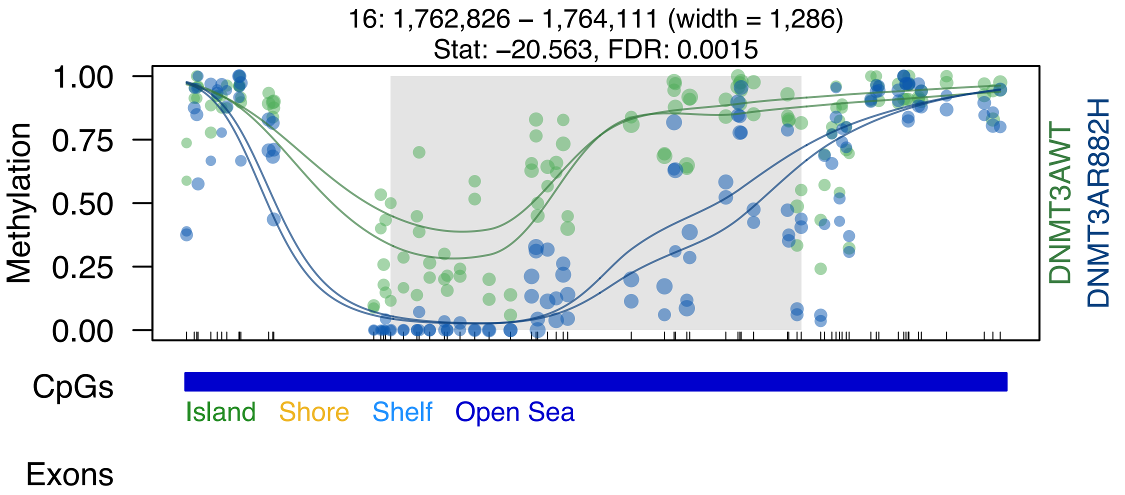
16: 1,762,826 – 1,764,111 (width = 1,286)
Stat: –20.563, FDR: 0.0015
Methylation
1.00
0.75
0.50
0.25
0.00
DNMT3AWT
DNMT3AR882H
CpGs
Island
Shore
Shelf
Open Sea
Exons

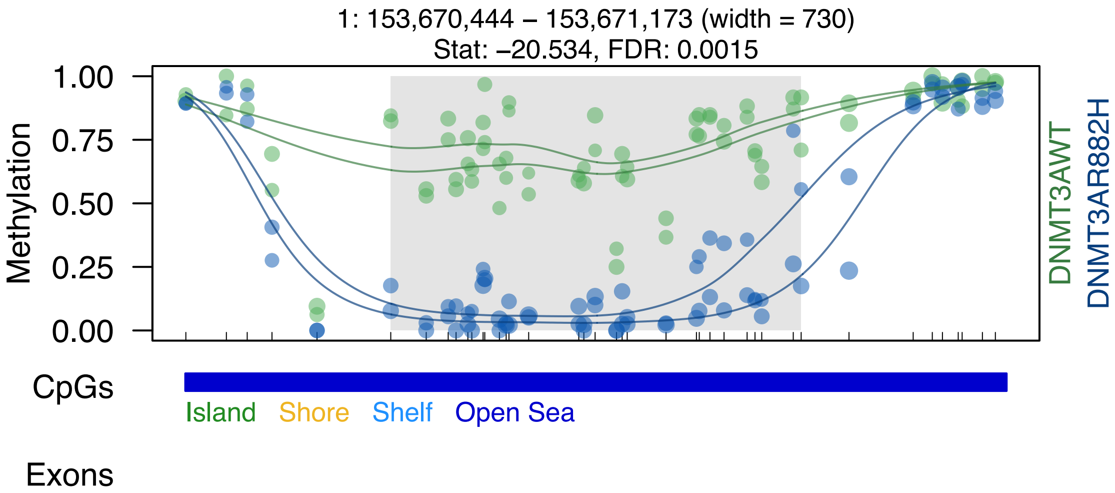
1: 153,670,444 – 153,671,173 (width = 730)
Stat: –20.534, FDR: 0.0015
Methylation
1.00
0.75
0.50
0.25
0.00
DNMT3AWT
DNMT3AR882H
CpGs
Island
Shore
Shelf
Open Sea
Exons

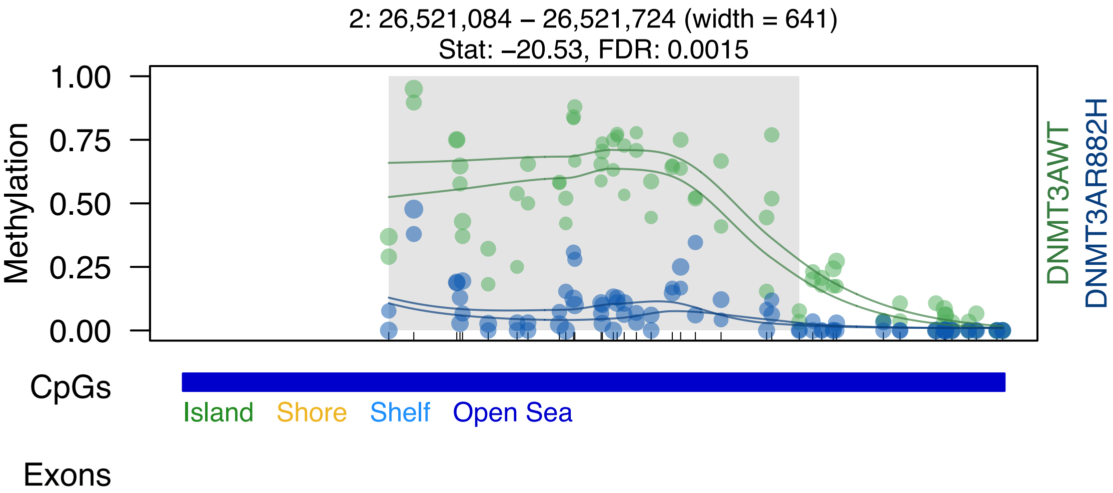

2: 26,521,084 – 26,521,724 (width = 641)
Stat: –20.53, FDR: 0.0015
Methylation
1.00
0.75
0.50
0.25
0.00
DNMT3AWT
DNMT3AR882H
CpGs
Island
Shore
Shelf
Open Sea
Exons

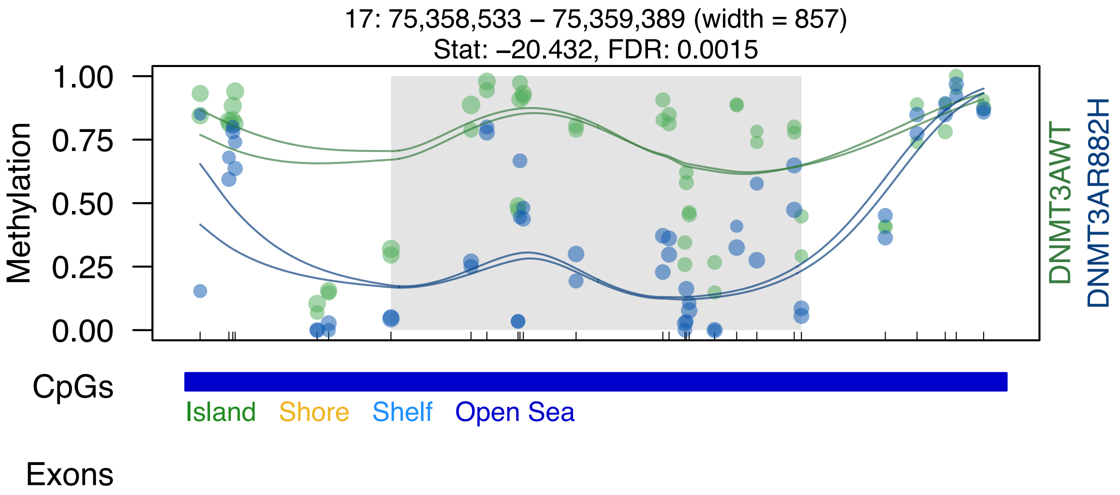

17: 75,358,533 – 75,359,389 (width = 857)
Stat: –20.432, FDR: 0.0015
1.00
0.75
0.50
0.25
0.00
Methylation
DNMT3AWT
DNMT3AR882H
CpGs
Island
Shore
Shelf
Open Sea
Exons

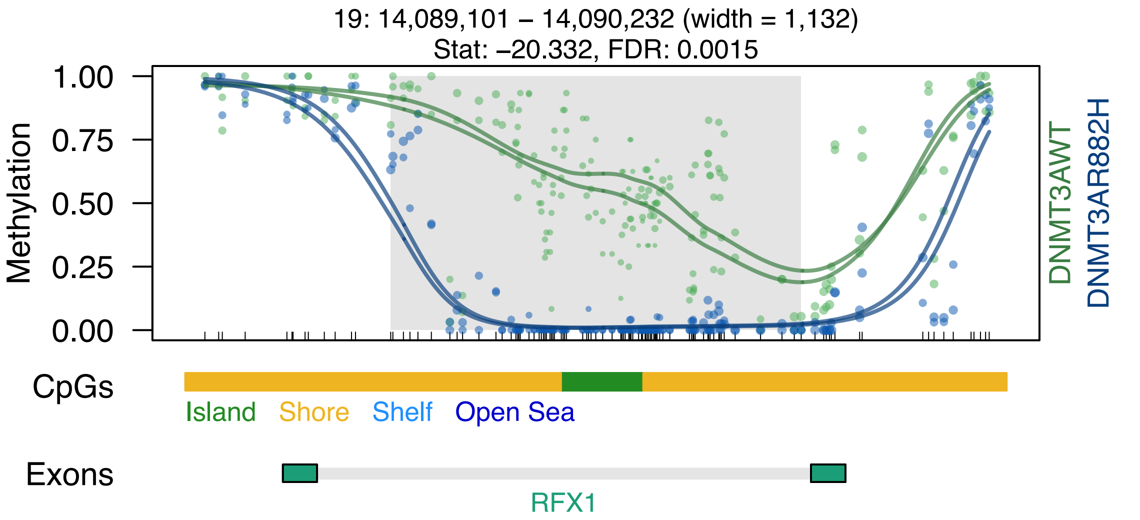

19: 14,089,101 – 14,090,232 (width = 1,132)
Stat: –20.332, FDR: 0.0015
Methylation
1.00
0.75
0.50
0.25
0.00
DNMT3AWT
DNMT3AR882H
CpGs
Island
Shore
Shelf
Open Sea
Exons
RFX1

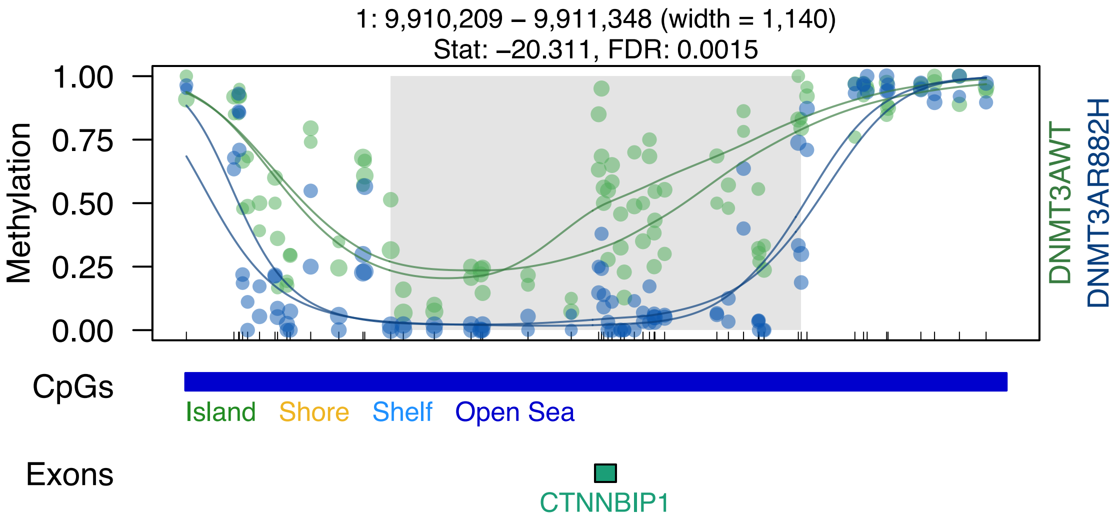
1: 9,910,209 – 9,911,348 (width = 1,140)
Stat: –20.311, FDR: 0.0015
Methylation
1.00
0.75
0.50
0.25
0.00
DNMT3AWT
DNMT3AR882H
CpGs
Island
Shore
Shelf
Open Sea
Exons
CTNNBIP1

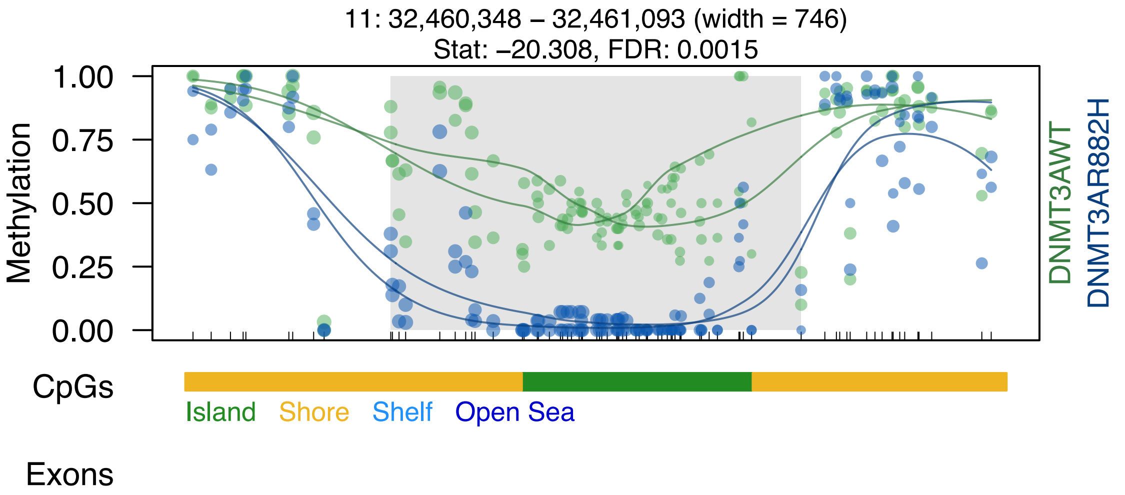

11: 32,460,348 – 32,461,093 (width = 746)
Stat: –20.308, FDR: 0.0015
Methylation
1.00
0.75
0.50
0.25
0.00
DNMT3AWT
DNMT3AR882H
CpGs
Island
Shore
Shelf
Open Sea
Exons

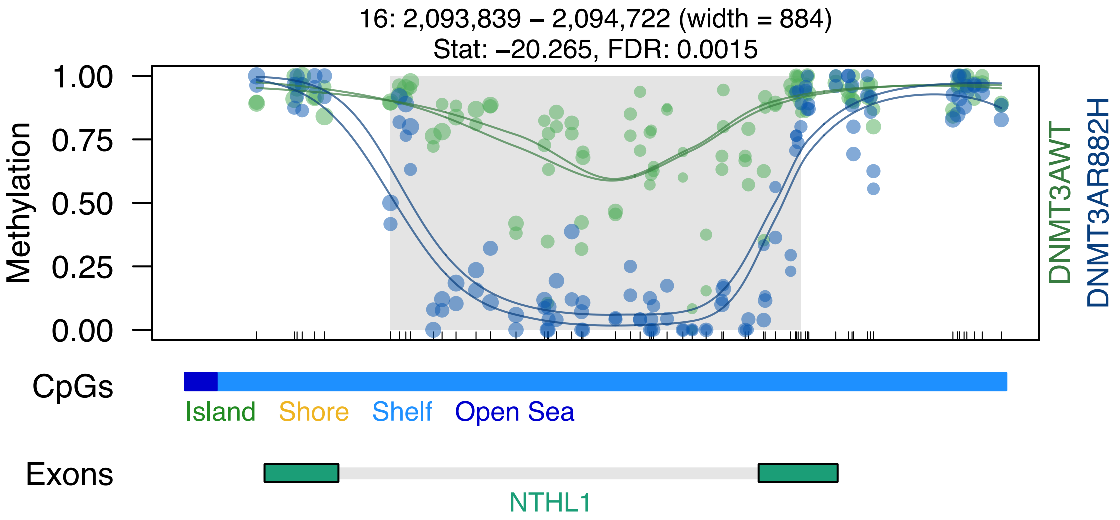
16: 2,093,839 – 2,094,722 (width = 884)
Stat: –20.265, FDR: 0.0015
Methylation
1.00
0.75
0.50
0.25
0.00
DNMT3AWT
DNMT3AR882H
CpGs
Island
Shore
Shelf
Open Sea
Exons
NTHL1

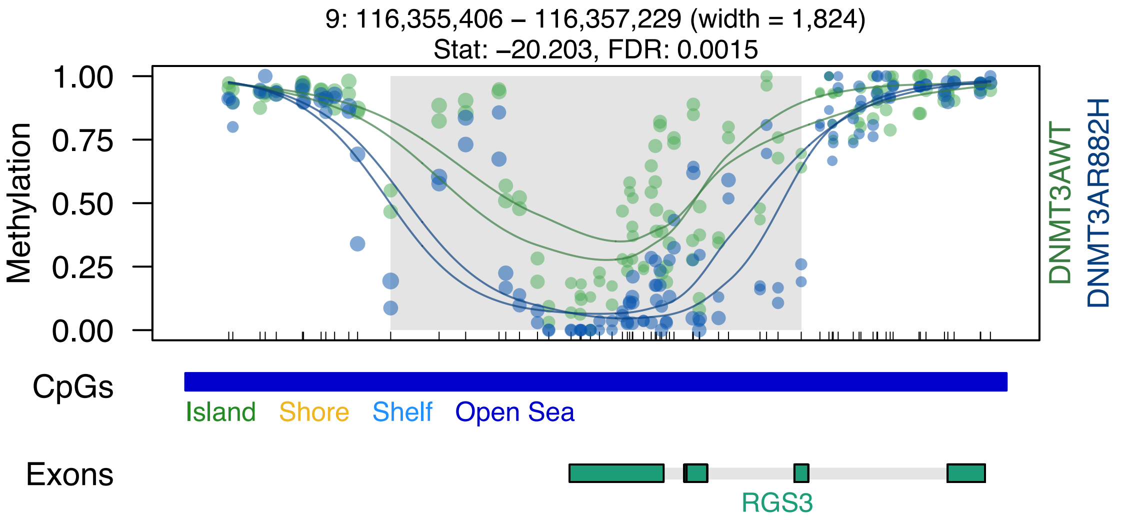

9: 116,355,406 – 116,357,229 (width = 1,824)
Stat: –20.203, FDR: 0.0015
1.00
0.75
0.50
0.25
0.00
Methylation
DNMT3AWT
DNMT3AR882H
CpGs
Island
Shore
Shelf
Open Sea
Exons
RGS3

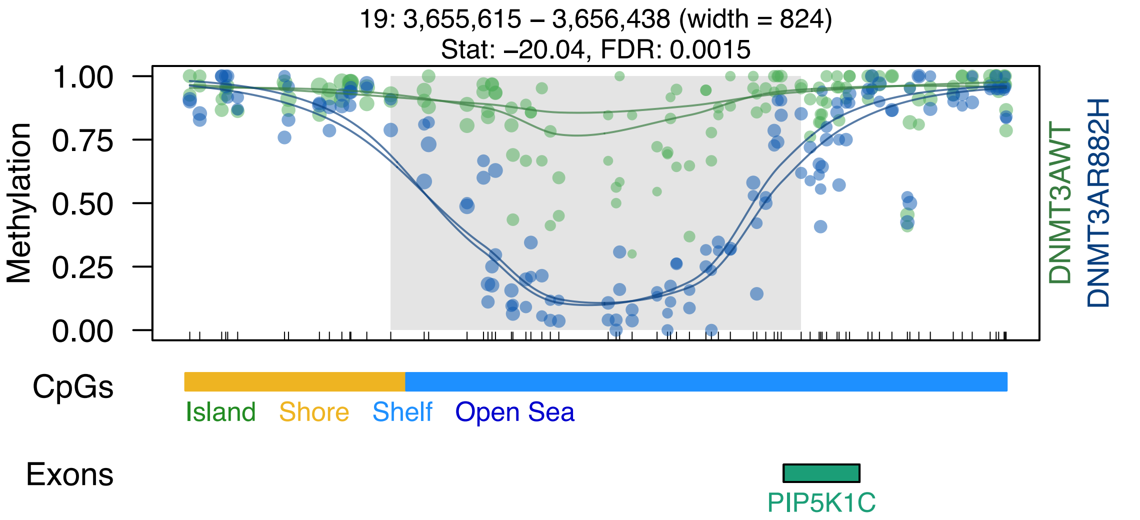
19: 3,655,615 – 3,656,438 (width = 824)
Stat: –20.04, FDR: 0.0015
1.00
0.75
0.50
0.25
0.00
Methylation
DNMT3AWT
DNMT3AR882H
CpGs
Island
Shore
Shelf
Open Sea
Exons
PIP5K1C

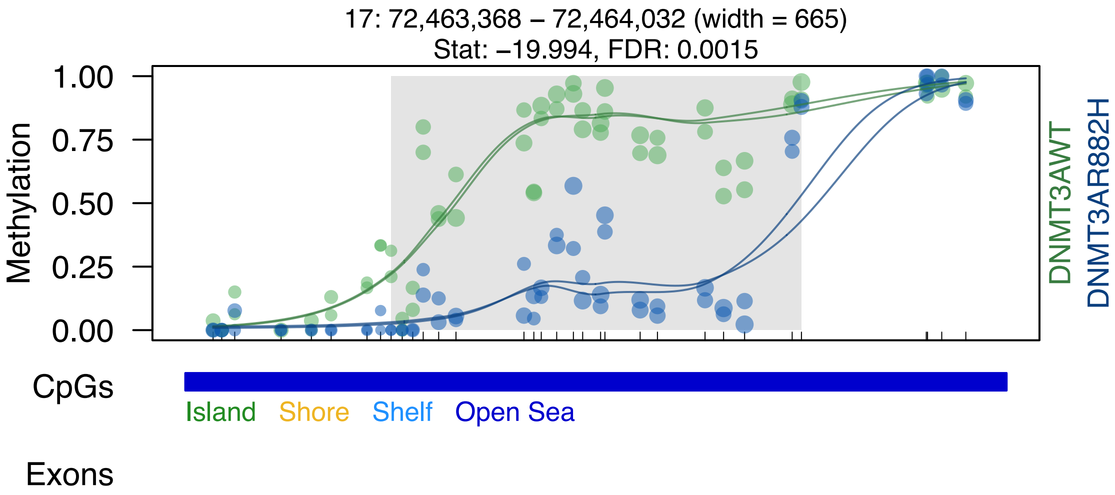

17: 72,463,368 – 72,464,032 (width = 665)
Stat: –19.994, FDR: 0.0015
Methylation
1.00
0.75
0.50
0.25
0.00
DNMT3AWT
DNMT3AR882H
CpGs
Island
Shore
Shelf
Open Sea
Exons

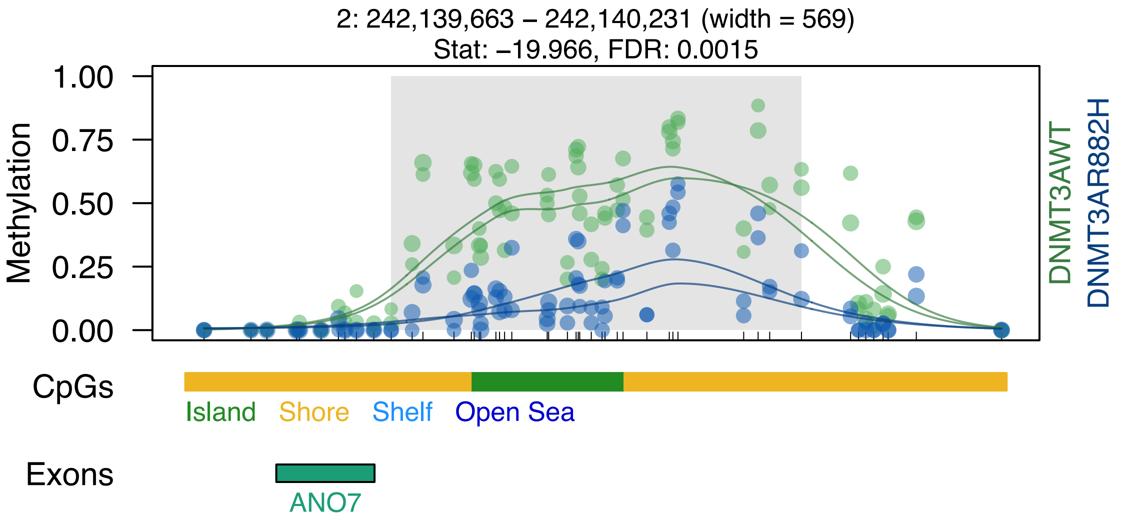
2: 242,139,663 – 242,140,231 (width = 569)
Stat: −19.966, FDR: 0.0015
Methylation
1.00
0.75
0.50
0.25
0.00
DNMT3AWT
DNMT3AR882H
CpGs
Island
Shore
Shelf
Open Sea
Exons
ANO7

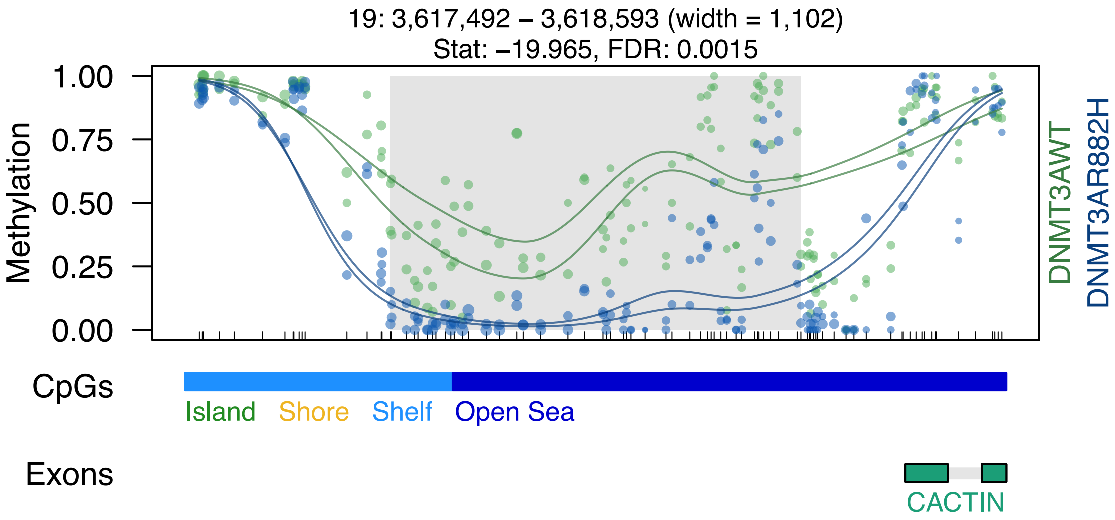

19: 3,617,492 – 3,618,593 (width = 1,102)
Stat: –19.965, FDR: 0.0015
Methylation
1.00
0.75
0.50
0.25
0.00
DNMT3AWT
DNMT3AR882H
CpGs
Island
Shore
Shelf
Open Sea
Exons
CACTIN

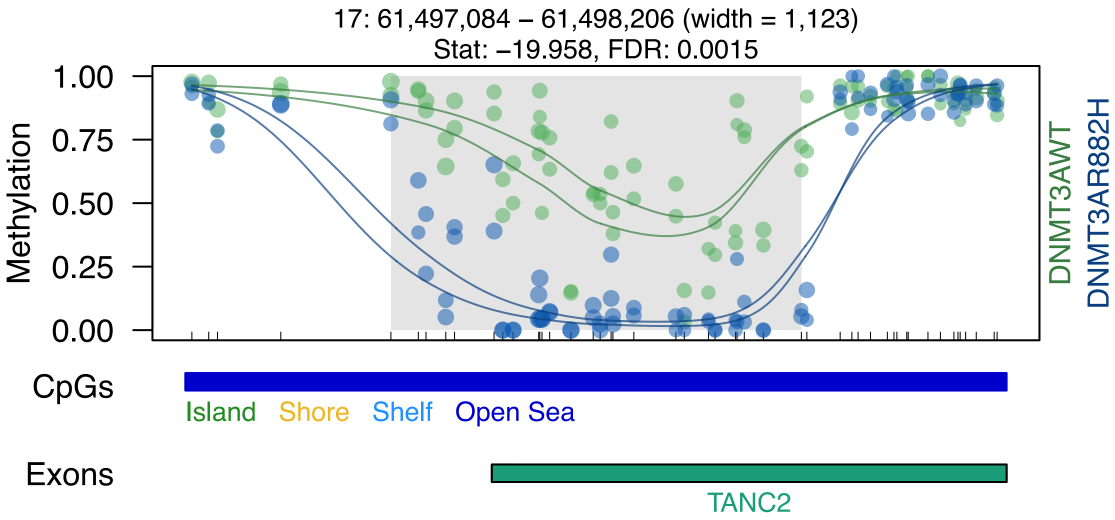
17: 61,497,084 – 61,498,206 (width = 1,123)
Stat: –19.958, FDR: 0.0015
Methylation
1.00
0.75
0.50
0.25
0.00
DNMT3AWT
DNMT3AR882H
CpGs
Island
Shore
Shelf
Open Sea
Exons
TANC2

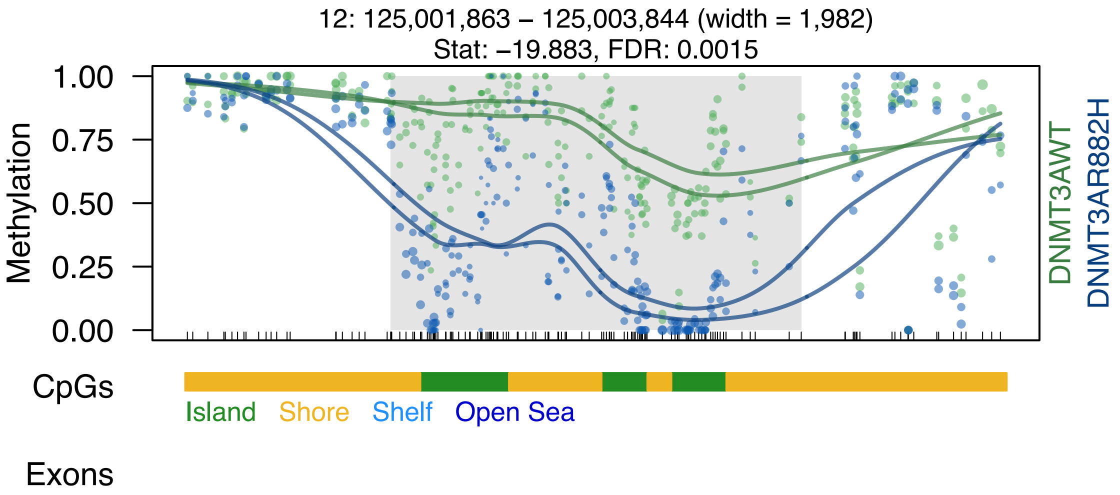
12: 125,001,863 – 125,003,844 (width = 1,982)
Stat: −19.883, FDR: 0.0015
Methylation
1.00
0.75
0.50
0.25
0.00
DNMT3AWT
DNMT3AR882H
CpGs
Island
Shore
Shelf
Open Sea
Exons

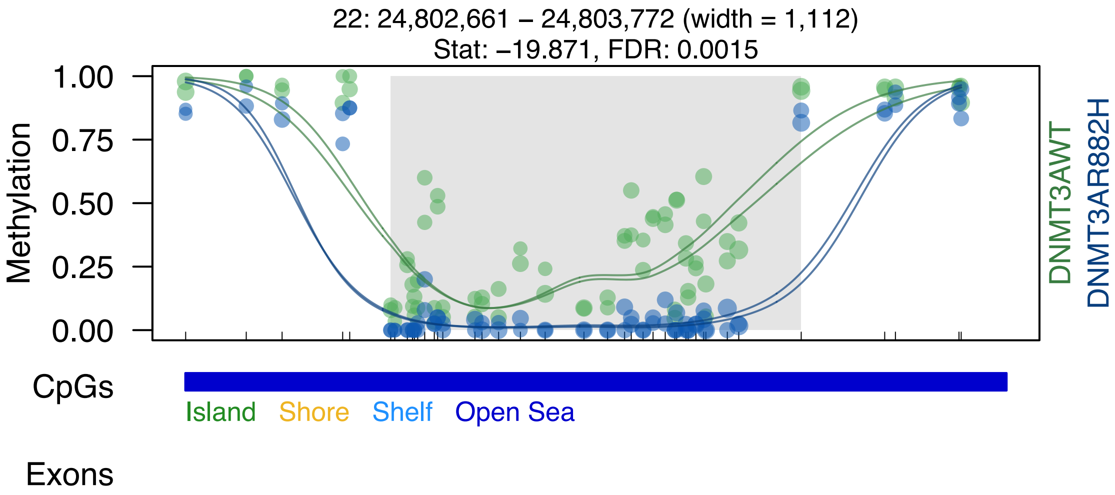
22: 24,802,661 – 24,803,772 (width = 1,112)
Stat: –19.871, FDR: 0.0015
Methylation
1.00
0.75
0.50
0.25
0.00
DNMT3AWT
DNMT3AR882H
CpGs
Island
Shore
Shelf
Open Sea
Exons

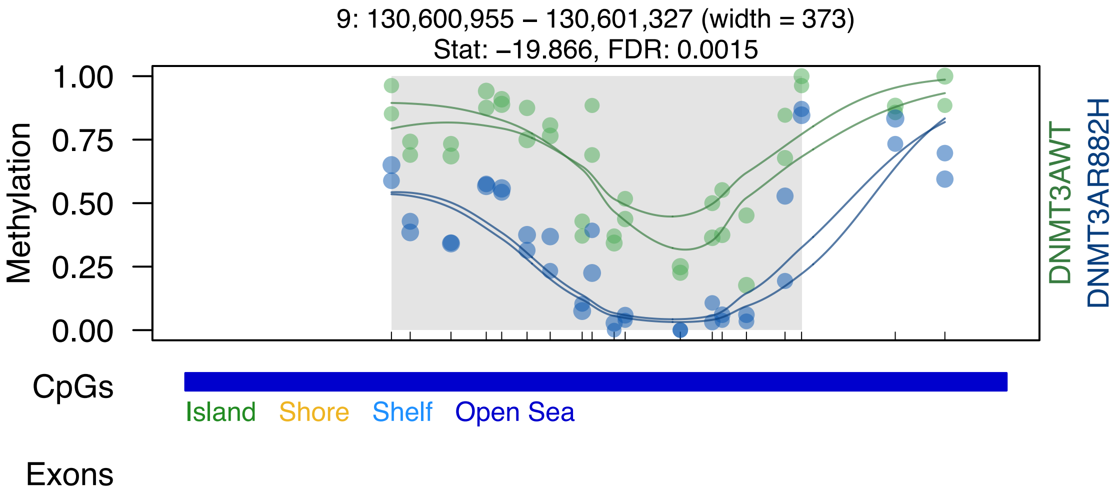

9: 130,600,955 – 130,601,327 (width = 373)
Stat: –19.866, FDR: 0.0015
Methylation
1.00
0.75
0.50
0.25
0.00
DNMT3AWT
DNMT3AR882H
CpGs
Island
Shore
Shelf
Open Sea
Exons

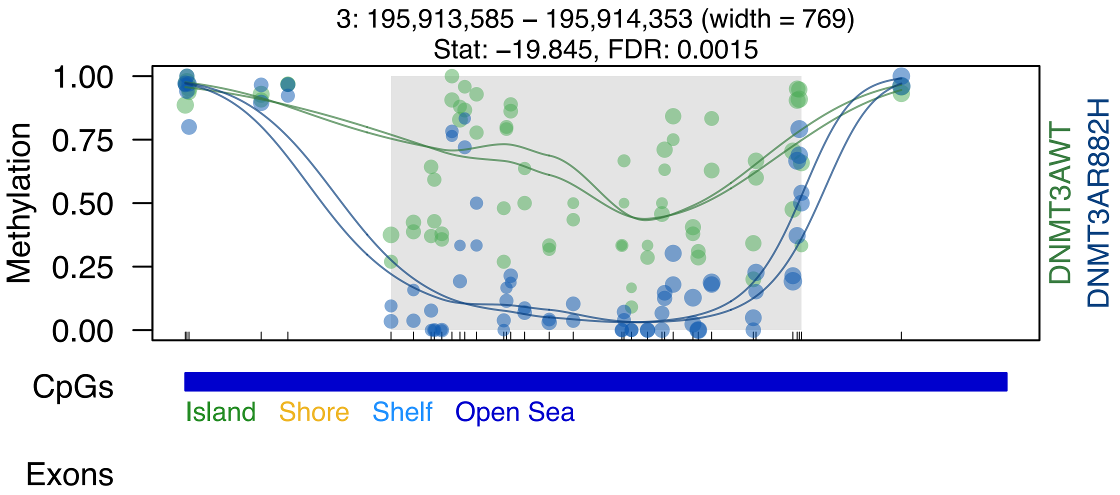
3: 195,913,585 – 195,914,353 (width = 769)
Stat: –19.845, FDR: 0.0015
Methylation
1.00
0.75
0.50
0.25
0.00
DNMT3AWT
DNMT3AR882H
CpGs
Island
Shore
Shelf
Open Sea
Exons

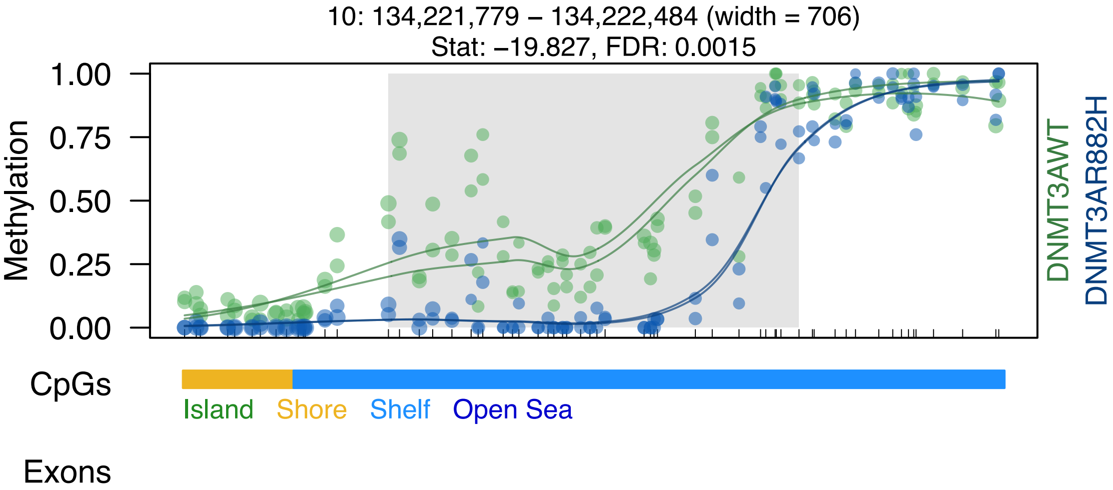

10: 134,221,779 – 134,222,484 (width = 706)
Stat: –19.827, FDR: 0.0015
Methylation
1.00
0.75
0.50
0.25
0.00
DNMT3AWT
DNMT3AR882H
CpGs
Island
Shore
Shelf
Open Sea
Exons

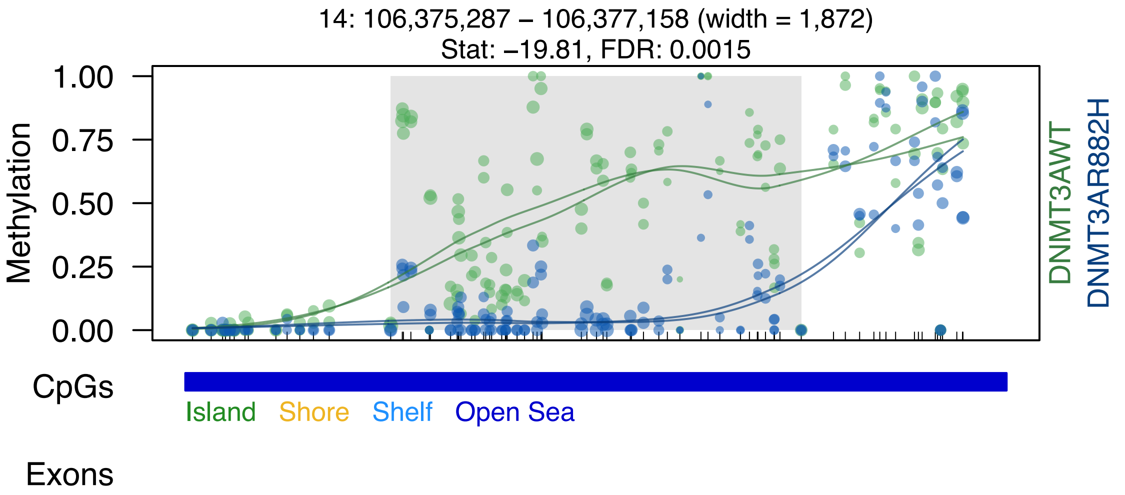
14: 106,375,287 – 106,377,158 (width = 1,872)
Stat: –19.81, FDR: 0.0015
Methylation
1.00
0.75
0.50
0.25
0.00
DNMT3AWT
DNMT3AR882H
CpGs
Island
Shore
Shelf
Open Sea
Exons

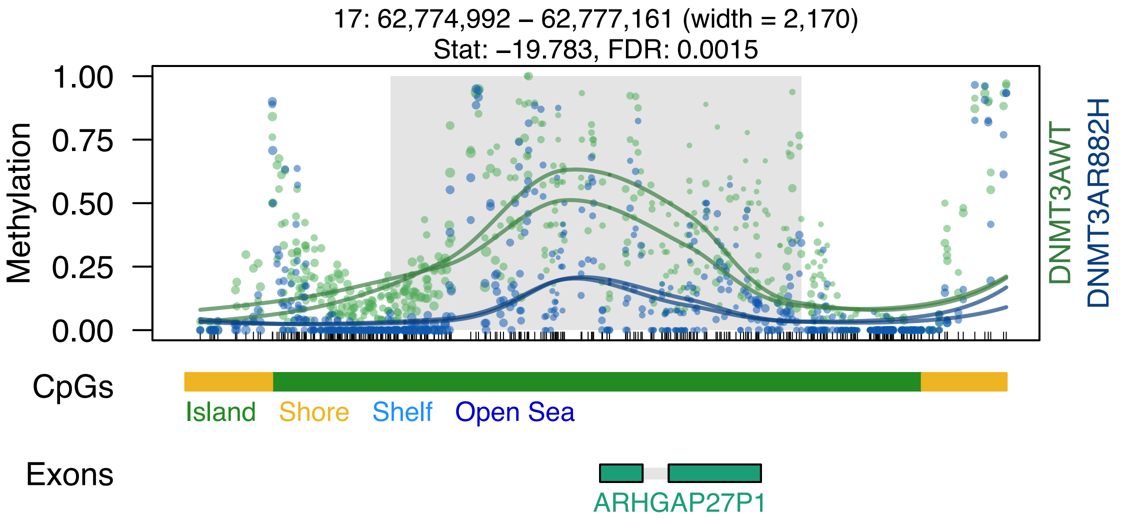
17: 62,774,992 – 62,777,161 (width = 2,170)
Stat: −19.783, FDR: 0.0015
Methylation
1.00
0.75
0.50
0.25
0.00
DNMT3AWT
DNMT3AR882H
CpGs
Island
Shore
Shelf
Open Sea
Exons
ARHGAP27P1

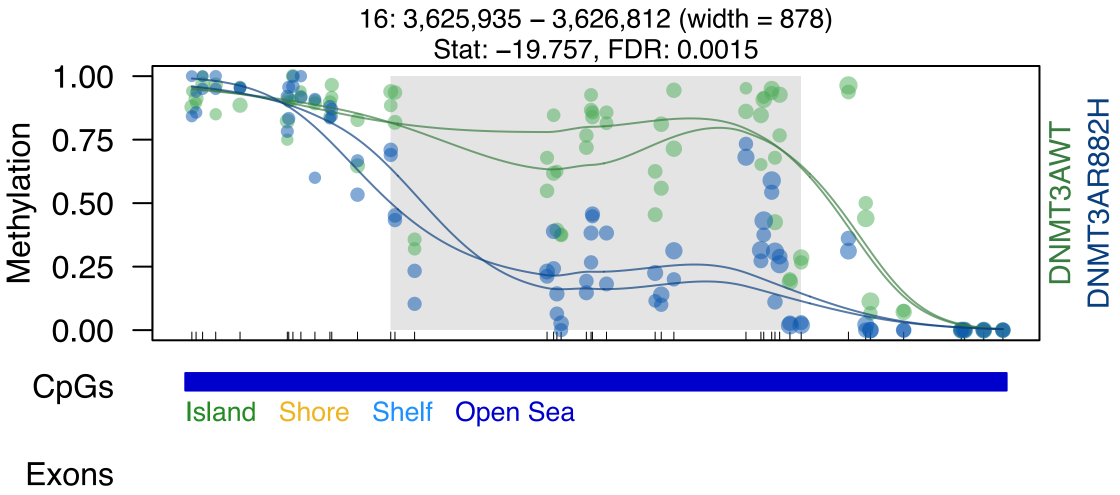
16: 3,625,935 – 3,626,812 (width = 878)
Stat: –19.757, FDR: 0.0015
1.00
0.75
0.50
0.25
0.00
Methylation
DNMT3AWT
DNMT3AR882H
CpGs
Island
Shore
Shelf
Open Sea
Exons

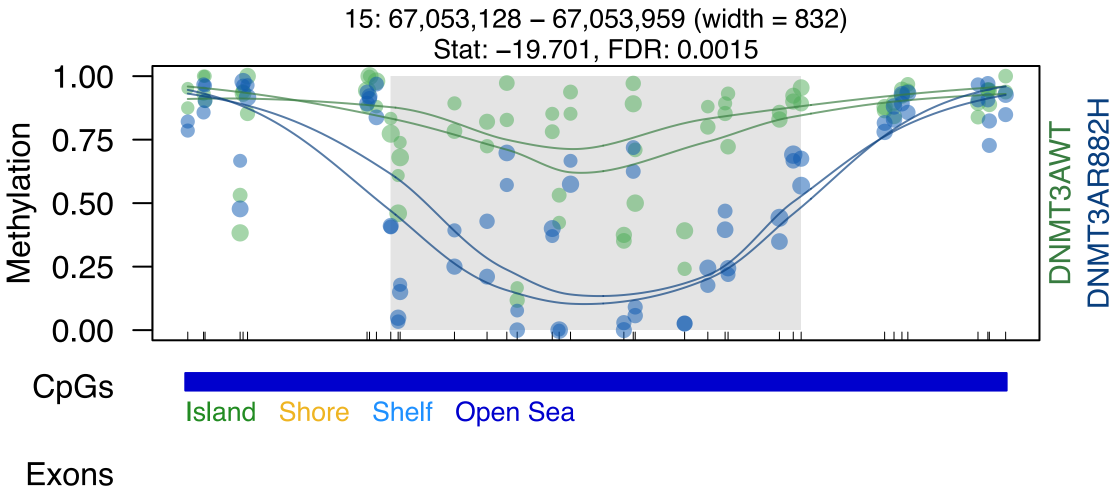
15: 67,053,128 – 67,053,959 (width = 832)
Stat: –19.701, FDR: 0.0015
1.00
0.75
0.50
0.25
0.00
Methylation
DNMT3AWT
DNMT3AR882H
CpGs
Island
Shore
Shelf
Open Sea
Exons

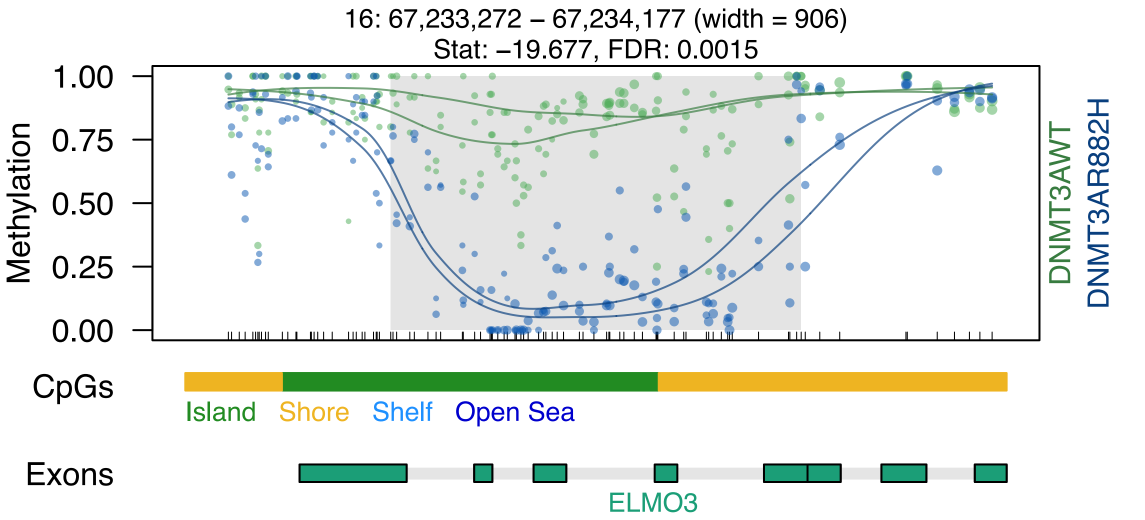
16: 67,233,272 – 67,234,177 (width = 906)
Stat: –19.677, FDR: 0.0015
Methylation
1.00
0.75
0.50
0.25
0.00
DNMT3AWT
DNMT3AR882H
CpGs
Island
Shore
Shelf
Open Sea
Exons
ELMO3

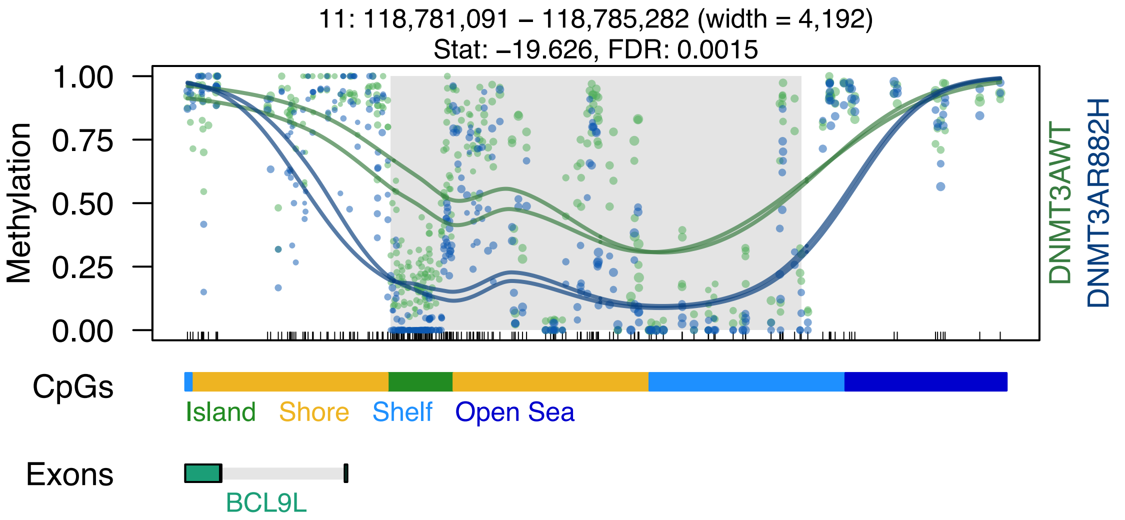

11: 118,781,091 – 118,785,282 (width = 4,192)
Stat: –19.626, FDR: 0.0015
1.00
0.75
0.50
0.25
0.00
Methylation
DNMT3AWT
DNMT3AR882H
CpGs
Island
Shore
Shelf
Open Sea
Exons
BCL9L

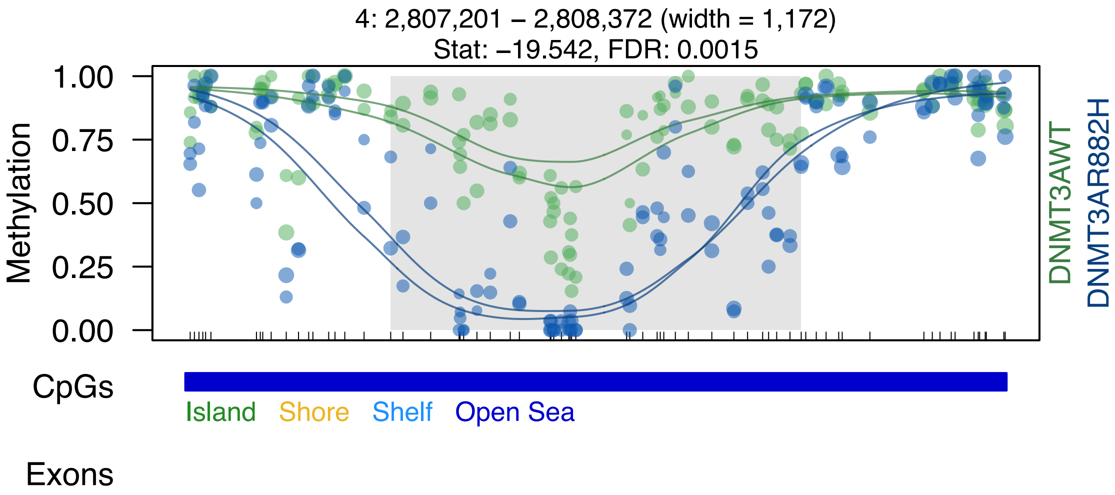

4: 2,807,201 – 2,808,372 (width = 1,172)
Stat: –19.542, FDR: 0.0015
Methylation
1.00
0.75
0.50
0.25
0.00
DNMT3AWT
DNMT3AR882H
CpGs
Island
Shore
Shelf
Open Sea
Exons

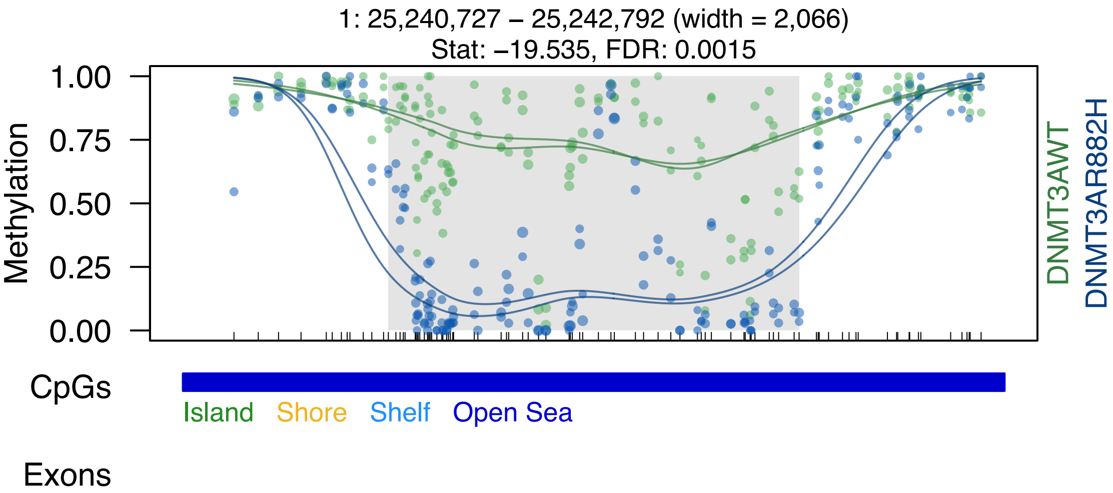
1: 25,240,727 – 25,242,792 (width = 2,066)
Stat: –19.535, FDR: 0.0015
Methylation
1.00
0.75
0.50
0.25
0.00
DNMT3AWT
DNMT3AR882H
CpGs
Island
Shore
Shelf
Open Sea
Exons

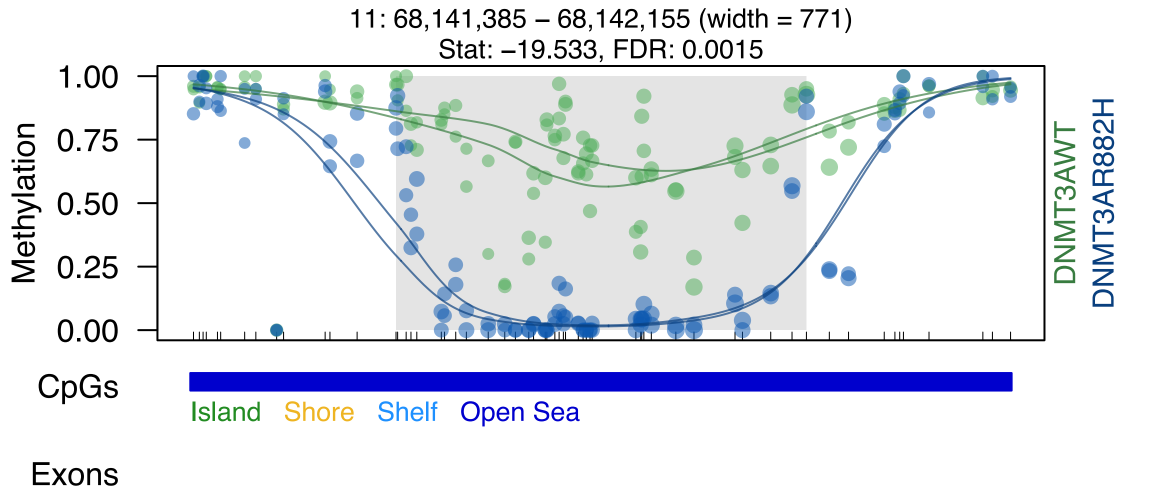

11: 68,141,385 – 68,142,155 (width = 771)
Stat: –19.533, FDR: 0.0015
Methylation
1.00
0.75
0.50
0.25
0.00
DNMT3AWT
DNMT3AR882H
CpGs
Island
Shore
Shelf
Open Sea
Exons

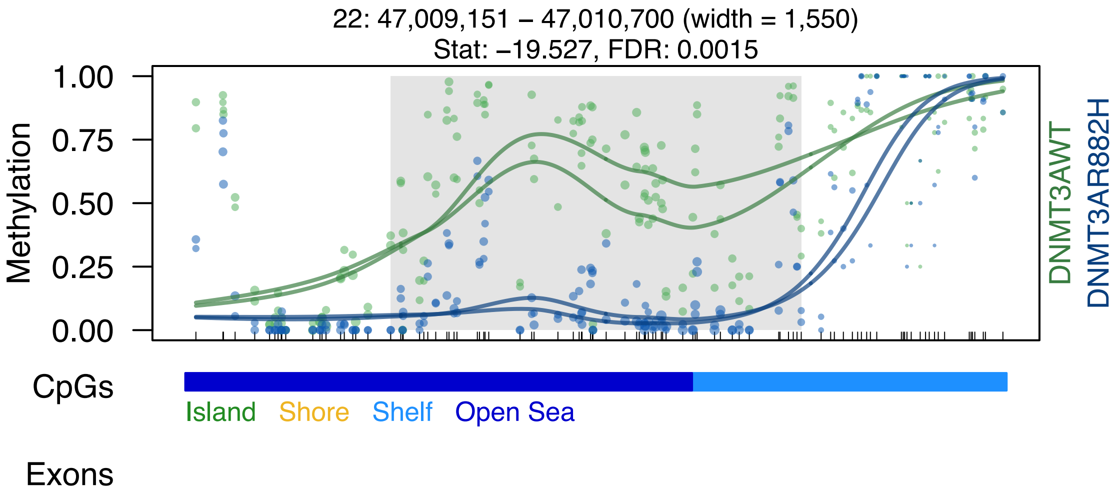

22: 47,009,151 – 47,010,700 (width = 1,550)
Stat: –19.527, FDR: 0.0015
Methylation
1.00
0.75
0.50
0.25
0.00
DNMT3AWT
DNMT3AR882H
CpGs
Island
Shore
Shelf
Open Sea
Exons

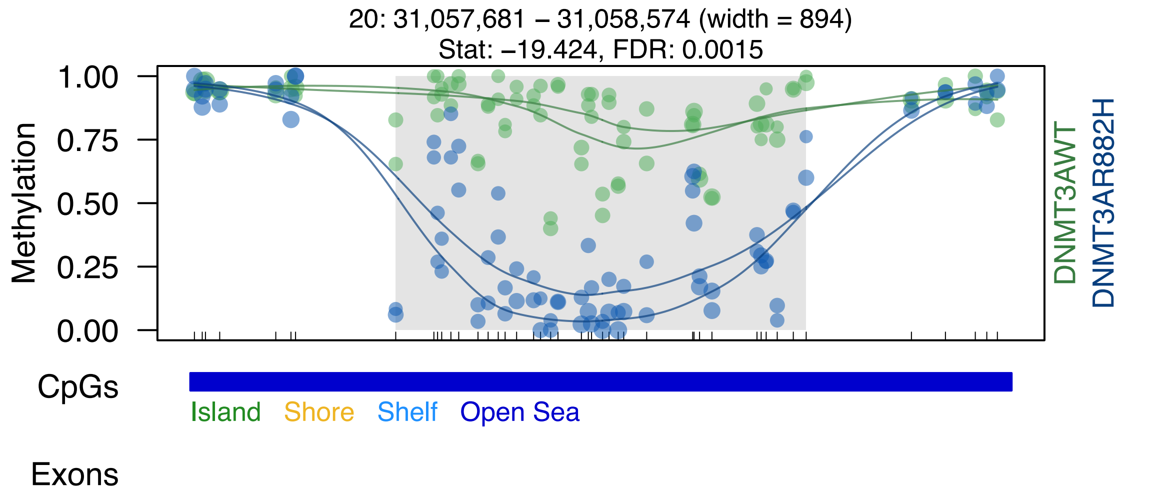
20: 31,057,681 – 31,058,574 (width = 894)
Stat: –19.424, FDR: 0.0015
Methylation
1.00
0.75
0.50
0.25
0.00
DNMT3AWT
DNMT3AR882H
CpGs
Island
Shore
Shelf
Open Sea
Exons

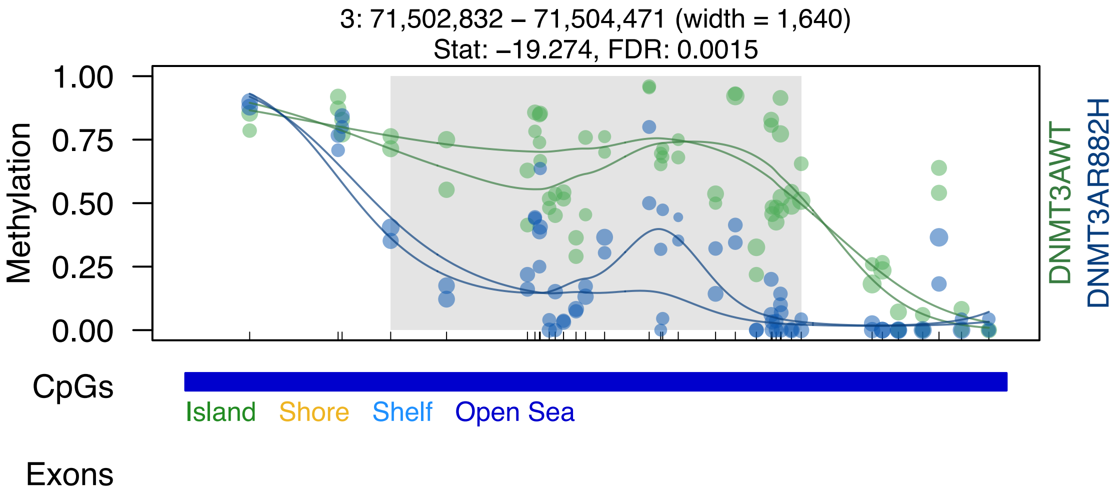
3: 71,502,832 – 71,504,471 (width = 1,640)
Stat: –19.274, FDR: 0.0015
Methylation
1.00
0.75
0.50
0.25
0.00
DNMT3AWT
DNMT3AR882H
CpGs
Island
Shore
Shelf
Open Sea
Exons

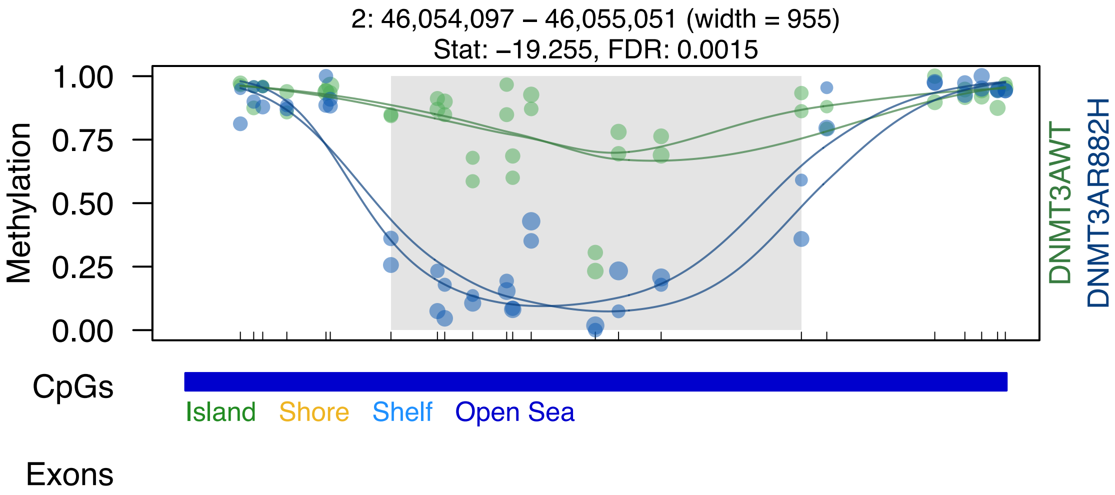
2: 46,054,097 – 46,055,051 (width = 955)
Stat: –19.255, FDR: 0.0015
Methylation
1.00
0.75
0.50
0.25
0.00
DNMT3AWT
DNMT3AR882H
CpGs
Island
Shore
Shelf
Open Sea
Exons

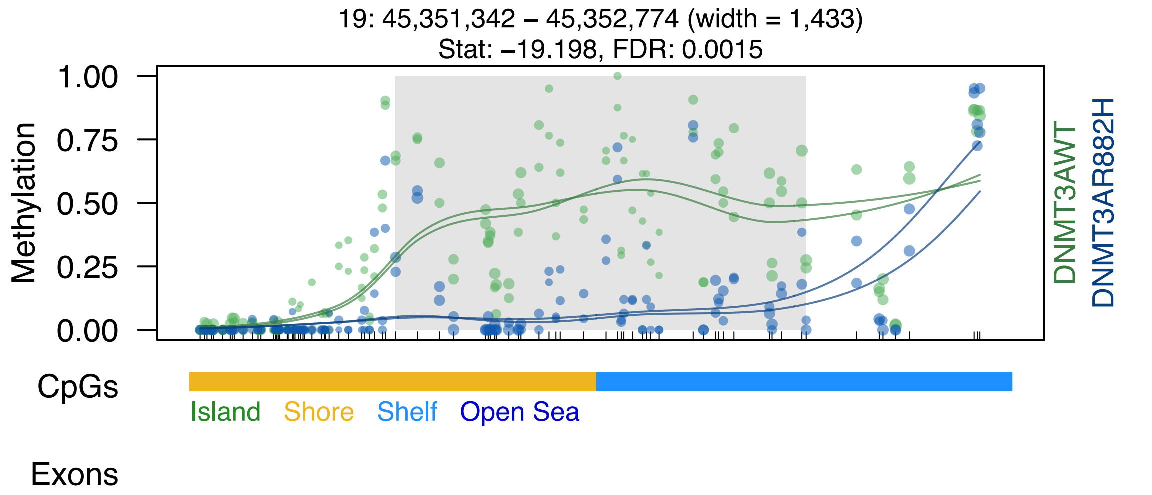
19: 45,351,342 – 45,352,774 (width = 1,433)
Stat: –19.198, FDR: 0.0015
Methylation
1.00
0.75
0.50
0.25
0.00
DNMT3AWT
DNMT3AR882H
CpGs
Island
Shore
Shelf
Open Sea
Exons

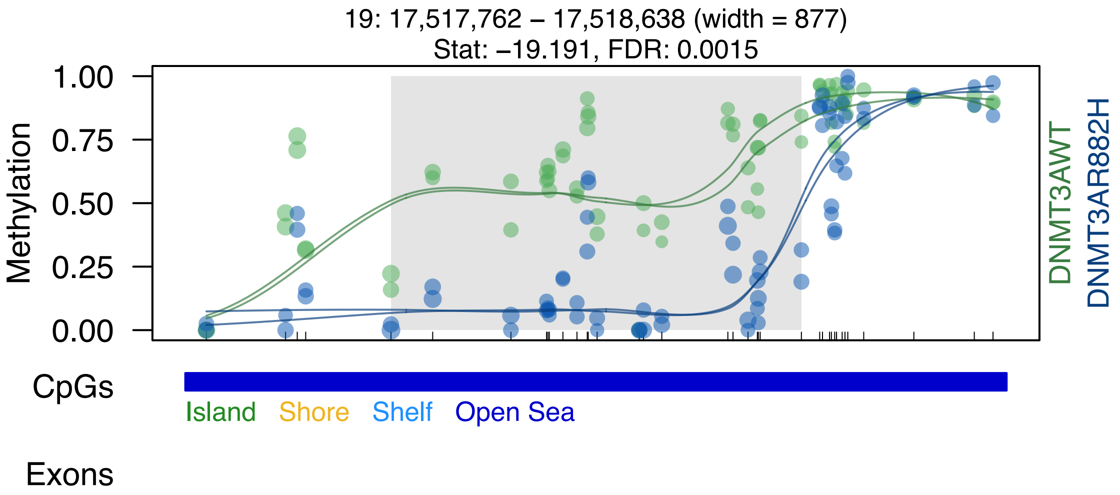

19: 17,517,762 – 17,518,638 (width = 877)
Stat: –19.191, FDR: 0.0015
Methylation
1.00
0.75
0.50
0.25
0.00
DNMT3AWT
DNMT3AR882H
CpGs
Island
Shore
Shelf
Open Sea
Exons

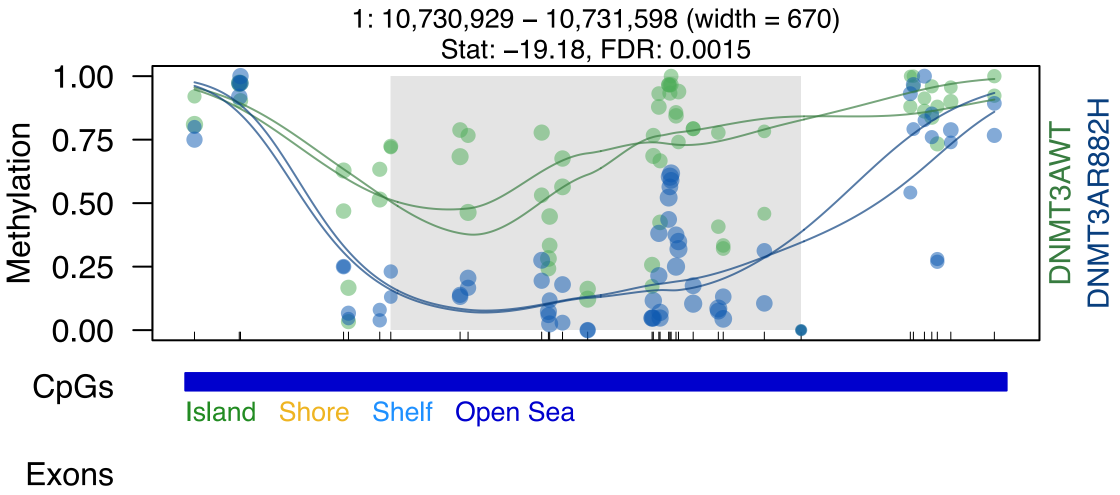
1: 10,730,929 – 10,731,598 (width = 670)
Stat: –19.18, FDR: 0.0015
Methylation
1.00
0.75
0.50
0.25
0.00
DNMT3AWT
DNMT3AR882H
CpGs
Island
Shore
Shelf
Open Sea
Exons

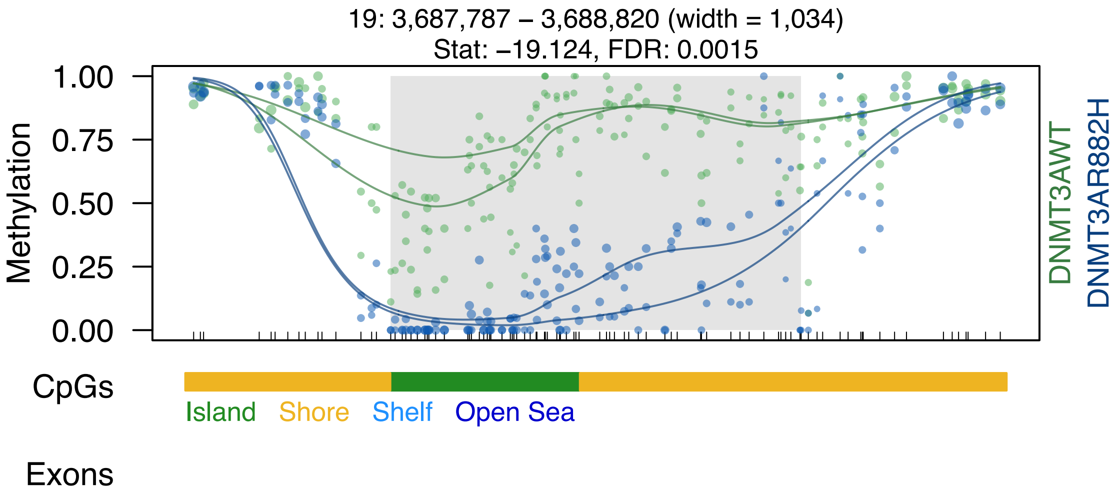

19: 3,687,787 – 3,688,820 (width = 1,034)
Stat: –19.124, FDR: 0.0015
Methylation
1.00
0.75
0.50
0.25
0.00
DNMT3AWT
DNMT3AR882H
CpGs
Island
Shore
Shelf
Open Sea
Exons

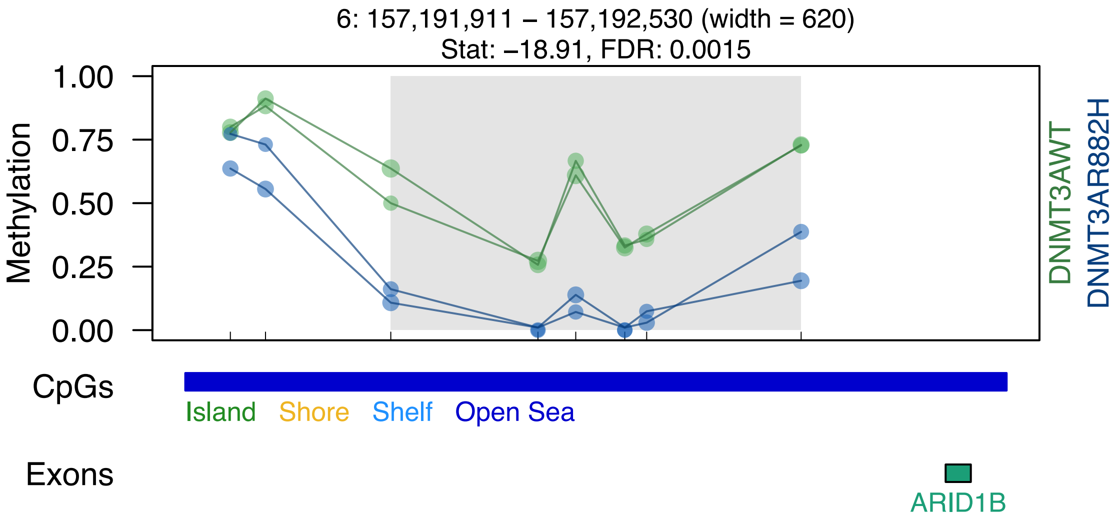
6: 157,191,911 – 157,192,530 (width = 620)
Stat: –18.91, FDR: 0.0015
Methylation
1.00
0.75
0.50
0.25
0.00
DNMT3AWT
DNMT3AR882H
CpGs
Island
Shore
Shelf
Open Sea
Exons
ARID1B

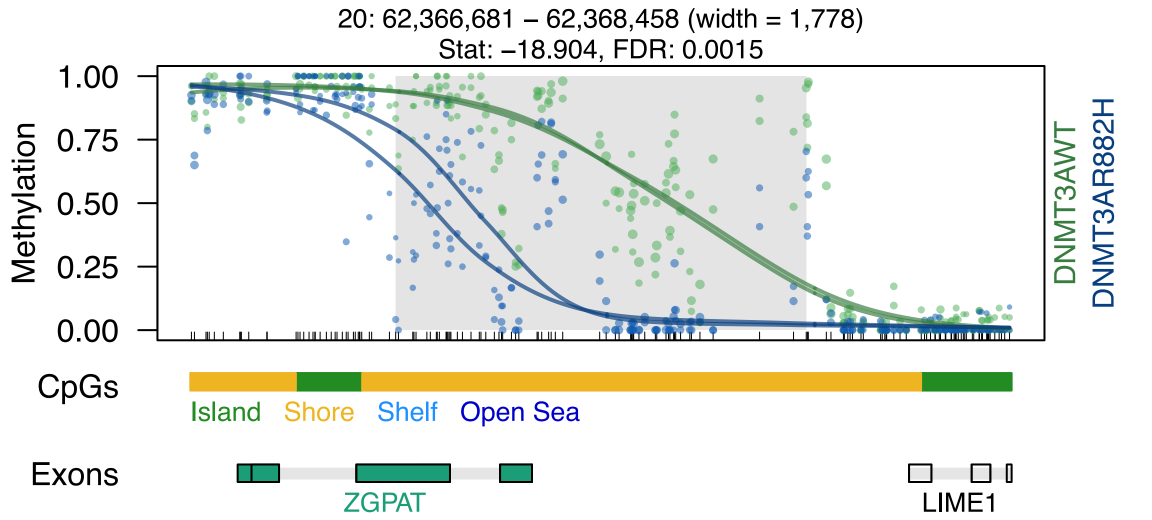
20: 62,366,681 – 62,368,458 (width = 1,778)
Stat: –18.904, FDR: 0.0015
1.00
0.75
0.50
0.25
0.00
Methylation
DNMT3AWT
DNMT3AR882H
CpGs
Island
Shore
Shelf
Open Sea
Exons
ZGPAT
LIME1

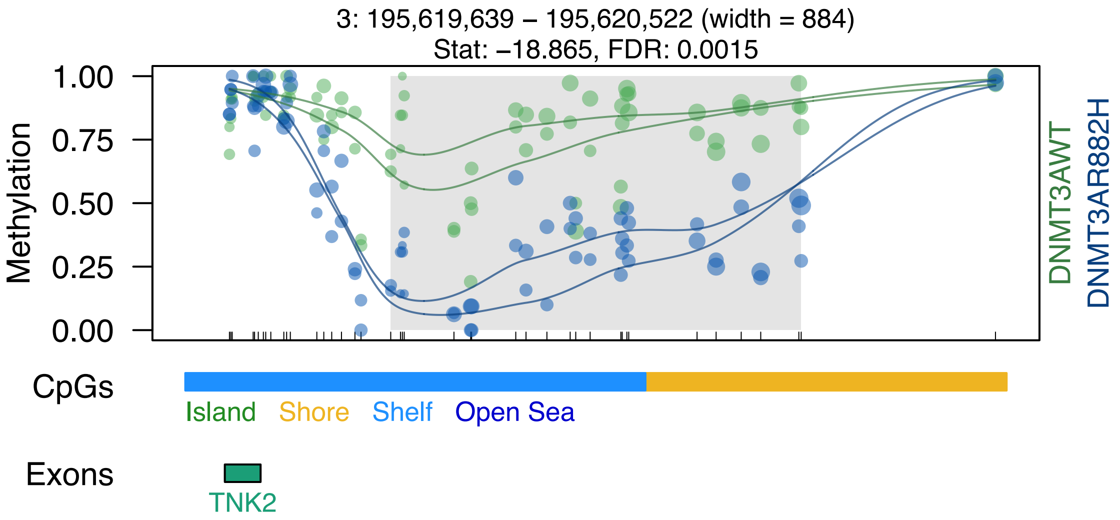
3: 195,619,639 – 195,620,522 (width = 884)
Stat: –18.865, FDR: 0.0015
Methylation
1.00
0.75
0.50
0.25
0.00
DNMT3AWT
DNMT3AR882H
CpGs
Island
Shore
Shelf
Open Sea
Exons
TNK2

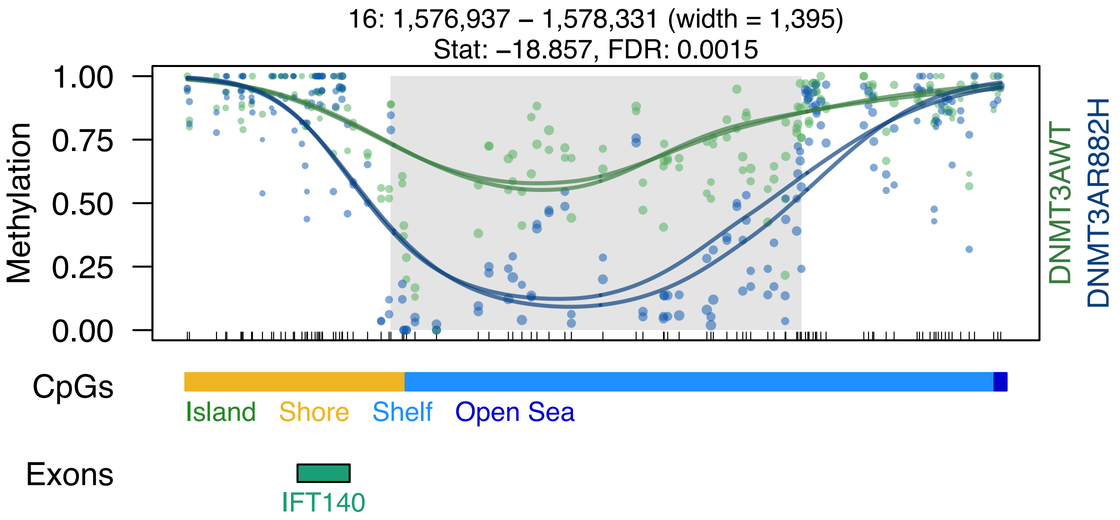

16: 1,576,937 – 1,578,331 (width = 1,395)
Stat: –18.857, FDR: 0.0015
Methylation
1.00
0.75
0.50
0.25
0.00
DNMT3AWT
DNMT3AR882H
CpGs
Island
Shore
Shelf
Open Sea
Exons
IFT140

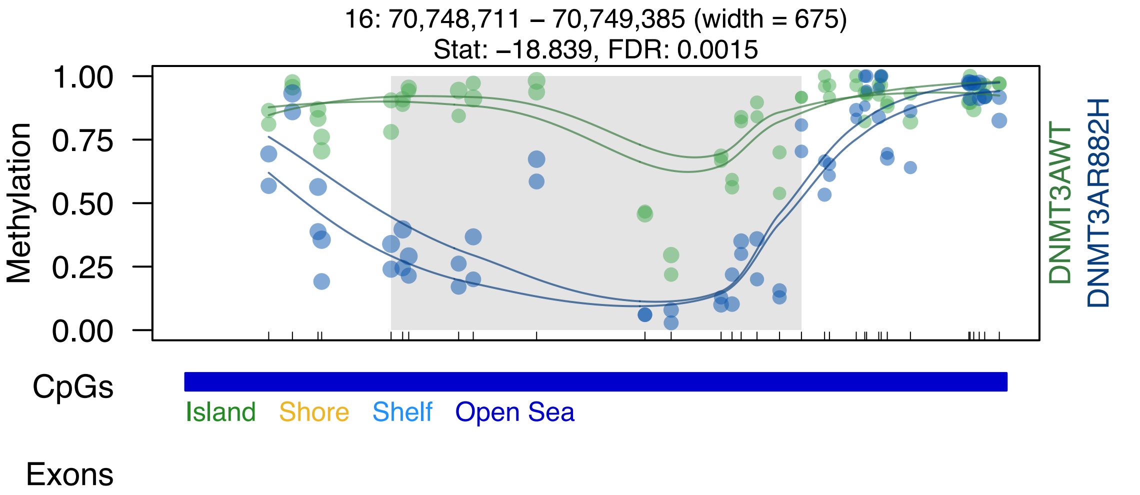

16: 70,748,711 – 70,749,385 (width = 675)
Stat: –18.839, FDR: 0.0015
Methylation
1.00
0.75
0.50
0.25
0.00
DNMT3AWT
DNMT3AR882H
CpGs
Island
Shore
Shelf
Open Sea
Exons

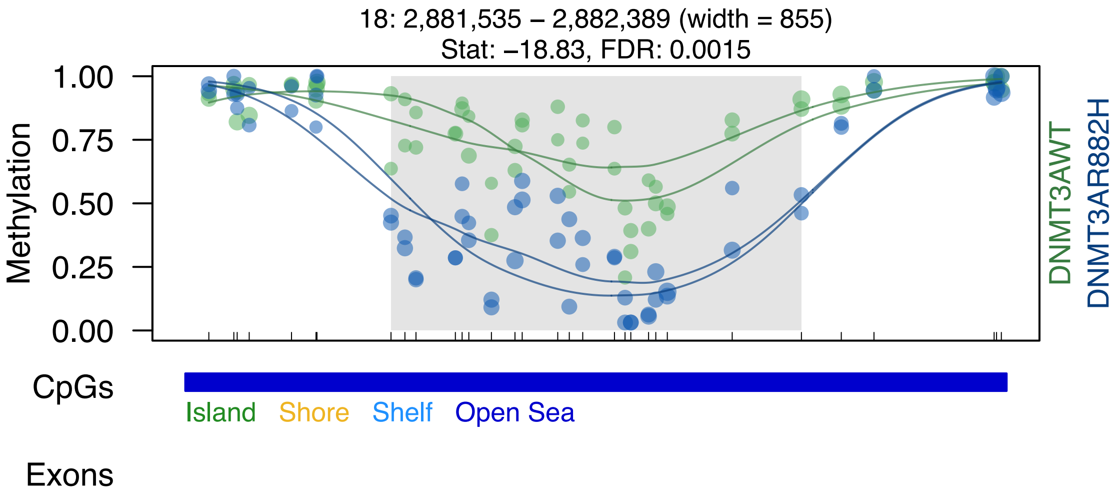
18: 2,881,535 – 2,882,389 (width = 855)
Stat: –18.83, FDR: 0.0015
Methylation
1.00
0.75
0.50
0.25
0.00
DNMT3AWT
DNMT3AR882H
CpGs
Island
Shore
Shelf
Open Sea
Exons

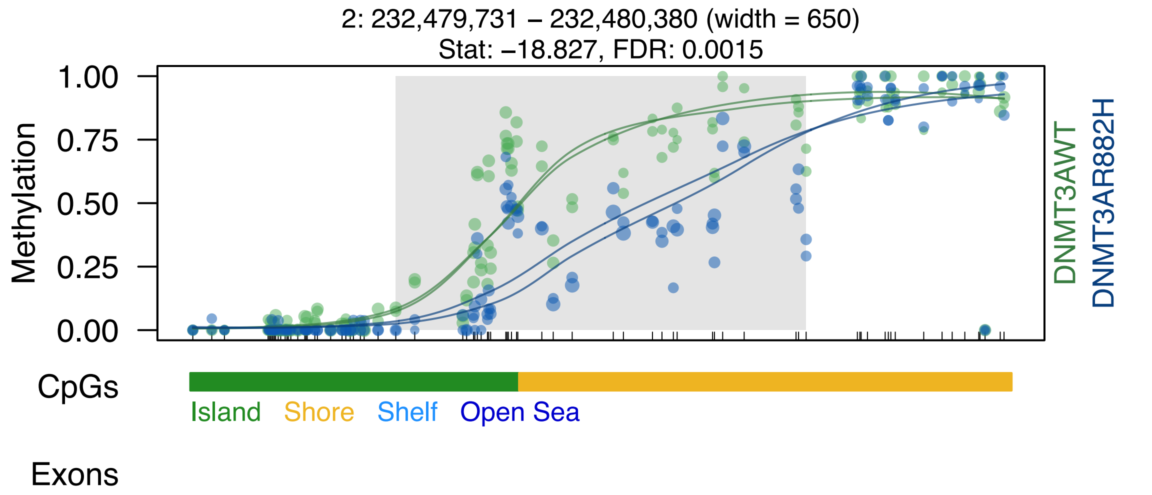
2: 232,479,731 – 232,480,380 (width = 650)
Stat: –18.827, FDR: 0.0015
Methylation
1.00
0.75
0.50
0.25
0.00
DNMT3AWT
DNMT3AR882H
CpGs
Island
Shore
Shelf
Open Sea
Exons

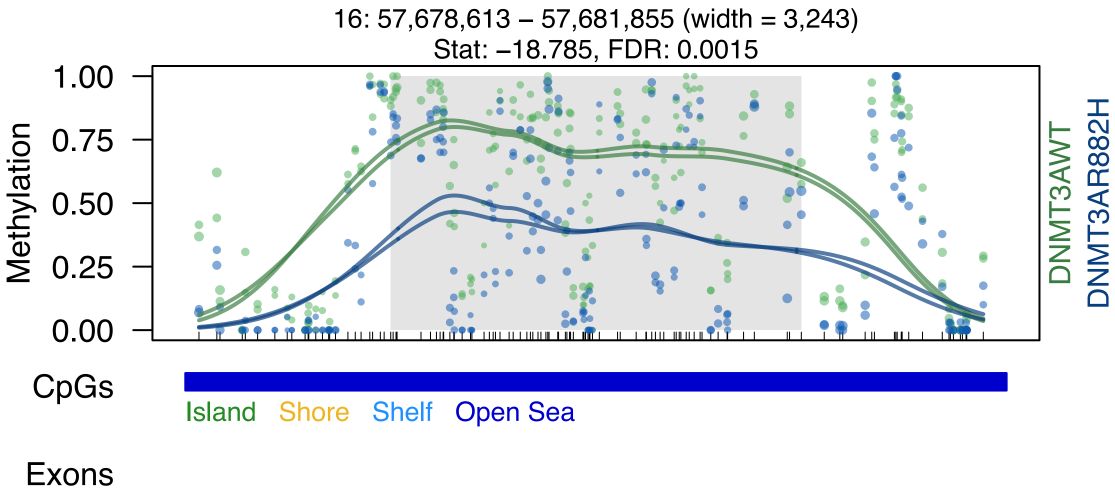

16: 57,678,613 – 57,681,855 (width = 3,243)
Stat: –18.785, FDR: 0.0015
Methylation
1.00
0.75
0.50
0.25
0.00
DNMT3AWT
DNMT3AR882H
CpGs
Island
Shore
Shelf
Open Sea
Exons

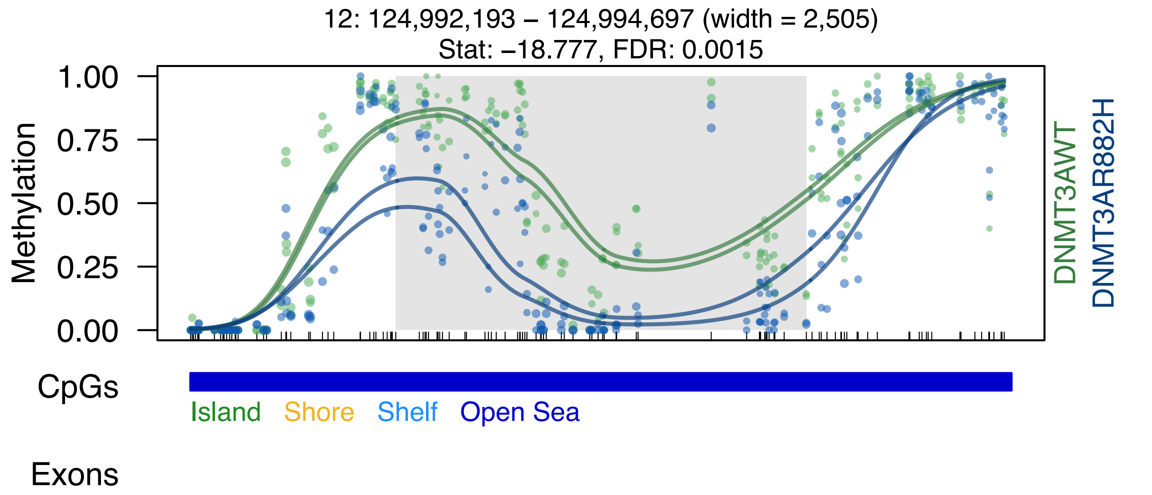

12: 124,992,193 – 124,994,697 (width = 2,505)
Stat: −18.777, FDR: 0.0015
Methylation
1.00
0.75
0.50
0.25
0.00
DNMT3AWT
DNMT3AR882H
CpGs
Island
Shore
Shelf
Open Sea
Exons

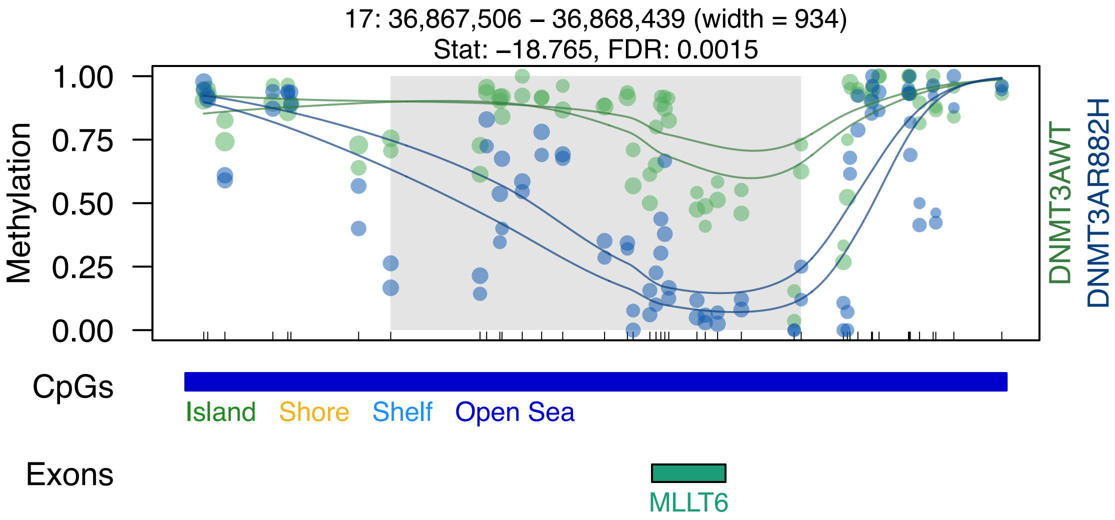

17: 36,867,506 – 36,868,439 (width = 934)
Stat: –18.765, FDR: 0.0015
Methylation
1.00
0.75
0.50
0.25
0.00
DNMT3AWT
DNMT3AR882H
CpGs
Island
Shore
Shelf
Open Sea
Exons
MLLT6

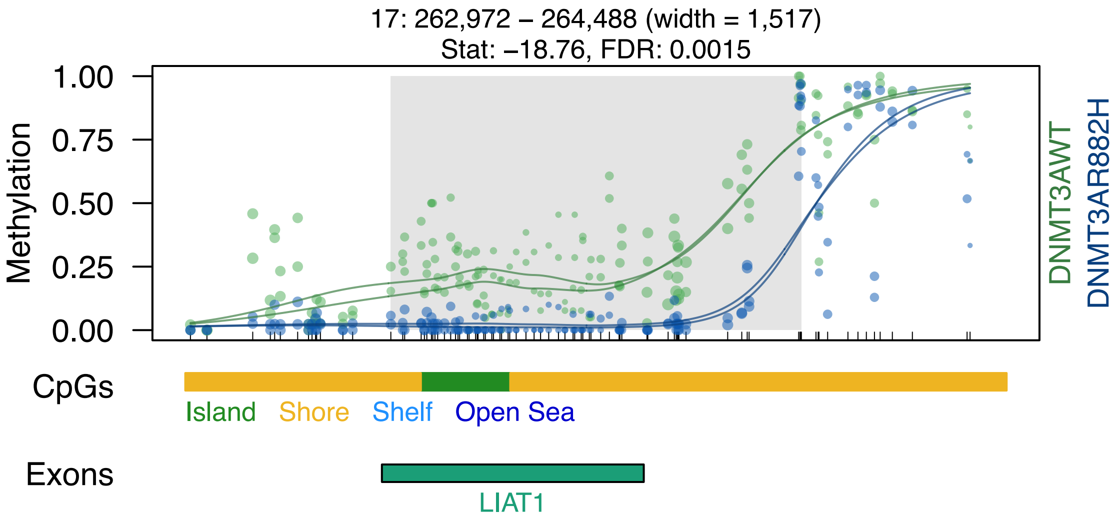

17: 262,972 – 264,488 (width = 1,517)
Stat: –18.76, FDR: 0.0015
1.00
0.75
0.50
0.25
0.00
Methylation
DNMT3AWT
DNMT3AR882H
CpGs
Island
Shore
Shelf
Open Sea
Exons
LIAT1

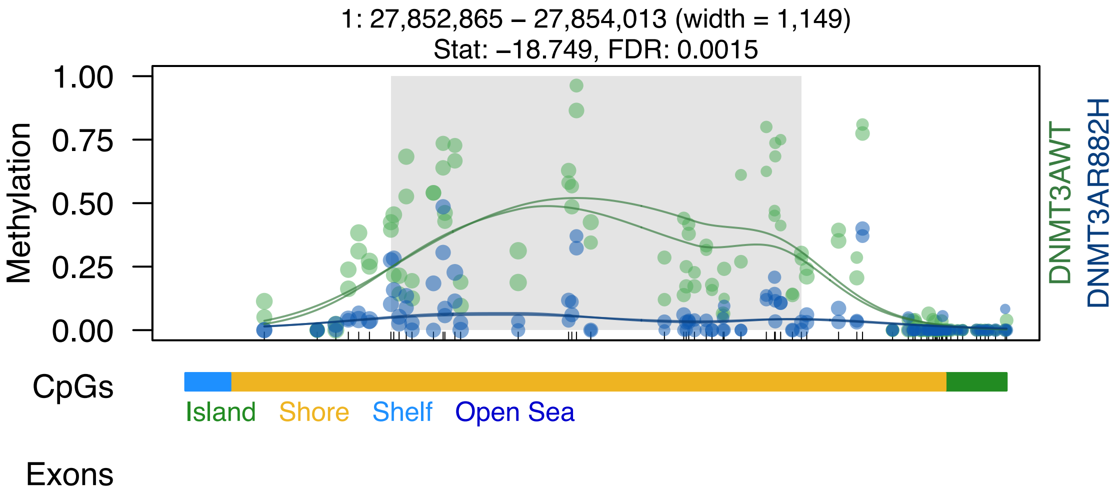
1: 27,852,865 – 27,854,013 (width = 1,149)
Stat: −18.749, FDR: 0.0015
1.00
0.75
0.50
0.25
0.00
Methylation
DNMT3AWT
DNMT3AR882H
CpGs
Island
Shore
Shelf
Open Sea
Exons

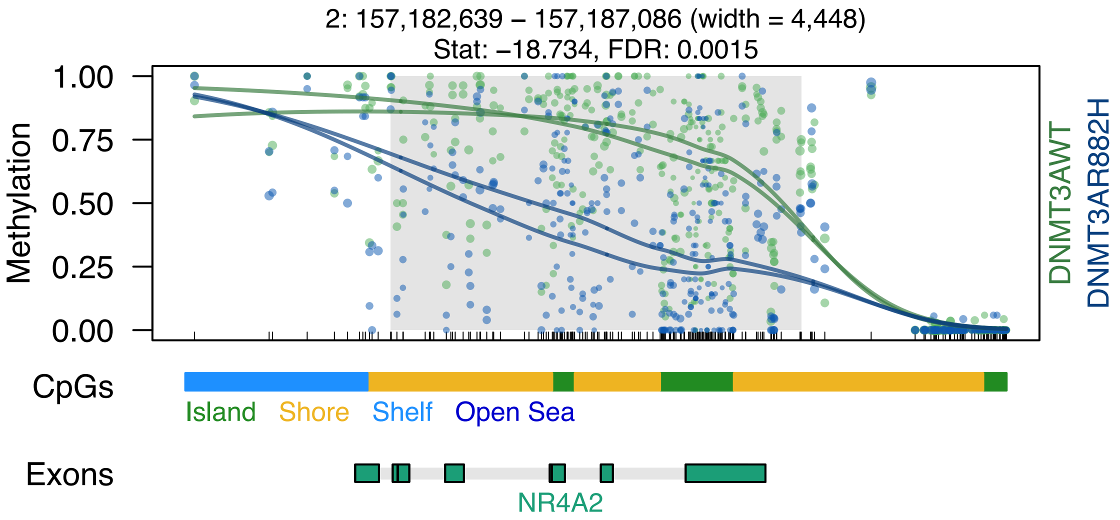
2: 157,182,639 – 157,187,086 (width = 4,448)
Stat: –18.734, FDR: 0.0015
Methylation
1.00
0.75
0.50
0.25
0.00
DNMT3AWT
DNMT3AR882H
CpGs
Island
Shore
Shelf
Open Sea
Exons
NR4A2

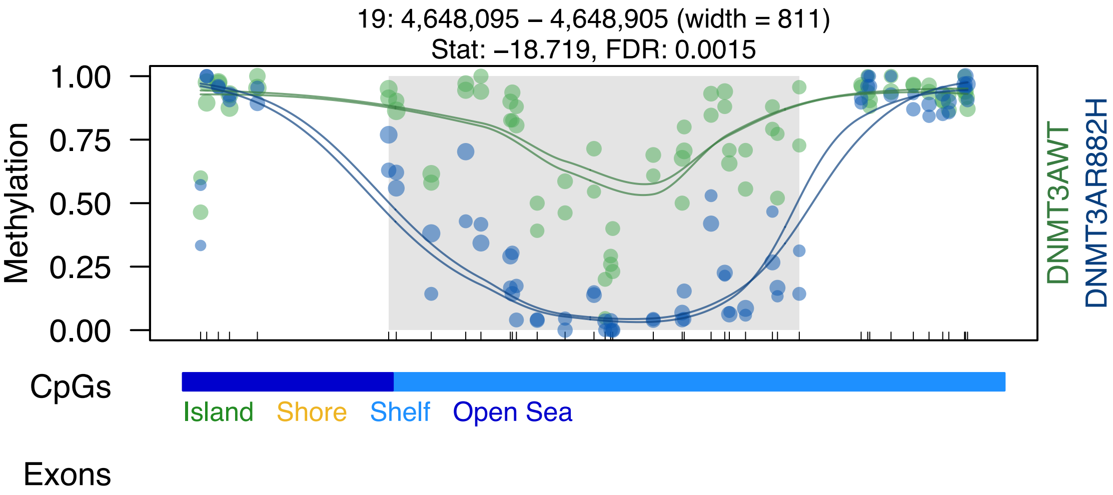

19: 4,648,095 – 4,648,905 (width = 811)
Stat: –18.719, FDR: 0.0015
Methylation
1.00
0.75
0.50
0.25
0.00
DNMT3AWT
DNMT3AR882H
CpGs
Island
Shore
Shelf
Open Sea
Exons
