## Supplemental Data S1-S7 for "DNMT3A^R882H^ Is Not Required for Disease Maintenance in Primary Human AML, but Is Associated With Increased Leukemia Stem Cell Frequency": DataS3_dmrSeq_SU575_top100DMRs.pdf

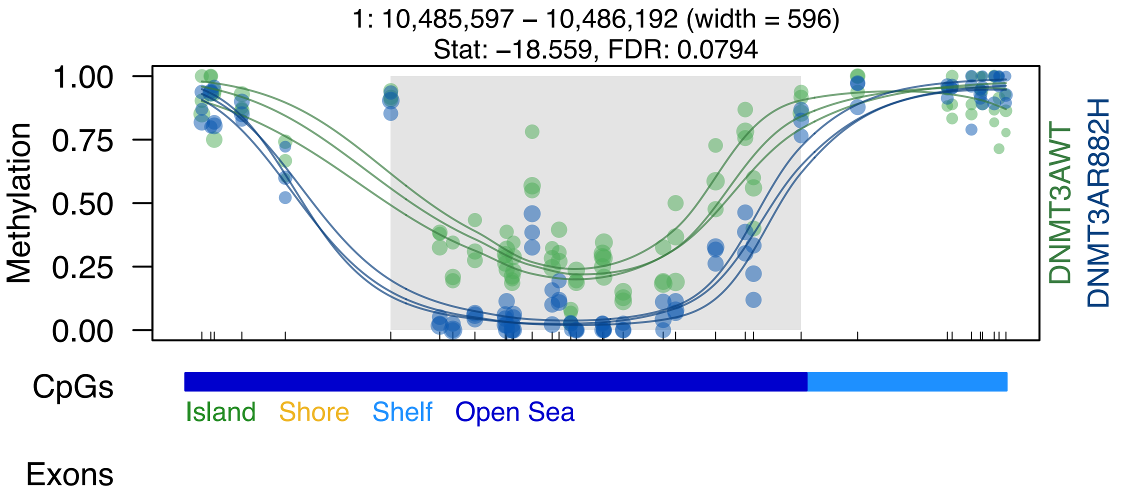

1: 10,485,597 – 10,486,192 (width = 596)
Stat: –18.559, FDR: 0.0794
Methylation
1.00
0.75
0.50
0.25
0.00
DNMT3AWT
DNMT3AR882H
CpGs
Island
Shore
Shelf
Open Sea
Exons

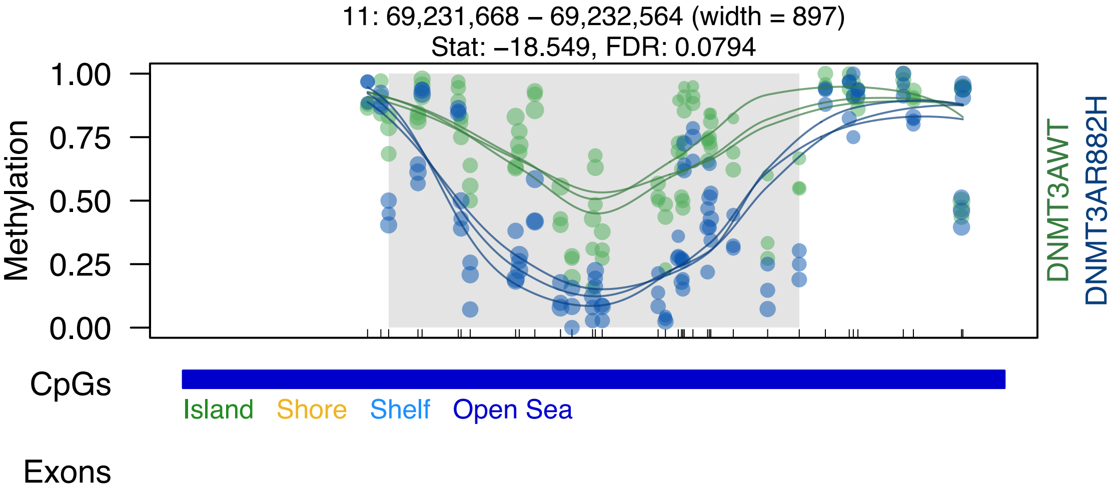
11: 69,231,668 – 69,232,564 (width = 897)
Stat: −18.549, FDR: 0.0794
Methylation
1.00
0.75
0.50
0.25
0.00
DNMT3AWT
DNMT3AR882H
CpGs
Island
Shore
Shelf
Open Sea
Exons

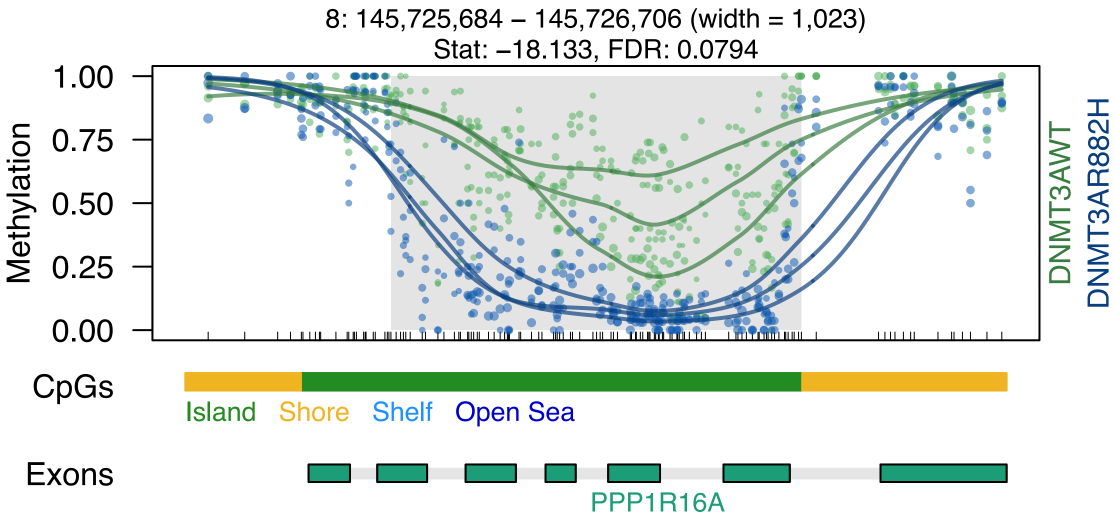
8: 145,725,684 – 145,726,706 (width = 1,023)
Stat: –18.133, FDR: 0.0794
1.00
0.75
0.50
0.25
0.00
Methylation
DNMT3AWT
DNMT3AR882H
CpGs
Island
Shore
Shelf
Open Sea
Exons
PPP1R16A

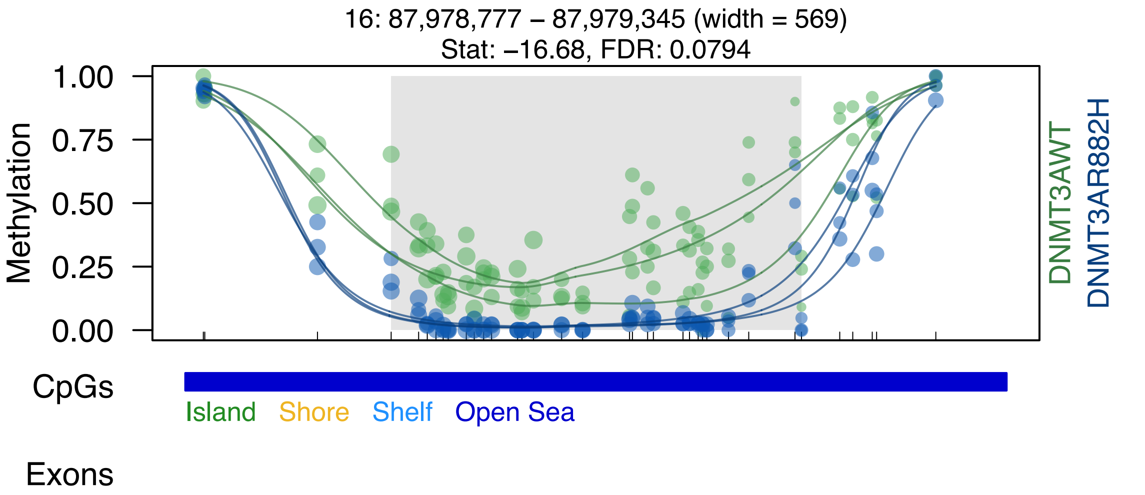

16: 87,978,777 – 87,979,345 (width = 569)
Stat: –16.68, FDR: 0.0794
Methylation
1.00
0.75
0.50
0.25
0.00
DNMT3AWT
DNMT3AR882H
CpGs
Island
Shore
Shelf
Open Sea
Exons

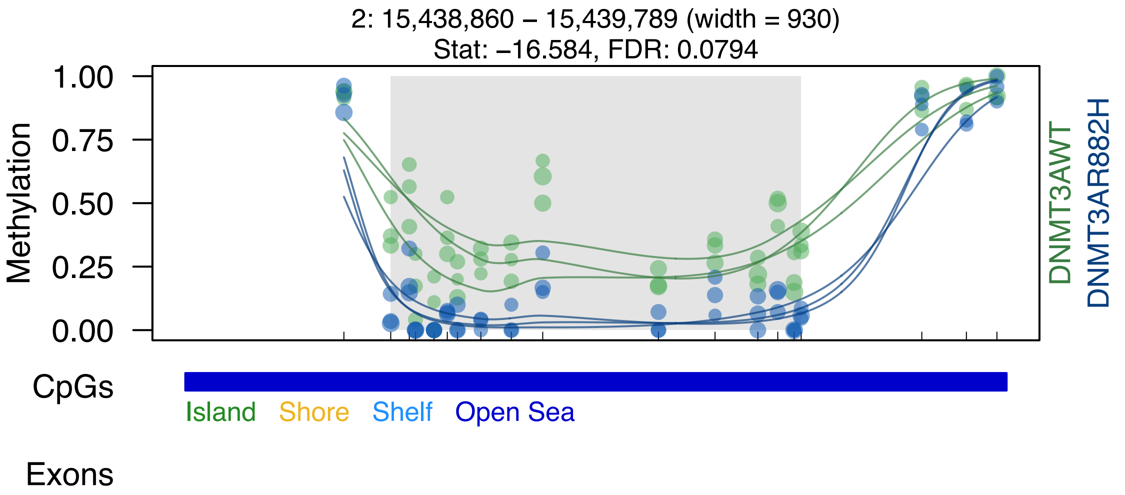
2: 15,438,860 – 15,439,789 (width = 930)
Stat: –16.584, FDR: 0.0794
Methylation
1.00
0.75
0.50
0.25
0.00
DNMT3AWT
DNMT3AR882H
CpGs
Island
Shore
Shelf
Open Sea
Exons

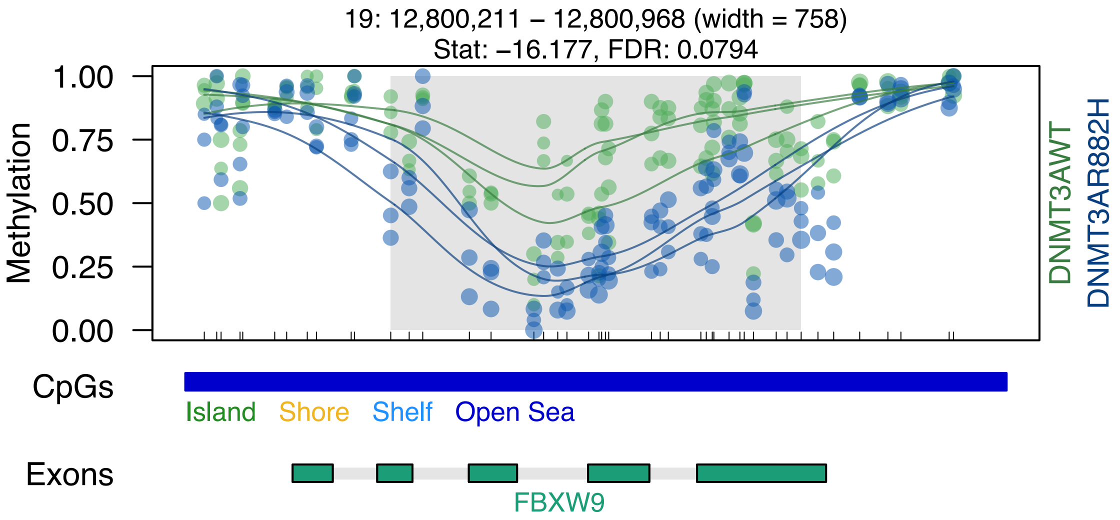
19: 12,800,211 – 12,800,968 (width = 758)
Stat: –16.177, FDR: 0.0794
Methylation
1.00
0.75
0.50
0.25
0.00
DNMT3AWT
DNMT3AR882H
CpGs
Island
Shore
Shelf
Open Sea
Exons
FBXW9

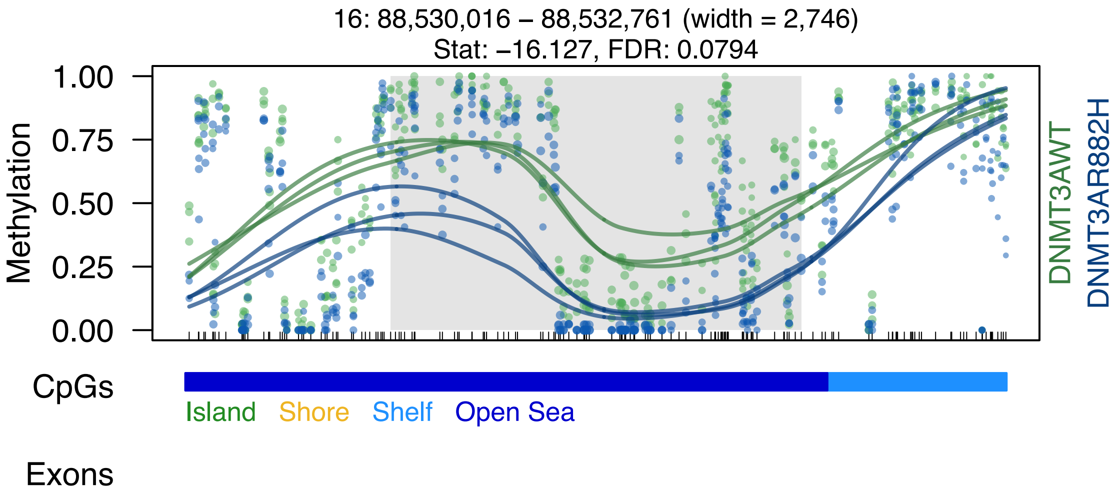
16: 88,530,016 – 88,532,761 (width = 2,746)
Stat: –16.127, FDR: 0.0794
1.00
0.75
0.50
0.25
0.00
Methylation
DNMT3AWT
DNMT3AR882H
CpGs
Island
Shore
Shelf
Open Sea
Exons

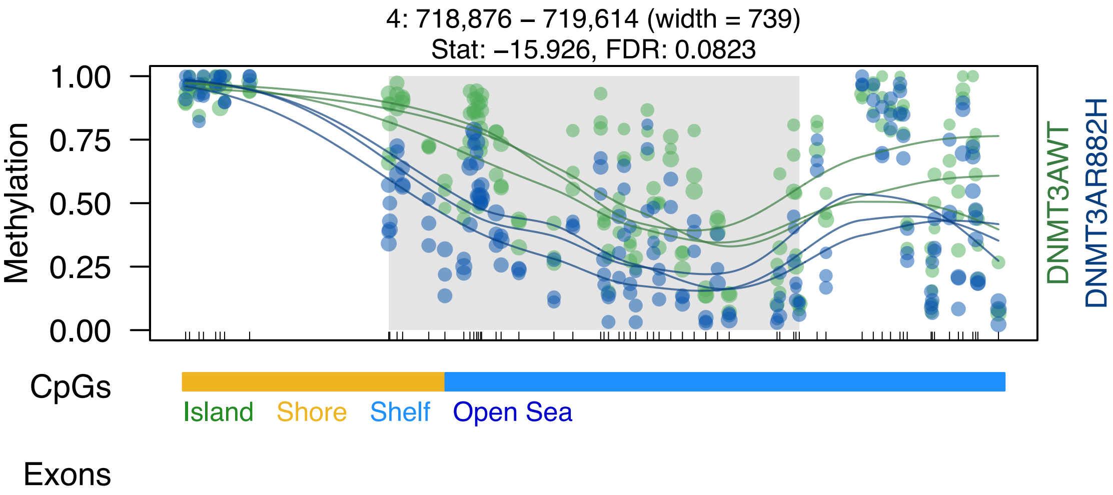
4: 718,876 – 719,614 (width = 739)
Stat: –15.926, FDR: 0.0823
1.00
0.75
0.50
0.25
0.00
Methylation
DNMT3AWT
DNMT3AR882H
CpGs
Island
Shore
Shelf
Open Sea
Exons

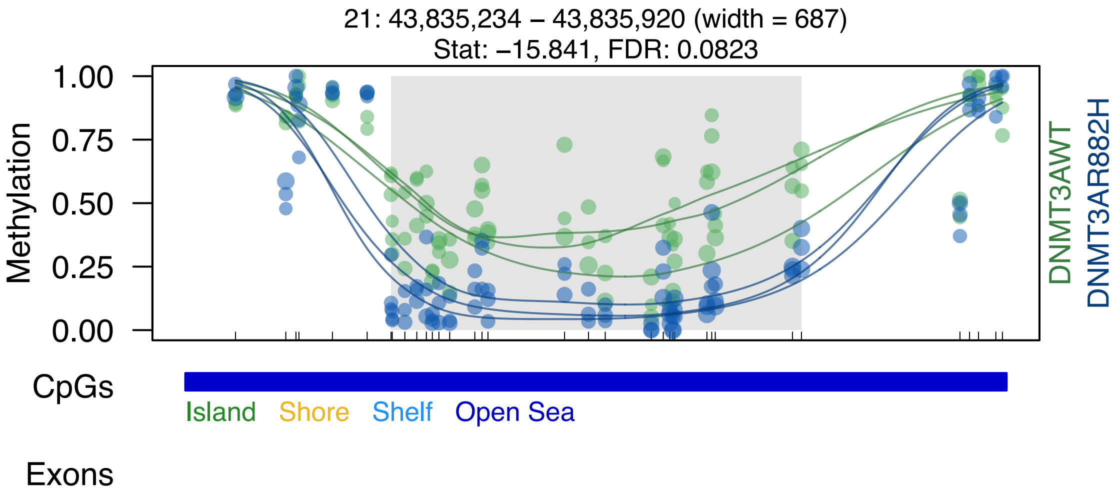
21: 43,835,234 – 43,835,920 (width = 687)
Stat: –15.841, FDR: 0.0823
Methylation
1.00
0.75
0.50
0.25
0.00
DNMT3AWT
DNMT3AR882H
CpGs
Island
Shore
Shelf
Open Sea
Exons

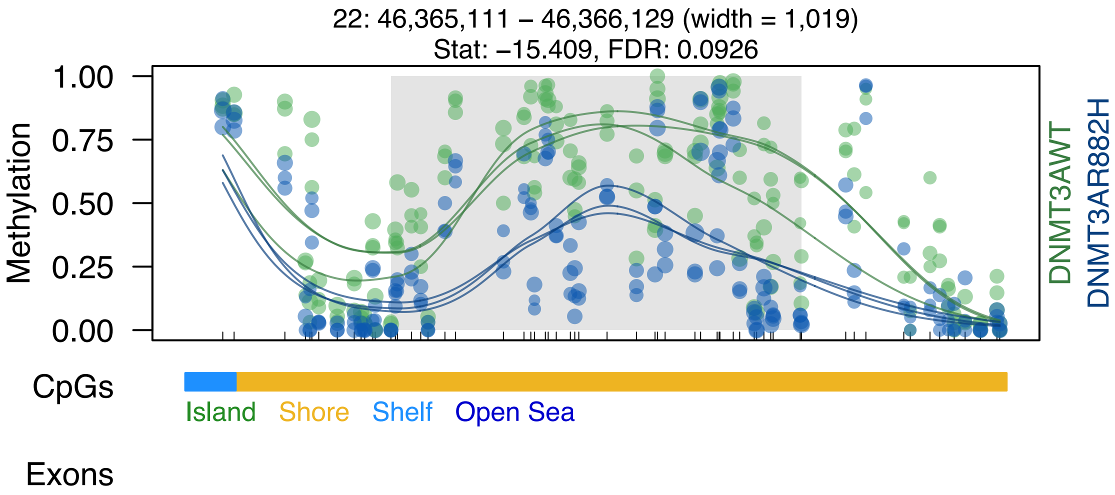
22: 46,365,111 – 46,366,129 (width = 1,019)
Stat: –15.409, FDR: 0.0926
Methylation
1.00
0.75
0.50
0.25
0.00
DNMT3AWT
DNMT3AR882H
CpGs
Island
Shore
Shelf
Open Sea
Exons

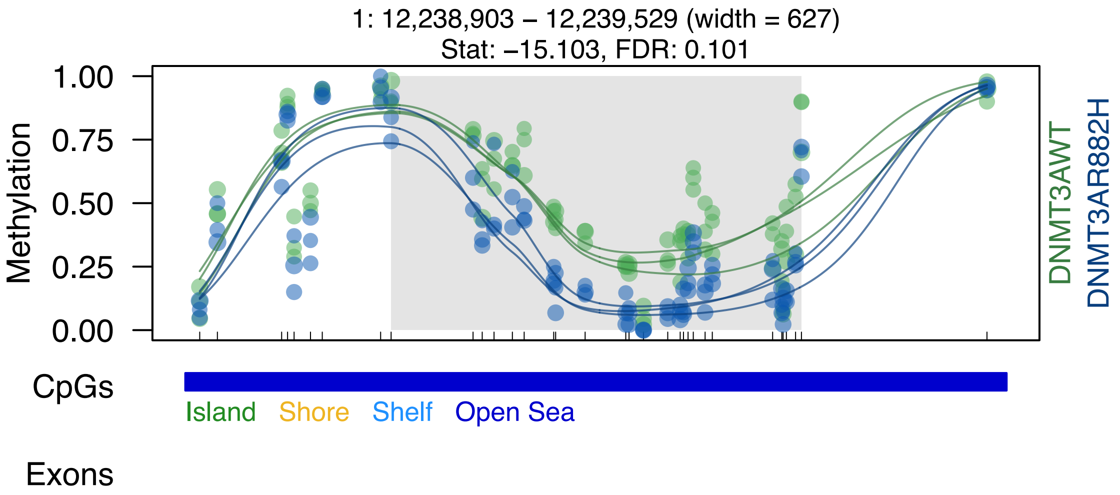

1: 12,238,903 – 12,239,529 (width = 627)
Stat: –15.103, FDR: 0.101
Methylation
1.00
0.75
0.50
0.25
0.00
DNMT3AWT
DNMT3AR882H
CpGs
Island
Shore
Shelf
Open Sea
Exons

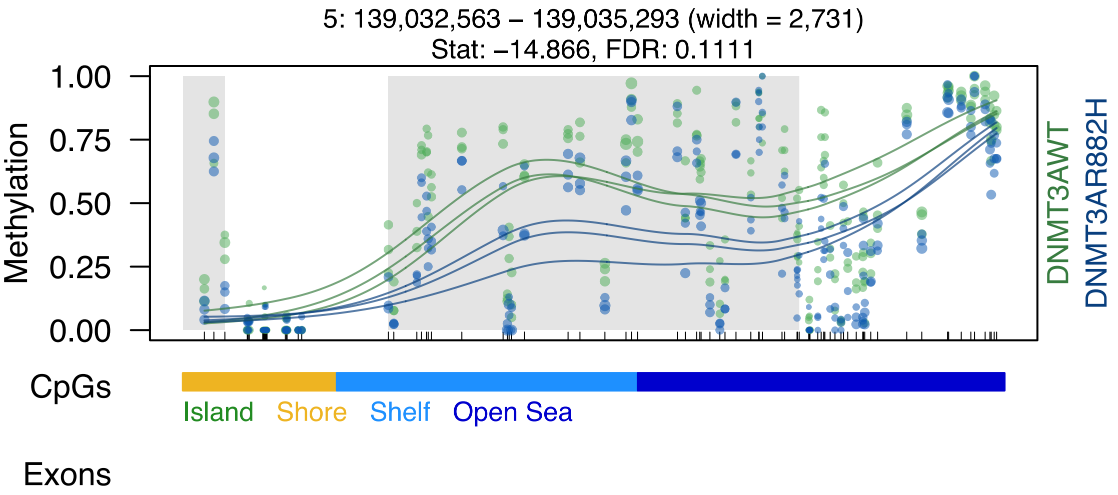
5: 139,032,563 – 139,035,293 (width = 2,731)
Stat: −14.866, FDR: 0.1111
Methylation
1.00
0.75
0.50
0.25
0.00
DNMT3AWT
DNMT3AR882H
CpGs
Island
Shore
Shelf
Open Sea
Exons

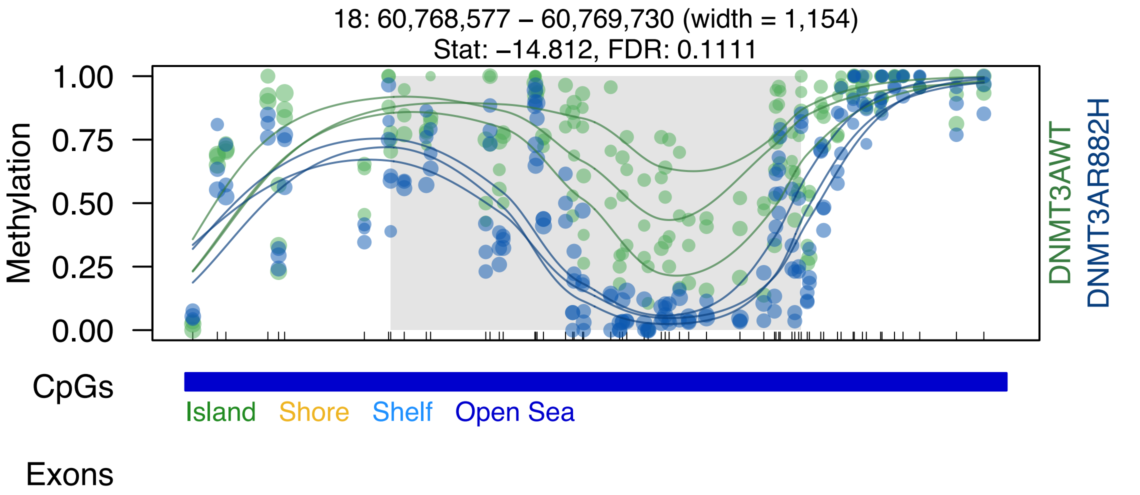

18: 60,768,577 – 60,769,730 (width = 1,154)
Stat: –14.812, FDR: 0.1111
Methylation
1.00
0.75
0.50
0.25
0.00
DNMT3AWT
DNMT3AR882H
CpGs
Island
Shore
Shelf
Open Sea
Exons

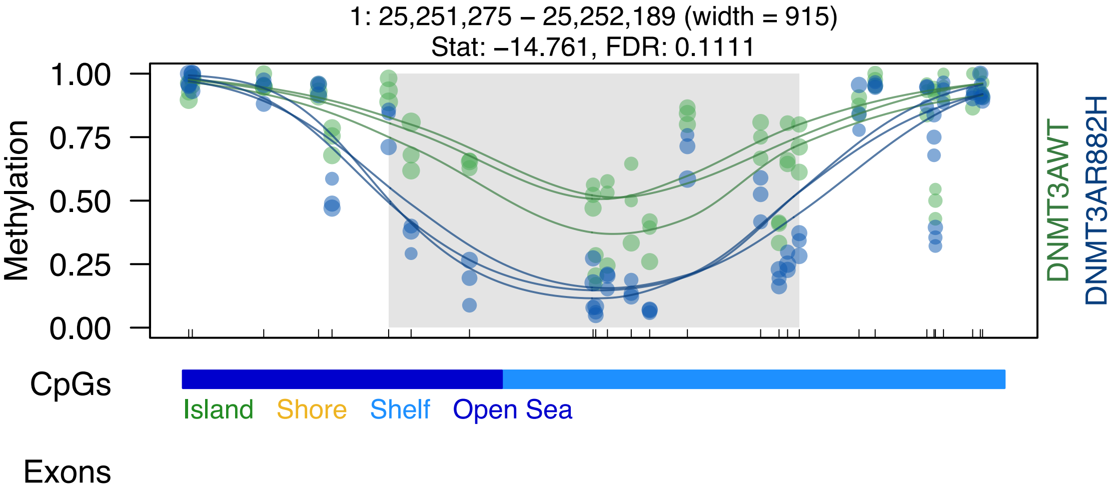

1: 25,251,275 – 25,252,189 (width = 915)
Stat: –14.761, FDR: 0.1111
1.00
0.75
0.50
0.25
0.00
Methylation
DNMT3AWT
DNMT3AR882H
CpGs
Island
Shore
Shelf
Open Sea
Exons

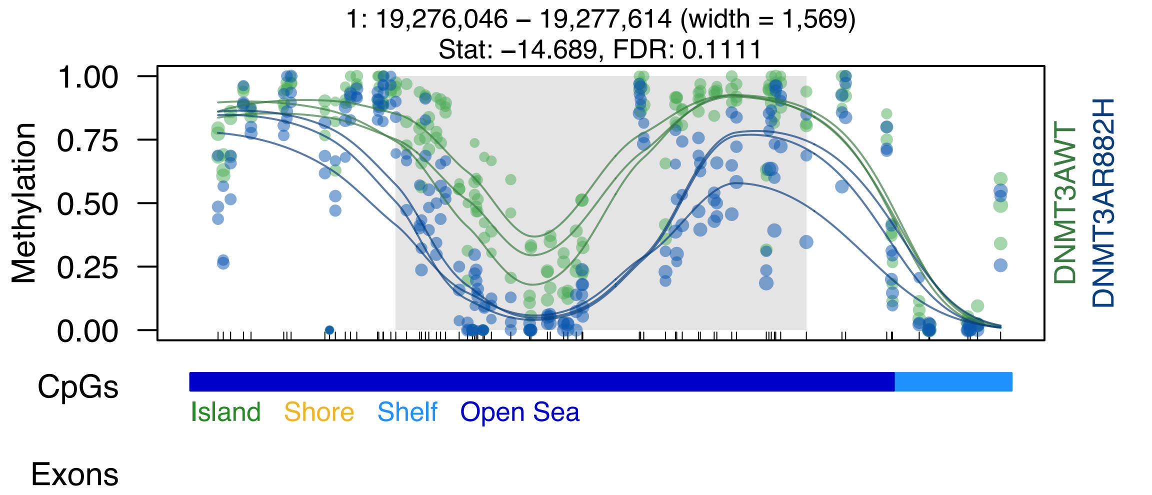
1: 19,276,046 – 19,277,614 (width = 1,569)
Stat: –14.689, FDR: 0.1111
Methylation
1.00
0.75
0.50
0.25
0.00
DNMT3AWT
DNMT3AR882H
CpGs
Island
Shore
Shelf
Open Sea
Exons

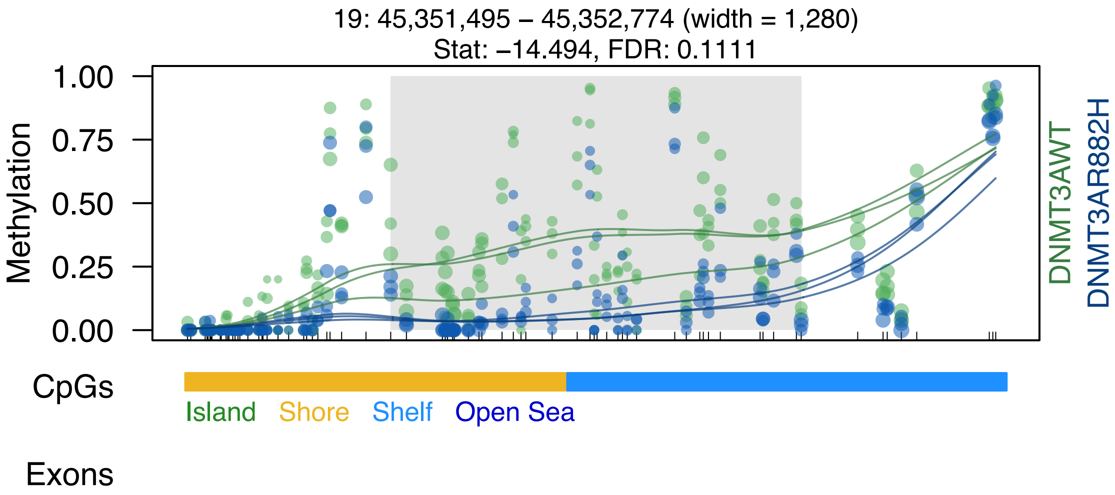
19: 45,351,495 – 45,352,774 (width = 1,280)
Stat: −14.494, FDR: 0.1111
Methylation
1.00
0.75
0.50
0.25
0.00
DNMT3AWT
DNMT3AR882H
CpGs
Island
Shore
Shelf
Open Sea
Exons

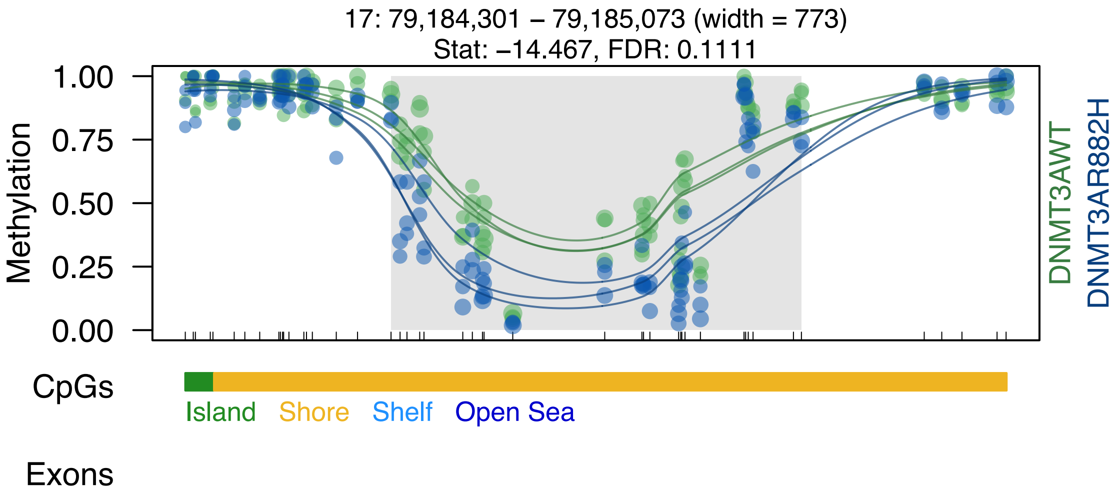

17: 79,184,301 – 79,185,073 (width = 773)
Stat: −14.467, FDR: 0.1111
1.00
0.75
0.50
0.25
0.00
Methylation
DNMT3AWT
DNMT3AR882H
CpGs
Island
Shore
Shelf
Open Sea
Exons

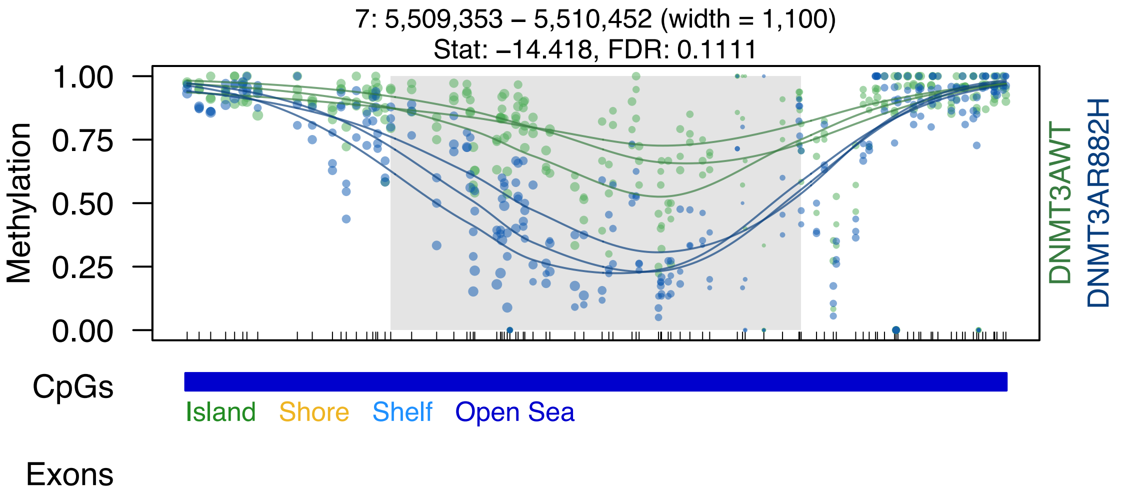

7: 5,509,353 – 5,510,452 (width = 1,100)
Stat: −14.418, FDR: 0.1111
Methylation
1.00
0.75
0.50
0.25
0.00
DNMT3AWT
DNMT3AR882H
CpGs
Island
Shore
Shelf
Open Sea
Exons

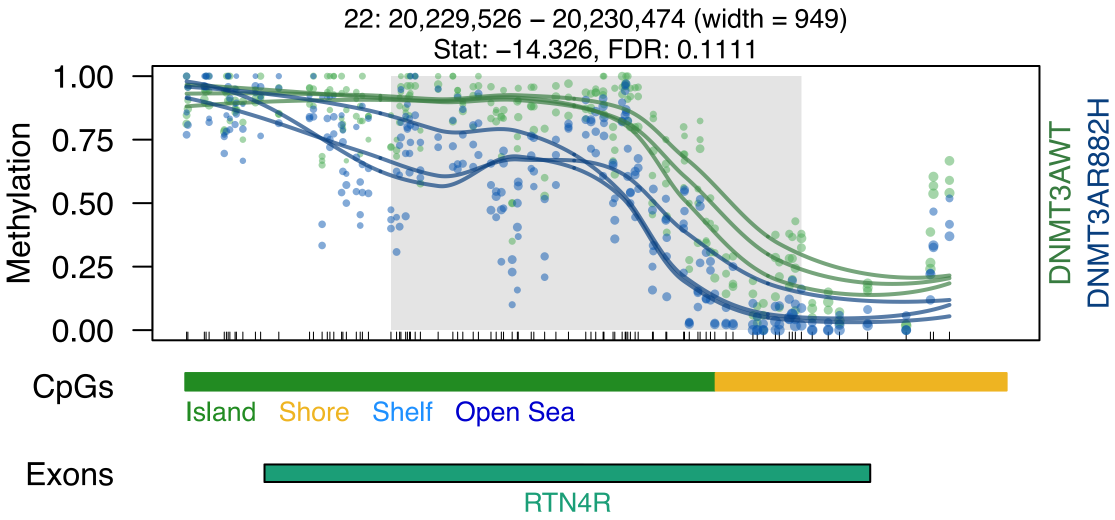

22: 20,229,526 – 20,230,474 (width = 949)
Stat: −14.326, FDR: 0.1111
Methylation
1.00
0.75
0.50
0.25
0.00
DNMT3AWT
DNMT3AR882H
CpGs
Island
Shore
Shelf
Open Sea
Exons
RTN4R

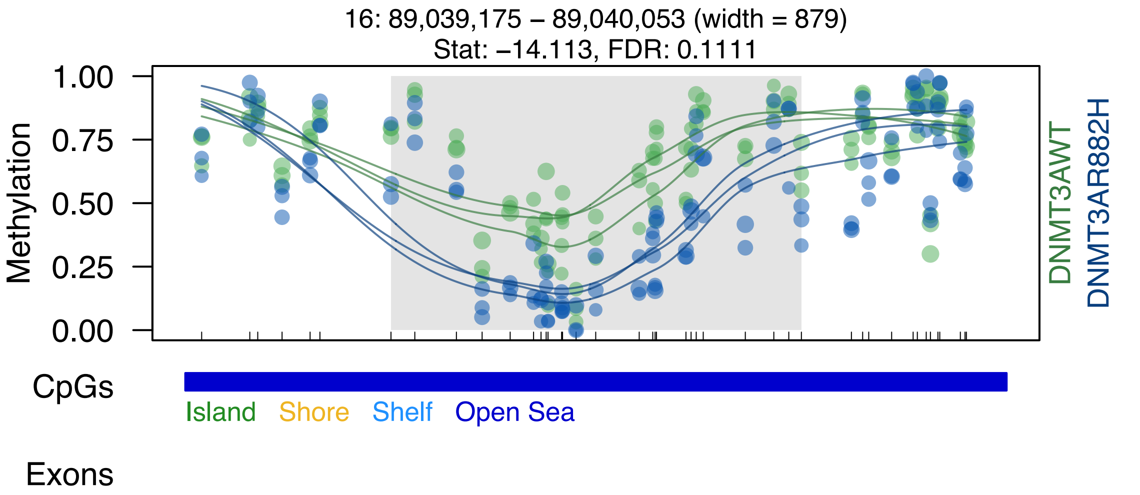

16: 89,039,175 – 89,040,053 (width = 879)
Stat: –14.113, FDR: 0.1111
Methylation
1.00
0.75
0.50
0.25
0.00
DNMT3AWT
DNMT3AR882H
CpGs
Island
Shore
Shelf
Open Sea
Exons

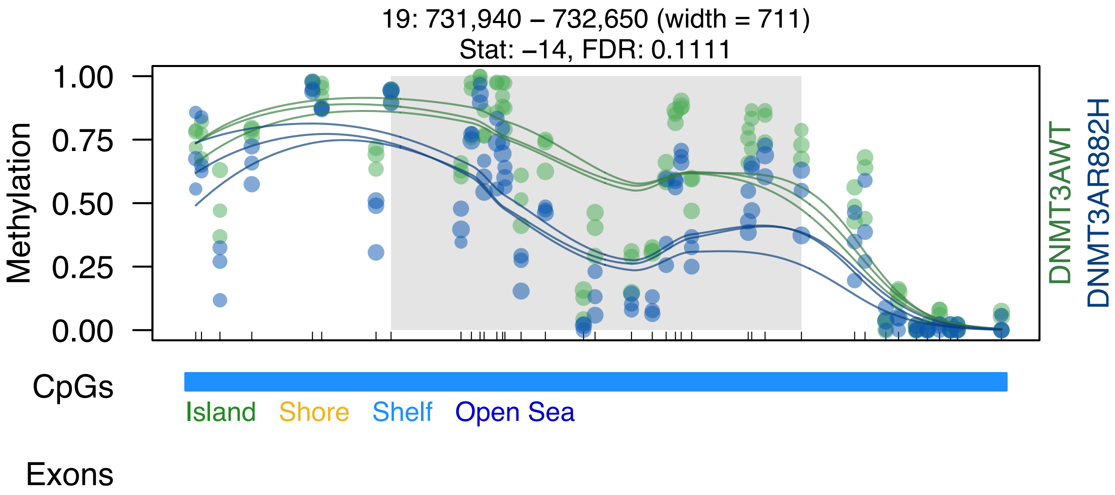

19: 731,940 – 732,650 (width = 711)
Stat: –14, FDR: 0.1111
Methylation
1.00
0.75
0.50
0.25
0.00
DNMT3AWT
DNMT3AR882H
CpGs
Island
Shore
Shelf
Open Sea
Exons

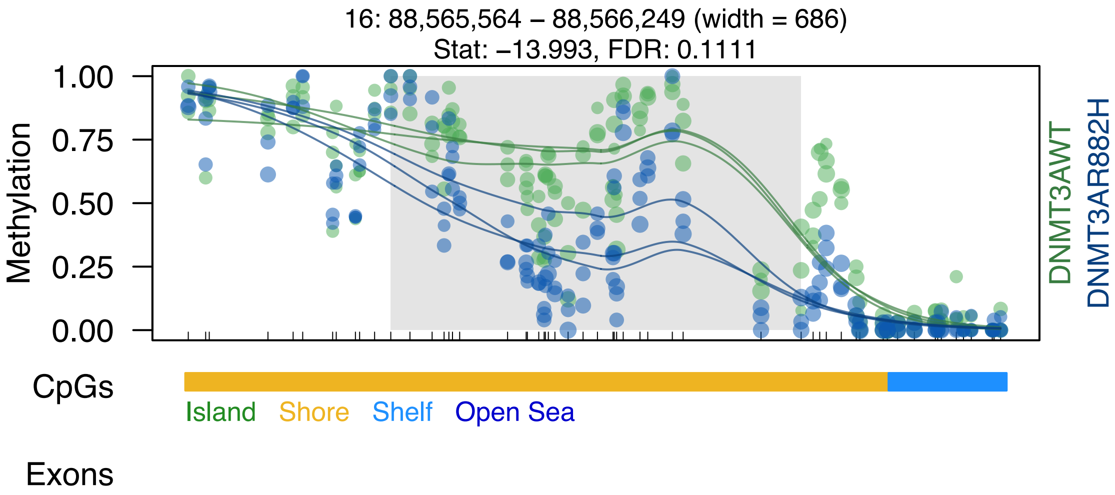
16: 88,565,564 – 88,566,249 (width = 686)
Stat: –13.993, FDR: 0.1111
Methylation
1.00
0.75
0.50
0.25
0.00
DNMT3AWT
DNMT3AR882H
CpGs
Island
Shore
Shelf
Open Sea
Exons

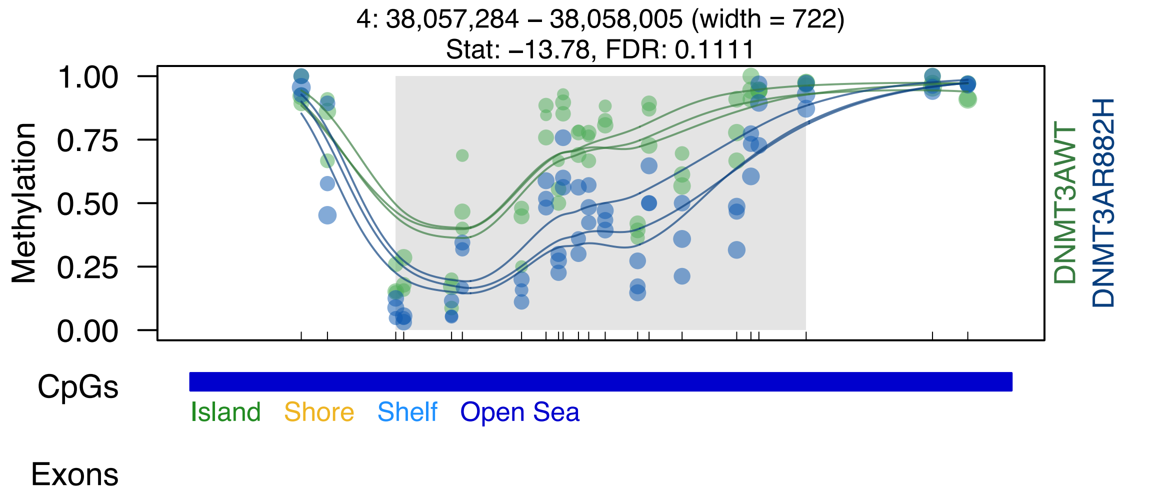
4: 38,057,284 – 38,058,005 (width = 722)
Stat: −13.78, FDR: 0.1111
1.00
0.75
0.50
0.25
0.00
Methylation
DNMT3AWT
DNMT3AR882H
CpGs
Island
Shore
Shelf
Open Sea
Exons

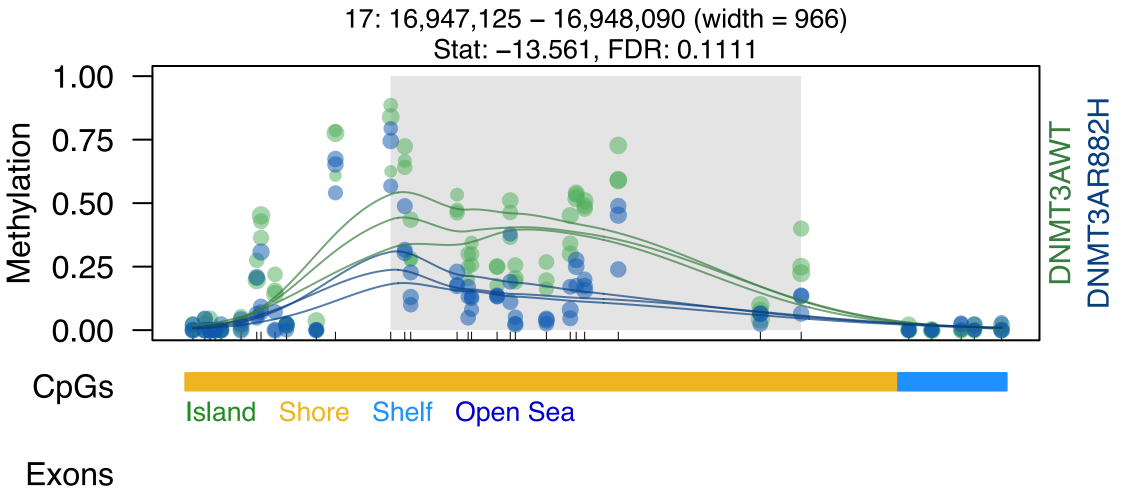
17: 16,947,125 – 16,948,090 (width = 966)
Stat: –13.561, FDR: 0.1111
Methylation
1.00
0.75
0.50
0.25
0.00
DNMT3AWT
DNMT3AR882H
CpGs
Island
Shore
Shelf
Open Sea
Exons

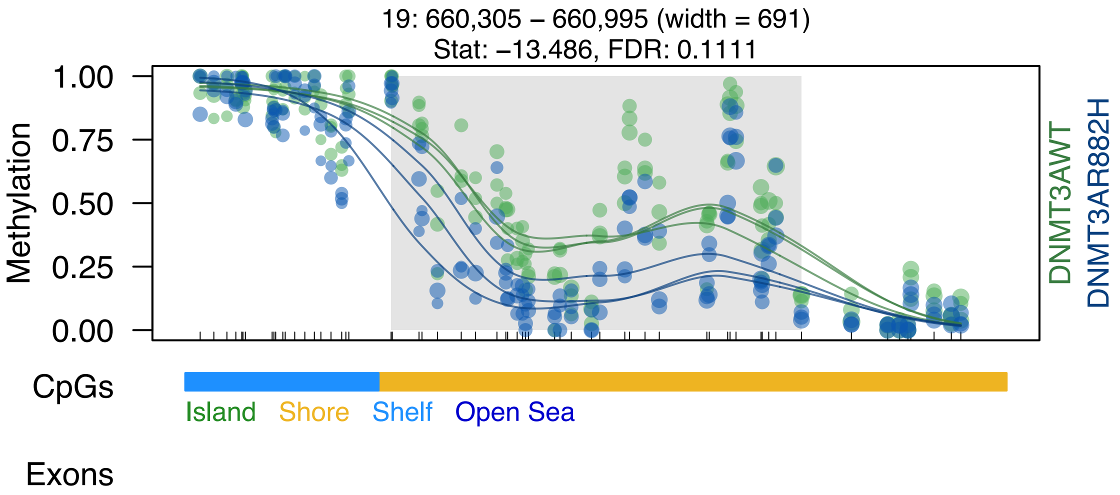

19: 660,305 – 660,995 (width = 691)
Stat: –13.486, FDR: 0.1111
1.00
0.75
0.50
0.25
0.00
Methylation
DNMT3AWT
DNMT3AR882H
CpGs
Island
Shore
Shelf
Open Sea
Exons

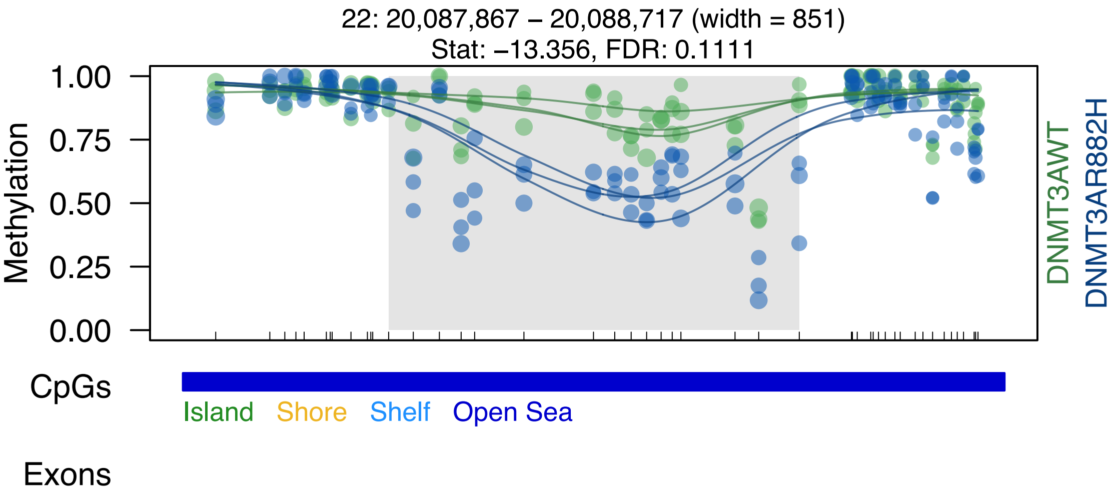

22: 20,087,867 – 20,088,717 (width = 851)
Stat: −13.356, FDR: 0.1111
Methylation
1.00
0.75
0.50
0.25
0.00
DNMT3AWT
DNMT3AR882H
CpGs
Island
Shore
Shelf
Open Sea
Exons

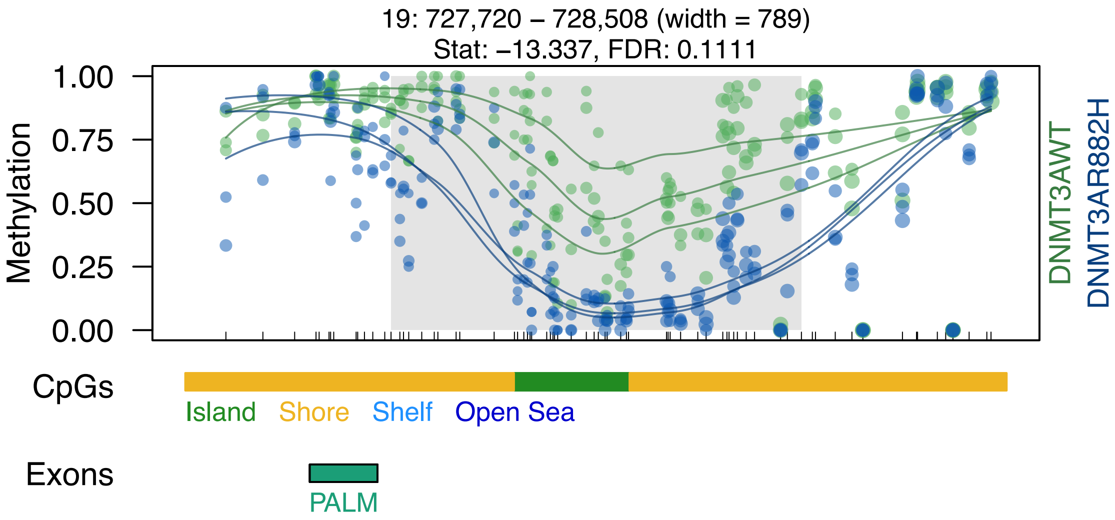

19: 727,720 – 728,508 (width = 789)
Stat: −13.337, FDR: 0.1111
Methylation
1.00
0.75
0.50
0.25
0.00
DNMT3AWT
DNMT3AR882H
CpGs
Island
Shore
Shelf
Open Sea
Exons
PALM

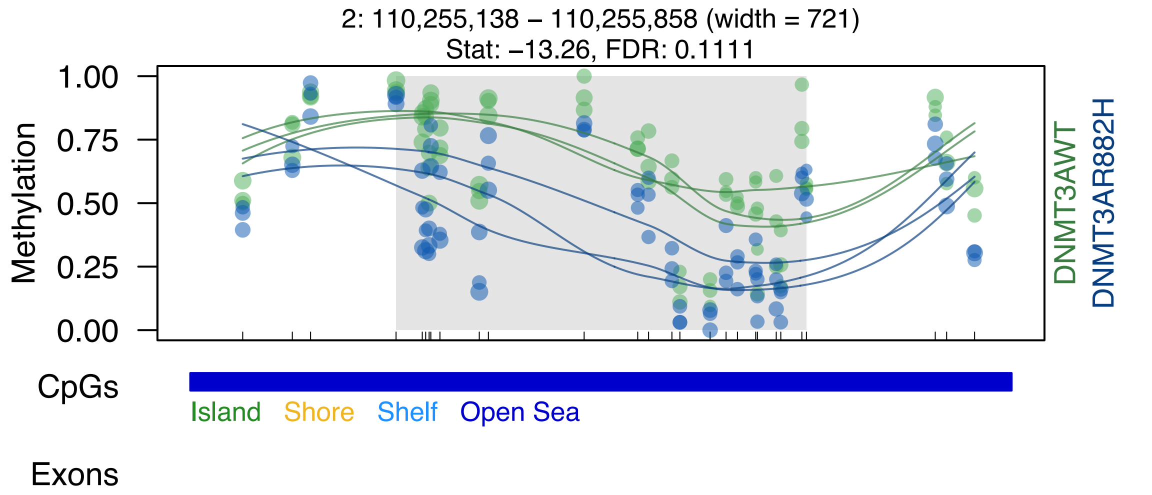

2: 110,255,138 – 110,255,858 (width = 721)
Stat: –13.26, FDR: 0.1111
Methylation
1.00
0.75
0.50
0.25
0.00
DNMT3AWT
DNMT3AR882H
CpGs
Island
Shore
Shelf
Open Sea
Exons

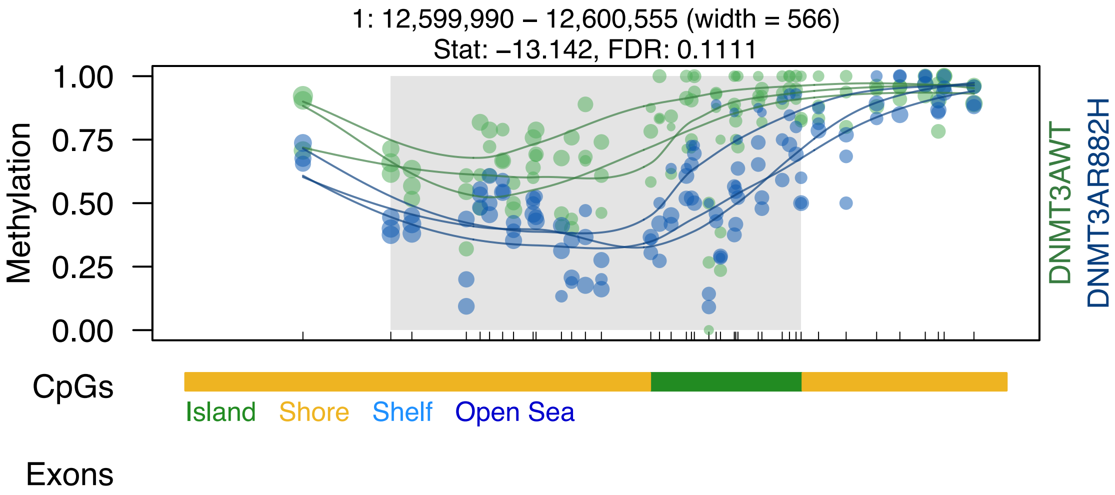

1: 12,599,990 – 12,600,555 (width = 566)
Stat: –13.142, FDR: 0.1111
Methylation
1.00
0.75
0.50
0.25
0.00
DNMT3AWT
DNMT3AR882H
CpGs
Island
Shore
Shelf
Open Sea
Exons

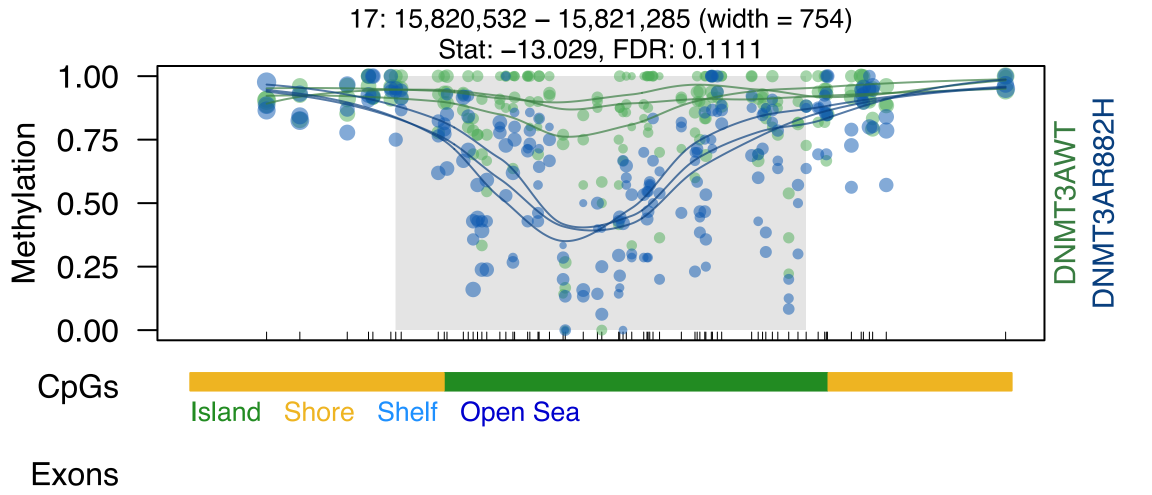

17: 15,820,532 – 15,821,285 (width = 754)
Stat: −13.029, FDR: 0.1111
Methylation
1.00
0.75
0.50
0.25
0.00
DNMT3AWT
DNMT3AR882H
CpGs
Island
Shore
Shelf
Open Sea
Exons

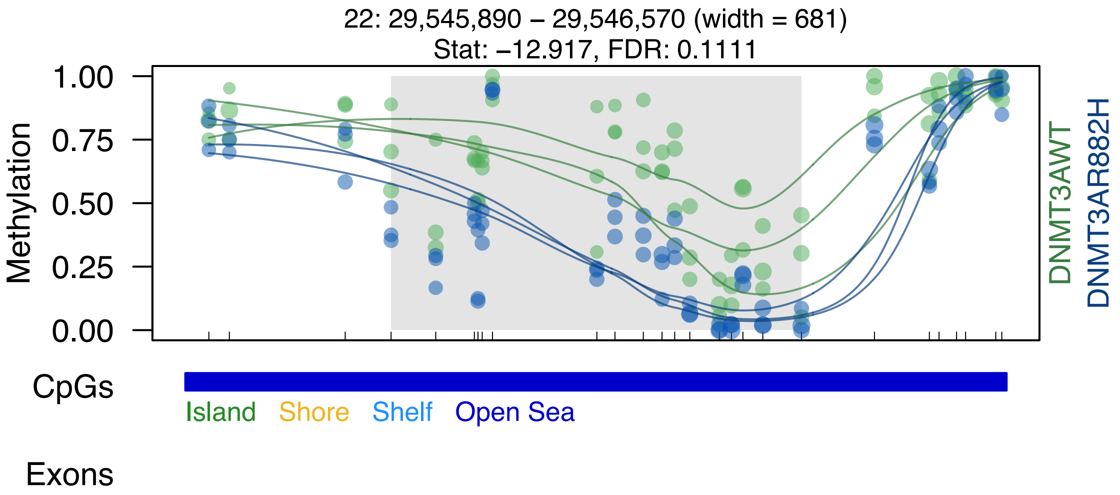

22: 29,545,890 – 29,546,570 (width = 681)
Stat: –12.917, FDR: 0.1111
Methylation
1.00
0.75
0.50
0.25
0.00
DNMT3AWT
DNMT3AR882H
CpGs
Island
Shore
Shelf
Open Sea
Exons

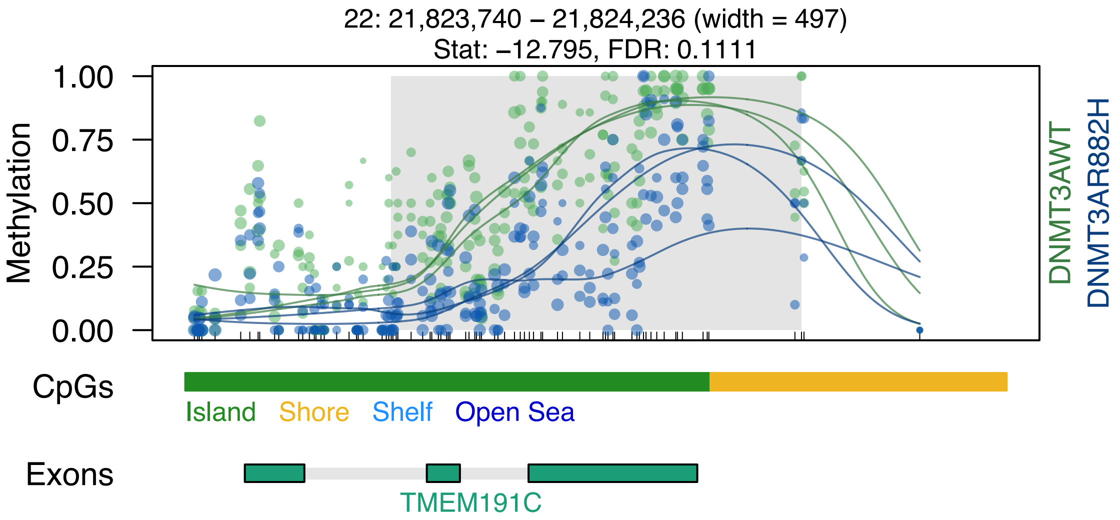

22: 21,823,740 – 21,824,236 (width = 497)
Stat: –12.795, FDR: 0.1111
Methylation
1.00
0.75
0.50
0.25
0.00
DNMT3AWT
DNMT3AR882H
CpGs
Island
Shore
Shelf
Open Sea
Exons
TMEM191C

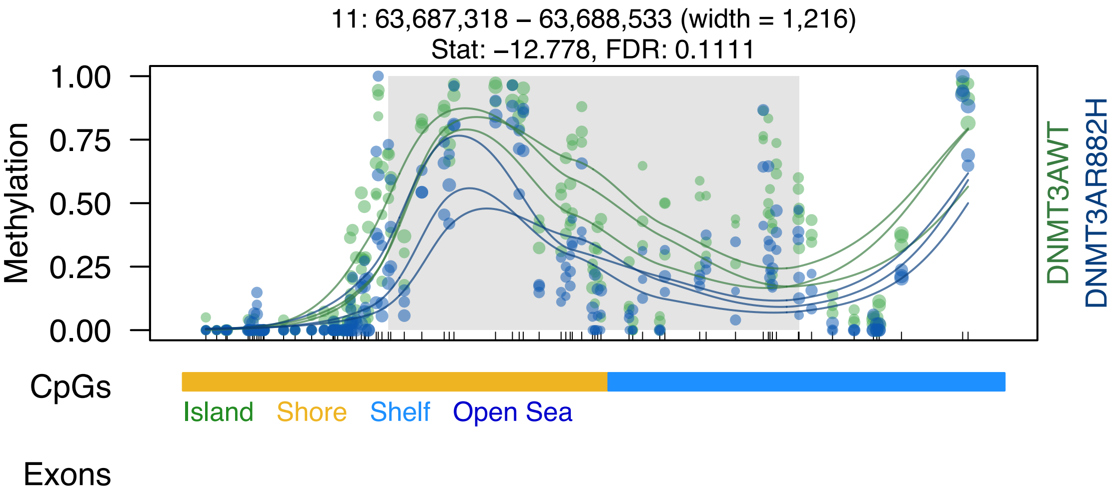

11: 63,687,318 – 63,688,533 (width = 1,216)
Stat: –12.778, FDR: 0.1111
Methylation
1.00
0.75
0.50
0.25
0.00
DNMT3AWT
DNMT3AR882H
CpGs
Island
Shore
Shelf
Open Sea
Exons

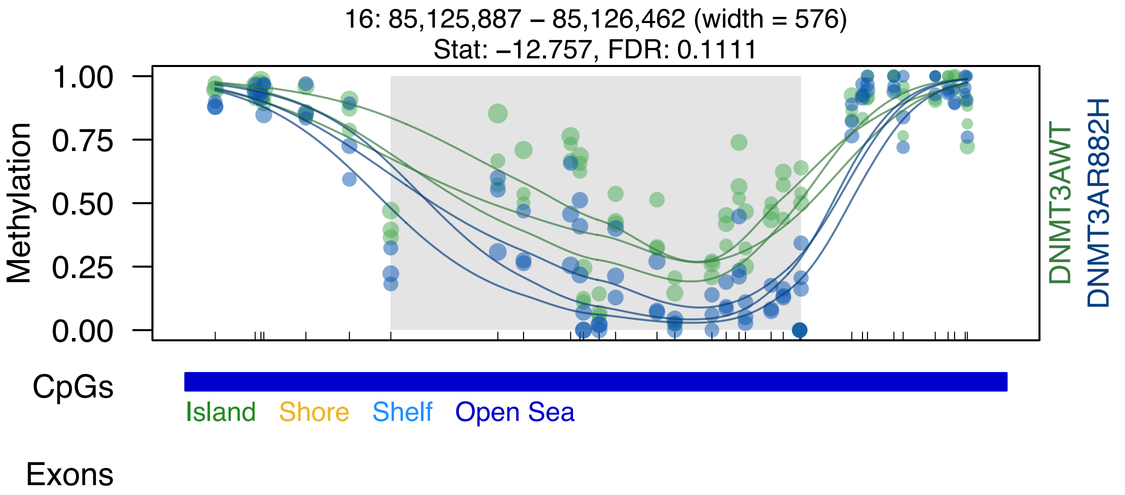
16: 85,125,887 – 85,126,462 (width = 576)
Stat: –12.757, FDR: 0.1111
1.00
0.75
0.50
0.25
0.00
Methylation
DNMT3AWT
DNMT3AR882H
CpGs
Island
Shore
Shelf
Open Sea
Exons

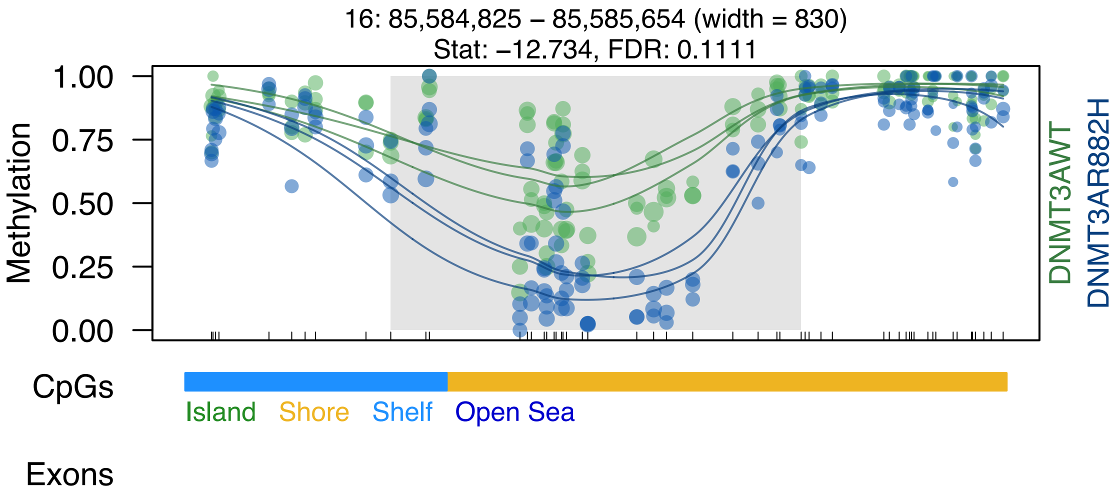

16: 85,584,825 – 85,585,654 (width = 830)
Stat: –12.734, FDR: 0.1111
Methylation
1.00
0.75
0.50
0.25
0.00
DNMT3AWT
DNMT3AR882H
CpGs
Island
Shore
Shelf
Open Sea
Exons

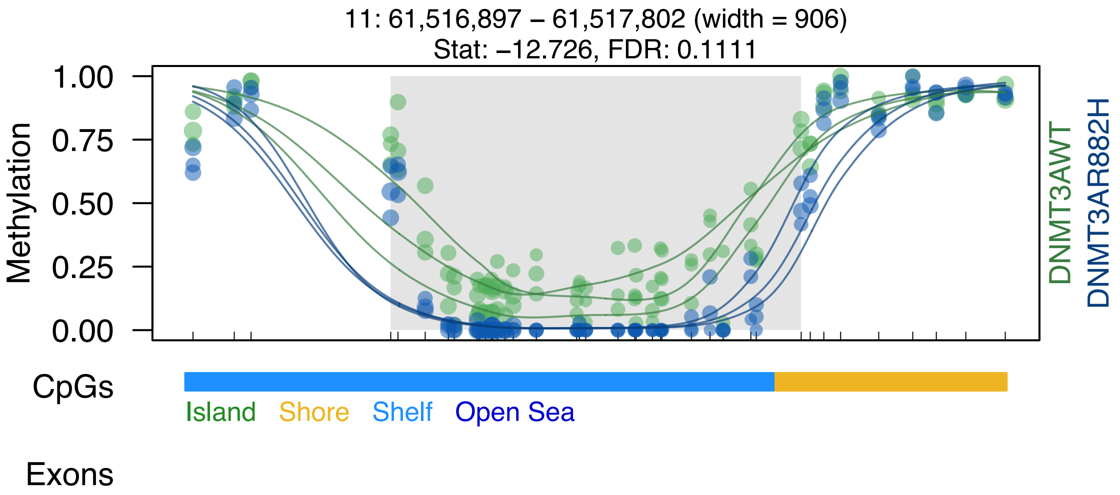

11: 61,516,897 – 61,517,802 (width = 906)
Stat: –12.726, FDR: 0.1111
Methylation
1.00
0.75
0.50
0.25
0.00
DNMT3AWT
DNMT3AR882H
CpGs
Island
Shore
Shelf
Open Sea
Exons

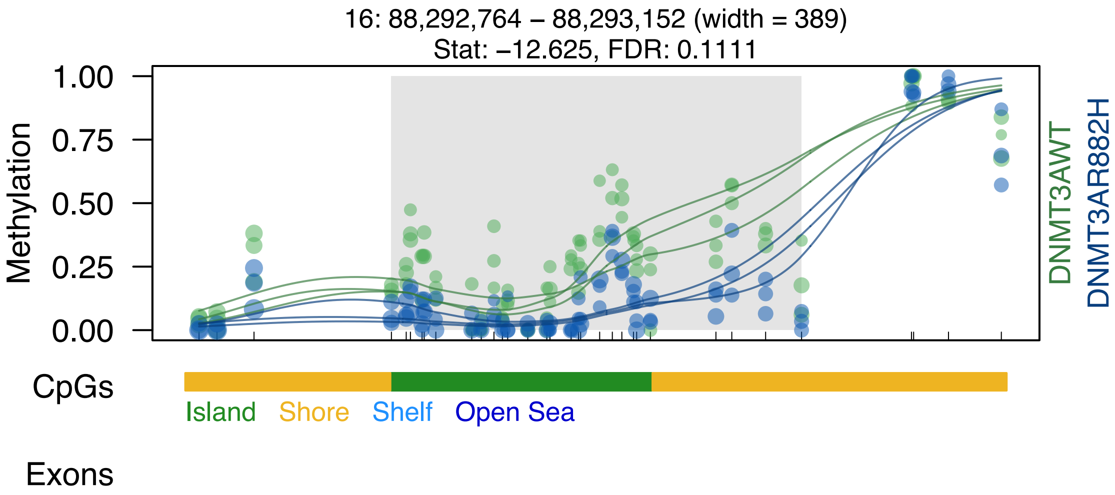
16: 88,292,764 – 88,293,152 (width = 389)
Stat: −12.625, FDR: 0.1111
Methylation
1.00
0.75
0.50
0.25
0.00
DNMT3AWT
DNMT3AR882H
CpGs
Island
Shore
Shelf
Open Sea
Exons

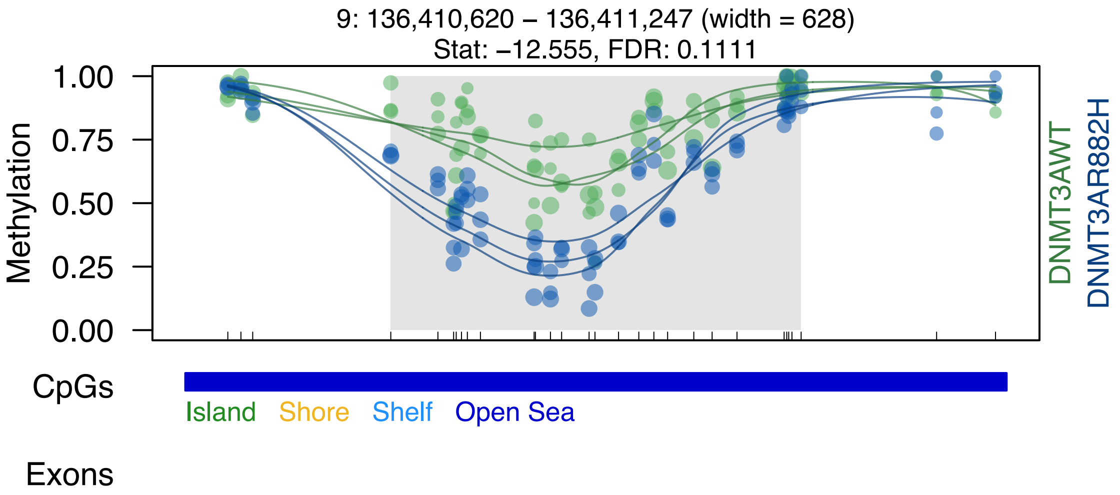
9: 136,410,620 – 136,411,247 (width = 628)
Stat: –12.555, FDR: 0.1111
Methylation
1.00
0.75
0.50
0.25
0.00
DNMT3AWT
DNMT3AR882H
CpGs
Island
Shore
Shelf
Open Sea
Exons

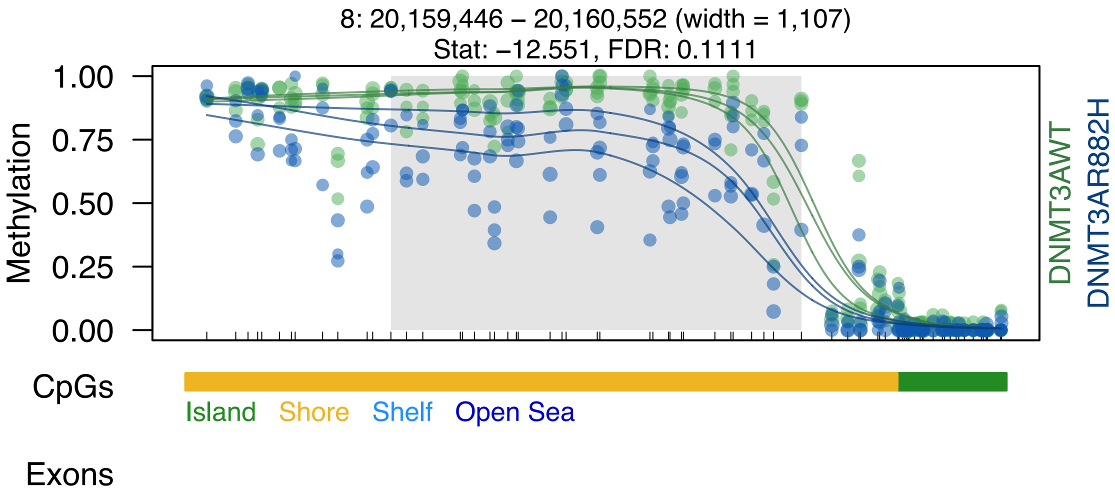

8: 20,159,446 – 20,160,552 (width = 1,107)
Stat: −12.551, FDR: 0.1111
Methylation
1.00
0.75
0.50
0.25
0.00
DNMT3AWT
DNMT3AR882H
CpGs
Island
Shore
Shelf
Open Sea
Exons

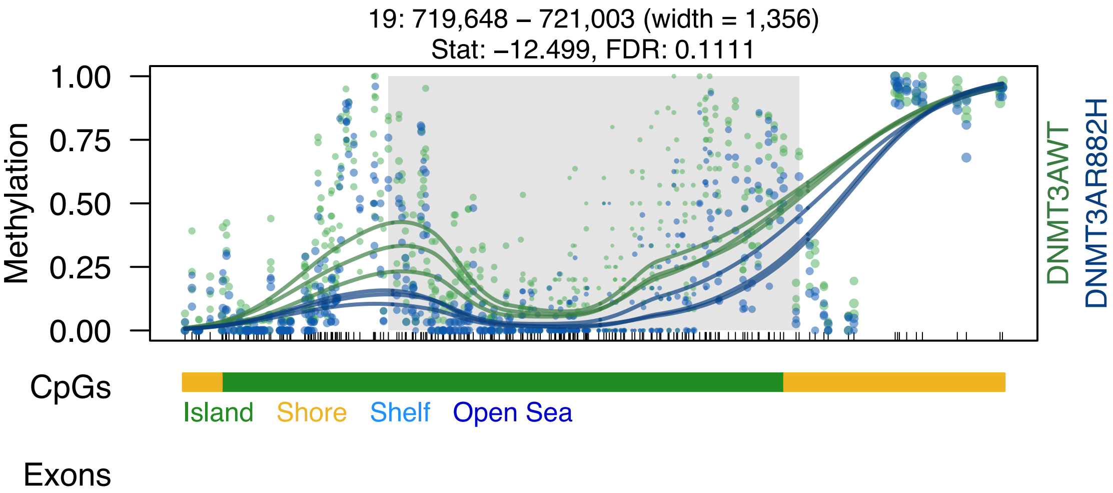
19: 719,648 – 721,003 (width = 1,356)
Stat: –12.499, FDR: 0.1111
Methylation
1.00
0.75
0.50
0.25
0.00
DNMT3AWT
DNMT3AR882H
CpGs
Island
Shore
Shelf
Open Sea
Exons

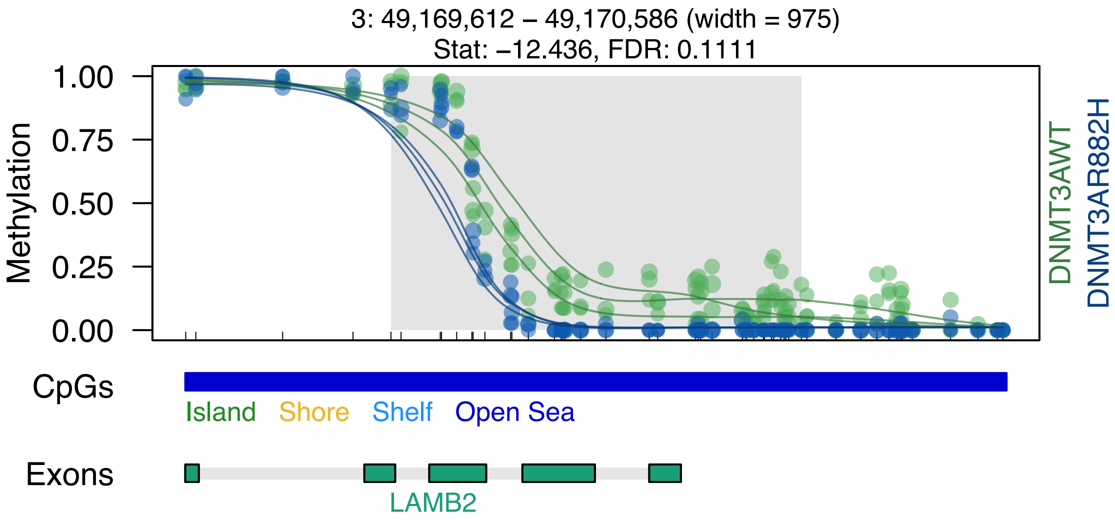

3: 49,169,612 – 49,170,586 (width = 975)
Stat: –12.436, FDR: 0.1111
Methylation
1.00
0.75
0.50
0.25
0.00
DNMT3AWT
DNMT3AR882H
CpGs
Island
Shore
Shelf
Open Sea
Exons
LAMB2

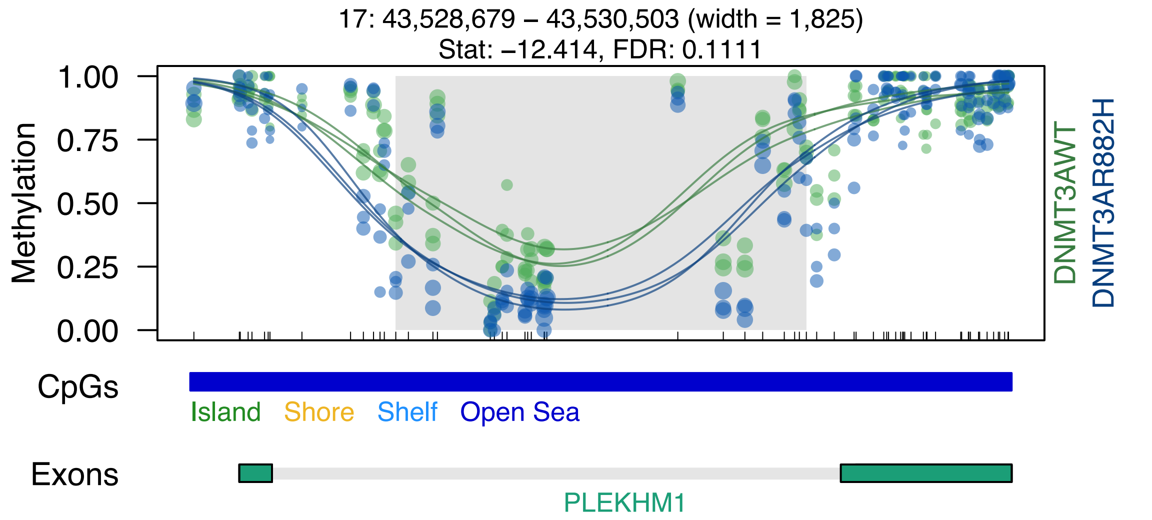

17: 43,528,679 – 43,530,503 (width = 1,825)
Stat: –12.414, FDR: 0.1111
Methylation
1.00
0.75
0.50
0.25
0.00
DNMT3AWT
DNMT3AR882H
CpGs
Island
Shore
Shelf
Open Sea
Exons
PLEKHM1

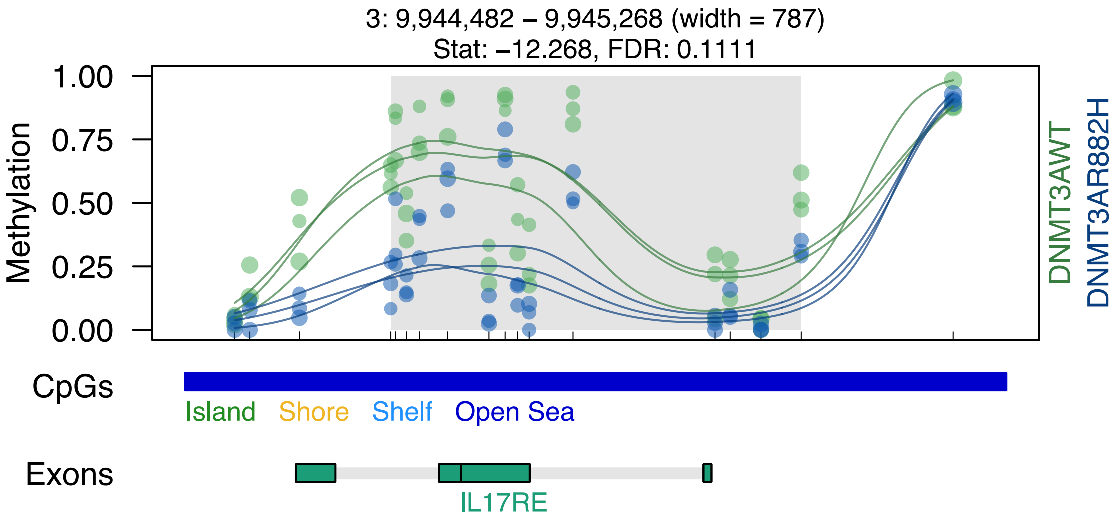
3: 9,944,482 – 9,945,268 (width = 787)
Stat: –12.268, FDR: 0.1111
1.00
0.75
0.50
0.25
0.00
Methylation
DNMT3AWT
DNMT3AR882H
CpGs
Island
Shore
Shelf
Open Sea
Exons
IL17RE

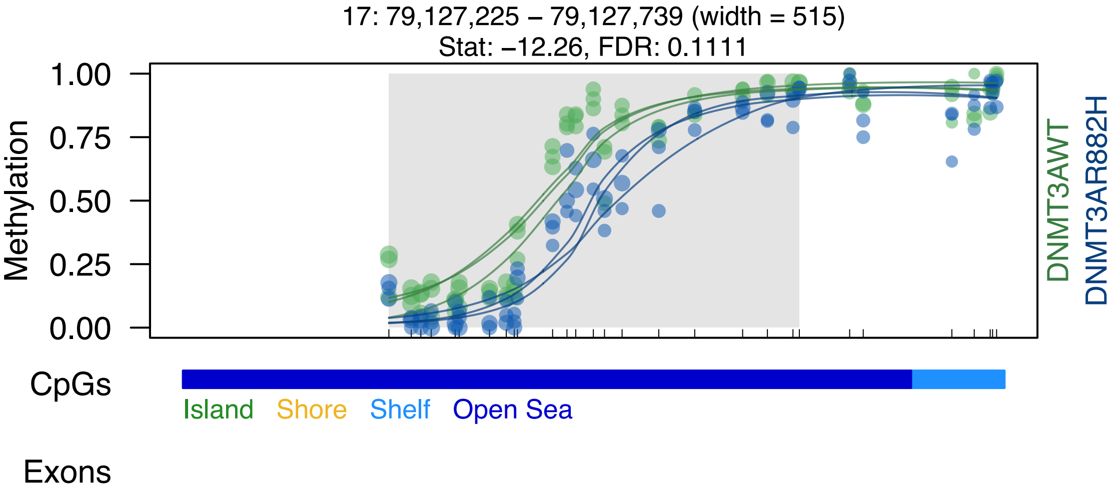
17: 79,127,225 – 79,127,739 (width = 515)
Stat: −12.26, FDR: 0.1111
1.00
0.75
0.50
0.25
0.00
Methylation
DNMT3AWT
DNMT3AR882H
CpGs
Island
Shore
Shelf
Open Sea
Exons

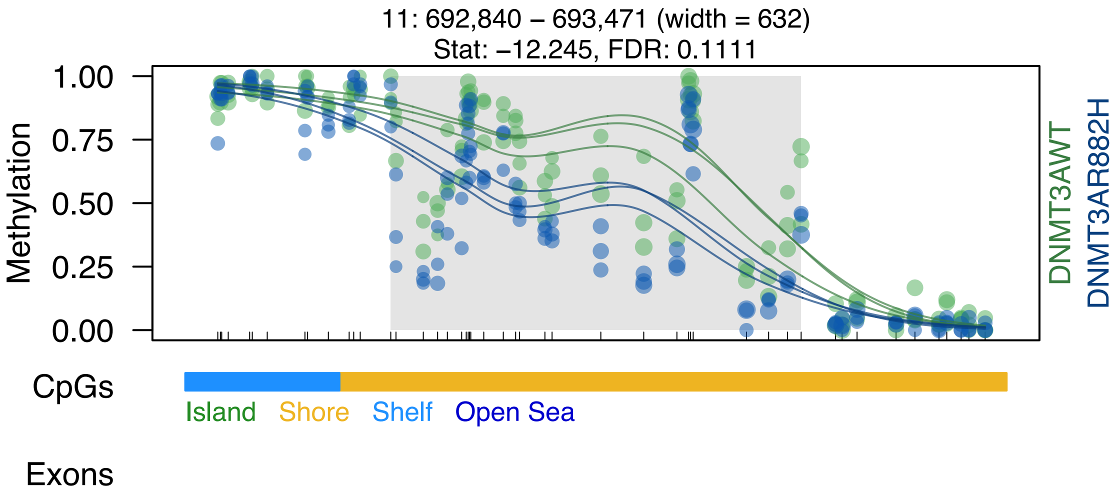

11: 692,840 – 693,471 (width = 632)
Stat: –12.245, FDR: 0.1111
1.00
0.75
0.50
0.25
0.00
Methylation
DNMT3AWT
DNMT3AR882H
CpGs
Island
Shore
Shelf
Open Sea
Exons

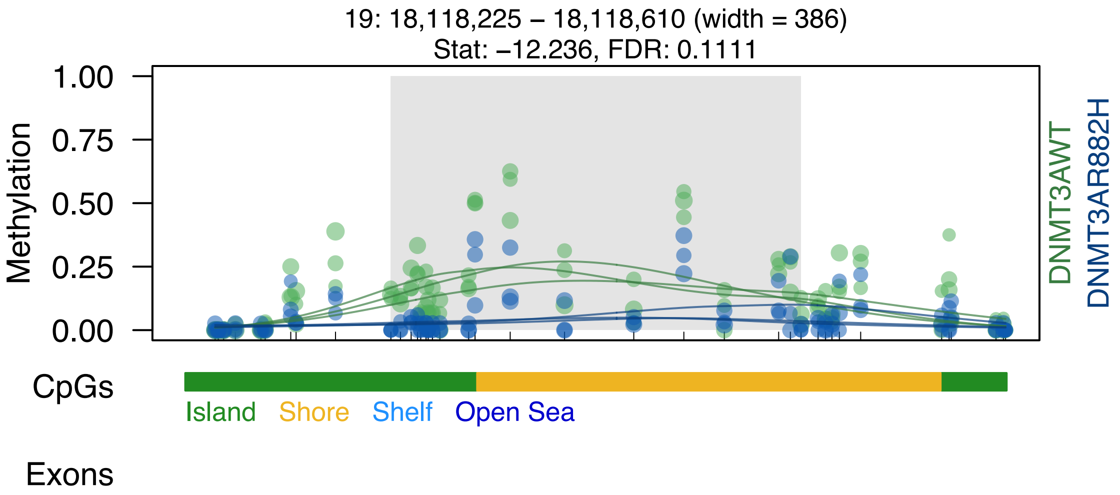
19: 18,118,225 – 18,118,610 (width = 386)
Stat: –12.236, FDR: 0.1111
Methylation
1.00
0.75
0.50
0.25
0.00
DNMT3AWT
DNMT3AR882H
CpGs
Island
Shore
Shelf
Open Sea
Exons

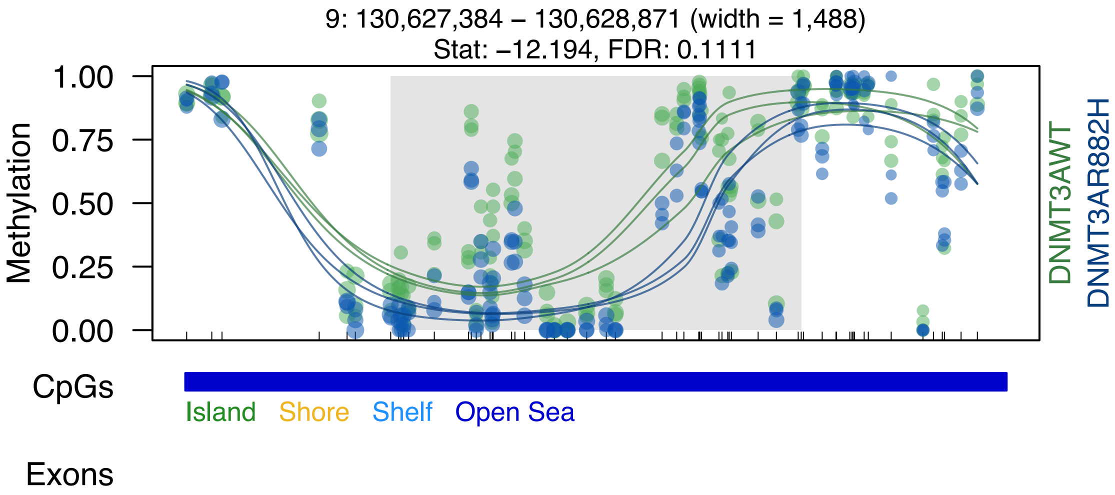
9: 130,627,384 – 130,628,871 (width = 1,488)
Stat: −12.194, FDR: 0.1111
1.00
0.75
0.50
0.25
0.00
Methylation
DNMT3AWT
DNMT3AR882H
CpGs
Island
Shore
Shelf
Open Sea
Exons

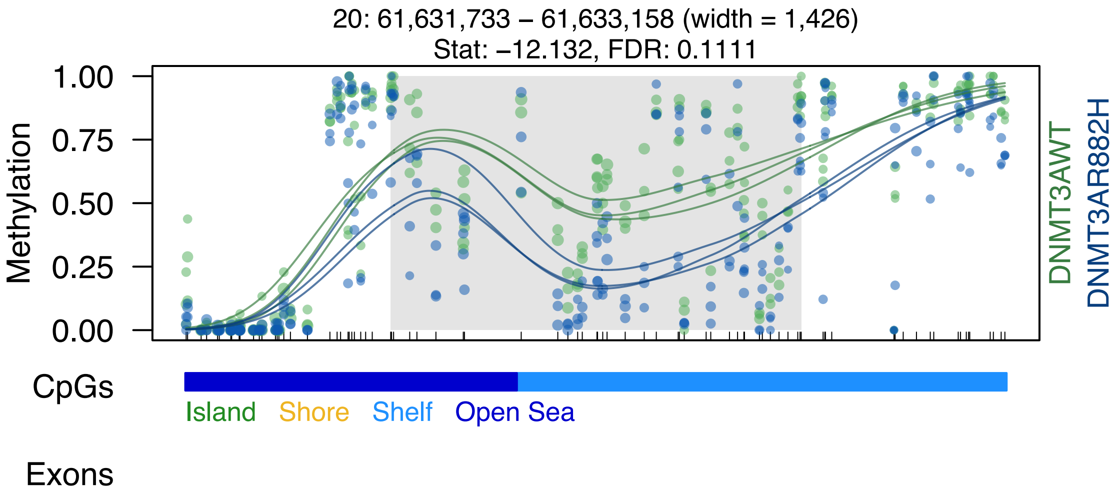
20: 61,631,733 – 61,633,158 (width = 1,426)
Stat: –12.132, FDR: 0.1111
Methylation
1.00
0.75
0.50
0.25
0.00
DNMT3AWT
DNMT3AR882H
CpGs
Island
Shore
Shelf
Open Sea
Exons

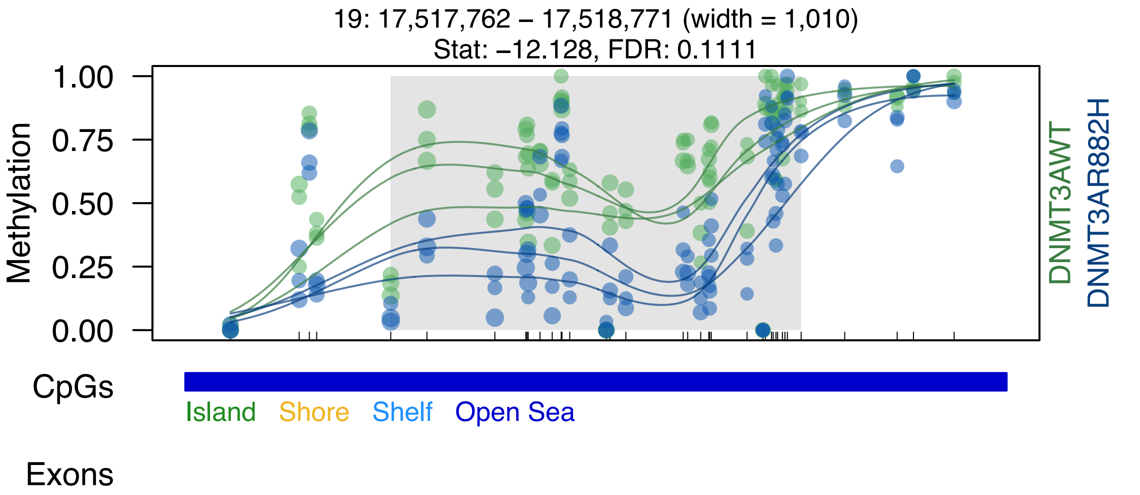

19: 17,517,762 – 17,518,771 (width = 1,010)
Stat: −12.128, FDR: 0.1111
Methylation
1.00
0.75
0.50
0.25
0.00
DNMT3AWT
DNMT3AR882H
CpGs
Island
Shore
Shelf
Open Sea
Exons

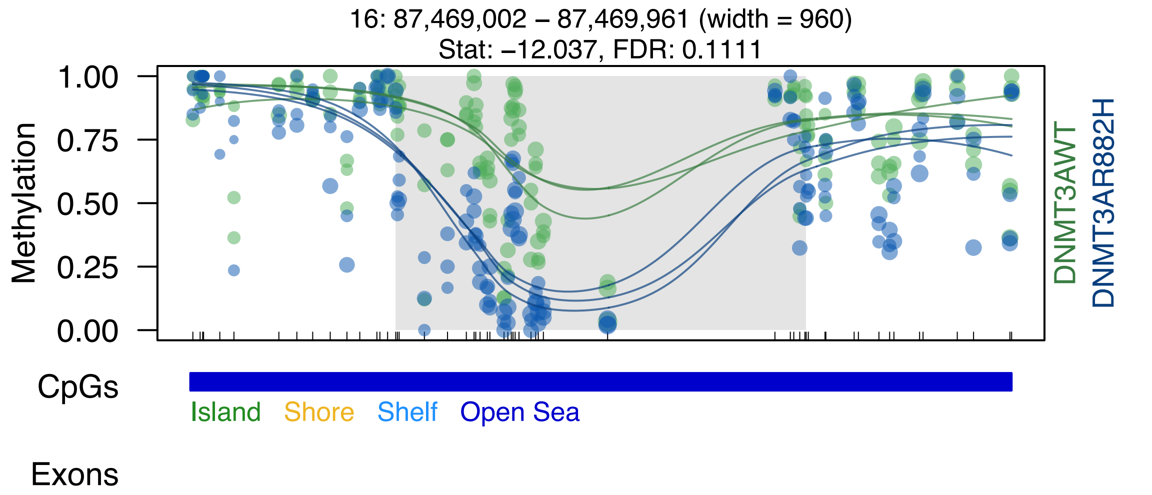
16: 87,469,002 – 87,469,961 (width = 960)
Stat: –12.037, FDR: 0.1111
Methylation
1.00
0.75
0.50
0.25
0.00
DNMT3AWT
DNMT3AR882H
CpGs
Island
Shore
Shelf
Open Sea
Exons

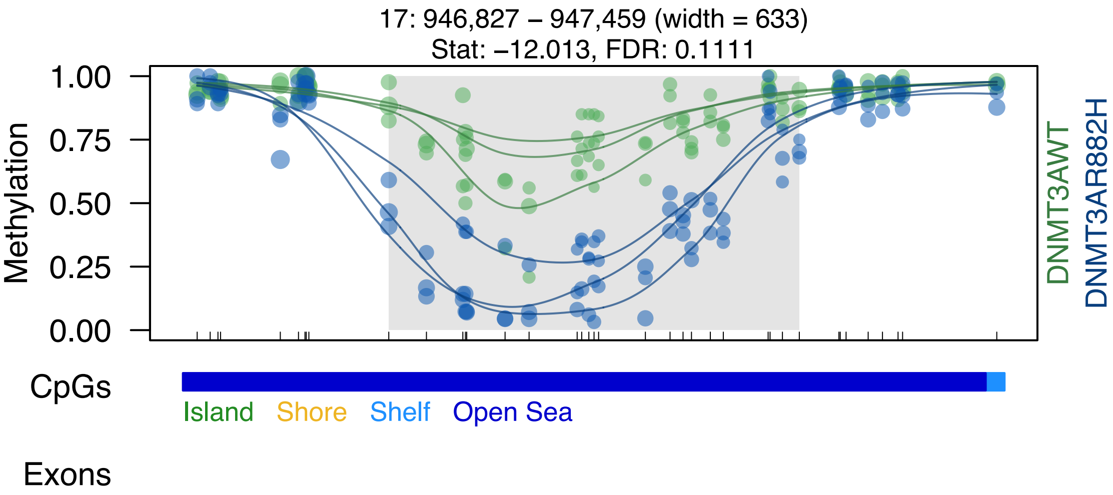
17: 946,827 – 947,459 (width = 633)
Stat: –12.013, FDR: 0.1111
Methylation
1.00
0.75
0.50
0.25
0.00
DNMT3AWT
DNMT3AR882H
CpGs
Island
Shore
Shelf
Open Sea
Exons

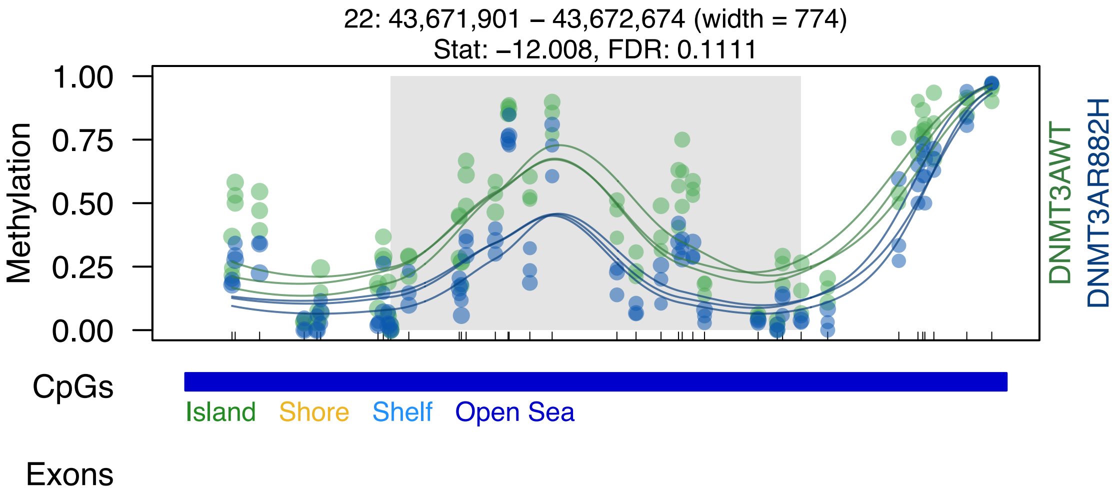
22: 43,671,901 – 43,672,674 (width = 774)
Stat: –12.008, FDR: 0.1111
Methylation
1.00
0.75
0.50
0.25
0.00
DNMT3AWT
DNMT3AR882H
CpGs
Island
Shore
Shelf
Open Sea
Exons

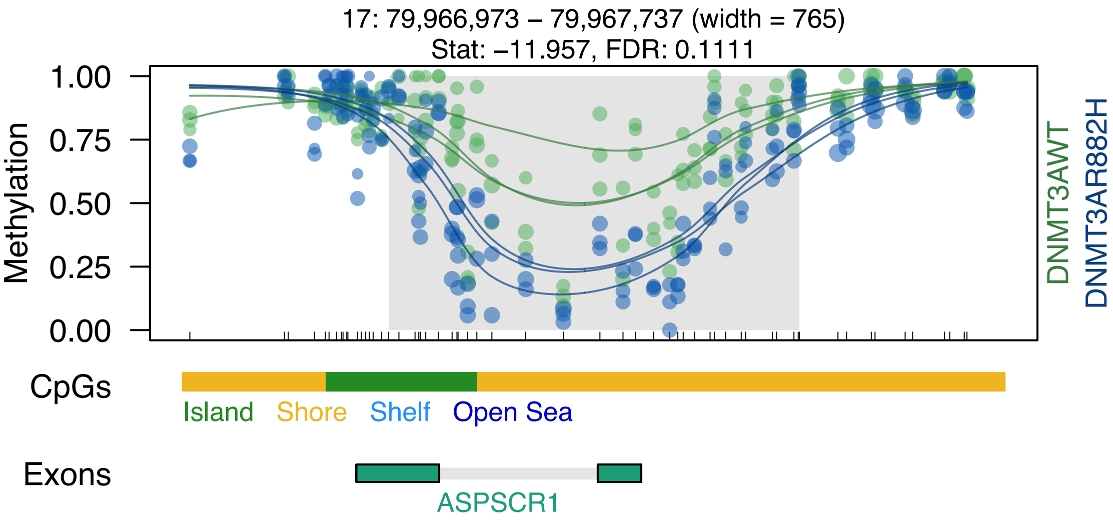
17: 79,966,973 – 79,967,737 (width = 765)
Stat: –11.957, FDR: 0.1111
Methylation
1.00
0.75
0.50
0.25
0.00
DNMT3AWT
DNMT3AR882H
CpGs
Island
Shore
Shelf
Open Sea
Exons
ASPSCR1

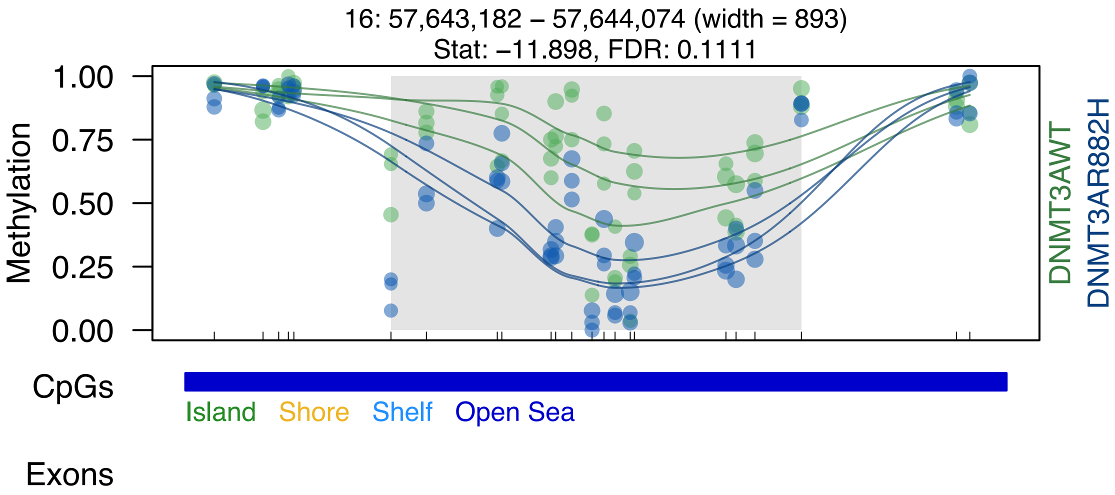

16: 57,643,182 – 57,644,074 (width = 893)
Stat: –11.898, FDR: 0.1111
Methylation
1.00
0.75
0.50
0.25
0.00
DNMT3AWT
DNMT3AR882H
CpGs
Island
Shore
Shelf
Open Sea
Exons

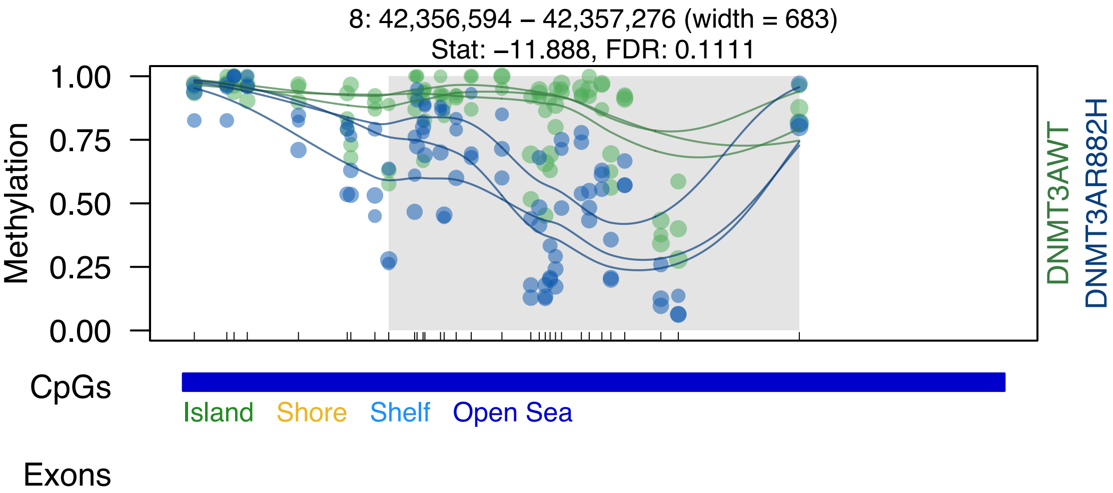

8: 42,356,594 – 42,357,276 (width = 683)
Stat: –11.888, FDR: 0.1111
Methylation
1.00
0.75
0.50
0.25
0.00
DNMT3AWT
DNMT3AR882H
CpGs
Island
Shore
Shelf
Open Sea
Exons

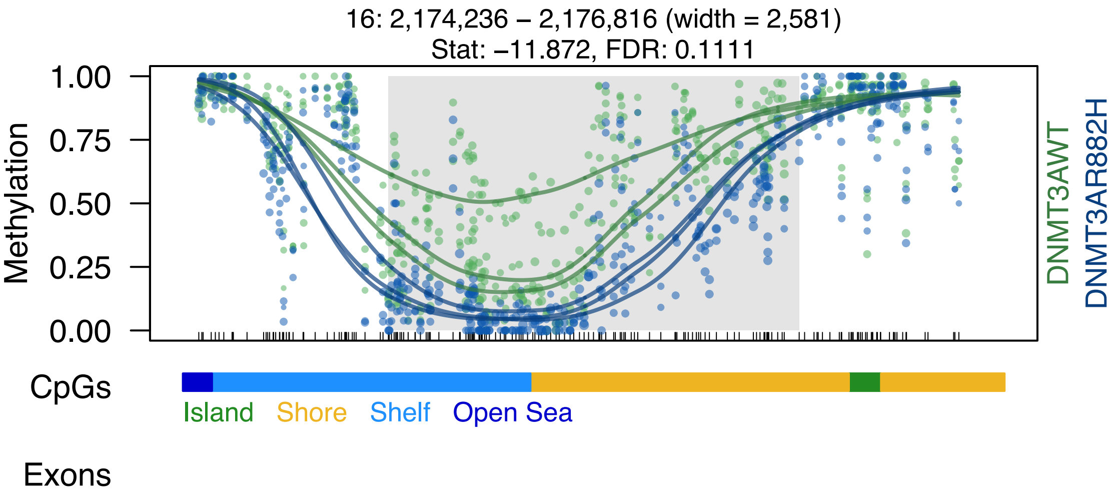
16: 2,174,236 – 2,176,816 (width = 2,581)
Stat: −11.872, FDR: 0.1111
Methylation
1.00
0.75
0.50
0.25
0.00
DNMT3AWT
DNMT3AR882H
CpGs
Island
Shore
Shelf
Open Sea
Exons

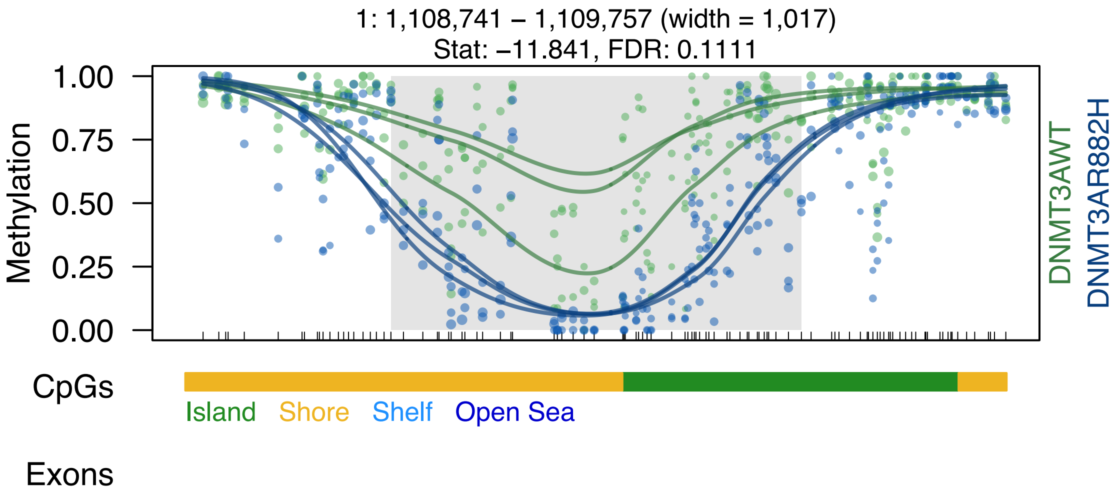
1: 1,108,741 – 1,109,757 (width = 1,017)
Stat: –11.841, FDR: 0.1111
Methylation
1.00
0.75
0.50
0.25
0.00
DNMT3AWT
DNMT3AR882H
CpGs
Island
Shore
Shelf
Open Sea
Exons

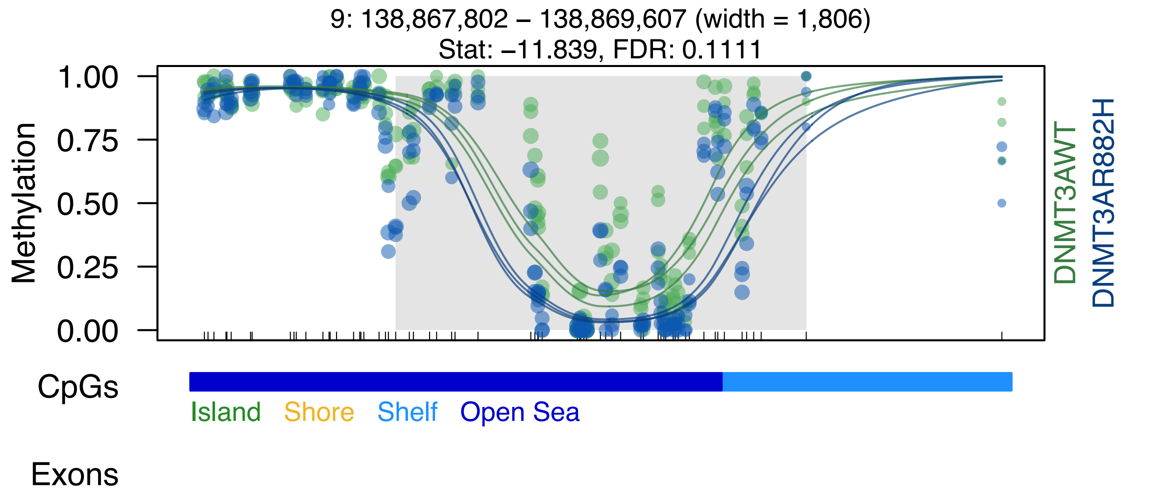
9: 138,867,802 – 138,869,607 (width = 1,806)
Stat: −11.839, FDR: 0.1111
Methylation
1.00
0.75
0.50
0.25
0.00
DNMT3AWT
DNMT3AR882H
CpGs
Island
Shore
Shelf
Open Sea
Exons

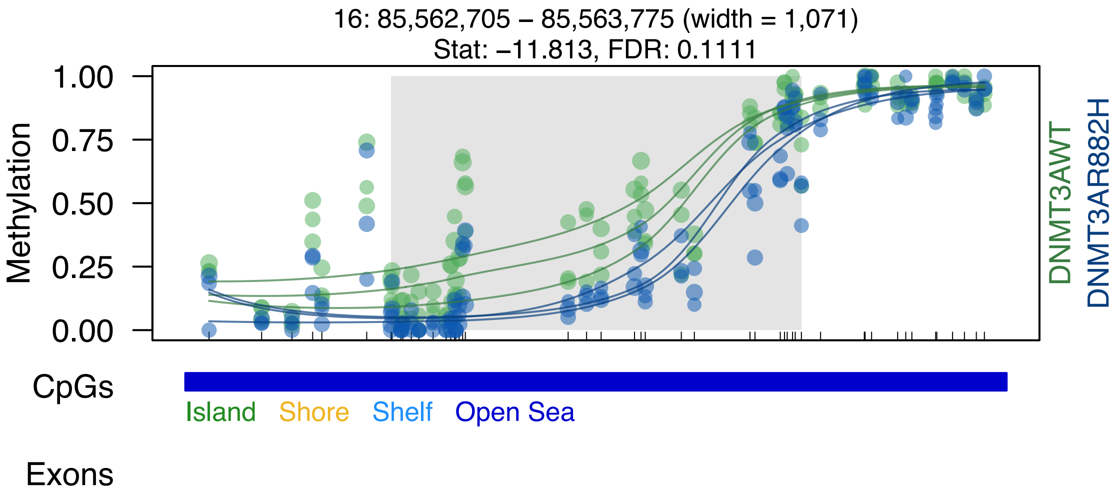

16: 85,562,705 – 85,563,775 (width = 1,071)
Stat: −11.813, FDR: 0.1111
Methylation
1.00
0.75
0.50
0.25
0.00
DNMT3AWT
DNMT3AR882H
CpGs
Island
Shore
Shelf
Open Sea
Exons

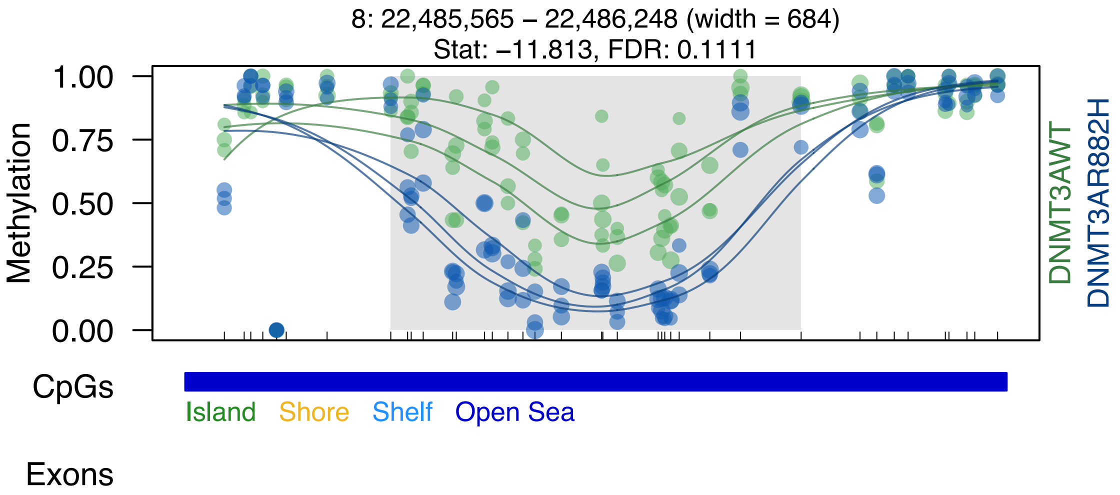
8: 22,485,565 – 22,486,248 (width = 684)
Stat: –11.813, FDR: 0.1111
Methylation
1.00
0.75
0.50
0.25
0.00
DNMT3AWT
DNMT3AR882H
CpGs
Island
Shore
Shelf
Open Sea
Exons

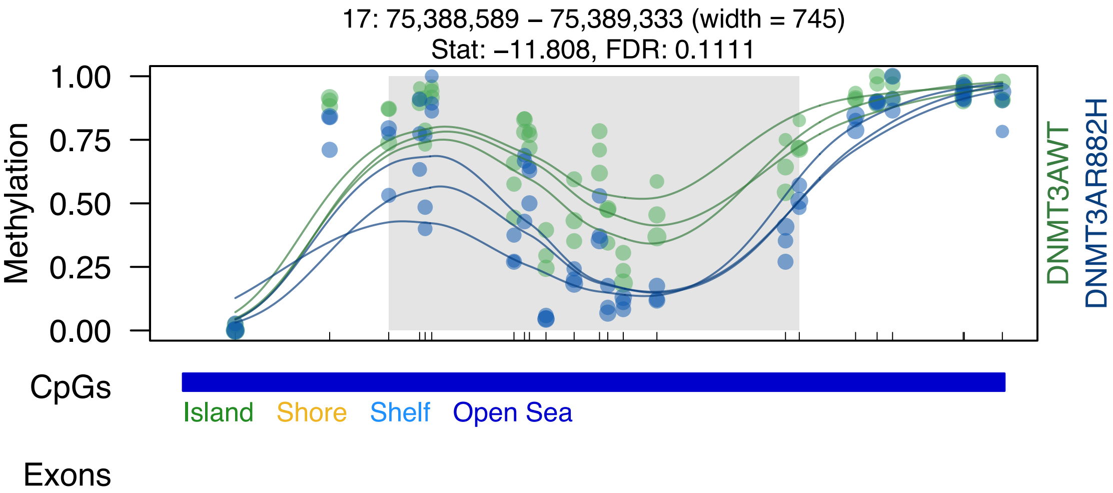

17: 75,388,589 – 75,389,333 (width = 745)
Stat: –11.808, FDR: 0.1111
Methylation
1.00
0.75
0.50
0.25
0.00
DNMT3AWT
DNMT3AR882H
CpGs
Island
Shore
Shelf
Open Sea
Exons

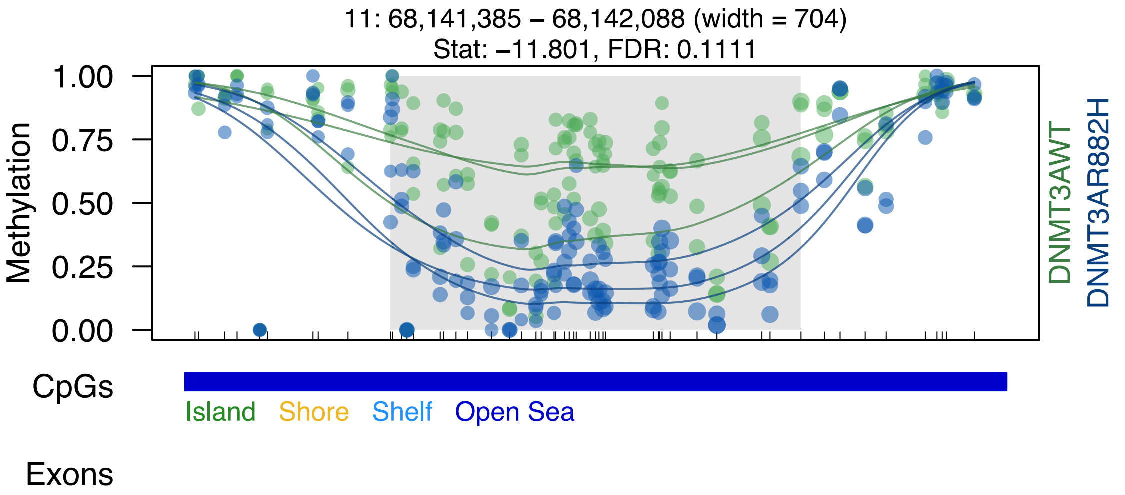

11: 68,141,385 – 68,142,088 (width = 704)
Stat: –11.801, FDR: 0.1111
Methylation
1.00
0.75
0.50
0.25
0.00
DNMT3AWT
DNMT3AR882H
CpGs
Island
Shore
Shelf
Open Sea
Exons

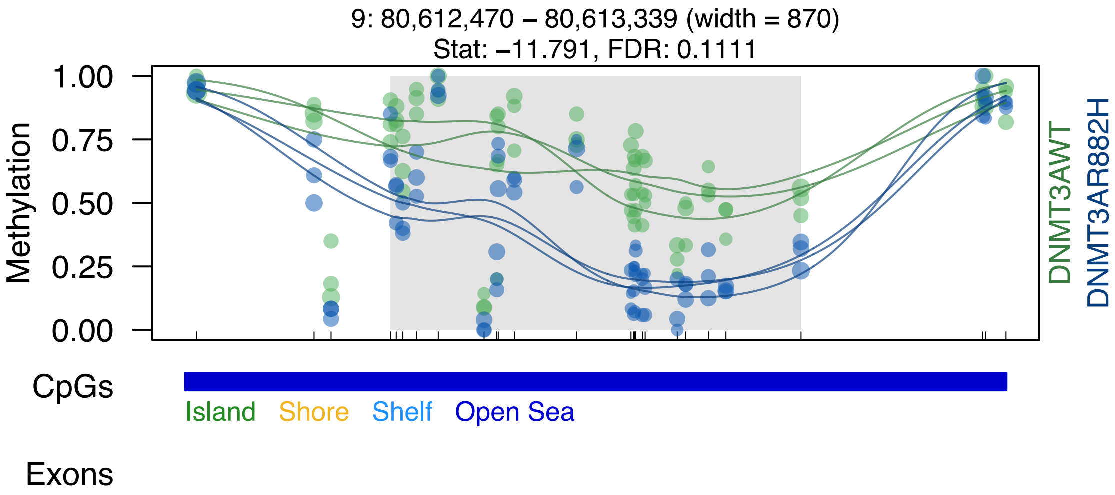
9: 80,612,470 – 80,613,339 (width = 870)
Stat: −11.791, FDR: 0.1111
Methylation
1.00
0.75
0.50
0.25
0.00
DNMT3AWT
DNMT3AR882H
CpGs
Island
Shore
Shelf
Open Sea
Exons

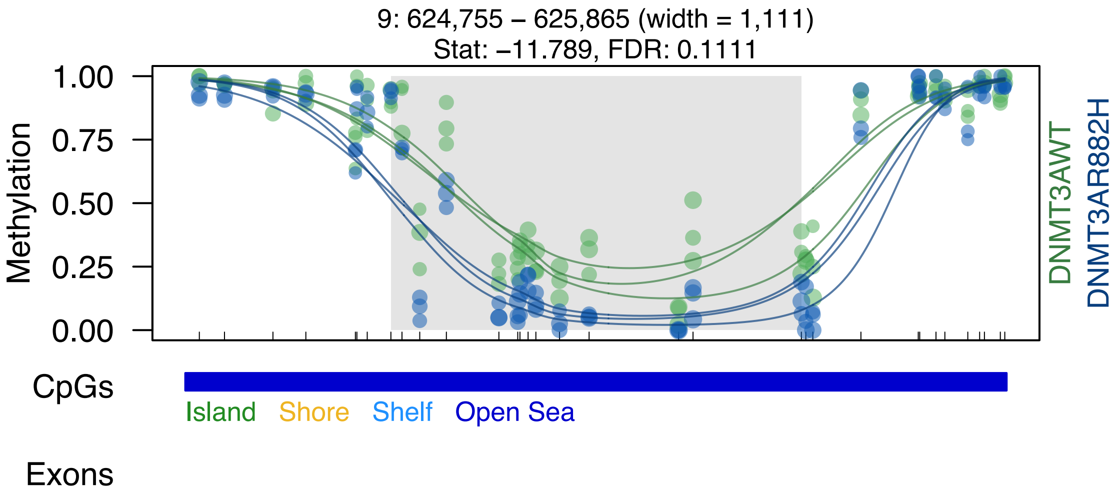

9: 624,755 – 625,865 (width = 1,111)
Stat: –11.789, FDR: 0.1111
Methylation
1.00
0.75
0.50
0.25
0.00
DNMT3AWT
DNMT3AR882H
CpGs
Island
Shore
Shelf
Open Sea
Exons

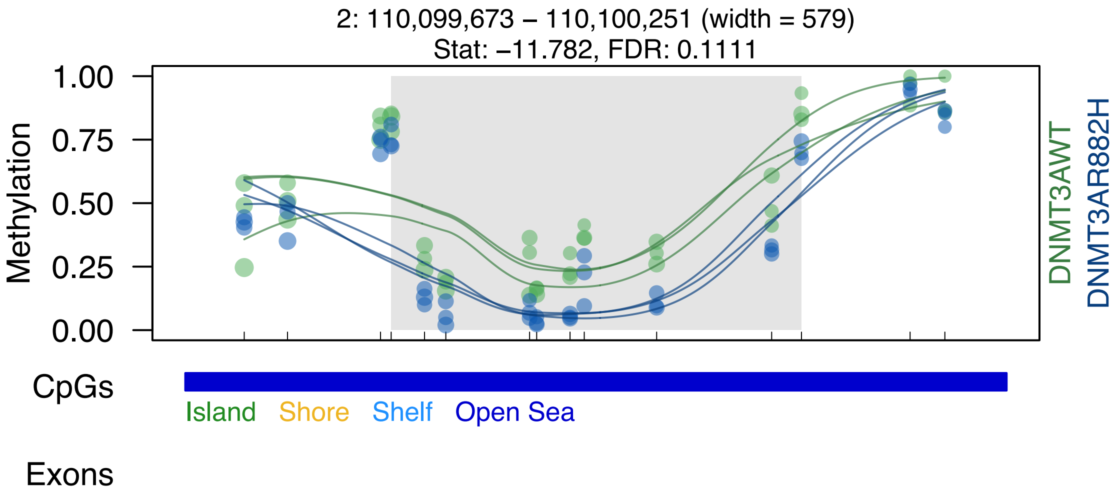

2: 110,099,673 – 110,100,251 (width = 579)
Stat: –11.782, FDR: 0.1111
Methylation
1.00
0.75
0.50
0.25
0.00
DNMT3AWT
DNMT3AR882H
CpGs
Island
Shore
Shelf
Open Sea
Exons

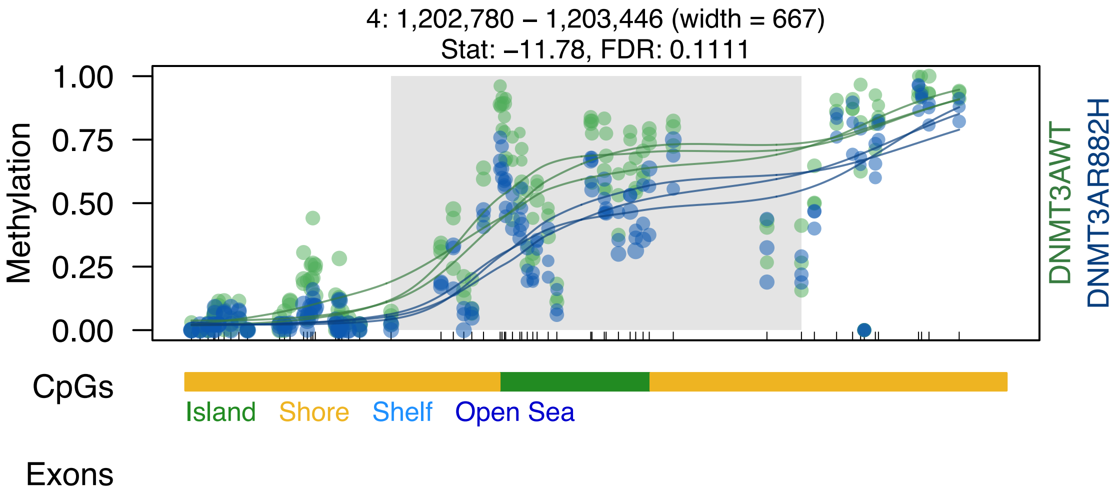

4: 1,202,780 – 1,203,446 (width = 667)
Stat: –11.78, FDR: 0.1111
Methylation
1.00
0.75
0.50
0.25
0.00
DNMT3AWT
DNMT3AR882H
CpGs
Island
Shore
Shelf
Open Sea
Exons

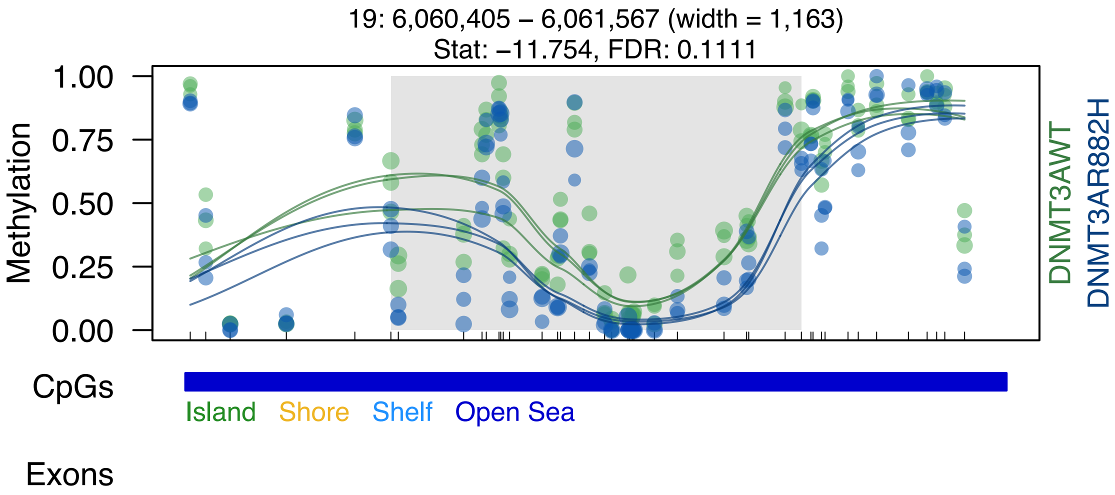

19: 6,060,405 – 6,061,567 (width = 1,163)
Stat: −11.754, FDR: 0.1111
Methylation
1.00
0.75
0.50
0.25
0.00
DNMT3AWT
DNMT3AR882H
CpGs
Island
Shore
Shelf
Open Sea
Exons

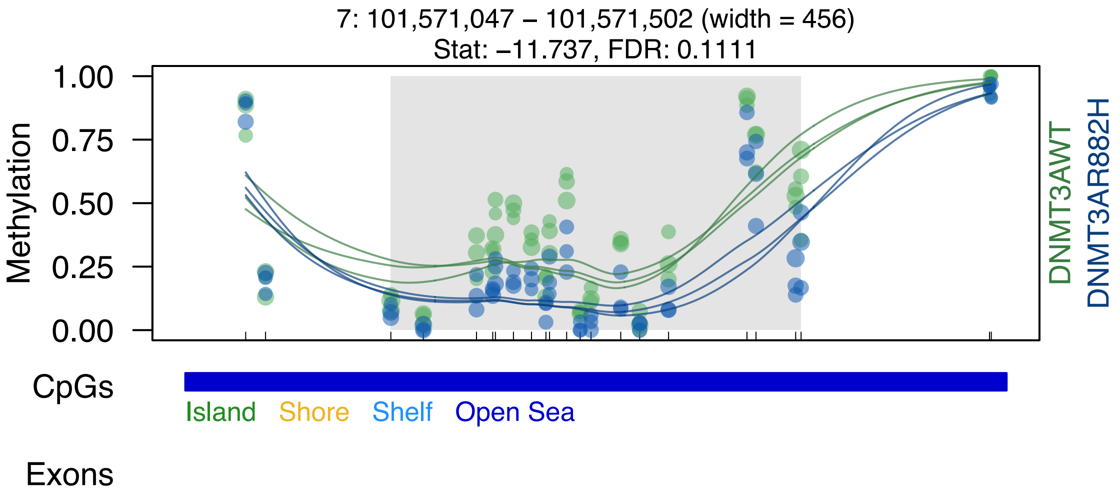

7: 101,571,047 – 101,571,502 (width = 456)
Stat: −11.737, FDR: 0.1111
Methylation
1.00
0.75
0.50
0.25
0.00
DNMT3AWT
DNMT3AR882H
CpGs
Island
Shore
Shelf
Open Sea
Exons

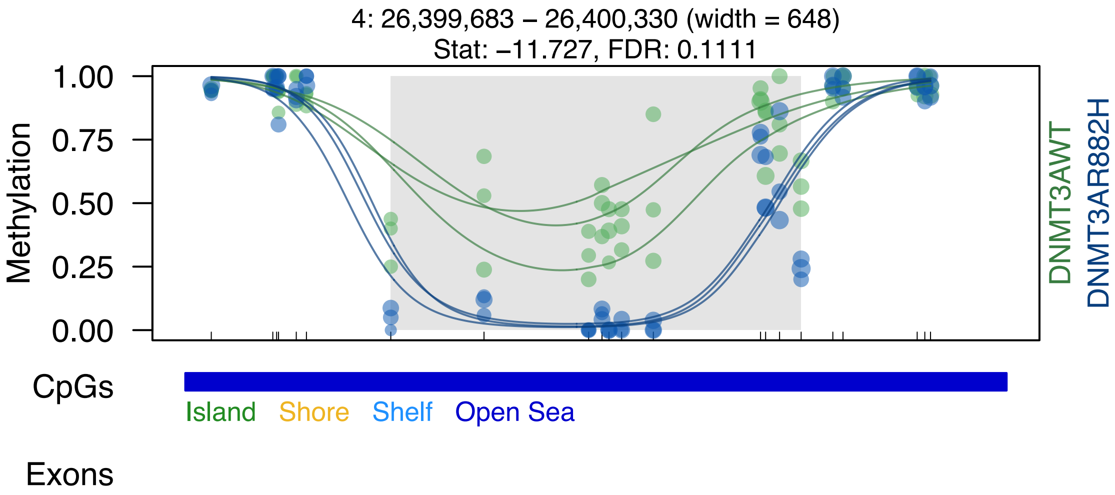

4: 26,399,683 – 26,400,330 (width = 648)
Stat: –11.727, FDR: 0.1111
Methylation
1.00
0.75
0.50
0.25
0.00
DNMT3AWT
DNMT3AR882H
CpGs
Island
Shore
Shelf
Open Sea
Exons

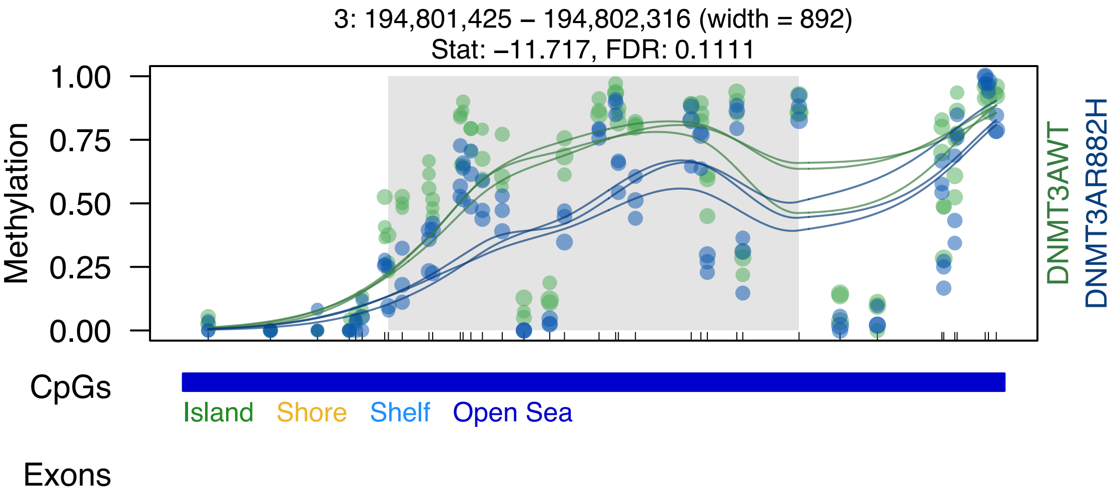

3: 194,801,425 – 194,802,316 (width = 892)
Stat: –11.717, FDR: 0.1111
Methylation
1.00
0.75
0.50
0.25
0.00
DNMT3AWT
DNMT3AR882H
CpGs
Island
Shore
Shelf
Open Sea
Exons

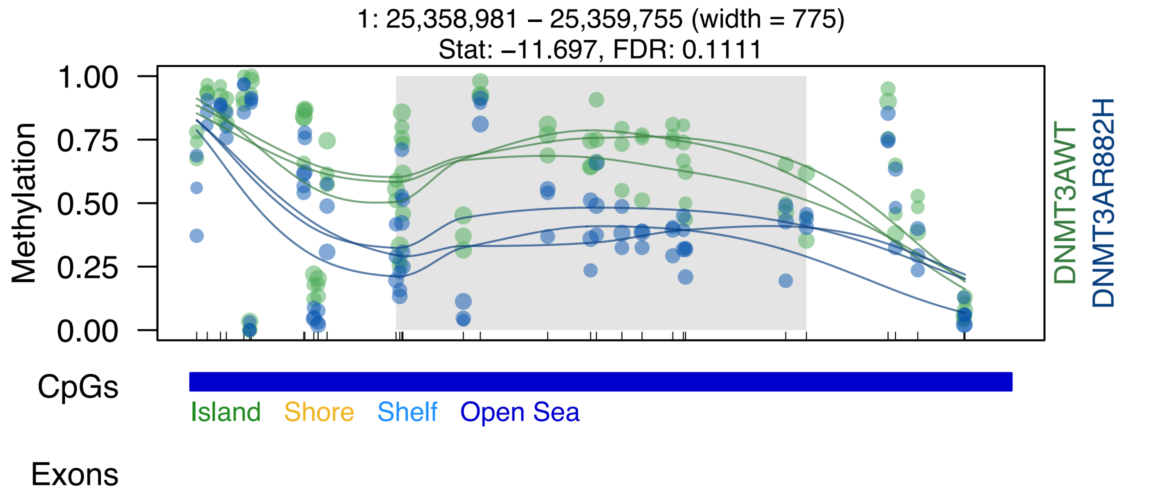
1: 25,358,981 – 25,359,755 (width = 775)
Stat: –11.697, FDR: 0.1111
Methylation
1.00
0.75
0.50
0.25
0.00
DNMT3AWT
DNMT3AR882H
CpGs
Island
Shore
Shelf
Open Sea
Exons

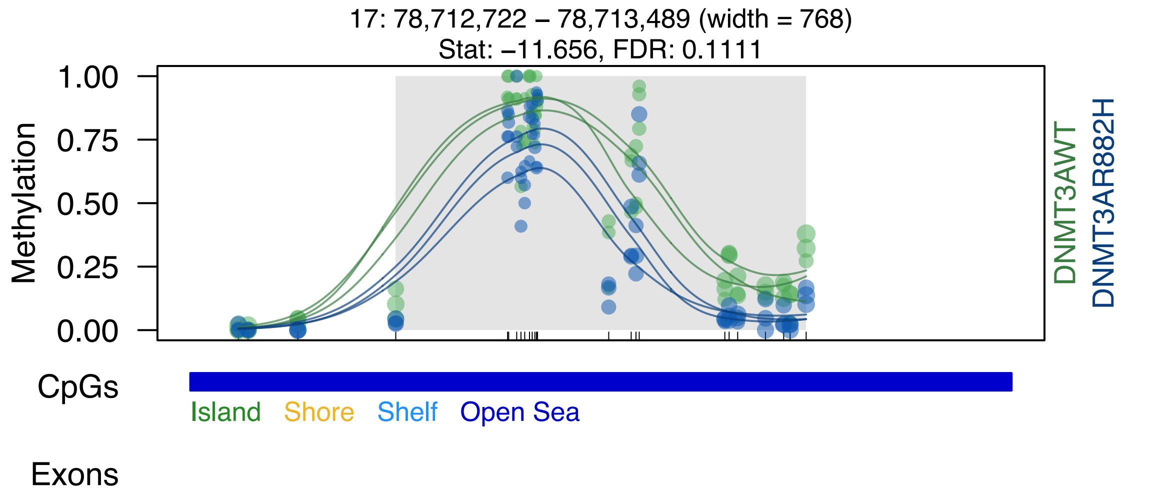
17: 78,712,722 – 78,713,489 (width = 768)
Stat: –11.656, FDR: 0.1111
Methylation
1.00
0.75
0.50
0.25
0.00
DNMT3AWT
DNMT3AR882H
CpGs
Island
Shore
Shelf
Open Sea
Exons

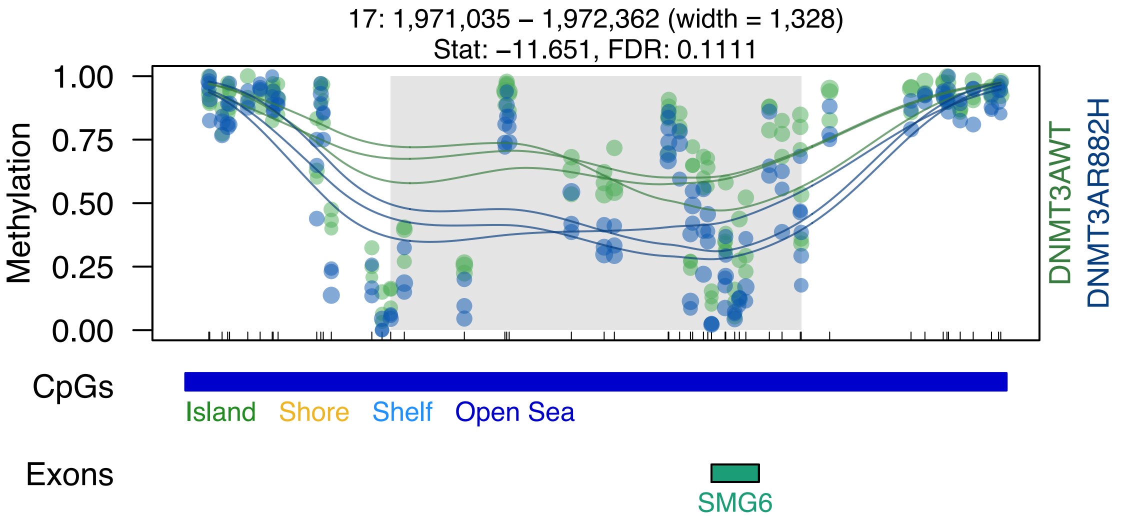
17: 1,971,035 – 1,972,362 (width = 1,328)
Stat: −11.651, FDR: 0.1111
Methylation
1.00
0.75
0.50
0.25
0.00
DNMT3AWT
DNMT3AR882H
CpGs
Island
Shore
Shelf
Open Sea
Exons
SMG6

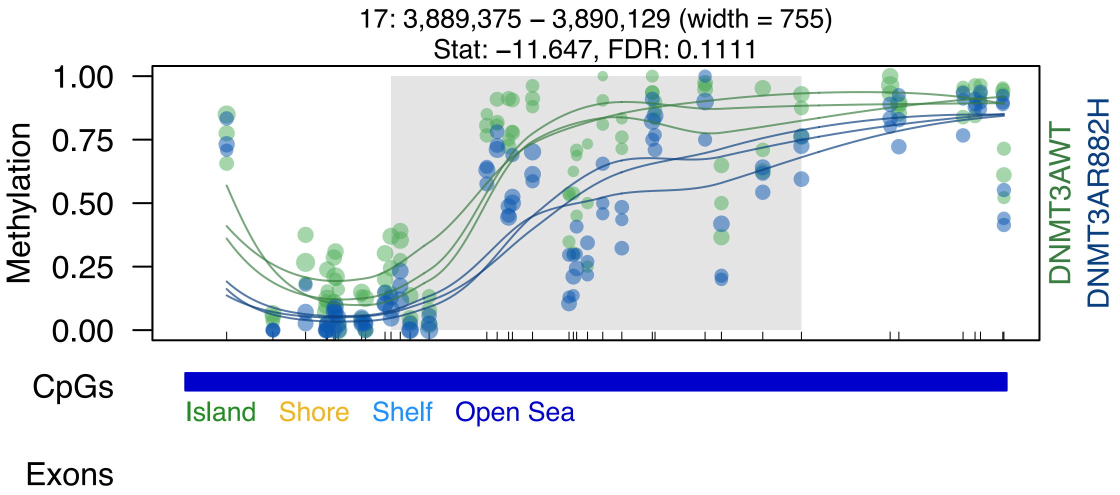
17: 3,889,375 – 3,890,129 (width = 755)
Stat: –11.647, FDR: 0.1111
Methylation
1.00
0.75
0.50
0.25
0.00
DNMT3AWT
DNMT3AR882H
CpGs
Island
Shore
Shelf
Open Sea
Exons

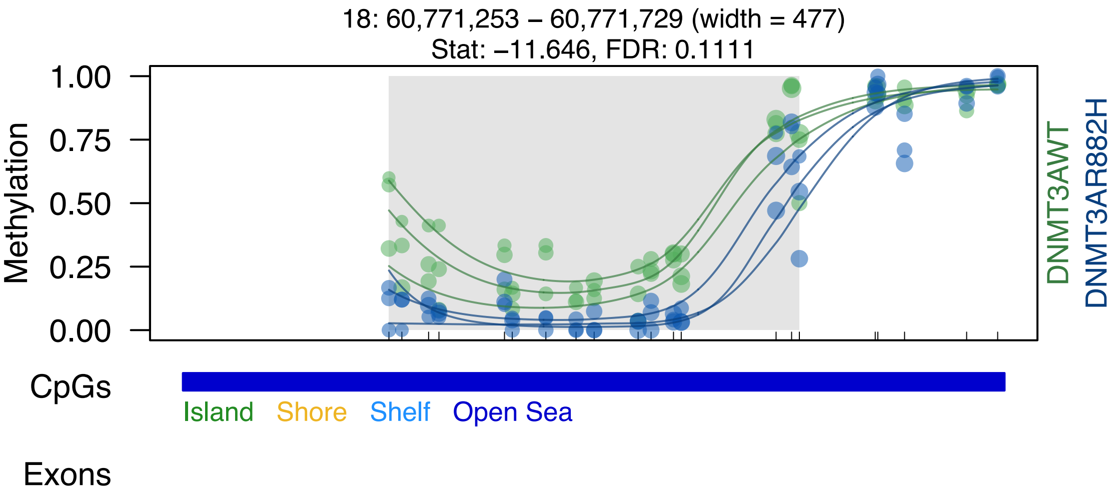

18: 60,771,253 – 60,771,729 (width = 477)
Stat: –11.646, FDR: 0.1111
Methylation
1.00
0.75
0.50
0.25
0.00
DNMT3AWT
DNMT3AR882H
CpGs
Island
Shore
Shelf
Open Sea
Exons

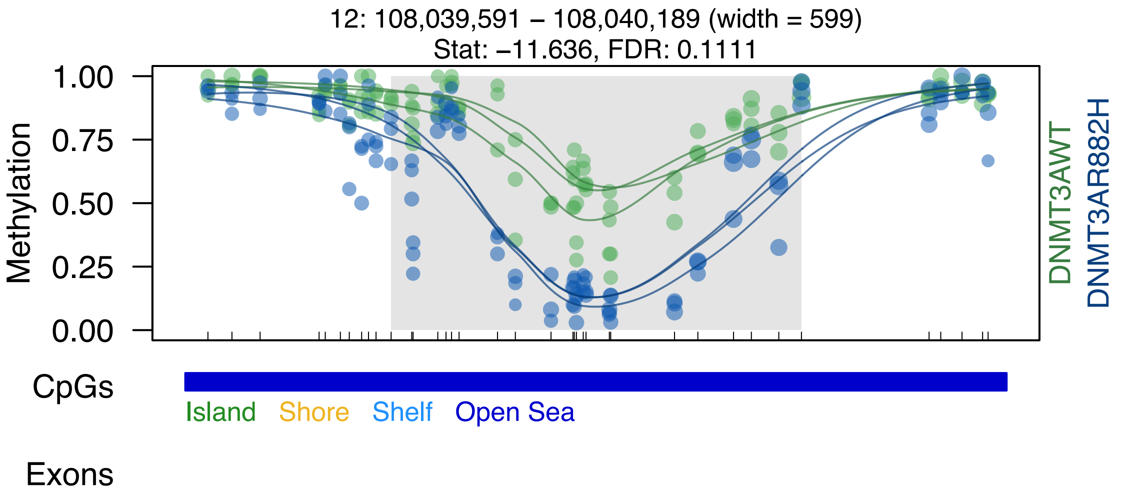

12: 108,039,591 – 108,040,189 (width = 599)
Stat: –11.636, FDR: 0.1111
1.00
0.75
0.50
0.25
0.00
Methylation
DNMT3AWT
DNMT3AR882H
CpGs
Island
Shore
Shelf
Open Sea
Exons

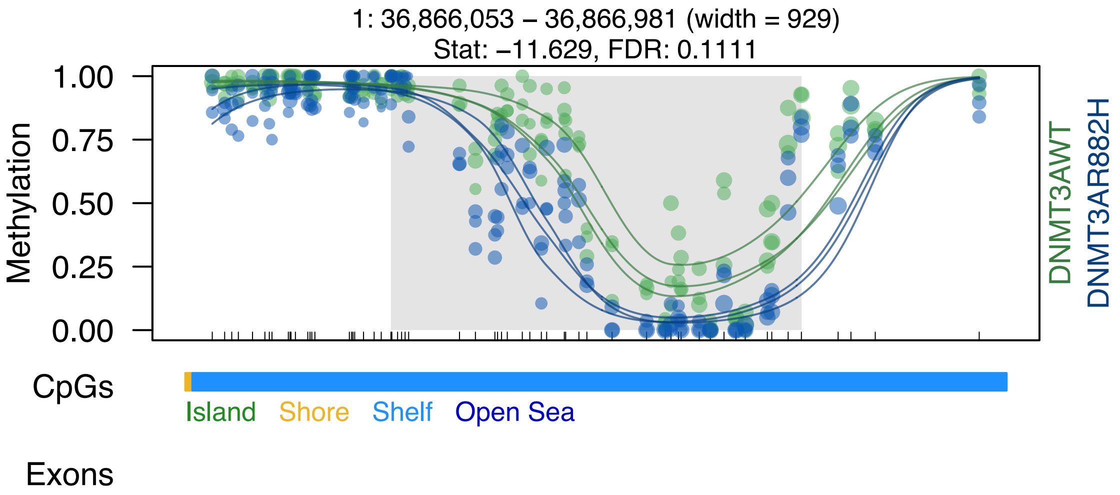

1: 36,866,053 – 36,866,981 (width = 929)
Stat: −11.629, FDR: 0.1111
Methylation
1.00
0.75
0.50
0.25
0.00
DNMT3AWT
DNMT3AR882H
CpGs
Island
Shore
Shelf
Open Sea
Exons

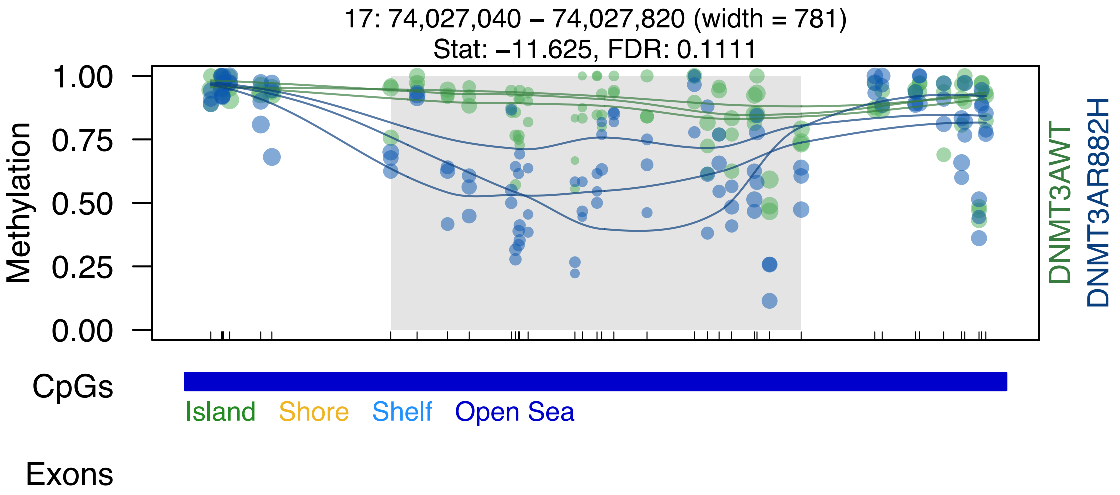

17: 74,027,040 – 74,027,820 (width = 781)
Stat: –11.625, FDR: 0.1111
1.00
0.75
0.50
0.25
0.00
Methylation
DNMT3AWT
DNMT3AR882H
CpGs
Island
Shore
Shelf
Open Sea
Exons

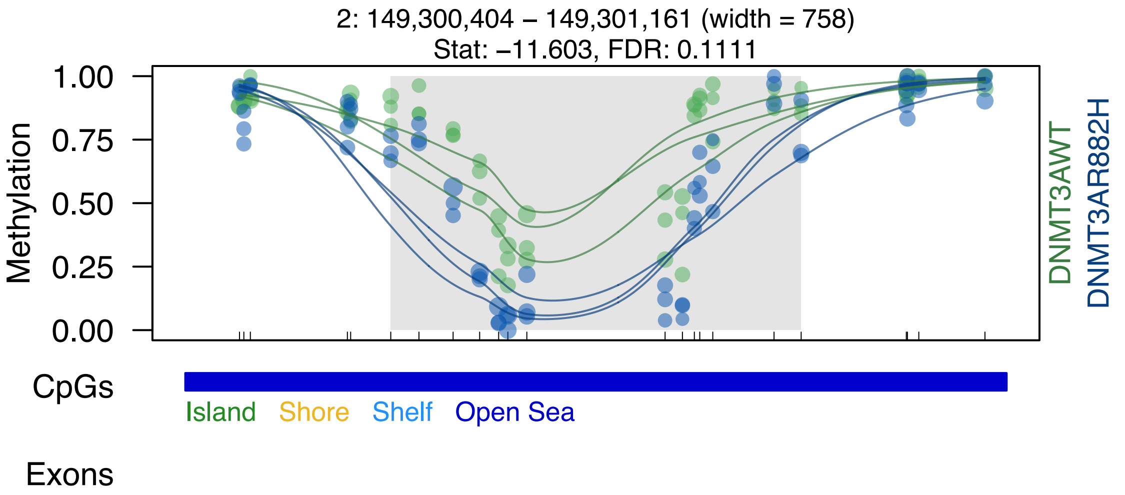
2: 149,300,404 – 149,301,161 (width = 758)
Stat: –11.603, FDR: 0.1111
Methylation
1.00
0.75
0.50
0.25
0.00
DNMT3AWT
DNMT3AR882H
CpGs
Island
Shore
Shelf
Open Sea
Exons

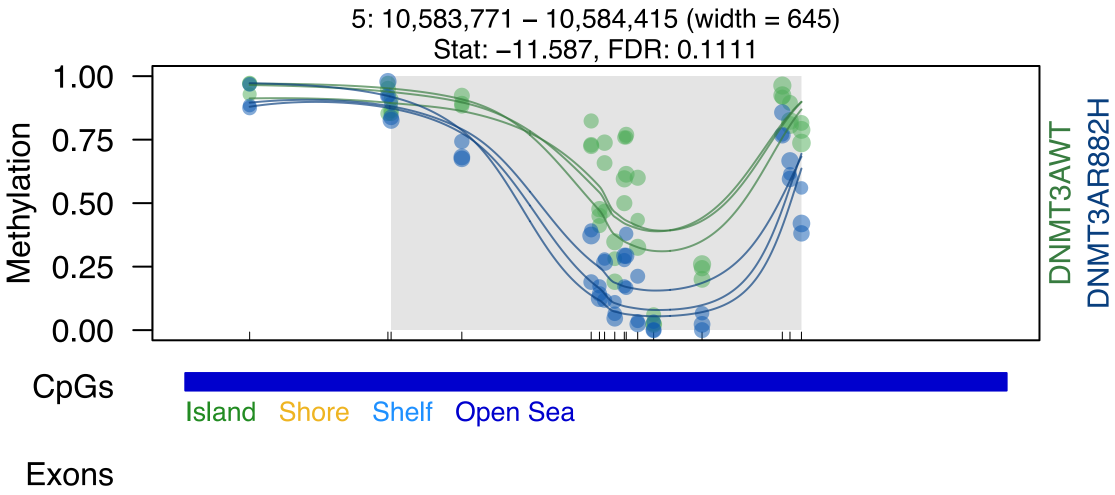

5: 10,583,771 – 10,584,415 (width = 645)
Stat: –11.587, FDR: 0.1111
Methylation
1.00
0.75
0.50
0.25
0.00
DNMT3AWT
DNMT3AR882H
CpGs
Island
Shore
Shelf
Open Sea
Exons

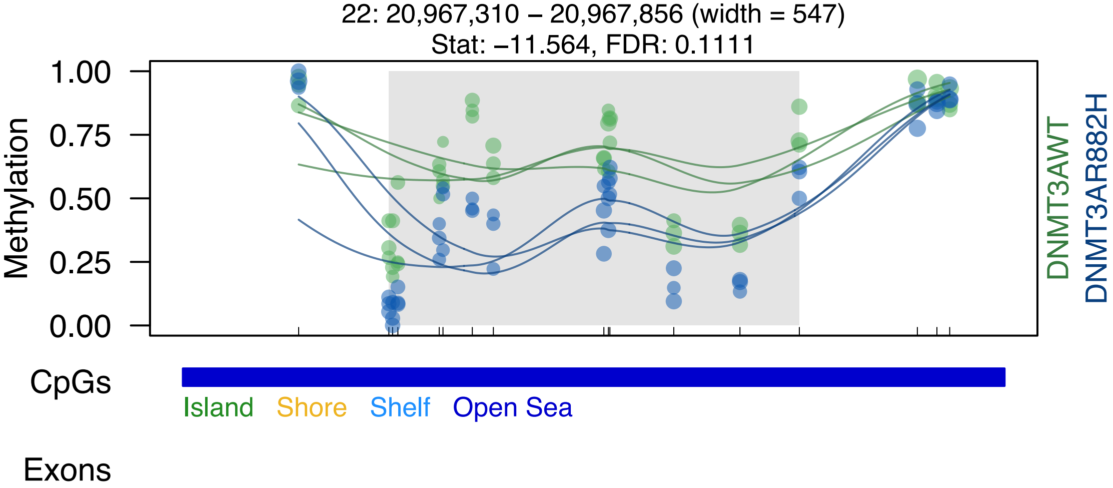
22: 20,967,310 – 20,967,856 (width = 547)
Stat: –11.564, FDR: 0.1111
Methylation
1.00
0.75
0.50
0.25
0.00
DNMT3AWT
DNMT3AR882H
CpGs
Island
Shore
Shelf
Open Sea
Exons

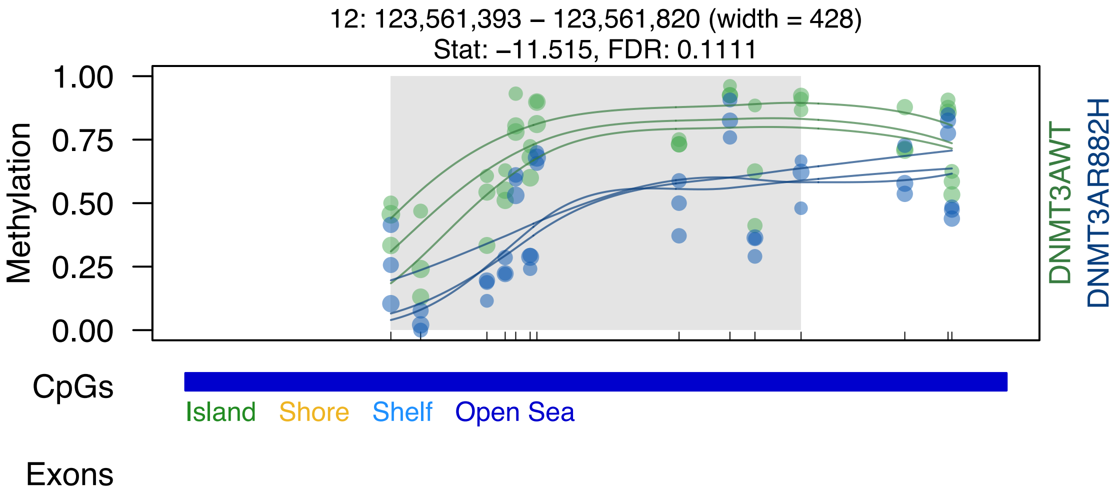
12: 123,561,393 – 123,561,820 (width = 428)
Stat: −11.515, FDR: 0.1111
Methylation
1.00
0.75
0.50
0.25
0.00
DNMT3AWT
DNMT3AR882H
CpGs
Island
Shore
Shelf
Open Sea
Exons

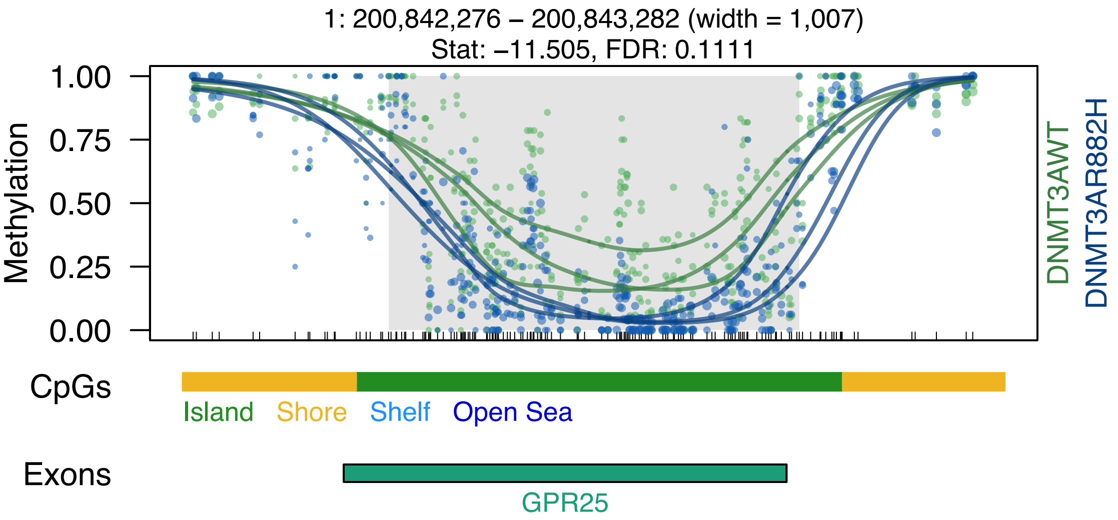
1: 200,842,276 – 200,843,282 (width = 1,007)
Stat: −11.505, FDR: 0.1111
1.00
0.75
0.50
0.25
0.00
Methylation
DNMT3AWT
DNMT3AR882H
CpGs
Island
Shore
Shelf
Open Sea
Exons
GPR25

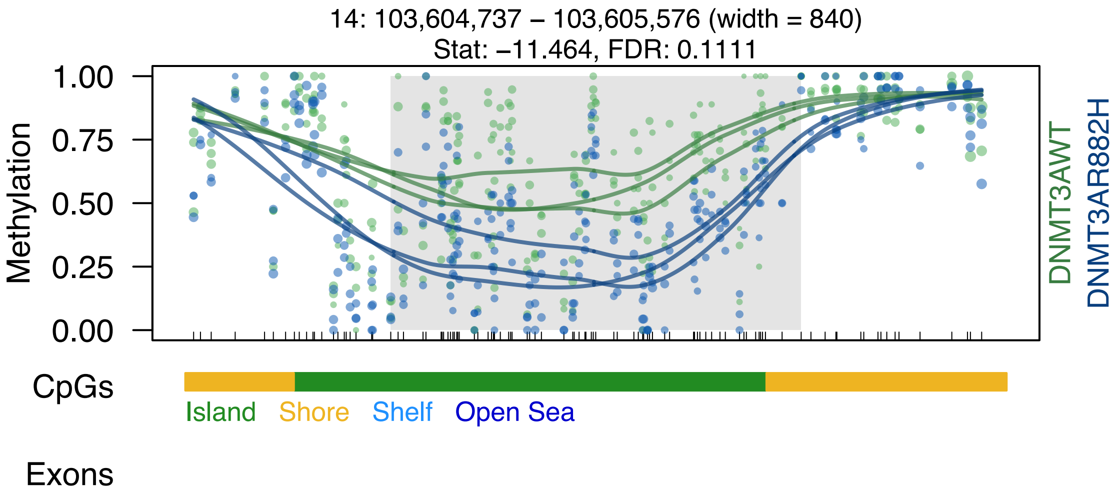

14: 103,604,737 – 103,605,576 (width = 840)
Stat: –11.464, FDR: 0.1111
Methylation
1.00
0.75
0.50
0.25
0.00
DNMT3AWT
DNMT3AR882H
CpGs
Island
Shore
Shelf
Open Sea
Exons

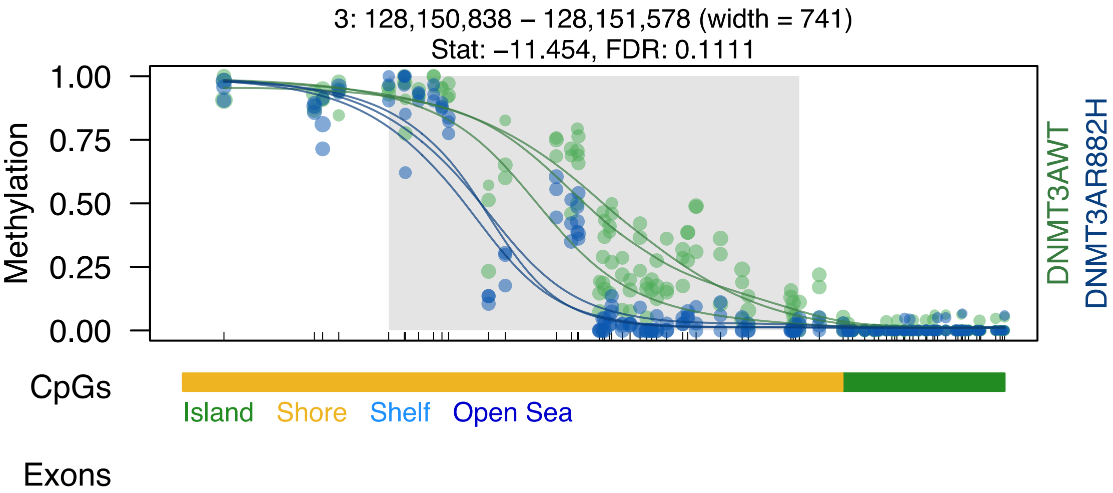
3: 128,150,838 – 128,151,578 (width = 741)
Stat: –11.454, FDR: 0.1111
Methylation
1.00
0.75
0.50
0.25
0.00
DNMT3AWT
DNMT3AR882H
CpGs
Island
Shore
Shelf
Open Sea
Exons

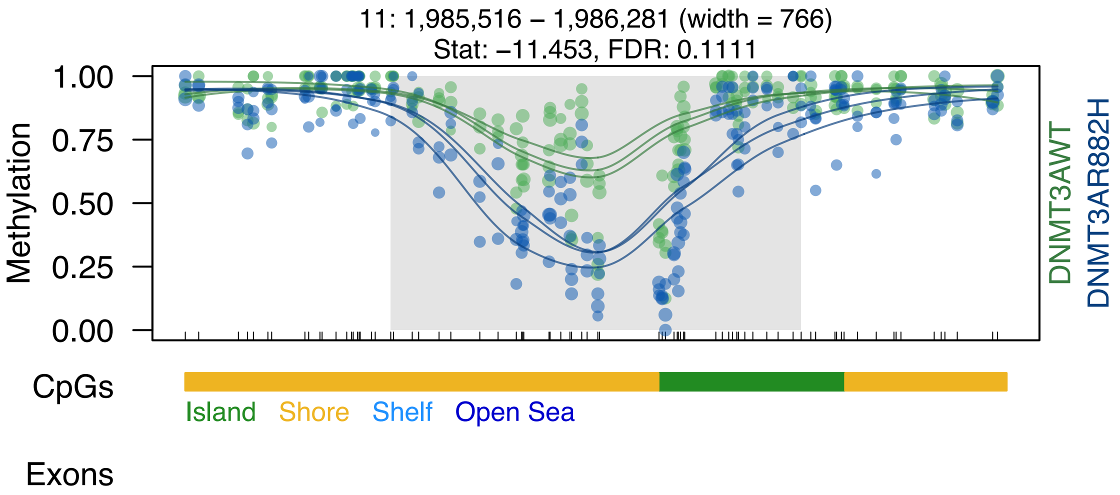
11: 1,985,516 – 1,986,281 (width = 766)
Stat: –11.453, FDR: 0.1111
Methylation
1.00
0.75
0.50
0.25
0.00
DNMT3AWT
DNMT3AR882H
CpGs
Island
Shore
Shelf
Open Sea
Exons

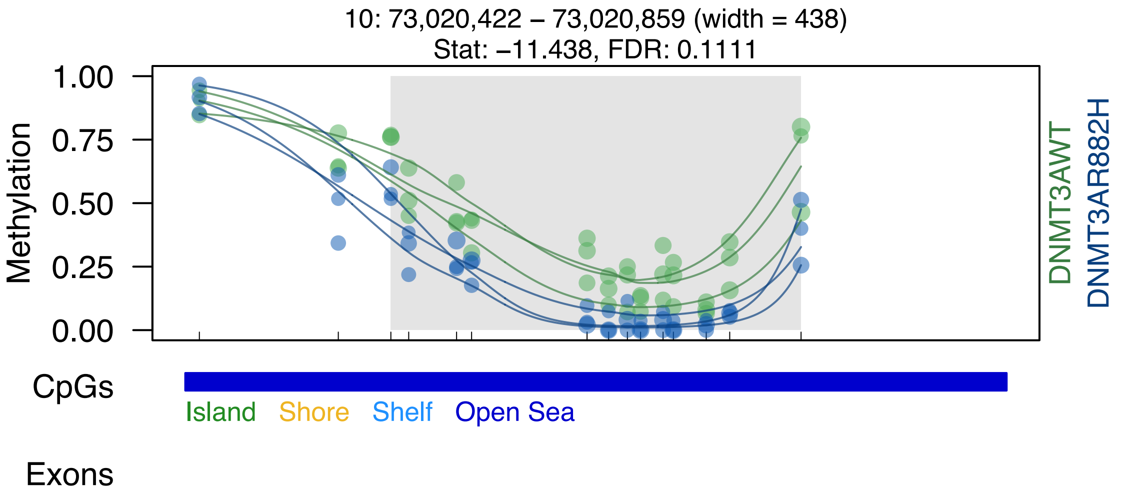

10: 73,020,422 – 73,020,859 (width = 438)
Stat: –11.438, FDR: 0.1111
Methylation
1.00
0.75
0.50
0.25
0.00
DNMT3AWT
DNMT3AR882H
CpGs
Island
Shore
Shelf
Open Sea
Exons

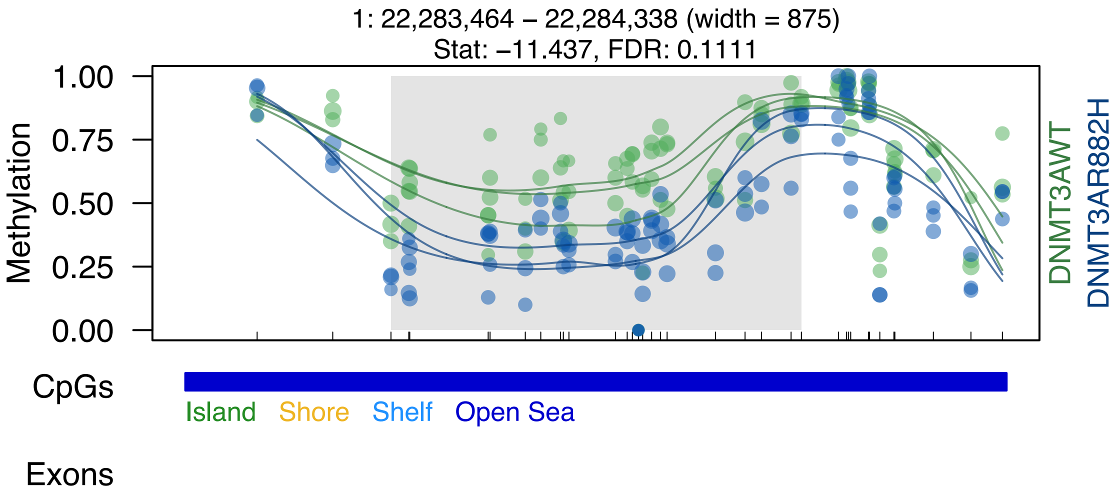

1: 22,283,464 – 22,284,338 (width = 875)
Stat: –11.437, FDR: 0.1111
Methylation
1.00
0.75
0.50
0.25
0.00
DNMT3AWT
DNMT3AR882H
CpGs
Island
Shore
Shelf
Open Sea
Exons

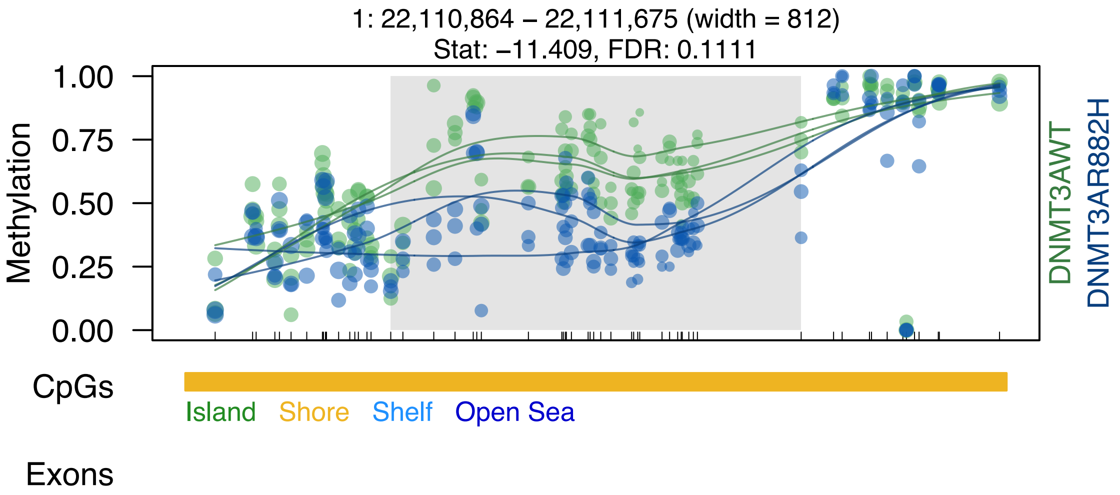
1: 22,110,864 – 22,111,675 (width = 812)
Stat: –11.409, FDR: 0.1111
Methylation
1.00
0.75
0.50
0.25
0.00
DNMT3AWT
DNMT3AR882H
CpGs
Island
Shore
Shelf
Open Sea
Exons

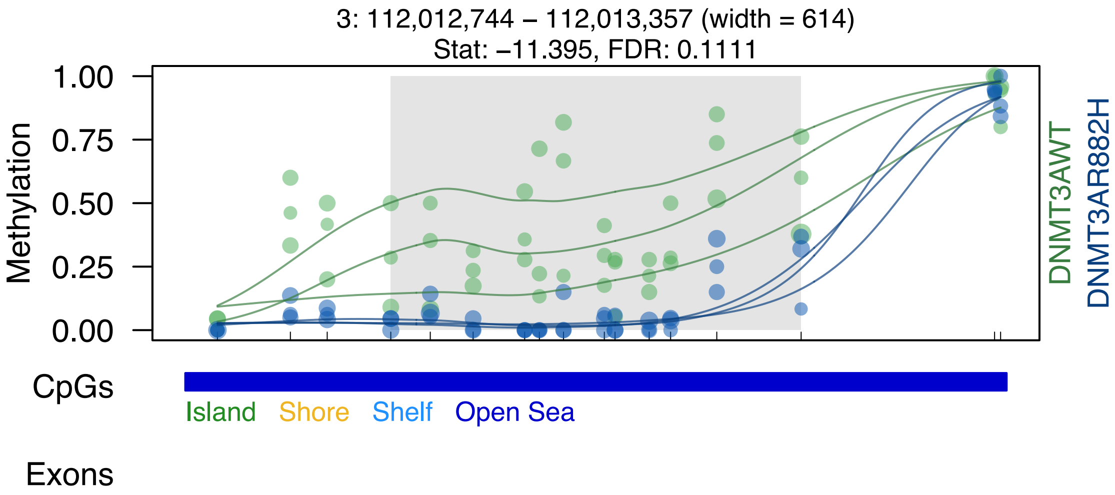
3: 112,012,744 – 112,013,357 (width = 614)
Stat: −11.395, FDR: 0.1111
Methylation
1.00
0.75
0.50
0.25
0.00
DNMT3AWT
DNMT3AR882H
CpGs
Island
Shore
Shelf
Open Sea
Exons

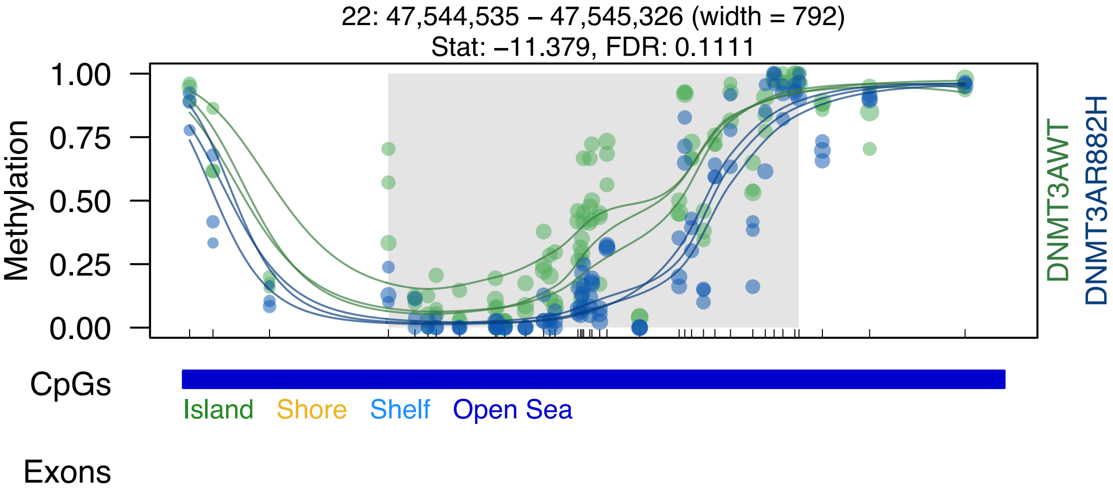

22: 47,544,535 – 47,545,326 (width = 792)
Stat: –11.379, FDR: 0.1111
Methylation
1.00
0.75
0.50
0.25
0.00
DNMT3AWT
DNMT3AR882H
CpGs
Island
Shore
Shelf
Open Sea
Exons

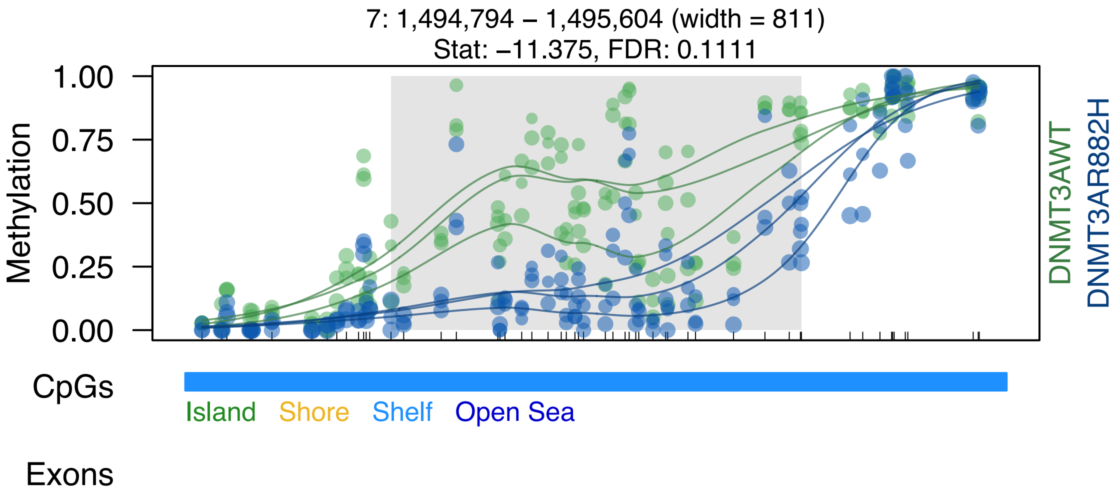
7: 1,494,794 – 1,495,604 (width = 811)
Stat: –11.375, FDR: 0.1111
Methylation
1.00
0.75
0.50
0.25
0.00
DNMT3AWT
DNMT3AR882H
CpGs
Island
Shore
Shelf
Open Sea
Exons

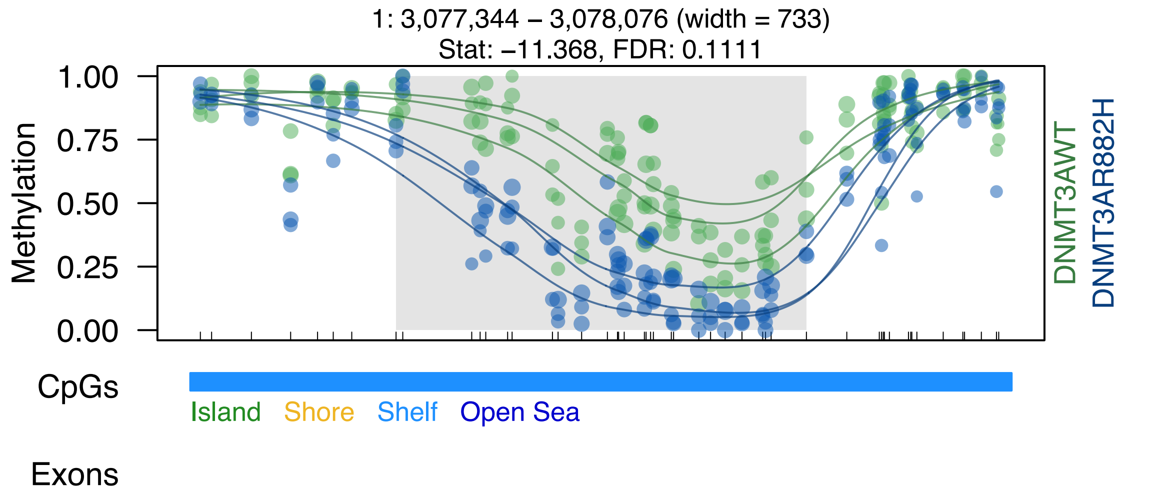
1: 3,077,344 – 3,078,076 (width = 733)
Stat: –11.368, FDR: 0.1111
1.00
0.75
0.50
0.25
0.00
Methylation
DNMT3AWT
DNMT3AR882H
CpGs
Island
Shore
Shelf
Open Sea
Exons

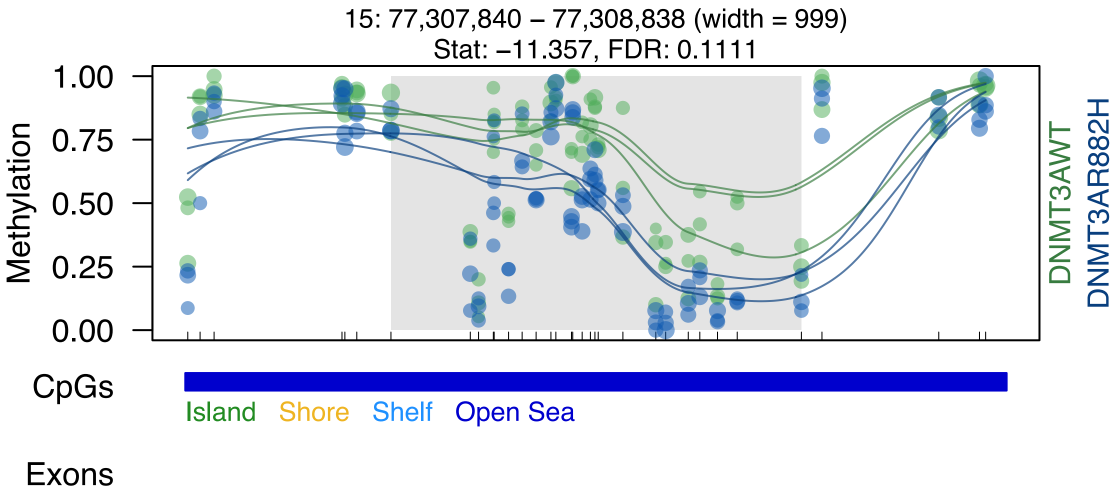
15: 77,307,840 – 77,308,838 (width = 999)
Stat: –11.357, FDR: 0.1111
Methylation
1.00
0.75
0.50
0.25
0.00
DNMT3AWT
DNMT3AR882H
CpGs
Island
Shore
Shelf
Open Sea
Exons

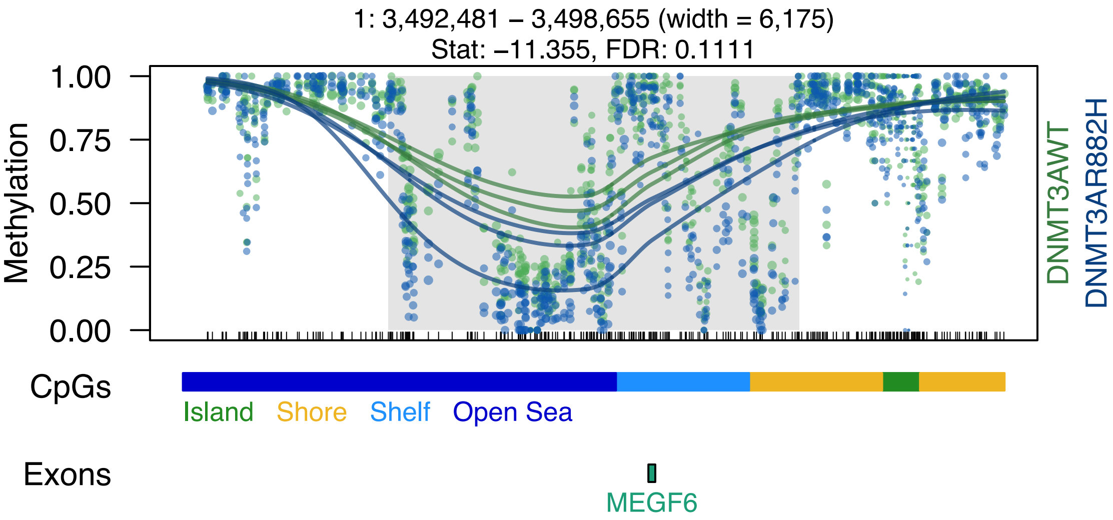

1: 3,492,481 – 3,498,655 (width = 6,175)
Stat: –11.355, FDR: 0.1111
Methylation
1.00
0.75
0.50
0.25
0.00
DNMT3AWT
DNMT3AR882H
CpGs
Island
Shore
Shelf
Open Sea
Exons
MEGF6

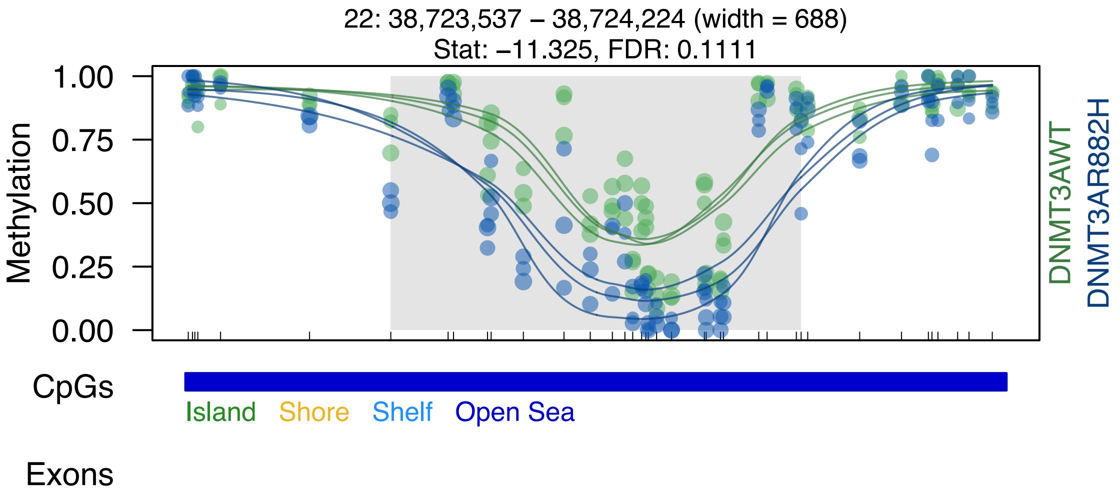
22: 38,723,537 – 38,724,224 (width = 688)
Stat: –11.325, FDR: 0.1111
1.00
0.75
0.50
0.25
0.00
Methylation
DNMT3AWT
DNMT3AR882H
CpGs
Island
Shore
Shelf
Open Sea
Exons

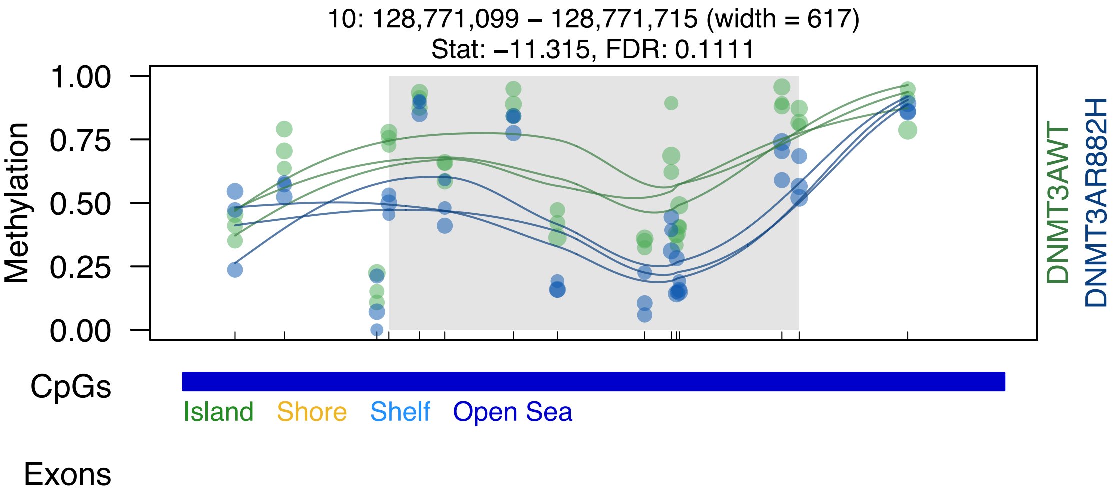

10: 128,771,099 – 128,771,715 (width = 617)
Stat: –11.315, FDR: 0.1111
Methylation
1.00
0.75
0.50
0.25
0.00
DNMT3AWT
DNMT3AR882H
CpGs
Island
Shore
Shelf
Open Sea
Exons

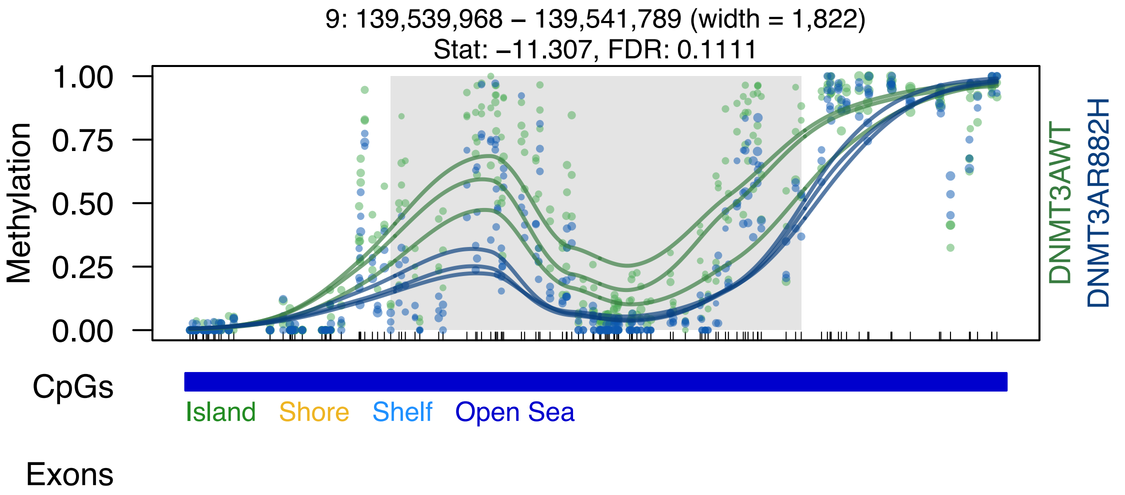
9: 139,539,968 – 139,541,789 (width = 1,822)
Stat: −11.307, FDR: 0.1111
Methylation
1.00
0.75
0.50
0.25
0.00
DNMT3AWT
DNMT3AR882H
CpGs
Island
Shore
Shelf
Open Sea
Exons

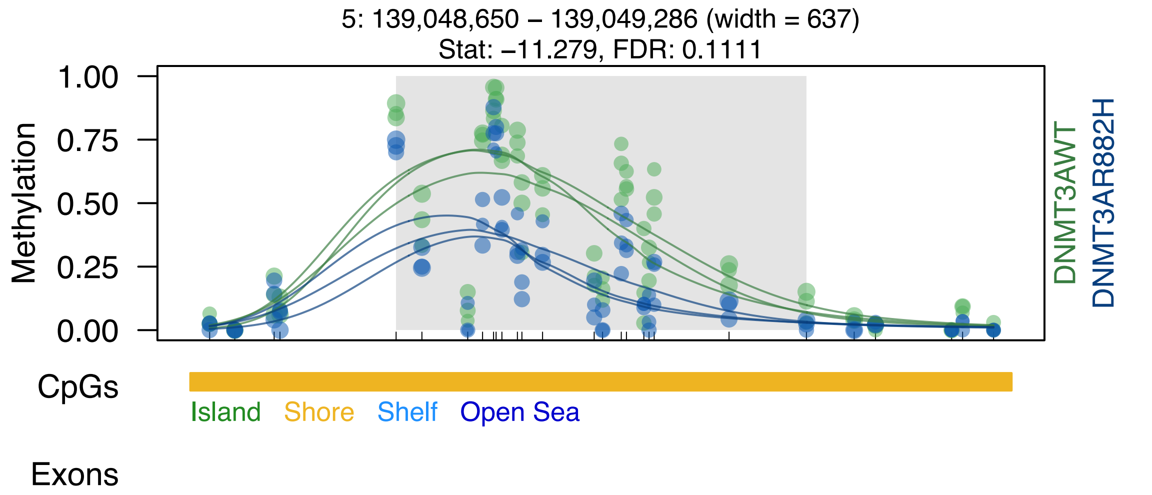

5: 139,048,650 – 139,049,286 (width = 637)
Stat: –11.279, FDR: 0.1111
1.00
0.75
0.50
0.25
0.00
Methylation
DNMT3AWT
DNMT3AR882H
CpGs
Island
Shore
Shelf
Open Sea
Exons

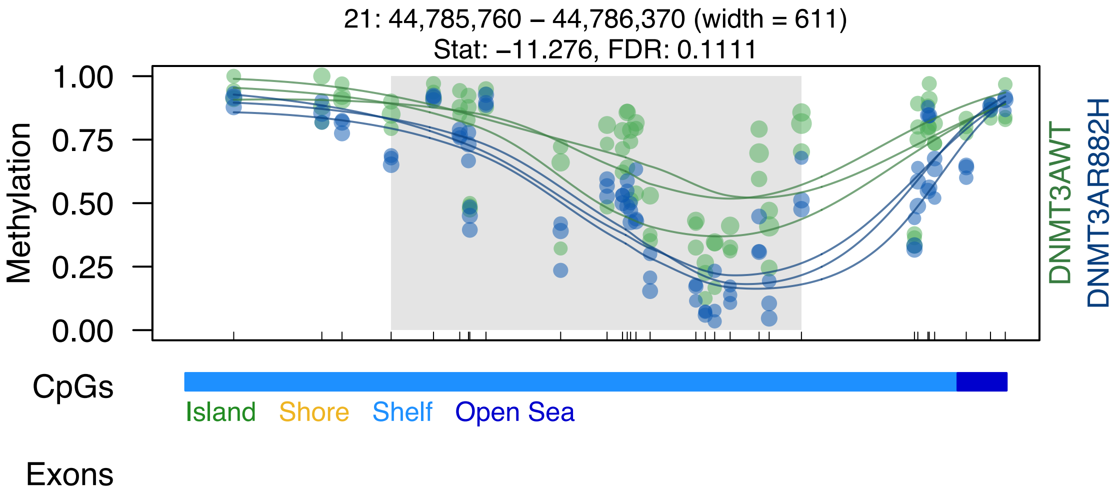

21: 44,785,760 – 44,786,370 (width = 611)
Stat: –11.276, FDR: 0.1111
Methylation
1.00
0.75
0.50
0.25
0.00
DNMT3AWT
DNMT3AR882H
CpGs
Island
Shore
Shelf
Open Sea
Exons
