## Supplemental Data S1-S7 for "DNMT3A^R882H^ Is Not Required for Disease Maintenance in Primary Human AML, but Is Associated With Increased Leukemia Stem Cell Frequency": DataS4_SU372C_top50dmrsOverlappingCanyons.pdf

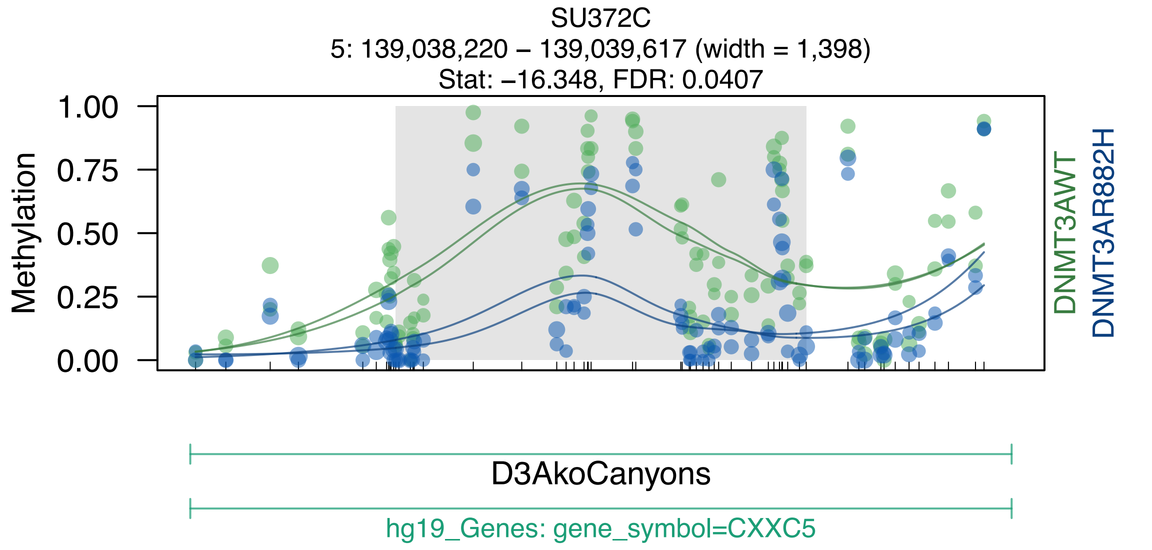

SU372C
5: 139,038,220 – 139,039,617 (width = 1,398)
Stat: –16.348, FDR: 0.0407
Methylation
1.00
0.75
0.50
0.25
0.00
DNMT3AWT
DNMT3AR882H
D3AkoCanyons
hg19_Genes: gene_symbol=CXXC5

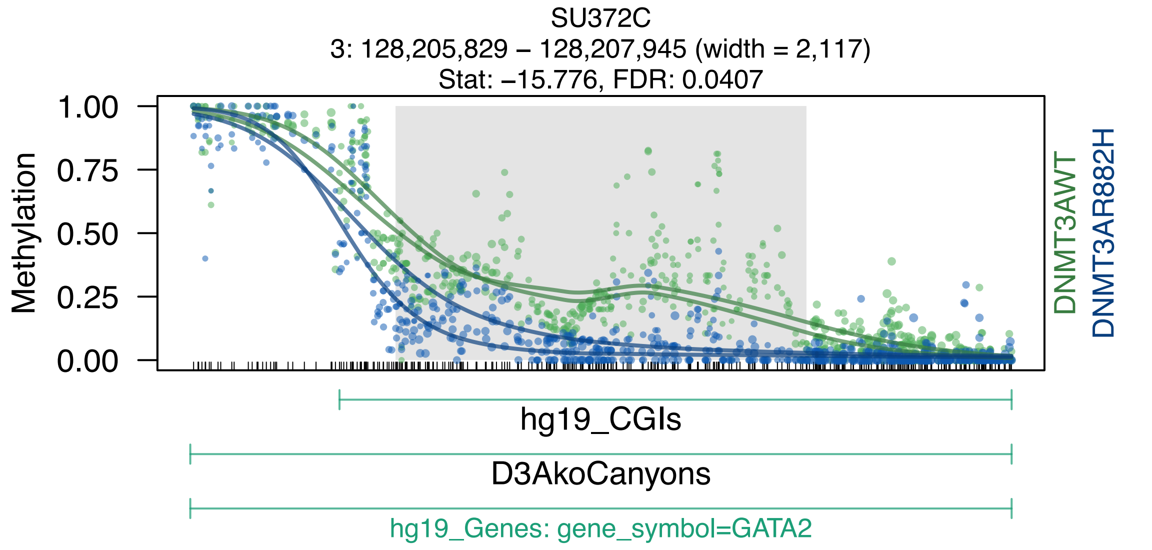
SU372C
3: 128,205,829 – 128,207,945 (width = 2,117)
Stat: –15.776, FDR: 0.0407
Methylation
1.00
0.75
0.50
0.25
0.00
DNMT3AWT
DNMT3AR882H
hg19_CGIs
D3AkoCanyons
hg19_Genes: gene_symbol=GATA2

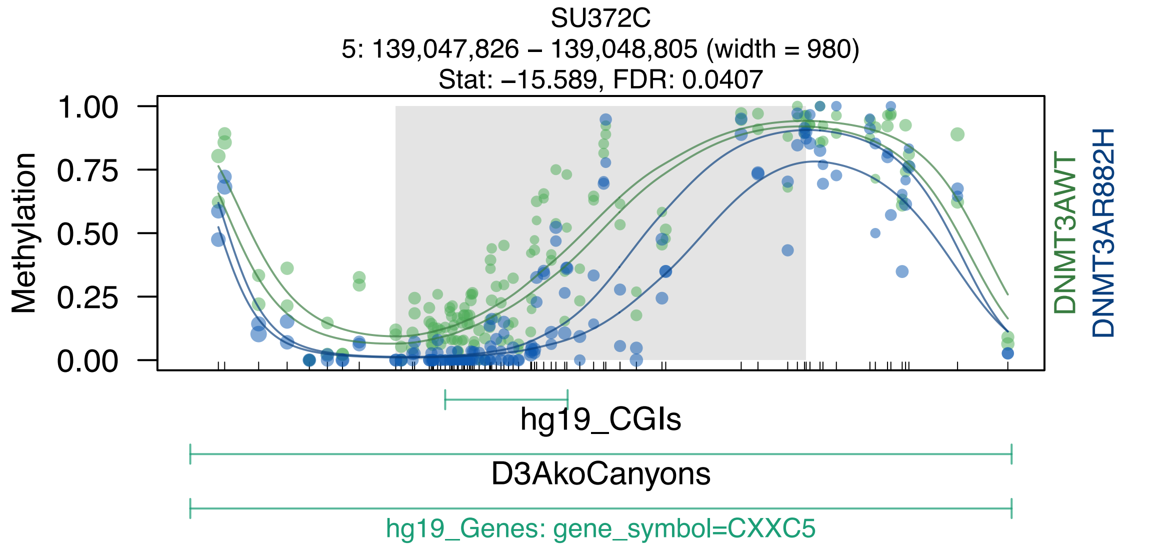
SU372C
5: 139,047,826 – 139,048,805 (width = 980)
Stat: –15.589, FDR: 0.0407
Methylation
1.00
0.75
0.50
0.25
0.00
DNMT3AWT
DNMT3AR882H
hg19_CGIs
D3AkoCanyons
hg19_Genes: gene_symbol=CXXC5

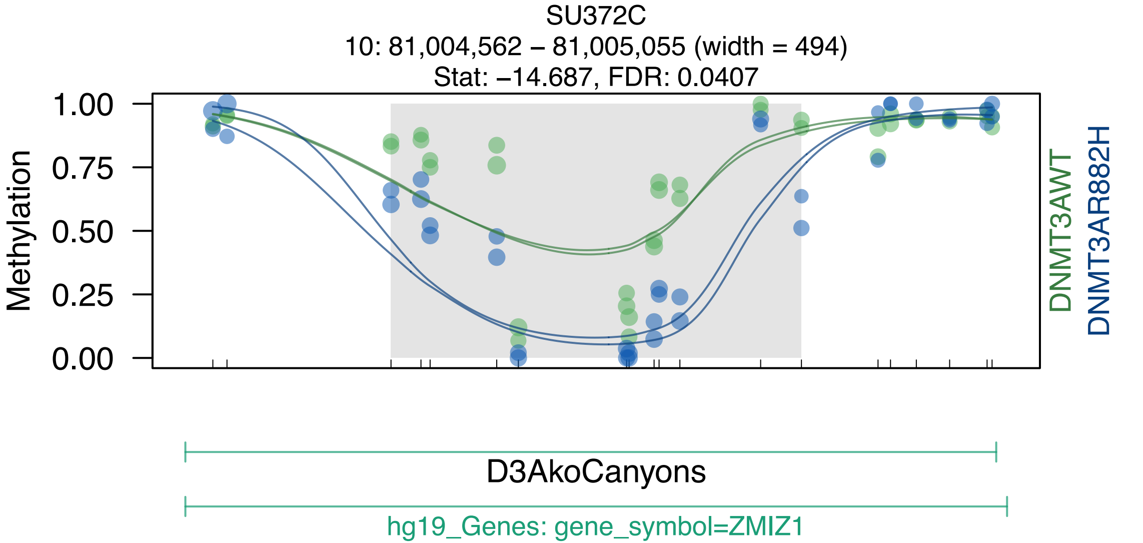
SU372C
10: 81,004,562 – 81,005,055 (width = 494)
Stat: −14.687, FDR: 0.0407
Methylation
1.00
0.75
0.50
0.25
0.00
DNMT3AWT
DNMT3AR882H
D3AkoCanyons
hg19_Genes: gene_symbol=ZMIZ1

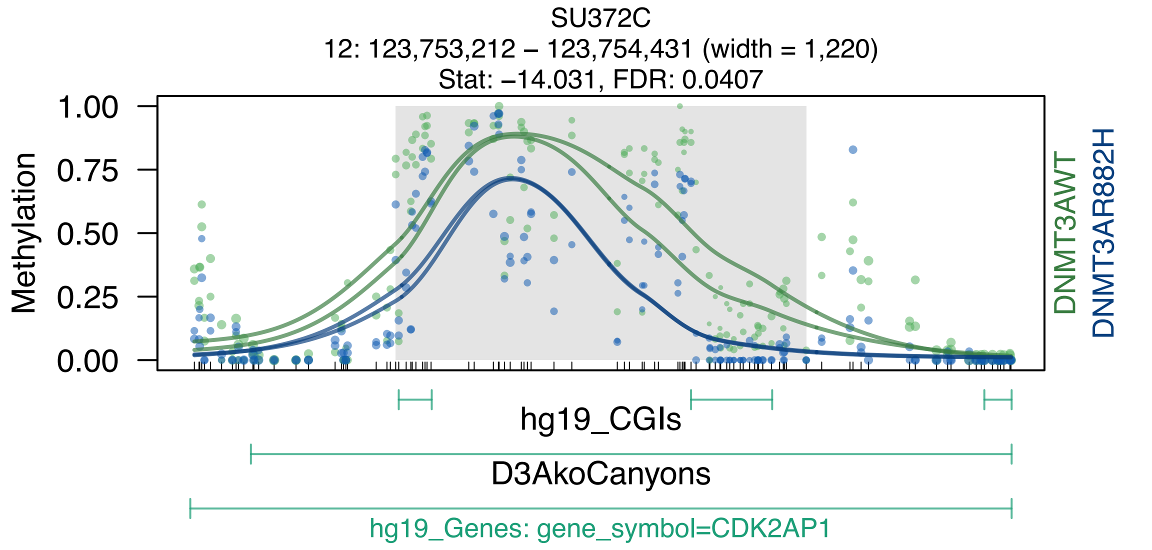
SU372C
12: 123,753,212 – 123,754,431 (width = 1,220)
Stat: –14.031, FDR: 0.0407
Methylation
1.00
0.75
0.50
0.25
0.00
DNMT3AWT
DNMT3AR882H
hg19_CGIs
D3AkoCanyons
hg19_Genes: gene_symbol=CDK2AP1

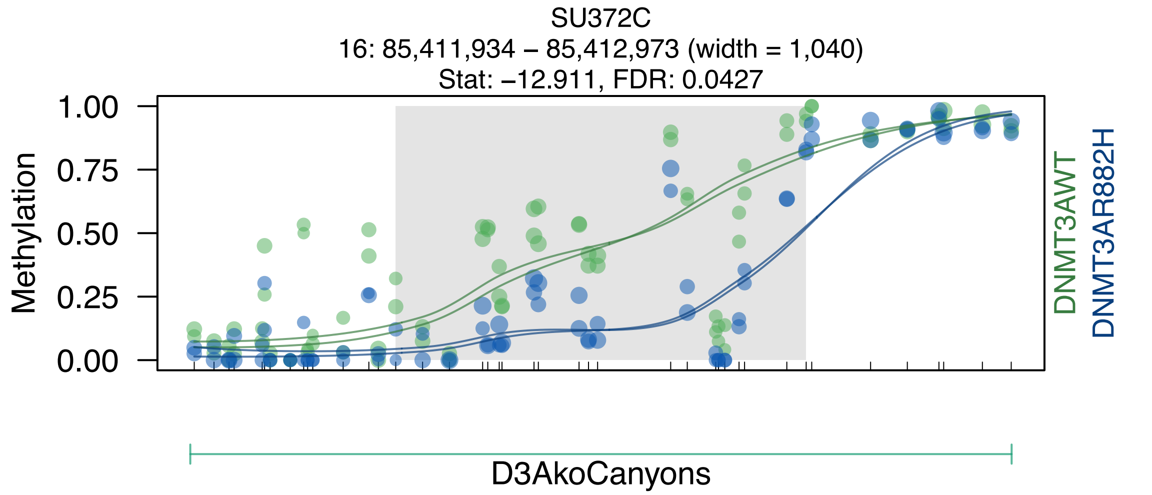
SU372C
16: 85,411,934 – 85,412,973 (width = 1,040)
Stat: –12.911, FDR: 0.0427
Methylation
1.00
0.75
0.50
0.25
0.00
DNMT3AWT
DNMT3AR882H
D3AkoCanyons

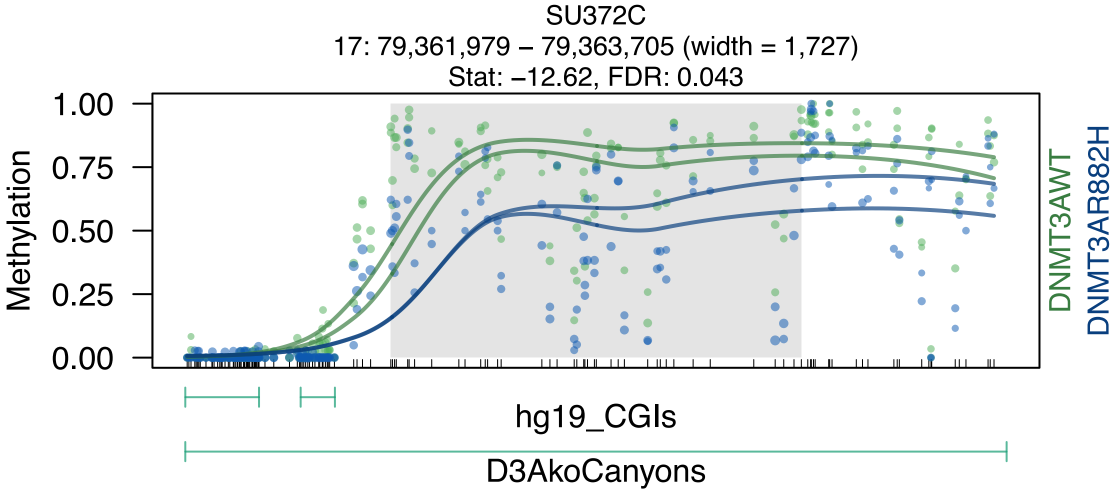

SU372C
17: 79,361,979 – 79,363,705 (width = 1,727)
Stat: –12.62, FDR: 0.043
Methylation
1.00
0.75
0.50
0.25
0.00
DNMT3AWT
DNMT3AR882H
hg19_CGIs
D3AkoCanyons

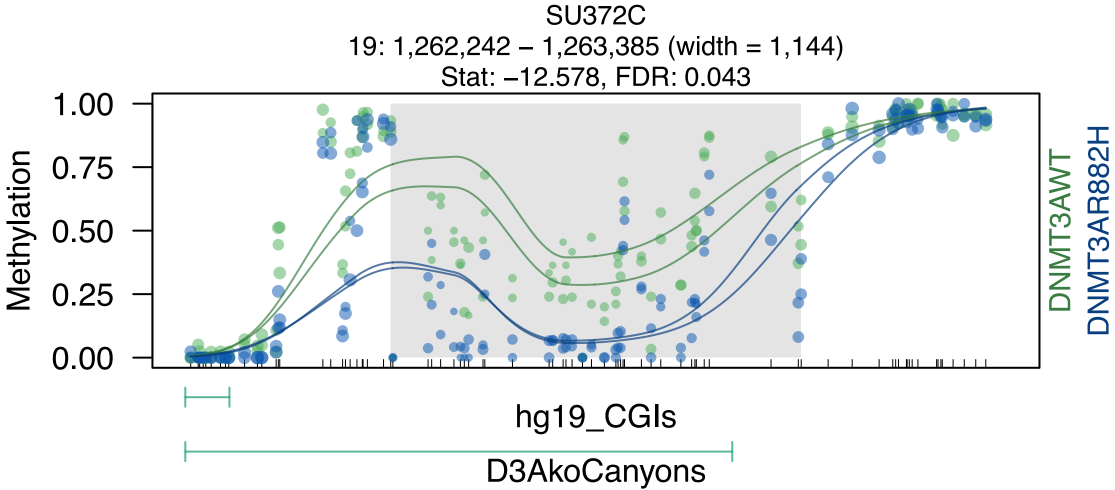
SU372C
19: 1,262,242 – 1,263,385 (width = 1,144)
Stat: –12.578, FDR: 0.043
Methylation
1.00
0.75
0.50
0.25
0.00
DNMT3AWT
DNMT3AR882H
hg19_CGIs
D3AkoCanyons

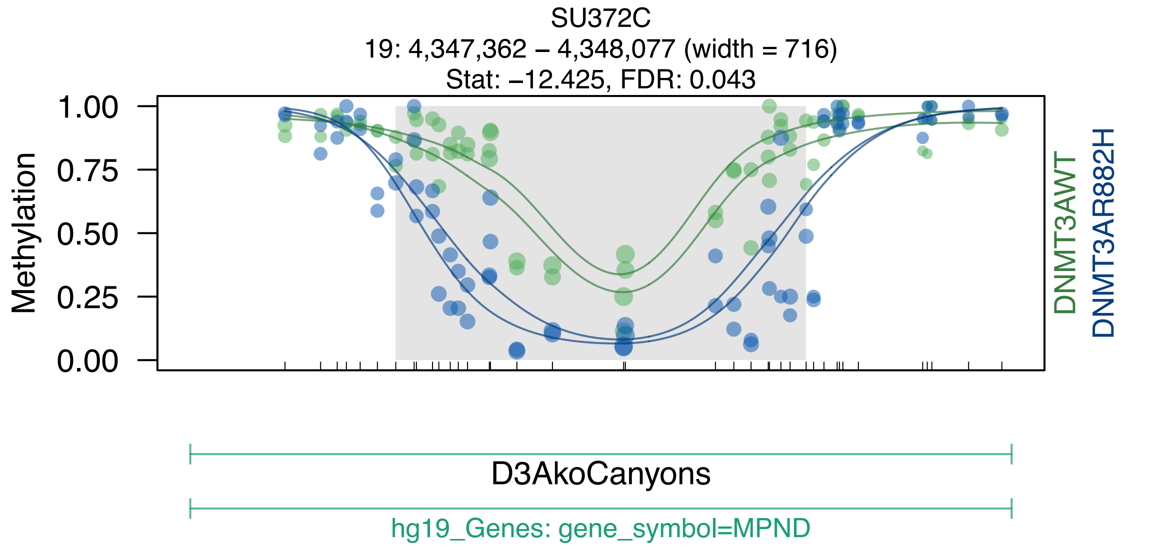
SU372C
19: 4,347,362 – 4,348,077 (width = 716)
Stat: –12.425, FDR: 0.043
Methylation
1.00
0.75
0.50
0.25
0.00
DNMT3AWT
DNMT3AR882H
D3AkoCanyons
hg19_Genes: gene_symbol=MPND

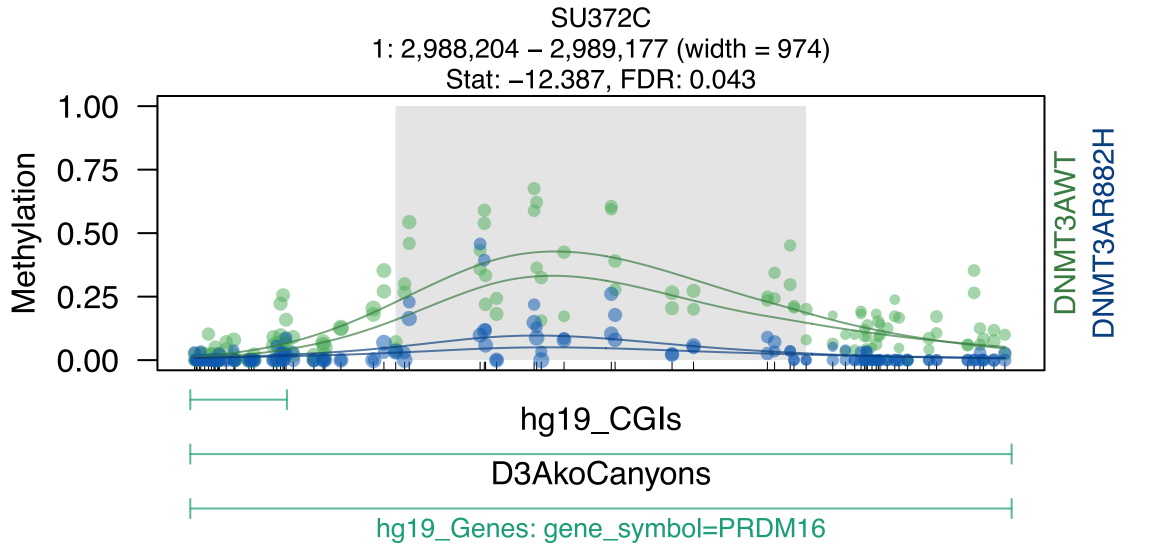
SU372C
1: 2,988,204 – 2,989,177 (width = 974)
Stat: –12.387, FDR: 0.043
Methylation
1.00
0.75
0.50
0.25
0.00
DNMT3AWT
DNMT3AR882H
hg19_CGIs
D3AkoCanyons
hg19_Genes: gene_symbol=PRDM16

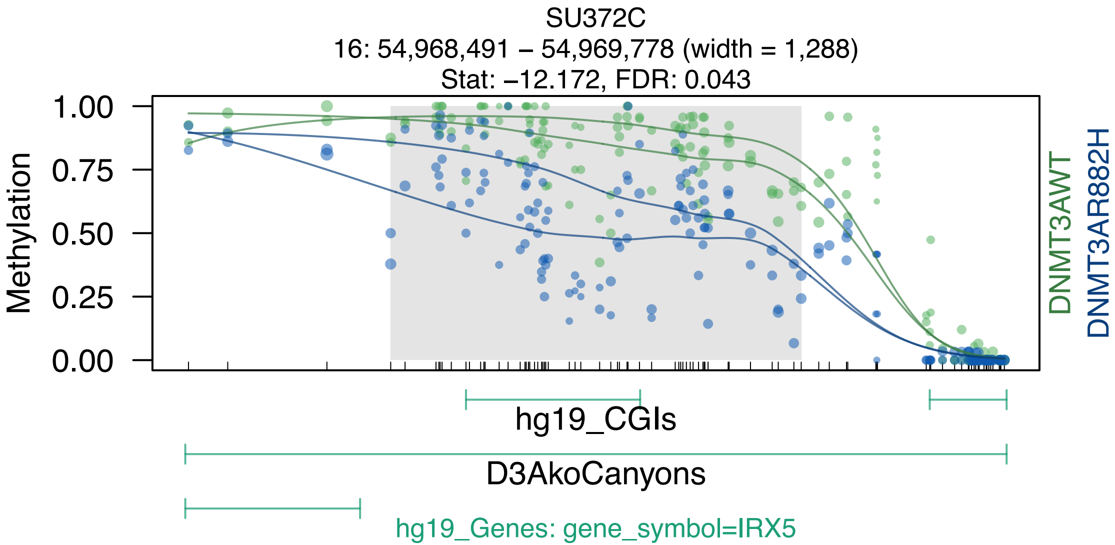
SU372C
16: 54,968,491 – 54,969,778 (width = 1,288)
Stat: −12.172, FDR: 0.043
Methylation
1.00
0.75
0.50
0.25
0.00
DNMT3AWT
DNMT3AR882H
hg19_CGIs
D3AkoCanyons
hg19_Genes: gene_symbol=IRX5

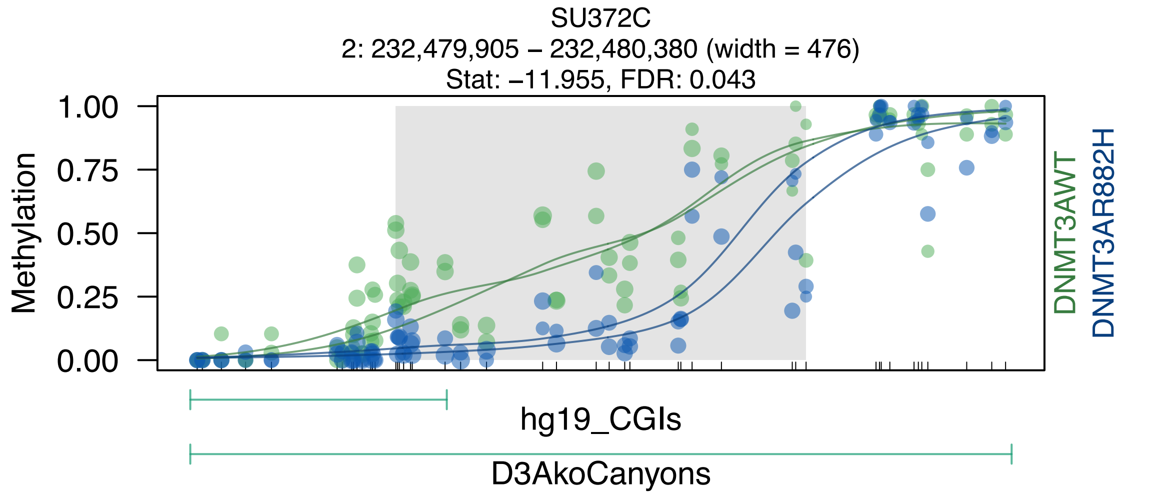
SU372C
2: 232,479,905 – 232,480,380 (width = 476)
Stat: –11.955, FDR: 0.043
Methylation
1.00
0.75
0.50
0.25
0.00
DNMT3AWT
DNMT3AR882H
hg19_CGIs
D3AkoCanyons

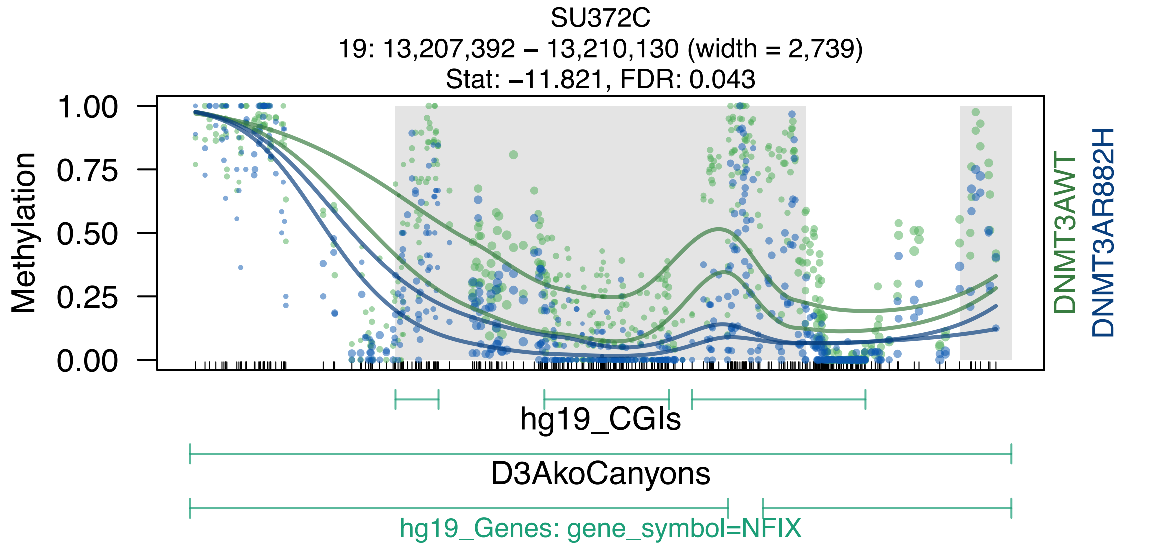
SU372C
19: 13,207,392 – 13,210,130 (width = 2,739)
Stat: –11.821, FDR: 0.043
1.00
0.75
0.50
0.25
0.00
Methylation
DNMT3AWT
DNMT3AR882H
hg19_CGIs
D3AkoCanyons
hg19_Genes: gene_symbol=NFIX

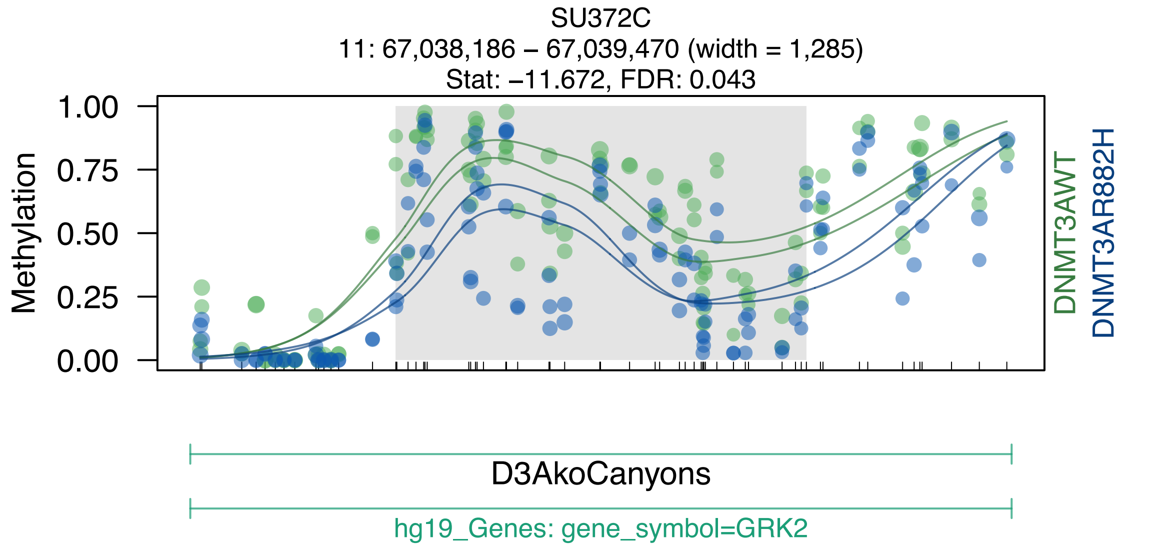
SU372C
11: 67,038,186 – 67,039,470 (width = 1,285)
Stat: –11.672, FDR: 0.043
Methylation
1.00
0.75
0.50
0.25
0.00
DNMT3AWT
DNMT3AR882H
D3AkoCanyons
hg19_Genes: gene_symbol=GRK2

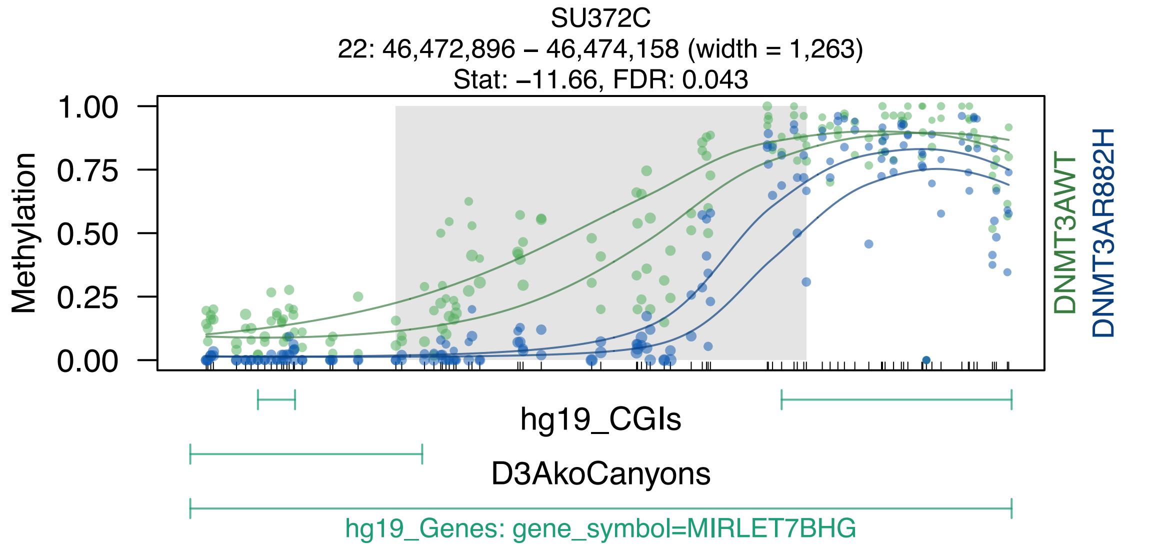
SU372C
22: 46,472,896 – 46,474,158 (width = 1,263)
Stat: −11.66, FDR: 0.043
Methylation
1.00
0.75
0.50
0.25
0.00
DNMT3AWT
DNMT3AR882H
hg19_CGIs
D3AkoCanyons
hg19_Genes: gene_symbol=MIRLET7BHG

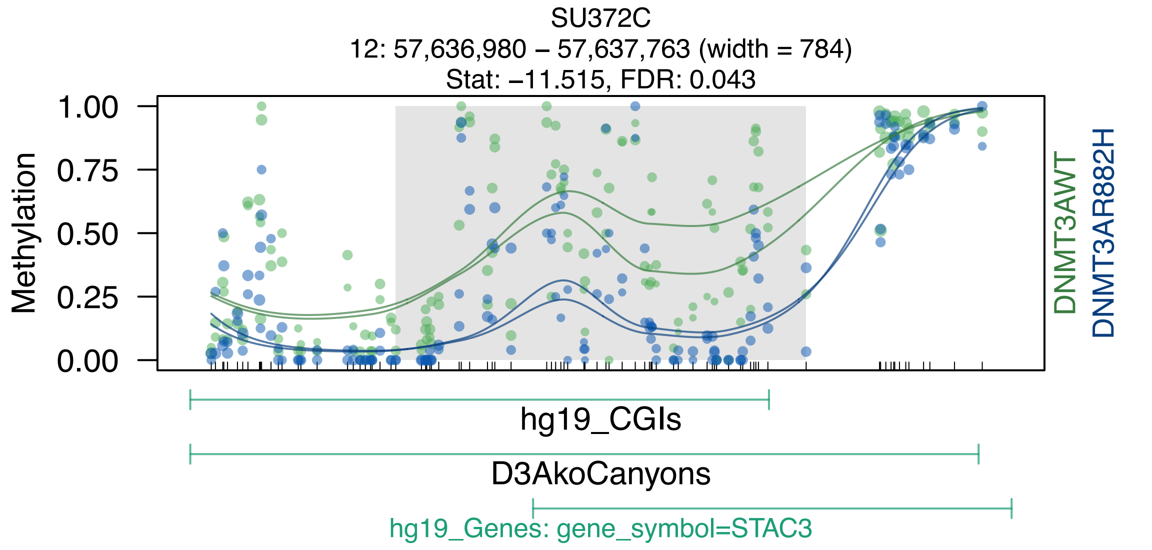
SU372C
12: 57,636,980 – 57,637,763 (width = 784)
Stat: –11.515, FDR: 0.043
Methylation
1.00
0.75
0.50
0.25
0.00
DNMT3AWT
DNMT3AR882H
hg19_CGIs
D3AkoCanyons
hg19_Genes: gene_symbol=STAC3

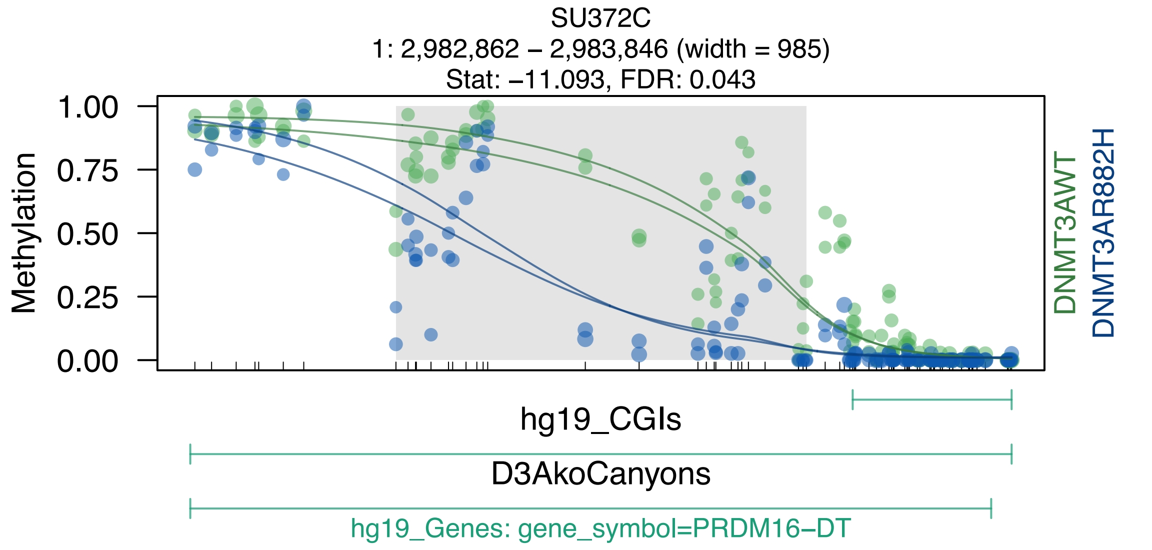

SU372C
1: 2,982,862 – 2,983,846 (width = 985)
Stat: −11.093, FDR: 0.043
Methylation
1.00
0.75
0.50
0.25
0.00
DNMT3AWT
DNMT3AR882H
hg19_CGIs
D3AkoCanyons
hg19_Genes: gene_symbol=PRDM16−DT

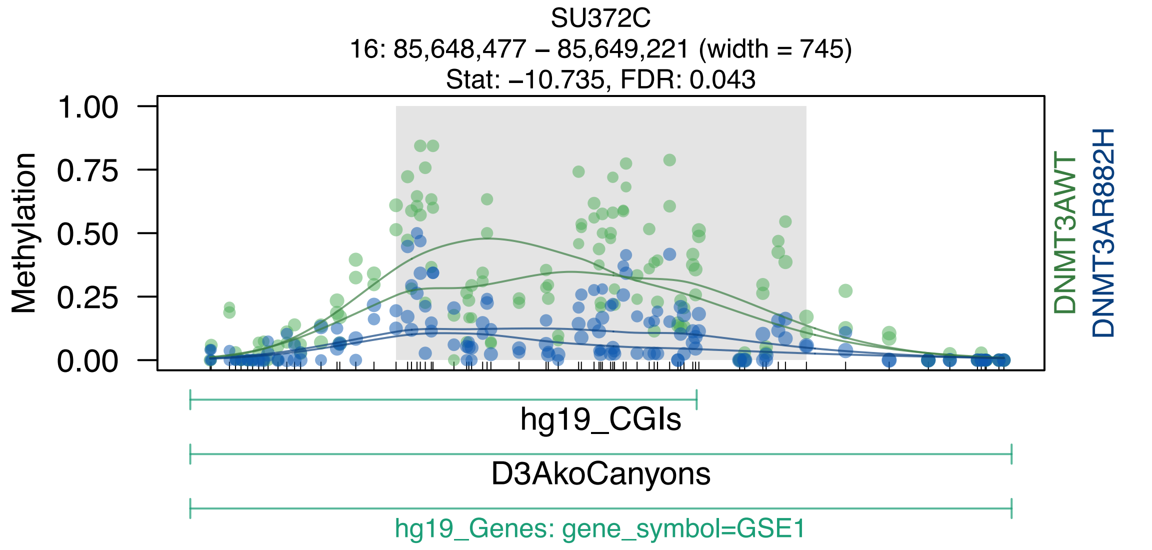

SU372C
16: 85,648,477 – 85,649,221 (width = 745)
Stat: –10.735, FDR: 0.043
Methylation
1.00
0.75
0.50
0.25
0.00
DNMT3AWT
DNMT3AR882H
hg19_CGIs
D3AkoCanyons
hg19_Genes: gene_symbol=GSE1

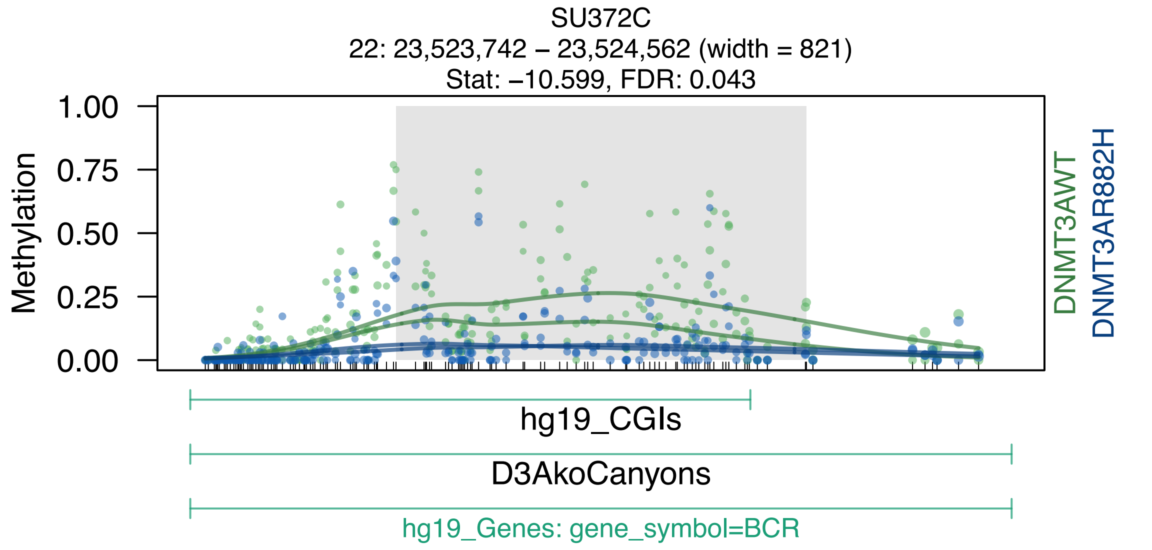

SU372C
22: 23,523,742 – 23,524,562 (width = 821)
Stat: –10.599, FDR: 0.043
Methylation
1.00
0.75
0.50
0.25
0.00
DNMT3AWT
DNMT3AR882H
hg19_CGIs
D3AkoCanyons
hg19_Genes: gene_symbol=BCR

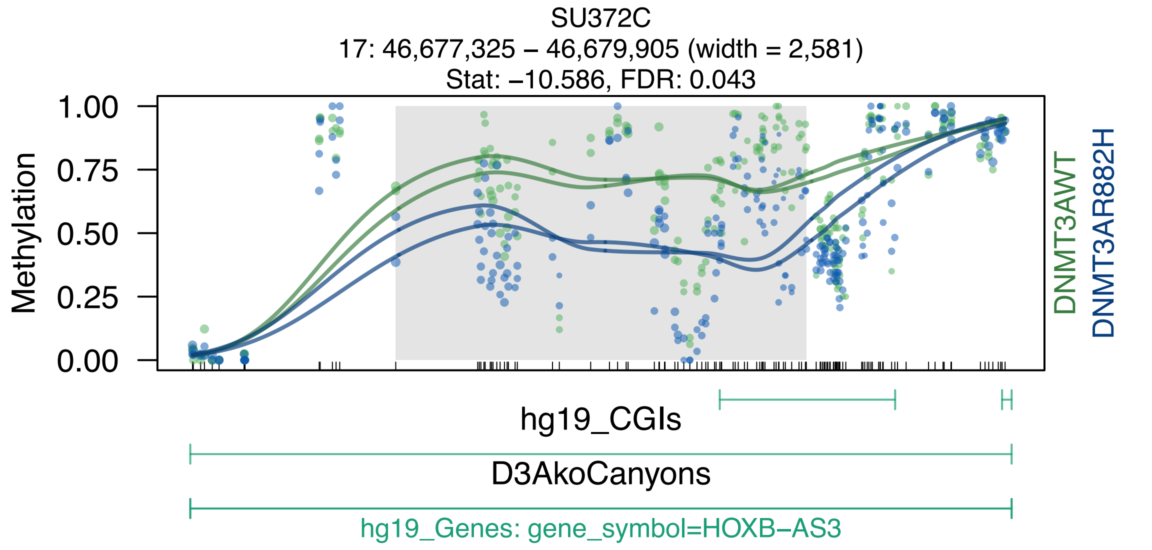
SU372C
17: 46,677,325 – 46,679,905 (width = 2,581)
Stat: −10.586, FDR: 0.043
Methylation
1.00
0.75
0.50
0.25
0.00
DNMT3AWT
DNMT3AR882H
hg19_CGIs
D3AkoCanyons
hg19_Genes: gene_symbol=HOXB−AS3

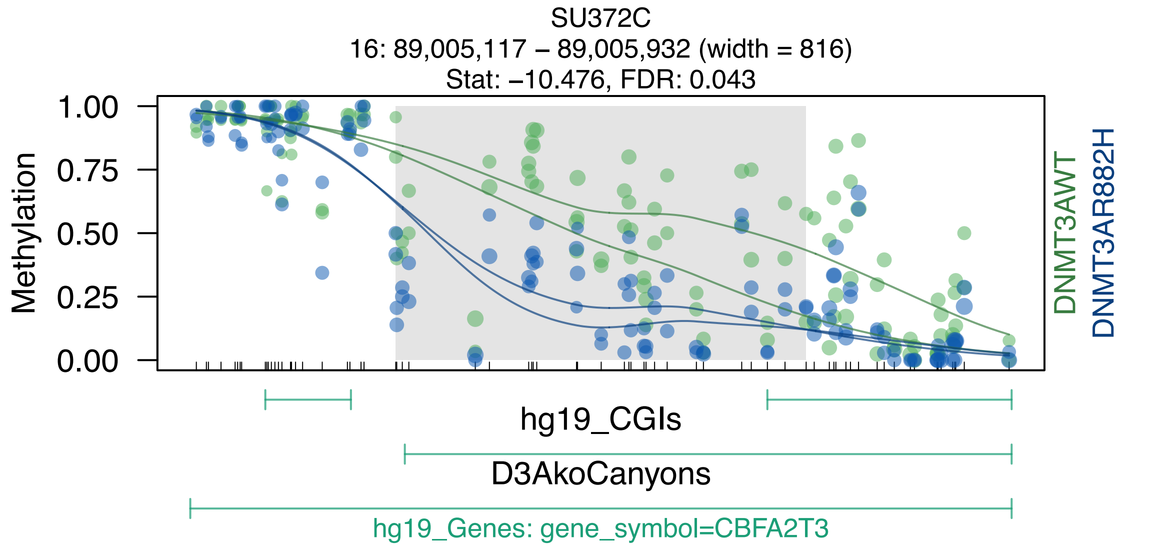

SU372C
16: 89,005,117 – 89,005,932 (width = 816)
Stat: –10.476, FDR: 0.043
Methylation
1.00
0.75
0.50
0.25
0.00
DNMT3AWT
DNMT3AR882H
hg19_CGIs
D3AkoCanyons
hg19_Genes: gene_symbol=CBFA2T3

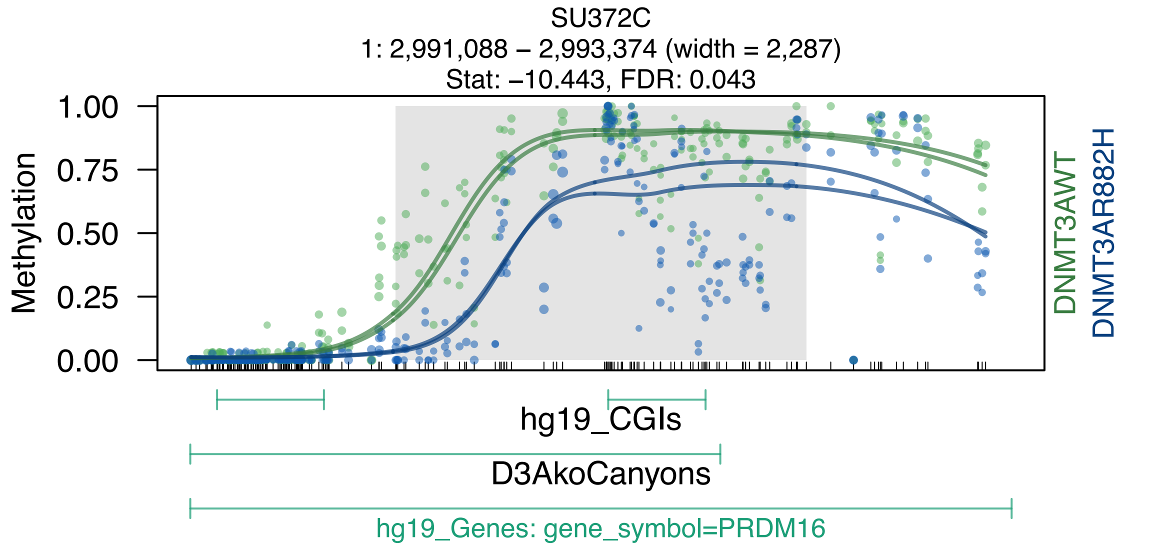
SU372C
1: 2,991,088 – 2,993,374 (width = 2,287)
Stat: –10.443, FDR: 0.043
Methylation
1.00
0.75
0.50
0.25
0.00
DNMT3AWT
DNMT3AR882H
hg19_CGIs
D3AkoCanyons
hg19_Genes: gene_symbol=PRDM16

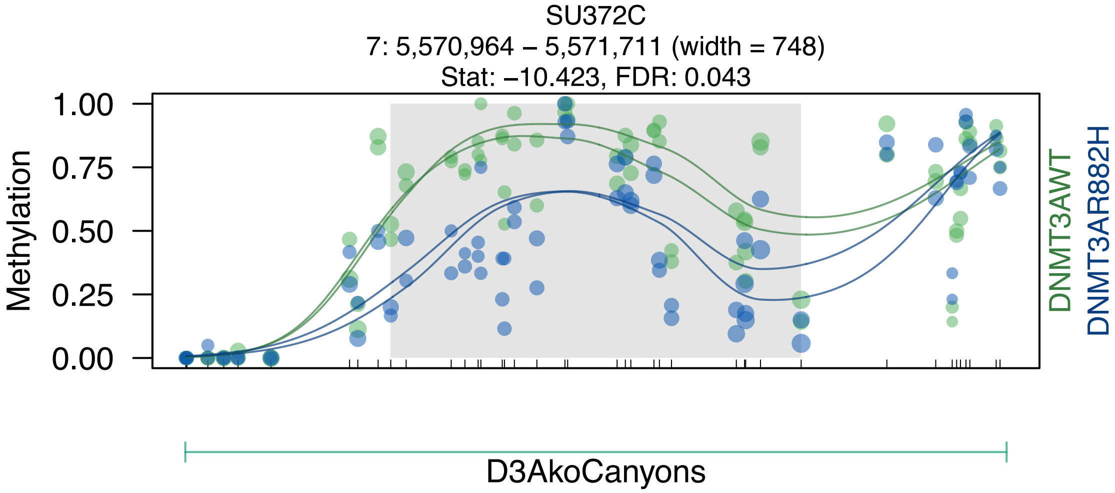
SU372C
7: 5,570,964 – 5,571,711 (width = 748)
Stat: −10.423, FDR: 0.043
Methylation
1.00
0.75
0.50
0.25
0.00
DNMT3AWT
DNMT3AR882H
D3AkoCanyons

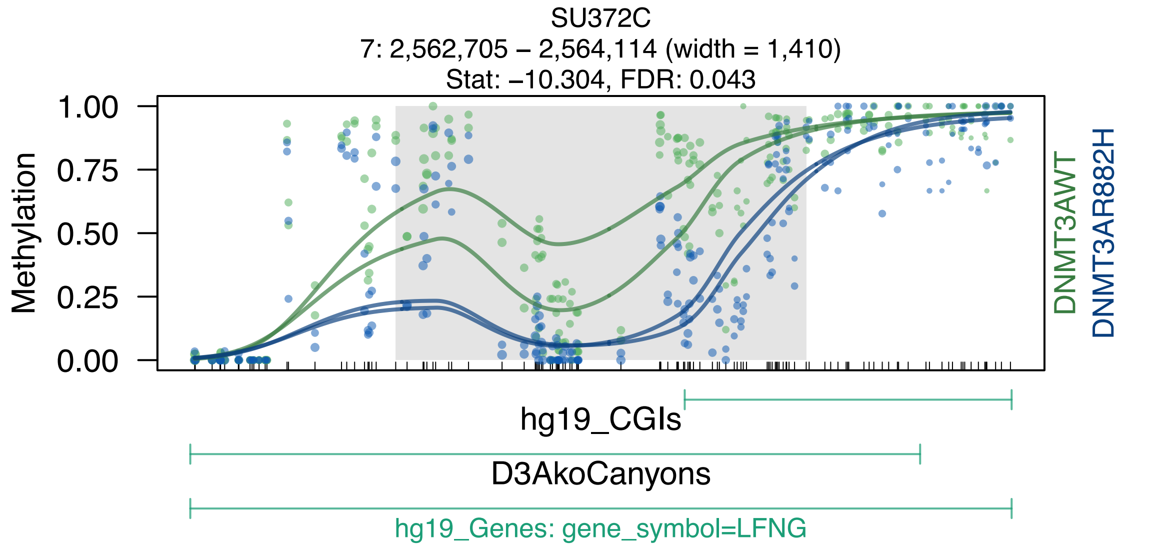
SU372C
7: 2,562,705 – 2,564,114 (width = 1,410)
Stat: –10.304, FDR: 0.043
Methylation
1.00
0.75
0.50
0.25
0.00
DNMT3AWT
DNMT3AR882H
hg19_CGIs
D3AkoCanyons
hg19_Genes: gene_symbol=LFNG

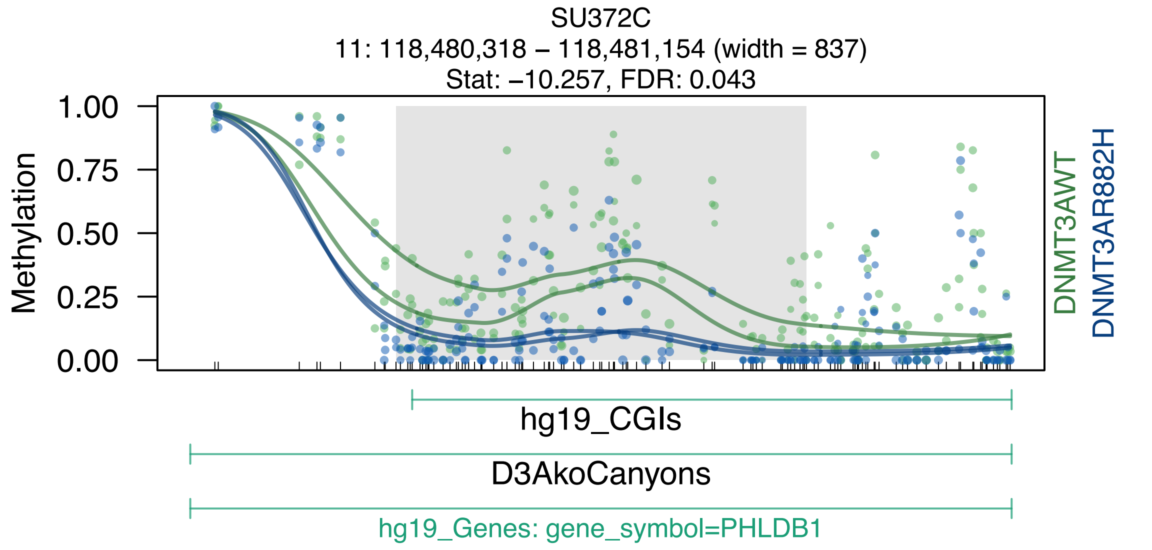
SU372C
11: 118,480,318 – 118,481,154 (width = 837)
Stat: –10.257, FDR: 0.043
Methylation
1.00
0.75
0.50
0.25
0.00
DNMT3AWT
DNMT3AR882H
hg19_CGIs
D3AkoCanyons
hg19_Genes: gene_symbol=PHLDB1

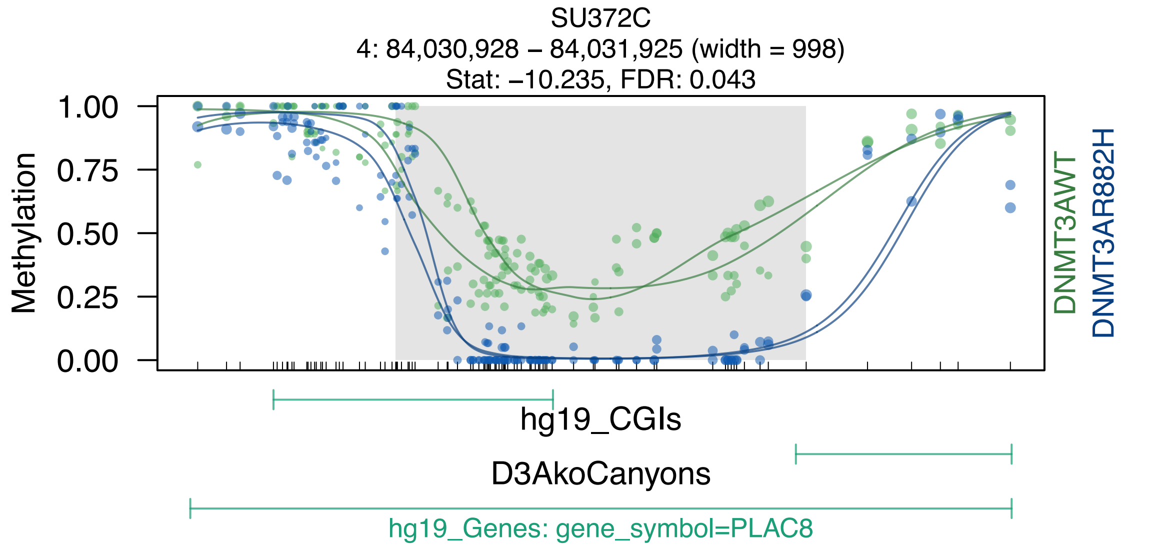

SU372C
4: 84,030,928 – 84,031,925 (width = 998)
Stat: –10.235, FDR: 0.043
Methylation
1.00
0.75
0.50
0.25
0.00
DNMT3AWT
DNMT3AR882H
hg19_CGIs
D3AkoCanyons
hg19_Genes: gene_symbol=PLAC8

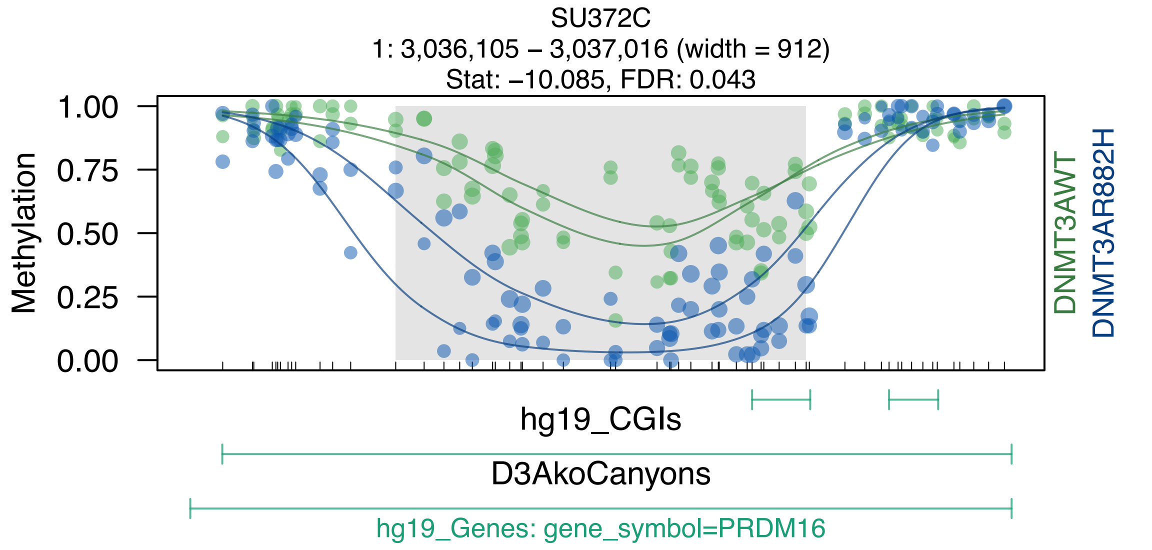

SU372C
1: 3,036,105 – 3,037,016 (width = 912)
Stat: –10.085, FDR: 0.043
1.00
0.75
0.50
0.25
0.00
Methylation
DNMT3AWT
DNMT3AR882H
hg19_CGIs
D3AkoCanyons
hg19_Genes: gene_symbol=PRDM16

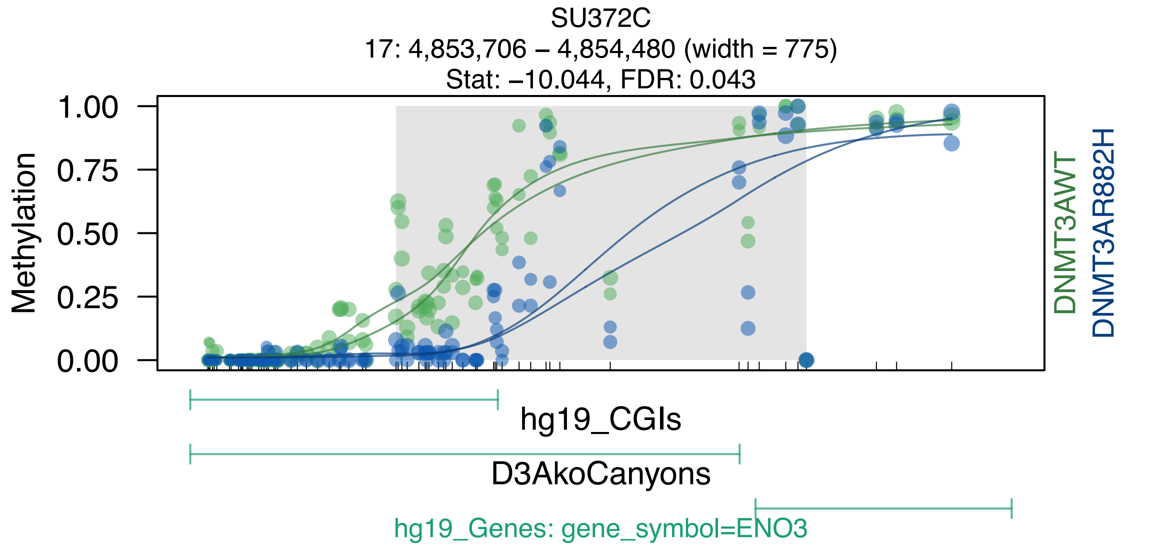
SU372C
17: 4,853,706 – 4,854,480 (width = 775)
Stat: –10.044, FDR: 0.043
Methylation
1.00
0.75
0.50
0.25
0.00
DNMT3AWT
DNMT3AR882H
hg19_CGIs
D3AkoCanyons
hg19_Genes: gene_symbol=ENO3

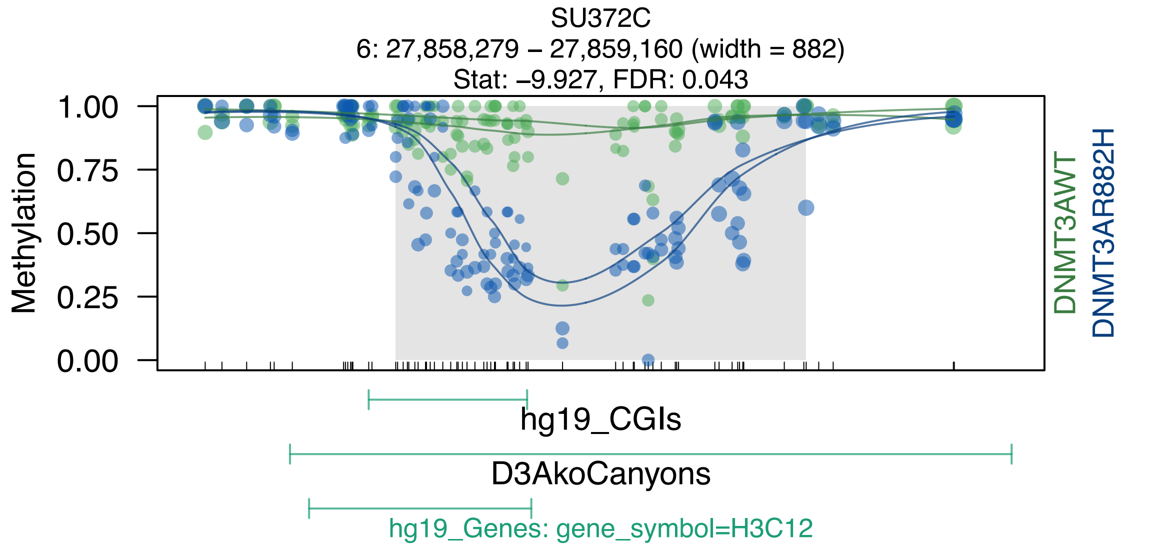
SU372C
6: 27,858,279 – 27,859,160 (width = 882)
Stat: –9.927, FDR: 0.043
Methylation
1.00
0.75
0.50
0.25
0.00
DNMT3AWT
DNMT3AR882H
hg19_CGIs
D3AkoCanyons
hg19_Genes: gene_symbol=H3C12

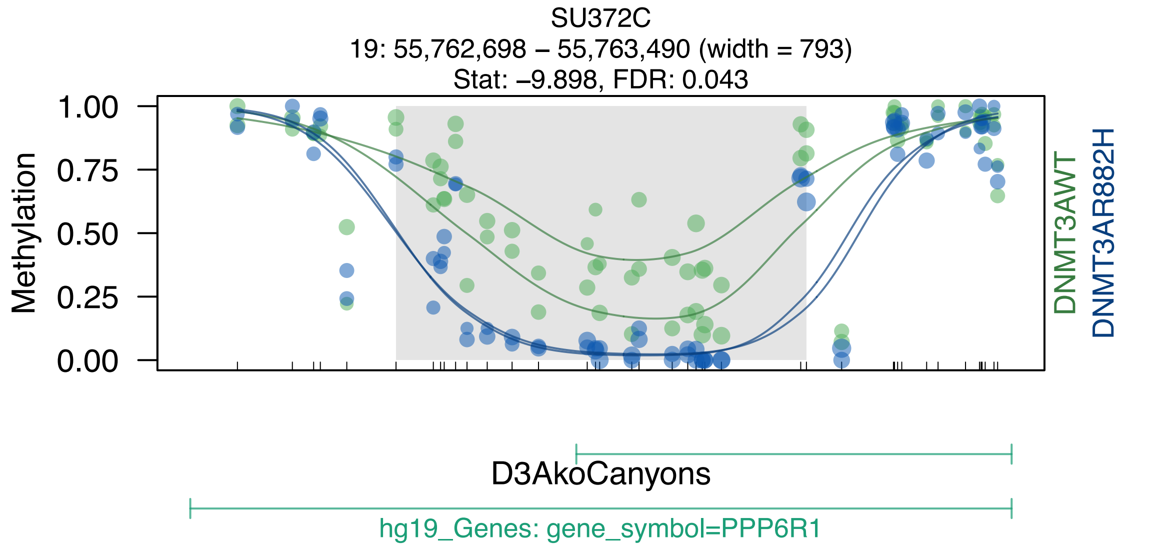
SU372C
19: 55,762,698 – 55,763,490 (width = 793)
Stat: –9.898, FDR: 0.043
Methylation
1.00
0.75
0.50
0.25
0.00
DNMT3AWT
DNMT3AR882H
D3AkoCanyons
hg19_Genes: gene_symbol=PPP6R1

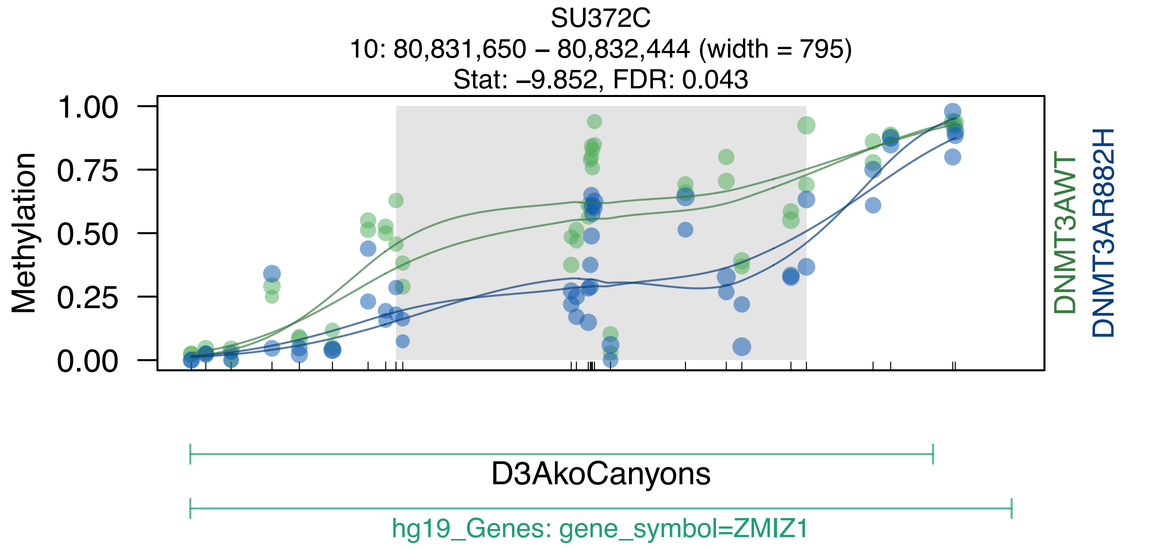
SU372C
10: 80,831,650 – 80,832,444 (width = 795)
Stat: –9.852, FDR: 0.043
Methylation
1.00
0.75
0.50
0.25
0.00
DNMT3AWT
DNMT3AR882H
D3AkoCanyons
hg19_Genes: gene_symbol=ZMIZ1

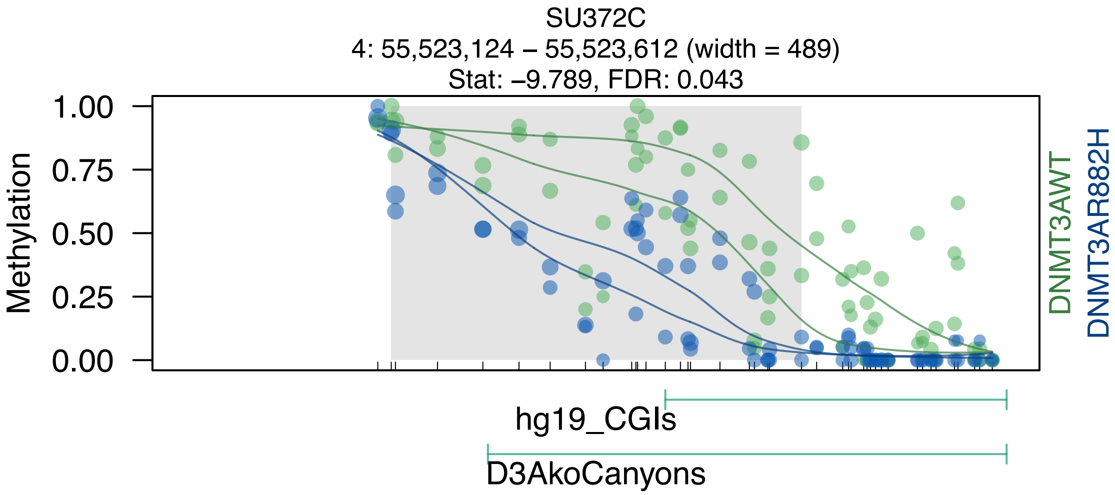

SU372C
4: 55,523,124 – 55,523,612 (width = 489)
Stat: –9.789, FDR: 0.043
1.00
0.75
0.50
0.25
0.00
Methylation
DNMT3AWT
DNMT3AR882H
hg19_CGIs
D3AkoCanyons

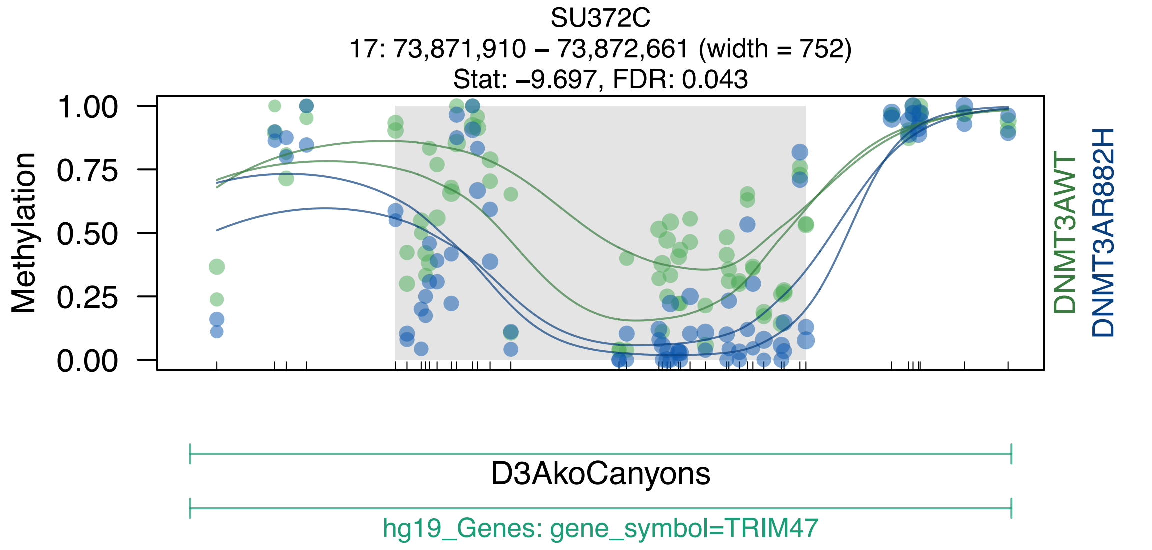
SU372C
17: 73,871,910 – 73,872,661 (width = 752)
Stat: –9.697, FDR: 0.043
Methylation
1.00
0.75
0.50
0.25
0.00
DNMT3AWT
DNMT3AR882H
D3AkoCanyons
hg19_Genes: gene_symbol=TRIM47

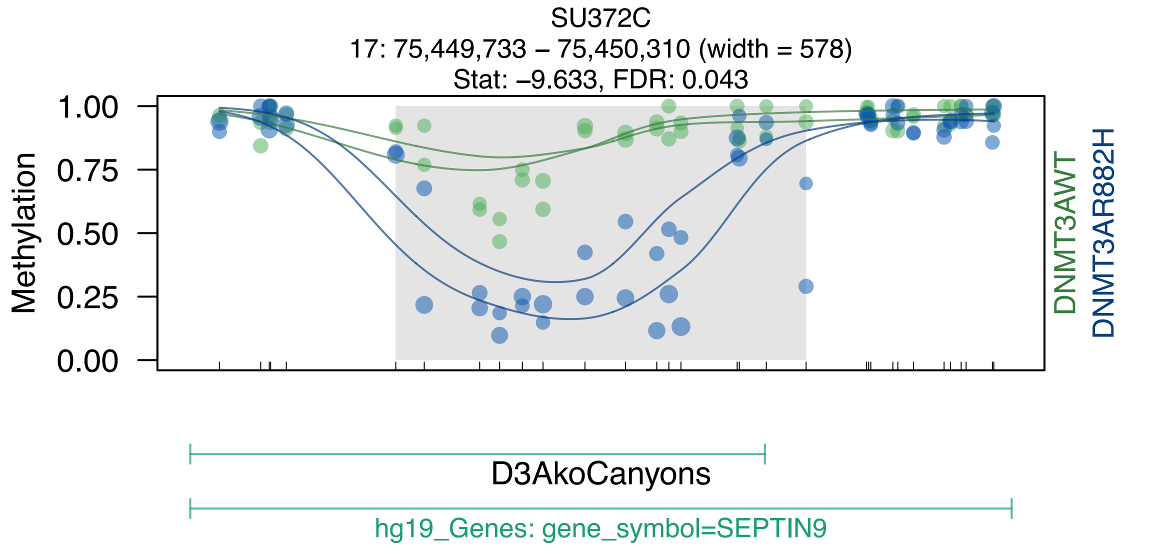
SU372C
17: 75,449,733 – 75,450,310 (width = 578)
Stat: –9.633, FDR: 0.043
Methylation
1.00
0.75
0.50
0.25
0.00
DNMT3AWT
DNMT3AR882H
D3AkoCanyons
hg19_Genes: gene_symbol=SEPTIN9

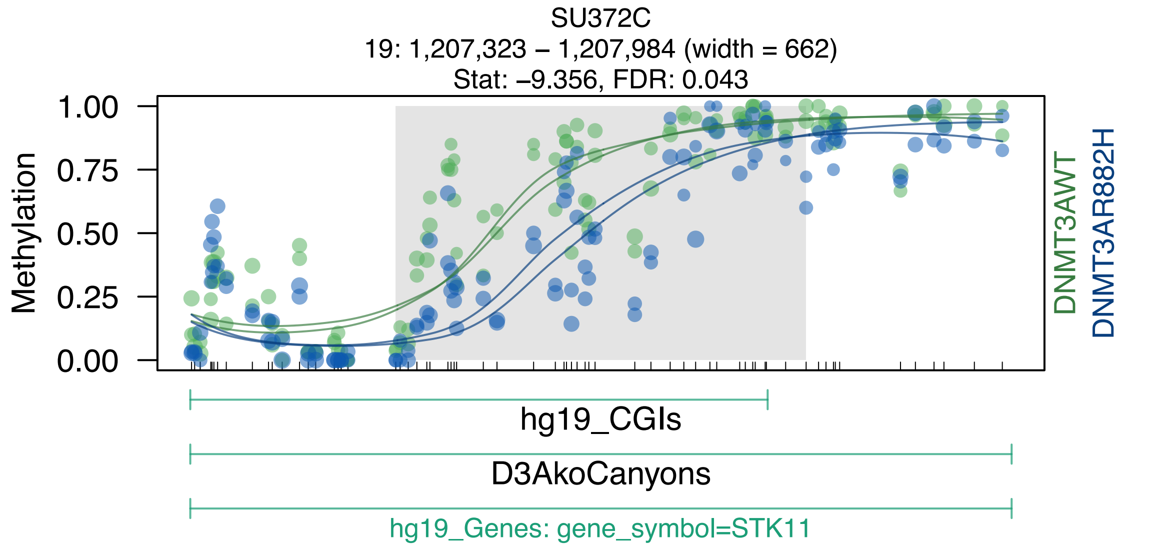

SU372C
19: 1,207,323 – 1,207,984 (width = 662)
Stat: –9.356, FDR: 0.043
Methylation
1.00
0.75
0.50
0.25
0.00
DNMT3AWT
DNMT3AR882H
hg19_CGIs
D3AkoCanyons
hg19_Genes: gene_symbol=STK11

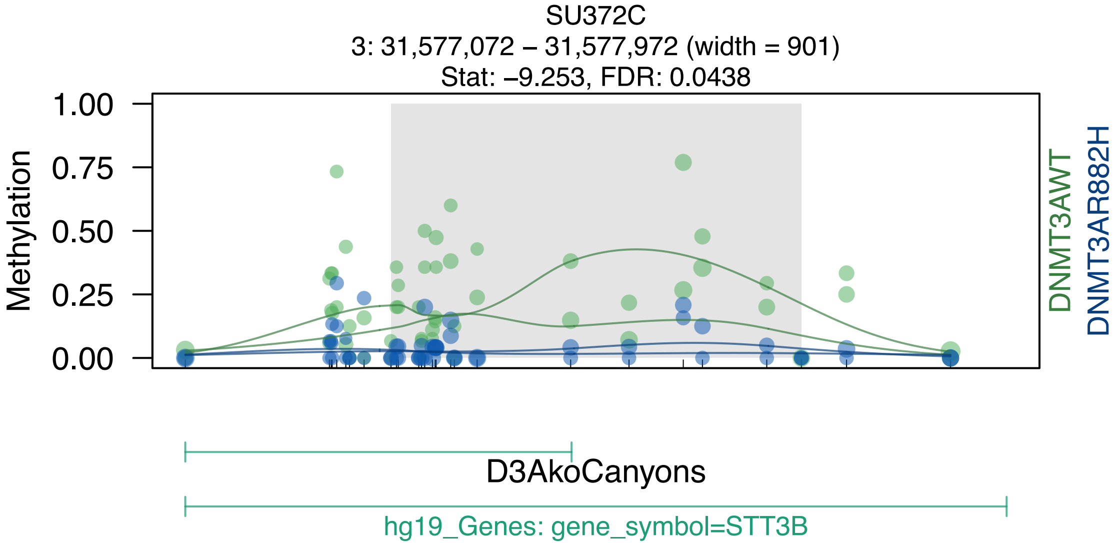
SU372C
3: 31,577,072 – 31,577,972 (width = 901)
Stat: –9.253, FDR: 0.0438
Methylation
1.00
0.75
0.50
0.25
0.00
DNMT3AWT
DNMT3AR882H
D3AkoCanyons
hg19_Genes: gene_symbol=STT3B

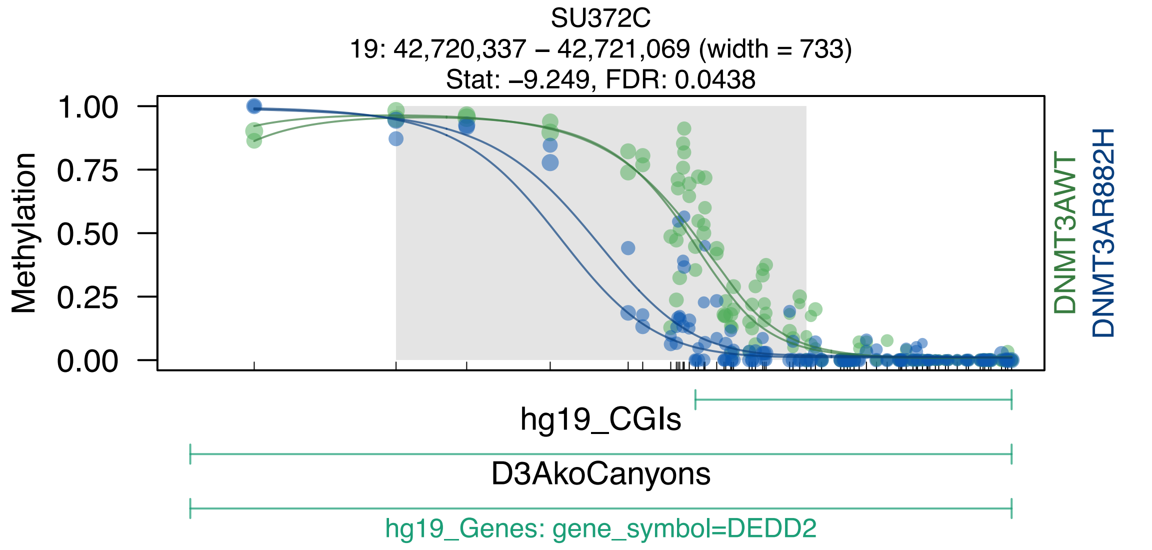
SU372C
19: 42,720,337 – 42,721,069 (width = 733)
Stat: –9.249, FDR: 0.0438
Methylation
1.00
0.75
0.50
0.25
0.00
DNMT3AWT
DNMT3AR882H
hg19_CGIs
D3AkoCanyons
hg19_Genes: gene_symbol=DEDD2

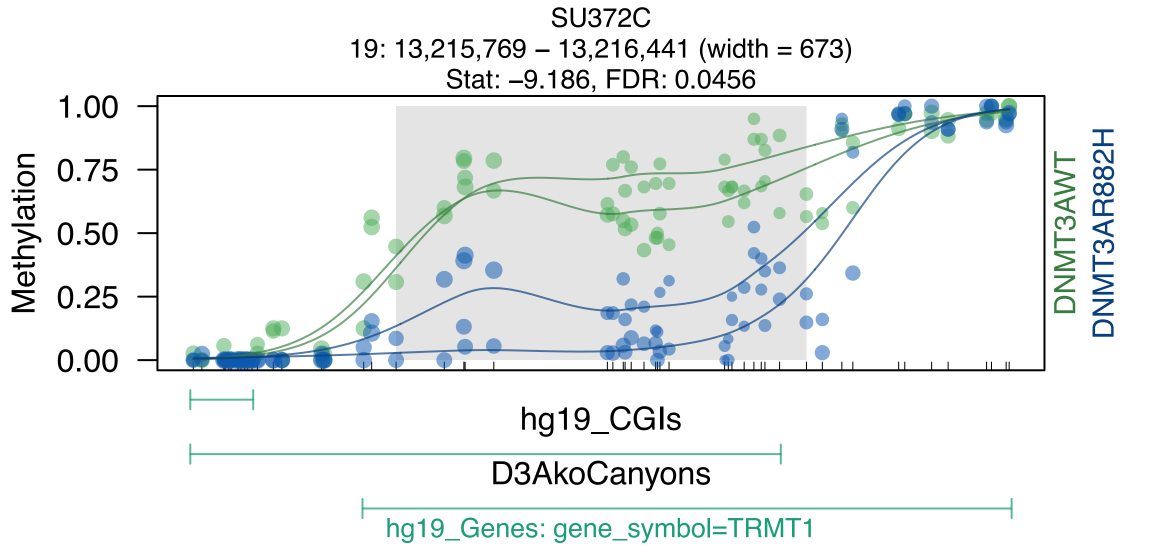
SU372C
19: 13,215,769 – 13,216,441 (width = 673)
Stat: –9.186, FDR: 0.0456
Methylation
1.00
0.75
0.50
0.25
0.00
DNMT3AWT
DNMT3AR882H
hg19_CGIs
D3AkoCanyons
hg19_Genes: gene_symbol=TRMT1

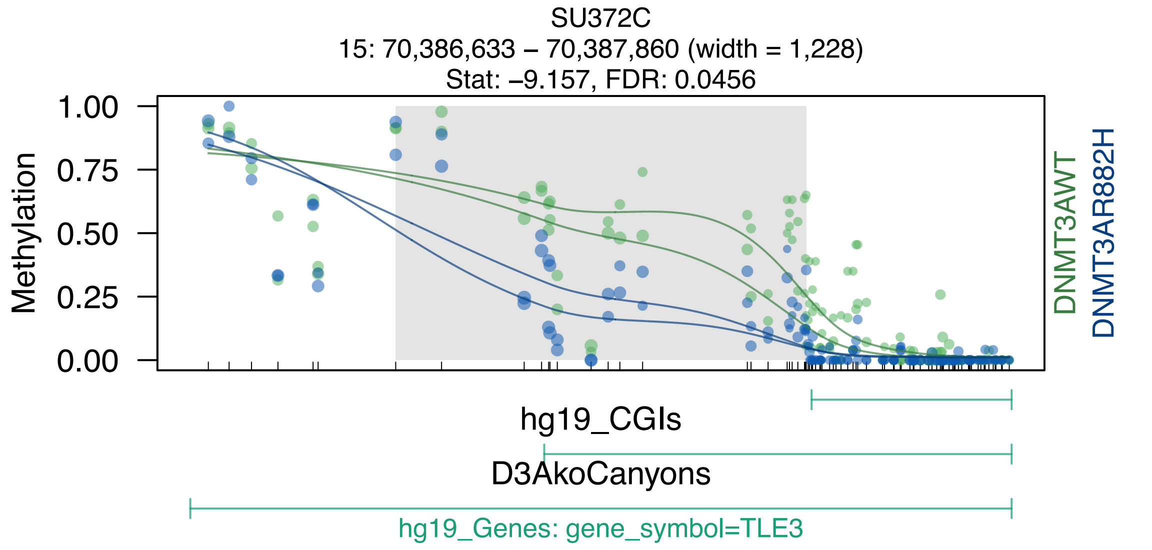
SU372C
15: 70,386,633 – 70,387,860 (width = 1,228)
Stat: –9.157, FDR: 0.0456
Methylation
1.00
0.75
0.50
0.25
0.00
DNMT3AWT
DNMT3AR882H
hg19_CGIs
D3AkoCanyons
hg19_Genes: gene_symbol=TLE3

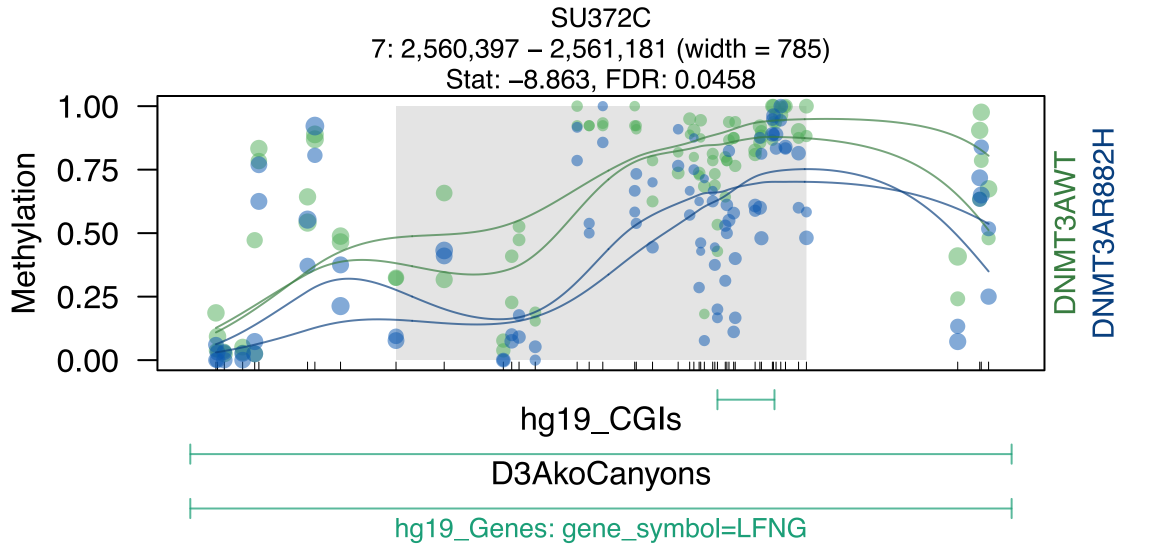
SU372C
7: 2,560,397 – 2,561,181 (width = 785)
Stat: –8.863, FDR: 0.0458
Methylation
1.00
0.75
0.50
0.25
0.00
DNMT3AWT
DNMT3AR882H
hg19_CGIs
D3AkoCanyons
hg19_Genes: gene_symbol=LFNG

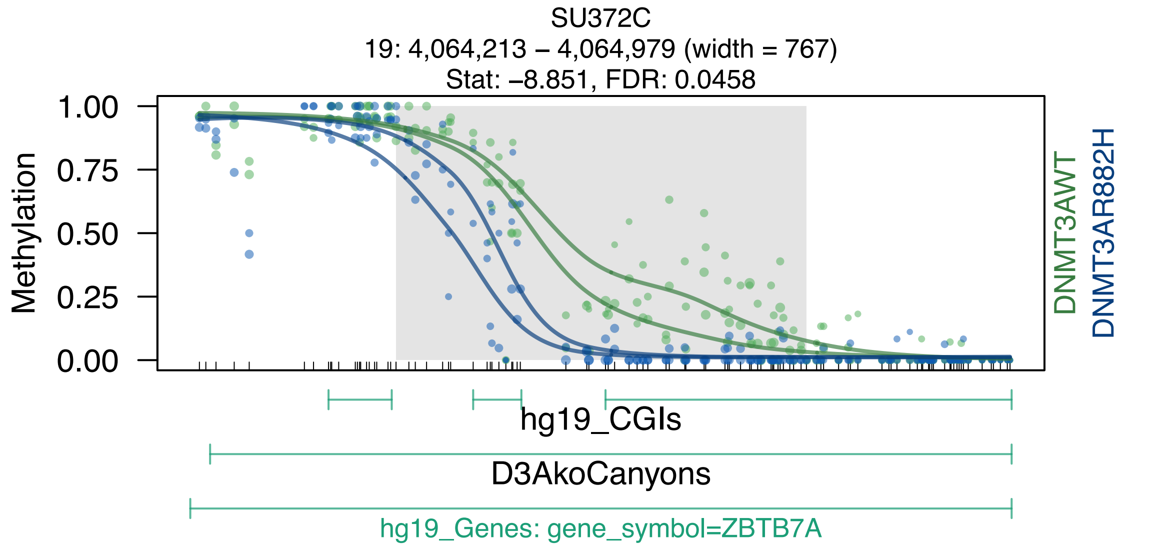
SU372C
19: 4,064,213 – 4,064,979 (width = 767)
Stat: −8.851, FDR: 0.0458
Methylation
1.00
0.75
0.50
0.25
0.00
DNMT3AWT
DNMT3AR882H
hg19_CGIs
D3AkoCanyons
hg19_Genes: gene_symbol=ZBTB7A

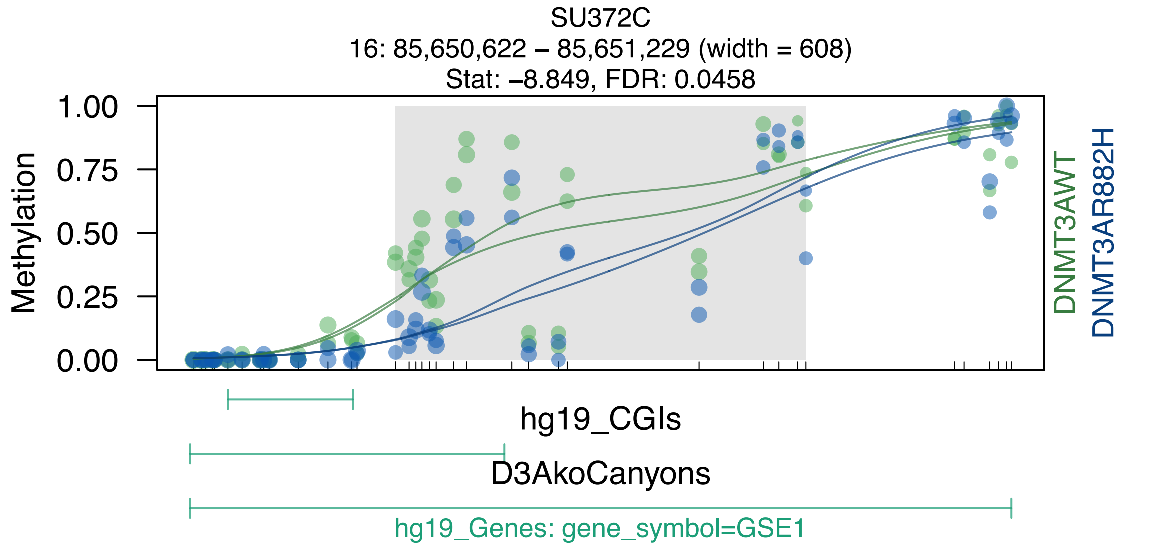

SU372C
16: 85,650,622 – 85,651,229 (width = 608)
Stat: –8.849, FDR: 0.0458
Methylation
1.00
0.75
0.50
0.25
0.00
DNMT3AWT
DNMT3AR882H
hg19_CGIs
D3AkoCanyons
hg19_Genes: gene_symbol=GSE1

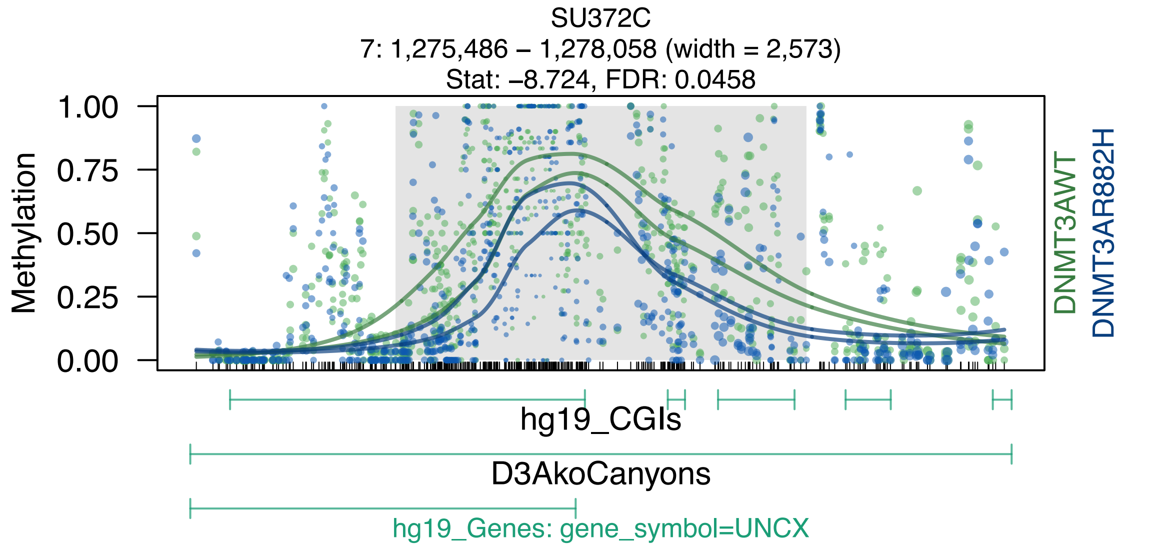
SU372C
7: 1,275,486 – 1,278,058 (width = 2,573)
Stat: –8.724, FDR: 0.0458
Methylation
1.00
0.75
0.50
0.25
0.00
DNMT3AWT
DNMT3AR882H
hg19_CGIs
D3AkoCanyons
hg19_Genes: gene_symbol=UNCX

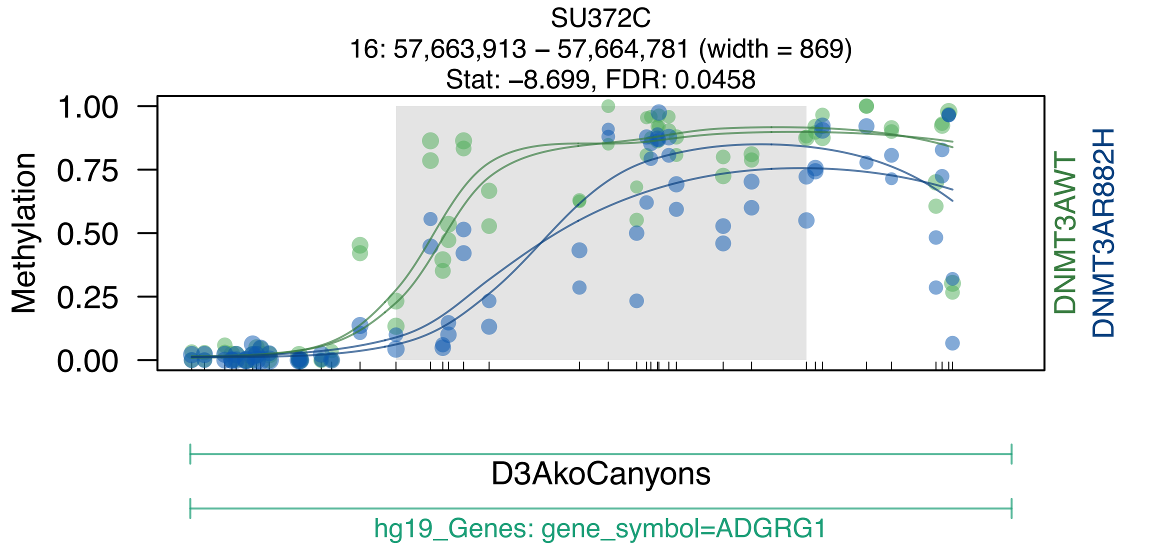
SU372C
16: 57,663,913 – 57,664,781 (width = 869)
Stat: –8.699, FDR: 0.0458
Methylation
1.00
0.75
0.50
0.25
0.00
DNMT3AWT
DNMT3AR882H
D3AkoCanyons
hg19_Genes: gene_symbol=ADGRG1

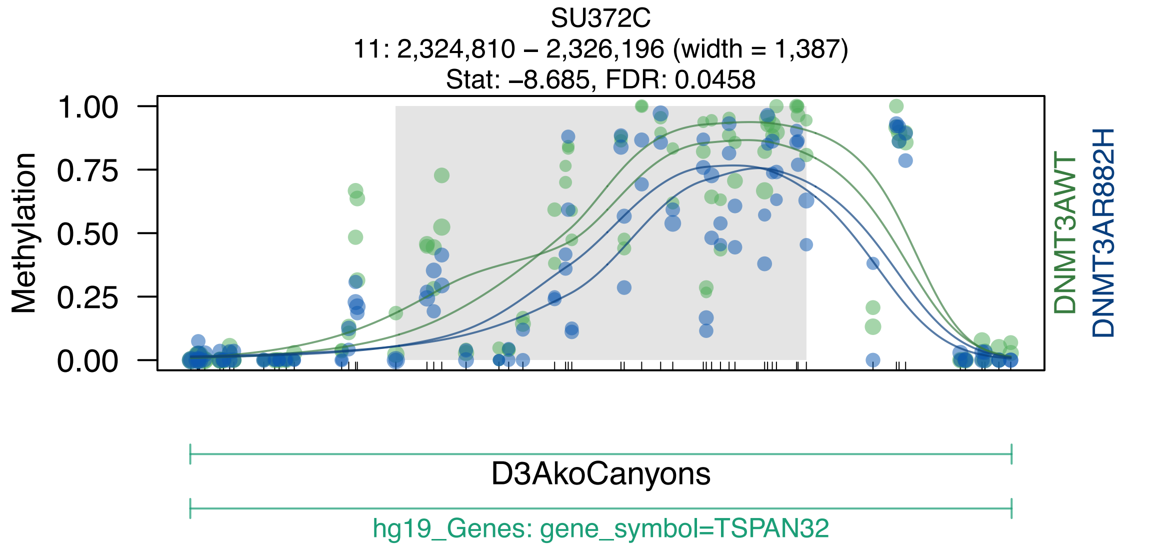

SU372C
11: 2,324,810 – 2,326,196 (width = 1,387)
Stat: –8.685, FDR: 0.0458
Methylation
1.00
0.75
0.50
0.25
0.00
DNMT3AWT
DNMT3AR882H
D3AkoCanyons
hg19_Genes: gene_symbol=TSPAN32

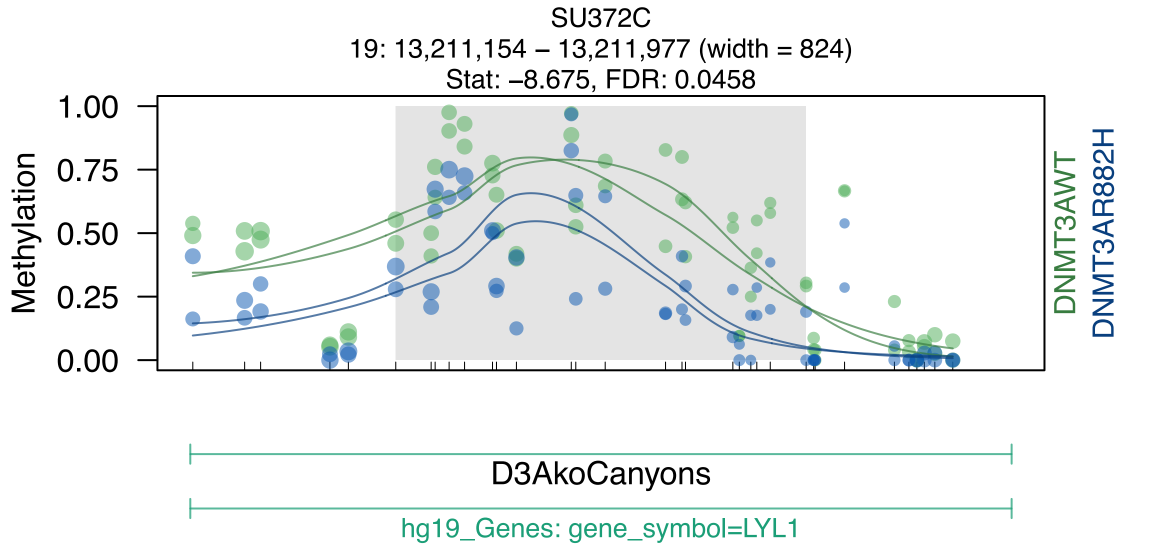
SU372C
19: 13,211,154 – 13,211,977 (width = 824)
Stat: –8.675, FDR: 0.0458
Methylation
1.00
0.75
0.50
0.25
0.00
DNMT3AWT
DNMT3AR882H
D3AkoCanyons
hg19_Genes: gene_symbol=LYL1

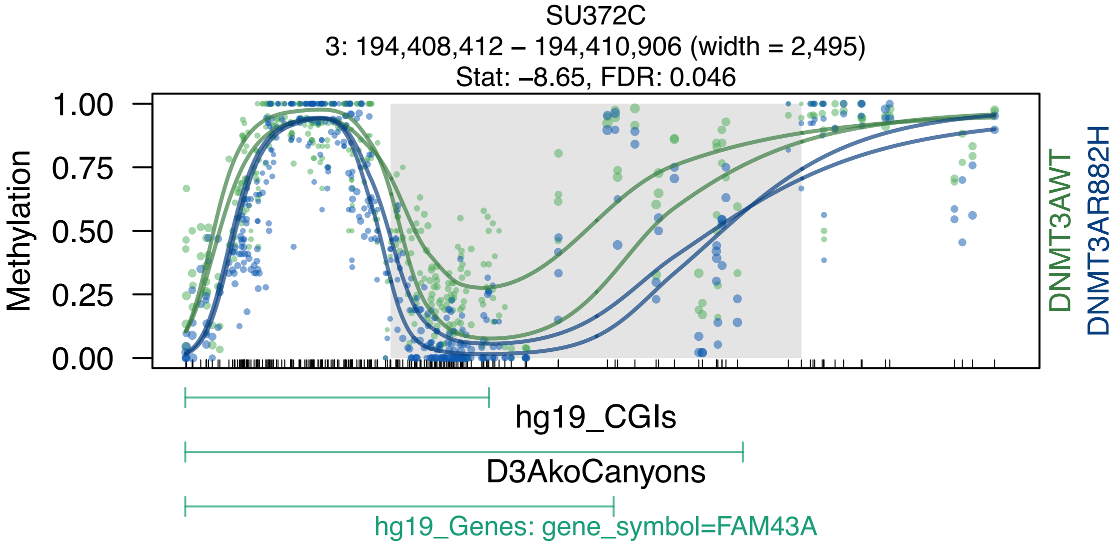
SU372C
3: 194,408,412 – 194,410,906 (width = 2,495)
Stat: –8.65, FDR: 0.046
Methylation
1.00
0.75
0.50
0.25
0.00
DNMT3AWT
DNMT3AR882H
hg19_CGIs
D3AkoCanyons
hg19_Genes: gene_symbol=FAM43A

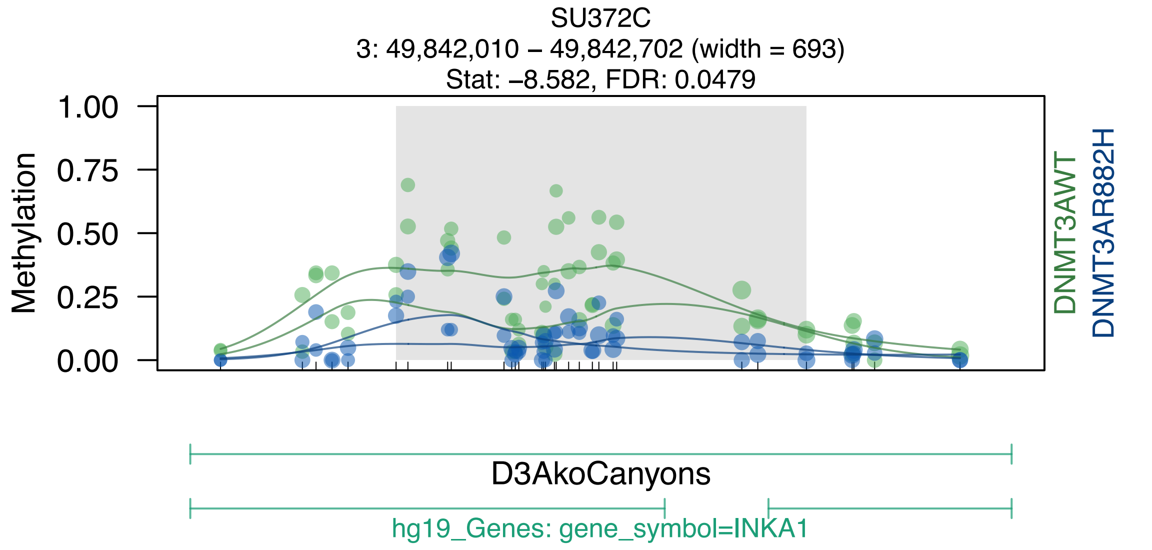

SU372C
3: 49,842,010 – 49,842,702 (width = 693)
Stat: –8.582, FDR: 0.0479
Methylation
1.00
0.75
0.50
0.25
0.00
DNMT3AWT
DNMT3AR882H
D3AkoCanyons
hg19_Genes: gene_symbol=INKA1

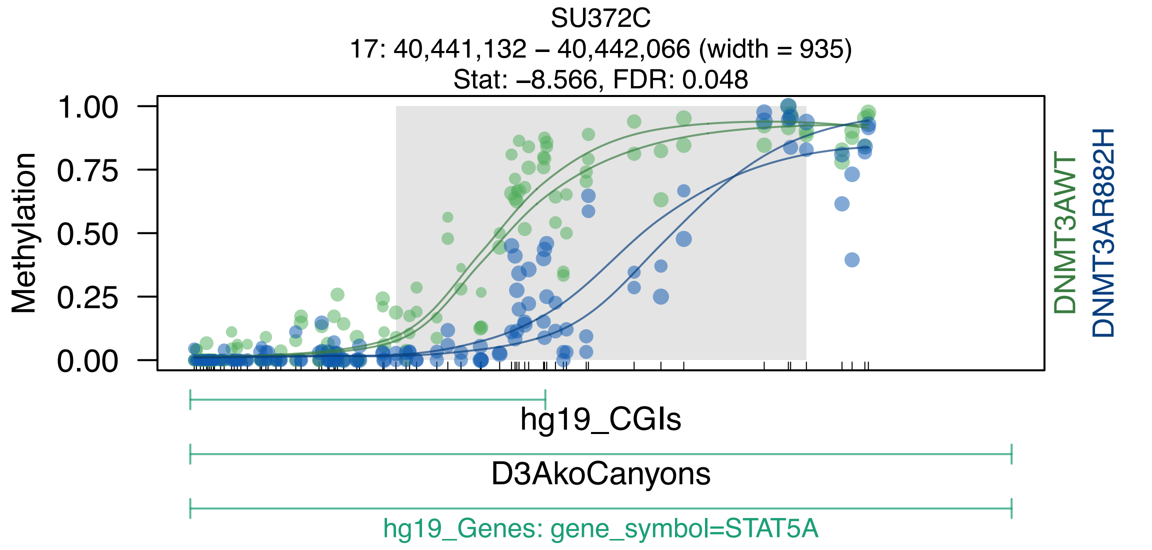

SU372C
17: 40,441,132 – 40,442,066 (width = 935)
Stat: –8.566, FDR: 0.048
Methylation
1.00
0.75
0.50
0.25
0.00
DNMT3AWT
DNMT3AR882H
hg19_CGIs
D3AkoCanyons
hg19_Genes: gene_symbol=STAT5A

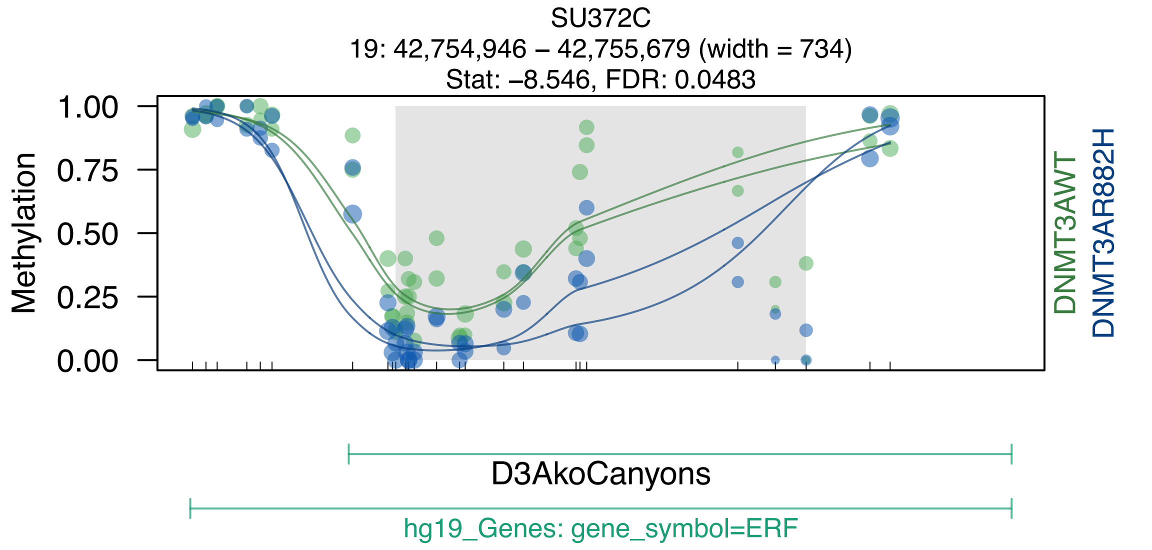
SU372C
19: 42,754,946 – 42,755,679 (width = 734)
Stat: –8.546, FDR: 0.0483
1.00
0.75
0.50
0.25
0.00
Methylation
DNMT3AWT
DNMT3AR882H
D3AkoCanyons
hg19_Genes: gene_symbol=ERF
