## Supplemental Data S1-S7 for "DNMT3A^R882H^ Is Not Required for Disease Maintenance in Primary Human AML, but Is Associated With Increased Leukemia Stem Cell Frequency": DataS5_SU540_top50dmrsOverlappingCanyons.pdf

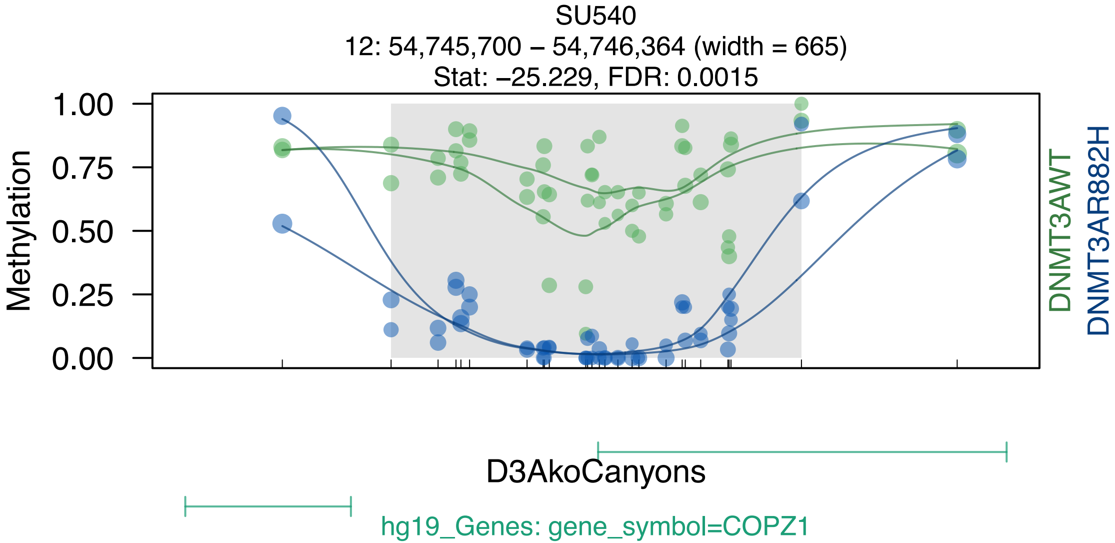
SU540
12: 54,745,700 – 54,746,364 (width = 665)
Stat: –25.229, FDR: 0.0015
Methylation
1.00
0.75
0.50
0.25
0.00
DNMT3AWT
DNMT3AR882H
D3AkoCanyons
hg19_Genes: gene_symbol=COPZ1

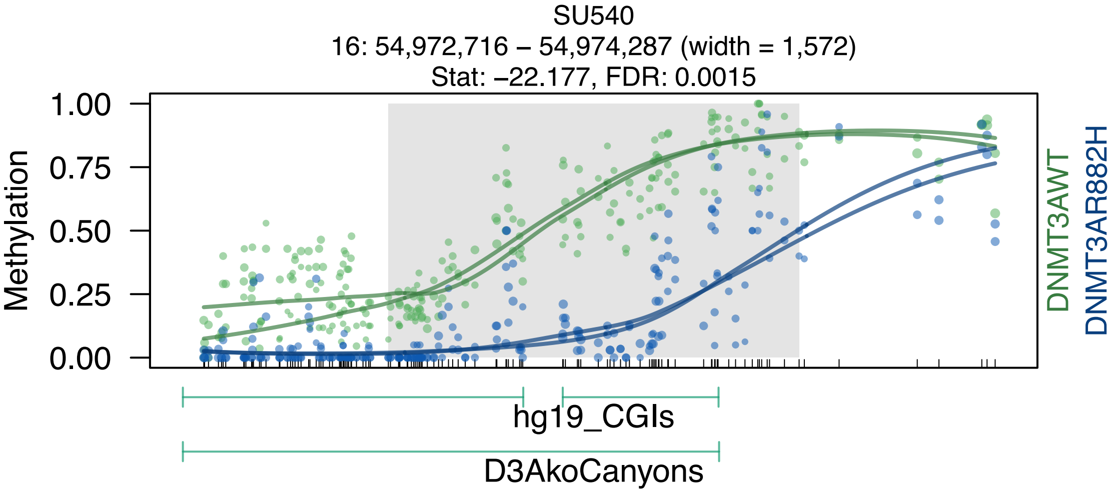
SU540
16: 54,972,716 – 54,974,287 (width = 1,572)
Stat: –22.177, FDR: 0.0015
Methylation
1.00
0.75
0.50
0.25
0.00
DNMT3AWT
DNMT3AR882H
hg19_CGIs
D3AkoCanyons

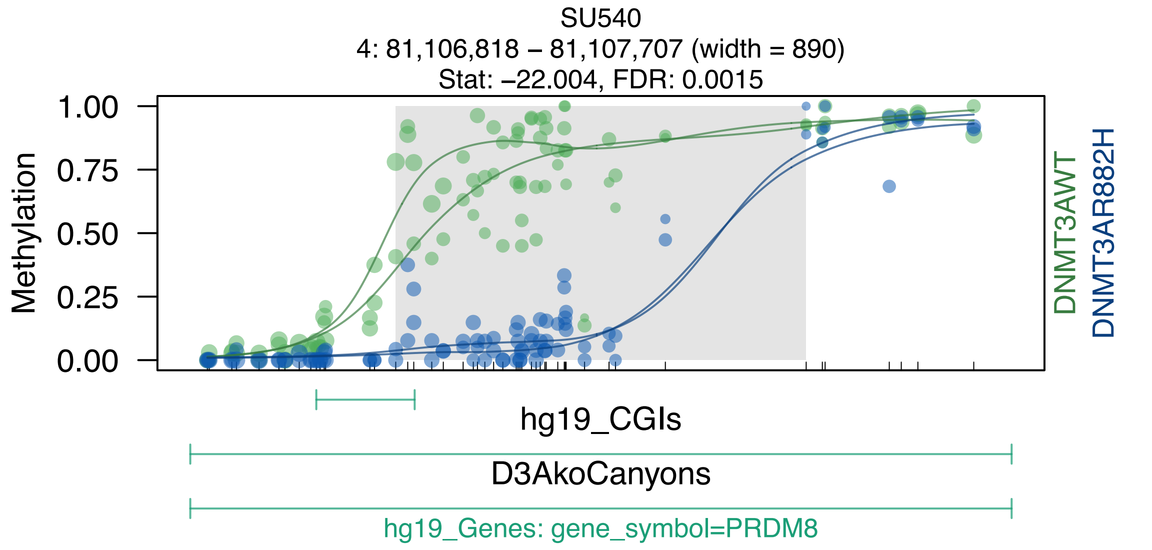
SU540
4: 81,106,818 – 81,107,707 (width = 890)
Stat: −22.004, FDR: 0.0015
1.00
0.75
0.50
0.25
0.00
Methylation
DNMT3AWT
DNMT3AR882H
hg19_CGIs
D3AkoCanyons
hg19_Genes: gene_symbol=PRDM8

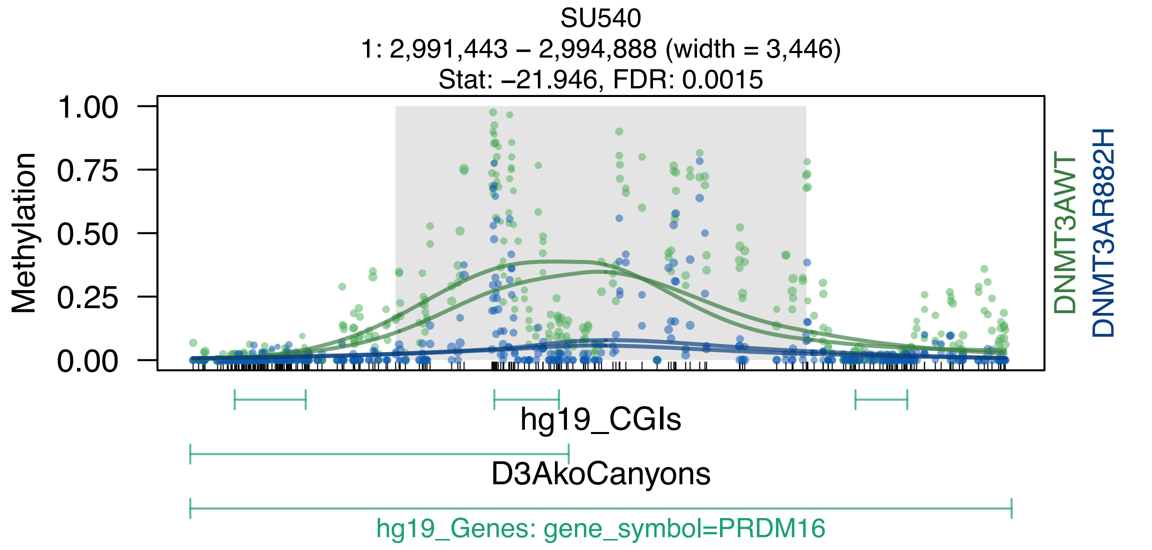
SU540
1: 2,991,443 – 2,994,888 (width = 3,446)
Stat: –21.946, FDR: 0.0015
Methylation
1.00
0.75
0.50
0.25
0.00
DNMT3AWT
DNMT3AR882H
hg19_CGIs
D3AkoCanyons
hg19_Genes: gene_symbol=PRDM16

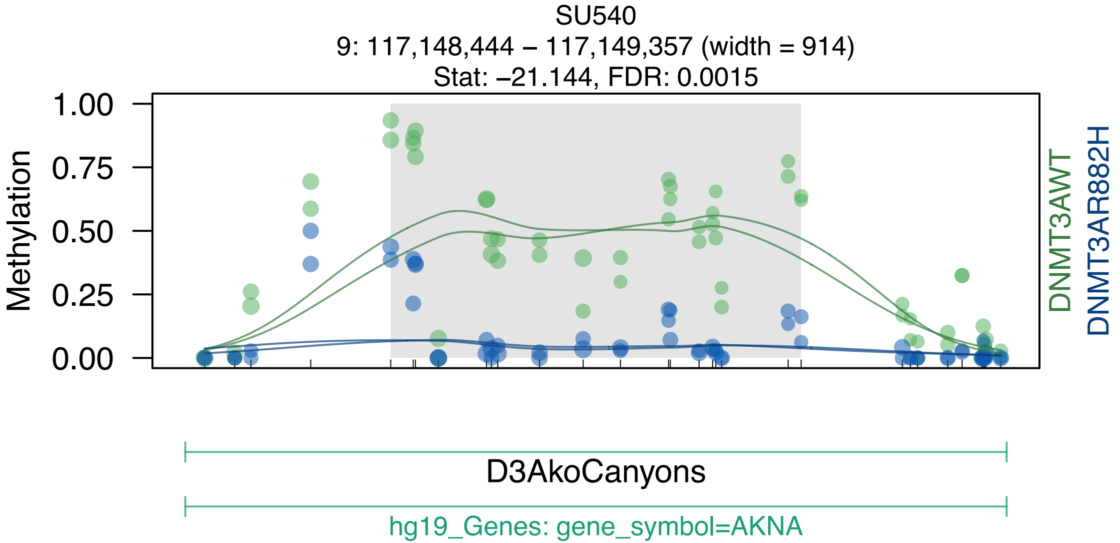
SU540
9: 117,148,444 – 117,149,357 (width = 914)
Stat: –21.144, FDR: 0.0015
Methylation
1.00
0.75
0.50
0.25
0.00
DNMT3AWT
DNMT3AR882H
D3AkoCanyons
hg19_Genes: gene_symbol=AKNA

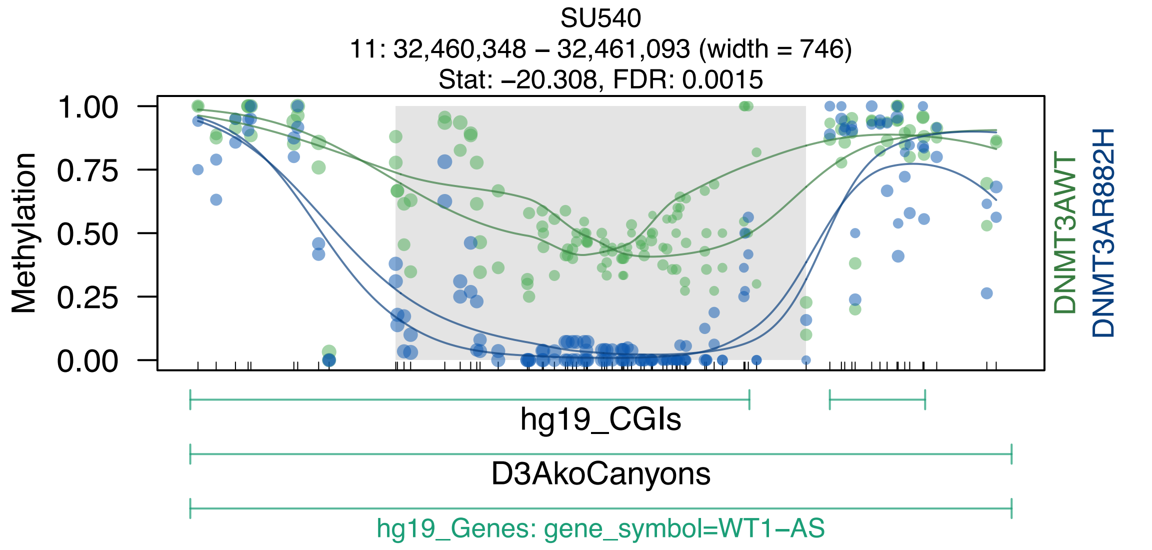
SU540
11: 32,460,348 – 32,461,093 (width = 746)
Stat: –20.308, FDR: 0.0015
Methylation
1.00
0.75
0.50
0.25
0.00
DNMT3AWT
DNMT3AR882H
hg19_CGIs
D3AkoCanyons
hg19_Genes: gene_symbol=WT1–AS

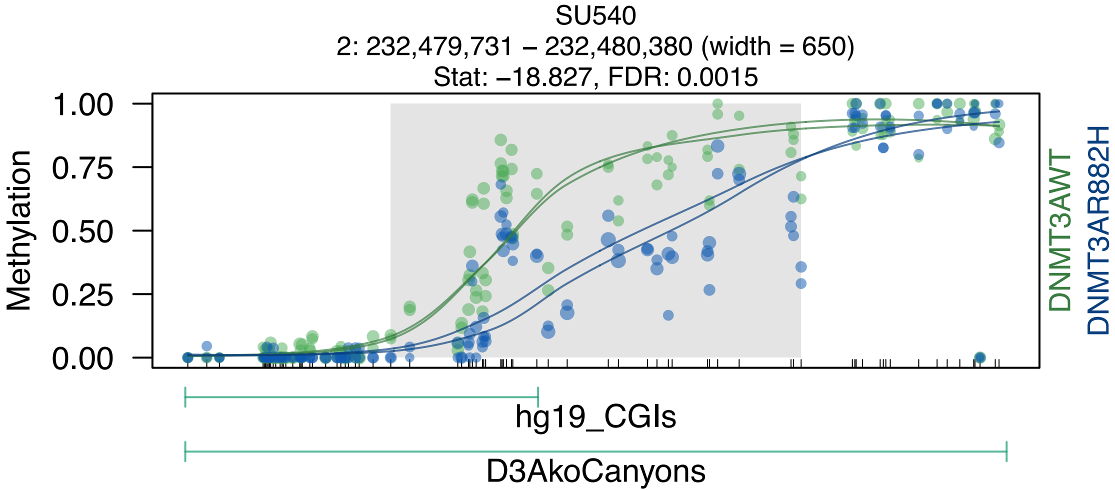
SU540
2: 232,479,731 – 232,480,380 (width = 650)
Stat: –18.827, FDR: 0.0015
1.00
0.75
0.50
0.25
0.00
Methylation
DNMT3AWT
DNMT3AR882H
hg19_CGIs
D3AkoCanyons

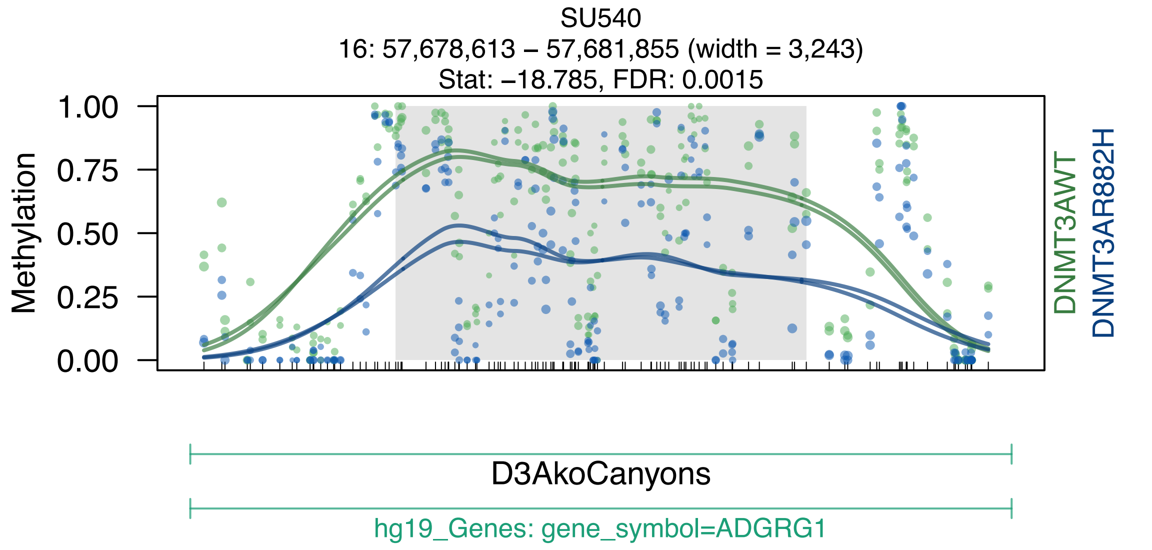
SU540
16: 57,678,613 – 57,681,855 (width = 3,243)
Stat: –18.785, FDR: 0.0015
Methylation
1.00
0.75
0.50
0.25
0.00
DNMT3AWT
DNMT3AR882H
D3AkoCanyons
hg19_Genes: gene_symbol=ADGRG1

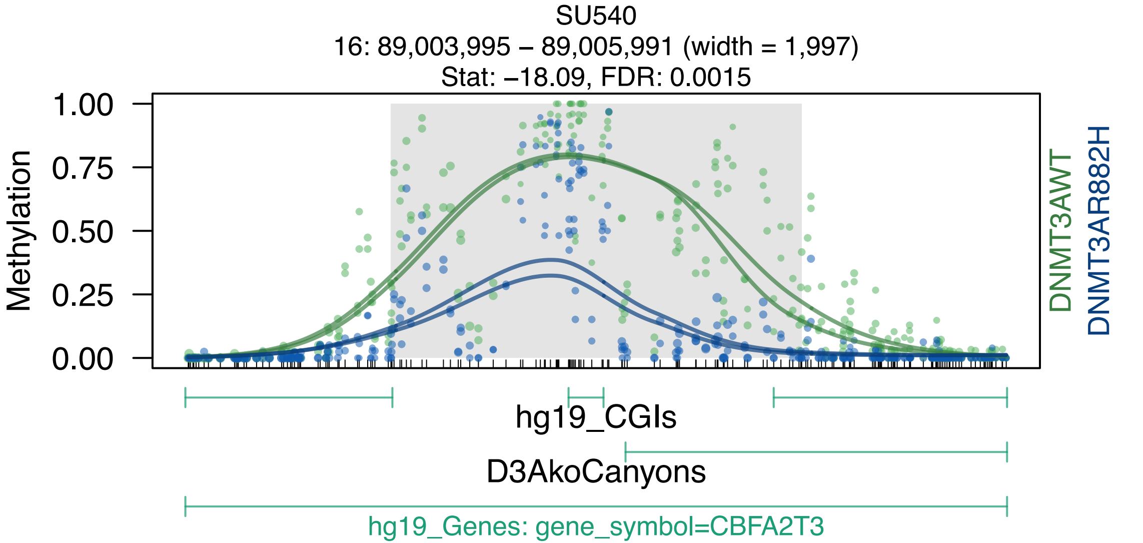

SU540
16: 89,003,995 – 89,005,991 (width = 1,997)
Stat: –18.09, FDR: 0.0015
Methylation
1.00
0.75
0.50
0.25
0.00
DNMT3AWT
DNMT3AR882H
hg19_CGIs
D3AkoCanyons
hg19_Genes: gene_symbol=CBFA2T3

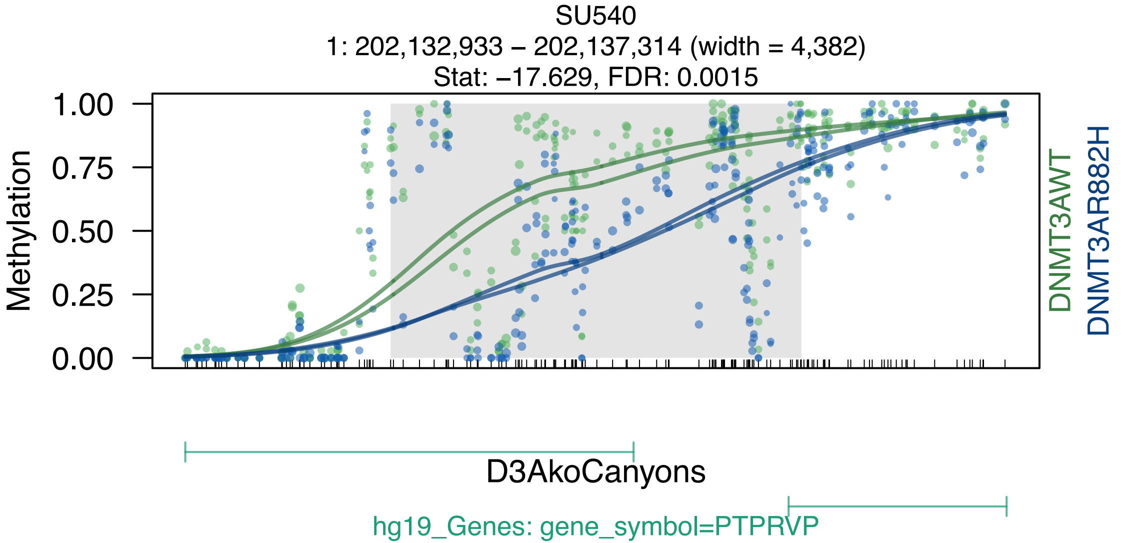
SU540
1: 202,132,933 – 202,137,314 (width = 4,382)
Stat: –17.629, FDR: 0.0015
Methylation
1.00
0.75
0.50
0.25
0.00
DNMT3AWT
DNMT3AR882H
D3AkoCanyons
hg19_Genes: gene_symbol=PTPRVP

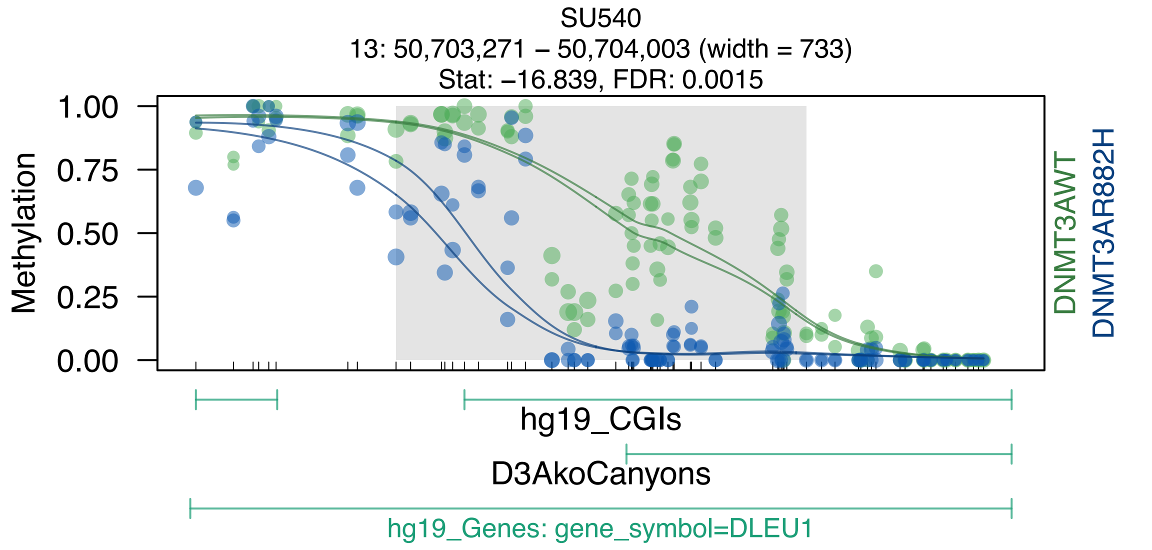
SU540
13: 50,703,271 – 50,704,003 (width = 733)
Stat: –16.839, FDR: 0.0015
Methylation
1.00
0.75
0.50
0.25
0.00
DNMT3AWT
DNMT3AR882H
hg19_CGIs
D3AkoCanyons
hg19_Genes: gene_symbol=DLEU1

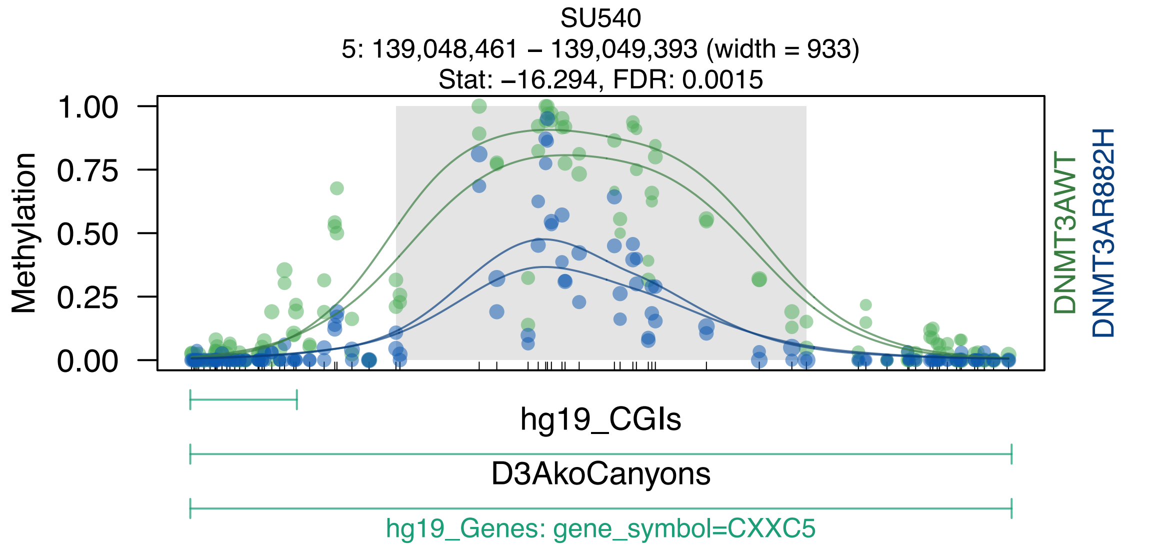
SU540
5: 139,048,461 – 139,049,393 (width = 933)
Stat: –16.294, FDR: 0.0015
1.00
0.75
0.50
0.25
0.00
Methylation
DNMT3AWT
DNMT3AR882H
hg19_CGIs
D3AkoCanyons
hg19_Genes: gene_symbol=CXXC5

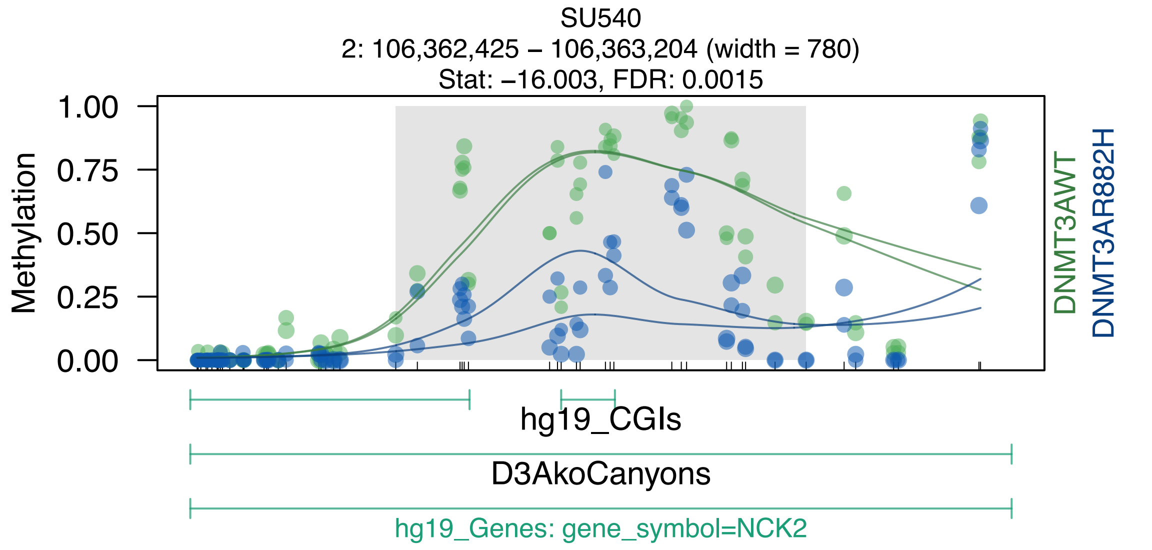
SU540
2: 106,362,425 – 106,363,204 (width = 780)
Stat: –16.003, FDR: 0.0015
Methylation
1.00
0.75
0.50
0.25
0.00
DNMT3AWT
DNMT3AR882H
hg19_CGIs
D3AkoCanyons
hg19_Genes: gene_symbol=NCK2

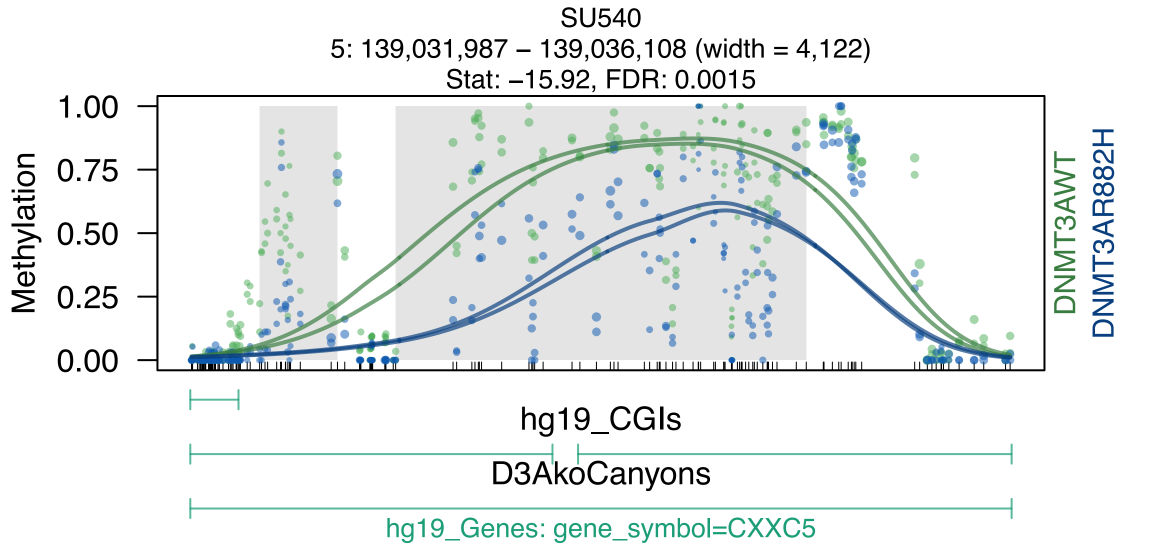

SU540
5: 139,031,987 – 139,036,108 (width = 4,122)
Stat: −15.92, FDR: 0.0015
Methylation
1.00
0.75
0.50
0.25
0.00
DNMT3AWT
DNMT3AR882H
hg19_CGIs
D3AkoCanyons
hg19_Genes: gene_symbol=CXXC5

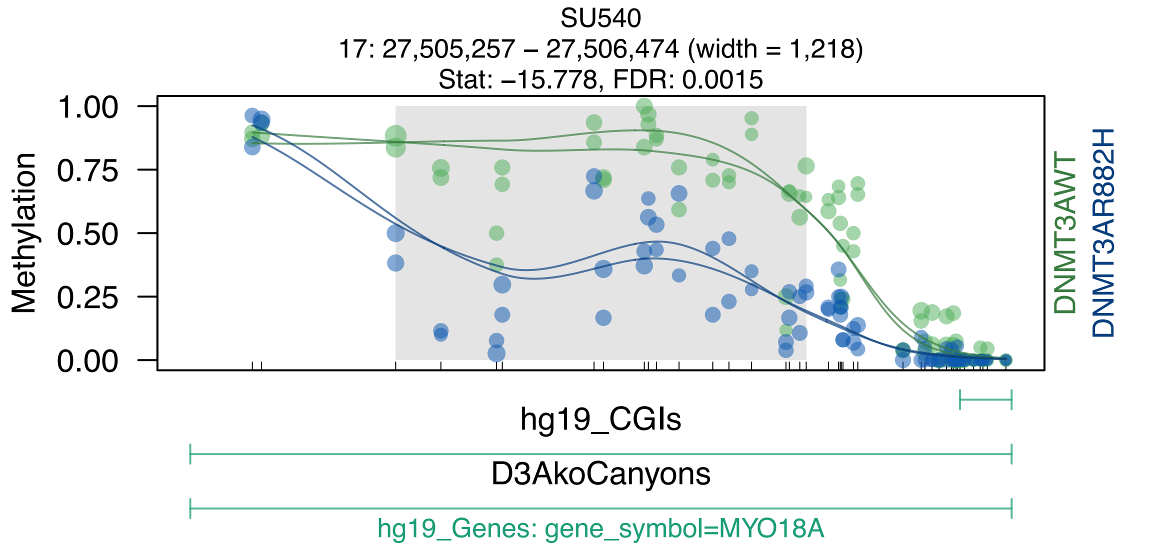
SU540
17: 27,505,257 – 27,506,474 (width = 1,218)
Stat: –15.778, FDR: 0.0015
Methylation
1.00
0.75
0.50
0.25
0.00
DNMT3AWT
DNMT3AR882H
hg19_CGIs
D3AkoCanyons
hg19_Genes: gene_symbol=MYO18A

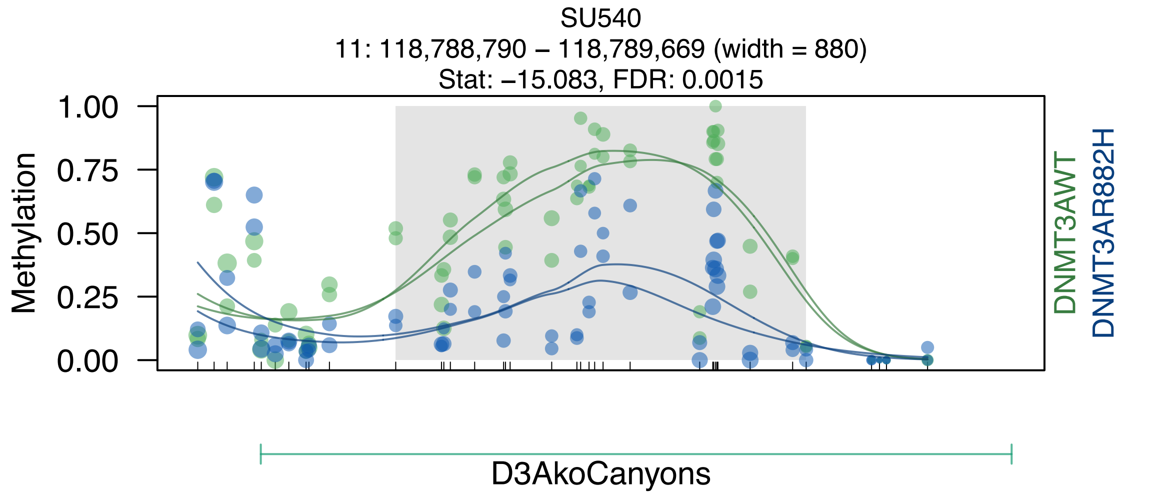

SU540
11: 118,788,790 – 118,789,669 (width = 880)
Stat: –15.083, FDR: 0.0015
Methylation
1.00
0.75
0.50
0.25
0.00
DNMT3AWT
DNMT3AR882H
D3AkoCanyons

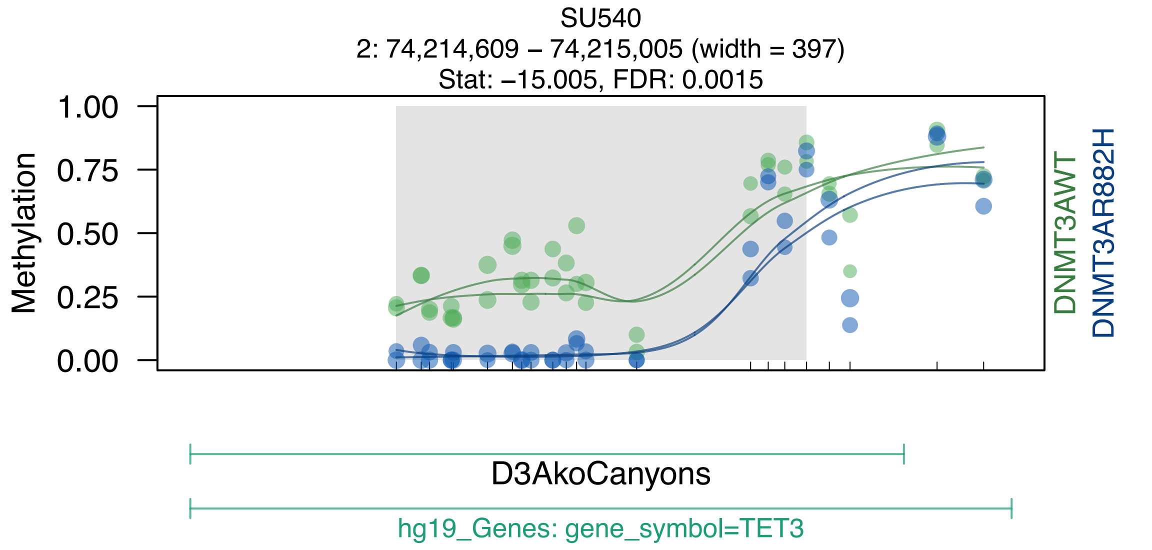
SU540
2: 74,214,609 – 74,215,005 (width = 397)
Stat: –15.005, FDR: 0.0015
1.00
0.75
0.50
0.25
0.00
Methylation
DNMT3AWT
DNMT3AR882H
D3AkoCanyons
hg19_Genes: gene_symbol=TET3

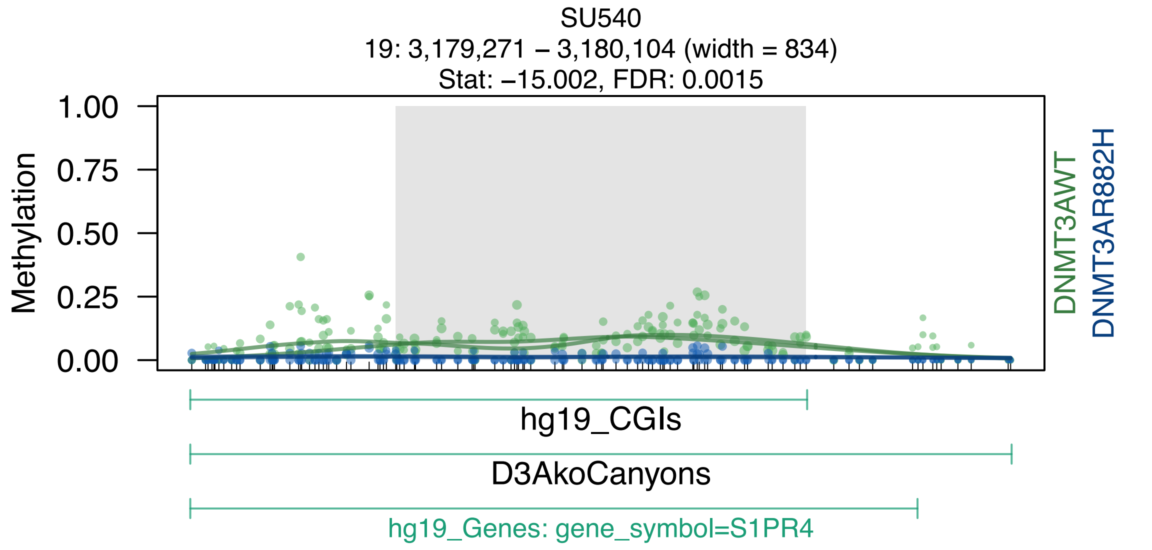
SU540
19: 3,179,271 – 3,180,104 (width = 834)
Stat: –15.002, FDR: 0.0015
Methylation
1.00
0.75
0.50
0.25
0.00
DNMT3AWT
DNMT3AR882H
hg19_CGIs
D3AkoCanyons
hg19_Genes: gene_symbol=S1PR4

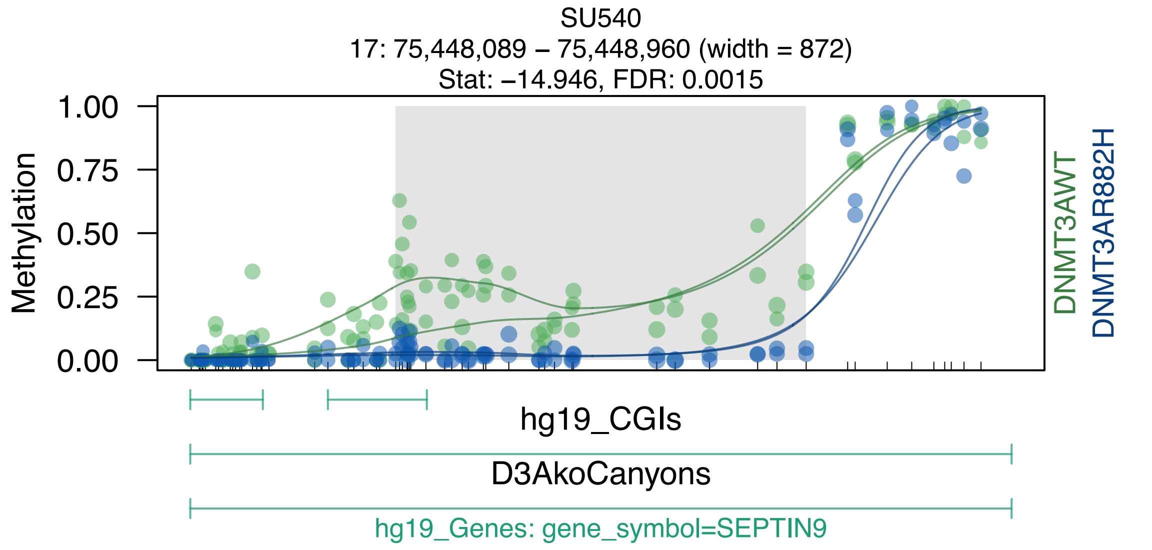
SU540
17: 75,448,089 – 75,448,960 (width = 872)
Stat: –14.946, FDR: 0.0015
Methylation
1.00
0.75
0.50
0.25
0.00
DNMT3AWT
DNMT3AR882H
hg19_CGIs
D3AkoCanyons
hg19_Genes: gene_symbol=SEPTIN9

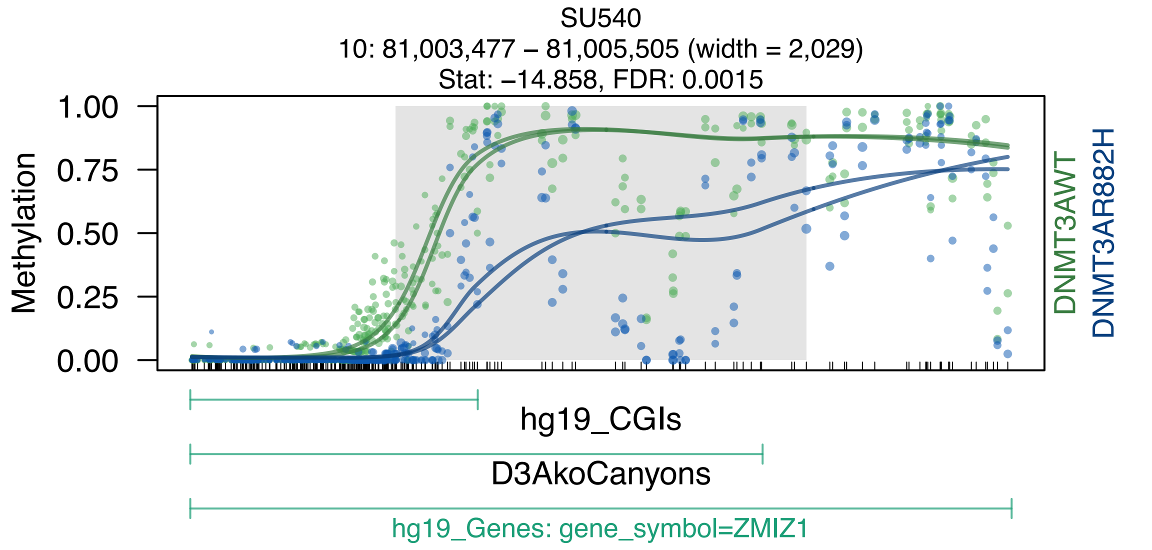
SU540
10: 81,003,477 – 81,005,505 (width = 2,029)
Stat: –14.858, FDR: 0.0015
Methylation
1.00
0.75
0.50
0.25
0.00
DNMT3AWT
DNMT3AR882H
hg19_CGIs
D3AkoCanyons
hg19_Genes: gene_symbol=ZMIZ1

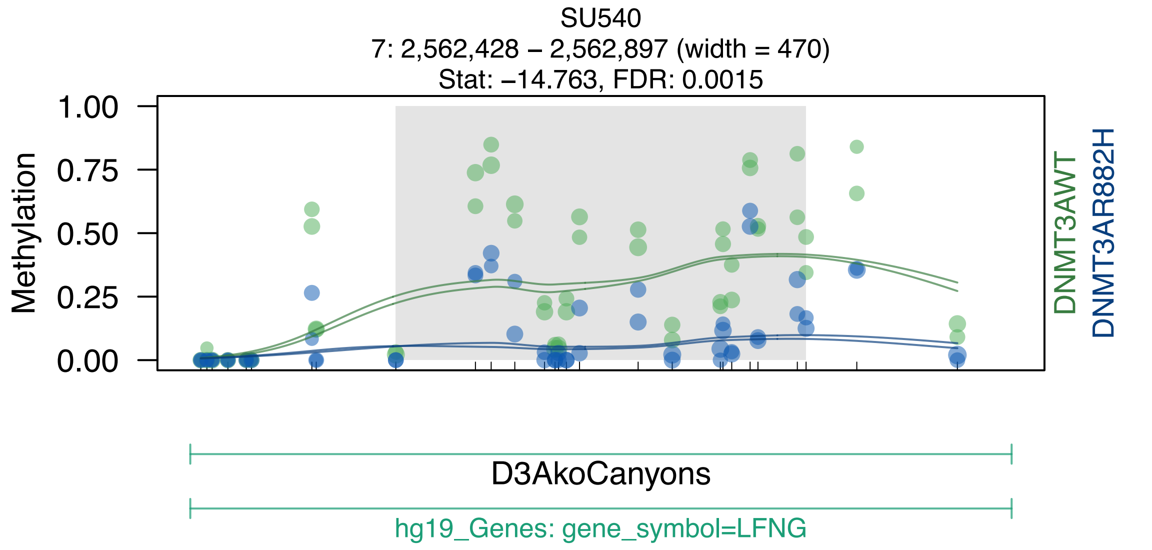
SU540
7: 2,562,428 – 2,562,897 (width = 470)
Stat: –14.763, FDR: 0.0015
Methylation
1.00
0.75
0.50
0.25
0.00
DNMT3AWT
DNMT3AR882H
D3AkoCanyons
hg19_Genes: gene_symbol=LFNG

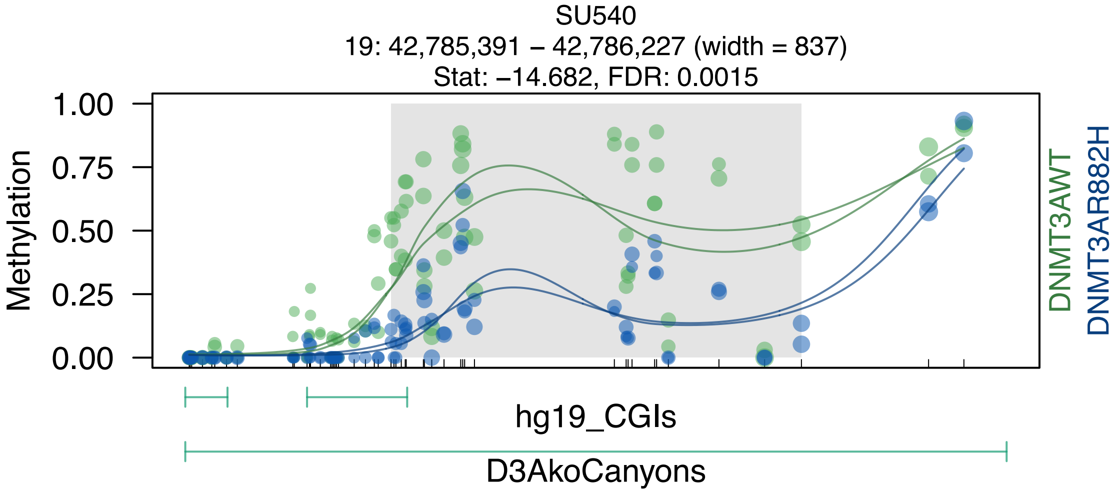

SU540
19: 42,785,391 – 42,786,227 (width = 837)
Stat: –14.682, FDR: 0.0015
Methylation
1.00
0.75
0.50
0.25
0.00
DNMT3AWT
DNMT3AR882H
hg19_CGIs
D3AkoCanyons

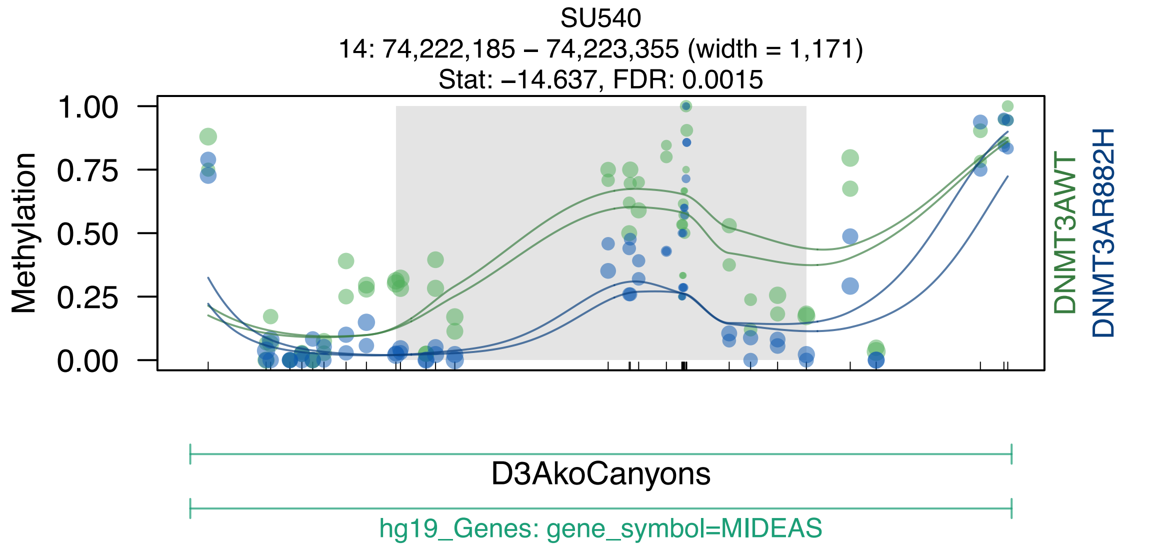
SU540
14: 74,222,185 – 74,223,355 (width = 1,171)
Stat: −14.637, FDR: 0.0015
Methylation
1.00
0.75
0.50
0.25
0.00
DNMT3AWT
DNMT3AR882H
D3AkoCanyons
hg19_Genes: gene_symbol=MIDEAS

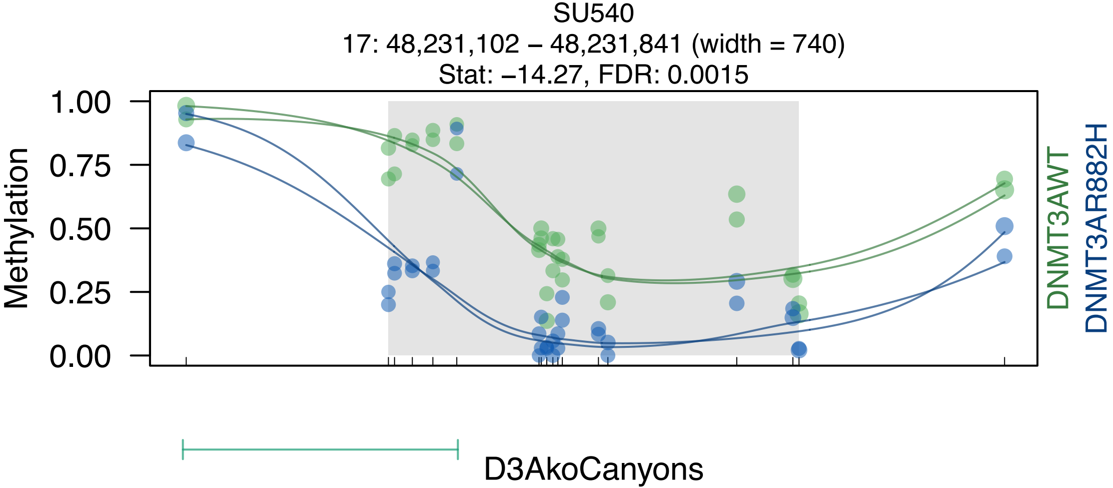
SU540
17: 48,231,102 – 48,231,841 (width = 740)
Stat: –14.27, FDR: 0.0015
Methylation
1.00
0.75
0.50
0.25
0.00
DNMT3AWT
DNMT3AR882H
D3AkoCanyons

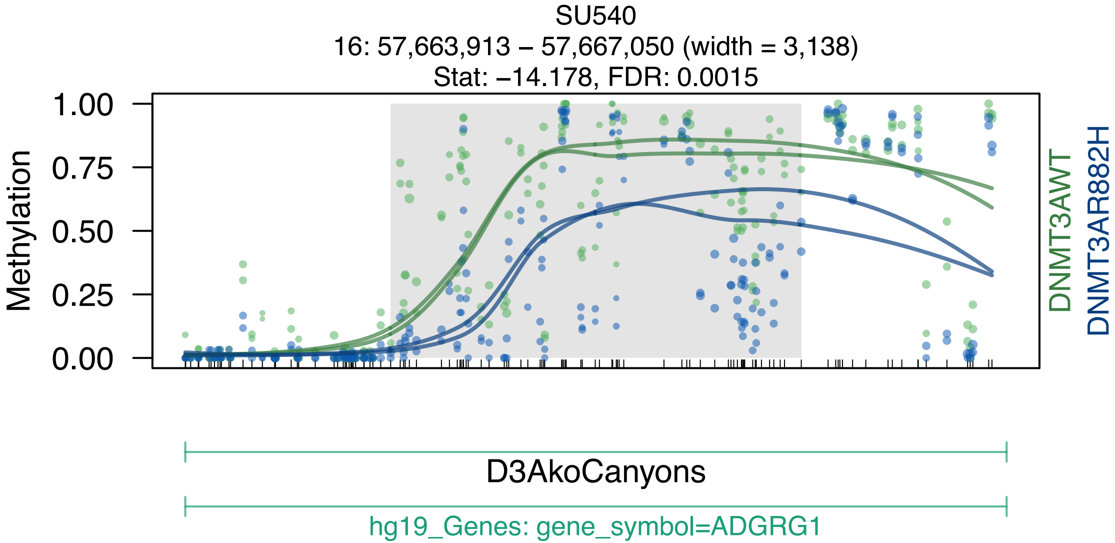
SU540
16: 57,663,913 – 57,667,050 (width = 3,138)
Stat: –14.178, FDR: 0.0015
Methylation
1.00
0.75
0.50
0.25
0.00
DNMT3AWT
DNMT3AR882H
D3AkoCanyons
hg19_Genes: gene_symbol=ADGRG1

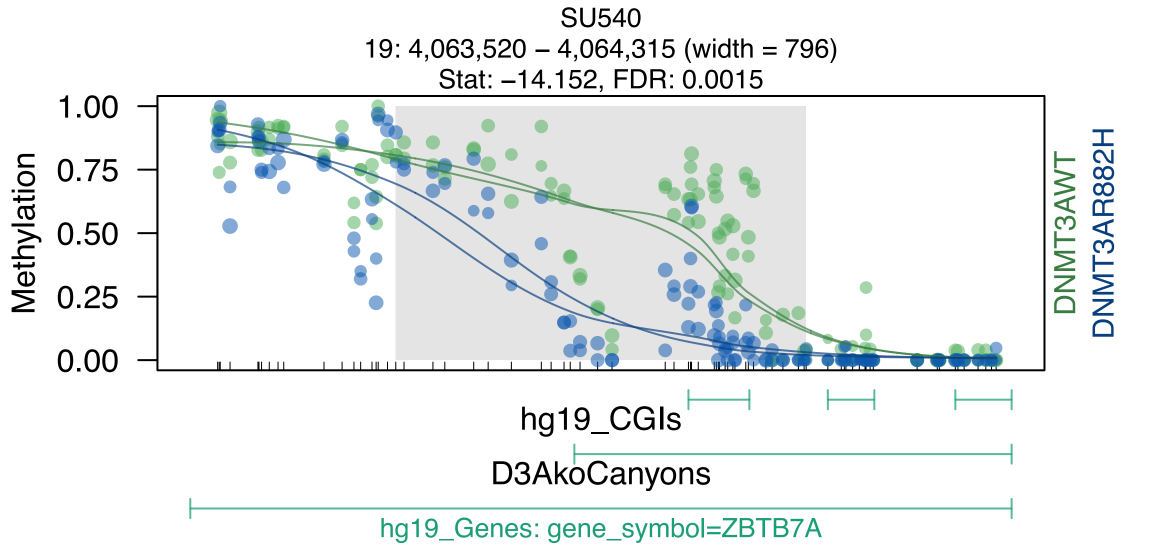
SU540
19: 4,063,520 – 4,064,315 (width = 796)
Stat: –14.152, FDR: 0.0015
1.00
0.75
0.50
0.25
0.00
Methylation
DNMT3AWT
DNMT3AR882H
hg19_CGIs
D3AkoCanyons
hg19_Genes: gene_symbol=ZBTB7A

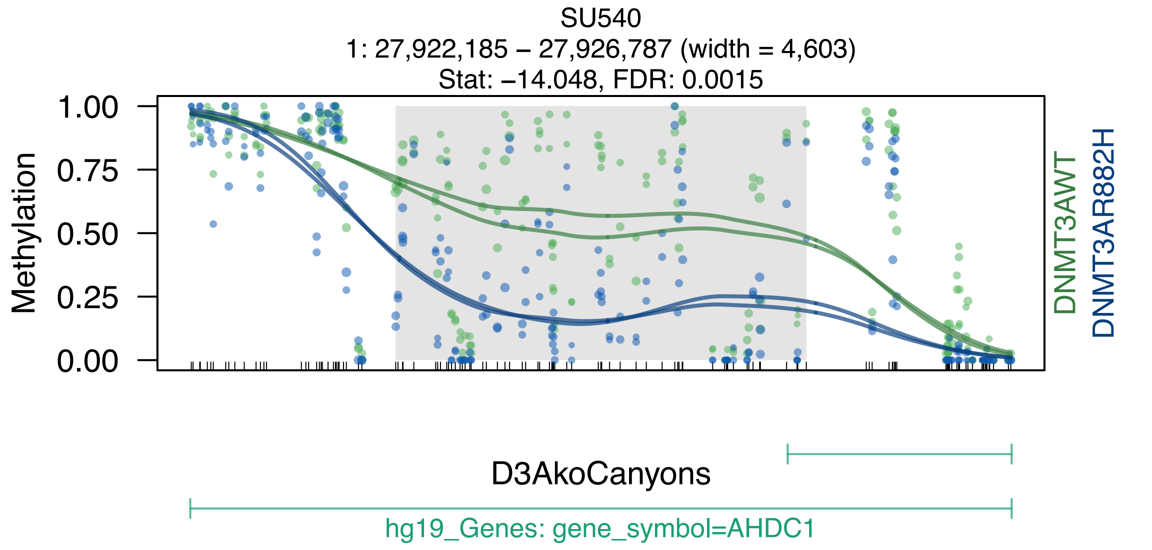
SU540
1: 27,922,185 – 27,926,787 (width = 4,603)
Stat: –14.048, FDR: 0.0015
Methylation
1.00
0.75
0.50
0.25
0.00
DNMT3AWT
DNMT3AR882H
D3AkoCanyons
hg19_Genes: gene_symbol=AHDC1

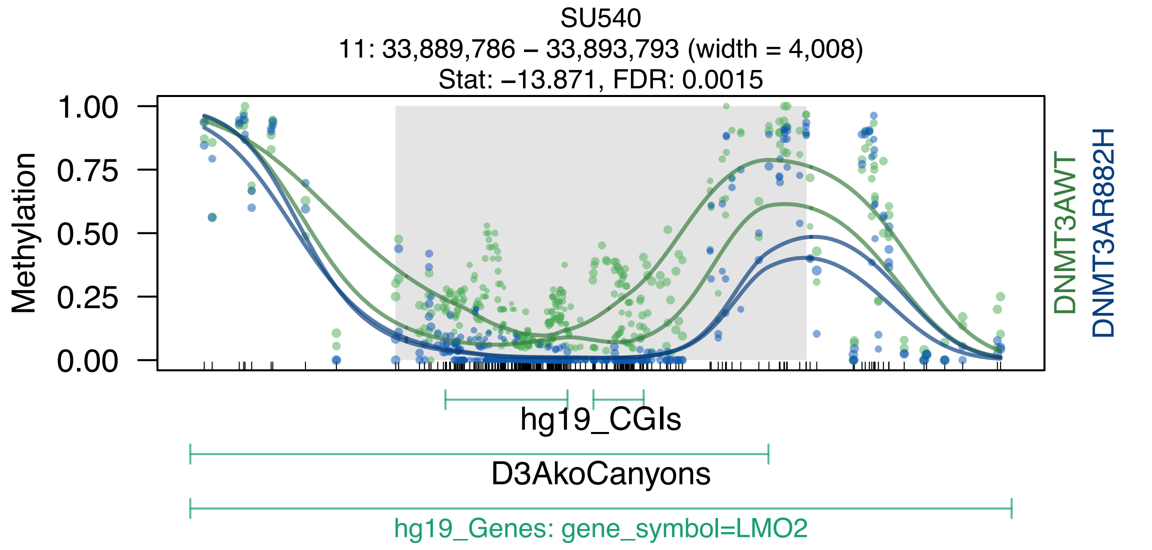
SU540
11: 33,889,786 – 33,893,793 (width = 4,008)
Stat: –13.871, FDR: 0.0015
Methylation
1.00
0.75
0.50
0.25
0.00
DNMT3AWT
DNMT3AR882H
hg19_CGIs
D3AkoCanyons
hg19_Genes: gene_symbol=LMO2

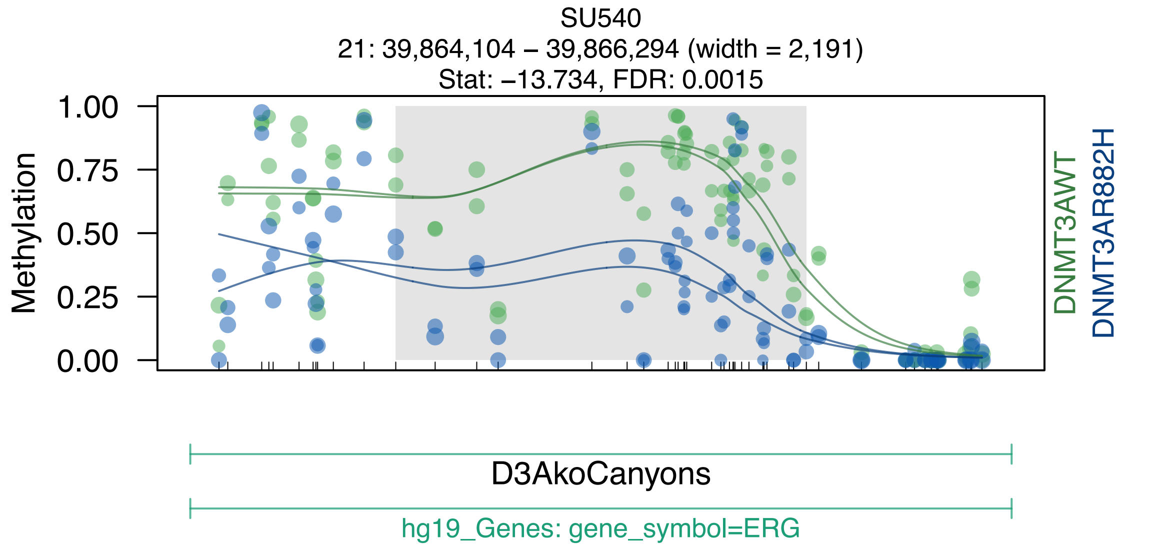
SU540
21: 39,864,104 – 39,866,294 (width = 2,191)
Stat: –13.734, FDR: 0.0015
Methylation
1.00
0.75
0.50
0.25
0.00
DNMT3AWT
DNMT3AR882H
D3AkoCanyons
hg19_Genes: gene_symbol=ERG

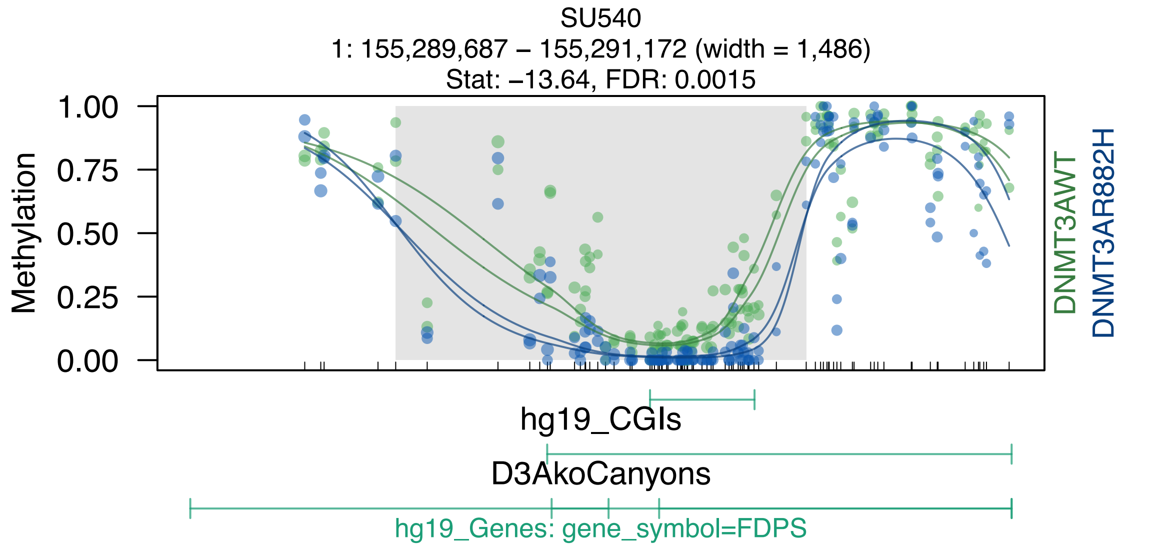
SU540
1: 155,289,687 – 155,291,172 (width = 1,486)
Stat: –13.64, FDR: 0.0015
Methylation
1.00
0.75
0.50
0.25
0.00
DNMT3AWT
DNMT3AR882H
hg19_CGIs
D3AkoCanyons
hg19_Genes: gene_symbol=FDPS

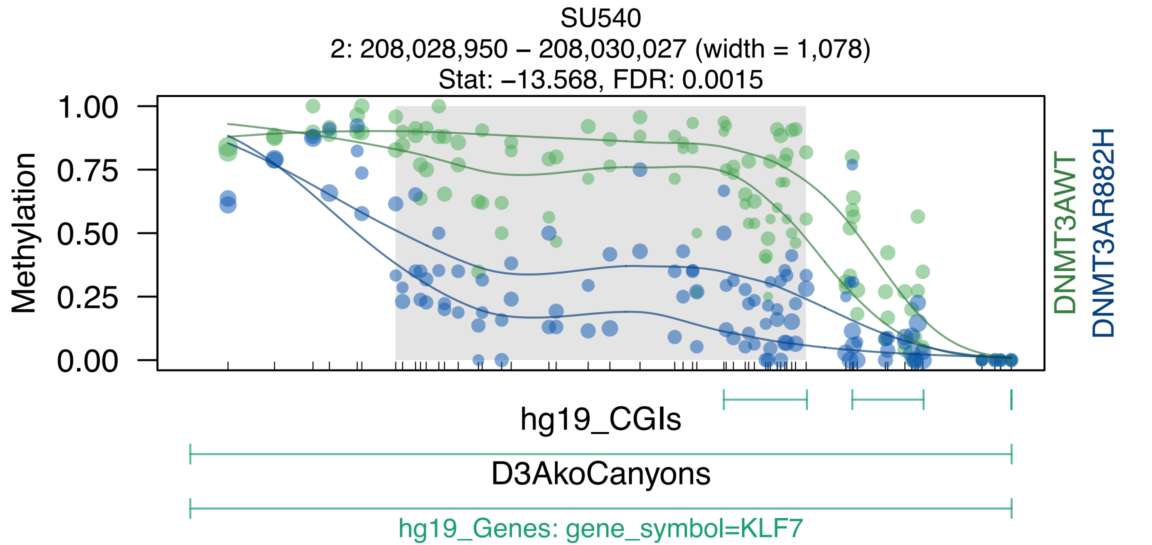
SU540
2: 208,028,950 – 208,030,027 (width = 1,078)
Stat: −13.568, FDR: 0.0015
Methylation
1.00
0.75
0.50
0.25
0.00
DNMT3AWT
DNMT3AR882H
hg19_CGIs
D3AkoCanyons
hg19_Genes: gene_symbol=KLF7

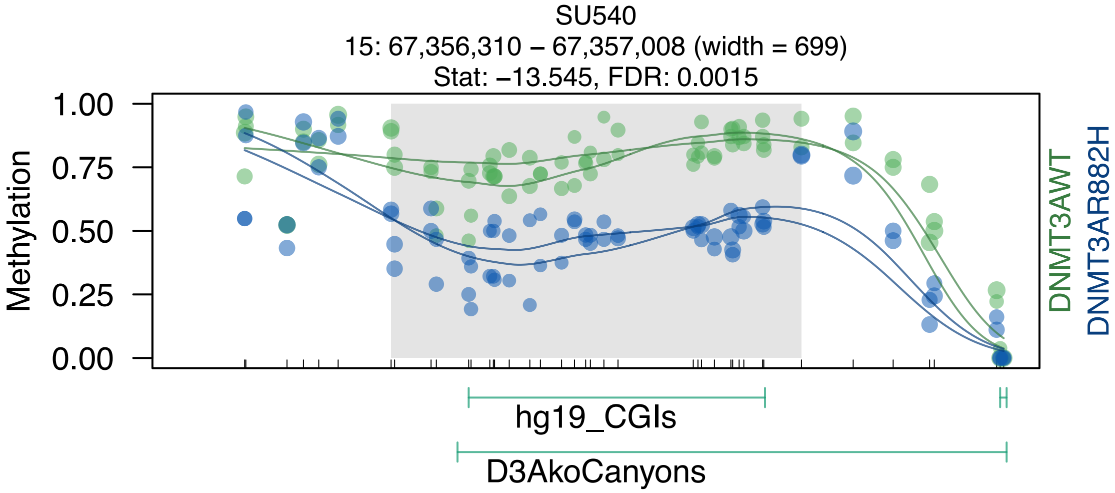
SU540
15: 67,356,310 – 67,357,008 (width = 699)
Stat: –13.545, FDR: 0.0015
Methylation
1.00
0.75
0.50
0.25
0.00
DNMT3AWT
DNMT3AR882H
hg19_CGIs
D3AkoCanyons

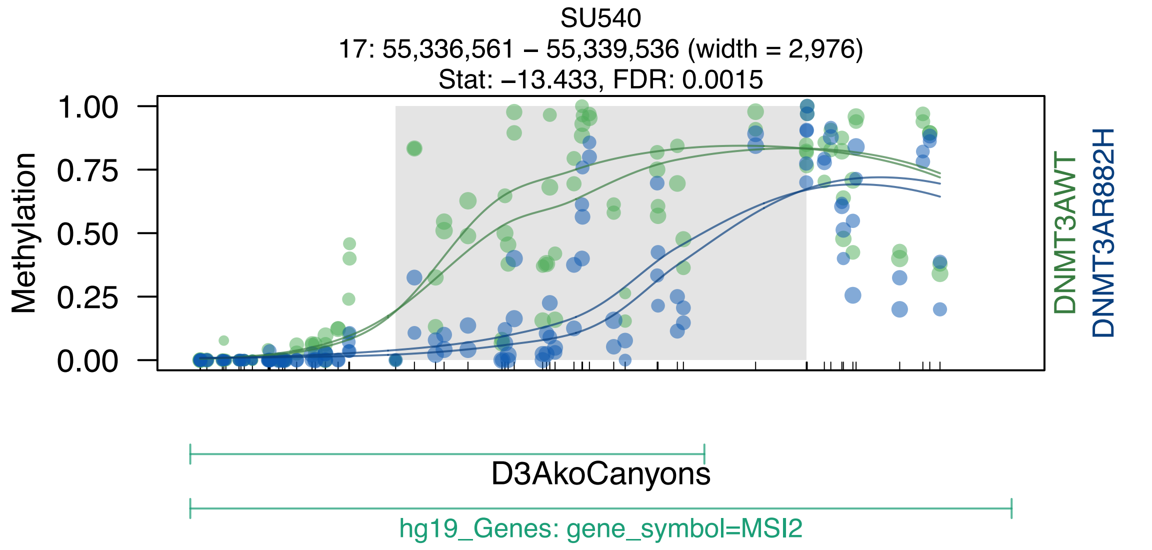
SU540
17: 55,336,561 – 55,339,536 (width = 2,976)
Stat: –13.433, FDR: 0.0015
Methylation
1.00
0.75
0.50
0.25
0.00
DNMT3AWT
DNMT3AR882H
D3AkoCanyons
hg19_Genes: gene_symbol=MSI2

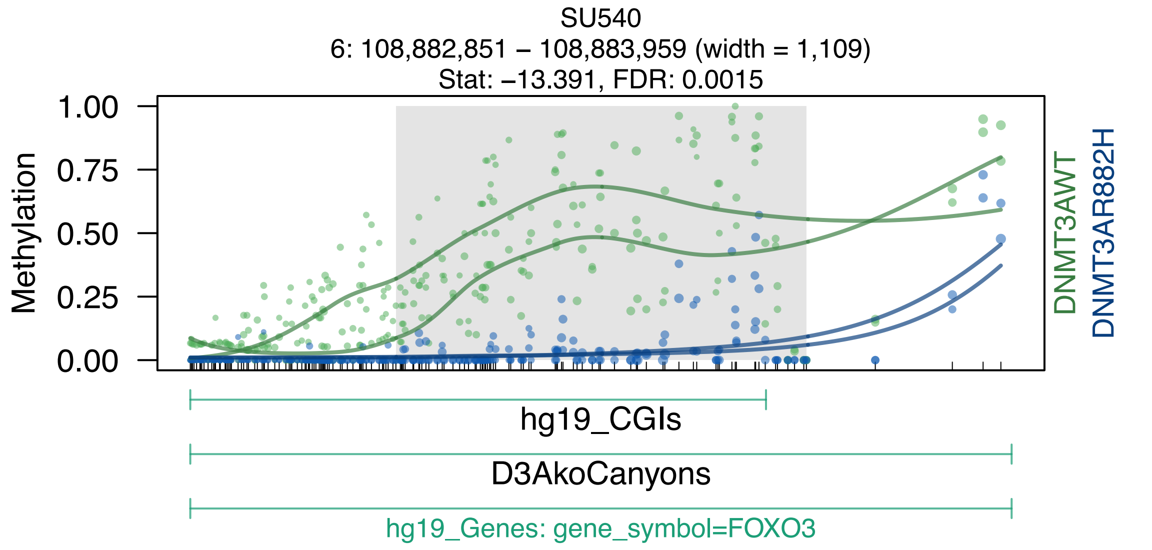
SU540
6: 108,882,851 – 108,883,959 (width = 1,109)
Stat: –13.391, FDR: 0.0015
Methylation
1.00
0.75
0.50
0.25
0.00
DNMT3AWT
DNMT3AR882H
hg19_CGIs
D3AkoCanyons
hg19_Genes: gene_symbol=FOXO3

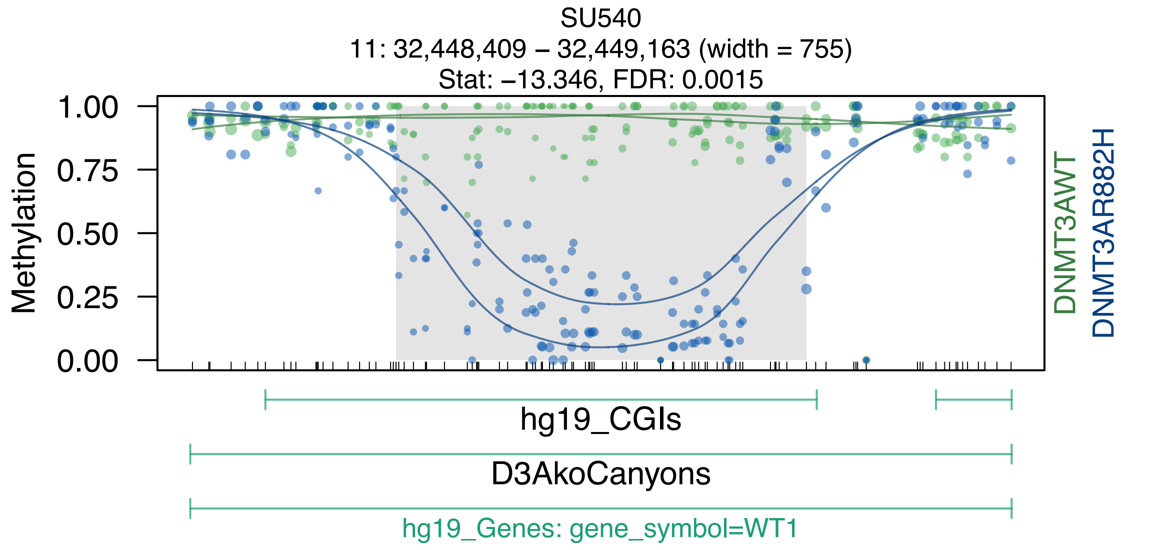

SU540
11: 32,448,409 – 32,449,163 (width = 755)
Stat: –13.346, FDR: 0.0015
Methylation
1.00
0.75
0.50
0.25
0.00
DNMT3AWT
DNMT3AR882H
hg19_CGIs
D3AkoCanyons
hg19_Genes: gene_symbol=WT1

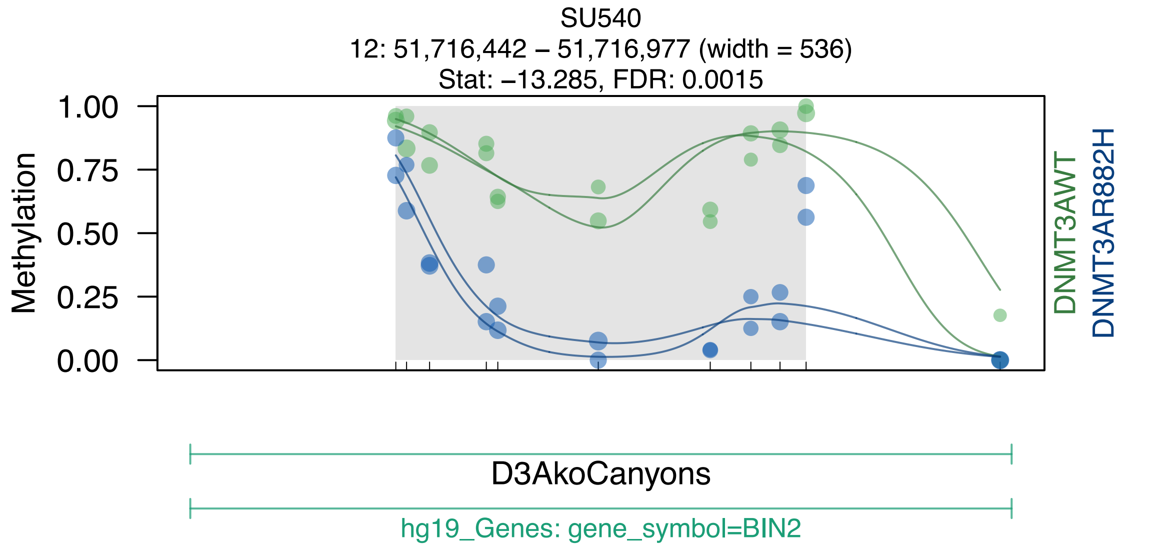
SU540
12: 51,716,442 – 51,716,977 (width = 536)
Stat: –13.285, FDR: 0.0015
Methylation
1.00
0.75
0.50
0.25
0.00
DNMT3AWT
DNMT3AR882H
D3AkoCanyons
hg19_Genes: gene_symbol=BIN2

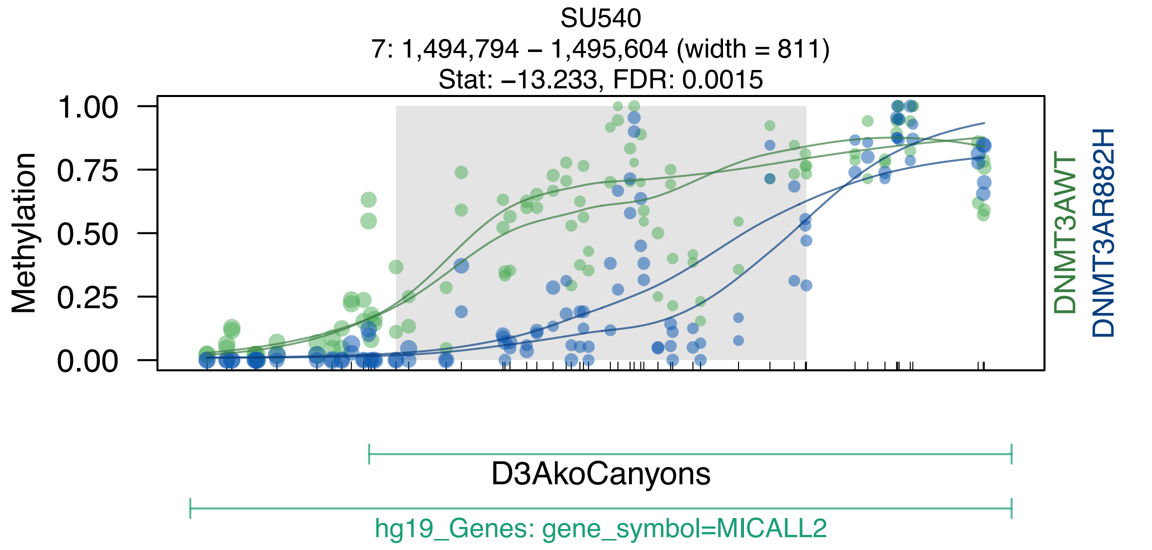
SU540
7: 1,494,794 – 1,495,604 (width = 811)
Stat: –13.233, FDR: 0.0015
Methylation
1.00
0.75
0.50
0.25
0.00
DNMT3AWT
DNMT3AR882H
D3AkoCanyons
hg19_Genes: gene_symbol=MICALL2

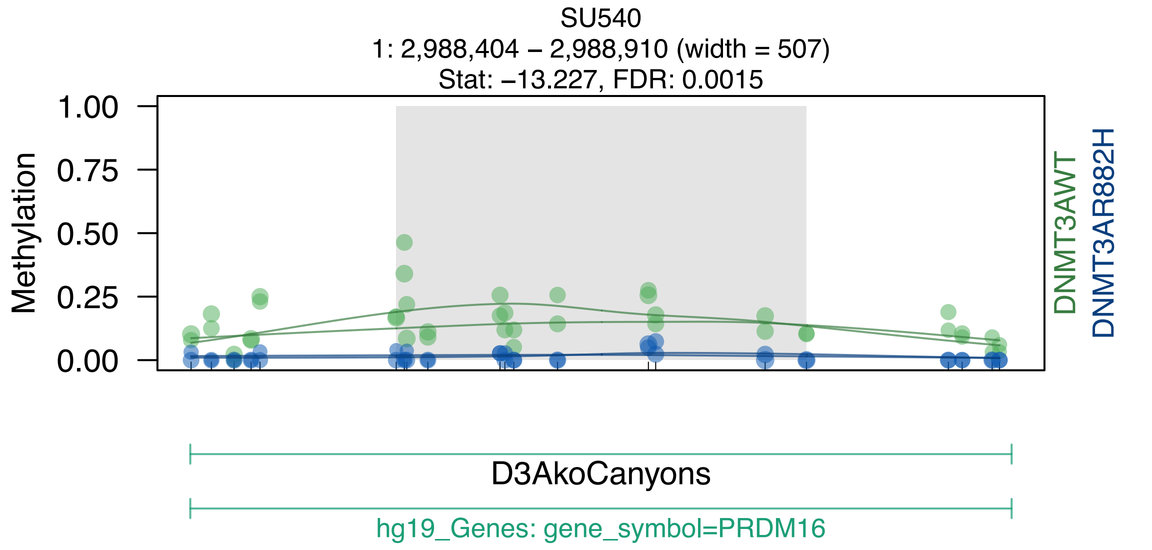

SU540
1: 2,988,404 – 2,988,910 (width = 507)
Stat: –13.227, FDR: 0.0015
1.00
0.75
0.50
0.25
0.00
Methylation
DNMT3AWT
DNMT3AR882H
D3AkoCanyons
hg19_Genes: gene_symbol=PRDM16

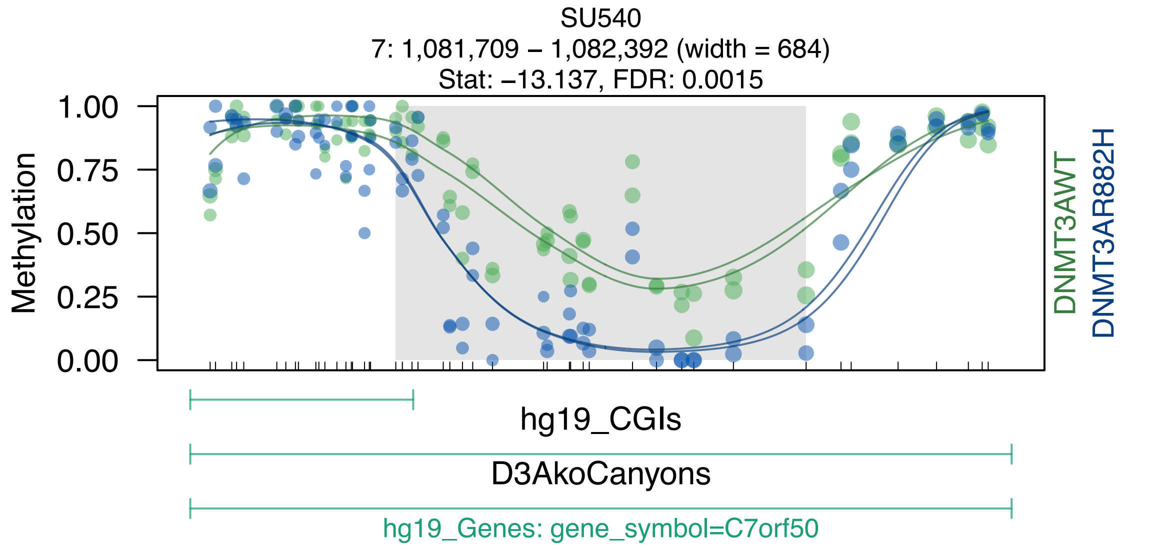
SU540
7: 1,081,709 – 1,082,392 (width = 684)
Stat: −13.137, FDR: 0.0015
Methylation
1.00
0.75
0.50
0.25
0.00
DNMT3AWT
DNMT3AR882H
hg19_CGIs
D3AkoCanyons
hg19_Genes: gene_symbol=C7orf50

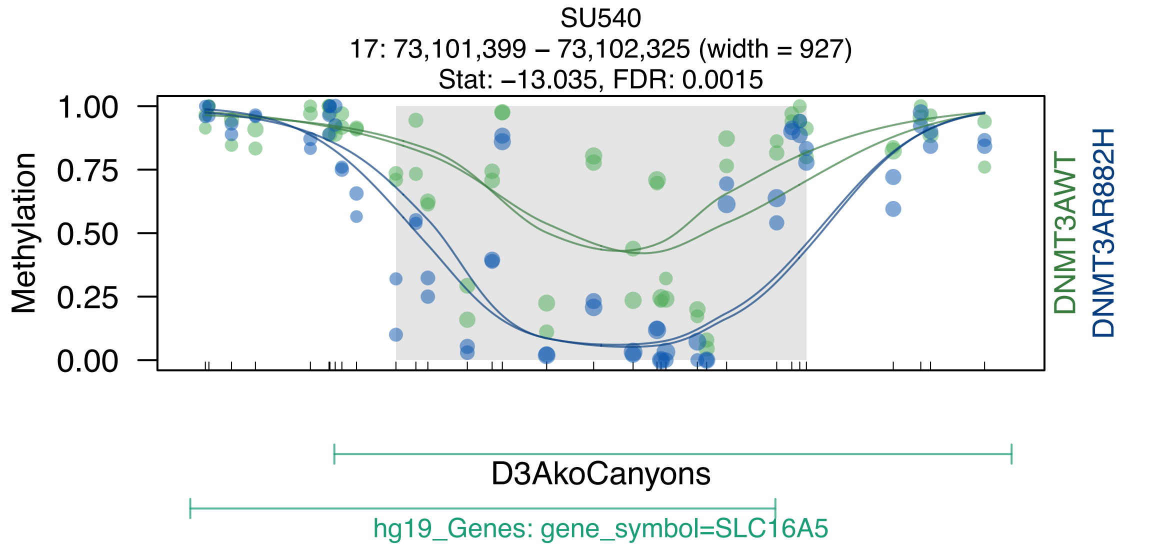
SU540
17: 73,101,399 – 73,102,325 (width = 927)
Stat: −13.035, FDR: 0.0015
Methylation
1.00
0.75
0.50
0.25
0.00
DNMT3AWT
DNMT3AR882H
D3AkoCanyons
hg19_Genes: gene_symbol=SLC16A5

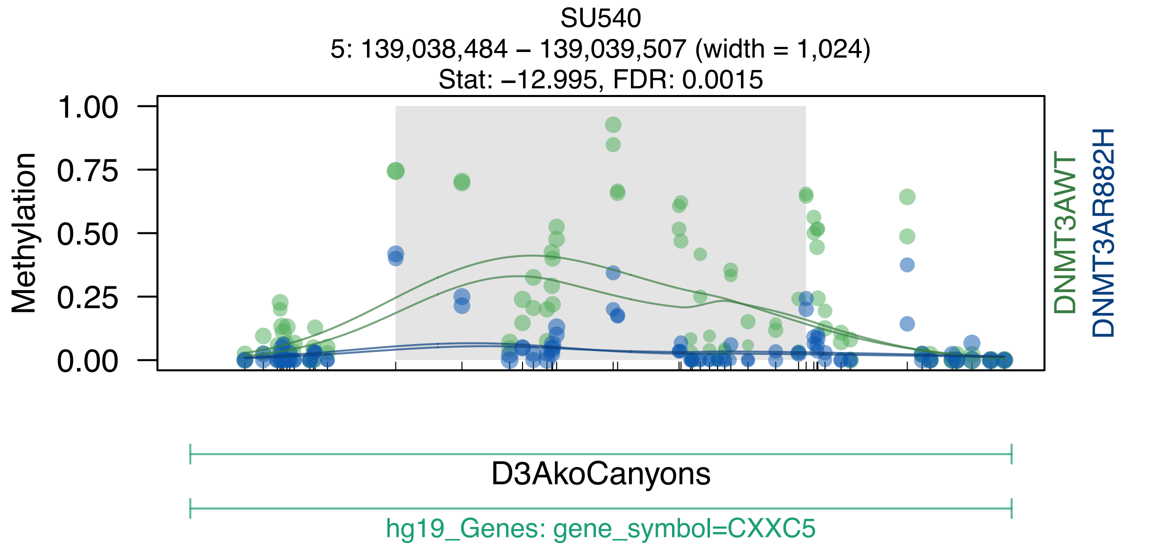
SU540
5: 139,038,484 – 139,039,507 (width = 1,024)
Stat: –12.995, FDR: 0.0015
Methylation
1.00
0.75
0.50
0.25
0.00
DNMT3AWT
DNMT3AR882H
D3AkoCanyons
hg19_Genes: gene_symbol=CXXC5

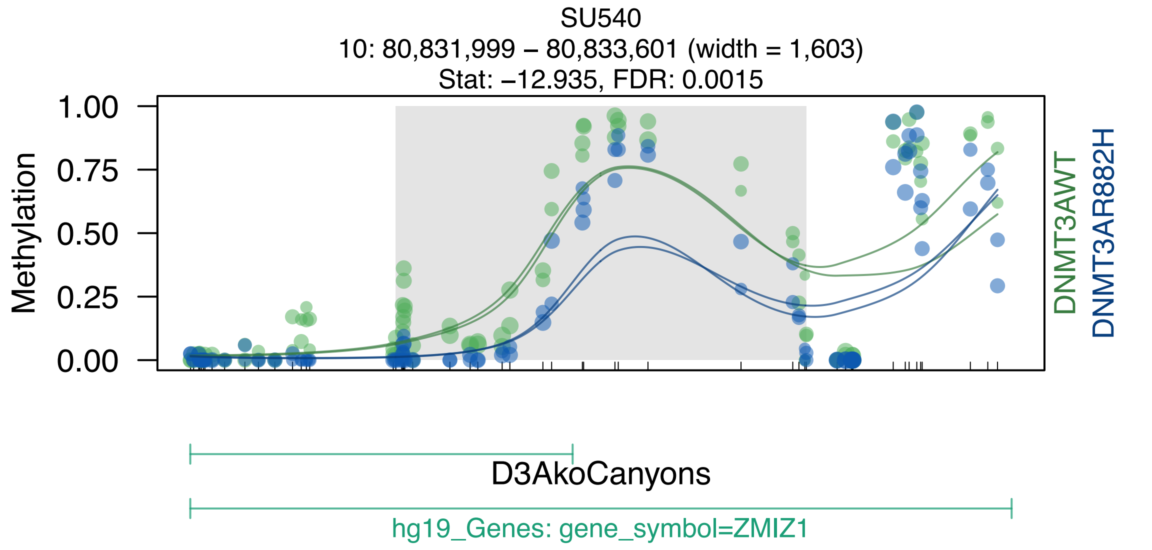
SU540
10: 80,831,999 – 80,833,601 (width = 1,603)
Stat: –12.935, FDR: 0.0015
Methylation
1.00
0.75
0.50
0.25
0.00
DNMT3AWT
DNMT3AR882H
D3AkoCanyons
hg19_Genes: gene_symbol=ZMIZ1

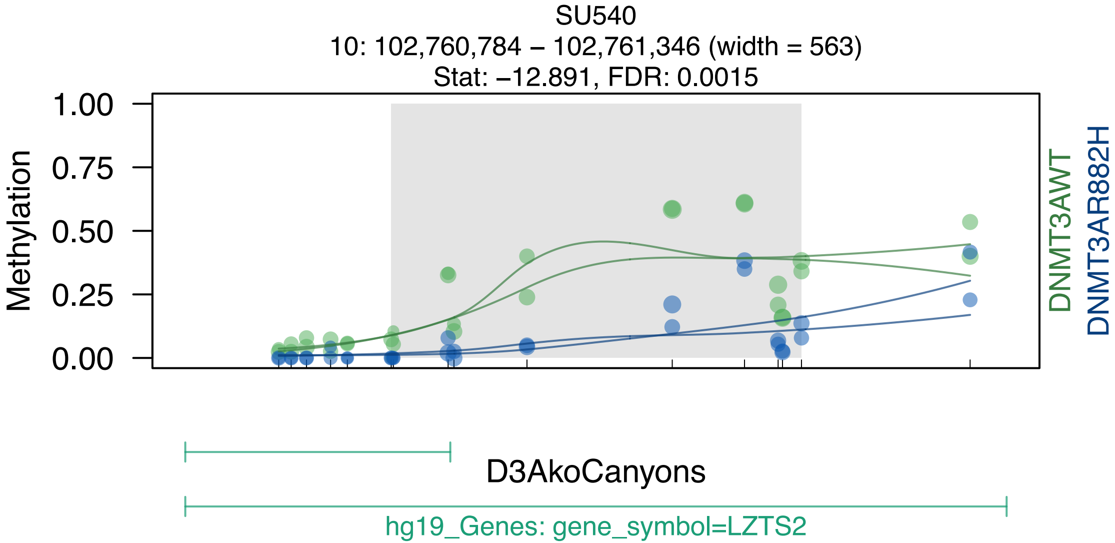
SU540
10: 102,760,784 – 102,761,346 (width = 563)
Stat: –12.891, FDR: 0.0015
Methylation
1.00
0.75
0.50
0.25
0.00
DNMT3AWT
DNMT3AR882H
D3AkoCanyons
hg19_Genes: gene_symbol=LZTS2

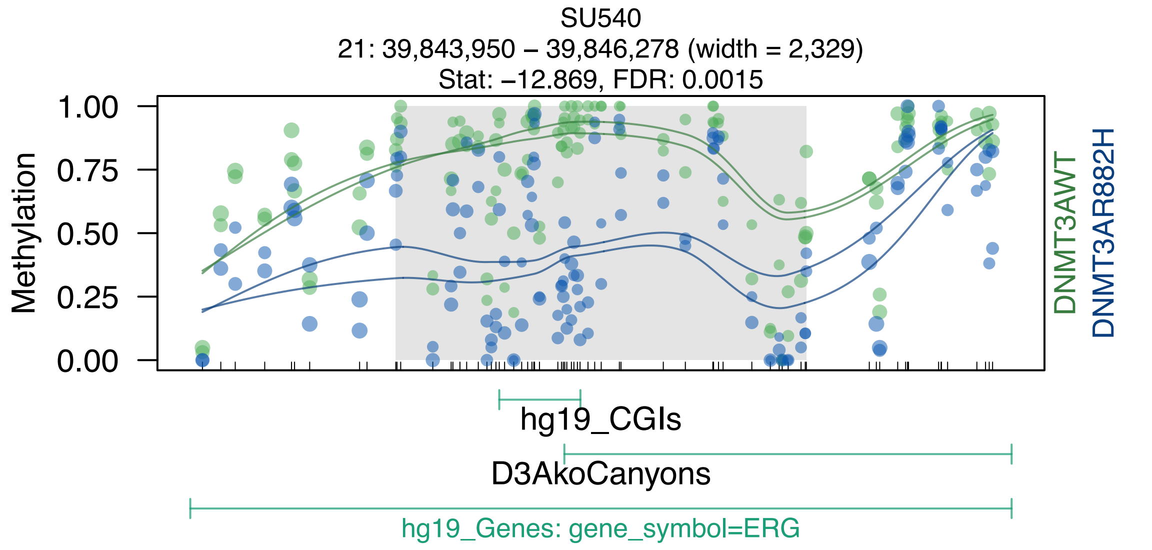
SU540
21: 39,843,950 – 39,846,278 (width = 2,329)
Stat: –12.869, FDR: 0.0015
Methylation
1.00
0.75
0.50
0.25
0.00
DNMT3AWT
DNMT3AR882H
hg19_CGIs
D3AkoCanyons
hg19_Genes: gene_symbol=ERG

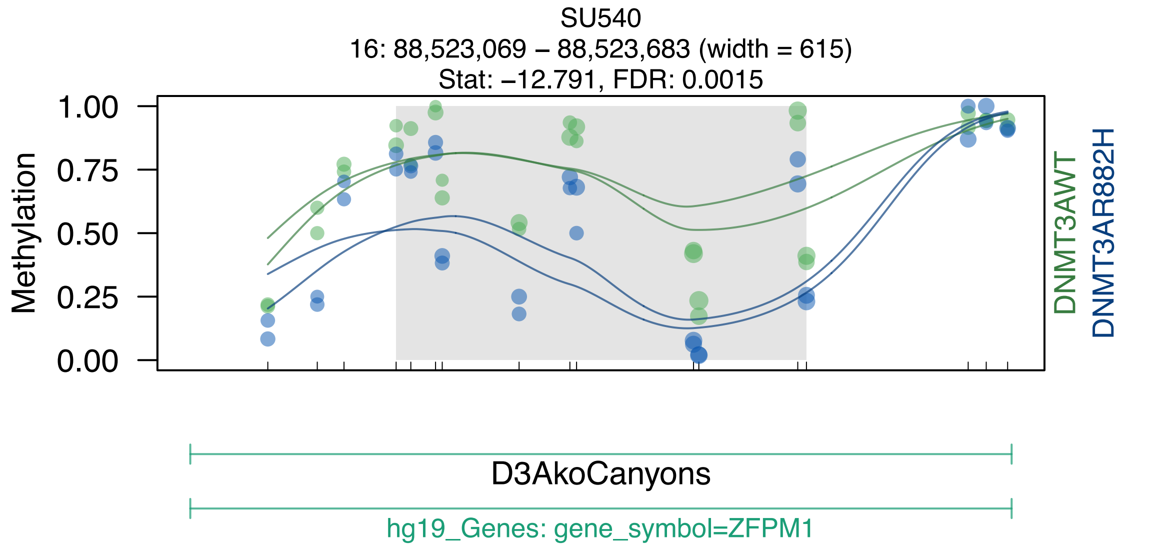
SU540
16: 88,523,069 – 88,523,683 (width = 615)
Stat: –12.791, FDR: 0.0015
Methylation
1.00
0.75
0.50
0.25
0.00
DNMT3AWT
DNMT3AR882H
D3AkoCanyons
hg19_Genes: gene_symbol=ZFPM1

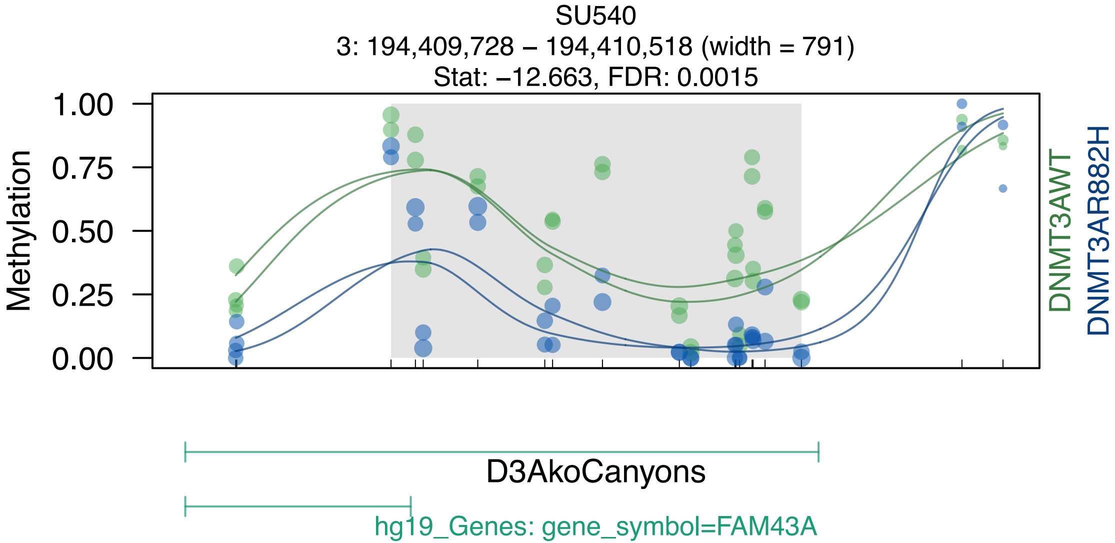
SU540
3: 194,409,728 – 194,410,518 (width = 791)
Stat: –12.663, FDR: 0.0015
Methylation
1.00
0.75
0.50
0.25
0.00
DNMT3AWT
DNMT3AR882H
D3AkoCanyons
hg19_Genes: gene_symbol=FAM43A

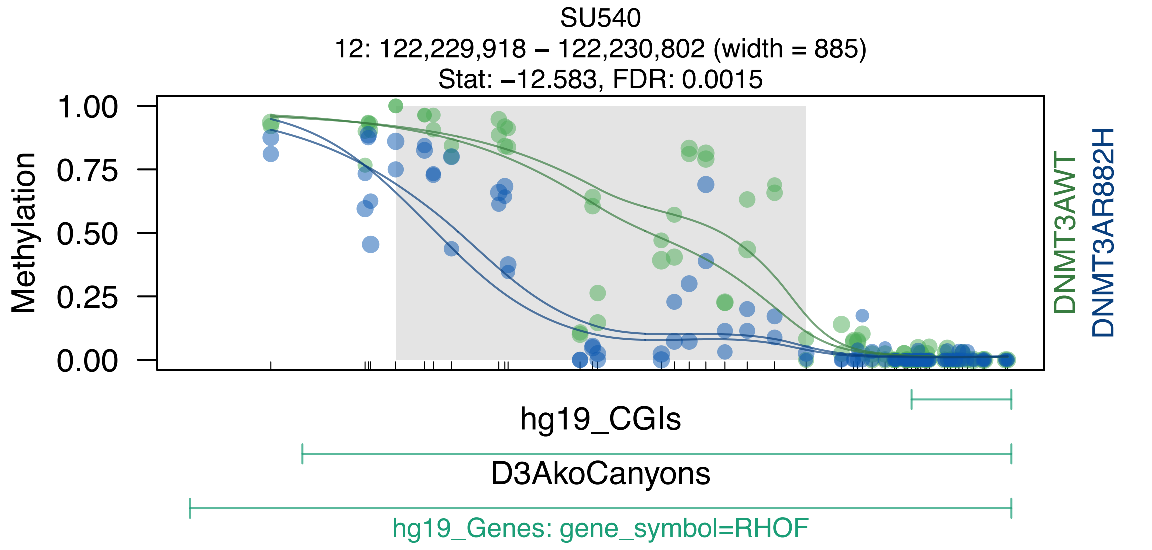

SU540
12: 122,229,918 – 122,230,802 (width = 885)
Stat: –12.583, FDR: 0.0015
Methylation
1.00
0.75
0.50
0.25
0.00
DNMT3AWT
DNMT3AR882H
hg19_CGIs
D3AkoCanyons
hg19_Genes: gene_symbol=RHOF

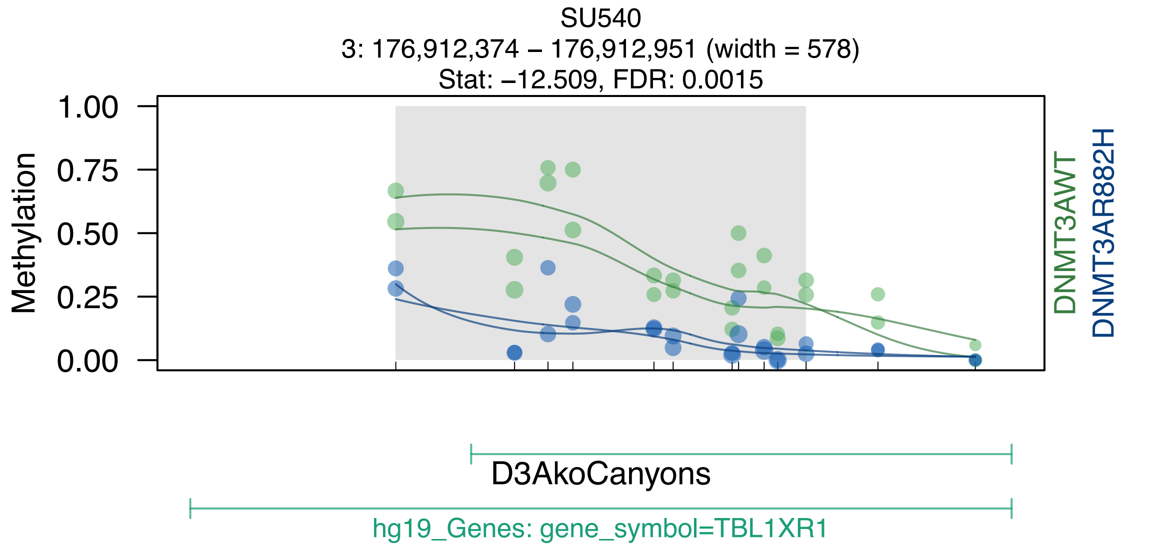
SU540
3: 176,912,374 – 176,912,951 (width = 578)
Stat: –12.509, FDR: 0.0015
Methylation
1.00
0.75
0.50
0.25
0.00
DNMT3AWT
DNMT3AR882H
D3AkoCanyons
hg19_Genes: gene_symbol=TBL1XR1

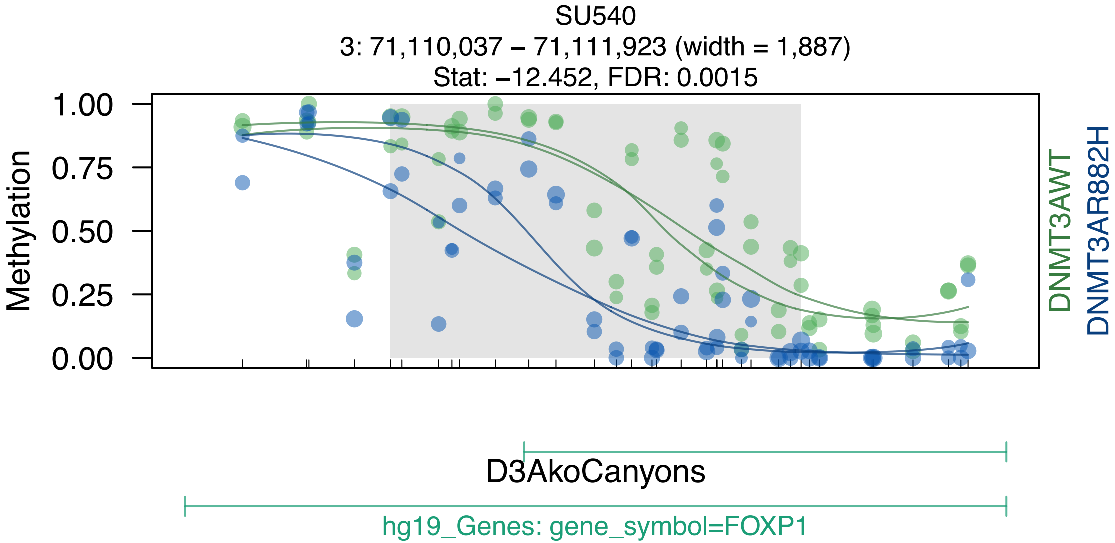
SU540
3: 71,110,037 – 71,111,923 (width = 1,887)
Stat: –12.452, FDR: 0.0015
Methylation
1.00
0.75
0.50
0.25
0.00
DNMT3AWT
DNMT3AR882H
D3AkoCanyons
hg19_Genes: gene_symbol=FOXP1

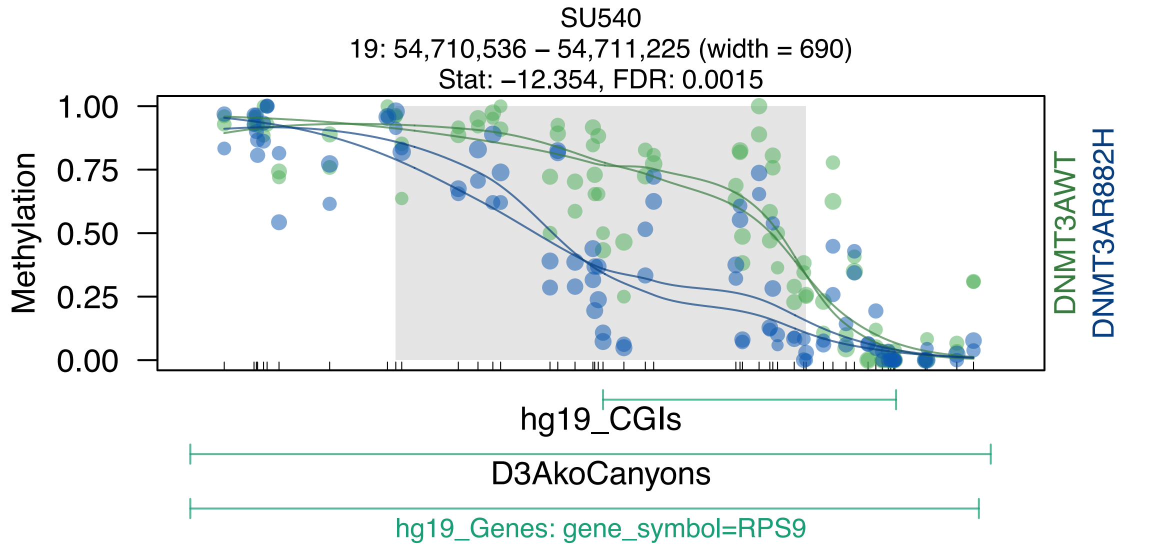
SU540
19: 54,710,536 – 54,711,225 (width = 690)
Stat: –12.354, FDR: 0.0015
Methylation
1.00
0.75
0.50
0.25
0.00
DNMT3AWT
DNMT3AR882H
hg19_CGIs
D3AkoCanyons
hg19_Genes: gene_symbol=RPS9
