## Supplemental Data S1-S7 for "DNMT3A^R882H^ Is Not Required for Disease Maintenance in Primary Human AML, but Is Associated With Increased Leukemia Stem Cell Frequency": DataS6_SU575_top50dmrsOverlappingCanyons.pdf

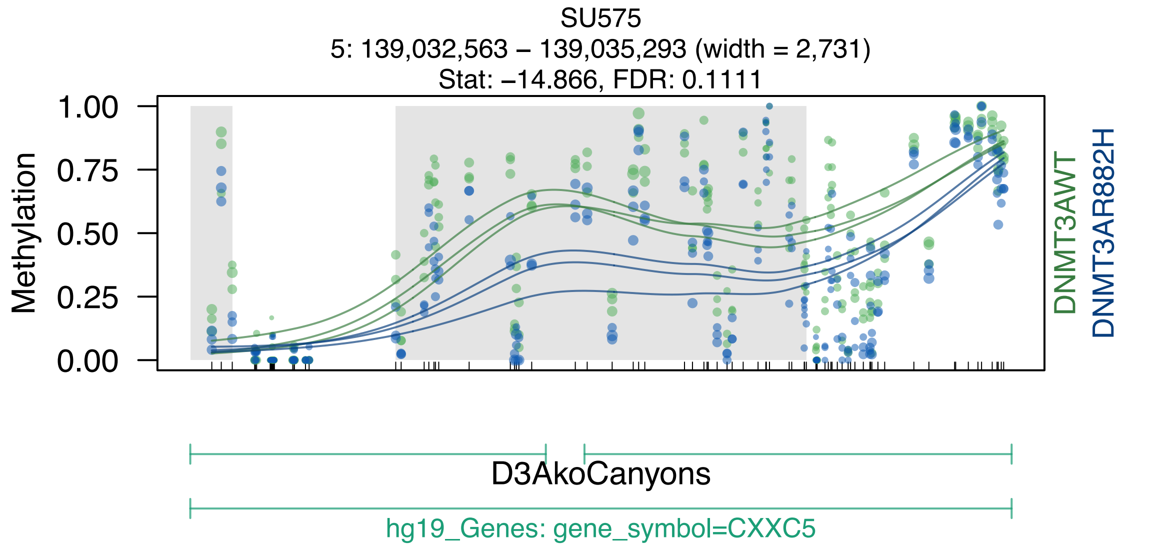
SU575
5: 139,032,563 – 139,035,293 (width = 2,731)
Stat: –14.866, FDR: 0.1111
Methylation
1.00
0.75
0.50
0.25
0.00
DNMT3AWT
DNMT3AR882H
D3AkoCanyons
hg19_Genes: gene_symbol=CXXC5

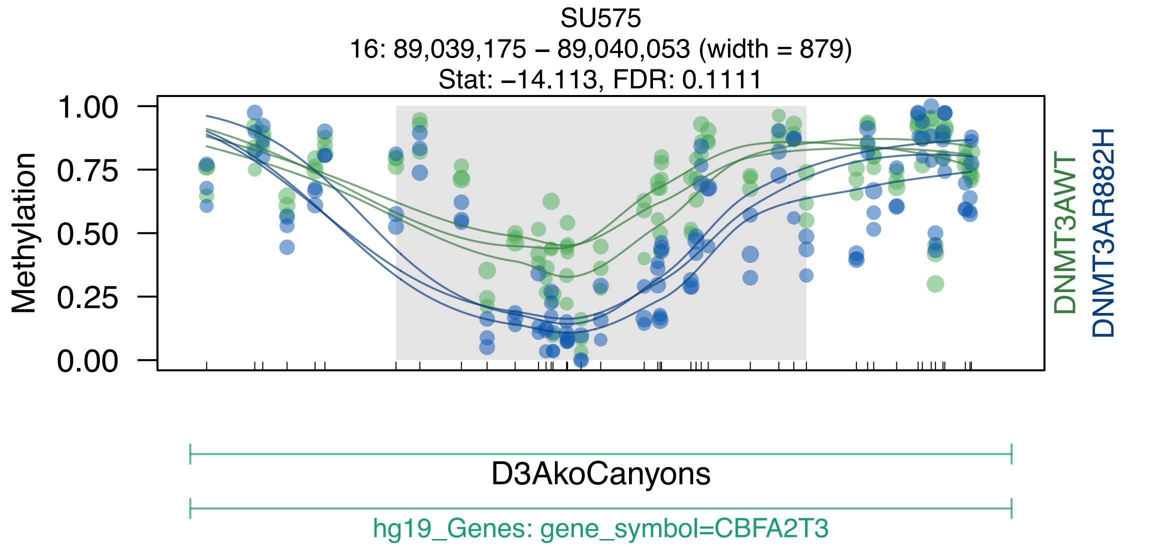
SU575
16: 89,039,175 – 89,040,053 (width = 879)
Stat: –14.113, FDR: 0.1111
Methylation
1.00
0.75
0.50
0.25
0.00
DNMT3AWT
DNMT3AR882H
D3AkoCanyons
hg19_Genes: gene_symbol=CBFA2T3

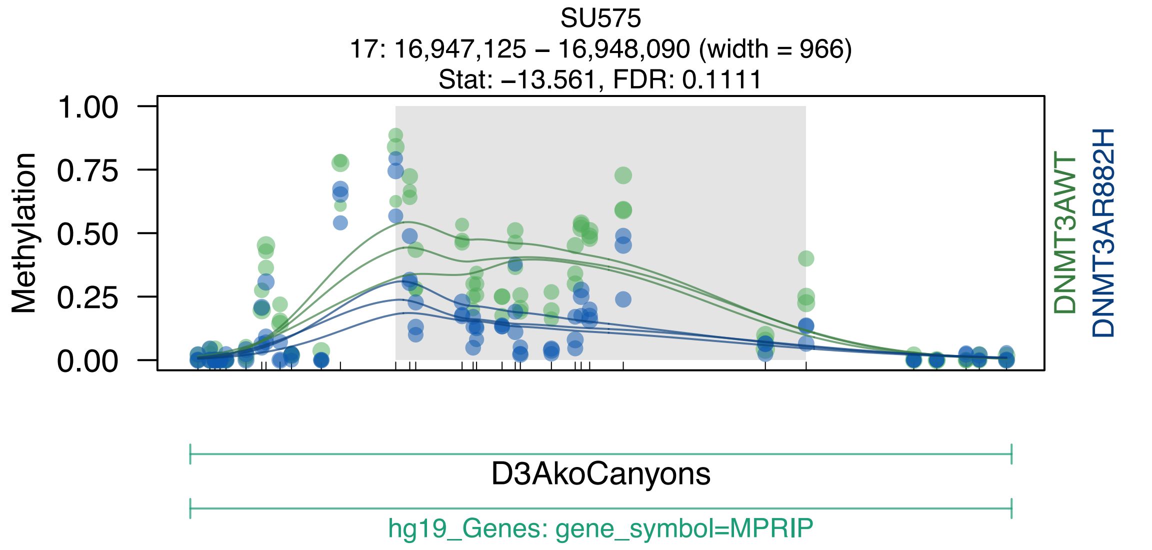
SU575
17: 16,947,125 – 16,948,090 (width = 966)
Stat: –13.561, FDR: 0.1111
Methylation
1.00
0.75
0.50
0.25
0.00
DNMT3AWT
DNMT3AR882H
D3AkoCanyons
hg19_Genes: gene_symbol=MPRIP

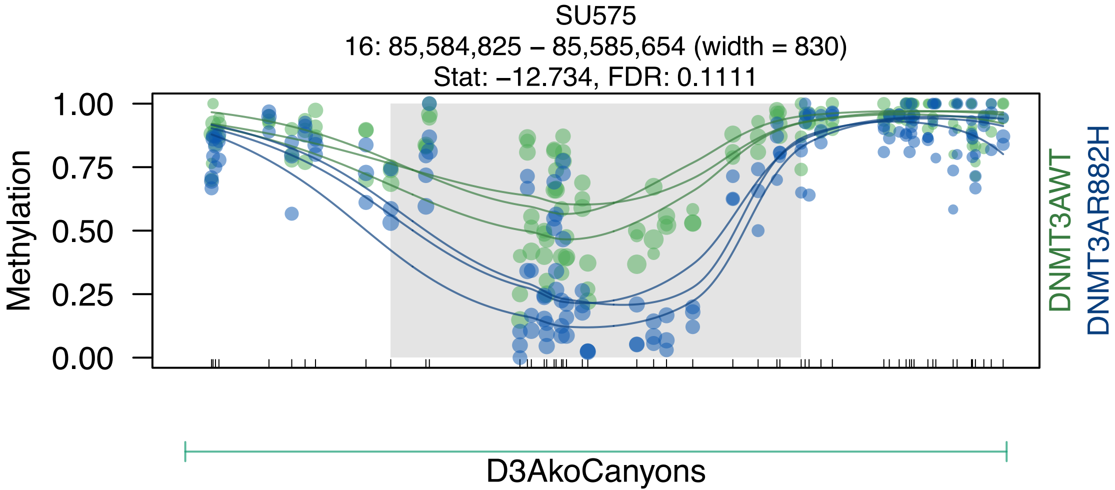
SU575
16: 85,584,825 – 85,585,654 (width = 830)
Stat: –12.734, FDR: 0.1111
Methylation
1.00
0.75
0.50
0.25
0.00
DNMT3AWT
DNMT3AR882H
D3AkoCanyons

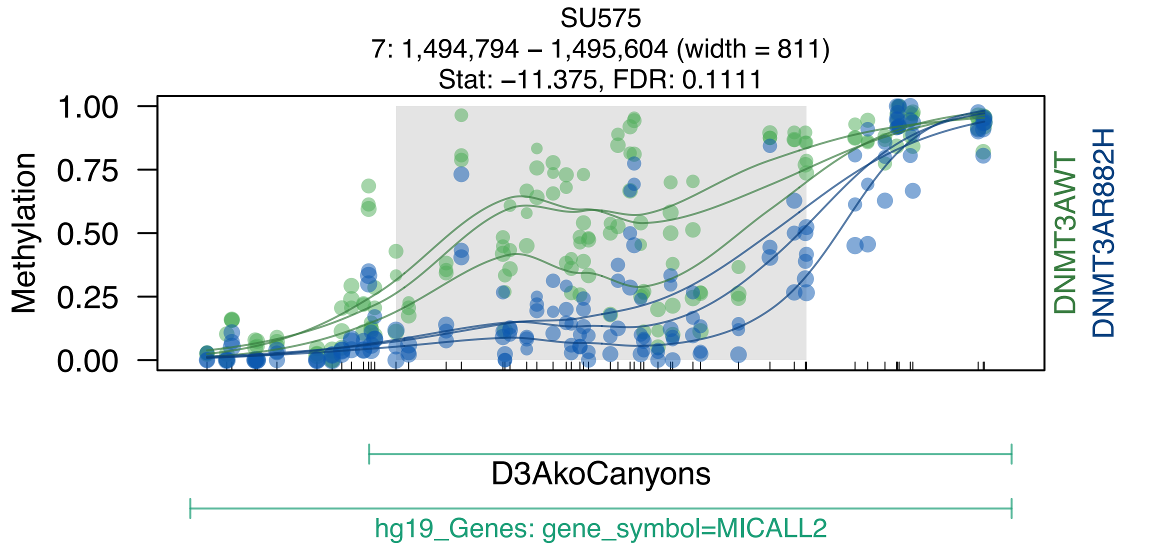
SU575
7: 1,494,794 – 1,495,604 (width = 811)
Stat: –11.375, FDR: 0.1111
Methylation
1.00
0.75
0.50
0.25
0.00
DNMT3AWT
DNMT3AR882H
D3AkoCanyons
hg19_Genes: gene_symbol=MICALL2

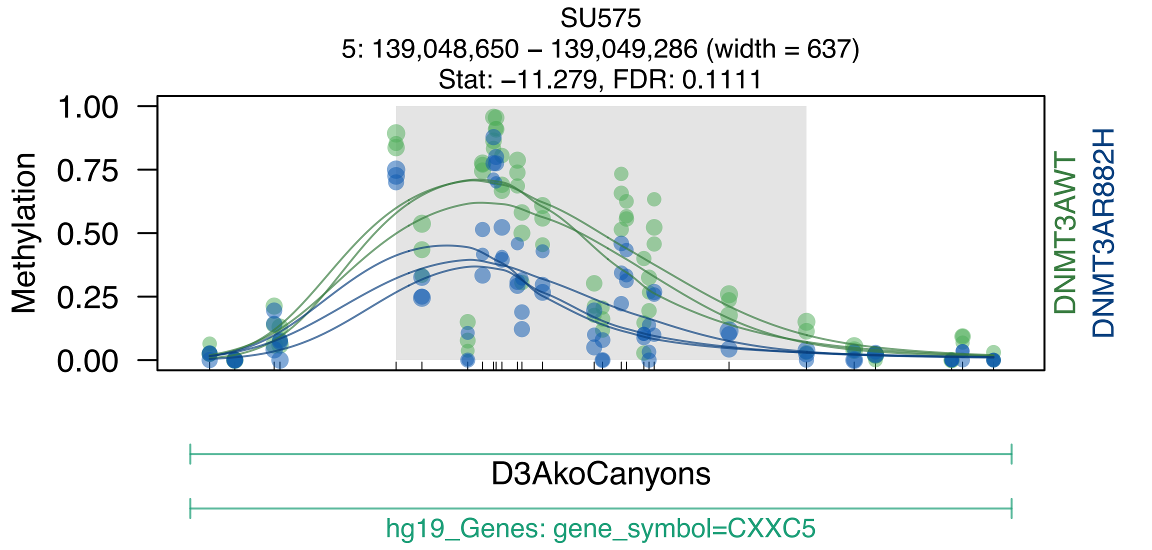
SU575
5: 139,048,650 – 139,049,286 (width = 637)
Stat: –11.279, FDR: 0.1111
Methylation
1.00
0.75
0.50
0.25
0.00
DNMT3AWT
DNMT3AR882H
D3AkoCanyons
hg19_Genes: gene_symbol=CXXC5

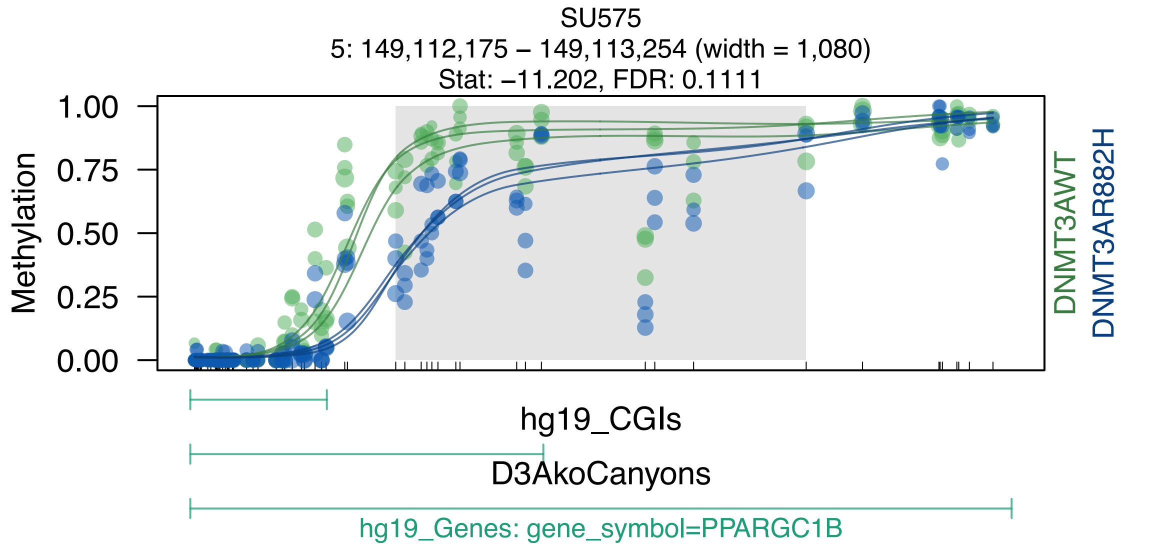
SU575
5: 149,112,175 – 149,113,254 (width = 1,080)
Stat: –11.202, FDR: 0.1111
Methylation
1.00
0.75
0.50
0.25
0.00
DNMT3AWT
DNMT3AR882H
hg19_CGIs
D3AkoCanyons
hg19_Genes: gene_symbol=PPARGC1B

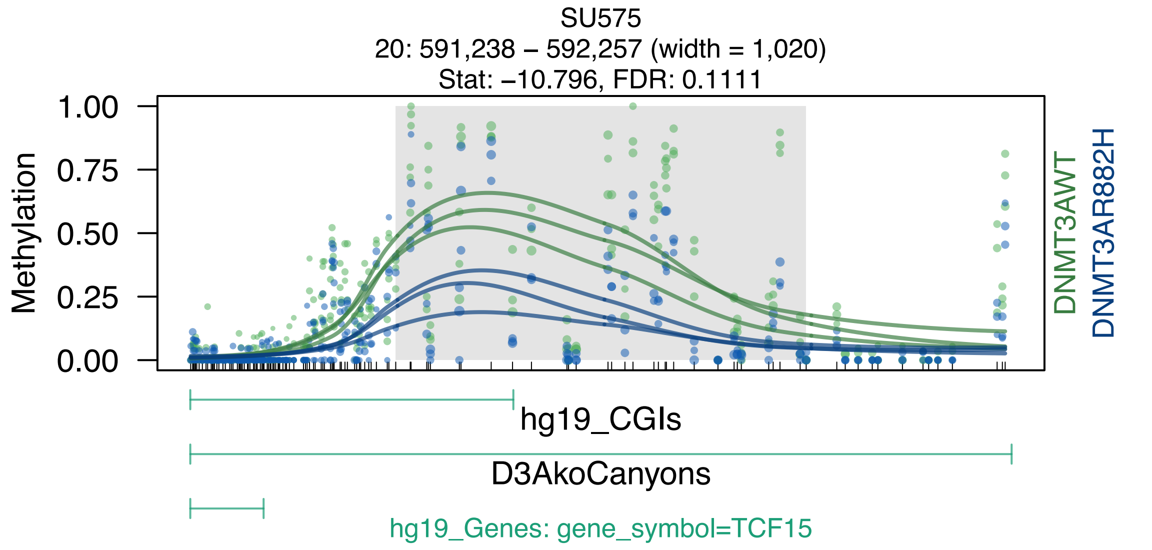
SU575
20: 591,238 – 592,257 (width = 1,020)
Stat: −10.796, FDR: 0.1111
Methylation
1.00
0.75
0.50
0.25
0.00
DNMT3AWT
DNMT3AR882H
hg19_CGIs
D3AkoCanyons
hg19_Genes: gene_symbol=TCF15

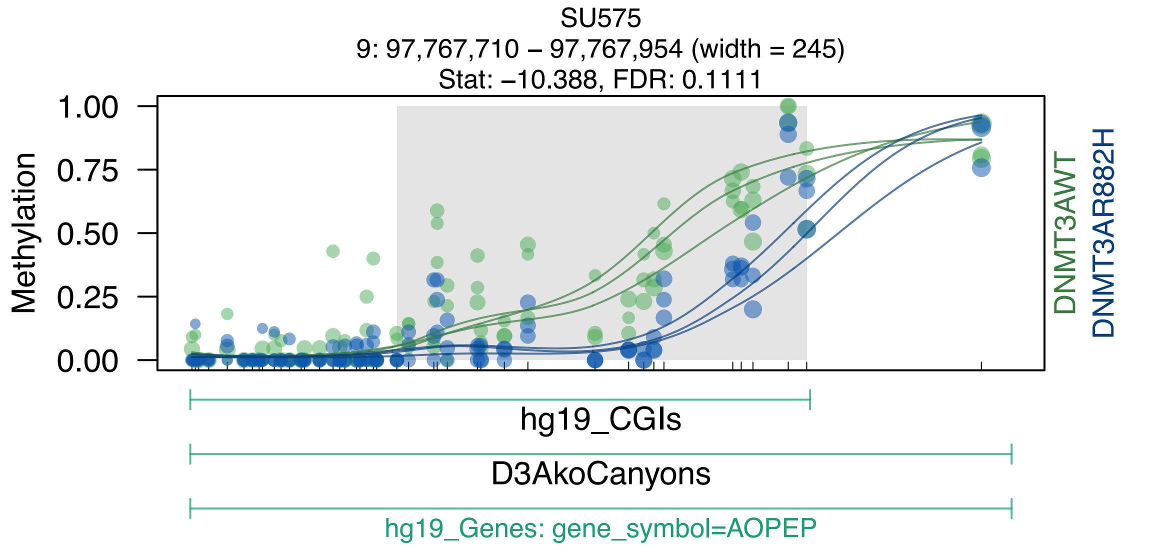
SU575
9: 97,767,710 – 97,767,954 (width = 245)
Stat: −10.388, FDR: 0.1111
Methylation
1.00
0.75
0.50
0.25
0.00
DNMT3AWT
DNMT3AR882H
hg19_CGIs
D3AkoCanyons
hg19_Genes: gene_symbol=AOPEP

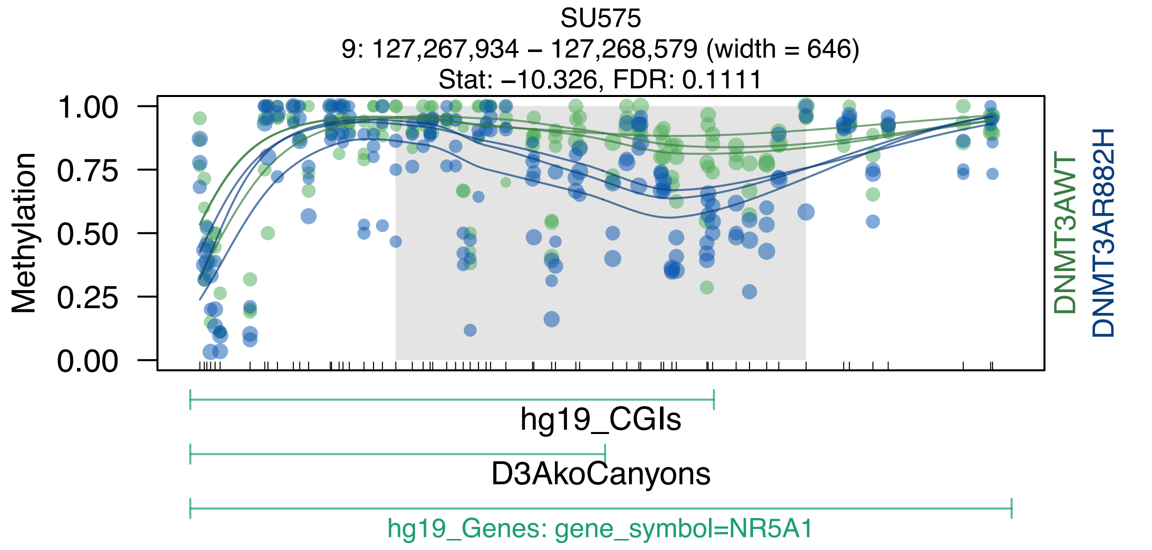
SU575
9: 127,267,934 – 127,268,579 (width = 646)
Stat: –10.326, FDR: 0.1111
Methylation
1.00
0.75
0.50
0.25
0.00
DNMT3AWT
DNMT3AR882H
hg19_CGIs
D3AkoCanyons
hg19_Genes: gene_symbol=NR5A1

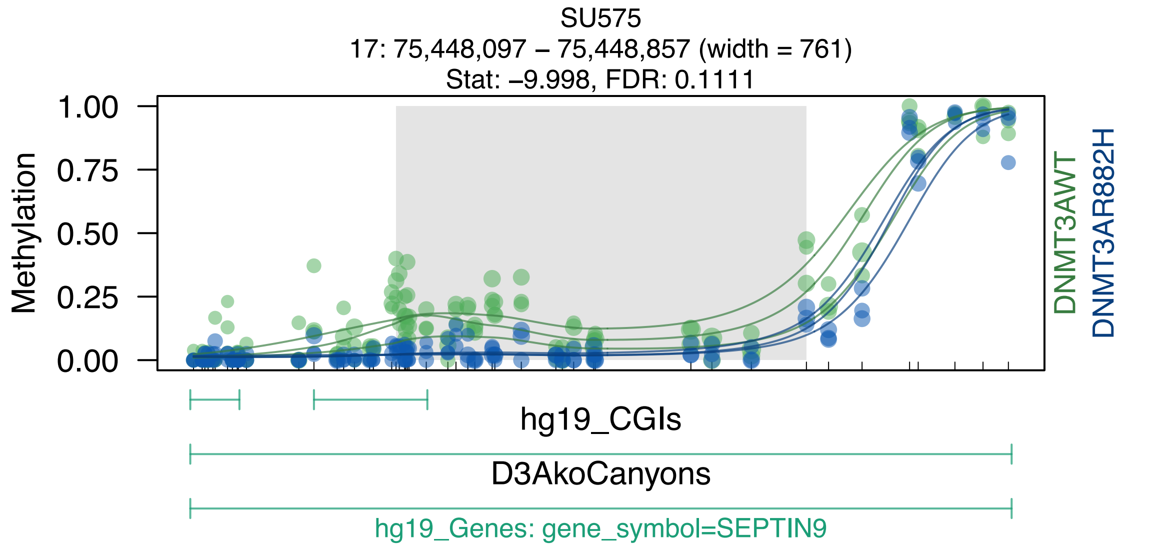
SU575
17: 75,448,097 – 75,448,857 (width = 761)
Stat: –9.998, FDR: 0.1111
Methylation
1.00
0.75
0.50
0.25
0.00
DNMT3AWT
DNMT3AR882H
hg19_CGIs
D3AkoCanyons
hg19_Genes: gene_symbol=SEPTIN9

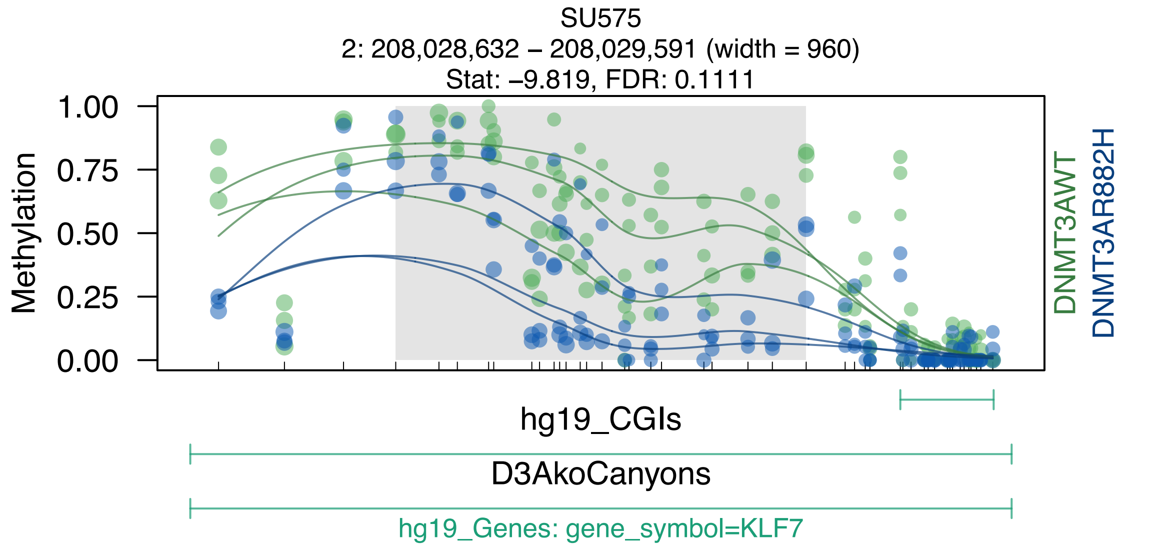

SU575
2: 208,028,632 – 208,029,591 (width = 960)
Stat: –9.819, FDR: 0.1111
Methylation
1.00
0.75
0.50
0.25
0.00
DNMT3AWT
DNMT3AR882H
hg19_CGIs
D3AkoCanyons
hg19_Genes: gene_symbol=KLF7

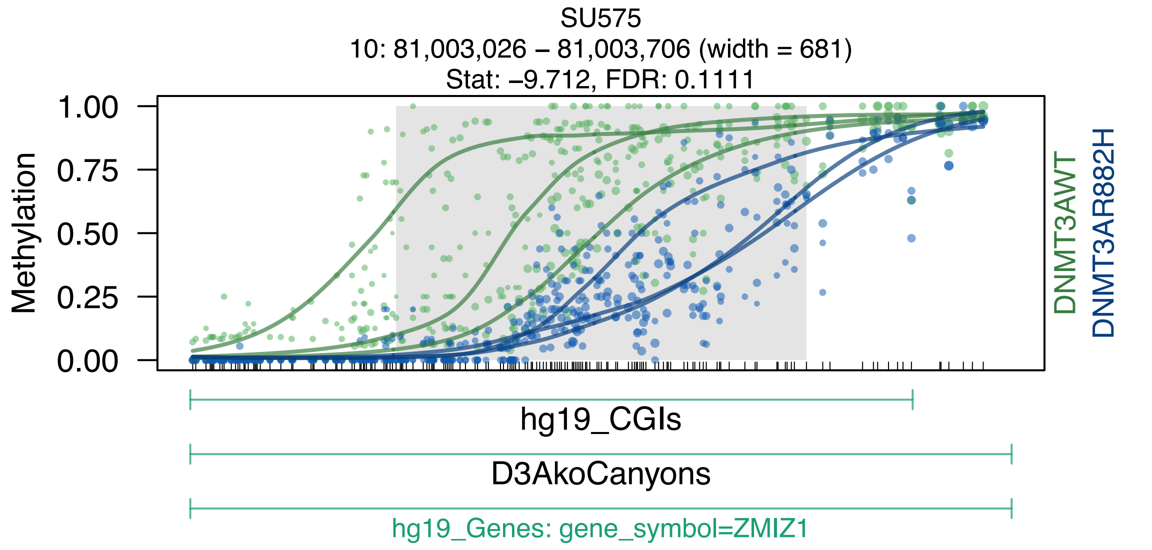
SU575
10: 81,003,026 – 81,003,706 (width = 681)
Stat: –9.712, FDR: 0.1111
Methylation
1.00
0.75
0.50
0.25
0.00
DNMT3AWT
DNMT3AR882H
hg19_CGIs
D3AkoCanyons
hg19_Genes: gene_symbol=ZMIZ1

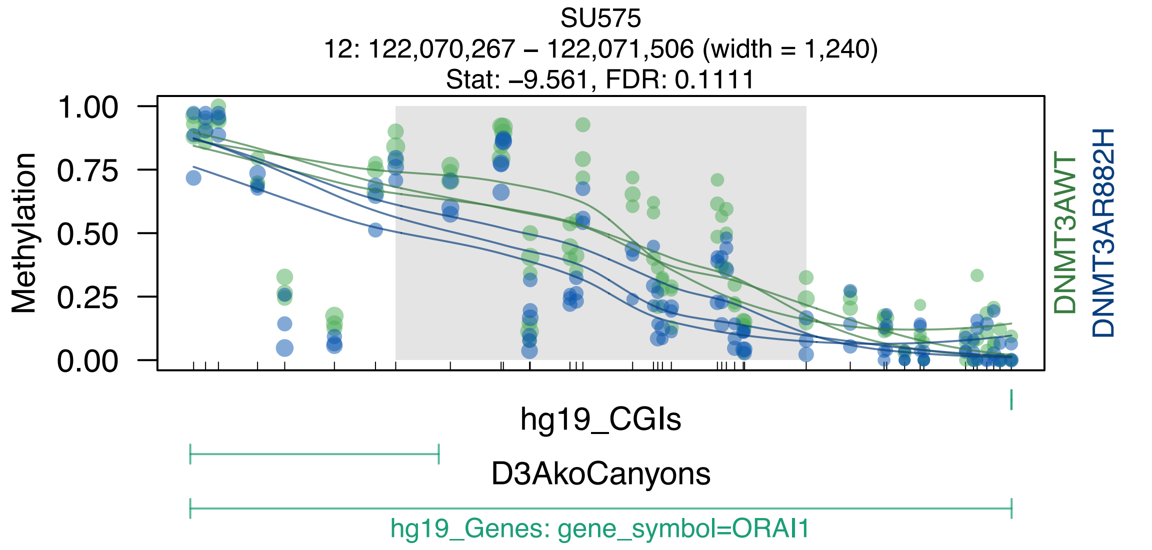

SU575
12: 122,070,267 – 122,071,506 (width = 1,240)
Stat: –9.561, FDR: 0.1111
Methylation
1.00
0.75
0.50
0.25
0.00
DNMT3AWT
DNMT3AR882H
hg19_CGIs
D3AkoCanyons
hg19_Genes: gene_symbol=ORAI1

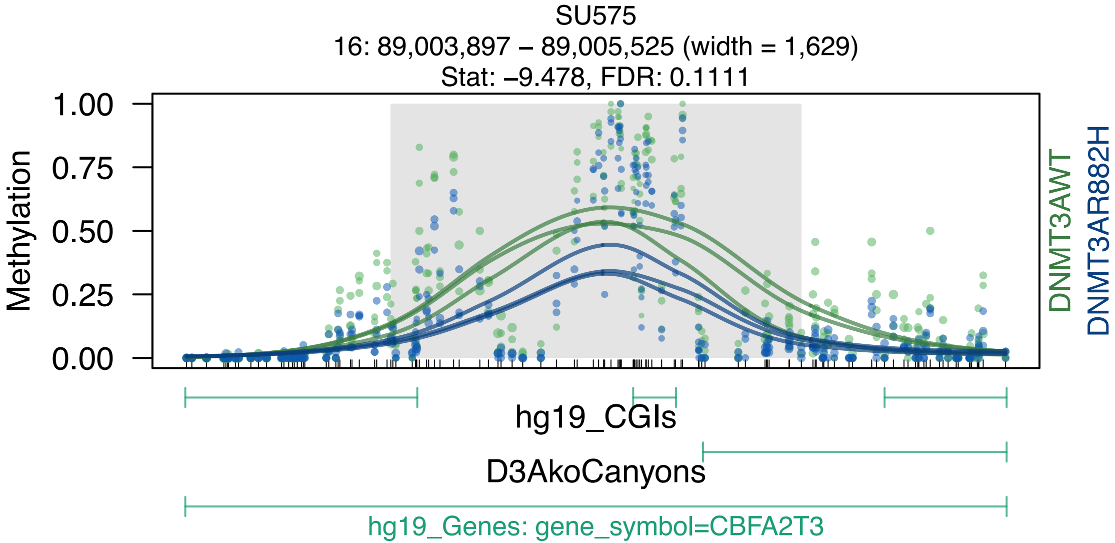
SU575
16: 89,003,897 – 89,005,525 (width = 1,629)
Stat: –9.478, FDR: 0.1111
Methylation
1.00
0.75
0.50
0.25
0.00
DNMT3AWT
DNMT3AR882H
hg19_CGIs
D3AkoCanyons
hg19_Genes: gene_symbol=CBFA2T3

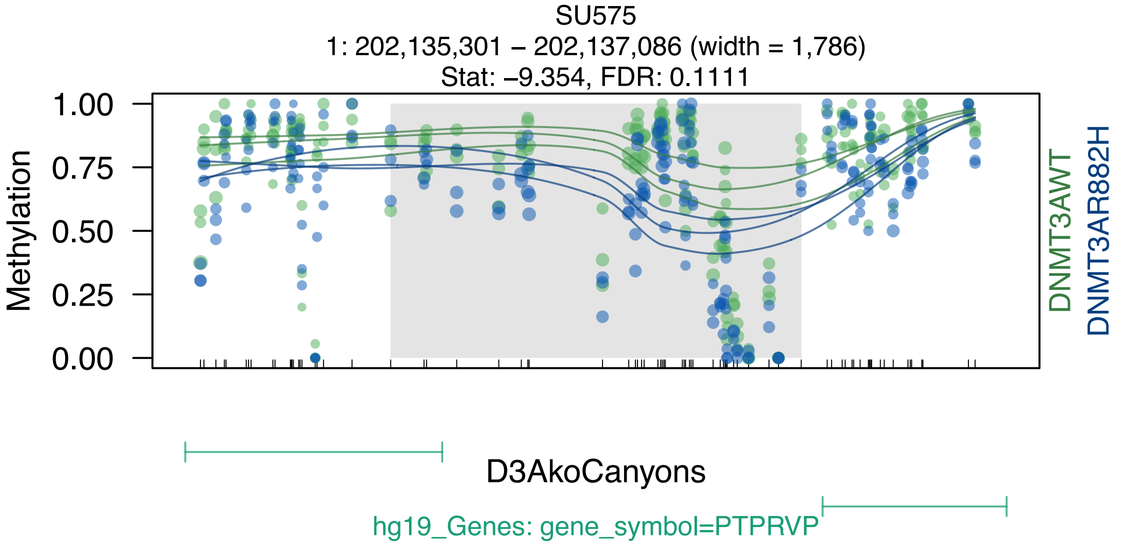
SU575
1: 202,135,301 – 202,137,086 (width = 1,786)
Stat: –9.354, FDR: 0.1111
Methylation
1.00
0.75
0.50
0.25
0.00
DNMT3AWT
DNMT3AR882H
D3AkoCanyons
hg19_Genes: gene_symbol=PTPRVP

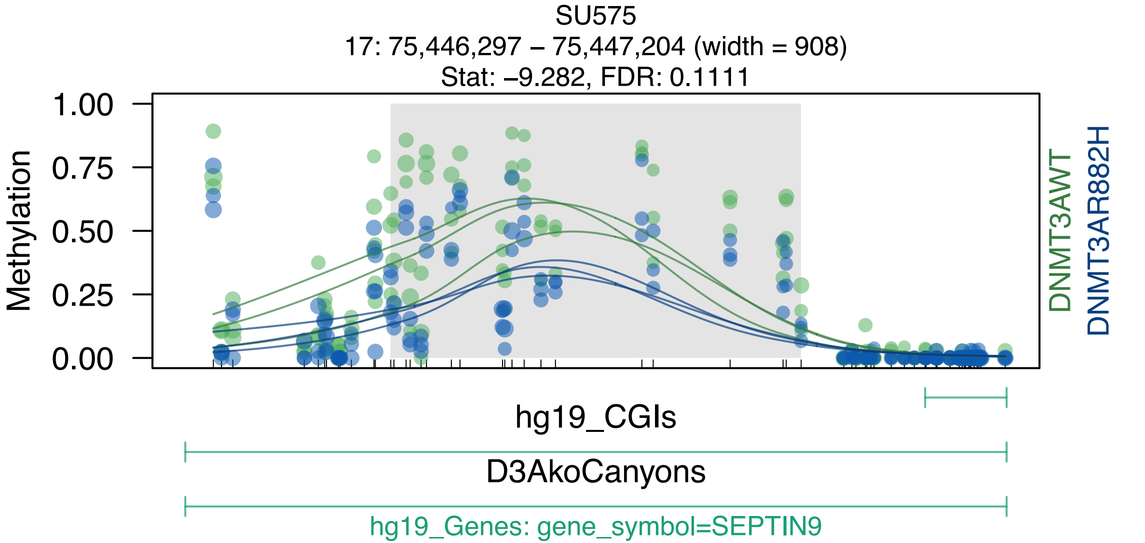

SU575
17: 75,446,297 – 75,447,204 (width = 908)
Stat: –9.282, FDR: 0.1111
Methylation
1.00
0.75
0.50
0.25
0.00
DNMT3AWT
DNMT3AR882H
hg19_CGIs
D3AkoCanyons
hg19_Genes: gene_symbol=SEPTIN9

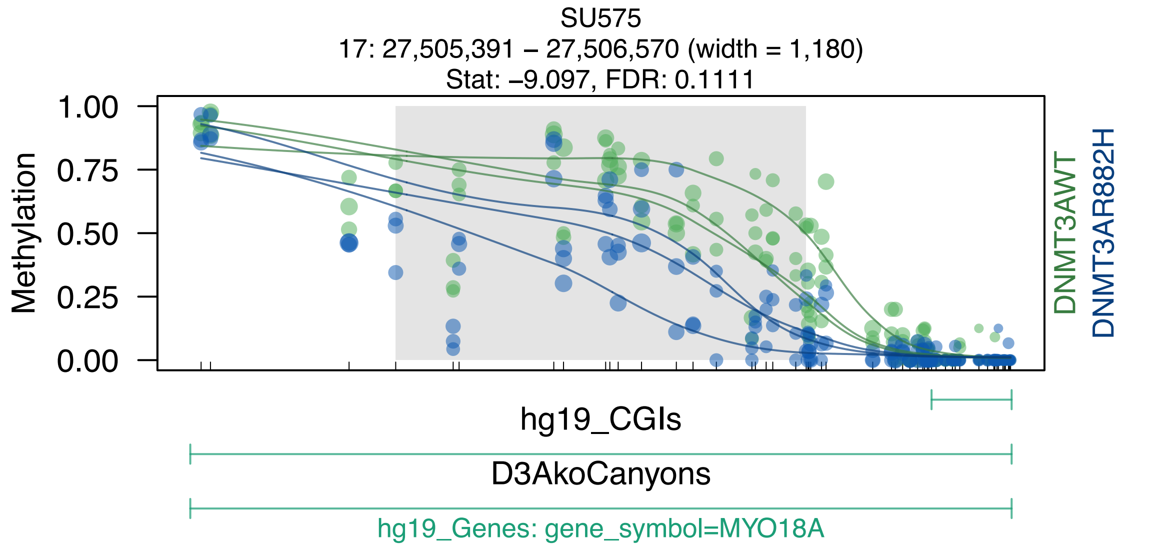

SU575
17: 27,505,391 – 27,506,570 (width = 1,180)
Stat: –9.097, FDR: 0.1111
Methylation
1.00
0.75
0.50
0.25
0.00
DNMT3AWT
DNMT3AR882H
hg19_CGIs
D3AkoCanyons
hg19_Genes: gene_symbol=MYO18A

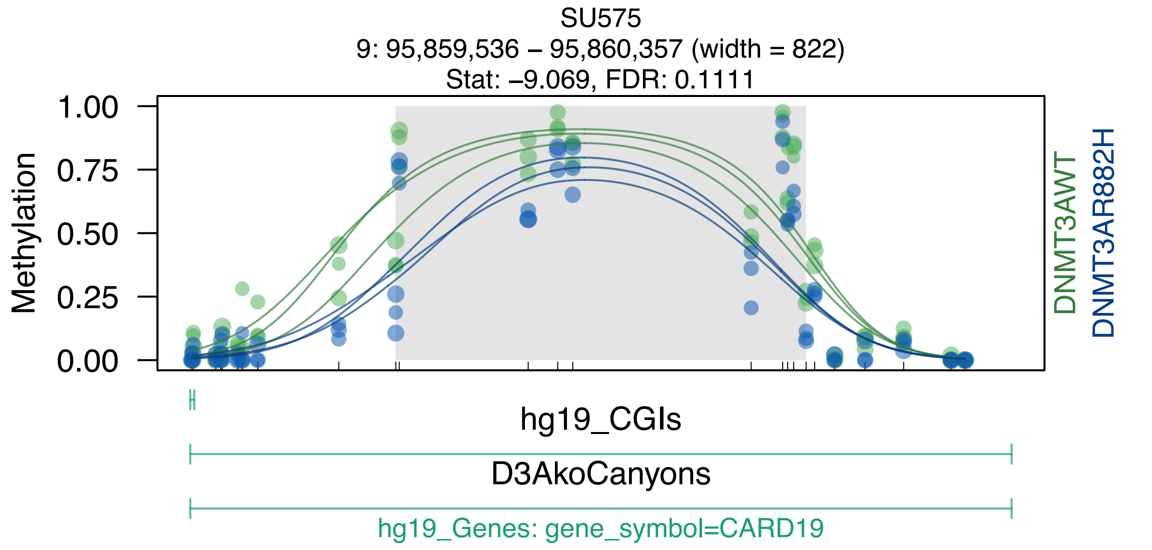

SU575
9: 95,859,536 – 95,860,357 (width = 822)
Stat: –9.069, FDR: 0.1111
Methylation
1.00
0.75
0.50
0.25
0.00
DNMT3AWT
DNMT3AR882H
hg19_CGIs
D3AkoCanyons
hg19_Genes: gene_symbol=CARD19

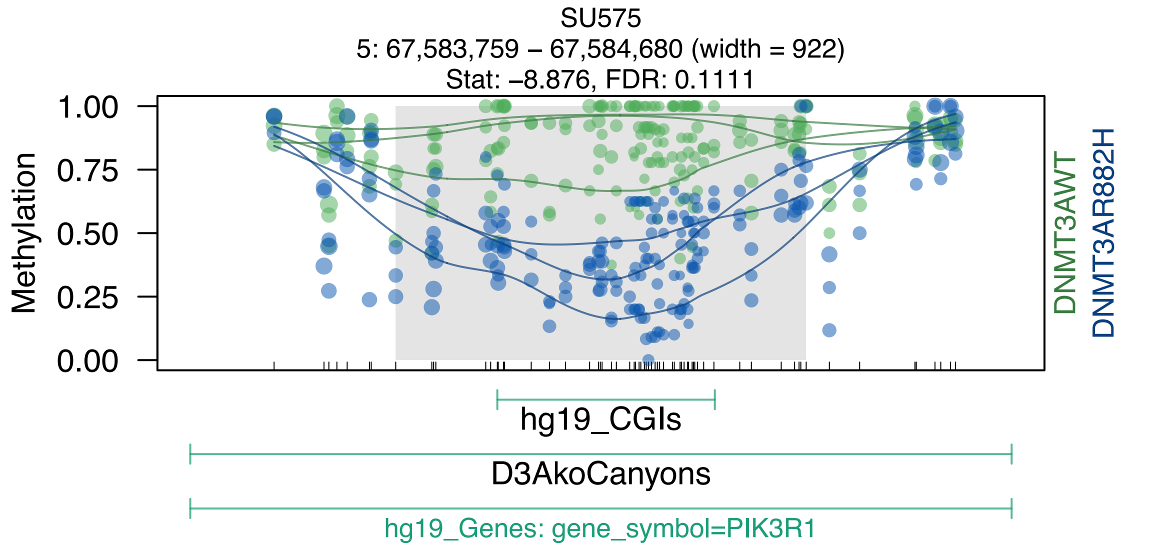

SU575
5: 67,583,759 – 67,584,680 (width = 922)
Stat: –8.876, FDR: 0.1111
1.00
0.75
0.50
0.25
0.00
Methylation
DNMT3AWT
DNMT3AR882H
hg19_CGIs
D3AkoCanyons
hg19_Genes: gene_symbol=PIK3R1

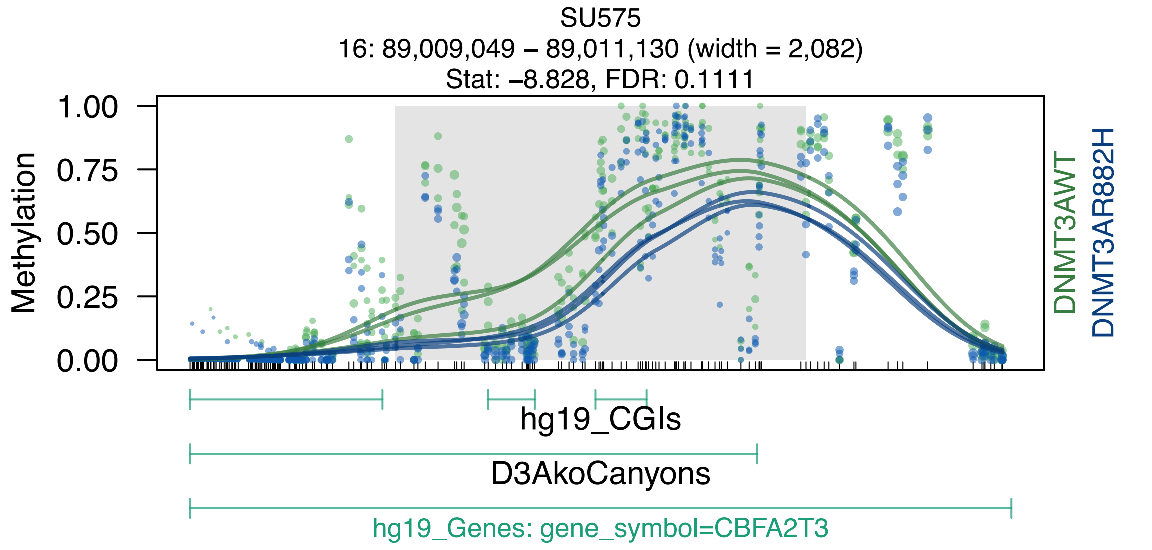

SU575
16: 89,009,049 – 89,011,130 (width = 2,082)
Stat: –8.828, FDR: 0.1111
Methylation
1.00
0.75
0.50
0.25
0.00
DNMT3AWT
DNMT3AR882H
hg19_CGIs
D3AkoCanyons
hg19_Genes: gene_symbol=CBFA2T3

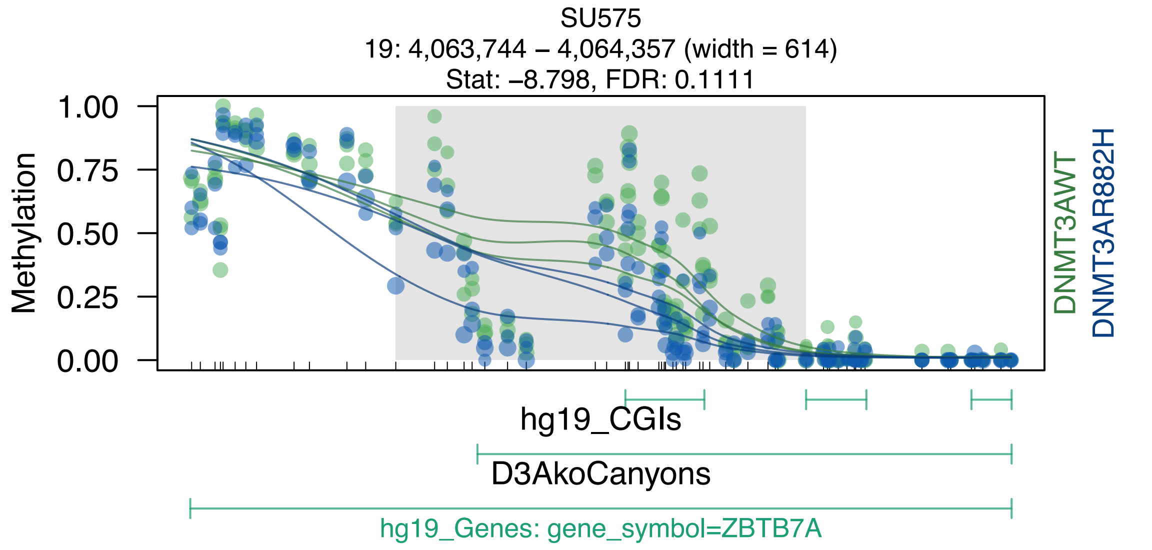
SU575
19: 4,063,744 – 4,064,357 (width = 614)
Stat: –8.798, FDR: 0.1111
Methylation
1.00
0.75
0.50
0.25
0.00
DNMT3AWT
DNMT3AR882H
hg19_CGIs
D3AkoCanyons
hg19_Genes: gene_symbol=ZBTB7A

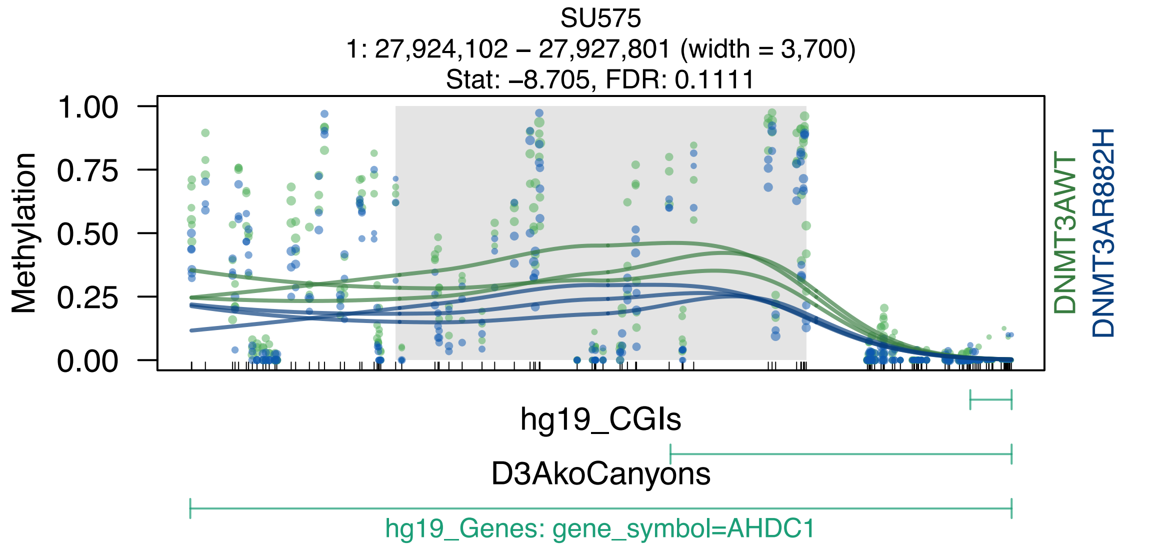

SU575
1: 27,924,102 – 27,927,801 (width = 3,700)
Stat: –8.705, FDR: 0.1111
Methylation
1.00
0.75
0.50
0.25
0.00
DNMT3AWT
DNMT3AR882H
hg19_CGIs
D3AkoCanyons
hg19_Genes: gene_symbol=AHDC1

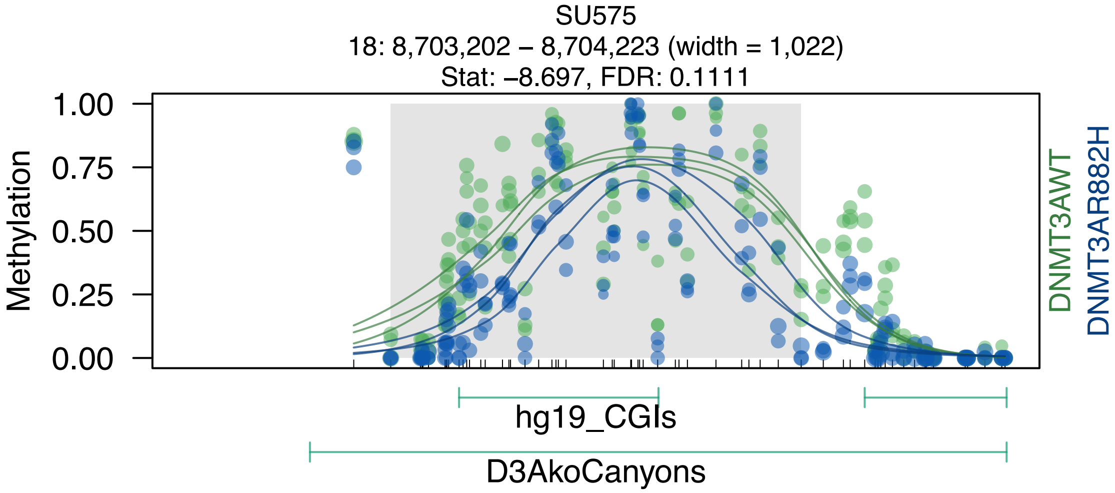

SU575
18: 8,703,202 – 8,704,223 (width = 1,022)
Stat: –8.697, FDR: 0.1111
Methylation
1.00
0.75
0.50
0.25
0.00
DNMT3AWT
DNMT3AR882H
hg19_CGIs
D3AkoCanyons

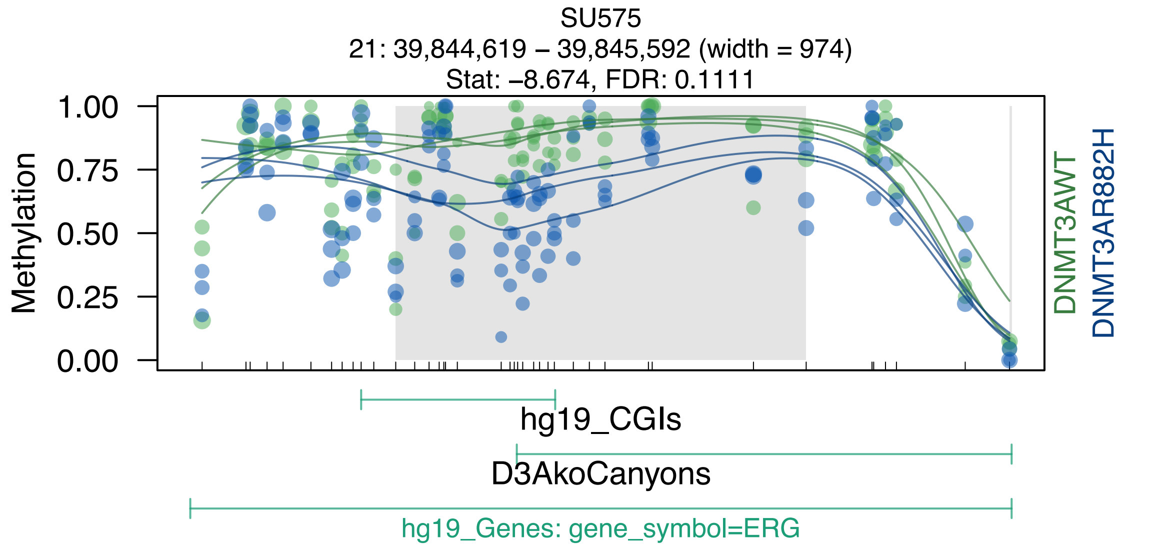
SU575
21: 39,844,619 – 39,845,592 (width = 974)
Stat: –8.674, FDR: 0.1111
Methylation
1.00
0.75
0.50
0.25
0.00
DNMT3AWT
DNMT3AR882H
hg19_CGIs
D3AkoCanyons
hg19_Genes: gene_symbol=ERG

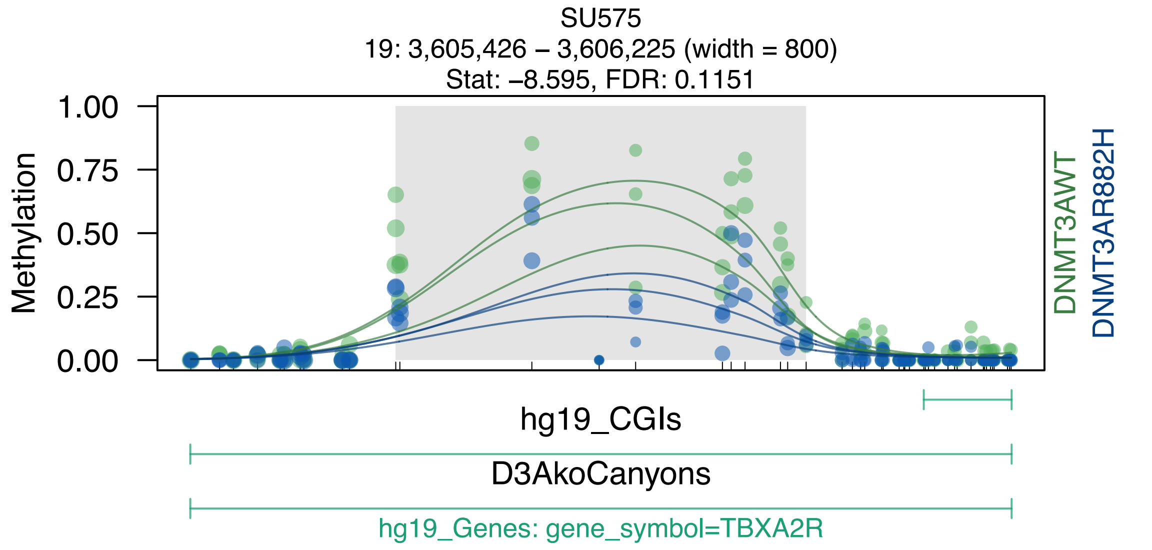

SU575
19: 3,605,426 – 3,606,225 (width = 800)
Stat: –8.595, FDR: 0.1151
Methylation
1.00
0.75
0.50
0.25
0.00
DNMT3AWT
DNMT3AR882H
hg19_CGIs
D3AkoCanyons
hg19_Genes: gene_symbol=TBXA2R

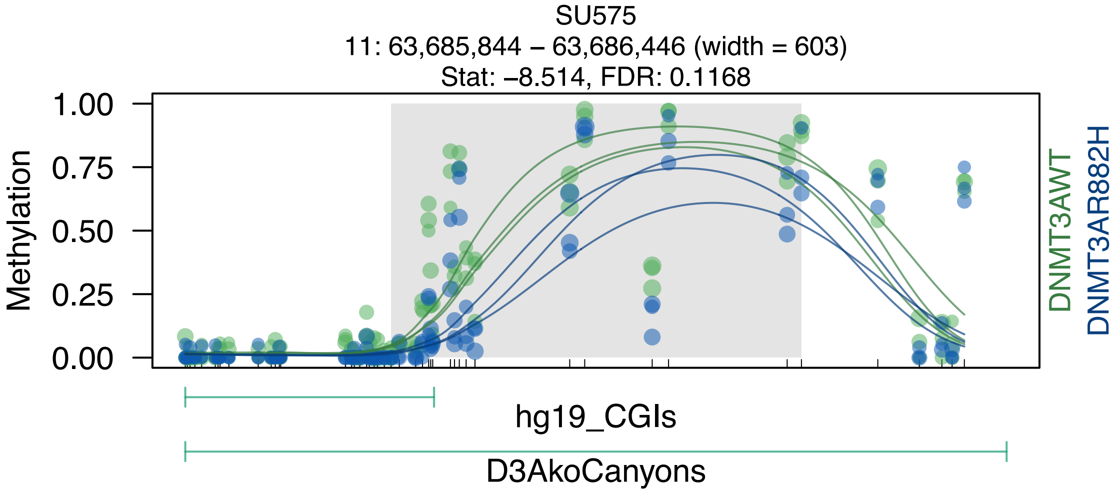
SU575
11: 63,685,844 – 63,686,446 (width = 603)
Stat: –8.514, FDR: 0.1168
Methylation
1.00
0.75
0.50
0.25
0.00
DNMT3AWT
DNMT3AR882H
hg19_CGIs
D3AkoCanyons

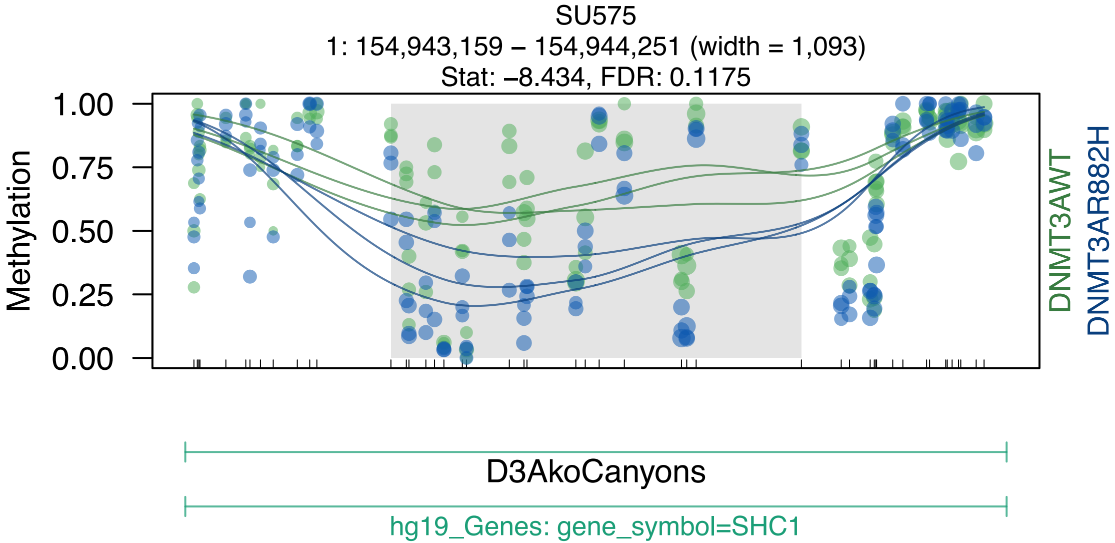
SU575
1: 154,943,159 – 154,944,251 (width = 1,093)
Stat: –8.434, FDR: 0.1175
Methylation
1.00
0.75
0.50
0.25
0.00
DNMT3AWT
DNMT3AR882H
D3AkoCanyons
hg19_Genes: gene_symbol=SHC1

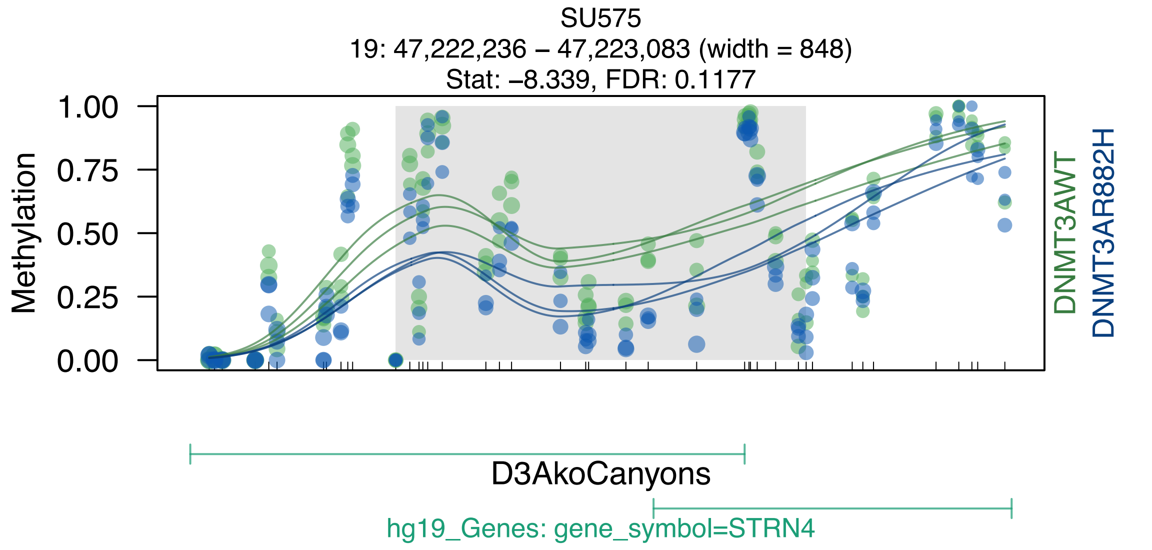

SU575
19: 47,222,236 – 47,223,083 (width = 848)
Stat: –8.339, FDR: 0.1177
Methylation
1.00
0.75
0.50
0.25
0.00
DNMT3AWT
DNMT3AR882H
D3AkoCanyons
hg19_Genes: gene_symbol=STRN4

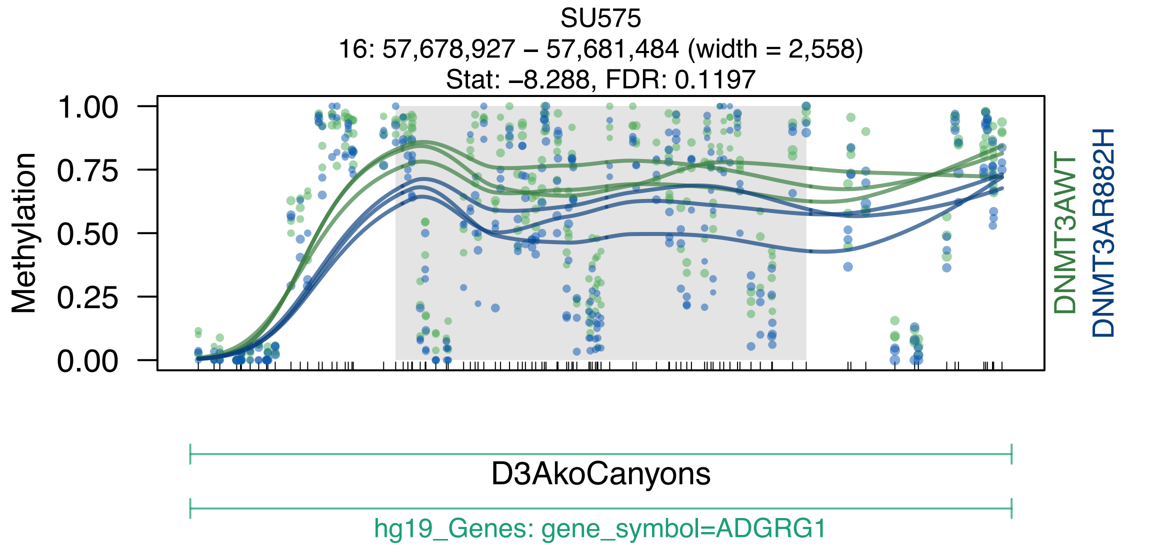

SU575
16: 57,678,927 – 57,681,484 (width = 2,558)
Stat: –8.288, FDR: 0.1197
Methylation
1.00
0.75
0.50
0.25
0.00
DNMT3AWT
DNMT3AR882H
D3AkoCanyons
hg19_Genes: gene_symbol=ADGRG1

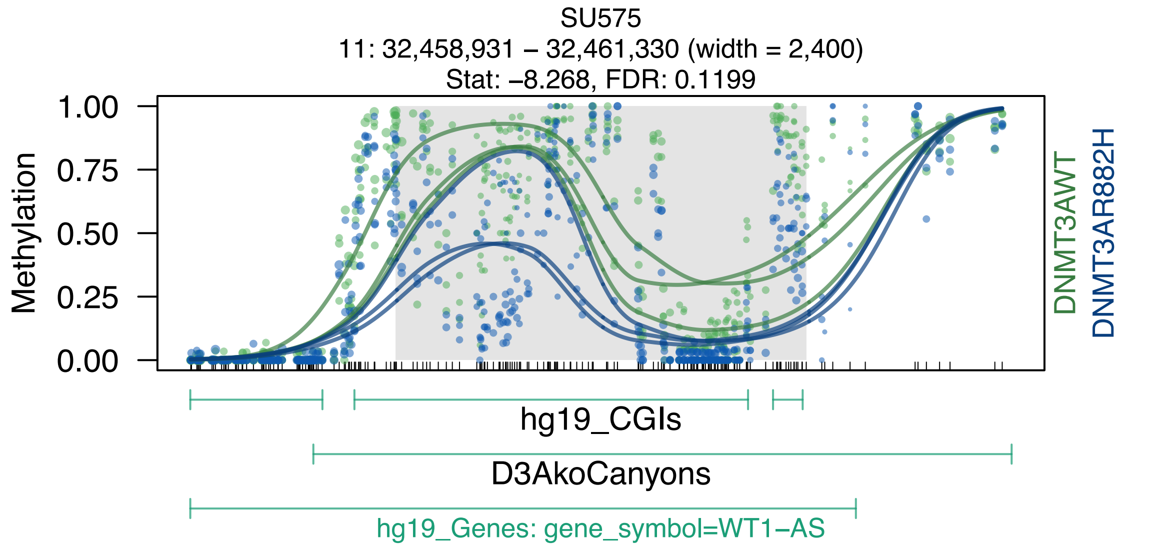
SU575
11: 32,458,931 – 32,461,330 (width = 2,400)
Stat: –8.268, FDR: 0.1199
Methylation
1.00
0.75
0.50
0.25
0.00
DNMT3AWT
DNMT3AR882H
hg19_CGIs
D3AkoCanyons
hg19_Genes: gene_symbol=WT1–AS

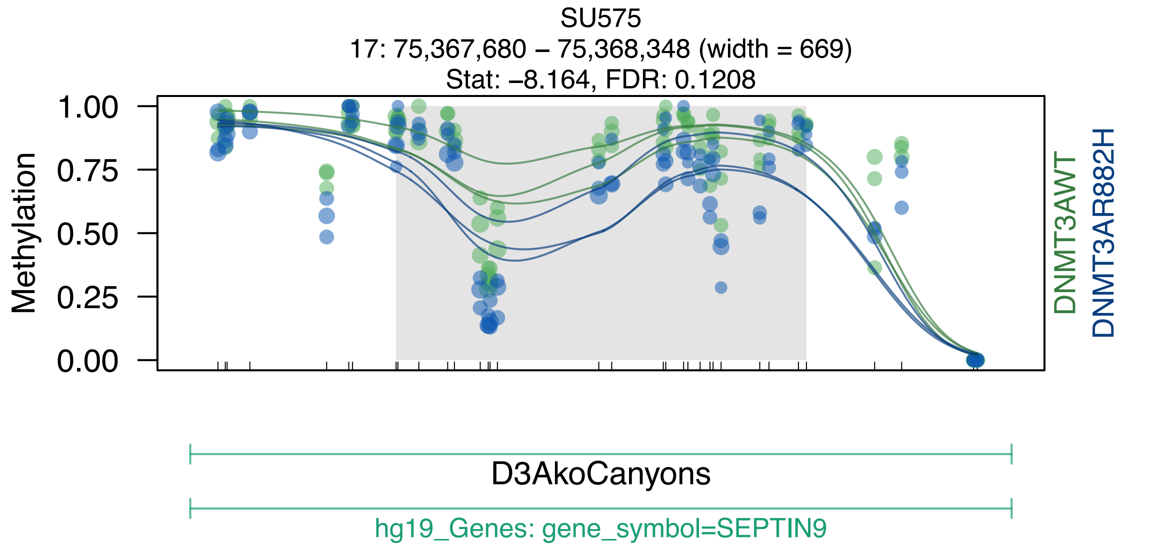
SU575
17: 75,367,680 – 75,368,348 (width = 669)
Stat: –8.164, FDR: 0.1208
Methylation
1.00
0.75
0.50
0.25
0.00
DNMT3AWT
DNMT3AR882H
D3AkoCanyons
hg19_Genes: gene_symbol=SEPTIN9

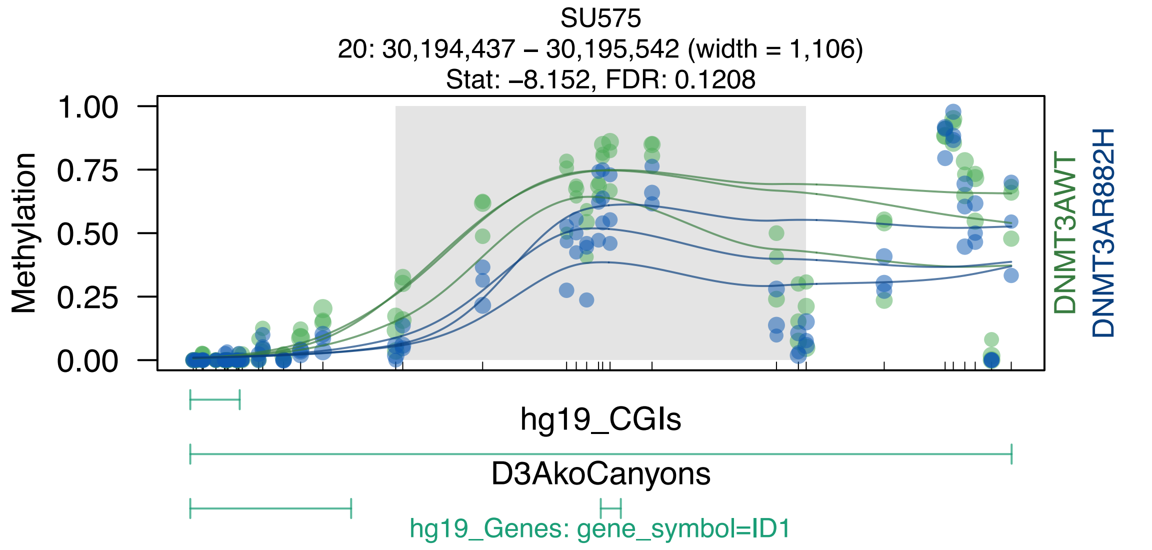
SU575
20: 30,194,437 – 30,195,542 (width = 1,106)
Stat: –8.152, FDR: 0.1208
Methylation
1.00
0.75
0.50
0.25
0.00
DNMT3AWT
DNMT3AR882H
hg19_CGIs
D3AkoCanyons
hg19_Genes: gene_symbol=ID1

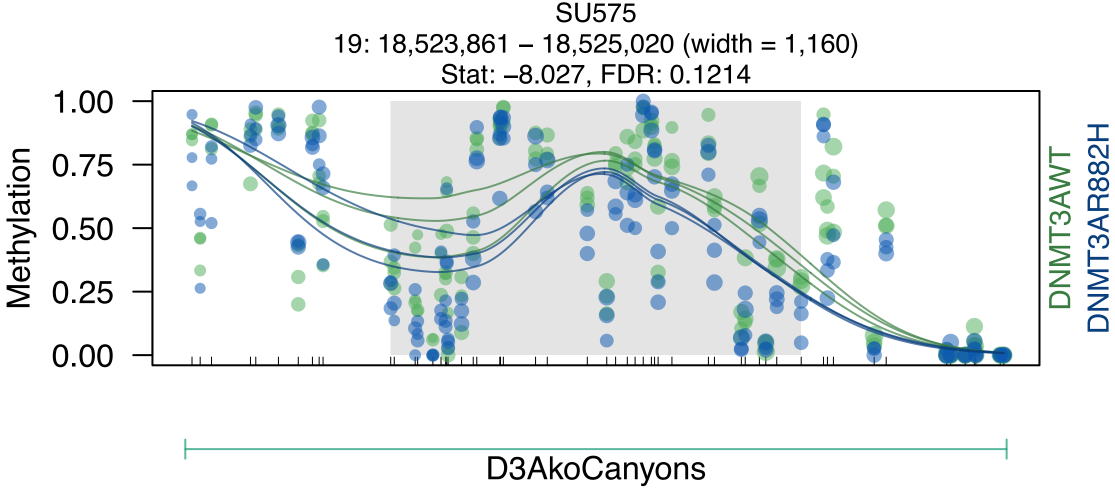
SU575
19: 18,523,861 – 18,525,020 (width = 1,160)
Stat: –8.027, FDR: 0.1214
Methylation
1.00
0.75
0.50
0.25
0.00
DNMT3AWT
DNMT3AR882H
D3AkoCanyons

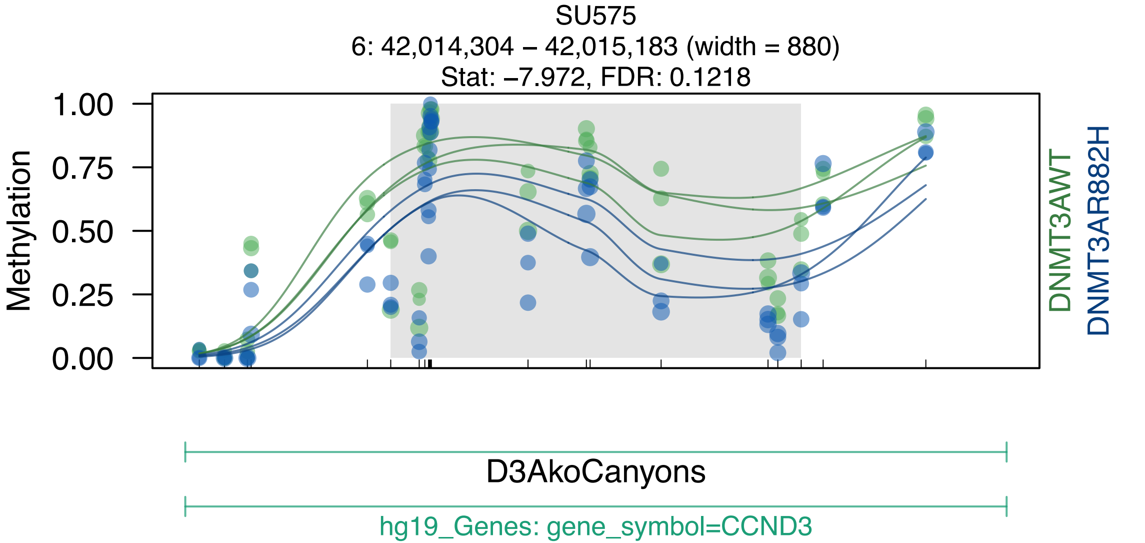
SU575
6: 42,014,304 – 42,015,183 (width = 880)
Stat: –7.972, FDR: 0.1218
Methylation
1.00
0.75
0.50
0.25
0.00
DNMT3AWT
DNMT3AR882H
D3AkoCanyons
hg19_Genes: gene_symbol=CCND3

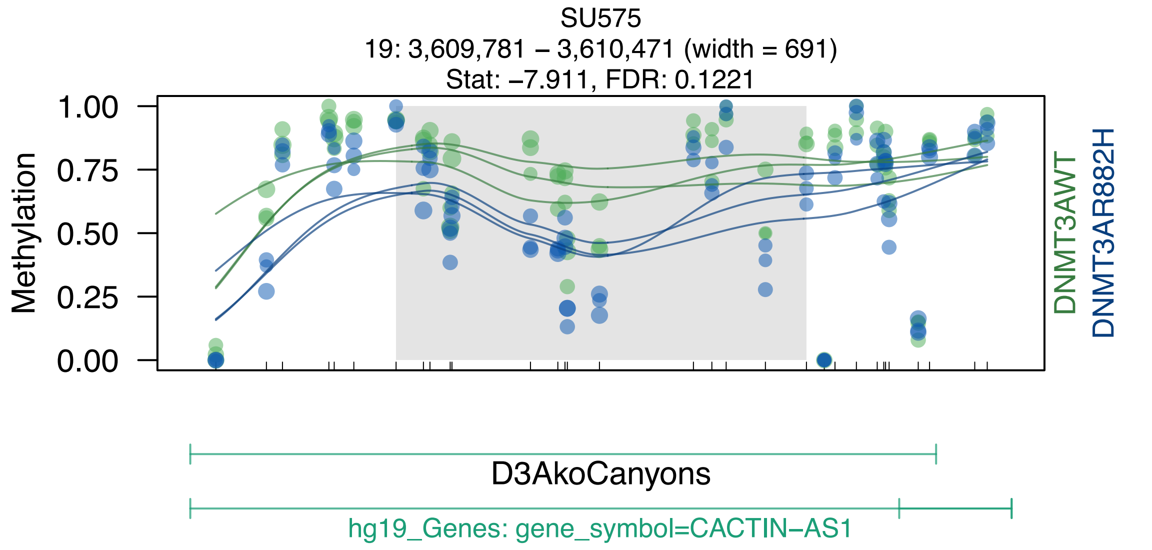
SU575
19: 3,609,781 – 3,610,471 (width = 691)
Stat: −7.911, FDR: 0.1221
Methylation
1.00
0.75
0.50
0.25
0.00
DNMT3AWT
DNMT3AR882H
D3AkoCanyons
hg19_Genes: gene_symbol=CACTIN−AS1

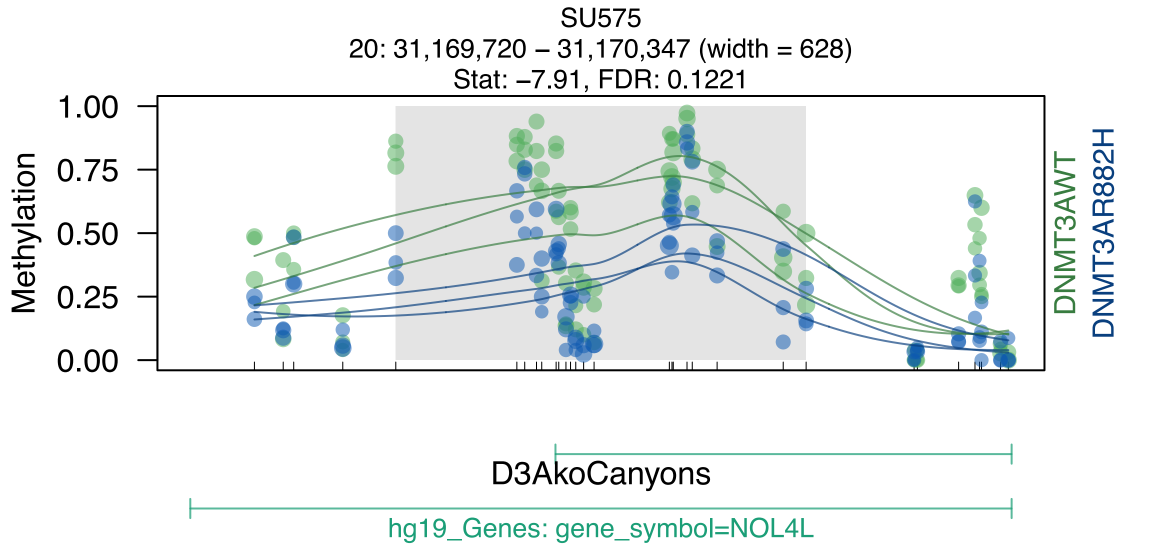
SU575
20: 31,169,720 – 31,170,347 (width = 628)
Stat: –7.91, FDR: 0.1221
Methylation
1.00
0.75
0.50
0.25
0.00
DNMT3AWT
DNMT3AR882H
D3AkoCanyons
hg19_Genes: gene_symbol=NOL4L

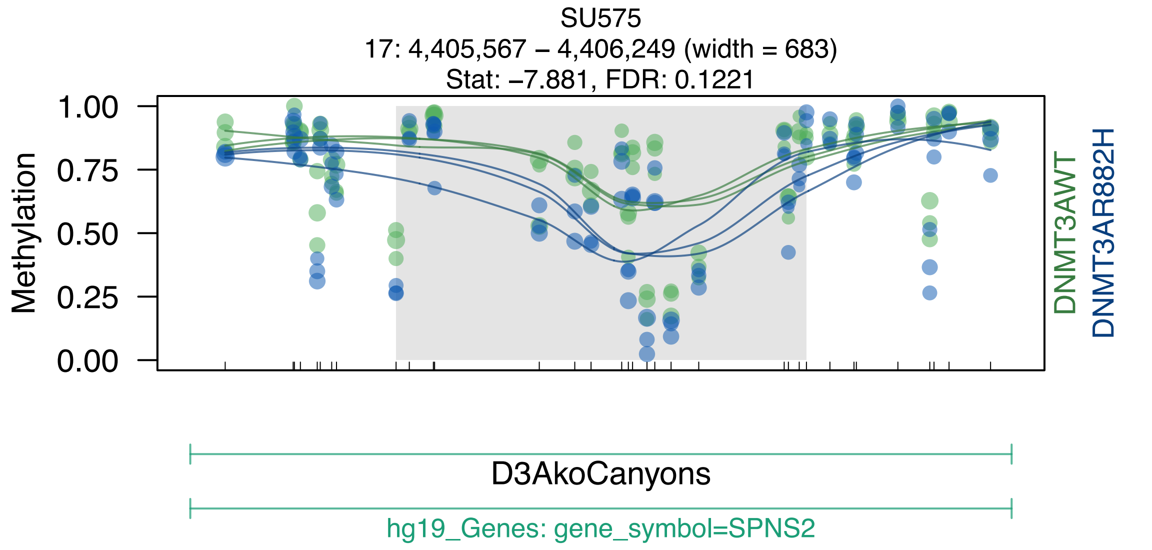
SU575
17: 4,405,567 – 4,406,249 (width = 683)
Stat: −7.881, FDR: 0.1221
Methylation
1.00
0.75
0.50
0.25
0.00
DNMT3AWT
DNMT3AR882H
D3AkoCanyons
hg19_Genes: gene_symbol=SPNS2

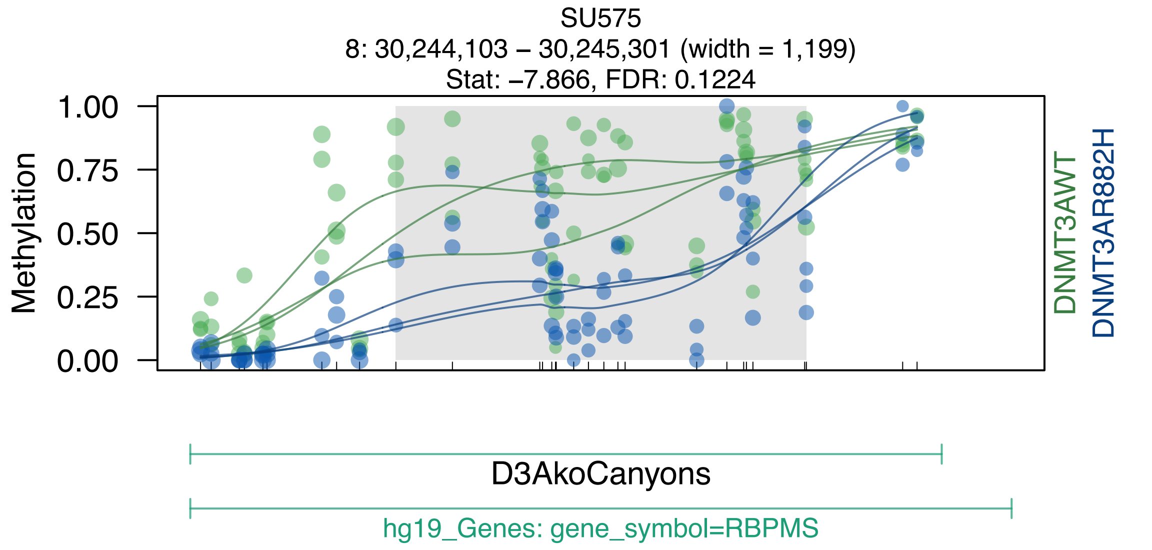
SU575
8: 30,244,103 – 30,245,301 (width = 1,199)
Stat: –7.866, FDR: 0.1224
Methylation
1.00
0.75
0.50
0.25
0.00
DNMT3AWT
DNMT3AR882H
D3AkoCanyons
hg19_Genes: gene_symbol=RBPMS

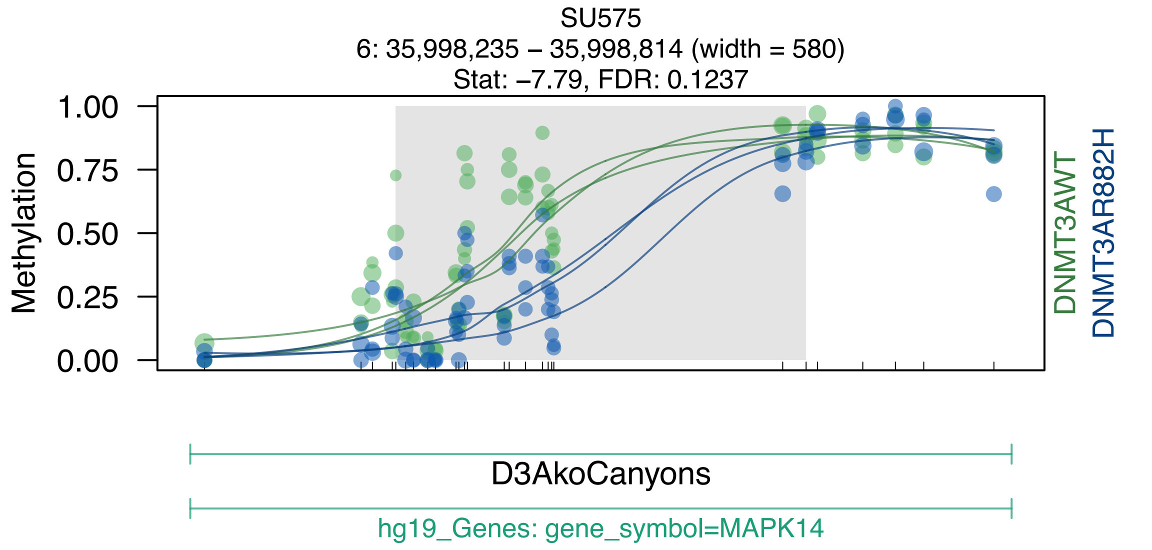
SU575
6: 35,998,235 – 35,998,814 (width = 580)
Stat: –7.79, FDR: 0.1237
Methylation
1.00
0.75
0.50
0.25
0.00
DNMT3AWT
DNMT3AR882H
D3AkoCanyons
hg19_Genes: gene_symbol=MAPK14

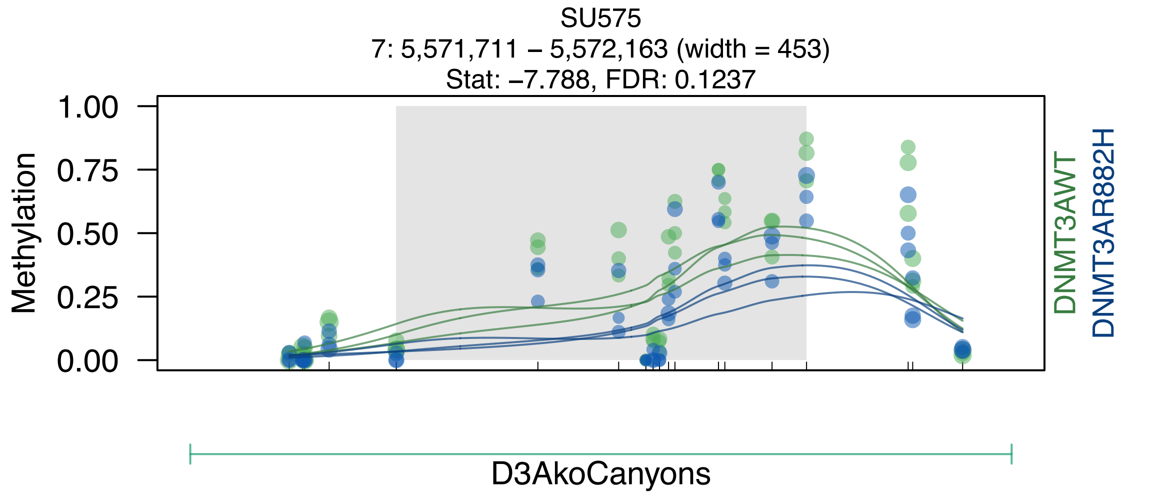

SU575
7: 5,571,711 – 5,572,163 (width = 453)
Stat: –7.788, FDR: 0.1237
Methylation
1.00
0.75
0.50
0.25
0.00
DNMT3AWT
DNMT3AR882H
D3AkoCanyons

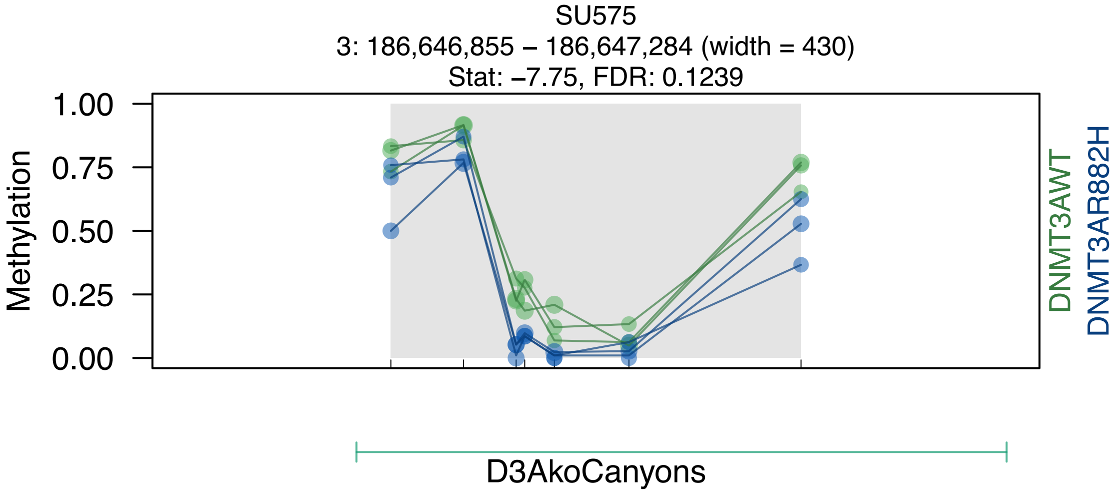
SU575
3: 186,646,855 – 186,647,284 (width = 430)
Stat: –7.75, FDR: 0.1239
Methylation
1.00
0.75
0.50
0.25
0.00
DNMT3AWT
DNMT3AR882H
D3AkoCanyons

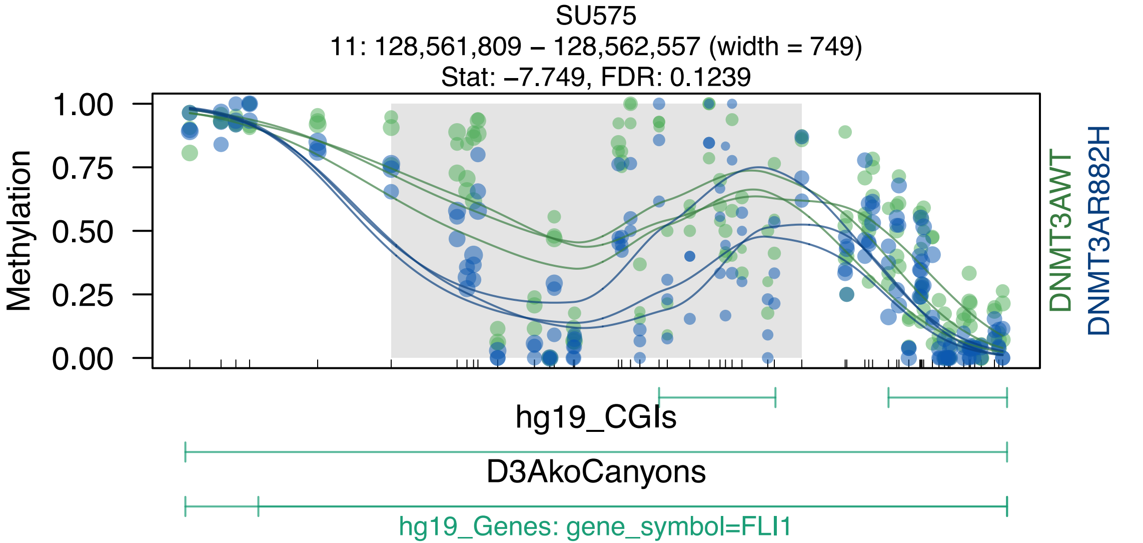

SU575
11: 128,561,809 – 128,562,557 (width = 749)
Stat: –7.749, FDR: 0.1239
Methylation
1.00
0.75
0.50
0.25
0.00
DNMT3AWT
DNMT3AR882H
hg19_CGIs
D3AkoCanyons
hg19_Genes: gene_symbol=FLI1

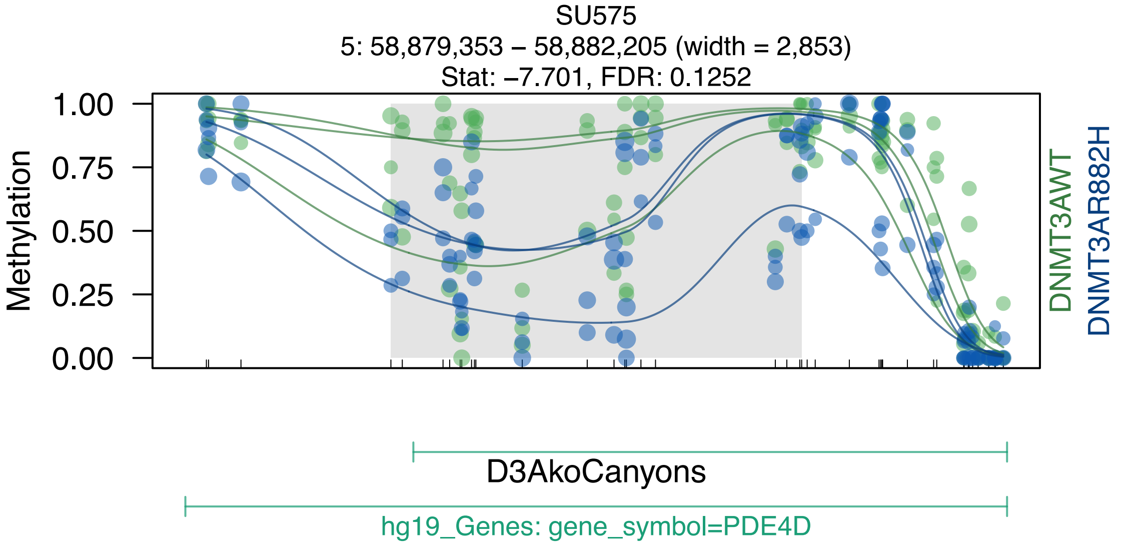
SU575
5: 58,879,353 – 58,882,205 (width = 2,853)
Stat: –7.701, FDR: 0.1252
Methylation
1.00
0.75
0.50
0.25
0.00
DNMT3AWT
DNMT3AR882H
D3AkoCanyons
hg19_Genes: gene_symbol=PDE4D

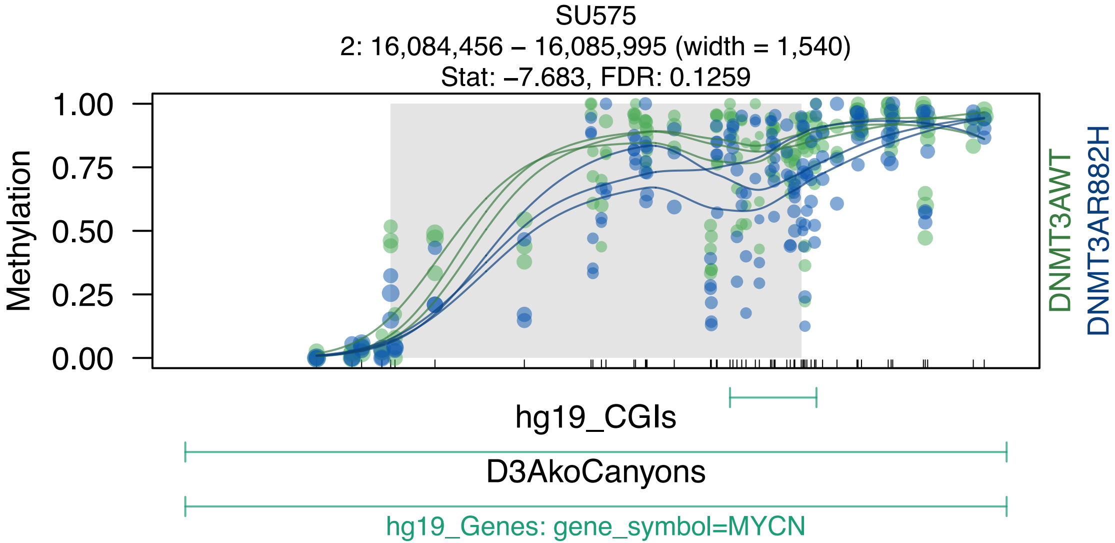
SU575
2: 16,084,456 – 16,085,995 (width = 1,540)
Stat: −7.683, FDR: 0.1259
Methylation
1.00
0.75
0.50
0.25
0.00
DNMT3AWT
DNMT3AR882H
hg19_CGIs
D3AkoCanyons
hg19_Genes: gene_symbol=MYCN

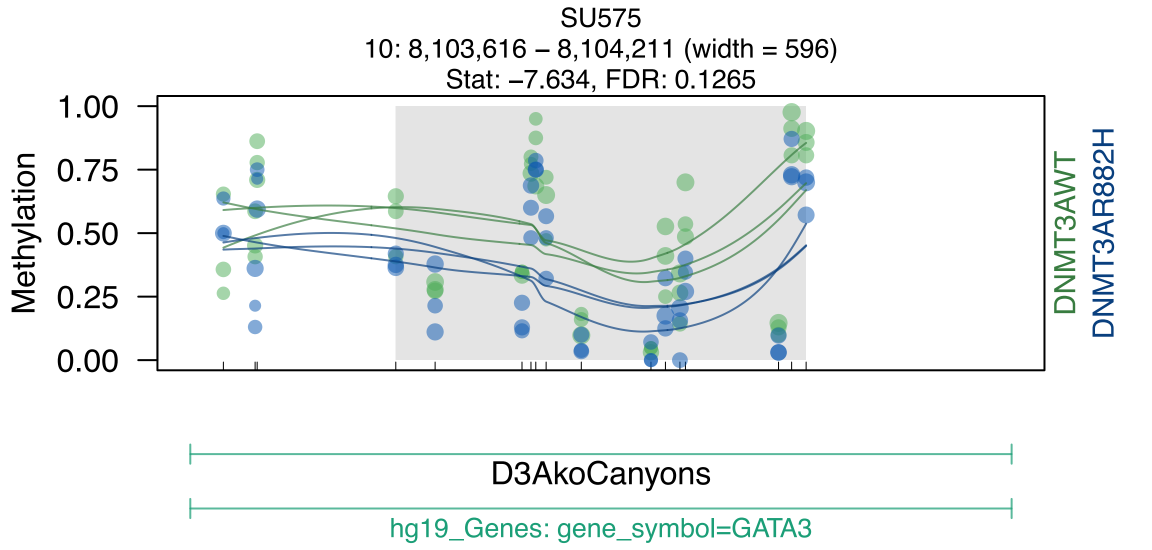
SU575
10: 8,103,616 – 8,104,211 (width = 596)
Stat: –7.634, FDR: 0.1265
Methylation
1.00
0.75
0.50
0.25
0.00
DNMT3AWT
DNMT3AR882H
D3AkoCanyons
hg19_Genes: gene_symbol=GATA3

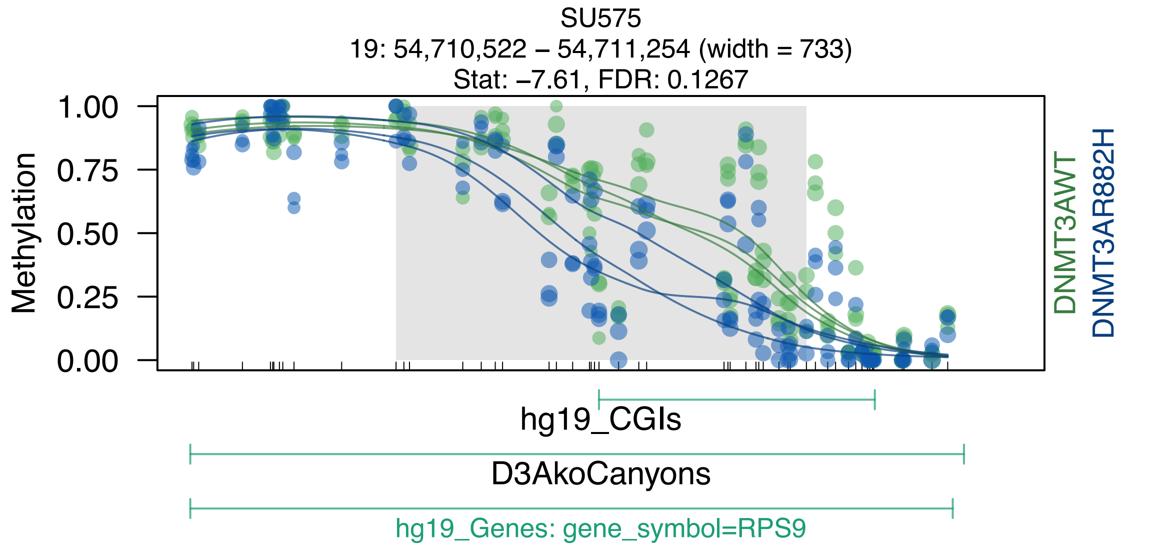
SU575
19: 54,710,522 – 54,711,254 (width = 733)
Stat: –7.61, FDR: 0.1267
Methylation
1.00
0.75
0.50
0.25
0.00
DNMT3AWT
DNMT3AR882H
hg19_CGIs
D3AkoCanyons
hg19_Genes: gene_symbol=RPS9

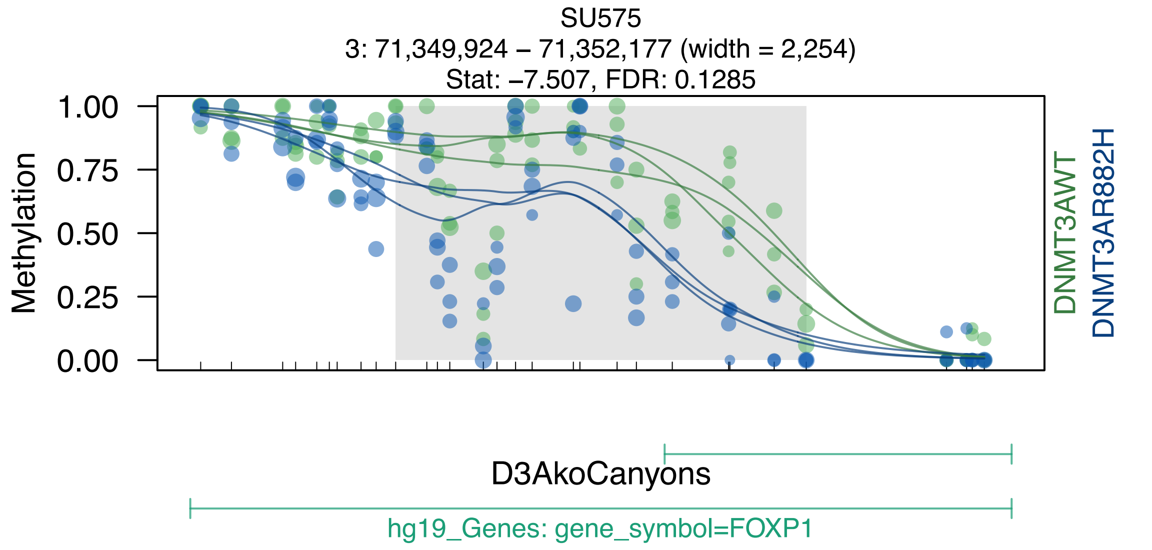

SU575
3: 71,349,924 – 71,352,177 (width = 2,254)
Stat: –7.507, FDR: 0.1285
Methylation
1.00
0.75
0.50
0.25
0.00
DNMT3AWT
DNMT3AR882H
D3AkoCanyons
hg19_Genes: gene_symbol=FOXP1

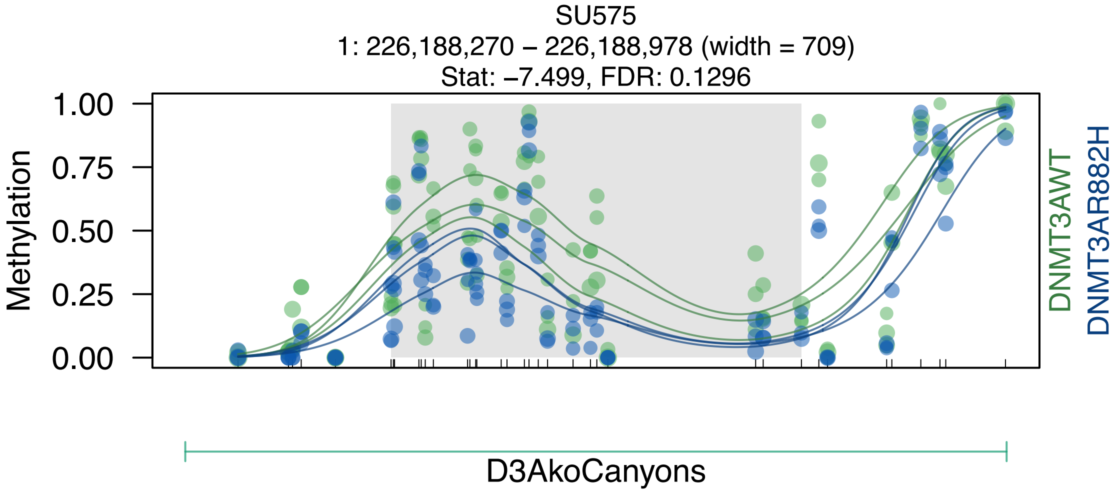
SU575
1: 226,188,270 – 226,188,978 (width = 709)
Stat: −7.499, FDR: 0.1296
Methylation
1.00
0.75
0.50
0.25
0.00
DNMT3AWT
DNMT3AR882H
D3AkoCanyons

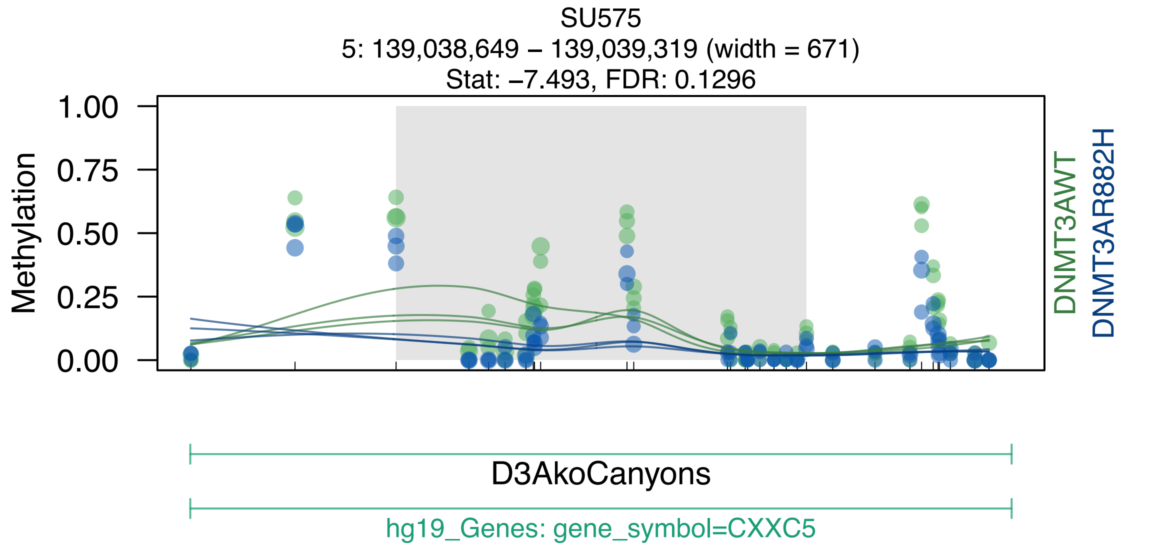
SU575
5: 139,038,649 – 139,039,319 (width = 671)
Stat: –7.493, FDR: 0.1296
Methylation
1.00
0.75
0.50
0.25
0.00
DNMT3AWT
DNMT3AR882H
D3AkoCanyons
hg19_Genes: gene_symbol=CXXC5
