## Supplemental Data S1-S7 for "DNMT3A^R882H^ Is Not Required for Disease Maintenance in Primary Human AML, but Is Associated With Increased Leukemia Stem Cell Frequency": DataS7_top100Canyons_AllSampAndCD34_MethOverCanyons.pdf

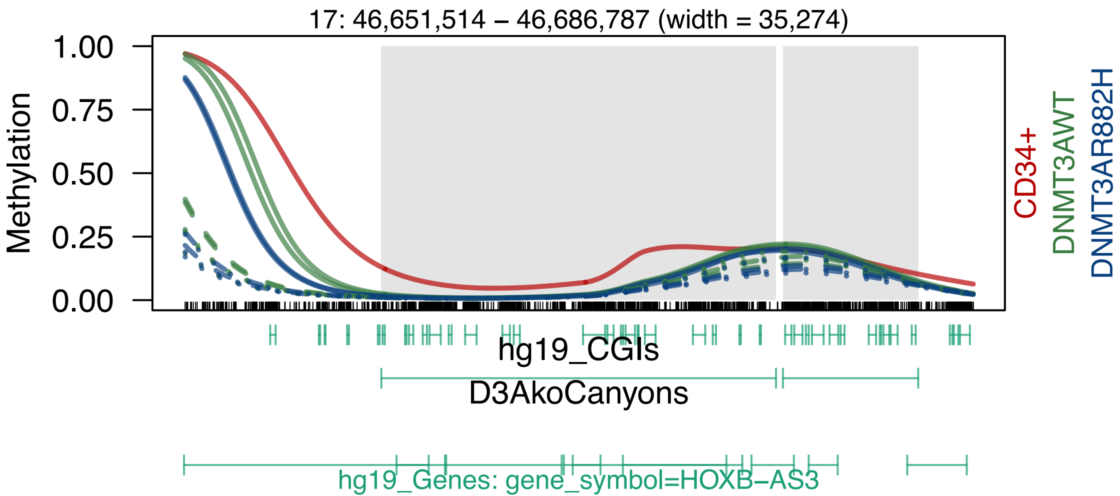
17: 46,651,514 – 46,686,787 (width = 35,274)
Methylation
1.00
0.75
0.50
0.25
0.00
CD34+
DNMT3AWT
DNMT3AR882H
hg19_CGIs
D3AkoCanyons
hg19_Genes: gene_symbol=HOXB–AS3

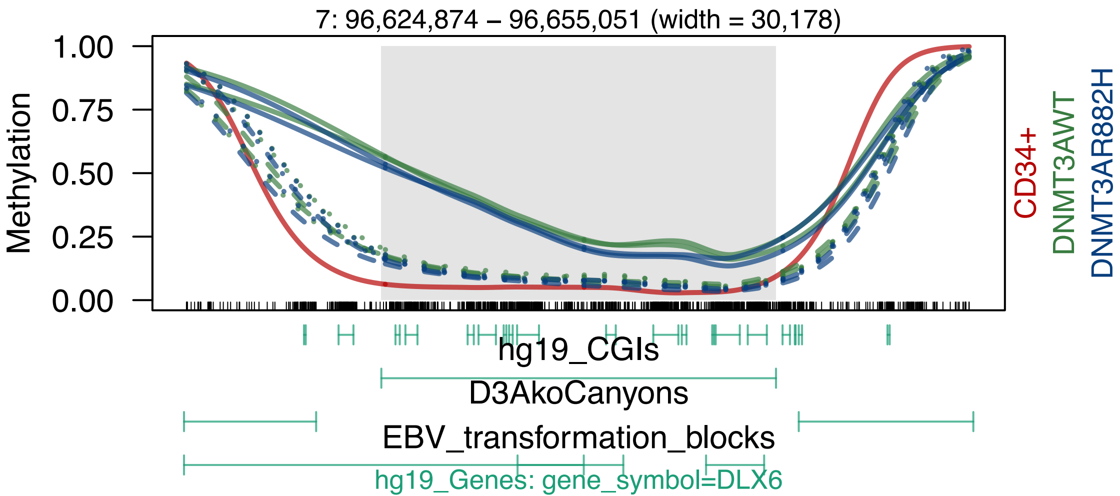

7: 96,624,874 – 96,655,051 (width = 30,178)
1.00
0.75
0.50
0.25
0.00
Methylation
CD34+
DNMT3AWT
DNMT3AR882H
hg19_CGIs
D3AkoCanyons
EBV_transformation_blocks
hg19_Genes: gene_symbol=DLX6

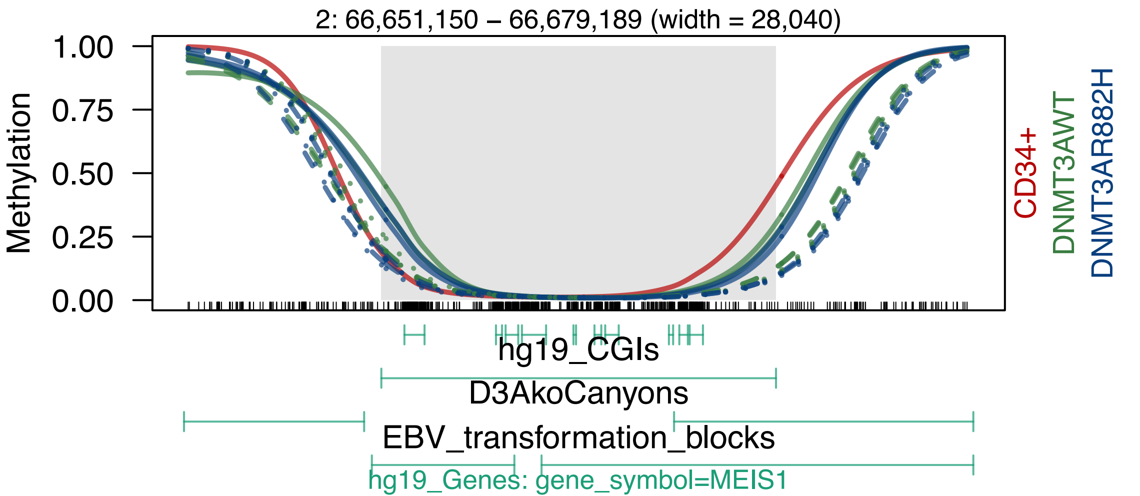

2: 66,651,150 – 66,679,189 (width = 28,040)
Methylation
1.00
0.75
0.50
0.25
0.00
CD34+
DNMT3AWT
DNMT3AR882H
hg19_CGIs
D3AkoCanyons
EBV_transformation_blocks
hg19_Genes: gene_symbol=MEIS1

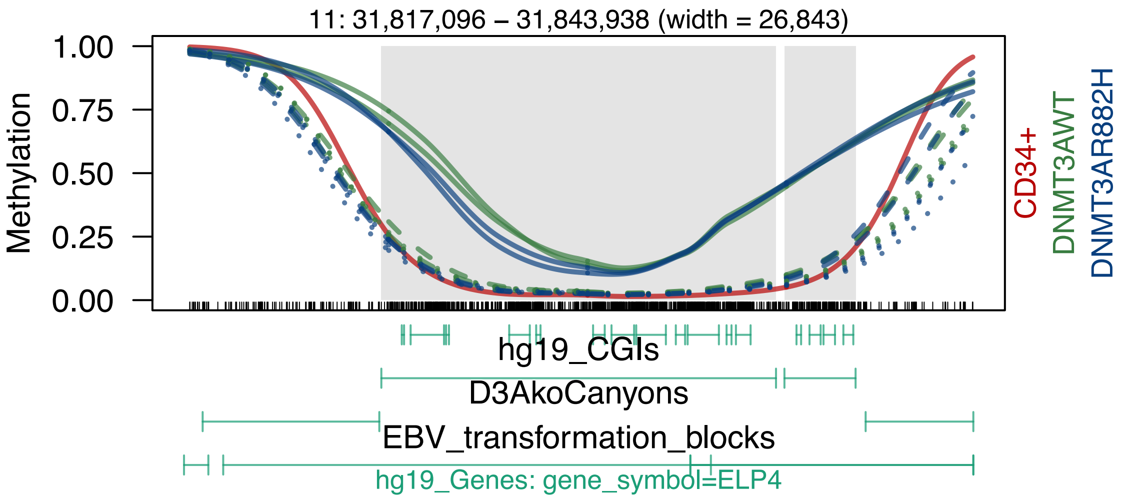
11: 31,817,096 – 31,843,938 (width = 26,843)
Methylation
1.00
0.75
0.50
0.25
0.00
CD34+
DNMT3AWT
DNMT3AR882H
hg19_CGIs
D3AkoCanyons
EBV_transformation_blocks
hg19_Genes: gene_symbol=ELP4

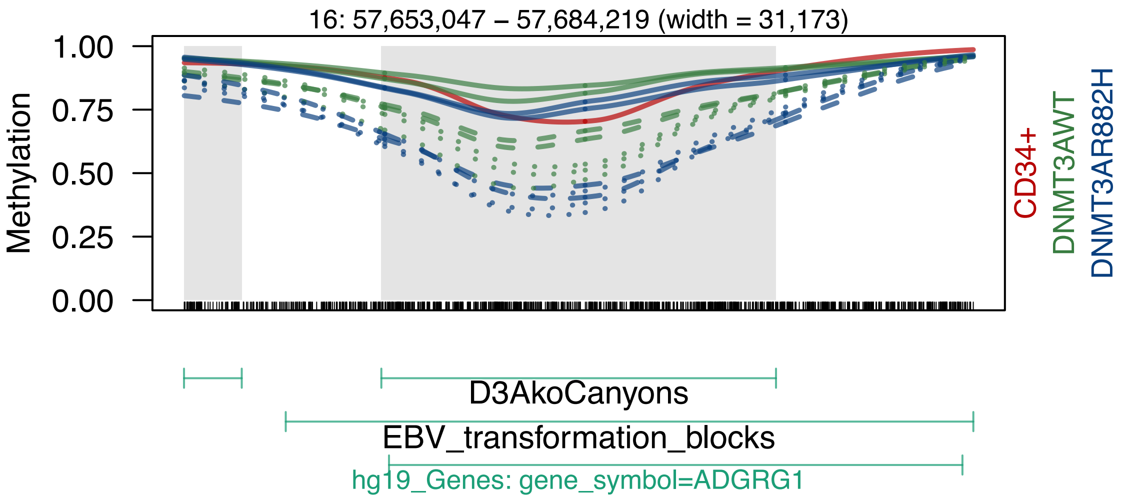
16: 57,653,047 – 57,684,219 (width = 31,173)
Methylation
1.00
0.75
0.50
0.25
0.00
CD34+
DNMT3AWT
DNMT3AR882H
D3AkoCanyons
EBV_transformation_blocks
hg19_Genes: gene_symbol=ADGRG1

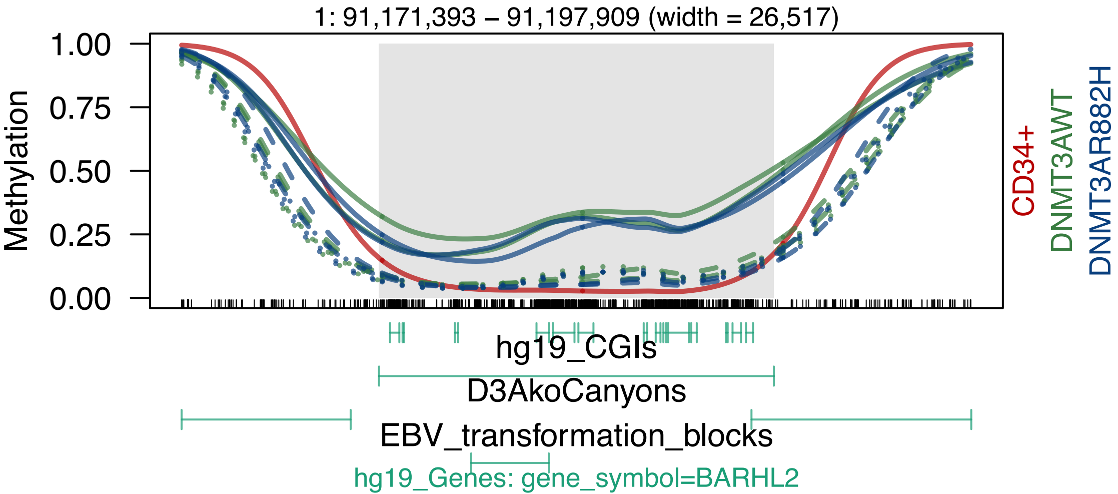

1: 91,171,393 – 91,197,909 (width = 26,517)
Methylation
1.00
0.75
0.50
0.25
0.00
CD34+
DNMT3AWT
DNMT3AR882H
hg19_CGIs
D3AkoCanyons
EBV_transformation_blocks
hg19_Genes: gene_symbol=BARHL2

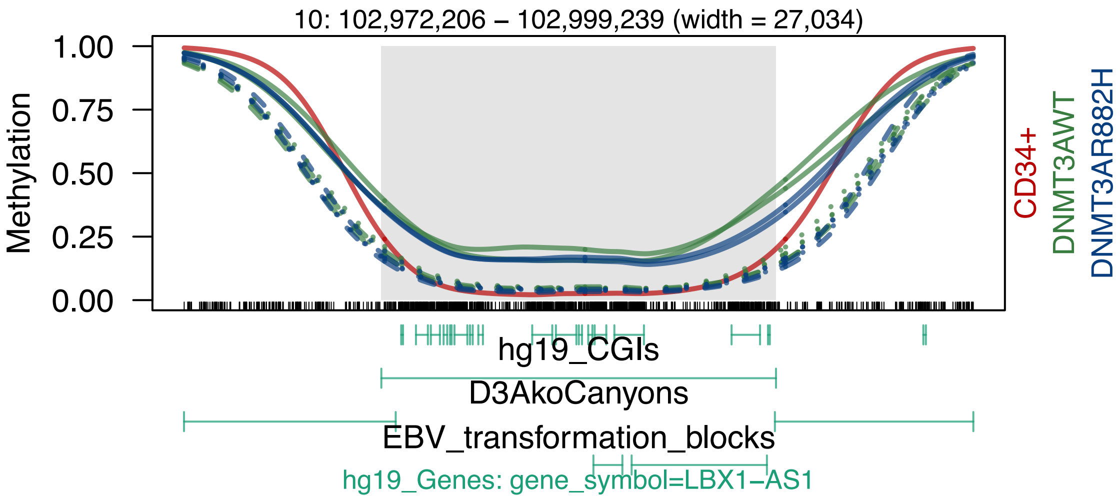
10: 102,972,206 – 102,999,239 (width = 27,034)
Methylation
1.00
0.75
0.50
0.25
0.00
CD34+
DNMT3AWT
DNMT3AR882H
hg19_CGIs
D3AkoCanyons
EBV_transformation_blocks
hg19_Genes: gene_symbol=LBX1−AS1

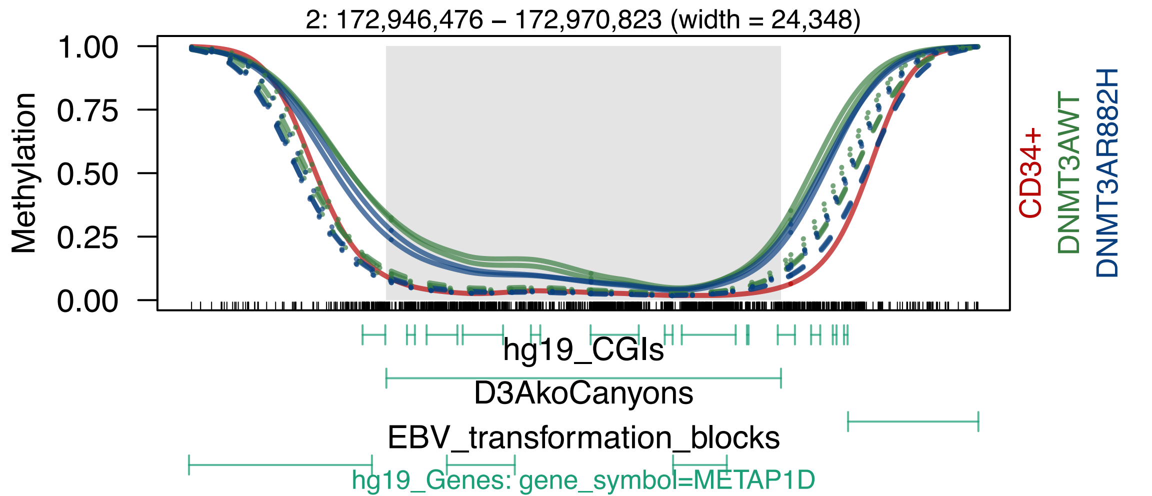

2: 172,946,476 – 172,970,823 (width = 24,348)
Methylation
1.00
0.75
0.50
0.25
0.00
CD34+
DNMT3AWT
DNMT3AR882H
hg19_CGIs
D3AkoCanyons
EBV_transformation_blocks
hg19_Genes: gene_symbol=METAP1D

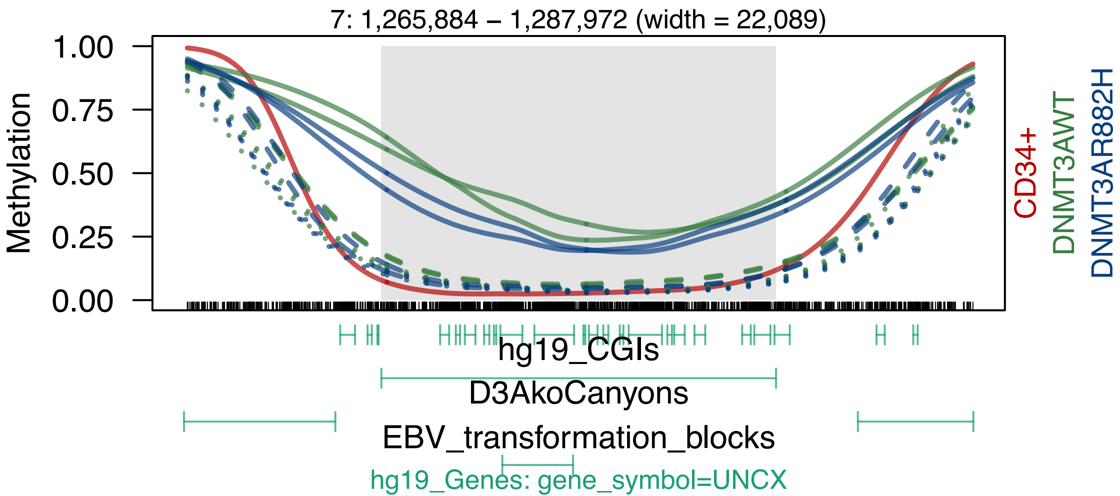
7: 1,265,884 – 1,287,972 (width = 22,089)
Methylation
1.00
0.75
0.50
0.25
0.00
CD34+
DNMT3AWT
DNMT3AR882H
hg19_CGIs
D3AkoCanyons
EBV_transformation_blocks
hg19_Genes: gene_symbol=UNCX

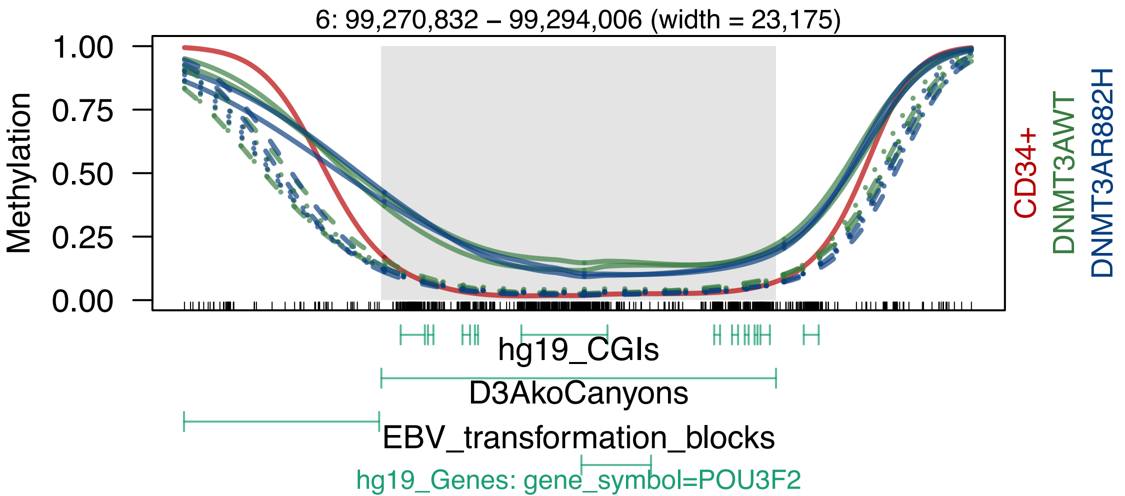

6: 99,270,832 – 99,294,006 (width = 23,175)
Methylation
1.00
0.75
0.50
0.25
0.00
CD34+
DNMT3AWT
DNMT3AR882H
hg19_CGIs
D3AkoCanyons
EBV_transformation_blocks
hg19_Genes: gene_symbol=POU3F2

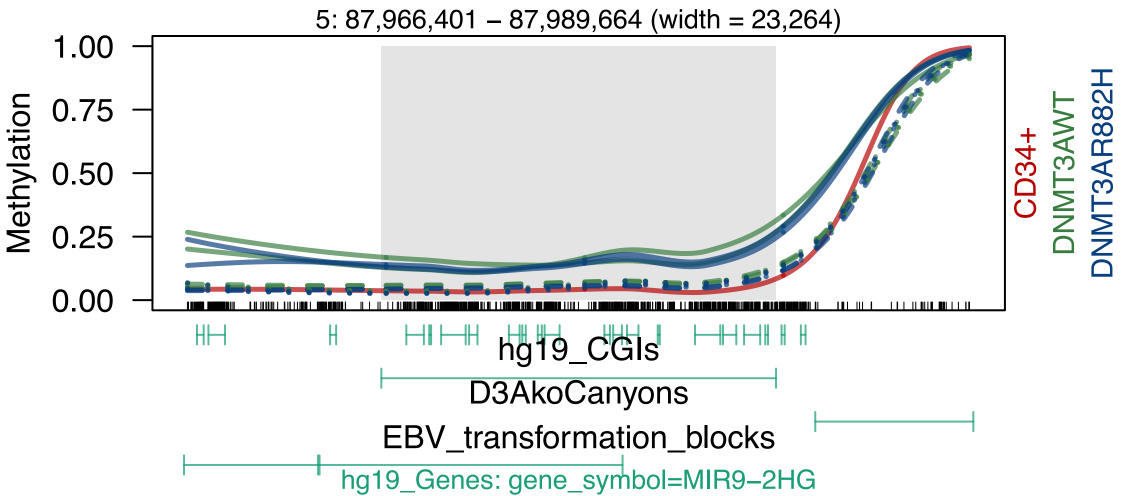
5: 87,966,401 – 87,989,664 (width = 23,264)
Methylation
1.00
0.75
0.50
0.25
0.00
CD34+
DNMT3AWT
DNMT3AR882H
hg19_CGIs
D3AkoCanyons
EBV_transformation_blocks
hg19_Genes: gene_symbol=MIR9−2HG

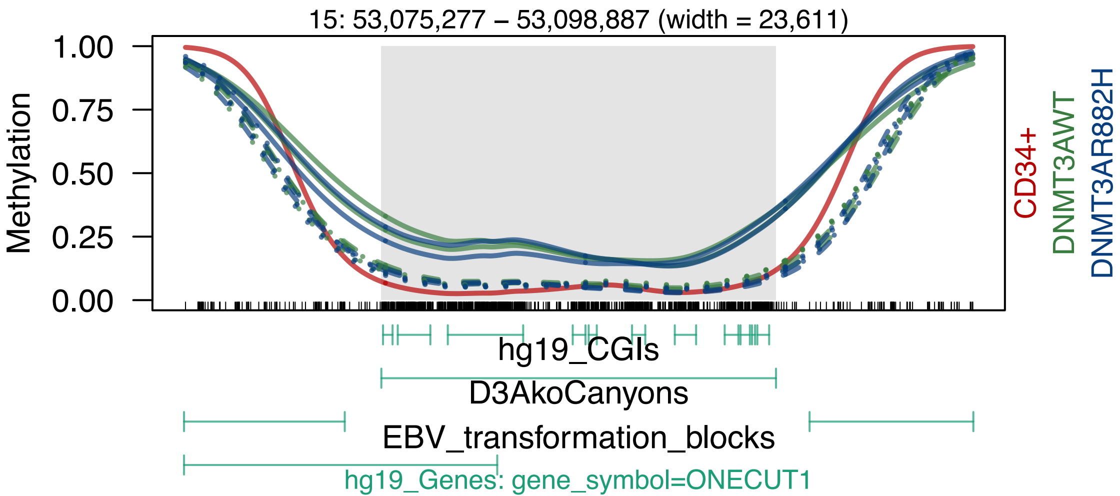
15: 53,075,277 – 53,098,887 (width = 23,611)
Methylation
1.00
0.75
0.50
0.25
0.00
CD34+
DNMT3AWT
DNMT3AR882H
hg19_CGIs
D3AkoCanyons
EBV_transformation_blocks
hg19_Genes: gene_symbol=ONECUT1

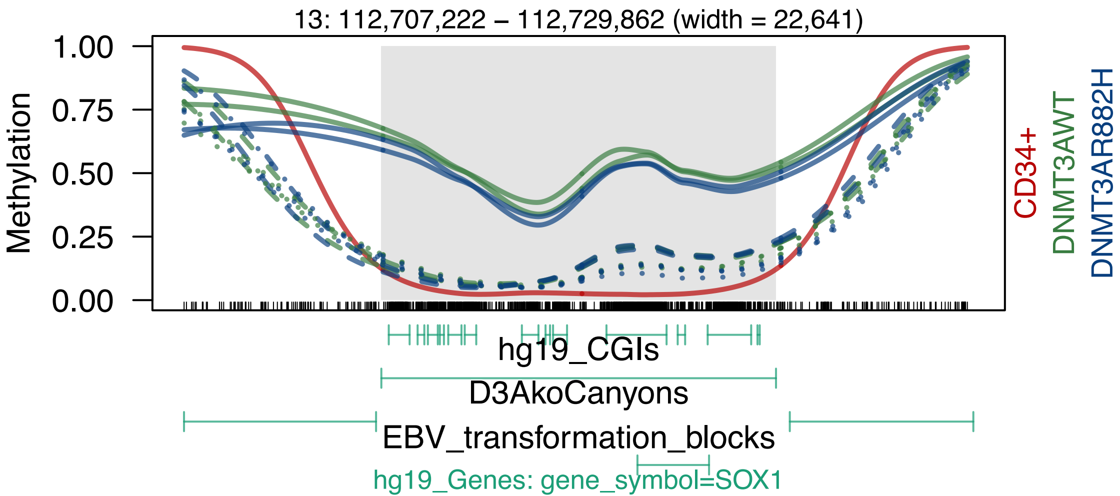

13: 112,707,222 – 112,729,862 (width = 22,641)
Methylation
1.00
0.75
0.50
0.25
0.00
CD34+
DNMT3AWT
DNMT3AR882H
hg19_CGIs
D3AkoCanyons
EBV_transformation_blocks
hg19_Genes: gene_symbol=SOX1

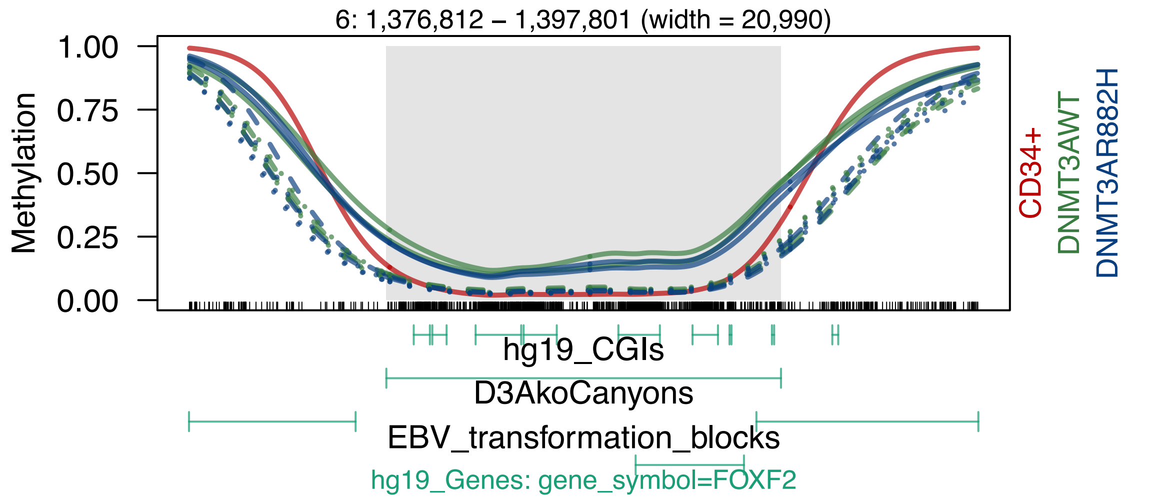
6: 1,376,812 – 1,397,801 (width = 20,990)
Methylation
1.00
0.75
0.50
0.25
0.00
CD34+
DNMT3AWT
DNMT3AR882H
hg19_CGIs
D3AkoCanyons
EBV_transformation_blocks
hg19_Genes: gene_symbol=FOXF2

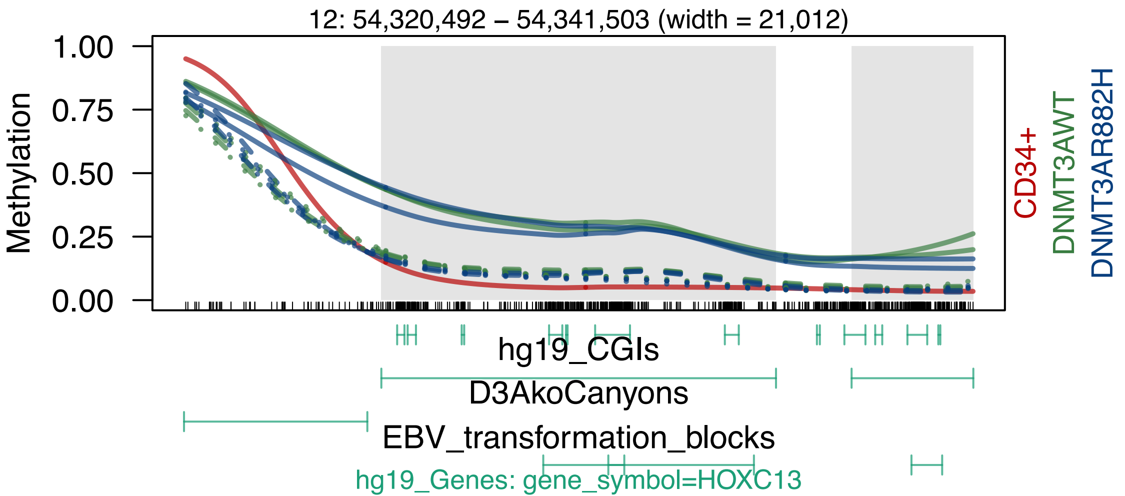
12: 54,320,492 – 54,341,503 (width = 21,012)
Methylation
1.00
0.75
0.50
0.25
0.00
CD34+
DNMT3AWT
DNMT3AR882H
hg19_CGIs
D3AkoCanyons
EBV_transformation_blocks
hg19_Genes: gene_symbol=HOXC13

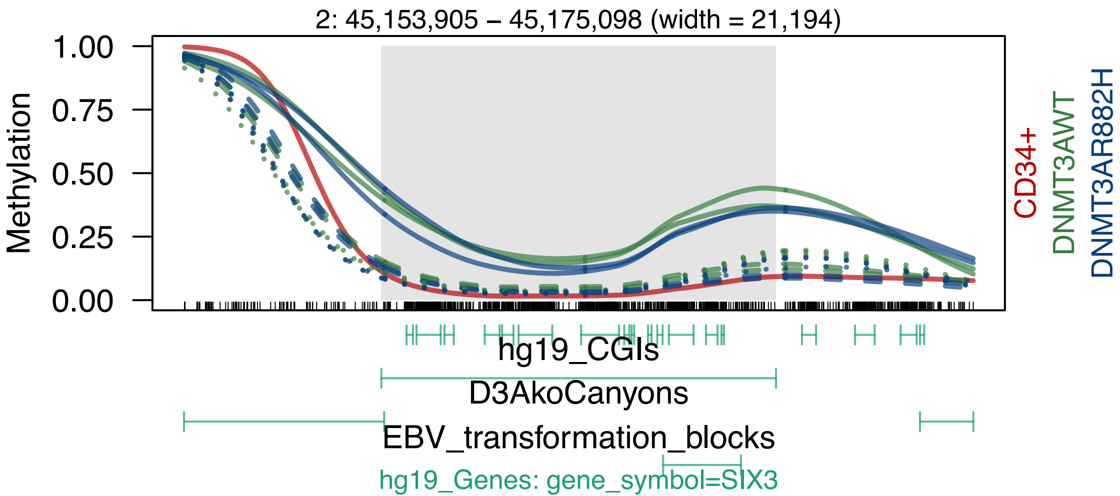
2: 45,153,905 – 45,175,098 (width = 21,194)
Methylation
1.00
0.75
0.50
0.25
0.00
CD34+
DNMT3AWT
DNMT3AR882H
hg19_CGIs
D3AkoCanyons
EBV_transformation_blocks
hg19_Genes: gene_symbol=SIX3

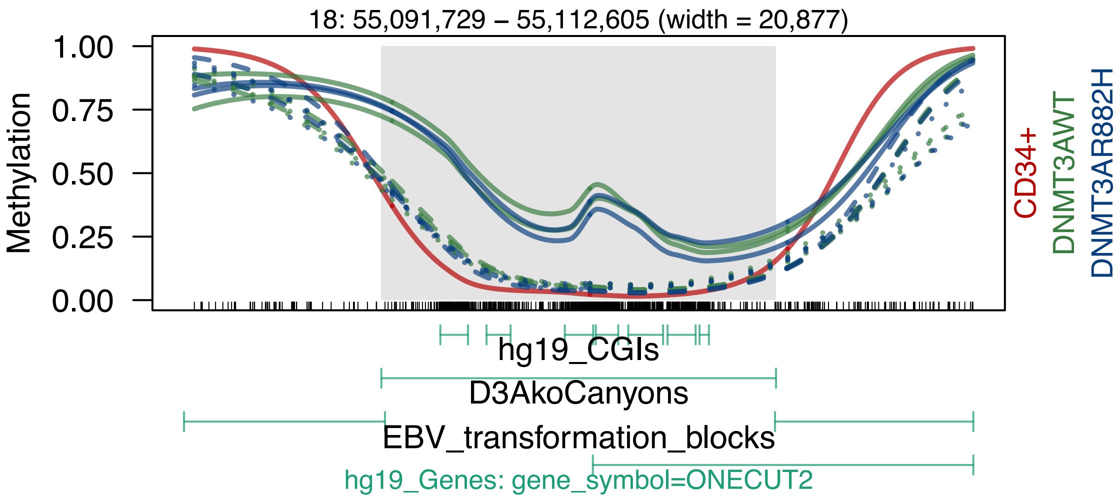
18: 55,091,729 – 55,112,605 (width = 20,877)
1.00
0.75
0.50
0.25
0.00
Methylation
CD34+
DNMT3AWT
DNMT3AR882H
hg19_CGIs
D3AkoCanyons
EBV_transformation_blocks
hg19_Genes: gene_symbol=ONECUT2

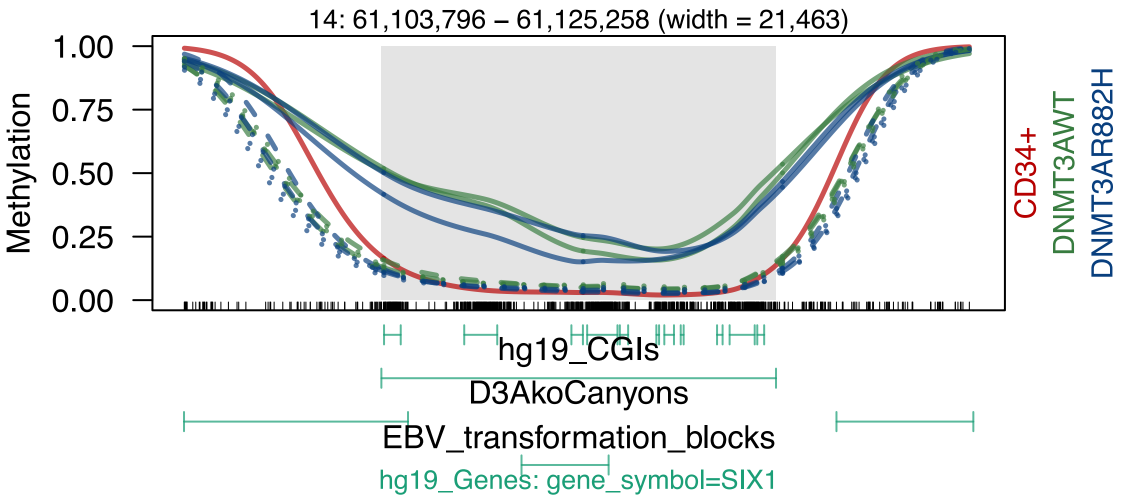
14: 61,103,796 – 61,125,258 (width = 21,463)
Methylation
1.00
0.75
0.50
0.25
0.00
CD34+
DNMT3AWT
DNMT3AR882H
hg19_CGIs
D3AkoCanyons
EBV_transformation_blocks
hg19_Genes: gene_symbol=SIX1

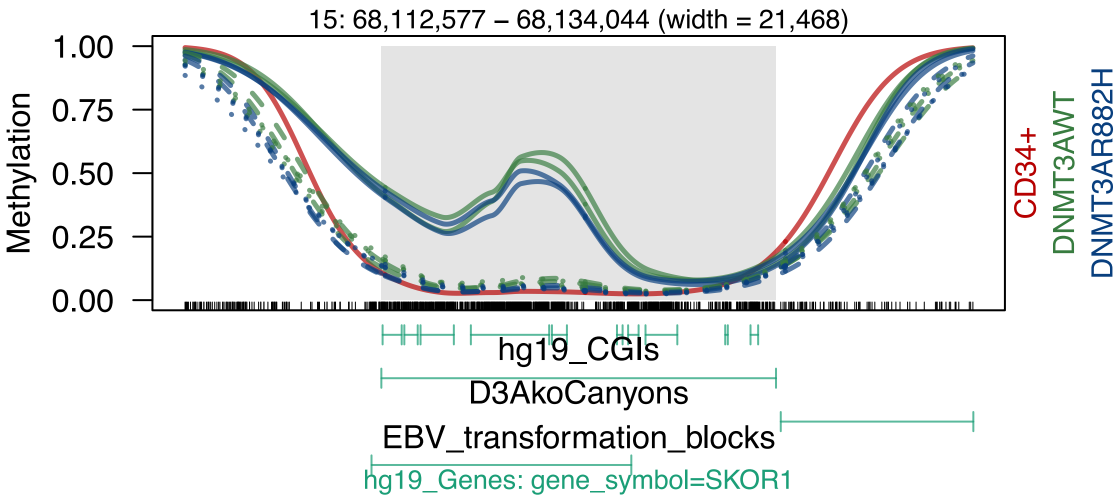
15: 68,112,577 – 68,134,044 (width = 21,468)
Methylation
1.00
0.75
0.50
0.25
0.00
CD34+
DNMT3AWT
DNMT3AR882H
hg19_CGIs
D3AkoCanyons
EBV_transformation_blocks
hg19_Genes: gene_symbol=SKOR1

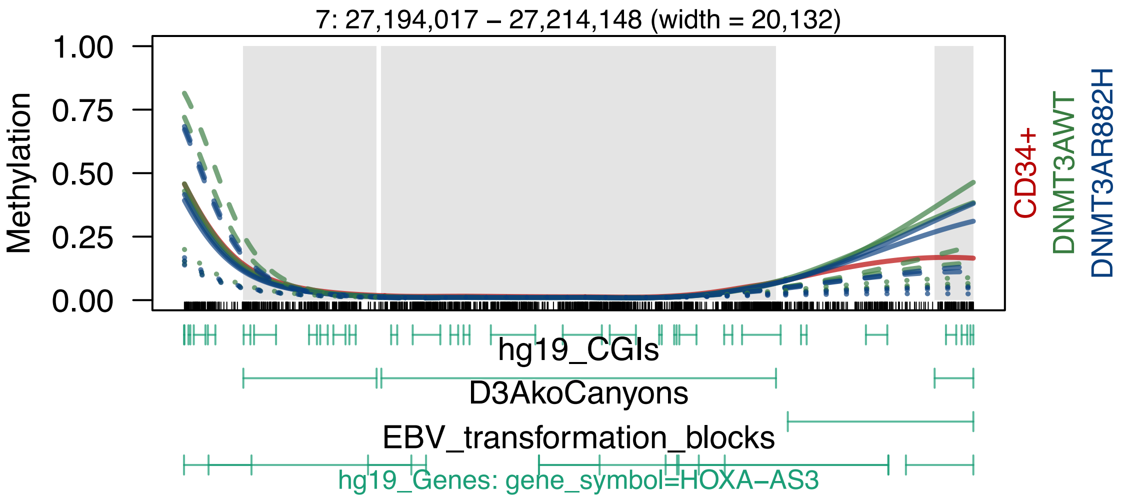
7: 27,194,017 – 27,214,148 (width = 20,132)
Methylation
1.00
0.75
0.50
0.25
0.00
CD34+
DNMT3AWT
DNMT3AR882H
hg19_CGIs
D3AkoCanyons
EBV_transformation_blocks
hg19_Genes: gene_symbol=HOXA–AS3

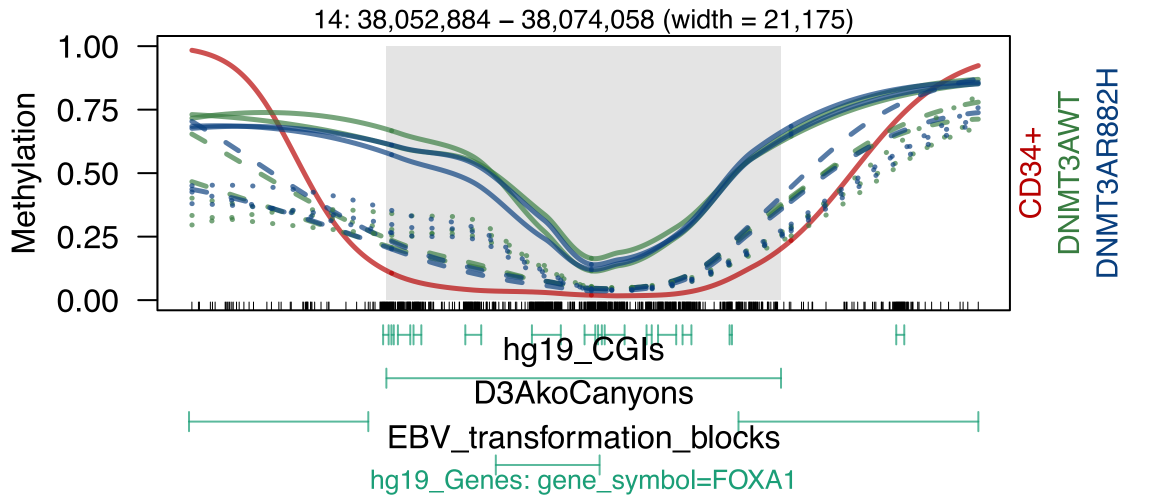

14: 38,052,884 – 38,074,058 (width = 21,175)
Methylation
1.00
0.75
0.50
0.25
0.00
CD34+
DNMT3AWT
DNMT3AR882H
hg19_CGIs
D3AkoCanyons
EBV_transformation_blocks
hg19_Genes: gene_symbol=FOXA1

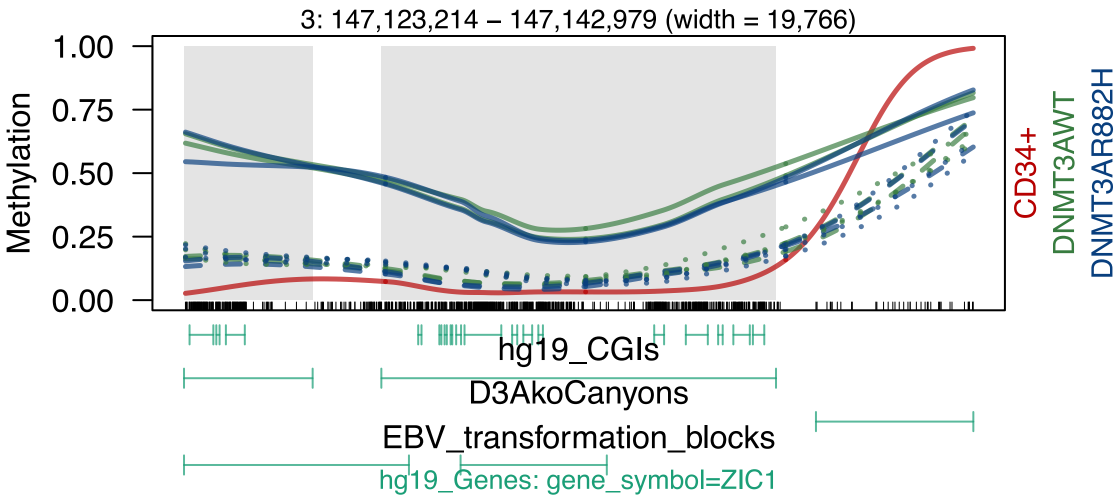
3: 147,123,214 – 147,142,979 (width = 19,766)
Methylation
1.00
0.75
0.50
0.25
0.00
CD34+
DNMT3AWT
DNMT3AR882H
hg19_CGIs
D3AkoCanyons
EBV_transformation_blocks
hg19_Genes: gene_symbol=ZIC1

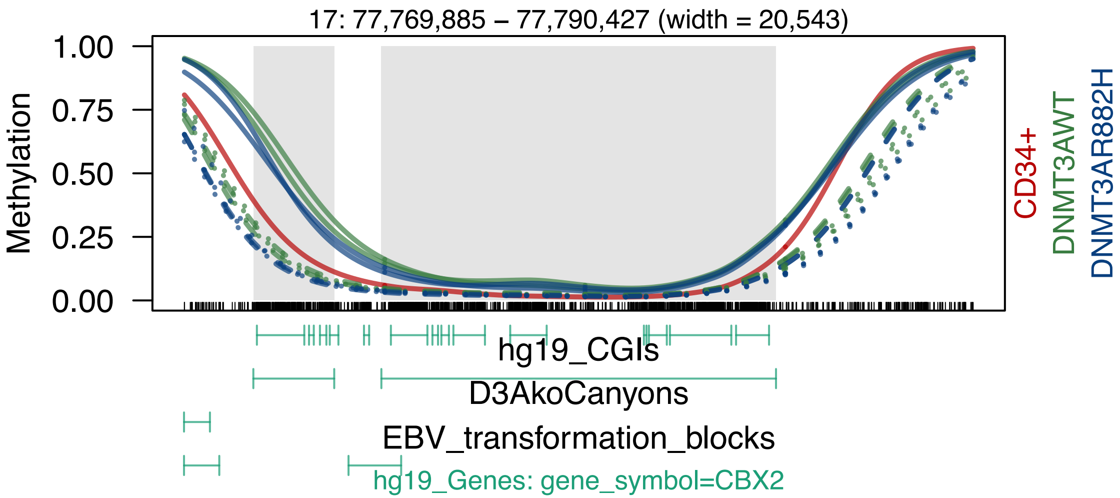
17: 77,769,885 – 77,790,427 (width = 20,543)
Methylation
1.00
0.75
0.50
0.25
0.00
CD34+
DNMT3AWT
DNMT3AR882H
hg19_CGIs
D3AkoCanyons
EBV_transformation_blocks
hg19_Genes: gene_symbol=CBX2

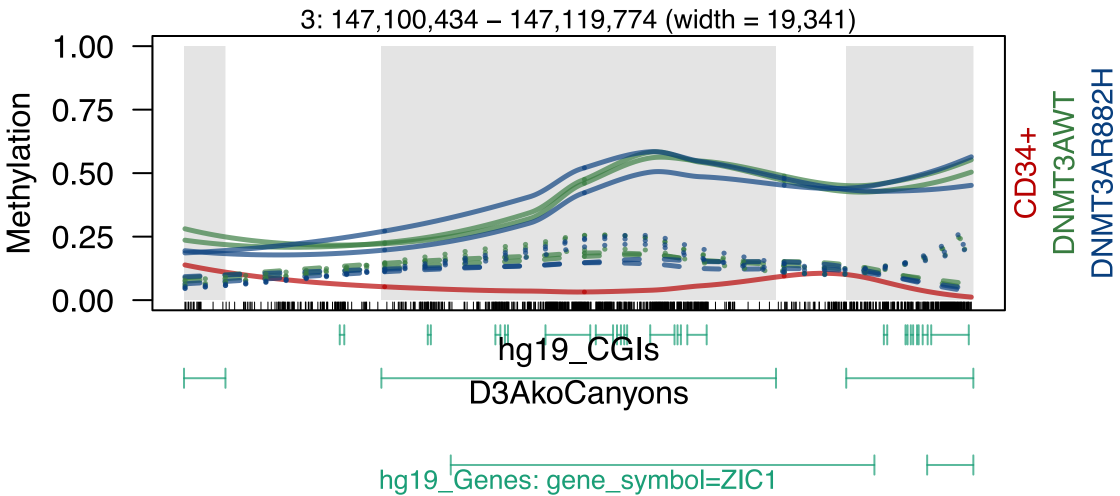
3: 147,100,434 – 147,119,774 (width = 19,341)
Methylation
1.00
0.75
0.50
0.25
0.00
CD34+
DNMT3AWT
DNMT3AR882H
hg19_CGIs
D3AkoCanyons
hg19_Genes: gene_symbol=ZIC1

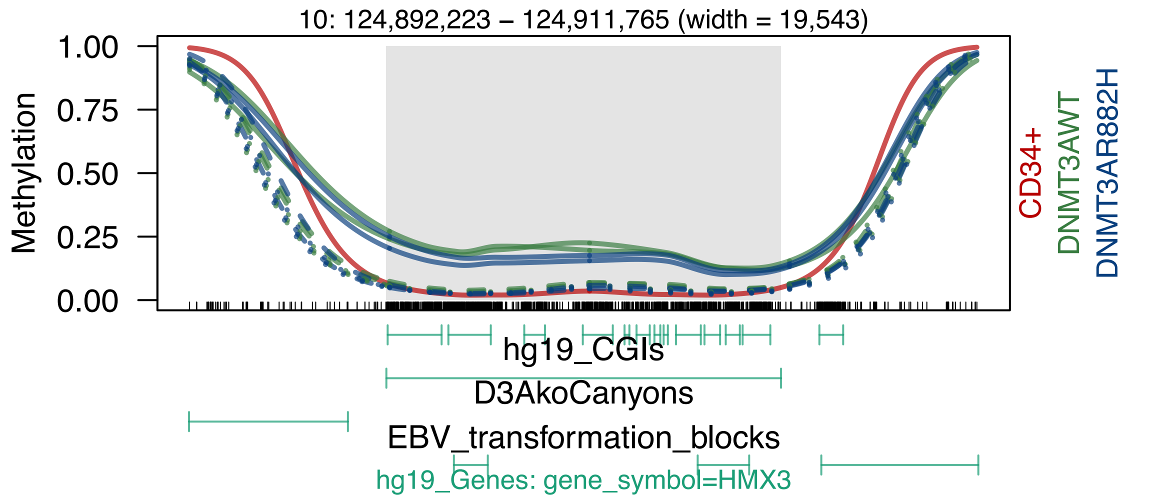
10: 124,892,223 – 124,911,765 (width = 19,543)
Methylation
1.00
0.75
0.50
0.25
0.00
CD34+
DNMT3AWT
DNMT3AR882H
hg19_CGIs
D3AkoCanyons
EBV_transformation_blocks
hg19_Genes: gene_symbol=HMX3

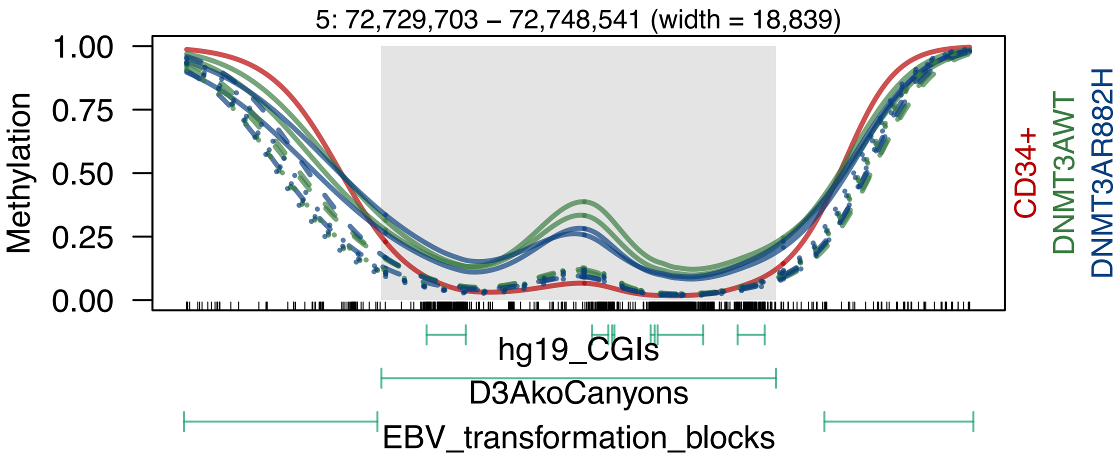

5: 72,729,703 – 72,748,541 (width = 18,839)
Methylation
1.00
0.75
0.50
0.25
0.00
CD34+
DNMT3AWT
DNMT3AR882H
hg19_CGIs
D3AkoCanyons
EBV_transformation_blocks

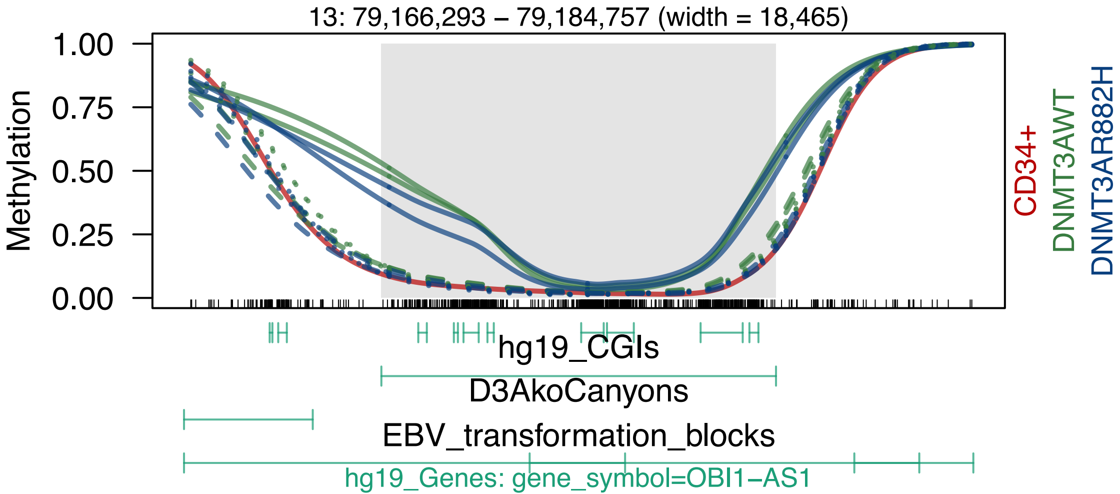
13: 79,166,293 – 79,184,757 (width = 18,465)
1.00
0.75
0.50
0.25
0.00
Methylation
CD34+
DNMT3AWT
DNMT3AR882H
hg19_CGIs
D3AkoCanyons
EBV_transformation_blocks
hg19_Genes: gene_symbol=OBI1−AS1

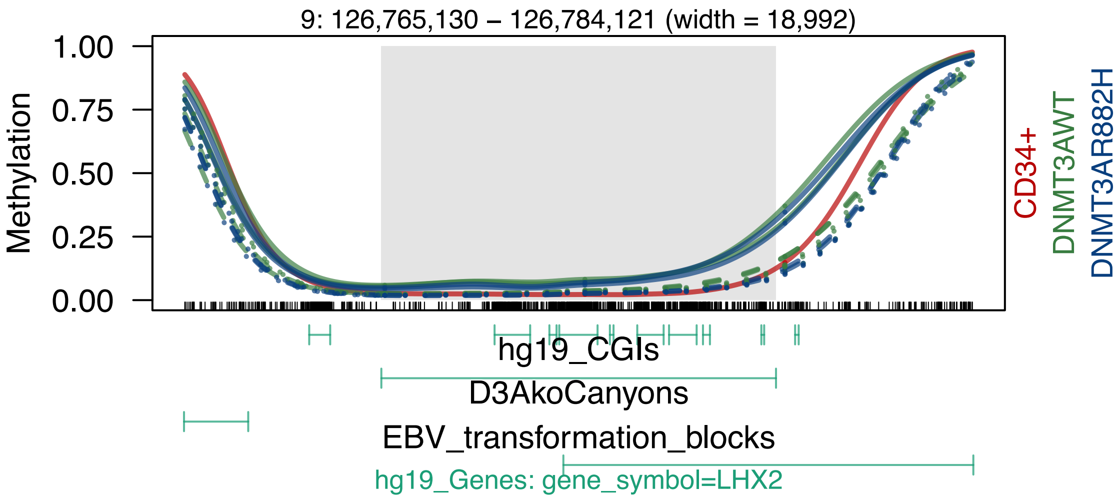
9: 126,765,130 – 126,784,121 (width = 18,992)
Methylation
1.00
0.75
0.50
0.25
0.00
CD34+
DNMT3AWT
DNMT3AR882H
hg19_CGIs
D3AkoCanyons
EBV_transformation_blocks
hg19_Genes: gene_symbol=LHX2

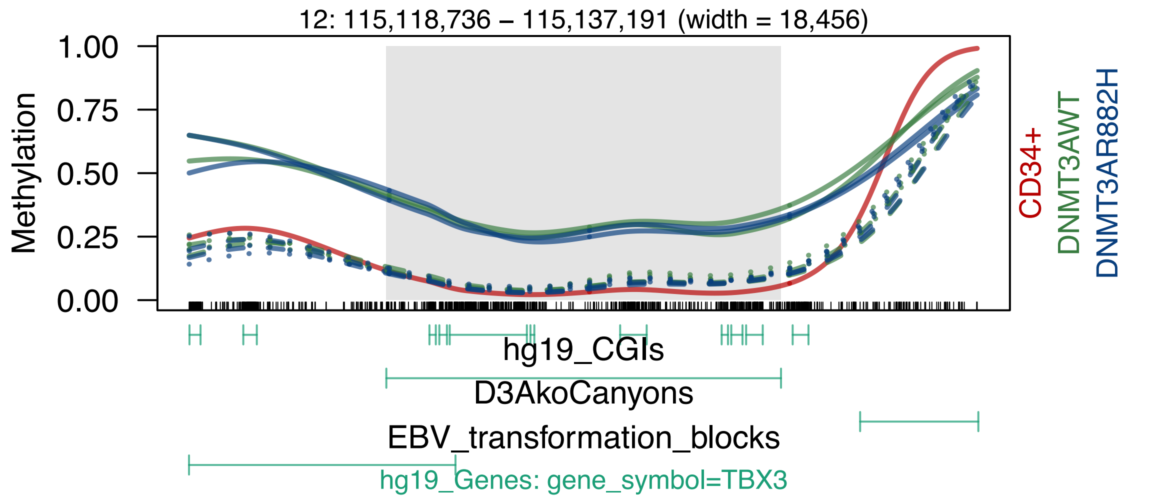
12: 115,118,736 – 115,137,191 (width = 18,456)
Methylation
1.00
0.75
0.50
0.25
0.00
CD34+
DNMT3AWT
DNMT3AR882H
hg19_CGIs
D3AkoCanyons
EBV_transformation_blocks
hg19_Genes: gene_symbol=TBX3

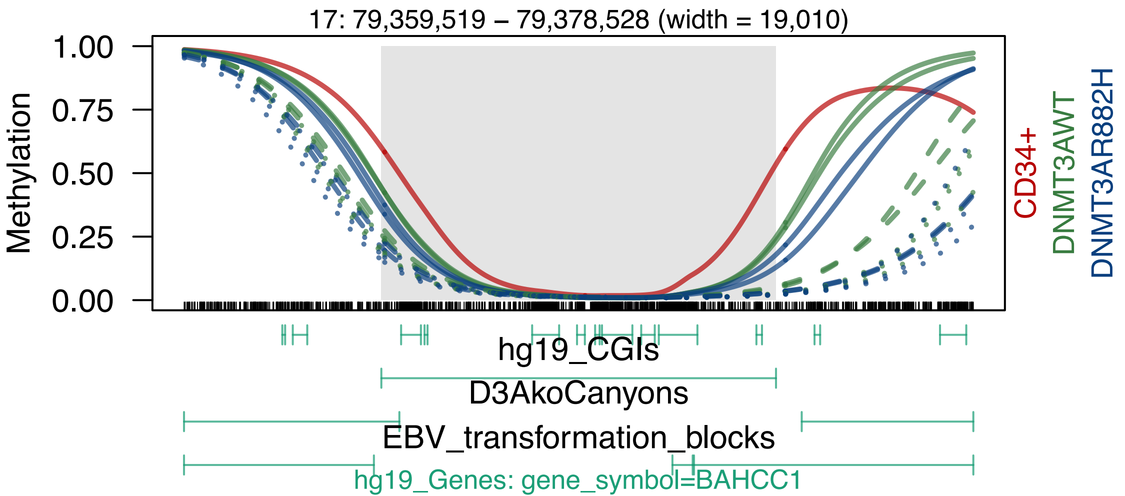
17: 79,359,519 – 79,378,528 (width = 19,010)
1.00
0.75
0.50
0.25
0.00
Methylation
CD34+
DNMT3AWT
DNMT3AR882H
hg19_CGIs
D3AkoCanyons
EBV_transformation_blocks
hg19_Genes: gene_symbol=BAHCC1

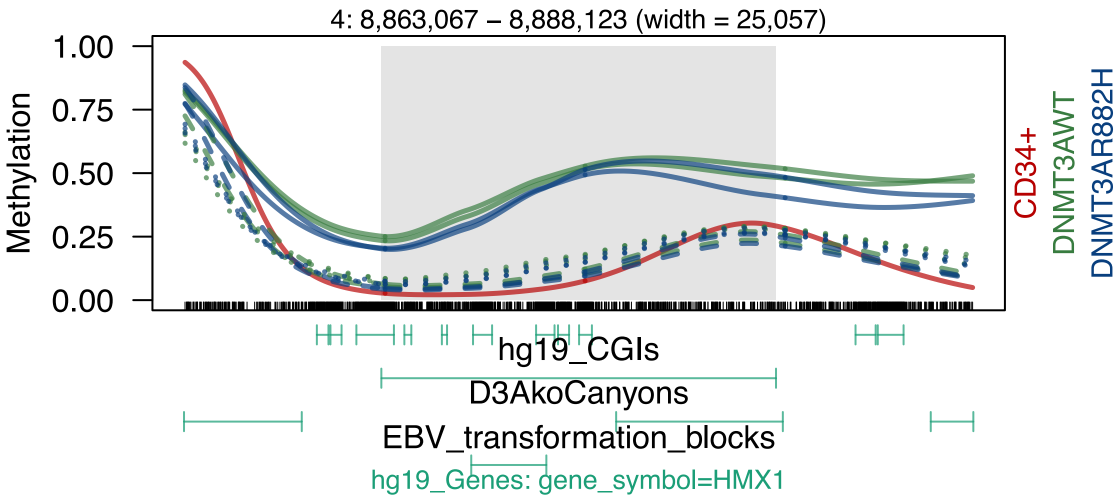

4: 8,863,067 – 8,888,123 (width = 25,057)
Methylation
1.00
0.75
0.50
0.25
0.00
CD34+
DNMT3AWT
DNMT3AR882H
hg19_CGIs
D3AkoCanyons
EBV_transformation_blocks
hg19_Genes: gene_symbol=HMX1

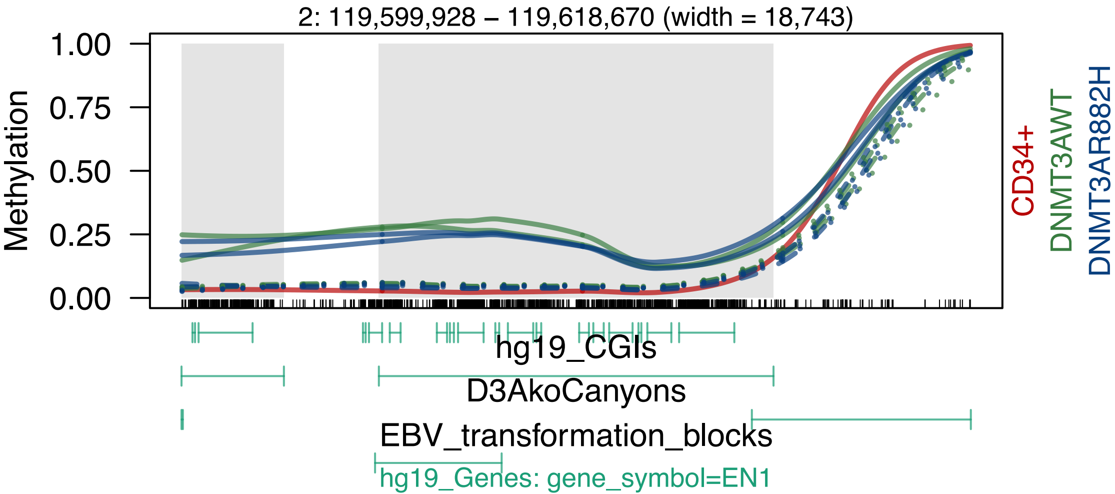

2: 119,599,928 – 119,618,670 (width = 18,743)
Methylation
1.00
0.75
0.50
0.25
0.00
CD34+
DNMT3AWT
DNMT3AR882H
hg19_CGIs
D3AkoCanyons
EBV_transformation_blocks
hg19_Genes: gene_symbol=EN1

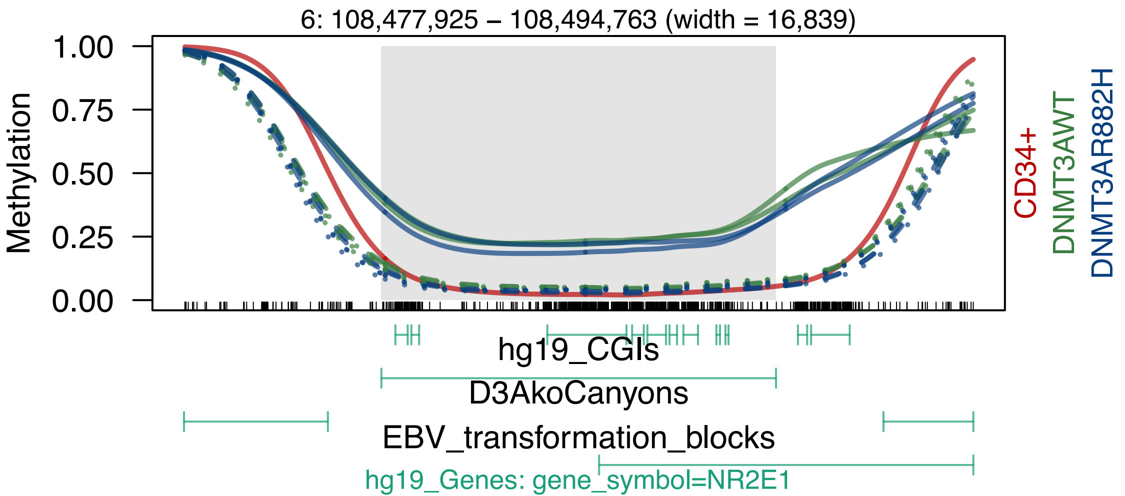
6: 108,477,925 – 108,494,763 (width = 16,839)
Methylation
1.00
0.75
0.50
0.25
0.00
CD34+
DNMT3AWT
DNMT3AR882H
hg19_CGIs
D3AkoCanyons
EBV_transformation_blocks
hg19_Genes: gene_symbol=NR2E1

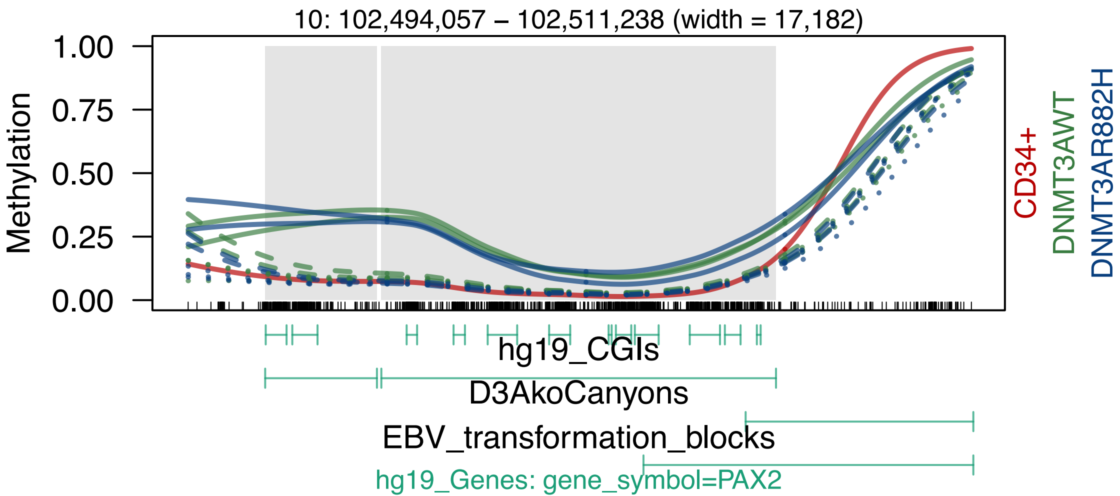

10: 102,494,057 – 102,511,238 (width = 17,182)
Methylation
1.00
0.75
0.50
0.25
0.00
CD34+
DNMT3AWT
DNMT3AR882H
hg19_CGIs
D3AkoCanyons
EBV_transformation_blocks
hg19_Genes: gene_symbol=PAX2

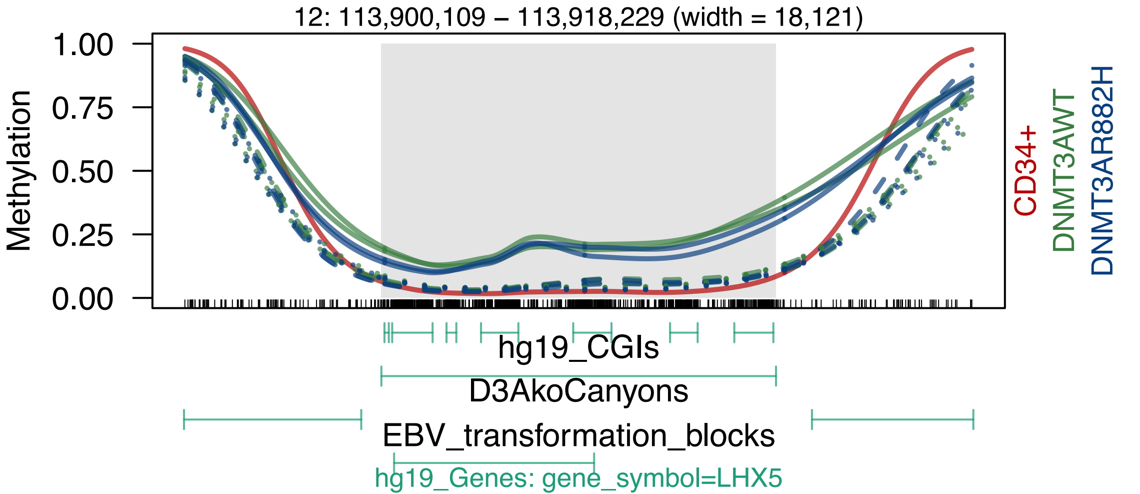
12: 113,900,109 – 113,918,229 (width = 18,121)
Methylation
1.00
0.75
0.50
0.25
0.00
CD34+
DNMT3AWT
DNMT3AR882H
hg19_CGIs
D3AkoCanyons
EBV_transformation_blocks
hg19_Genes: gene_symbol=LHX5

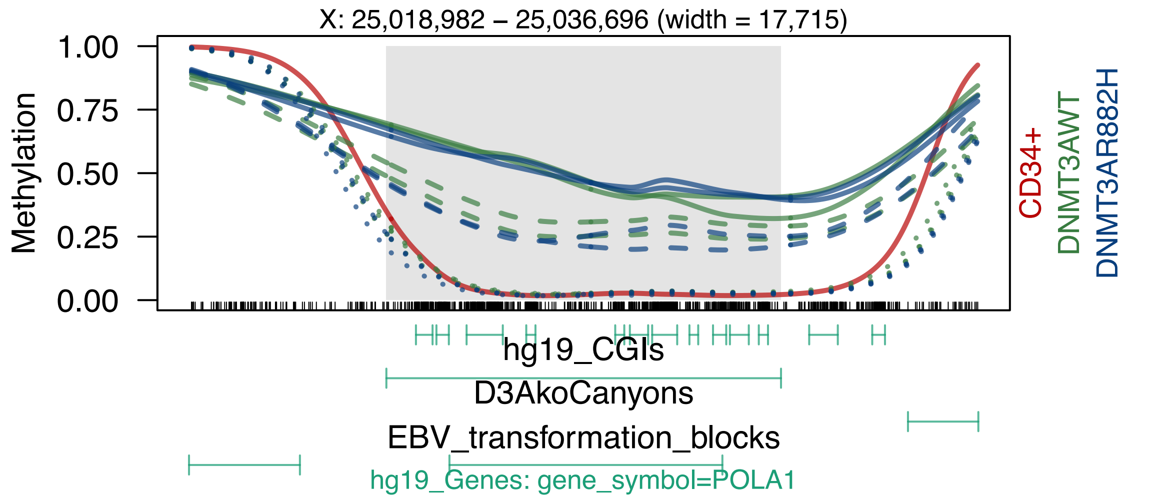

X: 25,018,982 – 25,036,696 (width = 17,715)
Methylation
1.00
0.75
0.50
0.25
0.00
CD34+
DNMT3AWT
DNMT3AR882H
hg19_CGIs
D3AkoCanyons
EBV_transformation_blocks
hg19_Genes: gene_symbol=POLA1

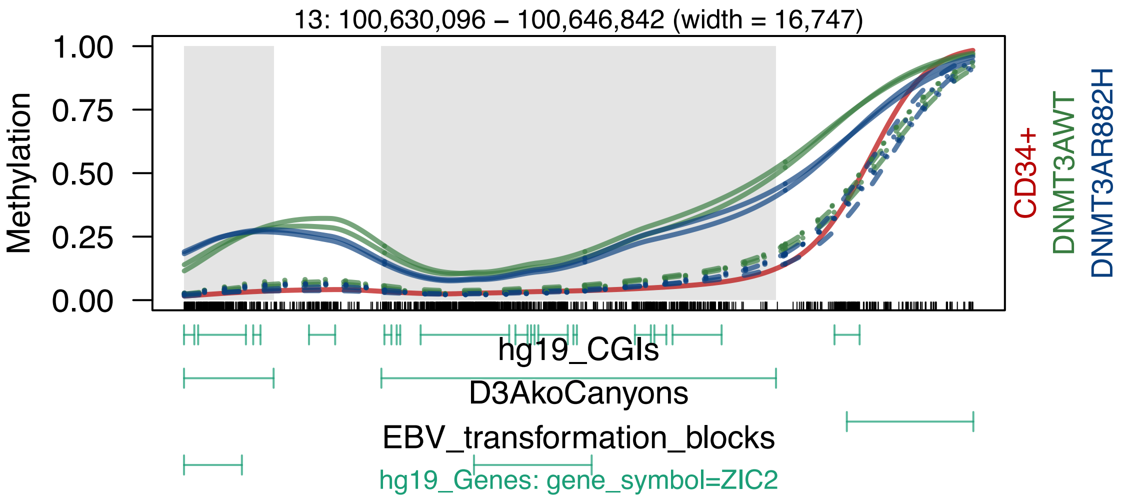

13: 100,630,096 – 100,646,842 (width = 16,747)
Methylation
1.00
0.75
0.50
0.25
0.00
CD34+
DNMT3AWT
DNMT3AR882H
hg19_CGIs
D3AkoCanyons
EBV_transformation_blocks
hg19_Genes: gene_symbol=ZIC2

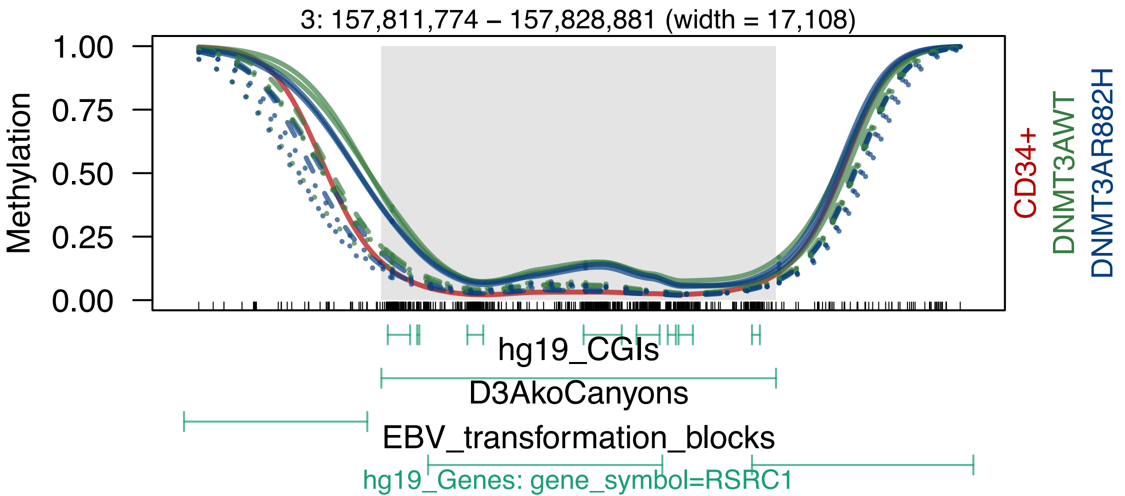
3: 157,811,774 – 157,828,881 (width = 17,108)
Methylation
1.00
0.75
0.50
0.25
0.00
CD34+
DNMT3AWT
DNMT3AR882H
hg19_CGIs
D3AkoCanyons
EBV_transformation_blocks
hg19_Genes: gene_symbol=RSRC1

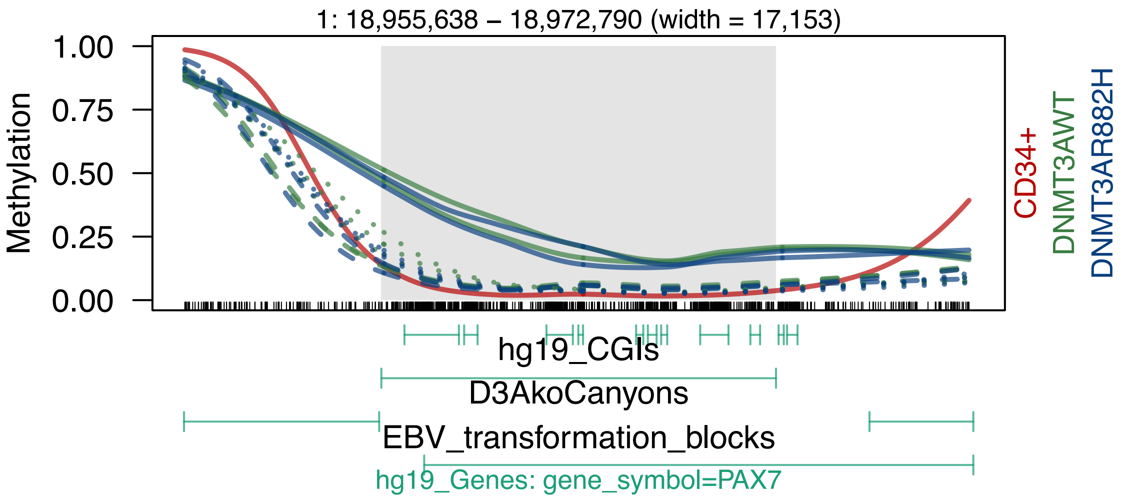
1: 18,955,638 – 18,972,790 (width = 17,153)
Methylation
1.00
0.75
0.50
0.25
0.00
CD34+
DNMT3AWT
DNMT3AR882H
hg19_CGIs
D3AkoCanyons
EBV_transformation_blocks
hg19_Genes: gene_symbol=PAX7

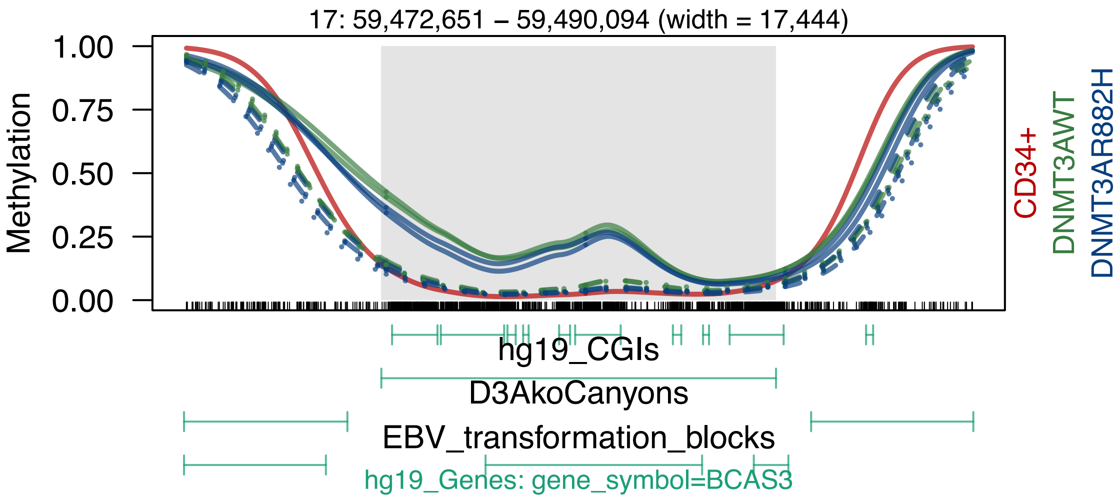
17: 59,472,651 – 59,490,094 (width = 17,444)
1.00
0.75
0.50
0.25
0.00
Methylation
CD34+
DNMT3AWT
DNMT3AR882H
hg19_CGIs
D3AkoCanyons
EBV_transformation_blocks
hg19_Genes: gene_symbol=BCAS3

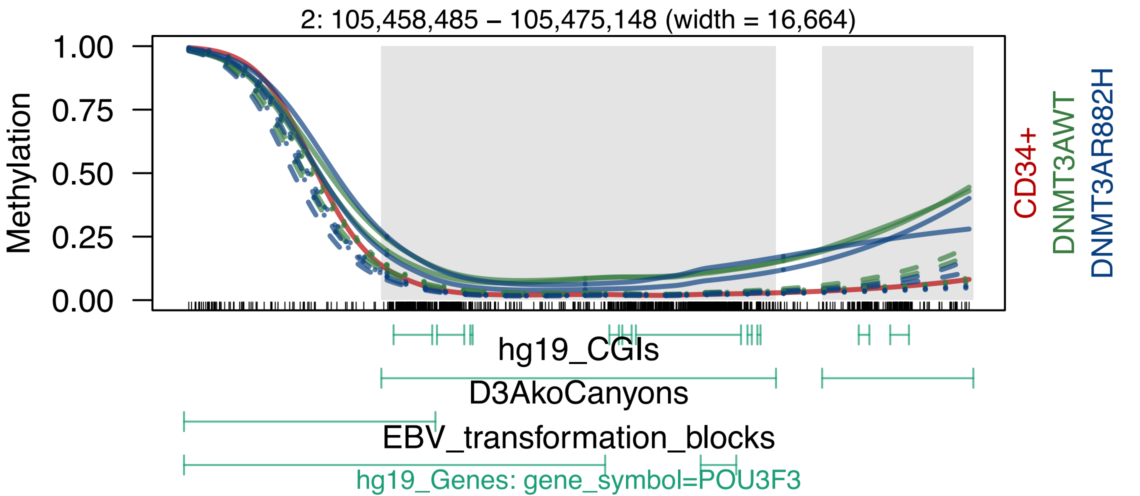

2: 105,458,485 – 105,475,148 (width = 16,664)
Methylation
1.00
0.75
0.50
0.25
0.00
CD34+
DNMT3AWT
DNMT3AR882H
hg19_CGIs
D3AkoCanyons
EBV_transformation_blocks
hg19_Genes: gene_symbol=POU3F3

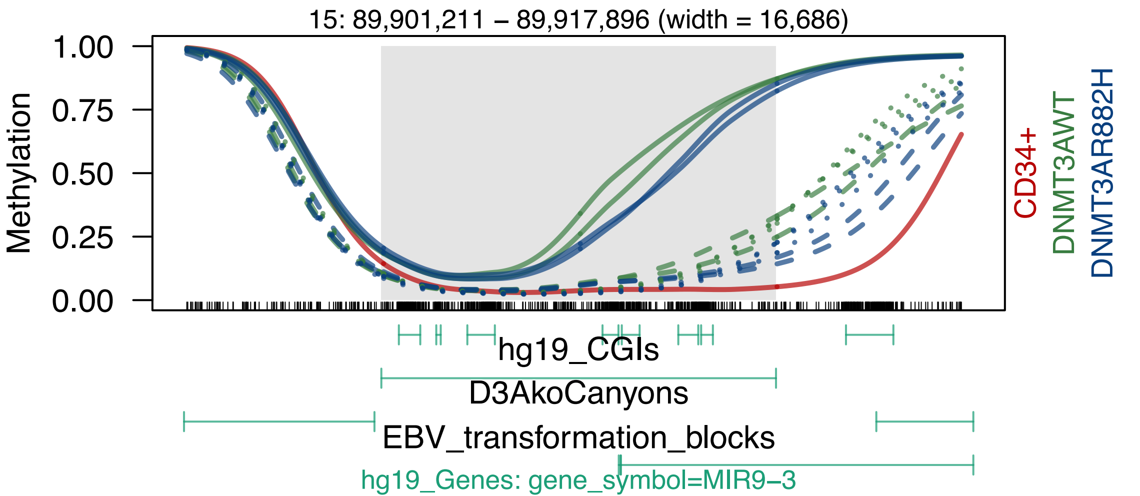

15: 89,901,211 – 89,917,896 (width = 16,686)
Methylation
1.00
0.75
0.50
0.25
0.00
CD34+
DNMT3AWT
DNMT3AR882H
hg19_CGIs
D3AkoCanyons
EBV_transformation_blocks
hg19_Genes: gene_symbol=MIR9–3

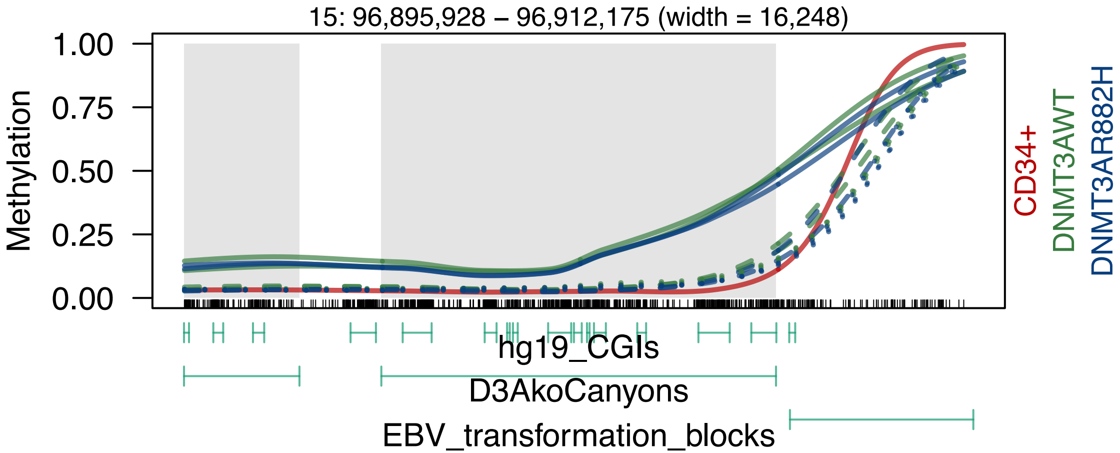

15: 96,895,928 – 96,912,175 (width = 16,248)
Methylation
1.00
0.75
0.50
0.25
0.00
CD34+
DNMT3AWT
DNMT3AR882H
hg19_CGIs
D3AkoCanyons
EBV_transformation_blocks

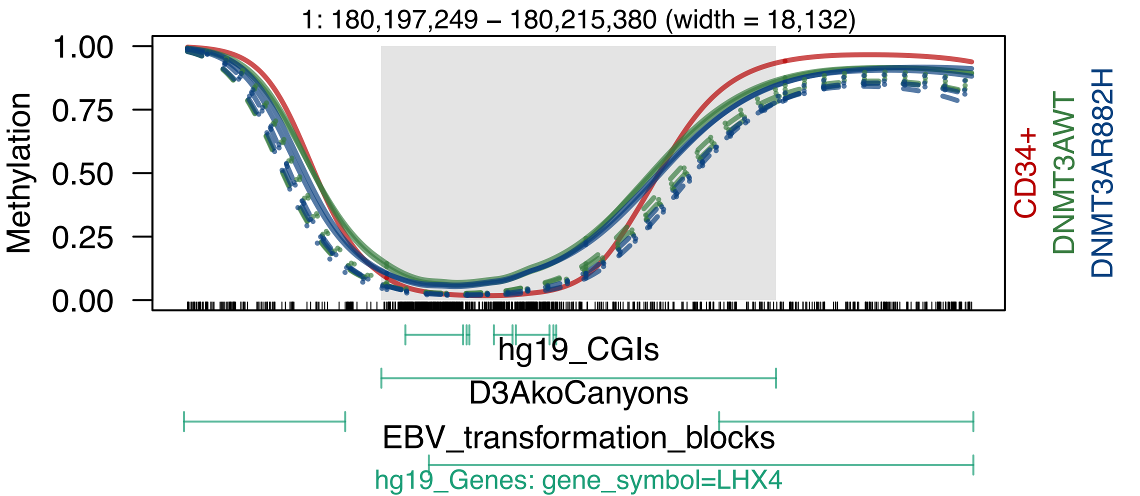
1: 180,197,249 – 180,215,380 (width = 18,132)
Methylation
1.00
0.75
0.50
0.25
0.00
CD34+
DNMT3AWT
DNMT3AR882H
hg19_CGIs
D3AkoCanyons
EBV_transformation_blocks
hg19_Genes: gene_symbol=LHX4

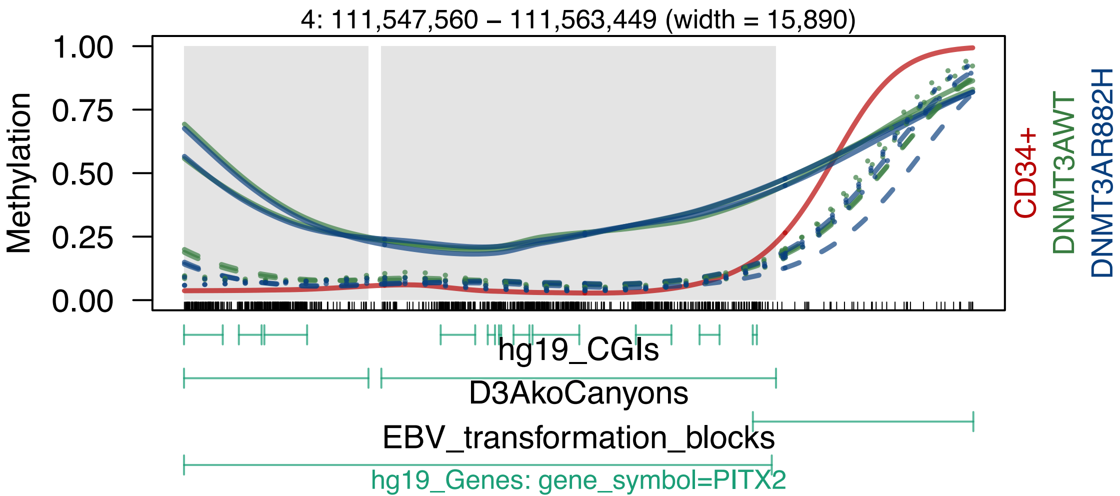

4: 111,547,560 – 111,563,449 (width = 15,890)
Methylation
1.00
0.75
0.50
0.25
0.00
CD34+
DNMT3AWT
DNMT3AR882H
hg19_CGIs
D3AkoCanyons
EBV_transformation_blocks
hg19_Genes: gene_symbol=PITX2

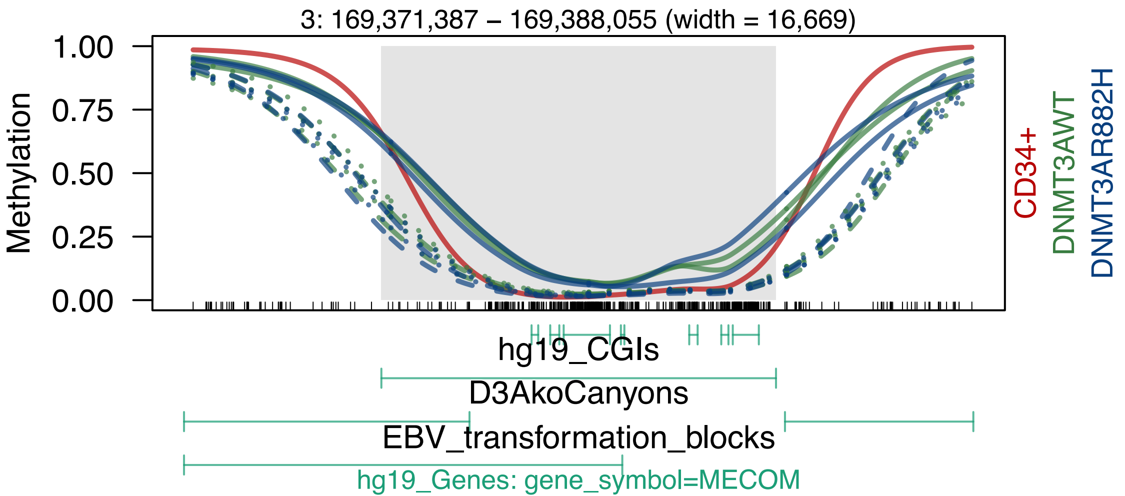
3: 169,371,387 – 169,388,055 (width = 16,669)
Methylation
1.00
0.75
0.50
0.25
0.00
CD34+
DNMT3AWT
DNMT3AR882H
hg19_CGIs
D3AkoCanyons
EBV_transformation_blocks
hg19_Genes: gene_symbol=MECOM

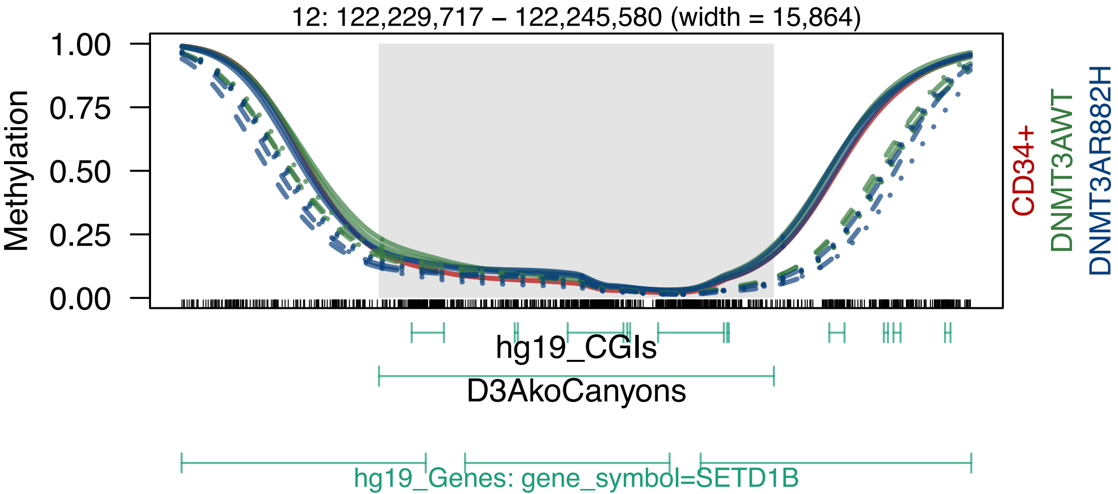
12: 122,229,717 – 122,245,580 (width = 15,864)
Methylation
1.00
0.75
0.50
0.25
0.00
CD34+
DNMT3AWT
DNMT3AR882H
hg19_CGIs
D3AkoCanyons
hg19_Genes: gene_symbol=SETD1B

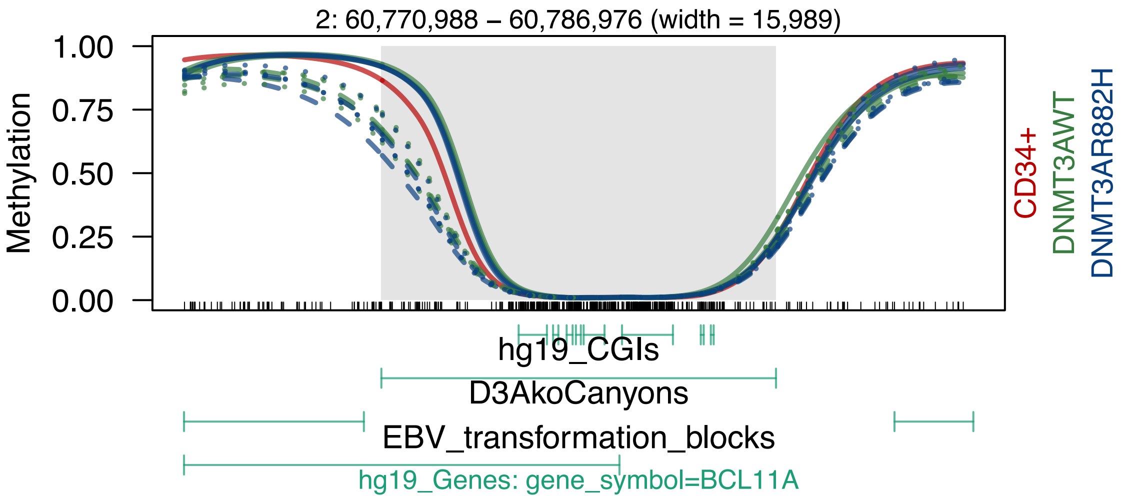
2: 60,770,988 – 60,786,976 (width = 15,989)
1.00
0.75
0.50
0.25
0.00
Methylation
CD34+
DNMT3AWT
DNMT3AR882H
hg19_CGIs
D3AkoCanyons
EBV_transformation_blocks
hg19_Genes: gene_symbol=BCL11A

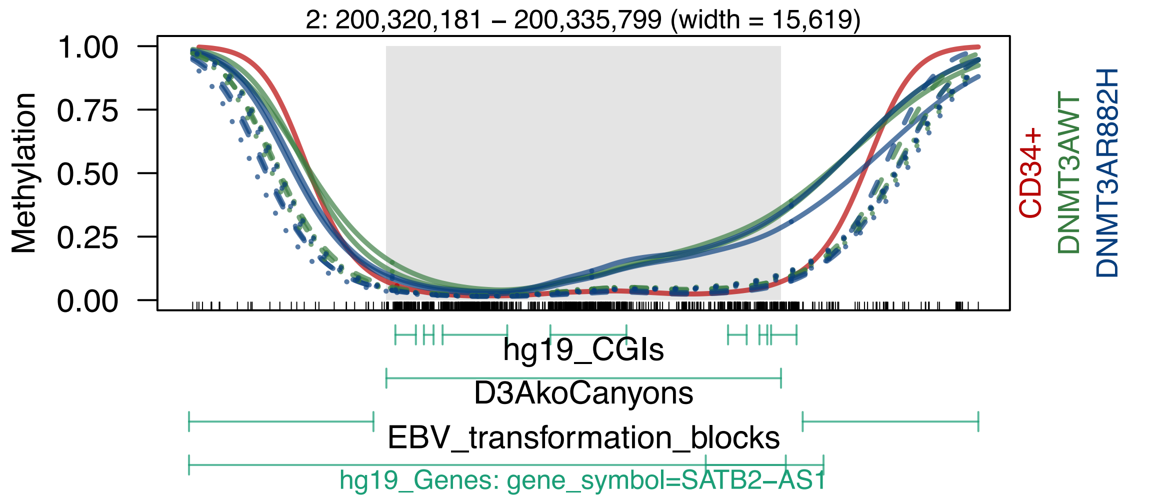

2: 200,320,181 – 200,335,799 (width = 15,619)
Methylation
1.00
0.75
0.50
0.25
0.00
CD34+
DNMT3AWT
DNMT3AR882H
hg19_CGIs
D3AkoCanyons
EBV_transformation_blocks
hg19_Genes: gene_symbol=SATB2−AS1

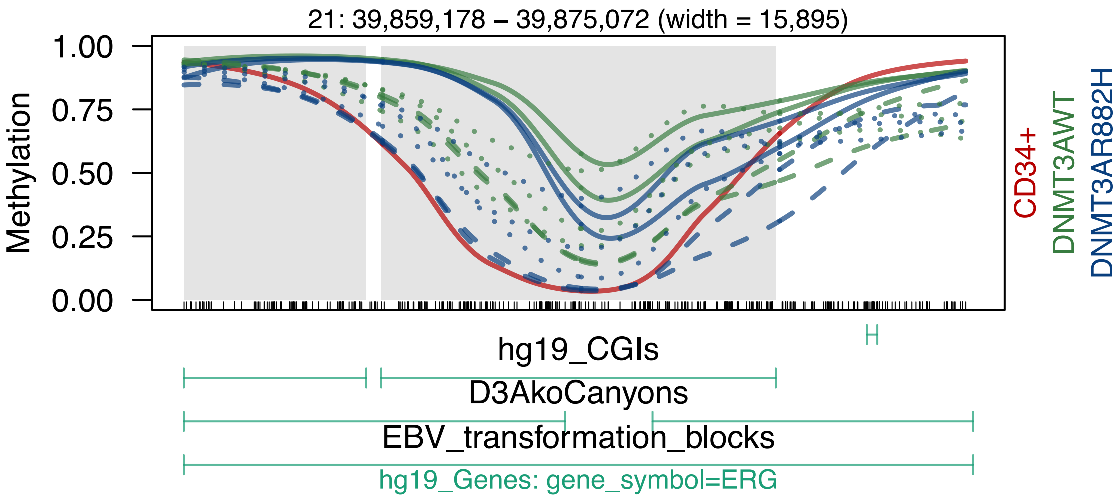
21: 39,859,178 – 39,875,072 (width = 15,895)
Methylation
1.00
0.75
0.50
0.25
0.00
CD34+
DNMT3AWT
DNMT3AR882H
hg19_CGIs
D3AkoCanyons
EBV_transformation_blocks
hg19_Genes: gene_symbol=ERG

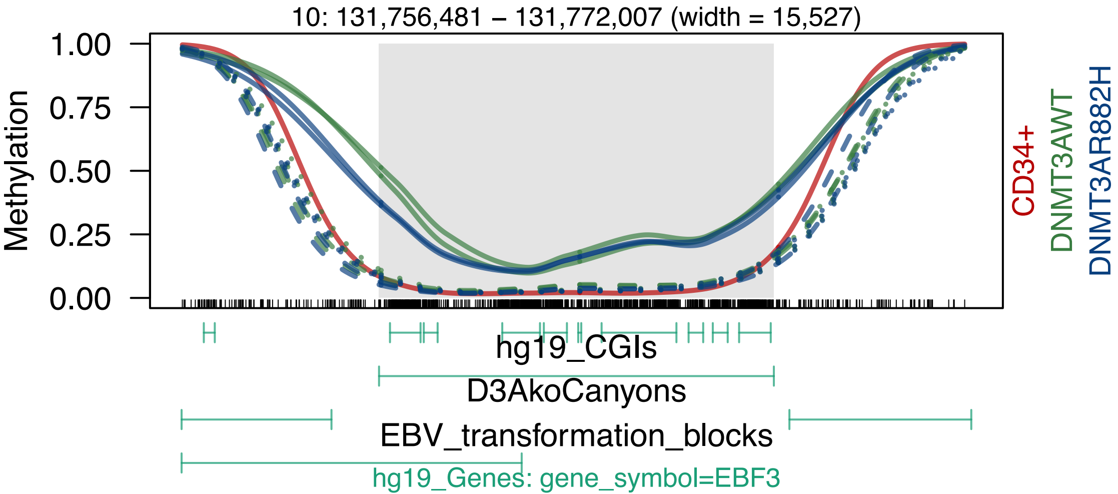
10: 131,756,481 – 131,772,007 (width = 15,527)
Methylation
1.00
0.75
0.50
0.25
0.00
CD34+
DNMT3AWT
DNMT3AR882H
hg19_CGIs
D3AkoCanyons
EBV_transformation_blocks
hg19_Genes: gene_symbol=EBF3

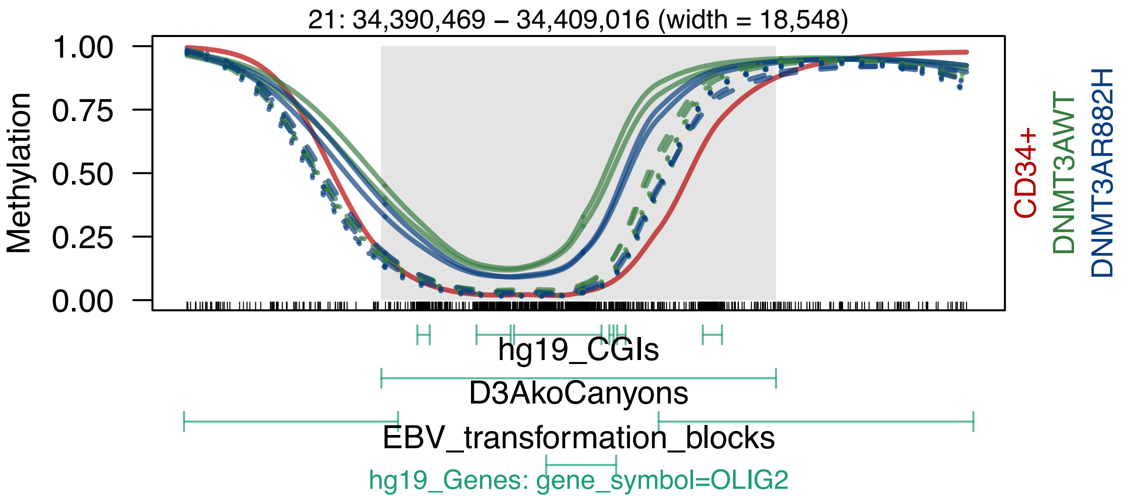
21: 34,390,469 – 34,409,016 (width = 18,548)
Methylation
1.00
0.75
0.50
0.25
0.00
CD34+
DNMT3AWT
DNMT3AR882H
hg19_CGIs
D3AkoCanyons
EBV_transformation_blocks
hg19_Genes: gene_symbol=OLIG2

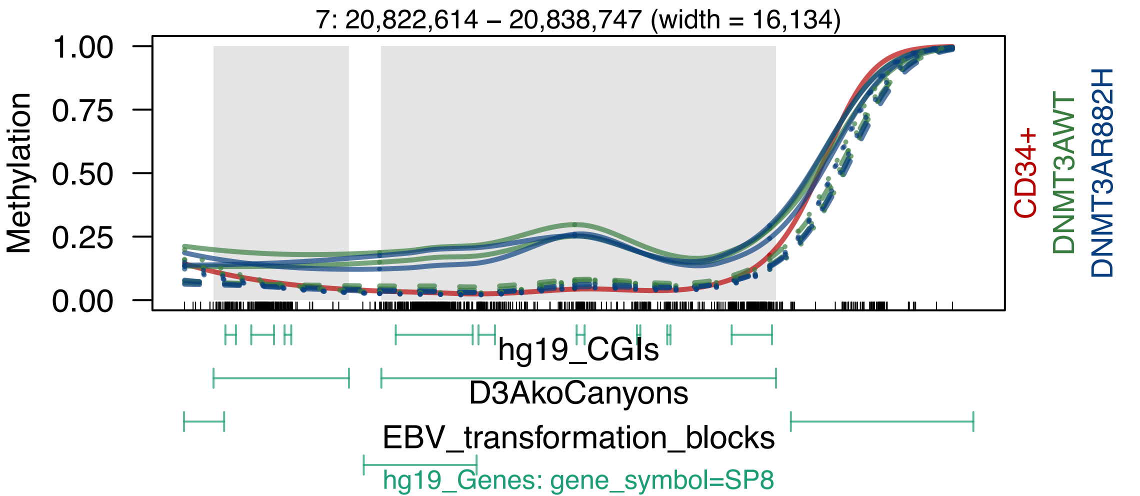

7: 20,822,614 – 20,838,747 (width = 16,134)
Methylation
1.00
0.75
0.50
0.25
0.00
CD34+
DNMT3AWT
DNMT3AR882H
hg19_CGIs
D3AkoCanyons
EBV_transformation_blocks
hg19_Genes: gene_symbol=SP8

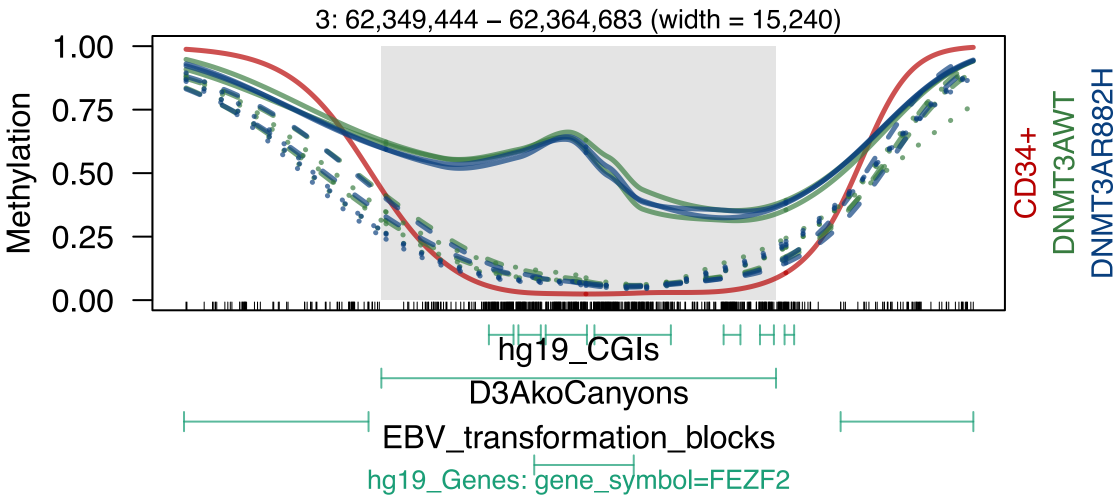

3: 62,349,444 – 62,364,683 (width = 15,240)
Methylation
1.00
0.75
0.50
0.25
0.00
CD34+
DNMT3AWT
DNMT3AR882H
hg19_CGIs
D3AkoCanyons
EBV_transformation_blocks
hg19_Genes: gene_symbol=FEZF2

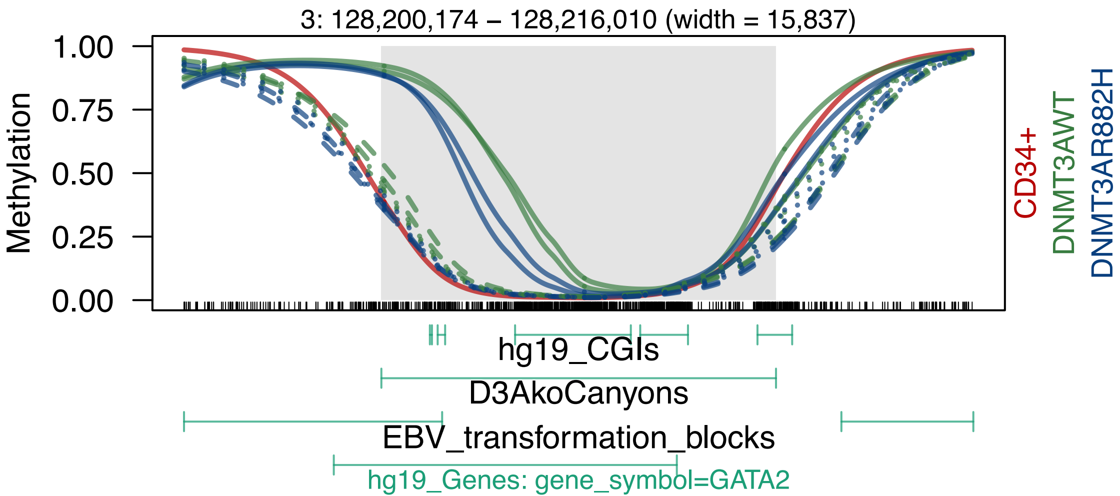

3: 128,200,174 – 128,216,010 (width = 15,837)
Methylation
1.00
0.75
0.50
0.25
0.00
CD34+
DNMT3AWT
DNMT3AR882H
hg19_CGIs
D3AkoCanyons
EBV_transformation_blocks
hg19_Genes: gene_symbol=GATA2

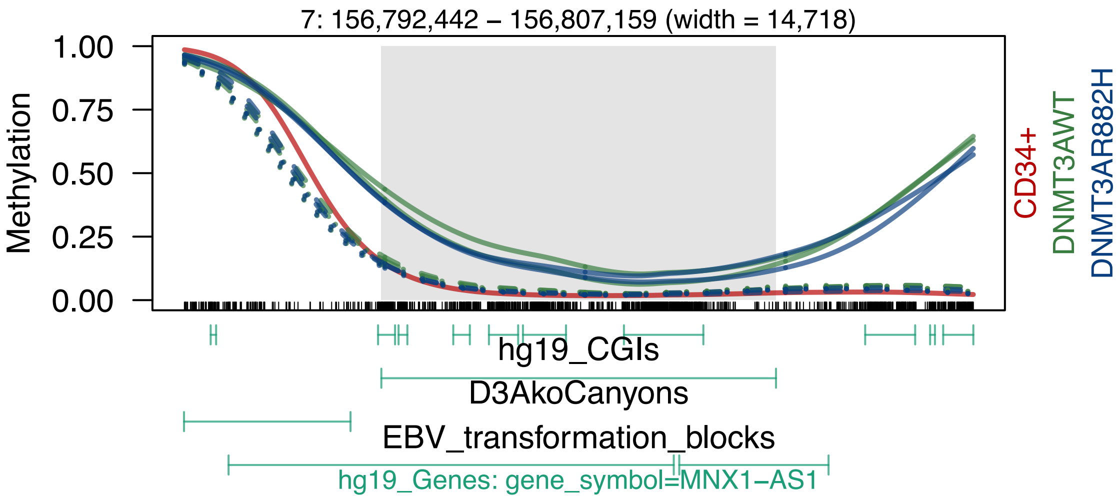

7: 156,792,442 – 156,807,159 (width = 14,718)
Methylation
1.00
0.75
0.50
0.25
0.00
CD34+
DNMT3AWT
DNMT3AR882H
hg19_CGIs
D3AkoCanyons
EBV_transformation_blocks
hg19_Genes: gene_symbol=MNX1−AS1

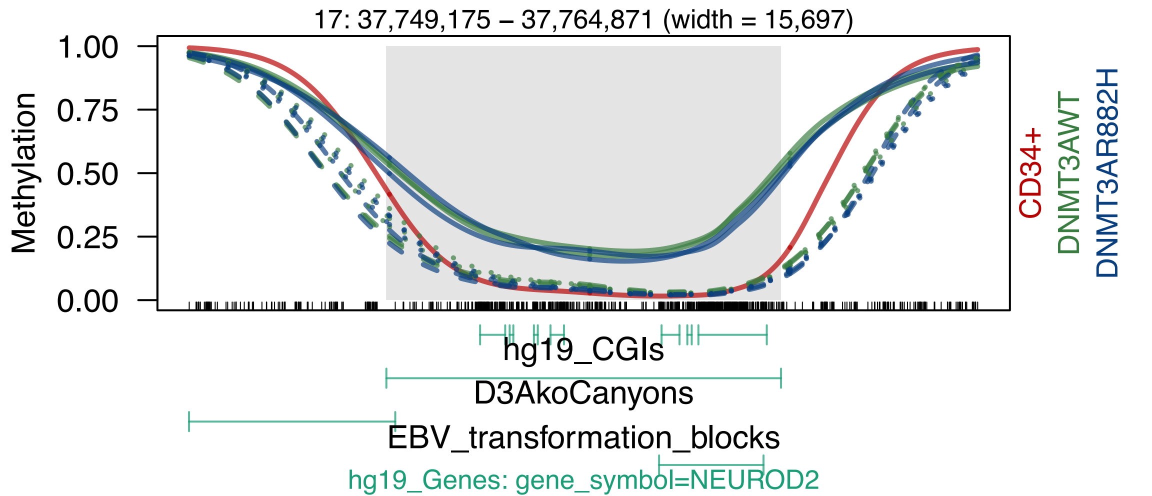

17: 37,749,175 – 37,764,871 (width = 15,697)
Methylation
1.00
0.75
0.50
0.25
0.00
CD34+
DNMT3AWT
DNMT3AR882H
hg19_CGIs
D3AkoCanyons
EBV_transformation_blocks
hg19_Genes: gene_symbol=NEUROD2

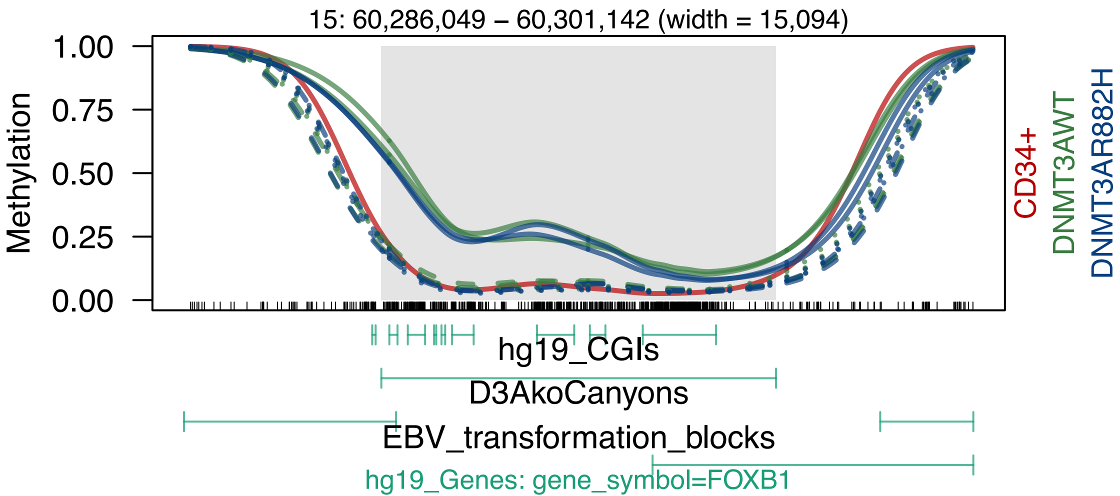

15: 60,286,049 – 60,301,142 (width = 15,094)
Methylation
1.00
0.75
0.50
0.25
0.00
CD34+
DNMT3AWT
DNMT3AR882H
hg19_CGIs
D3AkoCanyons
EBV_transformation_blocks
hg19_Genes: gene_symbol=FOXB1

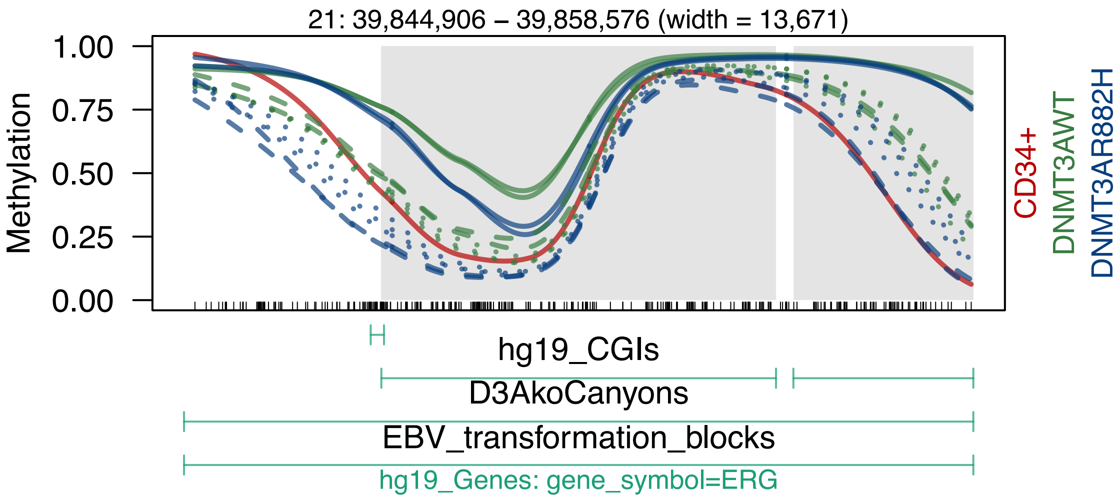

21: 39,844,906 – 39,858,576 (width = 13,671)
Methylation
1.00
0.75
0.50
0.25
0.00
CD34+
DNMT3AWT
DNMT3AR882H
hg19_CGIs
D3AkoCanyons
EBV_transformation_blocks
hg19_Genes: gene_symbol=ERG

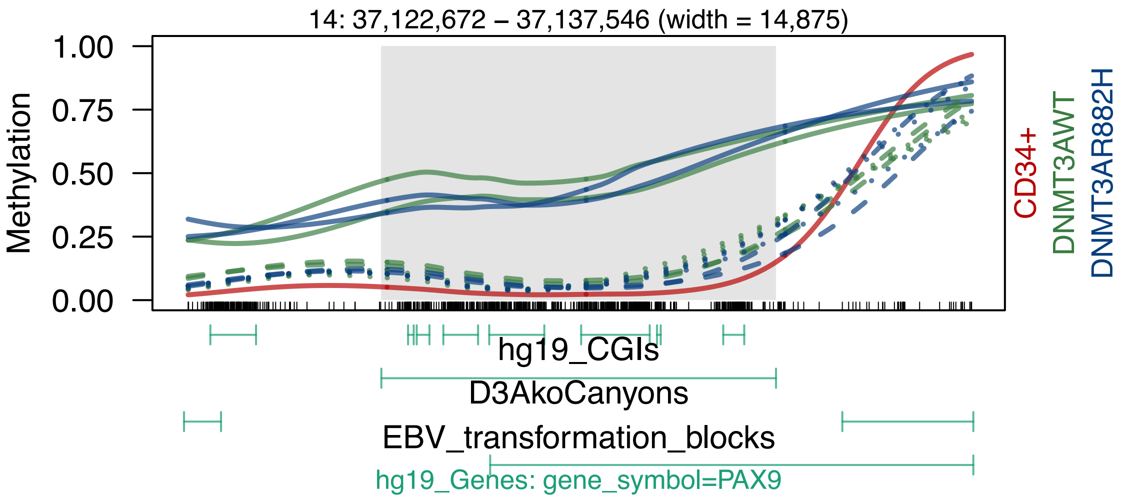
14: 37,122,672 – 37,137,546 (width = 14,875)
Methylation
1.00
0.75
0.50
0.25
0.00
CD34+
DNMT3AWT
DNMT3AR882H
hg19_CGIs
D3AkoCanyons
EBV_transformation_blocks
hg19_Genes: gene_symbol=PAX9

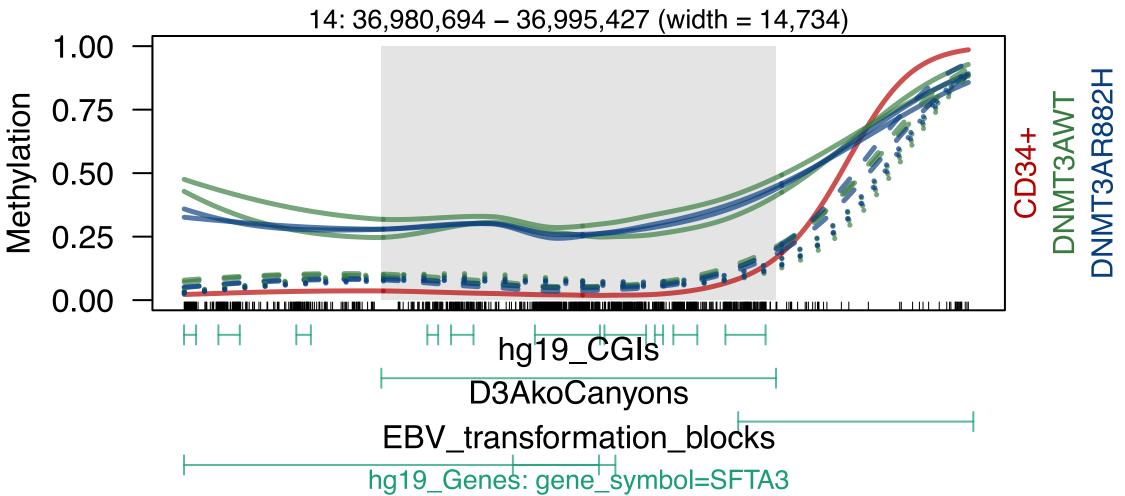

14: 36,980,694 – 36,995,427 (width = 14,734)
Methylation
1.00
0.75
0.50
0.25
0.00
CD34+
DNMT3AWT
DNMT3AR882H
hg19_CGIs
D3AkoCanyons
EBV_transformation_blocks
hg19_Genes: gene_symbol=SFTA3

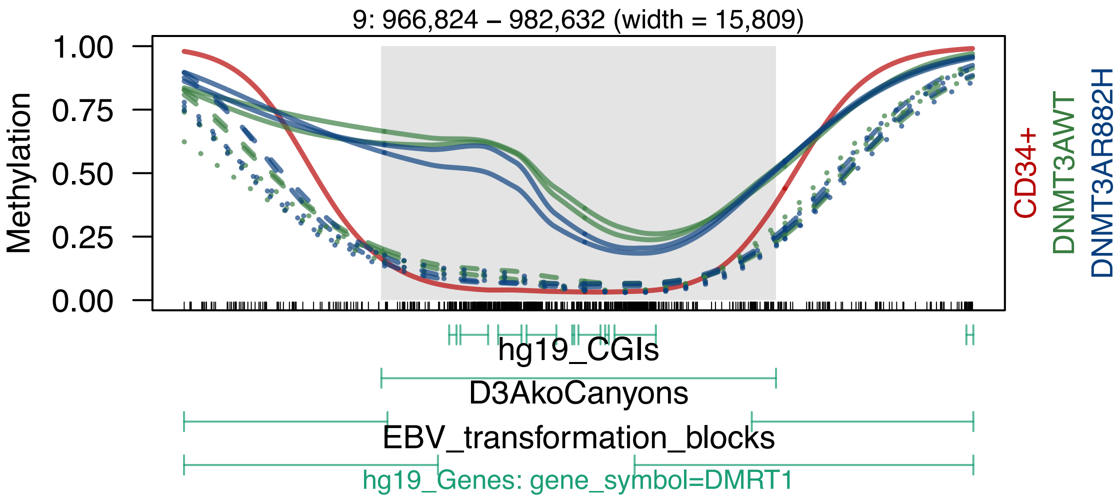

9: 966,824 – 982,632 (width = 15,809)
1.00
0.75
0.50
0.25
0.00
Methylation
CD34+
DNMT3AWT
DNMT3AR882H
hg19_CGIs
D3AkoCanyons
EBV_transformation_blocks
hg19_Genes: gene_symbol=DMRT1

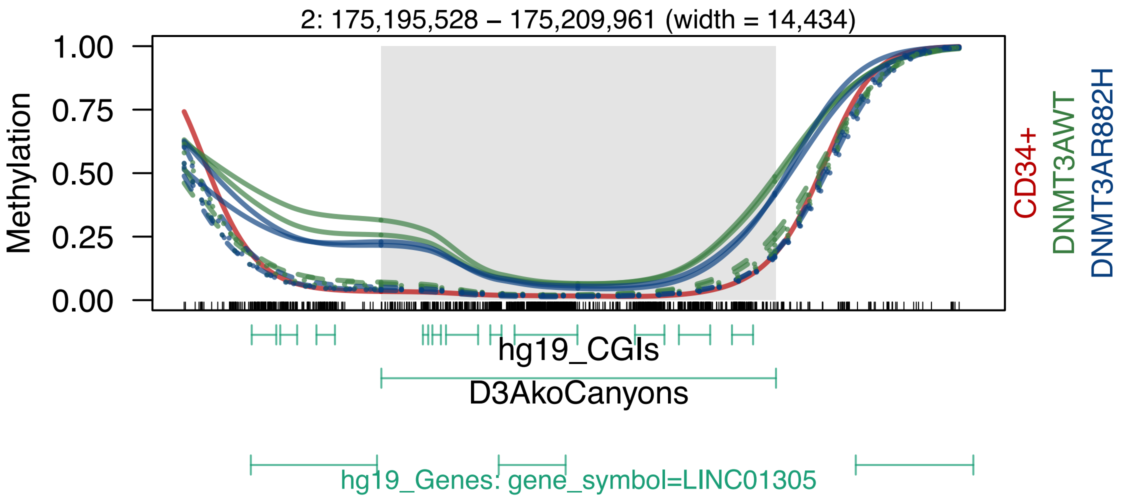
2: 175,195,528 – 175,209,961 (width = 14,434)
Methylation
1.00
0.75
0.50
0.25
0.00
CD34+
DNMT3AWT
DNMT3AR882H
hg19_CGIs
D3AkoCanyons
hg19_Genes: gene_symbol=LINC01305

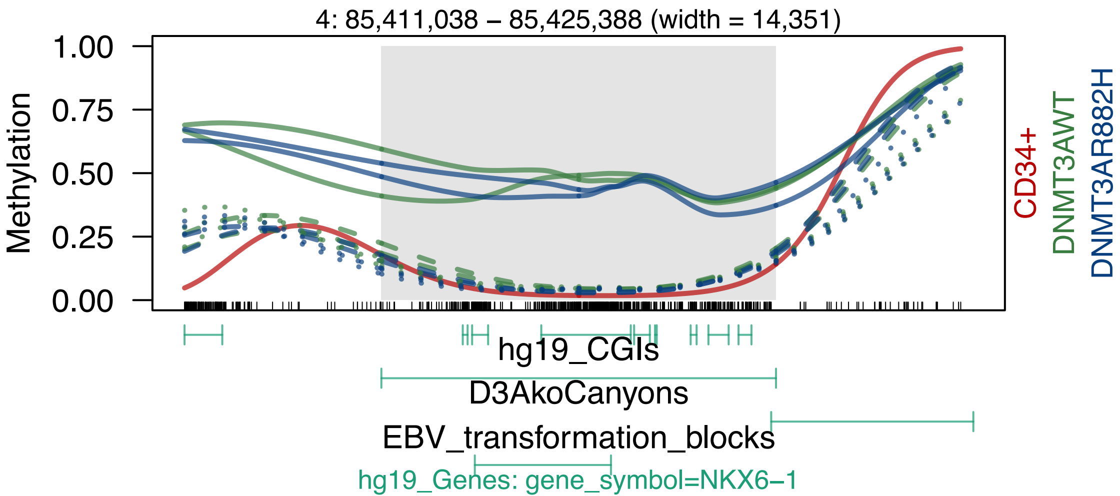
4: 85,411,038 – 85,425,388 (width = 14,351)
Methylation
1.00
0.75
0.50
0.25
0.00
CD34+
DNMT3AWT
DNMT3AR882H
hg19_CGIs
D3AkoCanyons
EBV_transformation_blocks
hg19_Genes: gene_symbol=NKX6−1

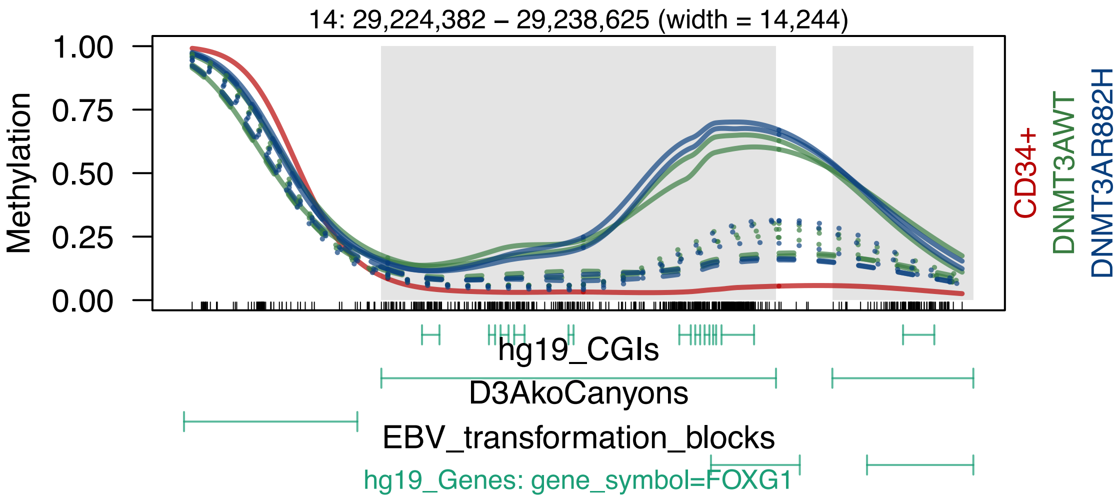

14: 29,224,382 – 29,238,625 (width = 14,244)
Methylation
1.00
0.75
0.50
0.25
0.00
CD34+
DNMT3AWT
DNMT3AR882H
hg19_CGIs
D3AkoCanyons
EBV_transformation_blocks
hg19_Genes: gene_symbol=FOXG1

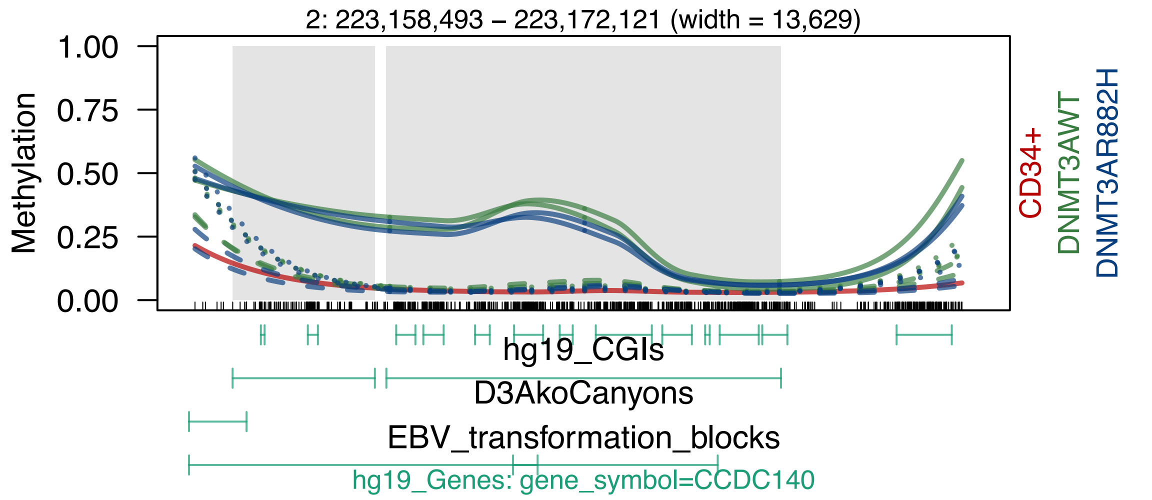

2: 223,158,493 – 223,172,121 (width = 13,629)
Methylation
1.00
0.75
0.50
0.25
0.00
CD34+
DNMT3AWT
DNMT3AR882H
hg19_CGIs
D3AkoCanyons
EBV_transformation_blocks
hg19_Genes: gene_symbol=CCDC140

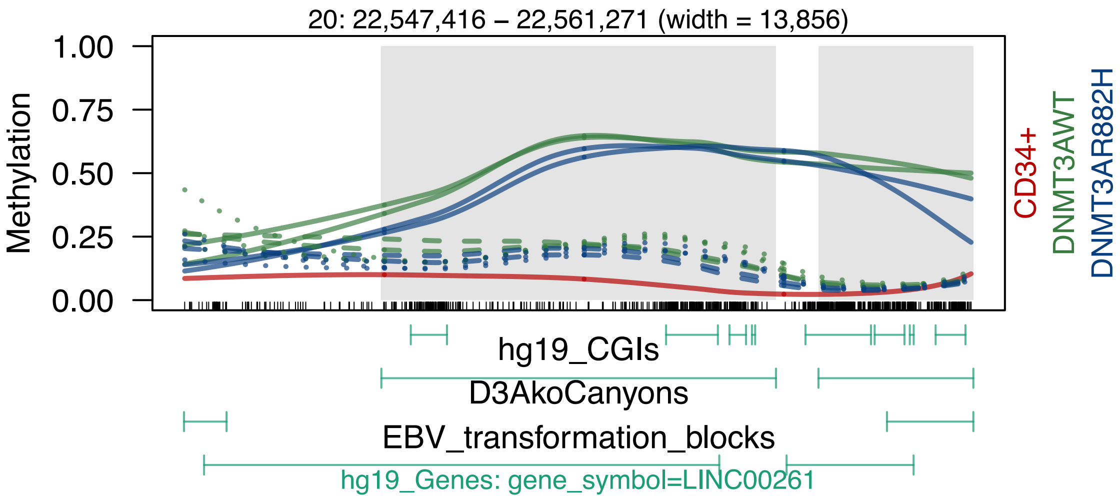
20: 22,547,416 – 22,561,271 (width = 13,856)
Methylation
1.00
0.75
0.50
0.25
0.00
CD34+
DNMT3AWT
DNMT3AR882H
hg19_CGIs
D3AkoCanyons
EBV_transformation_blocks
hg19_Genes: gene_symbol=LINC00261

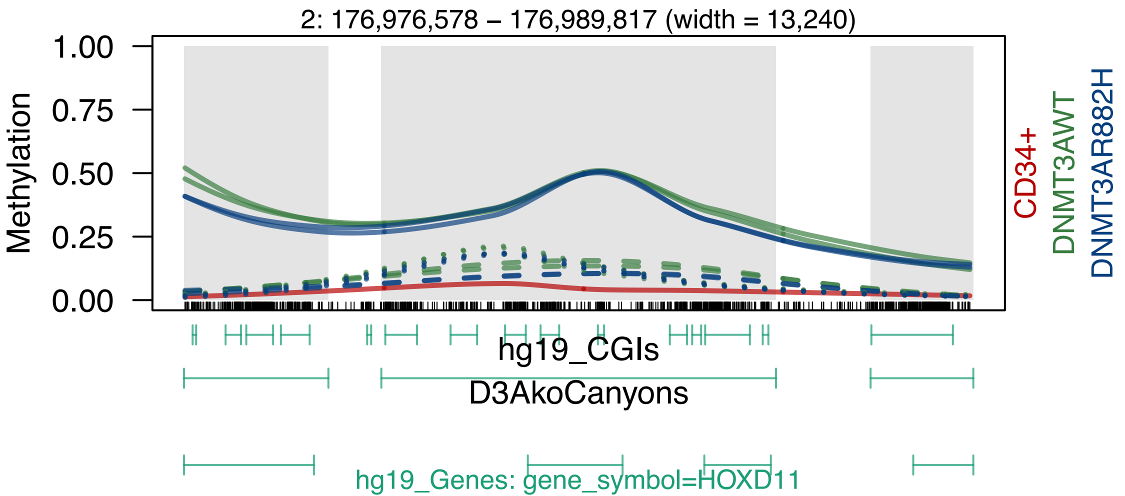
2: 176,976,578 – 176,989,817 (width = 13,240)
Methylation
1.00
0.75
0.50
0.25
0.00
CD34+
DNMT3AWT
DNMT3AR882H
hg19_CGIs
D3AkoCanyons
hg19_Genes: gene_symbol=HOXD11

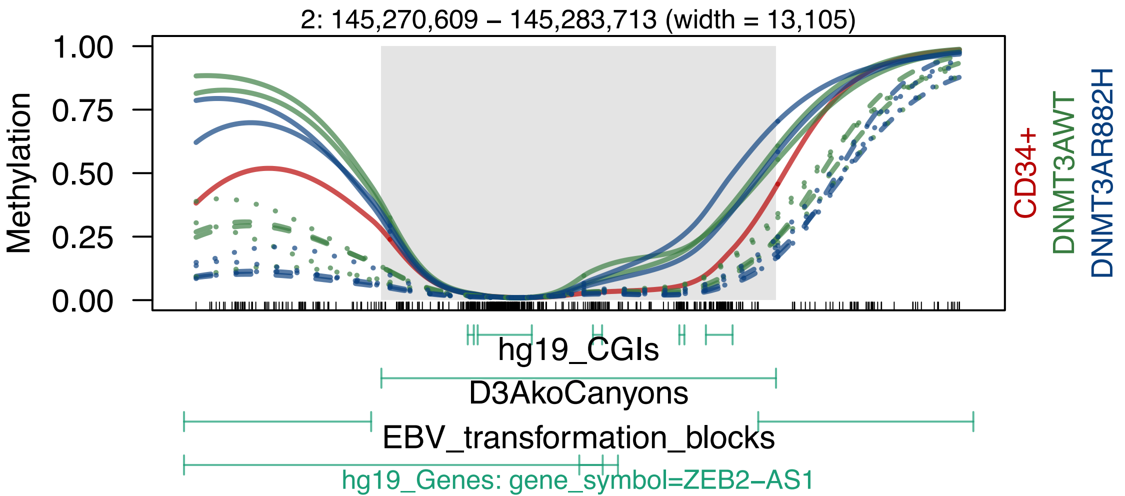
2: 145,270,609 – 145,283,713 (width = 13,105)
Methylation
1.00
0.75
0.50
0.25
0.00
CD34+
DNMT3AWT
DNMT3AR882H
hg19_CGIs
D3AkoCanyons
EBV_transformation_blocks
hg19_Genes: gene_symbol=ZEB2–AS1

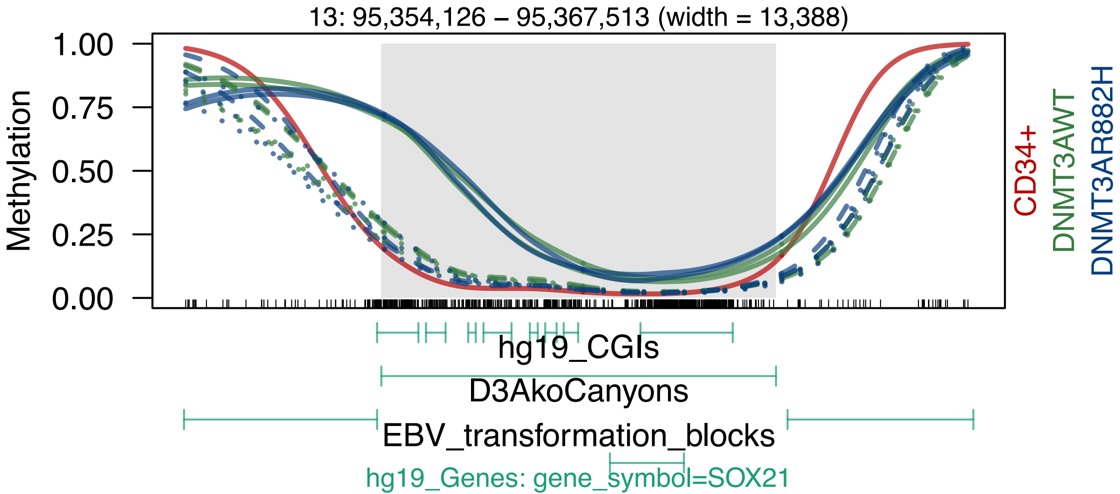

13: 95,354,126 – 95,367,513 (width = 13,388)
1.00
0.75
0.50
0.25
0.00
Methylation
CD34+
DNMT3AWT
DNMT3AR882H
hg19_CGIs
D3AkoCanyons
EBV_transformation_blocks
hg19_Genes: gene_symbol=SOX21

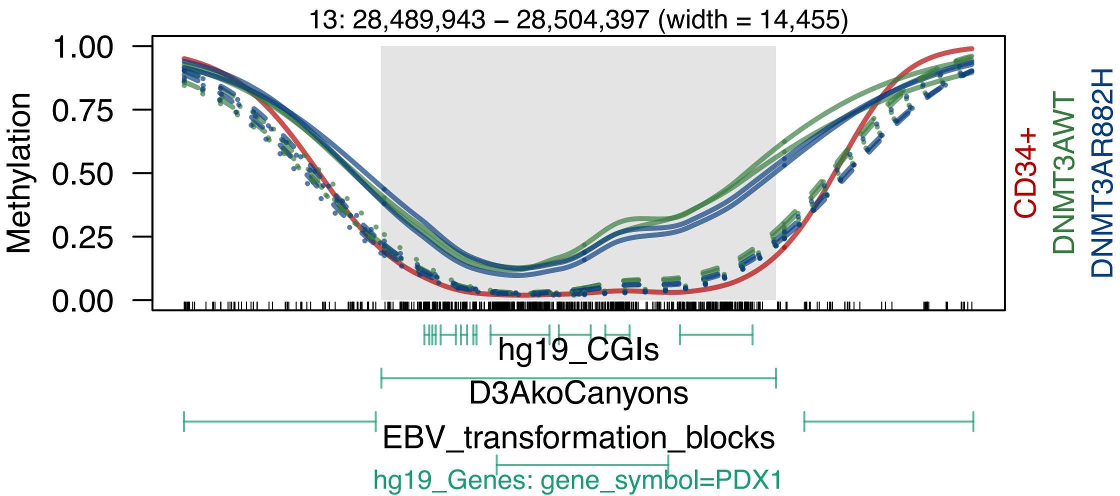
13: 28,489,943 – 28,504,397 (width = 14,455)
Methylation
1.00
0.75
0.50
0.25
0.00
CD34+
DNMT3AWT
DNMT3AR882H
hg19_CGIs
D3AkoCanyons
EBV_transformation_blocks
hg19_Genes: gene_symbol=PDX1

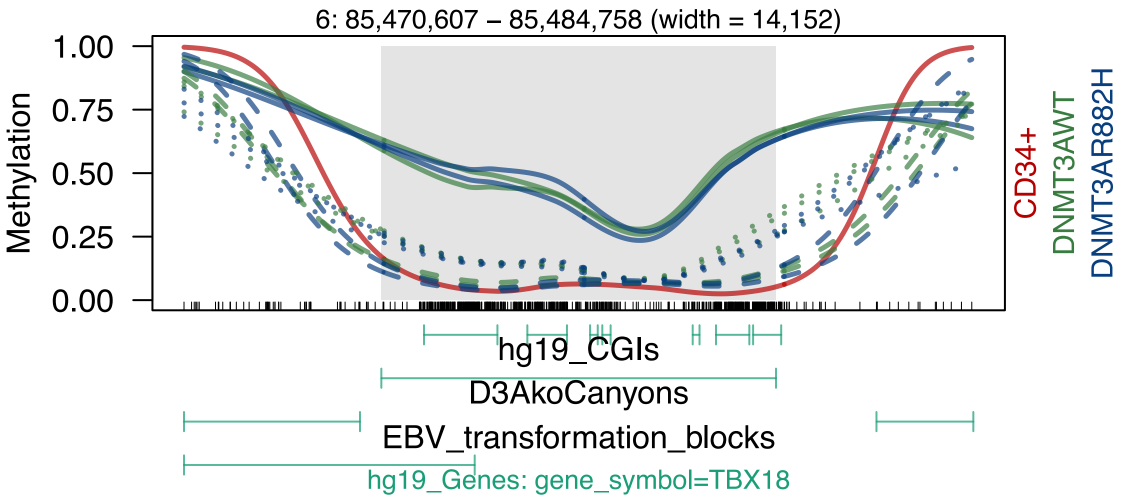
6: 85,470,607 – 85,484,758 (width = 14,152)
Methylation
1.00
0.75
0.50
0.25
0.00
CD34+
DNMT3AWT
DNMT3AR882H
hg19_CGIs
D3AkoCanyons
EBV_transformation_blocks
hg19_Genes: gene_symbol=TBX18

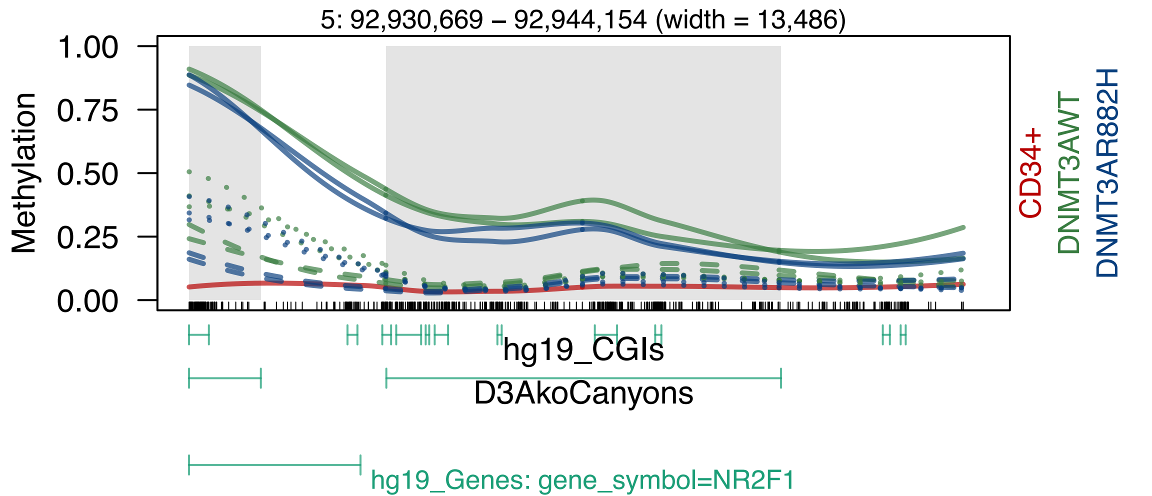

5: 92,930,669 – 92,944,154 (width = 13,486)
Methylation
1.00
0.75
0.50
0.25
0.00
CD34+
DNMT3AWT
DNMT3AR882H
hg19_CGIs
D3AkoCanyons
hg19_Genes: gene_symbol=NR2F1

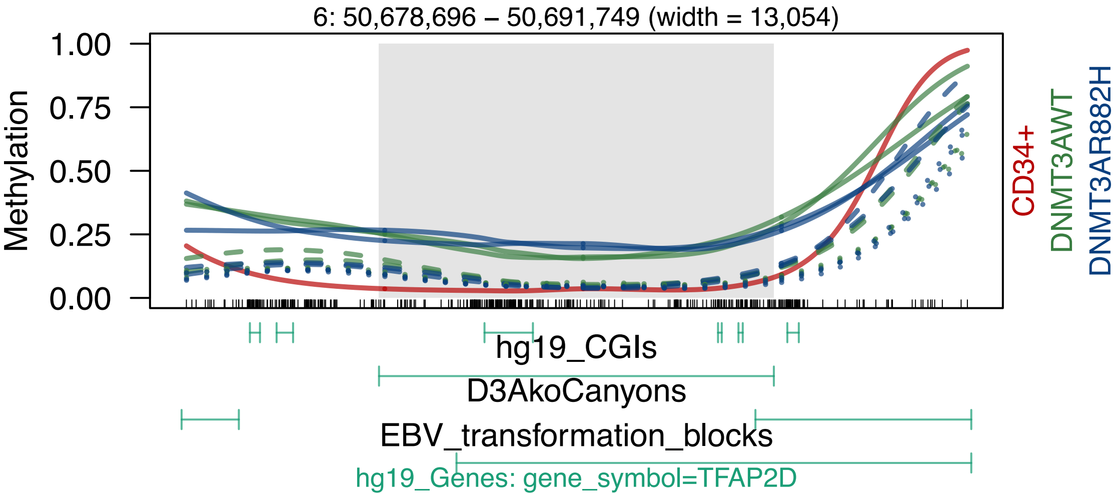

6: 50,678,696 – 50,691,749 (width = 13,054)
Methylation
1.00
0.75
0.50
0.25
0.00
CD34+
DNMT3AWT
DNMT3AR882H
hg19_CGIs
D3AkoCanyons
EBV_transformation_blocks
hg19_Genes: gene_symbol=TFAP2D

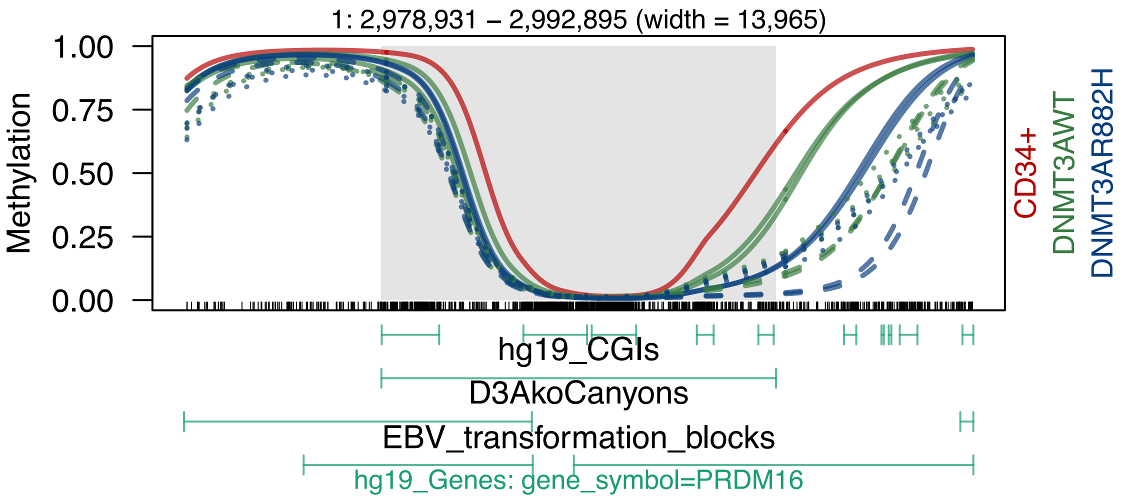

1: 2,978,931 – 2,992,895 (width = 13,965)
Methylation
1.00
0.75
0.50
0.25
0.00
CD34+
DNMT3AWT
DNMT3AR882H
hg19_CGIs
D3AkoCanyons
EBV_transformation_blocks
hg19_Genes: gene_symbol=PRDM16

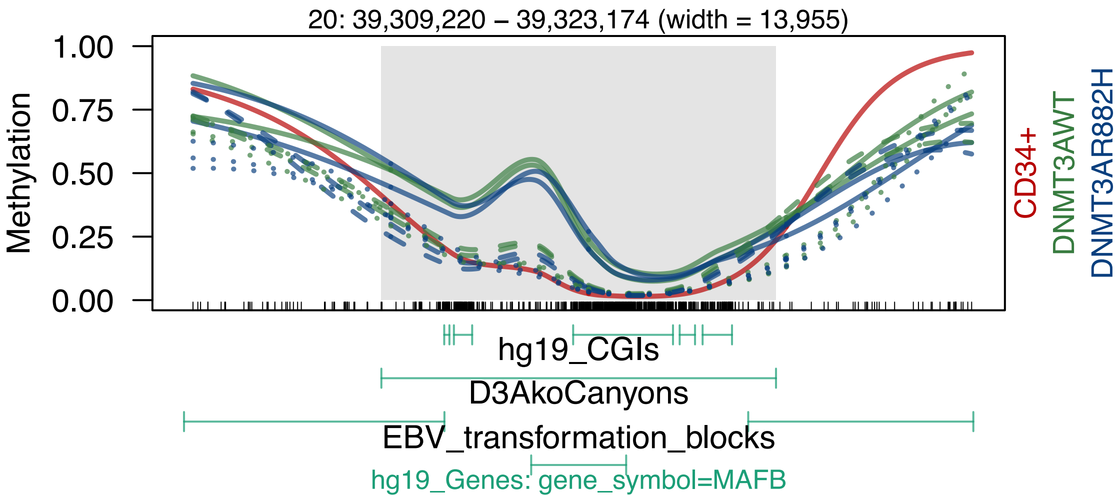
20: 39,309,220 – 39,323,174 (width = 13,955)
1.00
0.75
0.50
0.25
0.00
Methylation
CD34+
DNMT3AWT
DNMT3AR882H
hg19_CGIs
D3AkoCanyons
EBV_transformation_blocks
hg19_Genes: gene_symbol=MAFB

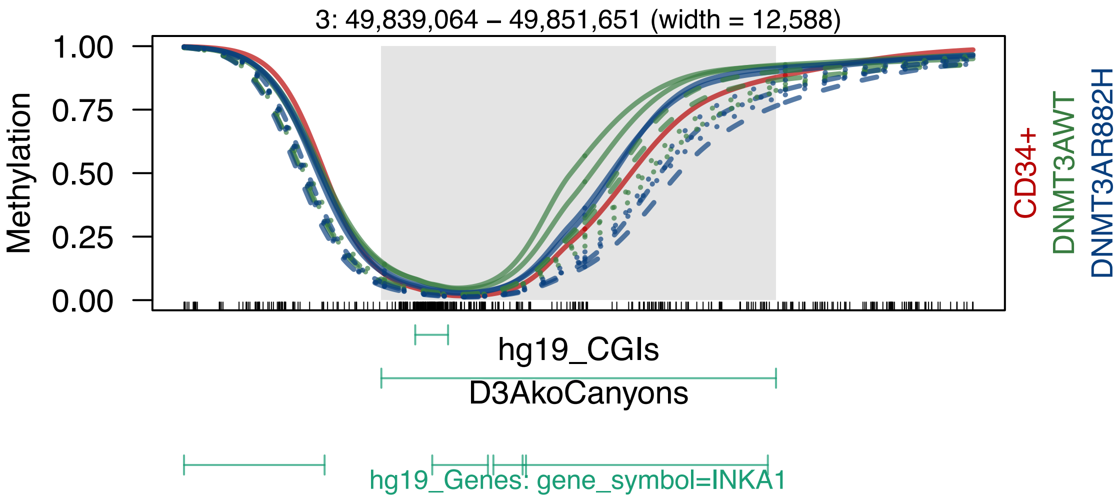
3: 49,839,064 – 49,851,651 (width = 12,588)
Methylation
1.00
0.75
0.50
0.25
0.00
CD34+
DNMT3AWT
DNMT3AR882H
hg19_CGIs
D3AkoCanyons
hg19_Genes: gene_symbol=INKA1

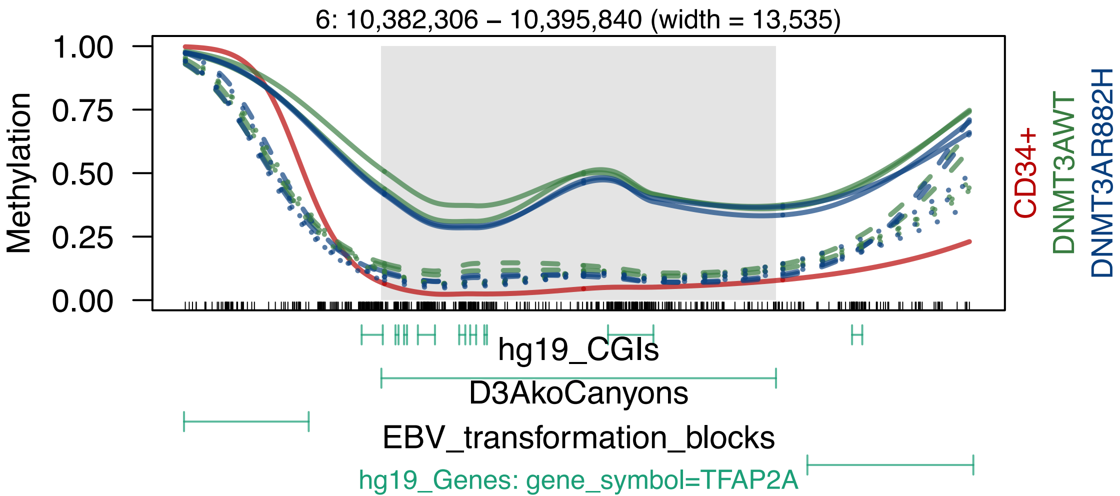
6: 10,382,306 – 10,395,840 (width = 13,535)
Methylation
1.00
0.75
0.50
0.25
0.00
CD34+
DNMT3AWT
DNMT3AR882H
hg19_CGIs
D3AkoCanyons
EBV_transformation_blocks
hg19_Genes: gene_symbol=TFAP2A

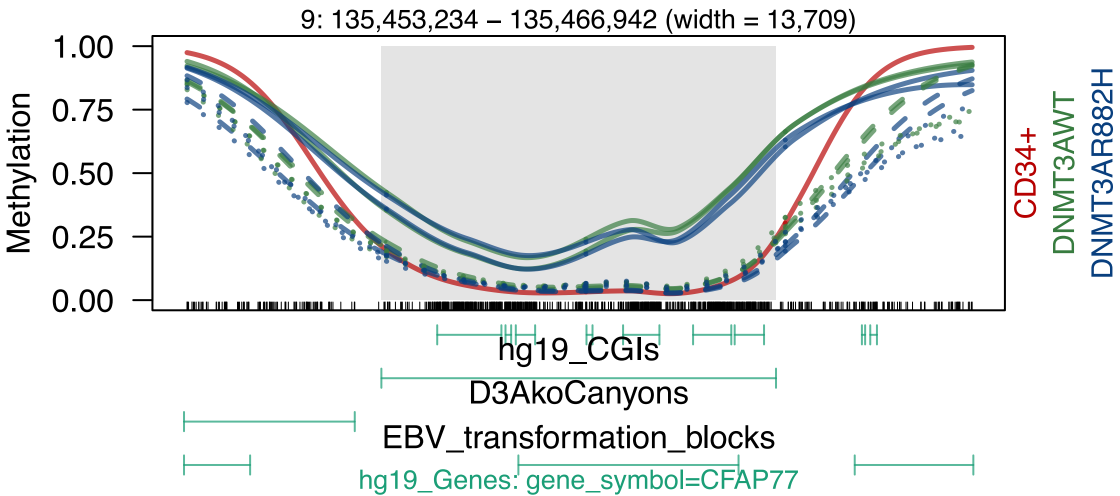
9: 135,453,234 – 135,466,942 (width = 13,709)
Methylation
1.00
0.75
0.50
0.25
0.00
CD34+
DNMT3AWT
DNMT3AR882H
hg19_CGIs
D3AkoCanyons
EBV_transformation_blocks
hg19_Genes: gene_symbol=CFAP77

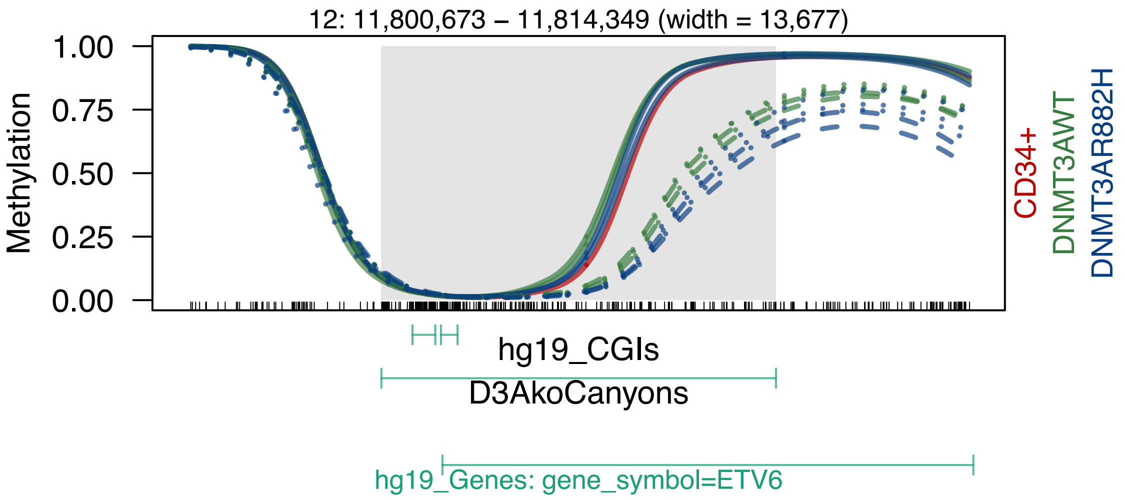
12: 11,800,673 – 11,814,349 (width = 13,677)
Methylation
1.00
0.75
0.50
0.25
0.00
CD34+
DNMT3AWT
DNMT3AR882H
hg19_CGIs
D3AkoCanyons
hg19_Genes: gene_symbol=ETV6

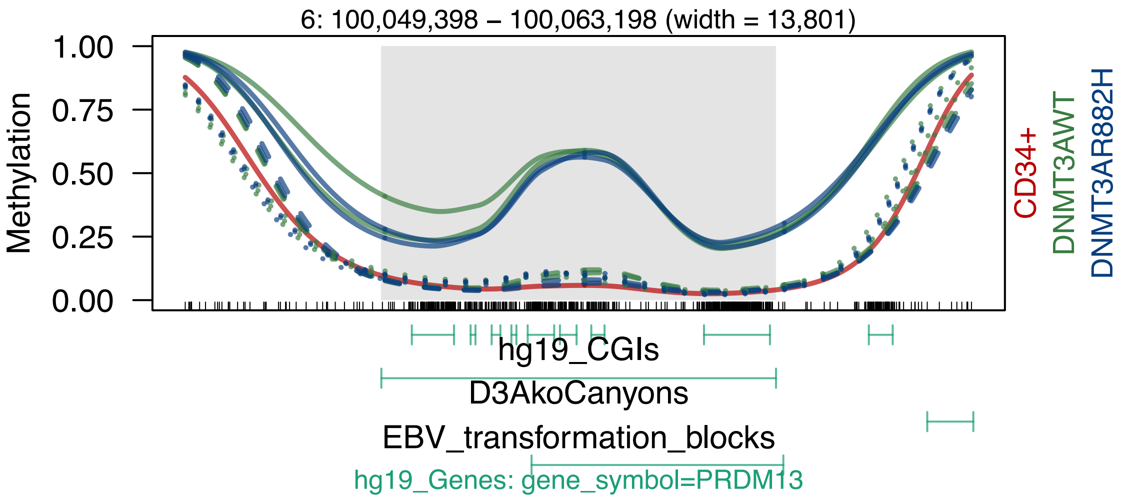
6: 100,049,398 – 100,063,198 (width = 13,801)
Methylation
1.00
0.75
0.50
0.25
0.00
CD34+
DNMT3AWT
DNMT3AR882H
hg19_CGIs
D3AkoCanyons
EBV_transformation_blocks
hg19_Genes: gene_symbol=PRDM13

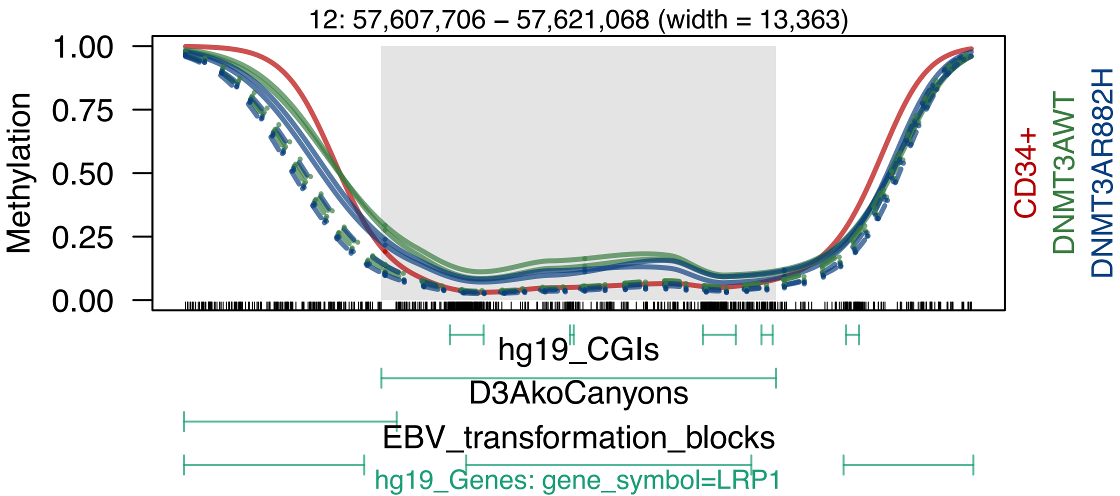
12: 57,607,706 – 57,621,068 (width = 13,363)
Methylation
1.00
0.75
0.50
0.25
0.00
CD34+
DNMT3AWT
DNMT3AR882H
hg19_CGIs
D3AkoCanyons
EBV_transformation_blocks
hg19_Genes: gene_symbol=LRP1

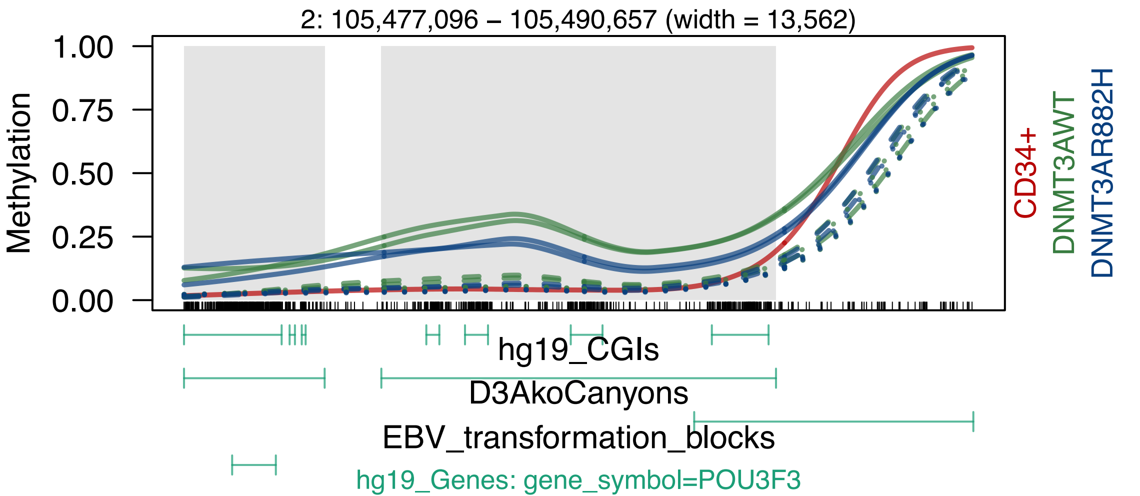
2: 105,477,096 – 105,490,657 (width = 13,562)
Methylation
1.00
0.75
0.50
0.25
0.00
CD34+
DNMT3AWT
DNMT3AR882H
hg19_CGIs
D3AkoCanyons
EBV_transformation_blocks
hg19_Genes: gene_symbol=POU3F3

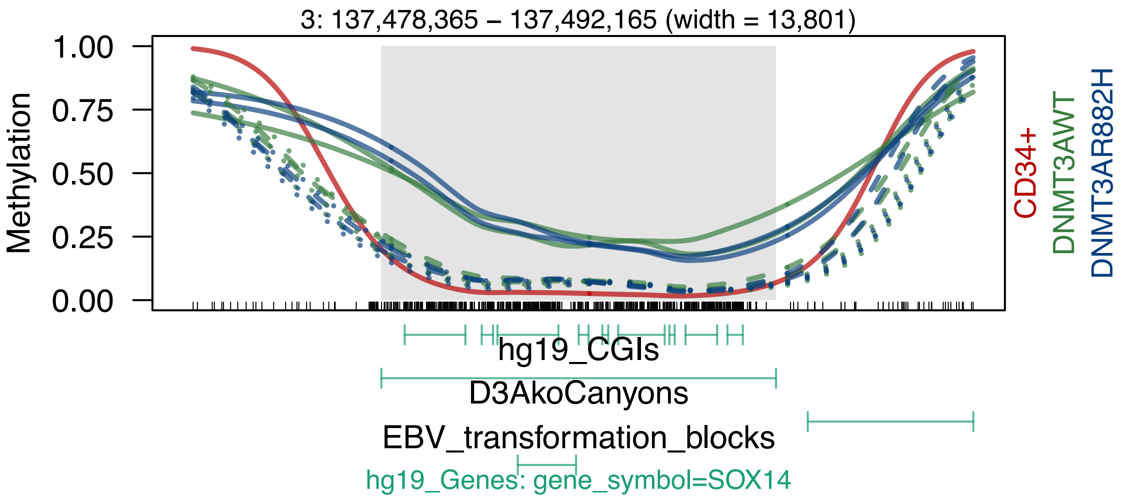
3: 137,478,365 – 137,492,165 (width = 13,801)
Methylation
1.00
0.75
0.50
0.25
0.00
CD34+
DNMT3AWT
DNMT3AR882H
hg19_CGIs
D3AkoCanyons
EBV_transformation_blocks
hg19_Genes: gene_symbol=SOX14

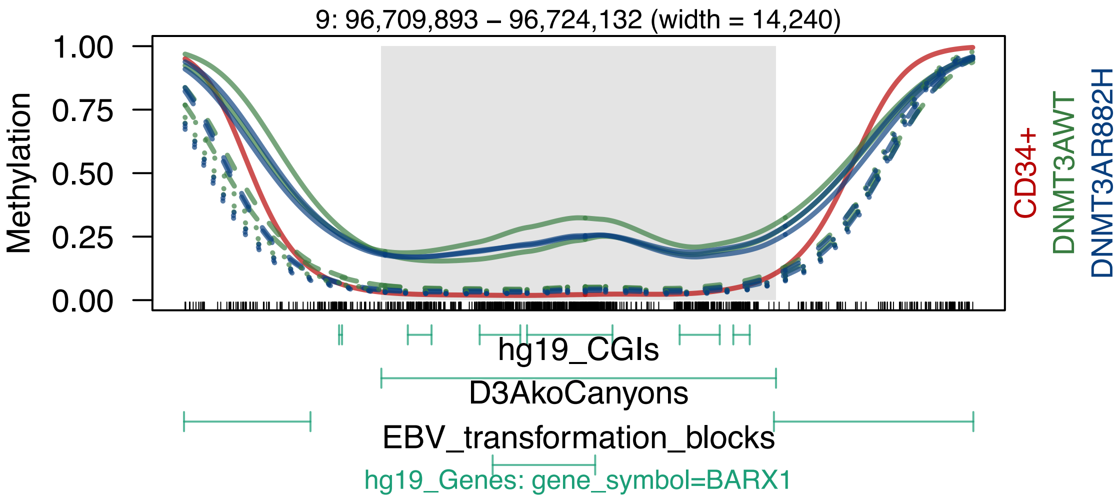
9: 96,709,893 – 96,724,132 (width = 14,240)
Methylation
1.00
0.75
0.50
0.25
0.00
CD34+
DNMT3AWT
DNMT3AR882H
hg19_CGIs
D3AkoCanyons
EBV_transformation_blocks
hg19_Genes: gene_symbol=BARX1

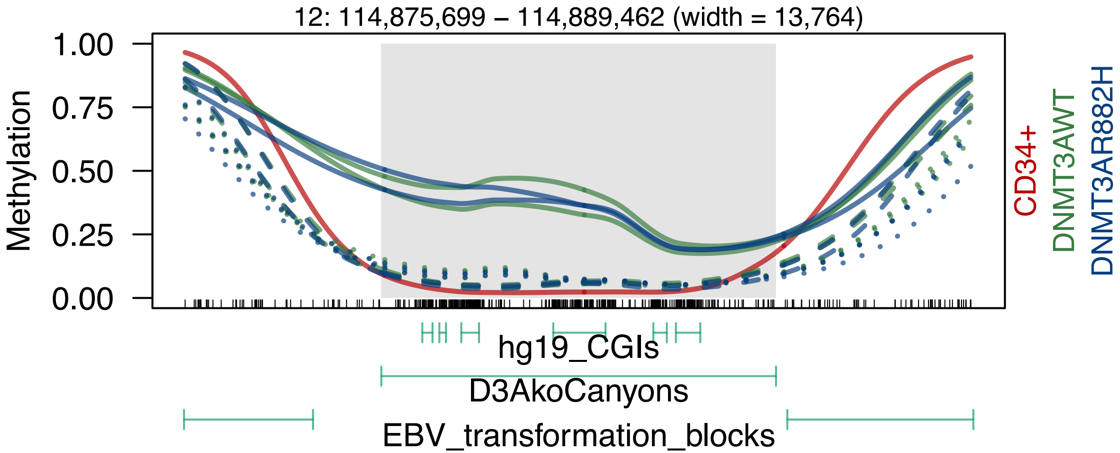
12: 114,875,699 – 114,889,462 (width = 13,764)
1.00
0.75
0.50
0.25
0.00
Methylation
CD34+
DNMT3AWT
DNMT3AR882H
hg19_CGIs
D3AkoCanyons
EBV_transformation_blocks

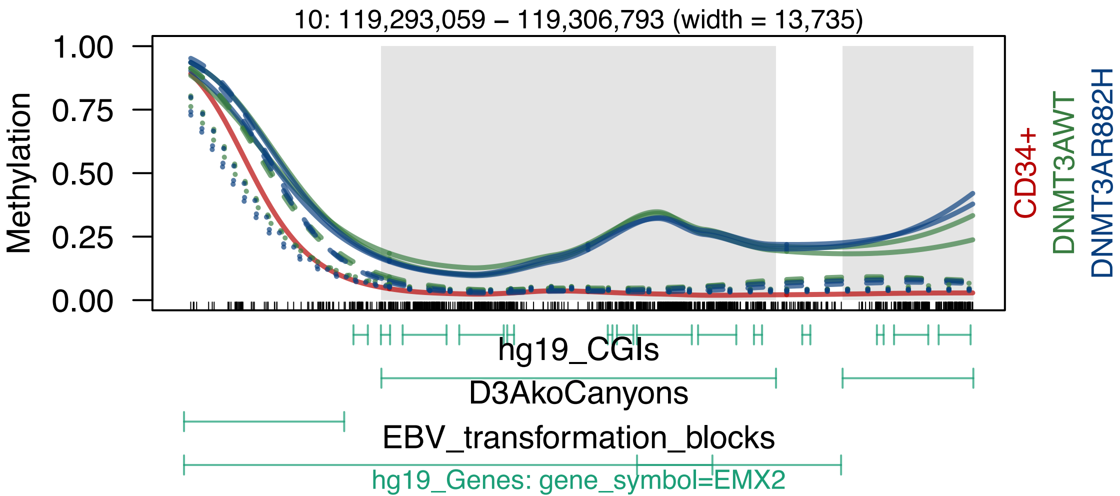

10: 119,293,059 – 119,306,793 (width = 13,735)
Methylation
1.00
0.75
0.50
0.25
0.00
CD34+
DNMT3AWT
DNMT3AR882H
hg19_CGIs
D3AkoCanyons
EBV_transformation_blocks
hg19_Genes: gene_symbol=EMX2

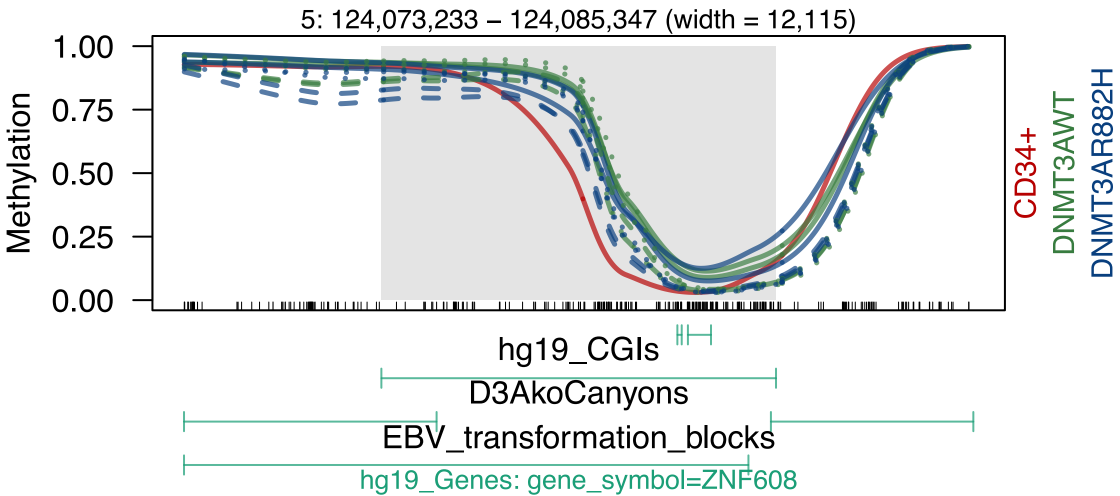
5: 124,073,233 – 124,085,347 (width = 12,115)
Methylation
1.00
0.75
0.50
0.25
0.00
CD34+
DNMT3AWT
DNMT3AR882H
hg19_CGIs
D3AkoCanyons
EBV_transformation_blocks
hg19_Genes: gene_symbol=ZNF608

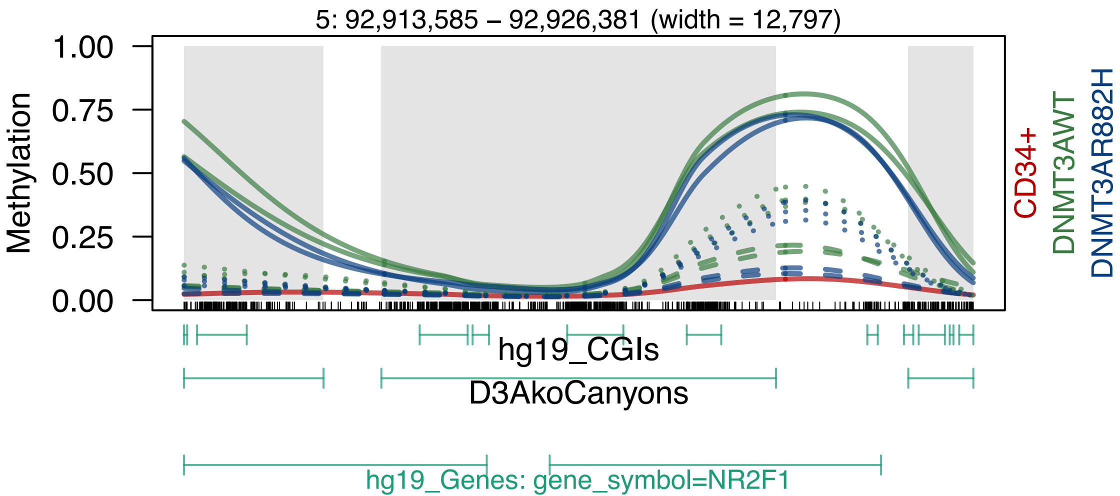
5: 92,913,585 – 92,926,381 (width = 12,797)
Methylation
1.00
0.75
0.50
0.25
0.00
CD34+
DNMT3AWT
DNMT3AR882H
hg19_CGIs
D3AkoCanyons
hg19_Genes: gene_symbol=NR2F1

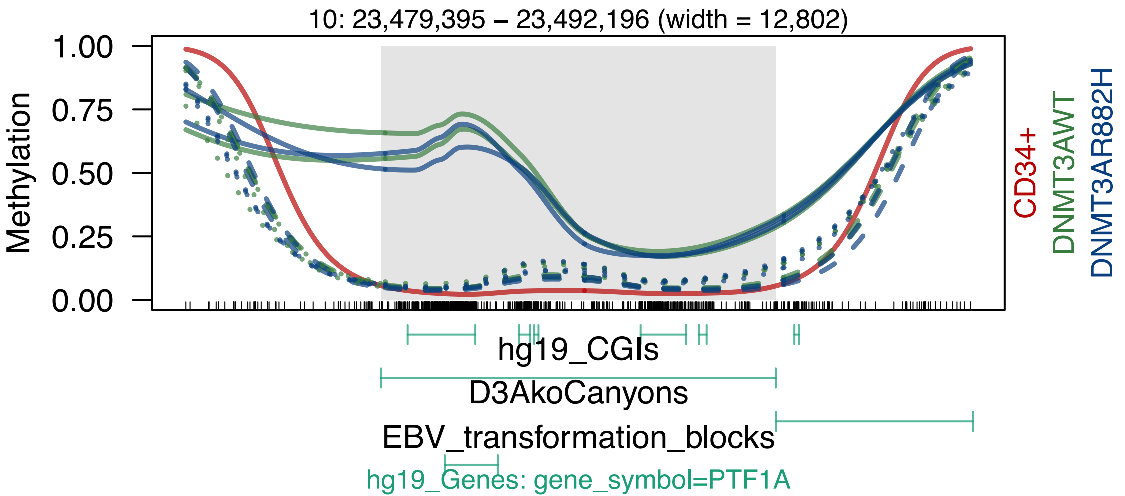
10: 23,479,395 – 23,492,196 (width = 12,802)
Methylation
1.00
0.75
0.50
0.25
0.00
CD34+
DNMT3AWT
DNMT3AR882H
hg19_CGIs
D3AkoCanyons
EBV_transformation_blocks
hg19_Genes: gene_symbol=PTF1A

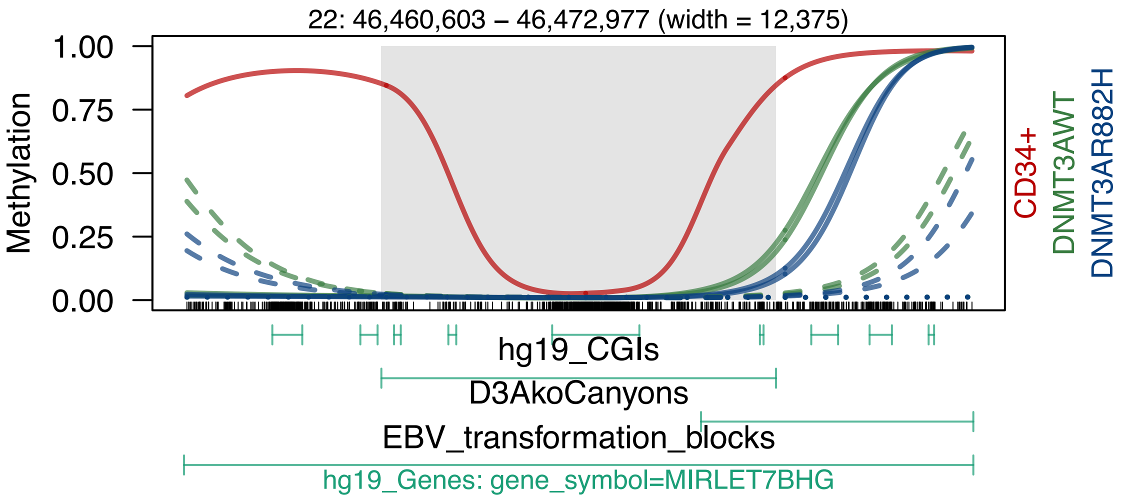
22: 46,460,603 – 46,472,977 (width = 12,375)
Methylation
1.00
0.75
0.50
0.25
0.00
CD34+
DNMT3AWT
DNMT3AR882H
hg19_CGIs
D3AkoCanyons
EBV_transformation_blocks
hg19_Genes: gene_symbol=MIRLET7BHG

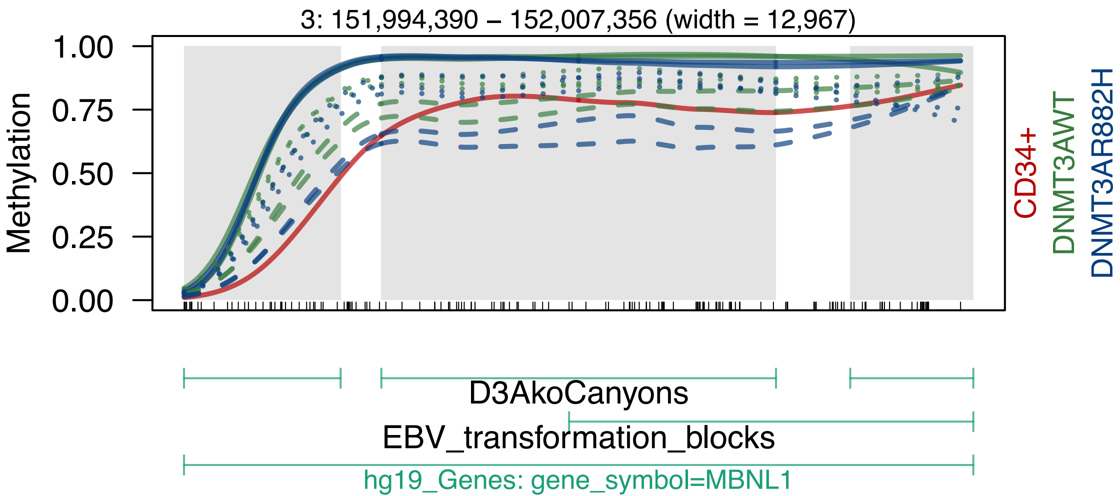

3: 151,994,390 – 152,007,356 (width = 12,967)
Methylation
1.00
0.75
0.50
0.25
0.00
CD34+
DNMT3AWT
DNMT3AR882H
D3AkoCanyons
EBV_transformation_blocks
hg19_Genes: gene_symbol=MBNL1

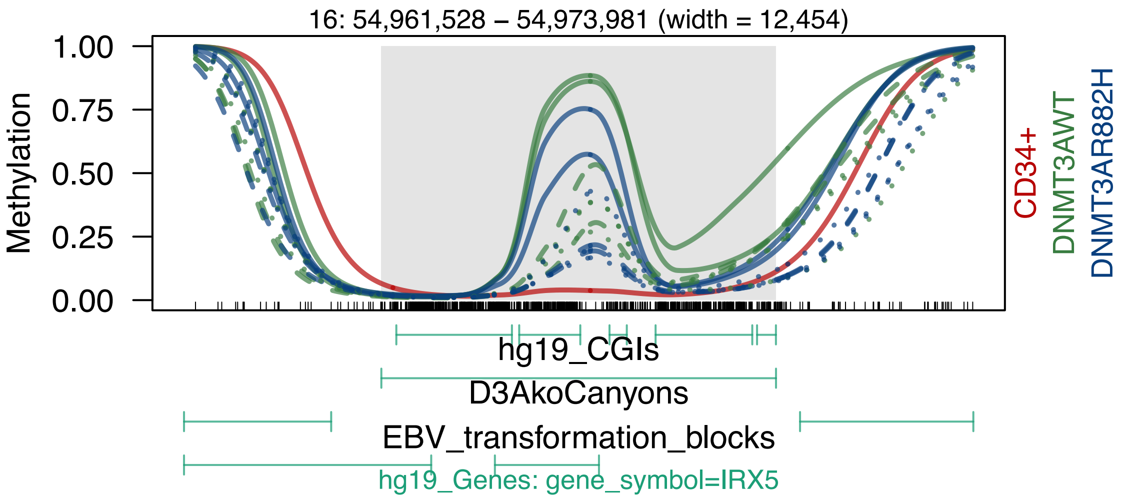
16: 54,961,528 – 54,973,981 (width = 12,454)
Methylation
1.00
0.75
0.50
0.25
0.00
CD34+
DNMT3AWT
DNMT3AR882H
hg19_CGIs
D3AkoCanyons
EBV_transformation_blocks
hg19_Genes: gene_symbol=IRX5

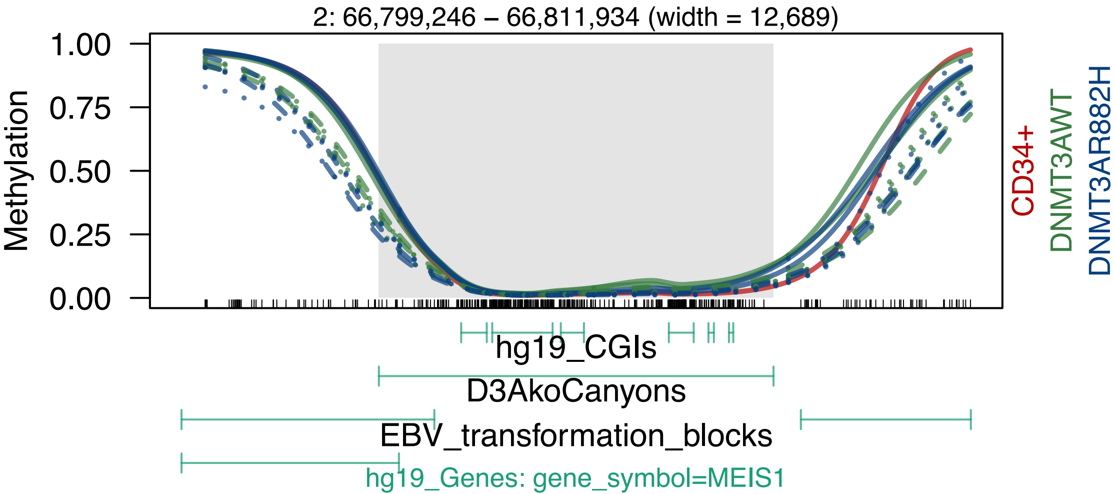

2: 66,799,246 – 66,811,934 (width = 12,689)
Methylation
1.00
0.75
0.50
0.25
0.00
CD34+
DNMT3AWT
DNMT3AR882H
hg19_CGIs
D3AkoCanyons
EBV_transformation_blocks
hg19_Genes: gene_symbol=MEIS1

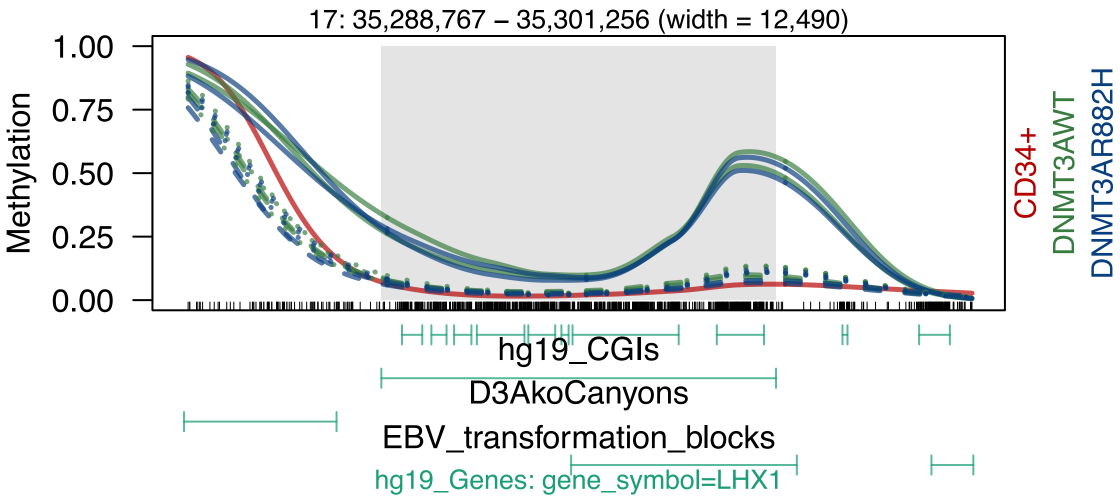
17: 35,288,767 – 35,301,256 (width = 12,490)
Methylation
1.00
0.75
0.50
0.25
0.00
CD34+
DNMT3AWT
DNMT3AR882H
hg19_CGIs
D3AkoCanyons
EBV_transformation_blocks
hg19_Genes: gene_symbol=LHX1

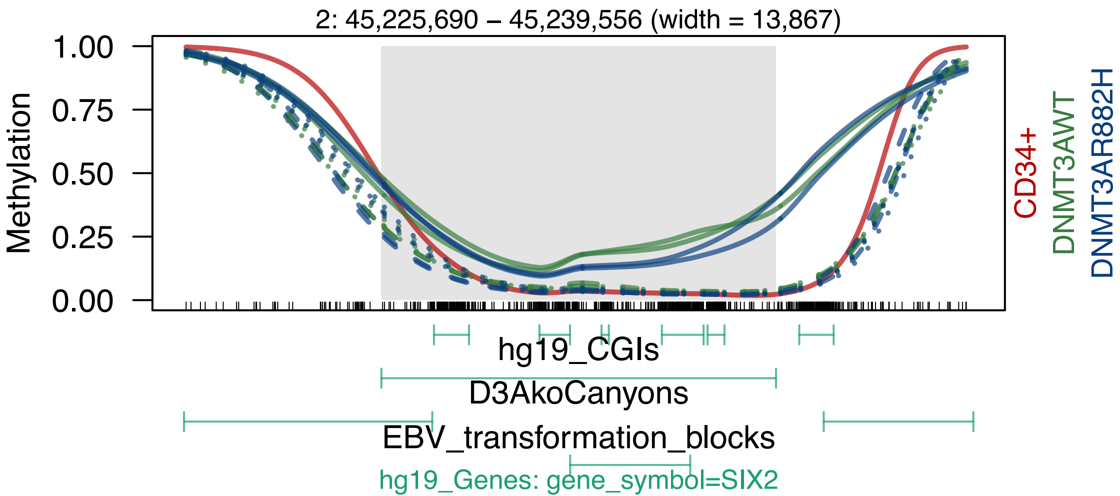
2: 45,225,690 – 45,239,556 (width = 13,867)
1.00
0.75
0.50
0.25
0.00
Methylation
CD34+
DNMT3AWT
DNMT3AR882H
hg19_CGIs
D3AkoCanyons
EBV_transformation_blocks
hg19_Genes: gene_symbol=SIX2

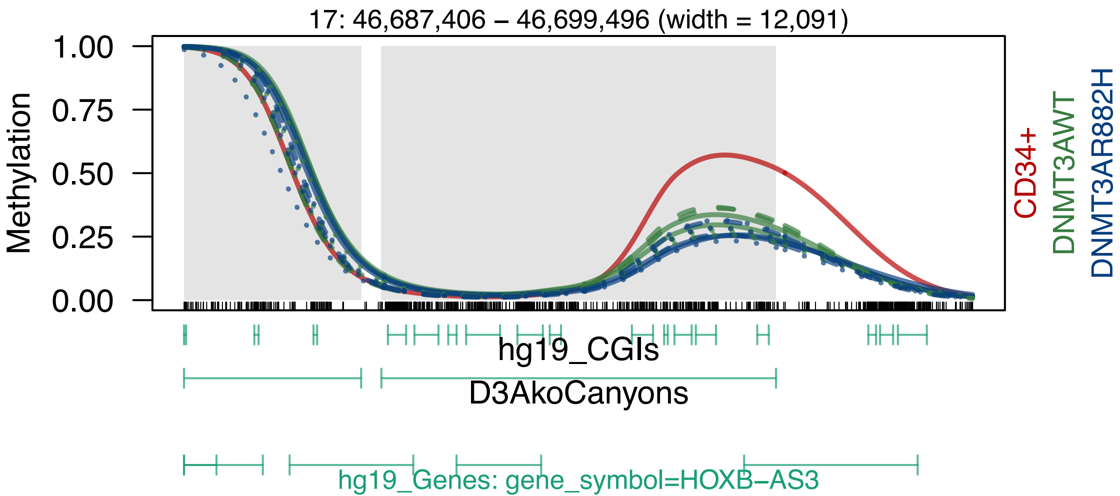

17: 46,687,406 – 46,699,496 (width = 12,091)
Methylation
1.00
0.75
0.50
0.25
0.00
CD34+
DNMT3AWT
DNMT3AR882H
hg19_CGIs
D3AkoCanyons
hg19_Genes: gene_symbol=HOXB–AS3

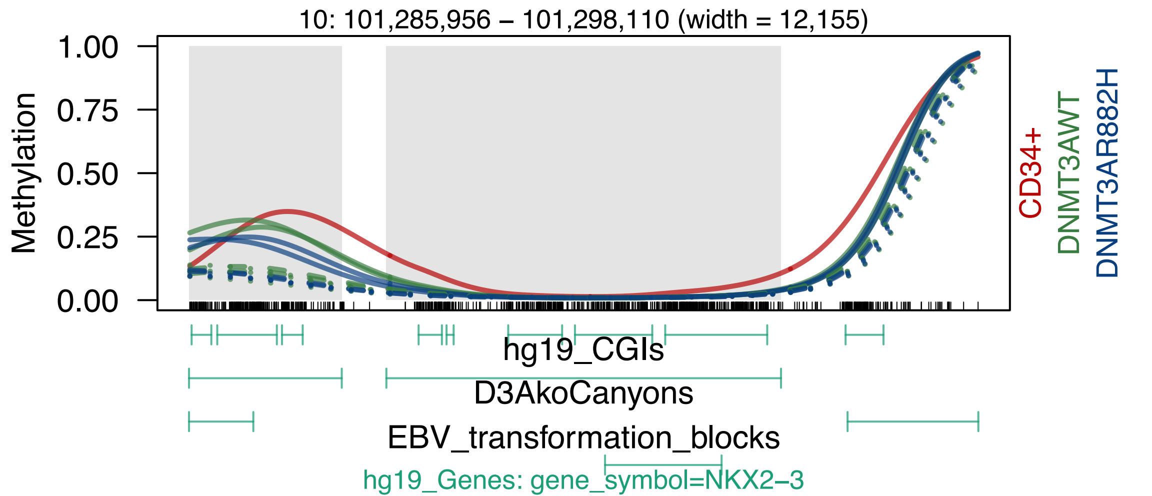
10: 101,285,956 – 101,298,110 (width = 12,155)
Methylation
1.00
0.75
0.50
0.25
0.00
CD34+
DNMT3AWT
DNMT3AR882H
hg19_CGIs
D3AkoCanyons
EBV_transformation_blocks
hg19_Genes: gene_symbol=NKX2–3

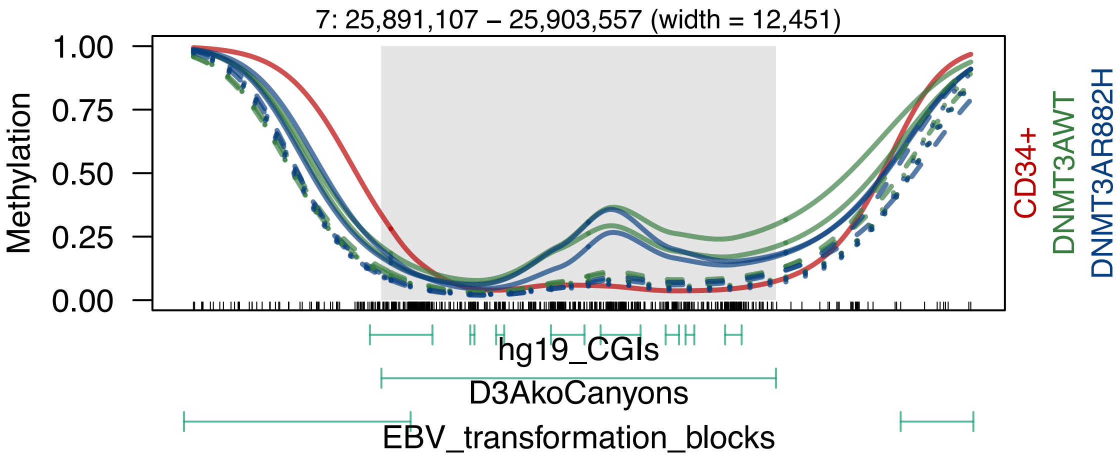
7: 25,891,107 – 25,903,557 (width = 12,451)
Methylation
1.00
0.75
0.50
0.25
0.00
CD34+
DNMT3AWT
DNMT3AR882H
hg19_CGIs
D3AkoCanyons
EBV_transformation_blocks

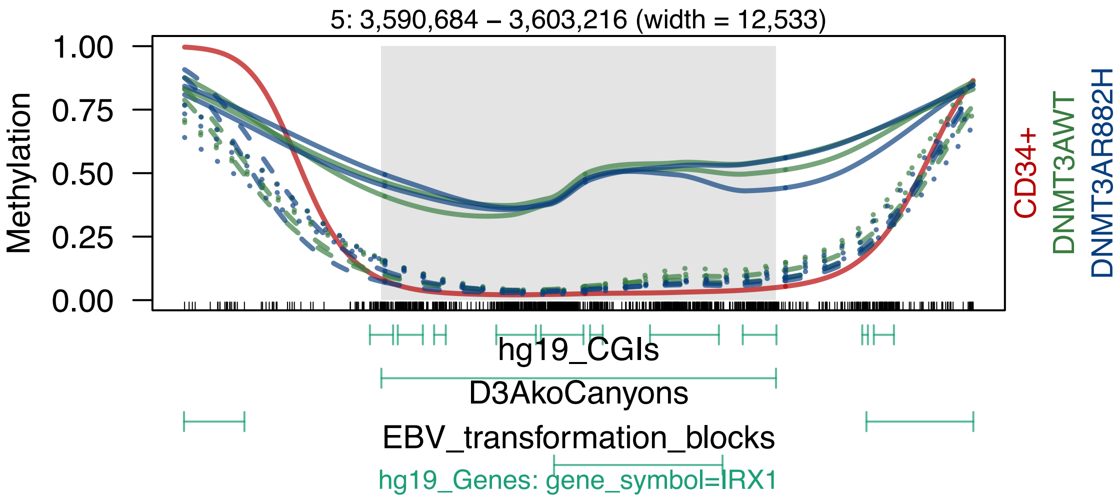
5: 3,590,684 – 3,603,216 (width = 12,533)
Methylation
1.00
0.75
0.50
0.25
0.00
CD34+
DNMT3AWT
DNMT3AR882H
hg19_CGIs
D3AkoCanyons
EBV_transformation_blocks
hg19_Genes: gene_symbol=IRX1
